# Synthetic HNK-1 containing glycans provide insight into binding properties of serum antibodies from MAG-neuropathy patients

**DOI:** 10.1101/2022.07.25.501369

**Authors:** Mehman Bunyatov, Margreet A. Wolfert, Ruth Huizinga, Marco W.J. Schreurs, Bart C. Jacobs, Geert-Jan Boons

**Affiliations:** Department of Chemical Biology and Drug Discovery, Utrecht Institute for Pharmaceutical Sciences, and Bijvoet Center for Biomolecular Research, Utrecht University, Utrecht, The Netherlands; Complex Carbohydrate Research Center, University of Georgia, Athens, GA, USA; Department of Immunology and University Medical Center, Rotterdam, The Netherlands; Department of Neurology, Erasmus MC, University Medical Center, Rotterdam, The Netherlands; Department of Chemistry, University of Georgia, Athens, GA, USA

**Author notes:** **Author Contributions.** M.B., M.A.W., R.H., B.C.J., and G.J.B. conceived the studies; M.B. and G.J.B. conceived the chemo-enzymatic approach; M.B. performed the chemoenzymatic synthesis; M.A.W. performed the array studies and associated data analysis; M.W.J.S. performed the Bühlmann ELISA assay; M.B., M.A.W., R.H., B.C.J., and G.J.B. wrote the paper.

**Keywords:** Chemoenzymatic synthesis, Glycans, Microarray, Auto-antibodies, Anti-MAG neuropathy

## Abstract

Anti-myelin-associated glycoprotein (anti-MAG) neuropathy is an autoimmune disease in which IgM autoantibodies target glycoconjugates of peripheral nerves resulting in progressive demyelination. To examine fine specificities of serum IgM autoantibodies and develop a more robust platform for diagnosis and disease monitoring, we describe here a chemoenzymatic approach that readily provided a panel of HNK-1 containing oligosaccharides presented on type 2 oligo-*N*-acetyl lactosamine (LacNAc) chains typical of glycosphingolipids. The compounds were prepared by a chemoenzymatic strategy in which an oligo-LacNAc structure was assembled enzymatically and then subjected to protecting group manipulation to chemically install a 3-*O*-sulfate glucuronic acid moiety. The synthetic strategy is highly divergent and made it possible to prepare from key precursors, additional compounds lacking sulfate of HNK-1 and derivatives in which the HNK-1 epitope is replaced by sulfate or sialic acid. The oligosaccharides were printed as a microarray to examine binding specificities of several monoclonal antibodies and serum antibodies of anti-MAG neuropathy patients. Surprisingly, three distinct patient subgroups were identified with variable dependance on the length of the LacNAc chain and sulfation of the glucuronyl moiety. In most cases, a lacto-*neo*hexaose backbone was required for binding indicating the antibodies target corresponding glycosphingolipids.

**Significance statement:** A chemoenzymatic strategy is introduced in which a glycan backbone is assembled by glycosyltransferases to give a core oligosaccharide that is subjected to protecting group manipulations and chemical glycosylations to install terminal epitopes. It addresses limitations of enzymatic synthesis when specific glycosyltransferases or glycan-modifying enzymes for terminal epitope synthesis are not readily available. It provided an unprecedented panel of HNK-1 containing oligosaccharides, which was used to develop a glycan microarray that uncovered distinct binding preferences of serum antibodies of anti-MAG patients. The clinical spectrum of IgM monoclonal gammopathy varies substantially and an understanding of binding properties of IgM auto-antibodies will provide opportunities to monitor disease progression and develop personalized treatment options.

## Introduction

Glycoconjugates play important roles in the development and maintenance of the nervous system and are involved in processes such as the formation and maintenance of myelin, neurogenesis, neurite outgrowth, and synaptic plasticity (1). The importance of glycans in nervous system development and function is highlighted by congenital disorders of glycosylation, which often are associated with psychomotor retardation, and other neuropathological symptoms (2, 3).

The human natural killer-1 (HNK-1) epitope is a trisaccharide composed of a 3-*O*-sulfated glucuronic acid linked to *N*-acetyl-lactosamine (HSO_3_-3GlcAß1-3Galß1-4GlcNAc-) (Fig. 1A). HNK-1 was first identified on human natural killer cells as a differentiation antigen (CD57) (4). Subsequent studies have shown that this glycan epitope is also abundantly expressed by Schwann cells and oligodendrocytes of the peripheral and central nervous system, respectively (5-8). It is a terminal epitope of *N*-glycans of myelin associated glycoprotein (MAG) and myelin proteins (P0, P22) (Fig. 1B) and is also found as part of glycosphingolipids of the lacto-series (Fig. 1C). Several studies have indicated that HNK-1 is important for nervous system development, plasticity and dendritic spine morphogenesis (1, 9, 10). It acts as neural-recognition molecules employing laminin, P- and L-selectins and galectins as receptors for cell adhesion (11-13).

**Figure 1.**
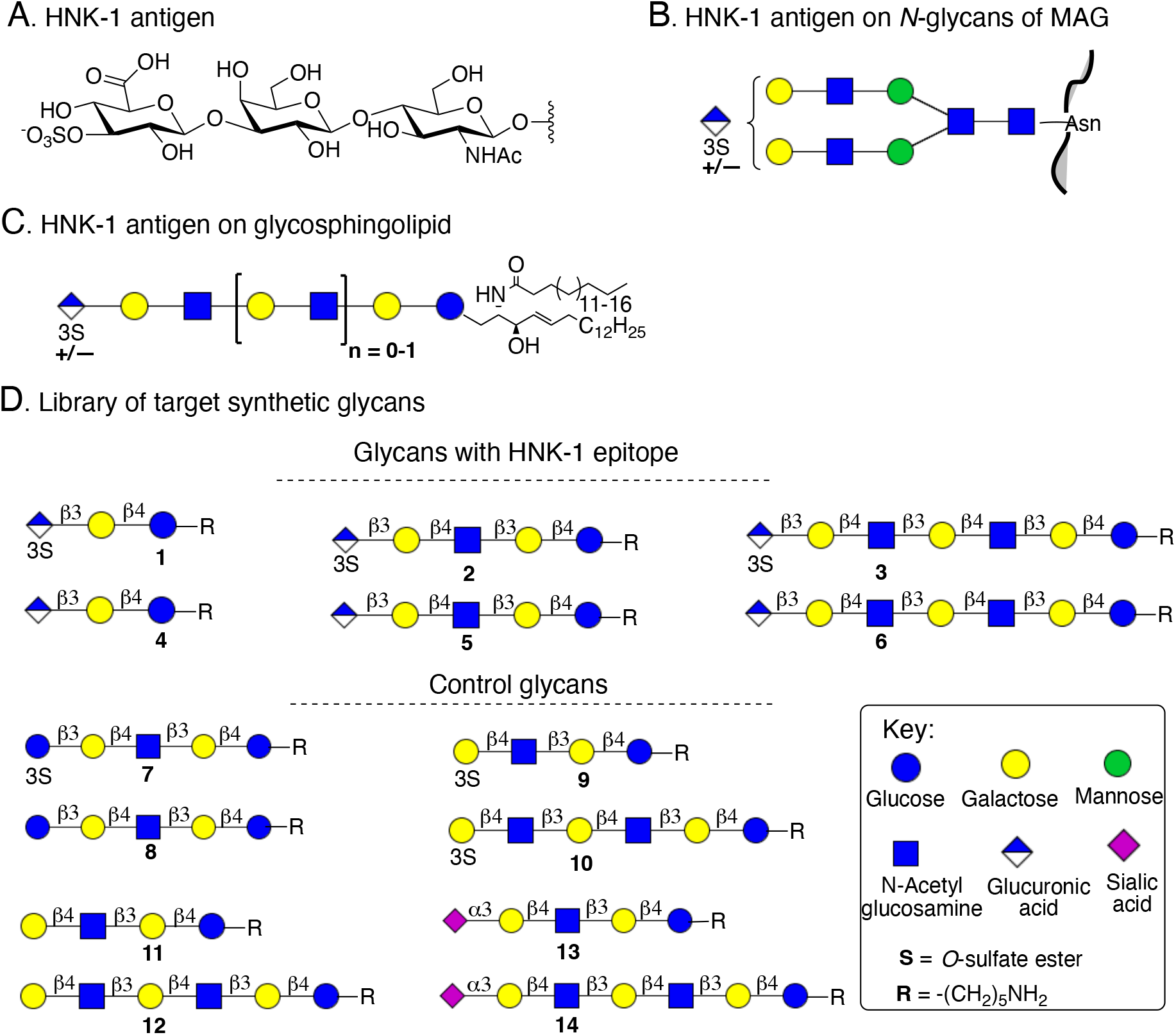
Chemical composition of HNK-1 antigen (**A**), its presentation on N-glycans of glycoprotein (**B**) and glycosphingolipids (**C**). Target library of glycans having various epitopes (**D**).

HNK-1 is also associated with disease and for example its expression is substantially reduced in brains of Alzheimer disease patients probably influencing β-amyloid protein (14). It can be an autoantigen and anti-myelin-associated glycoprotein (anti-MAG) neuropathy is caused by IgM monoclonal antibodies targeting HNK-1 containing glycoconjugates (6). It results in a chronic and slowly progressive demyelinating polyneuropathy mainly affecting large fibre sensory and motor nerves. Clinical symptoms include sensory ataxia with impaired gait, tremor, distal muscle weakness and neuropathic pain eventually leading to substantial disabilities.

Initial studies indicated that pathogenic serum autoantibodies from neuropathy patients target *N*-glycans of myelin associated glycoproteins (MAG) (15, 16). Subsequent studies pointed to glycosphingolipids as possible targets for the IgM auto-antibodies (17-20). For example, immunization of Lewis rats with sulfoglucuronyl paraglobosides caused clinical symptoms resembling anti-MAG neuropathy, whereas similar experiments with MAG did not result in any symptoms. Furthermore, passive transfer of patients’ anti-MAG antibodies and complement factors to experimental animals affected peripheral nerve myelin where HNK-1 is mainly presented as part of glycosphingolipids (21). Antibodies targeting glycosphingolipids have been implicated in other neuropathies such as Guillain-Barre syndrome where gangliosides are the primarily target (22, 23).

The structures of the underlying glycan moiety of the HNK-1 epitope on *N*-linked glycans and glycosphingolipids differ considerably. In case of *N*-linked glycans, the epitope is usually attached to the 1,6-linked mannoside of the core of biantennary glycans (Fig. 1B), whereas on glycosphingolipids it is attached to a type 2 oligo-*N*-acetyl lactosamine (LacNAc) moiety (Fig. 1B) (18, 24-26). These differences in presentation may influence binding properties of pathogenic auto-antibodies. Such properties have, however, been difficult to investigate due to a lack of well-defined HNK-1 containing oligosaccharides for binding studies (27-30). A detailed understanding of binding selectivities of pathogenic auto-antibodies is expected to provide opportunities for assay development for disease detection, predicting prognosis and for the development of personalized treatment options. Common diagnostic assays for IgM monoclonal gammopathy are based on Western blotting or ELISA using isolated MAG (31). These approaches give only 50-70% positive results for patients exhibiting demyelinating polyneuropathy. Assays that can detect anti-MAG as well as anti-glycosphingolipids antibodies will be more reliable and make it possible to identify distinct subsets of patients opening opportunities for predicting disease progression and personalizing treatment options.

Here, we report the synthesis of a series of glycans derived from sulfoglucuronyl paraglobosides (**1-3**) and similar compounds lacking the sulfate ester (**4-6**) (Fig. 1D). As control compounds, glucosyl paraglobosides with and without sulfate ester (**7, 8**), 3-*O*-sulfo-paraglobosides (**9, 10**), lacto-*N*-neotetraose and lacto-*N*-neohexaose (**11, 12**) and sialo-glycans (**13, 14**) were also prepared. The oligosaccharides were printed as a microarray, which was used to examine binding specificities of several monoclonal antibodies and serum antibodies of anti-MAG patients. Surprisingly, three distinct patient subgroups were identified. One group selectively bound to compounds containing an HNK-1 epitope, and in these cases, the antibodies showed a strong preference for a compound having two LacNAc moieties indicating they target glycosphingolipids. Another group of serum samples bound to compounds having a glucuronyl moiety lacking a sulfated ester, and in these instances, the dependance on chain length was modest. The remaining serum samples recognized sulfated as well as non-sulfated glucuronyl glycan epitopes with a preference for extended structures.

## Results and Discussion

### Chemoenzymatic synthesis of oligo-LacNAc precursor

Chemoenzymatic synthesis offers an attractive approach to prepare highly complex oligosaccharides (32-35). A common strategy is based on the chemical synthesis of a glycan structure followed by installation of various terminal epitopes using recombinant glycosyl transferases (Fig. 2A) (36-40). Progress in recombinant glycosyl transferase expression (41-43) is also making it possible to employ fully enzymatic methods to assemble complex glycans (44). In particular, the use of modified sugar nucleotides, which after transfer provide products that temporarily block specific sites from further enzymatic modification, provide opportunities to assemble oligosaccharides with complex branched architectures (45-47). Also, selective chemical modification of a complex glycan intermediate prepared by glycosyl transferases, allows to temporarily disable specific sites from further enzymatic modification (48, 49).

**Figure 2.**
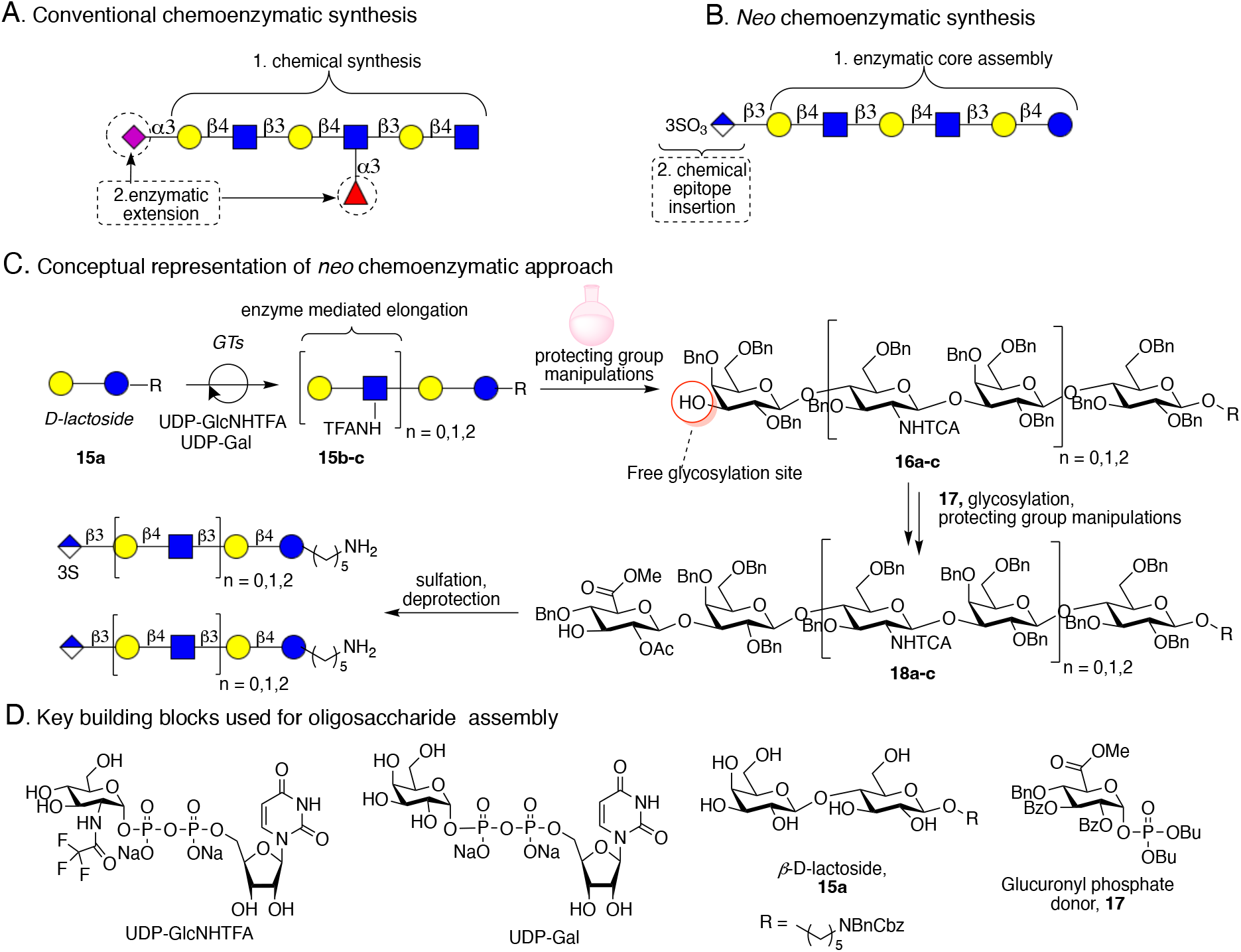
Conventional chemoenzymatic strategy (**A**), Neo-chemoenzymatic approach (**B**), and Conceptual representation of neo-chemoenzymatic strategy (**C**). Key building blocks used to synthesize the collection of glycans (**D**).

Despite the versatility of enzyme mediated oligosaccharide assembly, it has as a limitation that not all glycosyl transferases are readily available to install all natural occurring epitopes. Here, we introduce a chemoenzymatic strategy that we coined neo-chemoenzymatic synthesis in which glycosyl transferases are used to assemble a glycan structure that is subjected to protecting group manipulation to provide a compound that can be subjected to chemical glycosylations and further modifications (Fig. 2B). It made it possible to install terminal epitopes that could not readily be introduced enzymatically. The methodology was exploited to prepare a range of sulfoglucuronyl- (**1-3**) and glucuronyl paraglobosides (**4-6**). It entailed the enzymatic assembly of lacto-*neo*tetraose **15b** and lacto-*neo*hexaose **15c** using recombinantly expressed HpB3GnT and LgtB enzymes in combination with UDP-GlcNHTFA and UDP-Gal. The TFA moiety of **15b** and **15c** could readily be cleaved by mild base and the resulting amines converted into azides by azido transfer reaction to provide compounds amenable to protecting group manipulations providing access to glycosyl acceptors **16a-c**. The latter compounds could readily be glycosylated with glucuronyl donor **17** to give products that were subjected to further protecting group manipulations to give compounds **18a-c** having a free C-3 hydroxyl at the glucuronyl moiety. *O*-sulfation and deprotection of the latter compounds provided HNK-1 containing paraglobosides **1-3**. Deprotection of **18a-c** gave similar compounds lacking a sulfate (Fig. 2C). Key intermediates **16b**,**c** also provided access to other compounds, and for example made it possible to prepare sulfates **9** and **10**.

The synthesis of the targeted compounds started with lactosyl acceptor **15a** (Scheme 1A) having an anomeric aminopentyl linker important for microarray development. The amine of the compound was doubly protected by benzyl (Bn) and benzyloxycarbonyl to allow protecting group manipulations at a later stage of synthesis. Compound **15a** was subjected to recombinant HpB3GnT in the presence of UDP-GlcNHTFA, and fortunately the enzyme readily utilized the chemically modified sugar nucleotide donor to give a trisaccharide that was further extended by a galactosyl moiety using LgtB in the presence UDP-Gal to provide tetrasaccharide **15b**. The same enzyme module was used to install an additional LacNHTFA unit to provide hexasaccharide **15c**. The product of each enzymatic transformation was purified by Biogel P-2 size exclusion column chromatography and fully characterized by NMR and MALDI-TOF mass spectrometry. Next, the trifluoroacetamides of **15b** and **15c** were hydrolyzed by treatment with mild aqueous base (pH = 10) to furnish the corresponding glucosamine derivatives **19b** and **19c**, respectively that were subjected to an azido-transfer reaction employing imidazolonium sulfonyl azide (47, 50) to give azido-containing derivatives **20b**,**c**.

Next, a protecting group strategy was developed to converted **15a** and **20b**,**c** into glycosyl acceptors **16a-c** for chemical installation of a 3-*O*-sulfated glucuronic acid moiety. Benzyl ethers were selected as permanent protecting groups because these were expected to enhance the glycosyl accepting properties of **16a-c**. Furthermore, azides were used as masking groups of the amines because of their compatibility with the introduction of benzyl ethers under strong basic conditions. It has been reported that the C-3 hydroxyl of a terminal galactoside can selectively be alkylated by employing and intermediate stannylene acetal (51, 52), and this approach was exploited for regioselective allylation to give compounds **21a-c**. Thus, compounds **15a** and **20b-c** were heated under reflux in methanol in the presence of Bu_2_SnO to form a stannylene acetal. The methanol was removed by evaporation and the residue dissolved in DMF and then treated with allyl bromide and Bu_4_NI as an activator. The resulting product was purified by silica gel column chromatography to give selectively allylated derivatives **21a-c**. The remaining alcohols of **21a-c** were benzylated by reaction with benzyl bromide and NaH in DMF, which was followed by Staudinger reduction of the azides using trimethyl phosphine to give amines that were converted into trichloroacetamides (TCA) by reaction with trichloroacetyl chloride in the presence of triethylamine by reduction to give **22a-c**. Trichloroacetamides were selected for intermediate amino protection because during hydrogenation to remove benzyl and benzyloxycarbonyl protecting groups, they will also be reduced to natural acetamides. Finally, the allyl ether of the di-(**22a**), tetra-(**22b**), and hexasaccharide (**22c**) intermediates was cleaved by treated with catalytic PdCl_2_ in a mixture of methanol and chloroform to provide glycosyl acceptors **16a-c**.

### GlcA introduction, deprotection and further functionalization

Chemical glycosylation of glucuronate donors is often low yielding due to their poor reactivity. Anomeric phosphates have emerged as the most appropriate anomeric leaving group for glucuronyl donors, however, several cycles of glycosylation using an excess of donor is usually required to achieve high yielding glycosylation (28). In this study, we prepared 2,3-di-*O*-benzoyl substituted glucuronyl phosphate **17** (see *SI* for synthetic details), and discovered it has favorable glycosyl donor properties. Thus, TMSOTf-mediated glycosylation of glycosyl acceptors **16a-c** with donor **17** in DCM at -50 °C gave the corresponding tri-, penta- and heptasaccharides **23a-c** in high yield as only the β-anomer (Scheme 2). The linkage position and anomeric configuration of the newly installed GlcA moiety was confirmed by the chemical shift analysis which showed H-3 of the terminal galactoside at a more downfield region and measurement of ^13^C-^1^H coupling constant at 2D HSQC off-decoupled spectra, respectively.

The carboxylic acid of glucuronic acid can undergo intramolecular lactonization with its C-3 hydroxyl (30), and therefore we expected that such a reaction could be employed for selective protection of the C-3 hydroxyl important for the installation of a 3-O-sulfate ester. Thus, compounds **23a-c** were subjected to KOH (1M) in aqueous dioxane and then methanol to hydrolyze the methyl and benzoate esters. The crude product was dissolved in acetic anhydride and heated at 85 °C, which resulted in clean lactone formation. The reaction mixture was cooled to room temperature and then pyridine was added, which resulted in the acetylation of the remaining hydroxyl to give compounds **24a-c**. The 3,6-lactone forces the pyranosyl ring of glucuronic acid into a ^4^C_1_ conformation in which all *O-*substituents adopt an axial orientation. In accordance with this conformation, H-1 of glucuronic acid of **22a-c** shifted from 4.9 ppm to 5.5 ppm and appeared as a singlet. Furthermore, ^13^C−_1_H coupling constant at 2D HSQC off-decoupled spectrum shifted from 164 Hz to 174 Hz and 2D COSY spectrum showed a loss of correlation between H-1, H-2, and H-3, supporting a ^4^C_1_→^1^C_4_ conformational change (Fig. 3).

**Figure 3.**
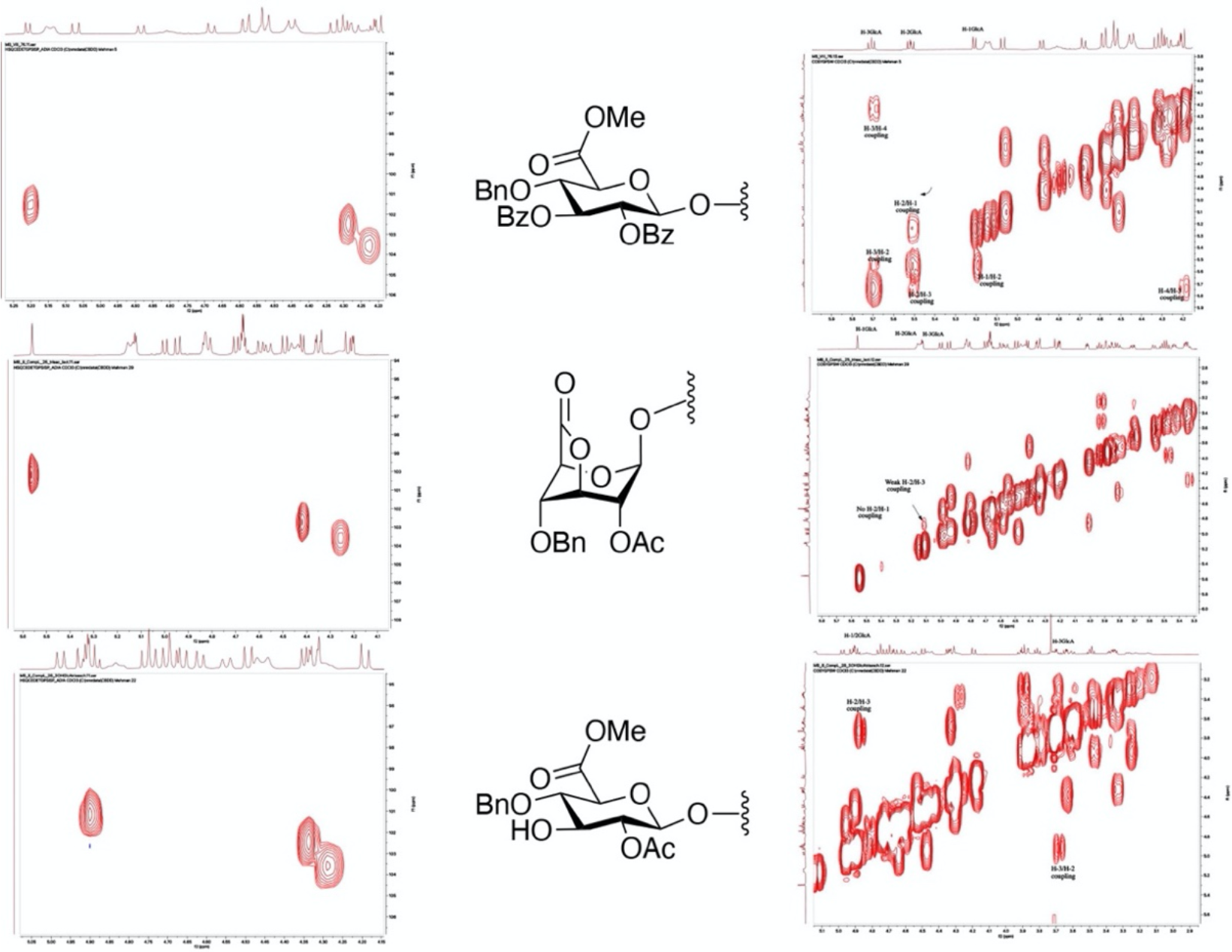
NMR characteristics of conformational change ^4^C_1_→^1^C_4_→^4^C_1_ of glucuronic acid pyranose ring induced by lactonization and delactonization.

Taking advantage of the ring strain of GlcA lactone, it was subjected to methanolysis using 100 mM methanolic solution of anhydrous sodium acetate resulting in the formation of a methyl ester and exposing a free C-3 hydroxyl at glucuronic acid moiety to give compounds **18a-c**. The resulting compounds were sulfated using SO_3_.Py complex in pyridine, followed by hydrogenation over Degussa type Pd(OH)_2_/C resulting in the removal of the benzyl and Cbz protecting groups and conversion of *N*-trichloroacetamides into acetamides to give the HNK-1 containing compounds **1-3**. Immediate deprotection of **18a-c** gave a similar ranger of compounds (**4-6**) lacking a sulfate ester. To examine the importance of the carboxylic acid of an HNK-1 epitope for auto-antibody binding, compounds **7** and **8** were prepared by hydrogenolysis of **18b** and its sulfated derivative over Degussa type Pd(OH)_2_/C followed by the reduction of the carboxylic acid methyl ester using NaBH_4_.

In addition to HNK-1 containing glycoconjugates, sialylated paragloboside (LM1) and sulfated galactosides have been implicated in auto-immune diseases (53-55), and therefore such derivatives were also prepared. Thus, treatment of **16b-c** with SO_3_.Py in pyridine followed by hydrogenation yielded sulfated paraglobosides **9** and **10** (Scheme 3). LM1 (**13**) and hexLM1 (**14**) were prepared by hydrogenolysis of **16b-c** to **11** and **12** followed by enzymatic sialylated using CMP-Neu5Ac and the microbial sialyl transferase PmST1.

### Glycan microarray development and screening

The synthetic compounds are ideally suited to examine binding properties of serum antibodies of MAG-neuropathy patients. It includes the HNK-1 alone (**1**) and this epitope presented on lacto-*N*-neotetraose (**2**) and lacto-N-neohexaose (**3**) typical of glycolipids, and thus make it possible to examine the importance of the underlying glycan for antibody binding. The compound series include similar compounds lacking the sulfate ester (**4-6**) and derivatives in which the HNK-1 epitope is replaced by sulfate (**9, 10**) or sialic acid (**13, 14**). Unnatural derivatives (**7, 8**) in which glucuronic acid was reduced to glucose were also synthesized to examine the importance of the carboxylic acid for antibody binding.

The synthetic glycans are equipped with an anomeric amino-pentyl linker allowing to construct a glycan microarray by piezoelectric printing on *N*-hydroxysuccinimde (NHS)-activated microarray glass slides. First, the glycan microarray was used to examine the binding selectivity of a mouse monoclonal anti-CD57 IgM antibody. The expression of HNK-1, which is also known as CD57, is a marker of a subset of natural killer (NK) cells that are in the final stages of maturation, and anti-CD57 antibodies are commonly employed to identify and isolate such cell populations (56). The anti-CD57 IgM antibody bound strongly to compounds **1-3**, confirming it requires a 3-*O*-sulfated glucuronic acid moiety for binding (Fig. 4A). Another HNK-1 carbohydrate-specific antibody (L2, IgG1) exhibited a similar binding profile (Fig. 4B). In each case, the responses for compounds **2a-c** were rather similar indicating the underlying glycan moiety has relatively little impact on binding.

**Figure 4.**
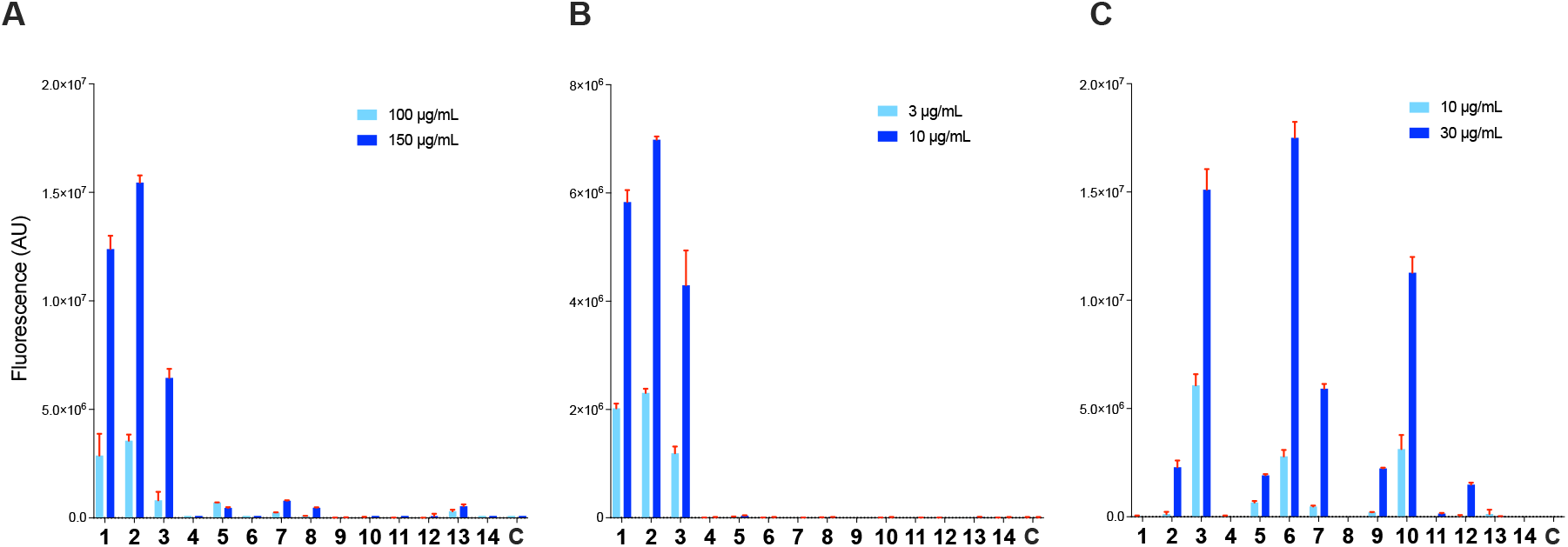
Microarray results of the synthetic library with mouse monoclonal IgM anti-human CD57 (**A**), mouse monoclonal IgG1 anti-human L2/HNK-1 carbohydrate (**B**), and galectin-3 (**C**) at the indicated concentrations. Bars represent the mean ± SD (n=4). C indicates buffer control.

Next, binding properties of serum antibodies from 10 patients with anti-MAG neuropathy and 3 healthy controls were examined. Antibody binding was detected using an appropriate anti-IgM secondary antibody tagged with AlexaFluor647. The healthy controls did not show substantial binding. The patients’ serum sample only recognized compounds carrying a glucuronic acid moiety. Surprisingly, differences were observed in the requirement of sulfation of the glucuronic acid moiety and the presence of the underlying oligo-LacNAc chain, resulting in three distinct groups of responders. One group of serum samples (SS02, SS08, SS09, SS10) exhibited a strong preference for a lacto-*neo*hexaose carrying an HNK-1 epitope (**3**) (Fig. 5A). Shorter sulfoglucuronosyl paragloboside **2**, and sulfated glucuronyl lactoside **1** showed much weaker binding. Corresponding compounds lacking a sulfate (**4, 6**, and **7**) exhibited no- or very weak reactivity. Thus, in these cases the serum antibodies have a strict requirement for HNK-1 epitopes presented on a lacto-*neo-*hexaose backbone which is typical of glycosphingolipids. In contrast, another group of serum samples (SS01 and SS07) showed a preference for glucuronic acid derivatives lacking a sulfate (**4, 6**, and **7**), and in these cases the underlying LacNAc moiety had only a small influence on antibody binding (Fig. 5B). The final group of serum samples recognized glucuronic acid derivatives with and without a sulfate. In one case (SS05), the two classes of compound were recognized with similar intensity with little preference for the presence of an extended LacNAc moiety. In other cases (SS03 and SS06), the HNK1 epitope presented on lacto-*neo*hexaose (**3**) exhibited the strongest binding, however, the corresponding compound lacking a sulfate did also show some reactivity. One sample had preference for a compound in which a glucuronic acid lacking sulfate (**6**) is presented on a lacto-*neo*hexaose backbone, but other glucuronic acid containing derivatives were also recognized (Fig. 5C).

**Figure 5.**
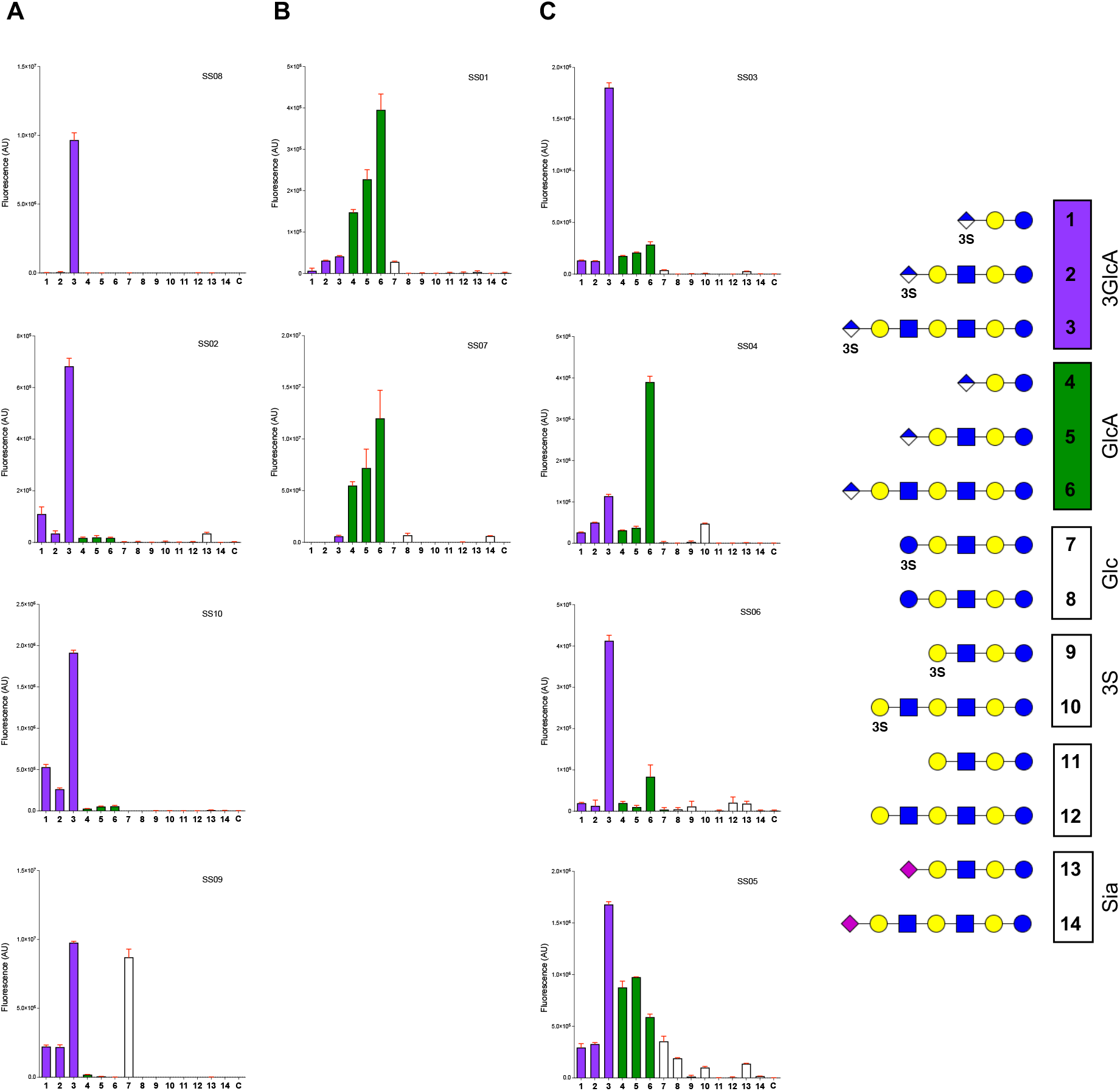
Microarray binding results of the synthetic library with serum antibodies of patients suffering from anti-MAG neuropathy divided over 3 groups, with a response mainly to HNK-1 (**A**), mainly to deS-HNK-1 (**B**), and a mixed response (**C**). Bars represent the mean ± SD (n=4). C indicates buffer control. Serum samples were diluted 1:250.

No correlation between reactivity in commercial Bühlmann ELISA assay using myelin associated glycoprotein and the newly developed microarray format was observed, which supports the notion that presentation of HNK-1 epitopes as part of a glycolipid and *N*-glycans results in different binding characteristics (Fig. S9).

The microarray was also employed to examine ligand requirements of several glycan binding proteins including P-and L-selectins and galectin-3. It has been reported that P- and L-selectin but not E-selectin can bind glycosphingolipids bearing HNK-1 (11). Microarray screening of L- and P-selectin resulted, however, only in very weak and promiscuous binding responses. It is possible that these proteins need to be presented in high multivalent forms to overcome low affinity monovalent binding. Galectin 3 gave interesting binding selectivities (Fig. 4C). As previously reported (57), it has a preference for extended LacNAc moieties, and strong binding was observed for lacto-*neo*hexaose **12**. 2,3-Sialylation of this compound (**14**) resulted in much weaker binding whereas modifications by 3-*O*-sulfate-glucuronic acid (**3**), glucuronic acid (**6**) or sulfate (**10**) were well tolerated. These findings indicate that the chemical nature of modification of the nonreducing galactoside of a poly-LacNAc chain can modulate galectin 3 binding, and notably a sulfate at a terminal C-3 position substantially increases binding.

## Conclusions

A chemoenzymatic methodology has been developed that can readily provide HNK-1 containing glycans derived from sulfoglucuronyl paraglobosides (**1-3**) and analogues lacking a sulfate ester (**4-6**). The approach also offered access to glycans in which glucuronic acid was replaced by sulfate (**9-10**) or sialic acid (**13-14**). The resulting glycans were printed as a microarray, which was employed to examine binding properties of serum antibodies of patients with anti-MAG neuropathy. Although the serum antibodies had a strict requirement for glucuronic acid, distinct groups were identified with unique binding characteristics. One group bound strongly to HNK-1 presented on a lacto-*neo*hexaose backbone supporting the notion that these antibodies target the corresponding glycosphingolipids. Another group of serum samples did not tolerate the presence of 3-*O*-sulfate at glucuronic acid, and in these cases, an extended LacNAc moiety did not substantially contribute to binding. It is possible that these antibodies recognize glycosphingolipids as well as *N*-glycans on glycoproteins such as MAG. Future efforts will focus on the synthesis of HNK-1 containing N-glycans to examine possible cross reactivities. A third group of serum samples bound to glucuronic acids with and without sulfate, and generally had a requirement for an extended LacNAc moiety. Thus, these serum antibodies also appear to target glycosphingolipids. The microarray screening highlights that anti-MAG neuropathy is not uniquely associated with serum antibodies targeting HNK-1 epitopes and may also involve glucuronyl containing glycoconjugates lacking sulfate.

ELISA and Western blotting using MAG are commonly employed to support a diagnosis of anti-MAG neuropathy in patients with related IgM monoclonal gammopathies (31). These methods give only 50-70% positive results for patients exhibiting a demyelinating and IgM monoclonal gammopathy related polyneuropathy. The array screening presented here indicates that oligosaccharides such as **1-3** and **4-6**, which are derived from glycosphingolipids, provide a more versatile substrate to investigate the presence of serum antibodies. The anti-MAG neuropathy is diverse with respect to clinical presentation and disease progression and the observation that the serum antibodies exhibit distinct binding patterns offers a prospect for more accurate disease diagnoses and stratification.

It is also the expectation that an understanding of binding properties of pathogenic serum IgM antibodies of anti-MAG neuropathy patients will give opportunities to develop more effective therapeutic strategies. The current therapeutic approach is based on treatment with an CD20^+^ B-cell-depleting antibody such as rituximab (58), which is non-selective and has only a minor therapeutic effect in a subset of patients. *In vitro* and *in vivo* studies have shown that polymers presenting HNK-1 epitopes can capture anti-MAG IgM antibodies and may provide a safer and more efficient therapeutic option (59, 60). The results presented here show that HNK-1 is a sub-optimal structure for binding pathogenic serum IgM antibodies of MAG neuropathy patients. The glycan array described here will make it possible to personalize such treatment by selecting a glycan structure optimal for antibody binding.

The HNK-1 epitope has been implicated in a multitude of biological processes such as cell recognition, adhesion, migration, synaptic plasticity, preferential motor re-innervation and regeneration after injury (61). Despite its importance, the molecular mechanisms by which HNK-1 containing glycoconjugates mediate function are poorly understood, which in part is due to a lack of well-defined HNK-1 containing glycoconjugates to examine interactions with glycan binding proteins and antibodies. Synthetic efforts have focused on this class of compounds, however, it has only resulted in small structural fragments or required very lengthy chemical approaches (27-30, 62). Here, we introduce a chemoenzymatic strategy that we coined neo-chemoenzymatic glycosylation, in which an oligo-LacNAc structure was assembled enzymatically and then subjected to protecting group manipulation to chemically install a 3-*O*-sulfate glucuronic acid moiety. The synthetic strategy is highly divergent and made it possible to synthesize from key intermediates other derivatives such a 3-*O*-sulfate galactosides and 2,3-linked sialosides. It addresses difficulties when specific glycosyl transferases or glycan modifying enzymes are not readily available. It is to be expected that neo-chemoenzymatic synthesis can be applied to the preparation of other glycoconjugates where targeted glycans cannot readily be obtained by enzymatic transformations alone. For example, an *N*-glycan obtained from egg yolk powder is an attractive starting material for enzymatic synthesis of multi-antennary glycans (47). An isolated *N*-glycan has also been used in chemical synthesis to provide more complex compounds (63). We anticipate that a combined use of an isolated *N*-glycan, enzymatic modification and chemical installation of terminal epitopes will, for example, give access to unique compounds such as HNK-1 containing *N*-glycans. Furthermore, chemoenzymatic approaches cannot easily be extended to carbohydrates of bacterial origin. The preparation of these synthetically challenging targets may be achieved by the application of neo-chemoenzymatic strategy where glycans isolated from a natural source can be turned into structurally more complex carbohydrate architectures by a combination of enzymatic and chemical manipulations.

See *SI Appendix* for synthetic procedures, MS and NMR characterization, and NMR spectra of compounds. Patient serum details, assay procedures, supplementary glycan microarray and ELISA data are provided.

## Data availability

All supporting data is available in *SI Appendix*.

## Acknowledgments and funding sources

This research was supported by TOP-PUNT grant 718.015.003 of the Netherlands Organization for Scientific Research (to G.J.B.) and ERC-ADG grant 101020769 of the European Commission (to G.J.B.). R.H. was funded by the GBS/CIDP Foundation International and the T2B collaboration project funded by PPP Allowance made available by Top Sector Life Sciences & Health to Samenwerkende Gezondheidsfondsen (SGF) under project number LSHM18055-SGF.

---

**Scheme 1.**
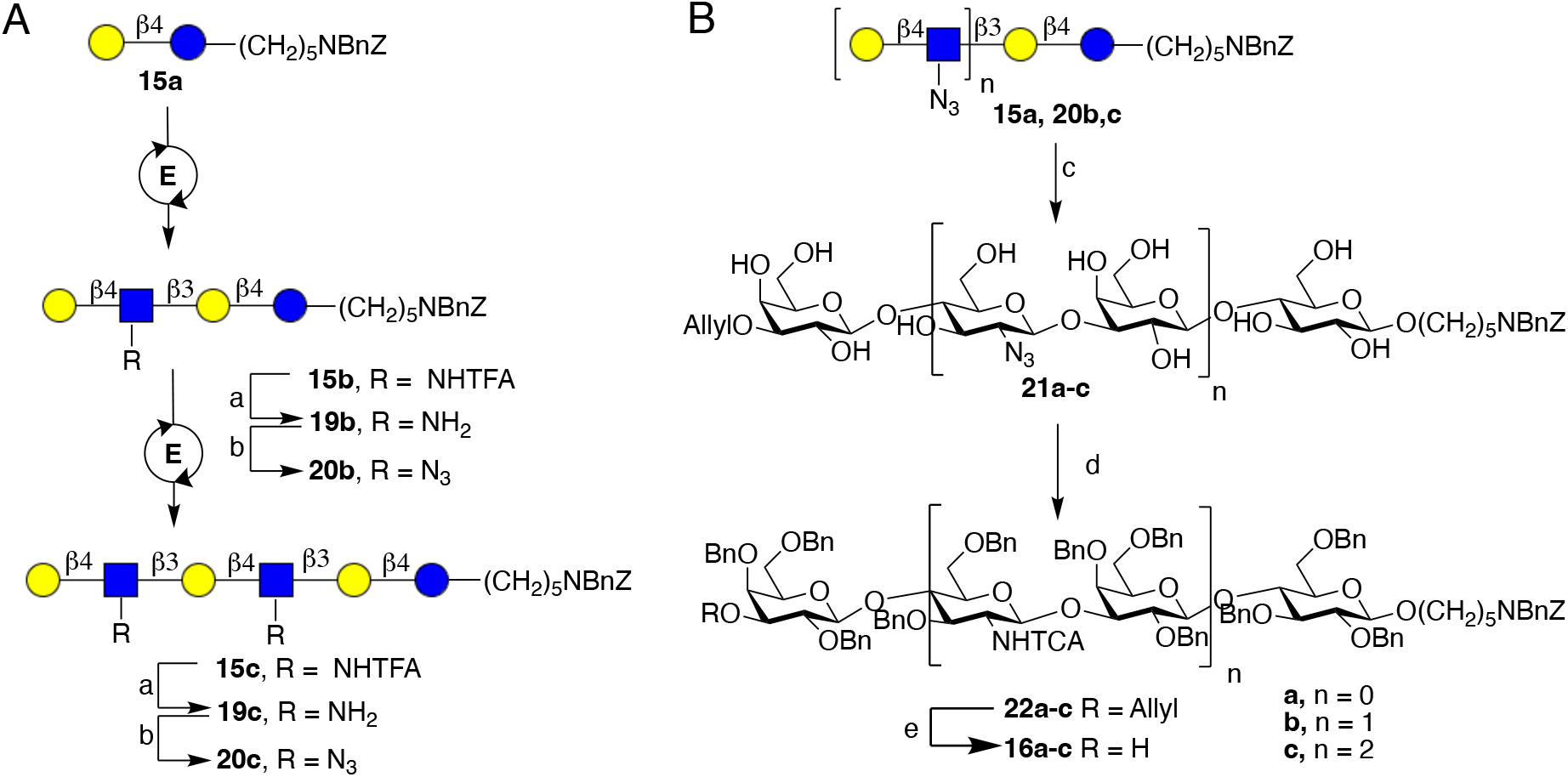
Chemoenzymatic synthesis of precursor compounds 20a-c. A. Enzymatic extension of lactoside using recombinant glycosyltransferases and unnatural sugar-nucleotides. Conditions: E-enzyme module for installation: UDP-GlcNHTFA, Tris·HCl buffer (pH = 7.5), MgCl_2_ (10 mM), KCl (100 mM), DTT (1 mM), HpB3GnT (1 mg/mmol acceptor) and CIAP (1U) for glucosamine moiety. UDP-Gal, Tris·HCl buffer (pH = 7.8), MgCl_2_ (10 mM), BSA (0.1% w/w acceptor), NmLgtB (1 mg/mmol acceptor) and CIAP (1U) for galactose transfer. ***a***. NaOH (pH = 10), ***b***. ImSO_2_N_3_·HCl (2 eq), CuSO_4_ (0.05 eq), K_2_CO_3_ (2.5 eq), H_2_O-MeOH. B. Preparation of acceptor oligosaccharides using protecting group manipulations. Conditions: ***c***. 1. Bu_2_SnO (1.2 eq), MeOH, reflux, 2. Allyl bromide (3 eq), DMF, TBAI (0.4 eq), ***d***. 1. NaH (2 eq/OH), BnBr (1.5 eq/OH), DMF, 0 °C, 2. PMe_3_ (5 eq/N_3_), THF-H_2_O, 3. Cl_3_CCOCl (1.5 eq/NH_2_), Et_3_N, DCM. ***e***. PdCl_2_ (0.2 eq), CHCl_3_/MeOH.

**Scheme 2.**
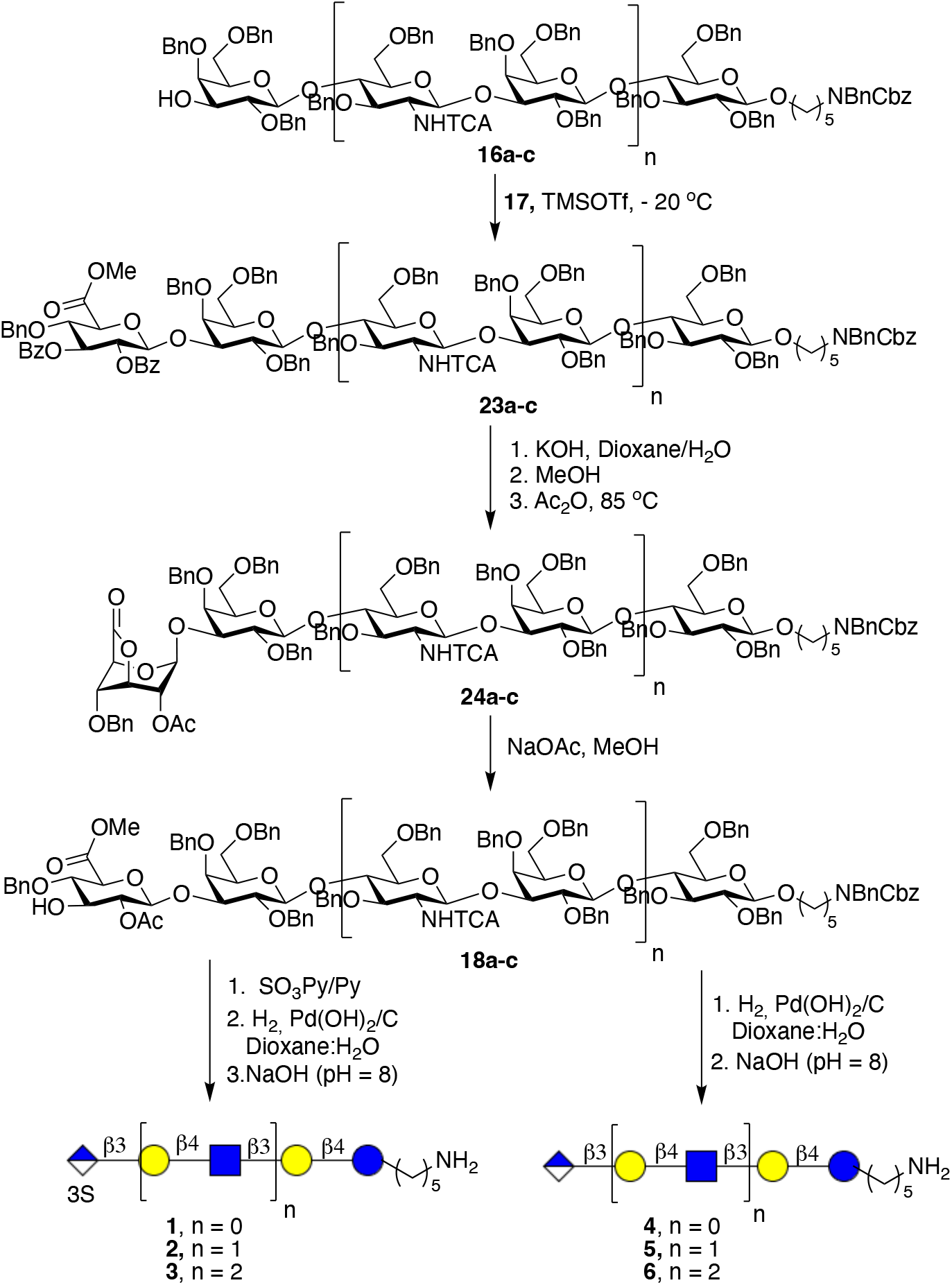
Glycosylation of acceptor oligosaccharides with glucuronic acid donor (**17**) and subsequent protecting group manipulations for the preparation of glycans having HNK-1 epitope (**1, 2, 3**) and its non-sulfated derivatives (**4, 5, 6**).

**Scheme 3.**
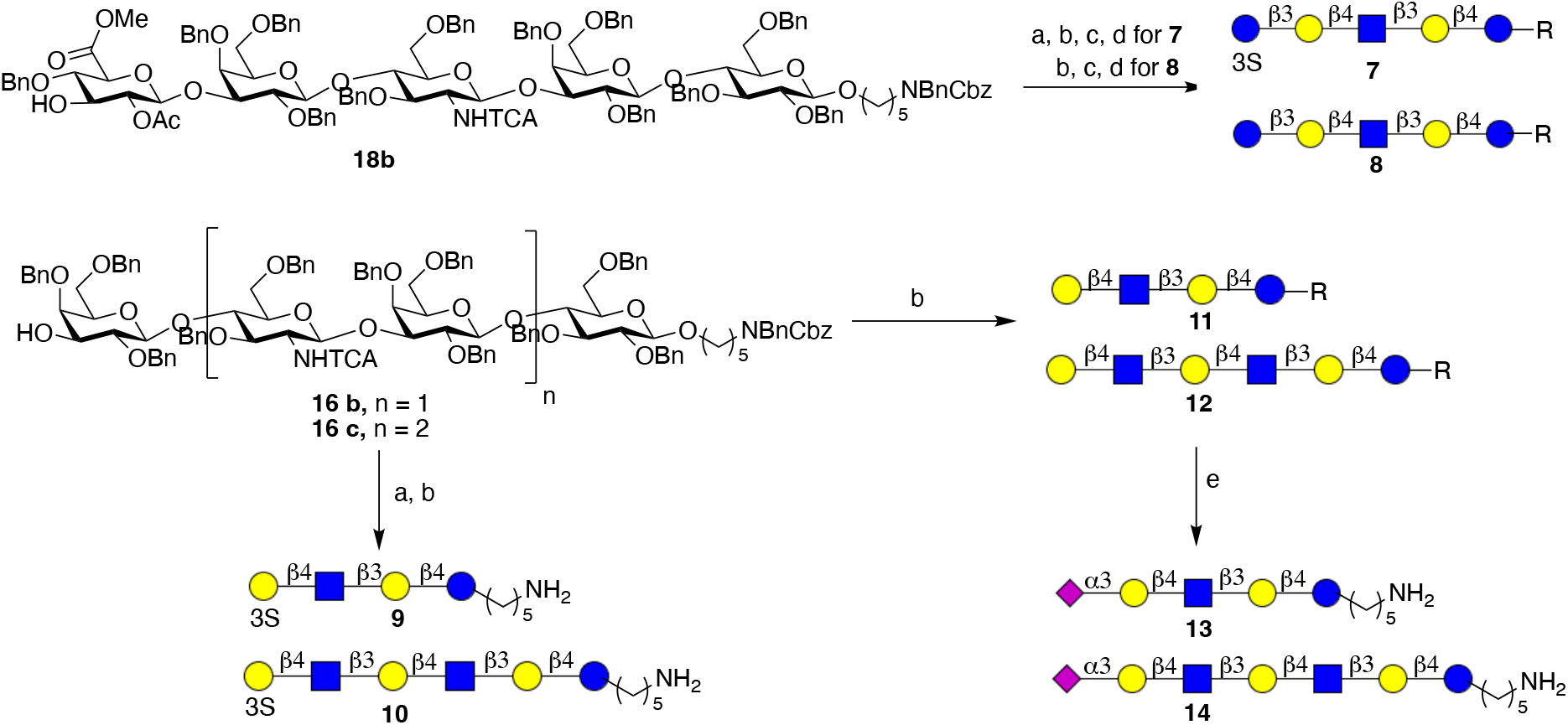
Synthesis of control compounds **7-14** from common precursors. Conditions: **a** – SO_3_Py/Py, **b** – H_2_/Pd(OH)_2_/C, dioxane:H_2_O, **c** – NaBH_4_/MeOH, **d** – NaOH (pH = 8), **e** – PmST1 (1% w/substrate), CMP-Neu5Ac (1.5 eq.), Tris (pH = 7.8, 100 mM), MgCl_2_ (10 mM), CIAP (0.1U), 37 °C.

