## Supplementary material for "Synthetic HNK-1 containing glycans provide insight into binding properties of serum antibodies from MAG-neuropathy patients": SI

Table of contents


1. Materials and methods
   Chemicals ........................................................................ S3
   Enzyme expression ........................................................................ S3
2. NMR nomenclature and analysis of target glycans
   Glycan chain numbering (Figure S1) ........................................................................ S4
   Key steps for structural elucidation (Figures S2-S6) ........................................................................ S4
3. Chemoenzymatic synthesis of oligosaccharide acceptors with various length
   Figure S7 ........................................................................ S9
   General protocol for the installation of unit using UDP-GlcNHTFA and
   UDP-Gal in combination with HpB3GnT and LgtB, respectively ........................................................................ S9
4. Chemical modification of enzymatically assembled di- and oligosaccharides
   General protocol for TFA removal ........................................................................ S13
   General procedure for azido-transfer reaction ........................................................................ S13
   General protocol for tin mediated allylation ........................................................................ S14
   General protocol for benzylation ........................................................................ S14
   General procedure for conversion of azide into NHTCA ........................................................................ S14
   General procedure for allyl ether removal ........................................................................ S15
5. Synthesis of target glycans with diverse terminal epitopes
   General procedure for glycosylation of lactosyl acceptors with glucuronic acid donor ........ S29
   General procedure for saponification and intralactone formation reactions ........................ S29
   General protocol for methanolysis of lactones ........................................................................ S30
   General protocol for *O*-sulfation ........................................................................ S30
   General procedure for global deprotection ........................................................................ S31
   General procedure for the sialylation of Lacto-N-neotetraose and
   Lacto-N-neohexaose ........................................................................ S54
6. Synthesis of key building blocks
   Chemoenzymatic synthesis of UDP-GlcNHTFA nucleotide donor (Scheme S1) ................ S57
   Synthesis of glucuronate phosphate donor (Scheme S2) ................................................ S59
7. Microarray
   Serology serum samples (Table S1) ........................................................................ S64
   Glycan array printing ........................................................................ S64
   Microarray binding assays and serum sample screening ................................................ S65
   Figure S8 ........................................................................ S66
   Figure S9 ........................................................................ S67
8. References ........................................................................ S68
9. Copies of NMR and MS spectra ........................................................................ S68

#### 1. Materials and methods

##### Chemicals

Unless otherwised stated, all chemicals were purchased from Sigma-Aldrich. Monosaccharide for building block synthesis were purchased from Carbosynth Limited (UK). Acetonitrile, dichloromethane, toluene, tetrahydrofuran and N,N-dimethylformamide used for synthesis were anhydrous grade and obtained from a solvent purifier (MB SPS 5). Other organic solvents for reactions were obtained from Biosolve Chemie. Organic solvents for work-up procedures were technical grade and obtained from VWR Chemicals. Deuterated solvents for NMR experiments were obtained from Cambridge Isotope Laboratories.

$^{1}$H and $^{13}$C NMR spectra were recorded on Bruker 600 UltraShield. Chemical shifts are reported in parts per million (ppm) relative to $CDCl_3$ as the internal standard. NMR data are presented as follows: Chemical shift, multiplicity (s = singlet, d = doublet, t = triplet, dd = doublet of doublets, m = multiplet, b = broad); coupling constants are reported in Hertz (Hz). All NMR signals were assigned on the basis of $^{1}$H NMR, COSY, HSQC, HMBC, TOCSY and NOESY experiments. Mass spectra were recorded on either on an Applied Biosystems SCIEX MALDI-TOF/TOF 5800 mass spectrometer, a Shimadzu Biotech Axima-CFR MALDI-TOF, or a high resolution 6500 Q-ToF MS (Agilent). Column chromatography was performed on silica gel G60 (Silicycle, 60-200 µm, 60 Å). TLC analysis was performed using precoated silica gel 60 F-254 plates (Merck) with detection by UV light (254 nm) where applicable, and by charring with 10% sulfuric acid in ethanol or a p-anisaldehyde staining solutionn ethanol (1% v/v). Unless specified otherwise, all moisture sensitive reactions were carried out under argon atmosphere and in the presence of activated molecular sieves. Unless otherwise stated, all reactions were carried out at room temperature (RT) in glassware with magnetic stirring. Solutions in organic solvents were dried with $Na_2SO_4$ and concentrated at 40 °C/2 kPa. Molecular sieves were flame-dried in vacuo immediate prior to use.

##### Enzyme expression

Glycosyl transferases were expressed according to published protocols (1), as soluble proteins in E. *coli* BL21 cell culture. All proteins contain a histidine tag, which allowed purification by $Ni^{2+}$-NTA affinity column chromatography. After concentration and buffer exchange, expression yield for enzymes HpB3GnT and NmLgtB were calculated as 7 mg/L and 2 mg/L of bacterial culture, respectively.

#### 2. NMR nomenclature and analysis of target glycans

##### Glycan chain numbering

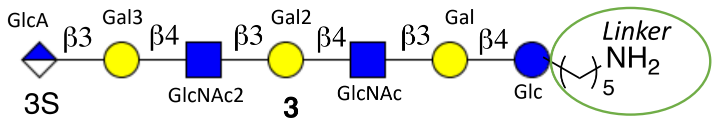


**Figure S1.** Labeling of monosaccharides for NMR assignment.

Numbering of glycan chain was from reducing end to non-reducing terminus. Assignments of chemical shifts of each proton of protected glycans started from the most downfield proton continuing until the most upfield proton. For deprotected glycans, carbohydrate and non-carbohydrate protons were assigned separately. Each monosaccharide was assigned separately (H-1 → H-6, for all, except for GlcA H-1 → H-5 and sialic acid H-1 → H-9).

##### Key steps for structural elucidation

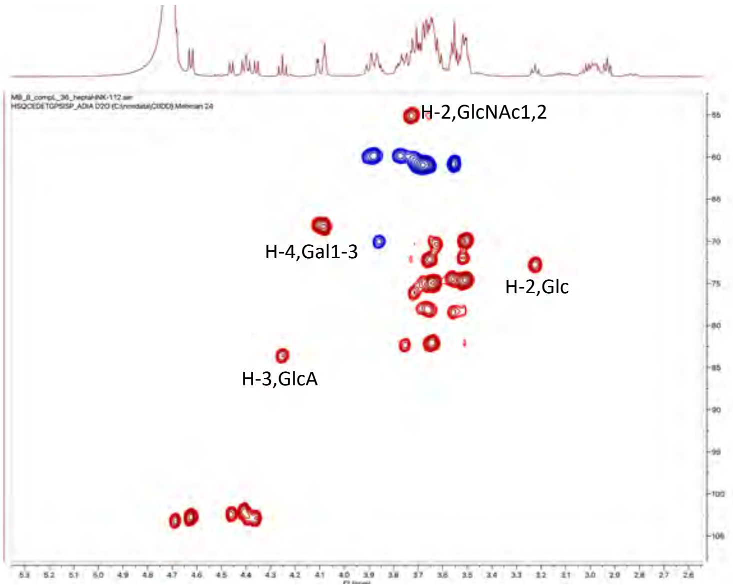


Glycan **3** and other glycans bearing similar structural motifs were assigned by employing 1D and 2D NMR spectroscopy techniques.

H-$1_{GlcNAc}$ is usually more downfield than the other anomeric protons and its $^1$H-$^1$H correlation with H-$2_{GlcNAc}$, which has a diagnostic chemical shift at 3.72 ppm ($^{13}$C = 55.5 ppm).

Another characteristic peak can be assigned to H-$2_{Glc}$ at 3.21 ppm, triplet with coupling constant of $J$ = 8.9 Hz. H-$1_{Glc}$ can be assigned from its correlation with H-2. H-$2_{Glc}$ gives another obvious correlation with H-$3_{Glc}$ at $^1$H-$^1$H spectrum. Furthermore, H-$3_{GlcA}$ is shifted to more downfield region as a result of deshielding effect of sulfate ester at C-3 of glucuronic acid. $^1H_{GlcA}$ can be assigned from H-$3_{GlcA}$→H-$2_{GlcA}$→H-$1_{GlcA}$ using COSY spectrum (Fig. S2).

Similarly, H-4 of galactose is usually one of the diagnostic protons, which commonly appears around 3.80 – 4.20 and characterized as a duplet with J ≈ 3 Hz. The most downfield H-$4_{Gal}$ can be assigned to H-4 of Gal3, since it has a TOCSY correlation with the most downfield H-$1_{Gal}$, resulting from the glucuronic acid which is linked to C-3 of penultimate Gal through glycosidic linkage. Furthermore, H-$3_{GlcA}$ has a through space correlation with H-$1_{GlcA}$, which has a similar NOE correlation with H-$3_{Gal}$ of Gal3. Similarly, H-$3_{Gal3}$→H-$1_{Gal3}$ through space correlation can be observed confirming the correct assignment of anomeric proton of Gal3 and correct position of glycosidic linkage between GlcA and Gal3. Other protons of a glycan chain can be assigned using the combination of TOCSY-HMBC-HSQC spectra. Anomeric protons of N-acetylglucosamine, glucuronic acid, glucose and penultimate galactose could be assigned using correlation with diagnostic peaks. On the other hand, in order to differentiate H-$1_{Gal1}$ and H-$1_{Gal2}$ the superimposition of 2D spectra such as HMBC-HSQC was applied. H-$1_{Glc}$ has a multiple bond correlation with $^{13}\underline{C}H_2$ of linker (which can also be observed from NOESY $^1$H-$1_{Glc}$ – OC$\underline{H_2}$linker), confirming its correct assignment. Moreover, correlation between H-$1_{Gal2,3}$ - $^{13}$C-$4_{GlcNAc}$ and H-$1_{Gal1}$ – $^{13}$C-$4_{Glc}$ indicates the chemical shift of Gal2 and Gal1 at 4.39 ppm and 4.35 ppm respectively.

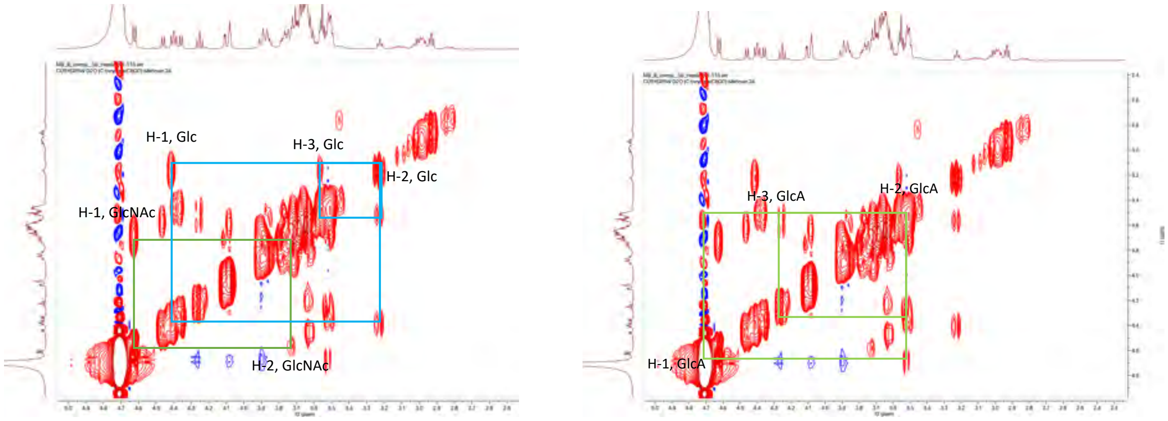


**Figure S2.** $^1$H-$^1$H COSY correlation between H-1 and H-2, H-3 protons of pyranose ring.

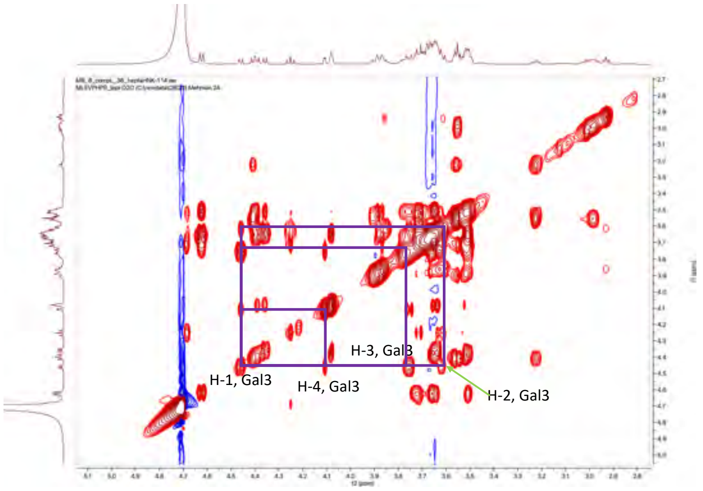


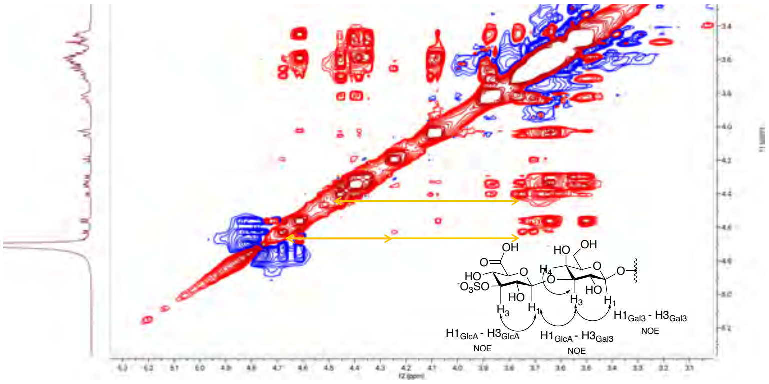


**Figure S3.** $^{1}H$-$^{1}H$ TOCSY and NOESY spectra of compound **3**.

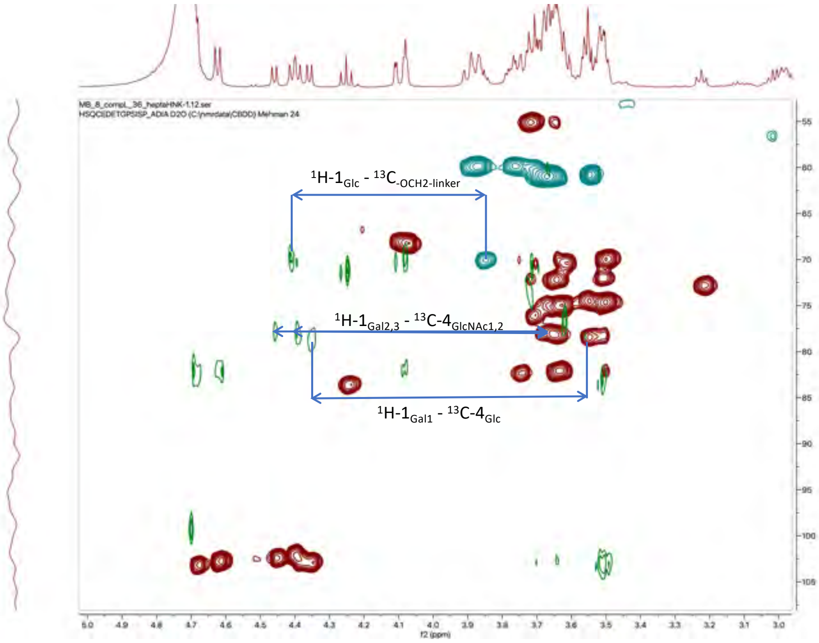


**Figure S4.** Superimposition of HMBC and HSQC spectra of compound **3**.

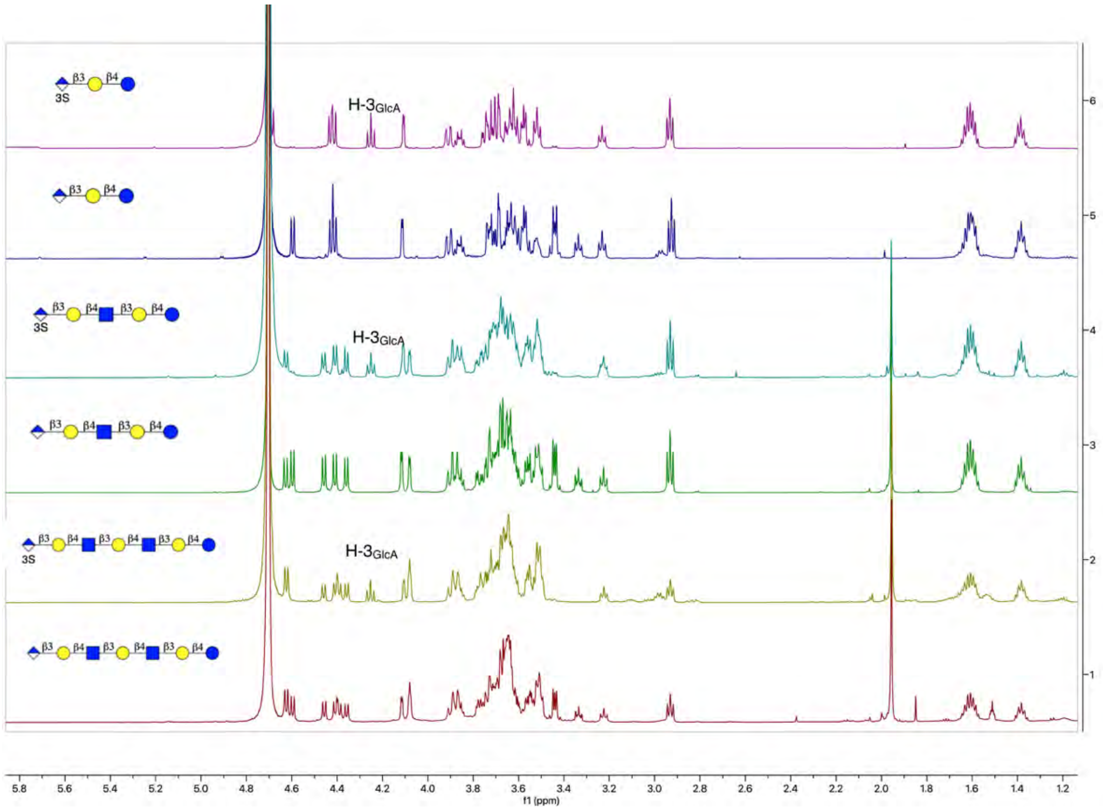


**Figure S5.** Stacked 1D $^1$H-NMR spectra of target compounds **1-6**. Illustration of deshielding effect of sulfate ester.

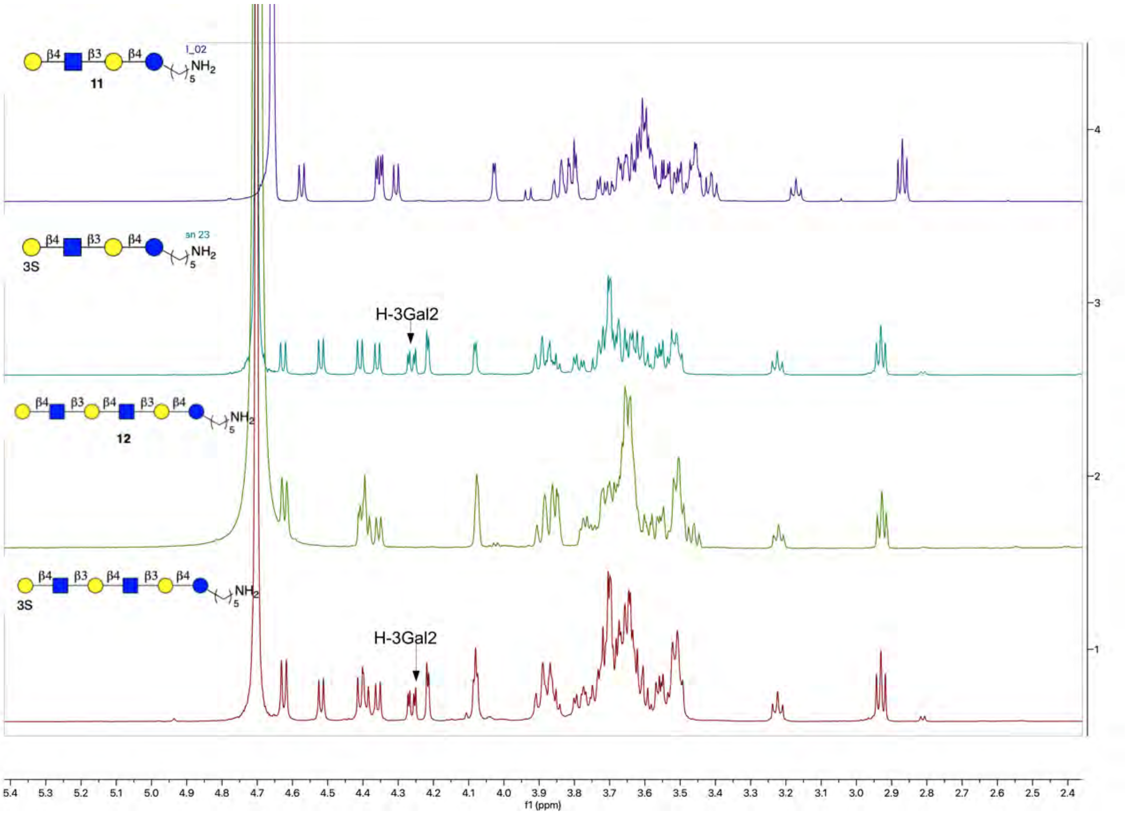


**Figure S6.** Stacked 1D $^1$H-NMR spectra of control compounds **9-12**.

#### 3. Chemoenzymatic synthesis of oligosaccharide acceptors with various length

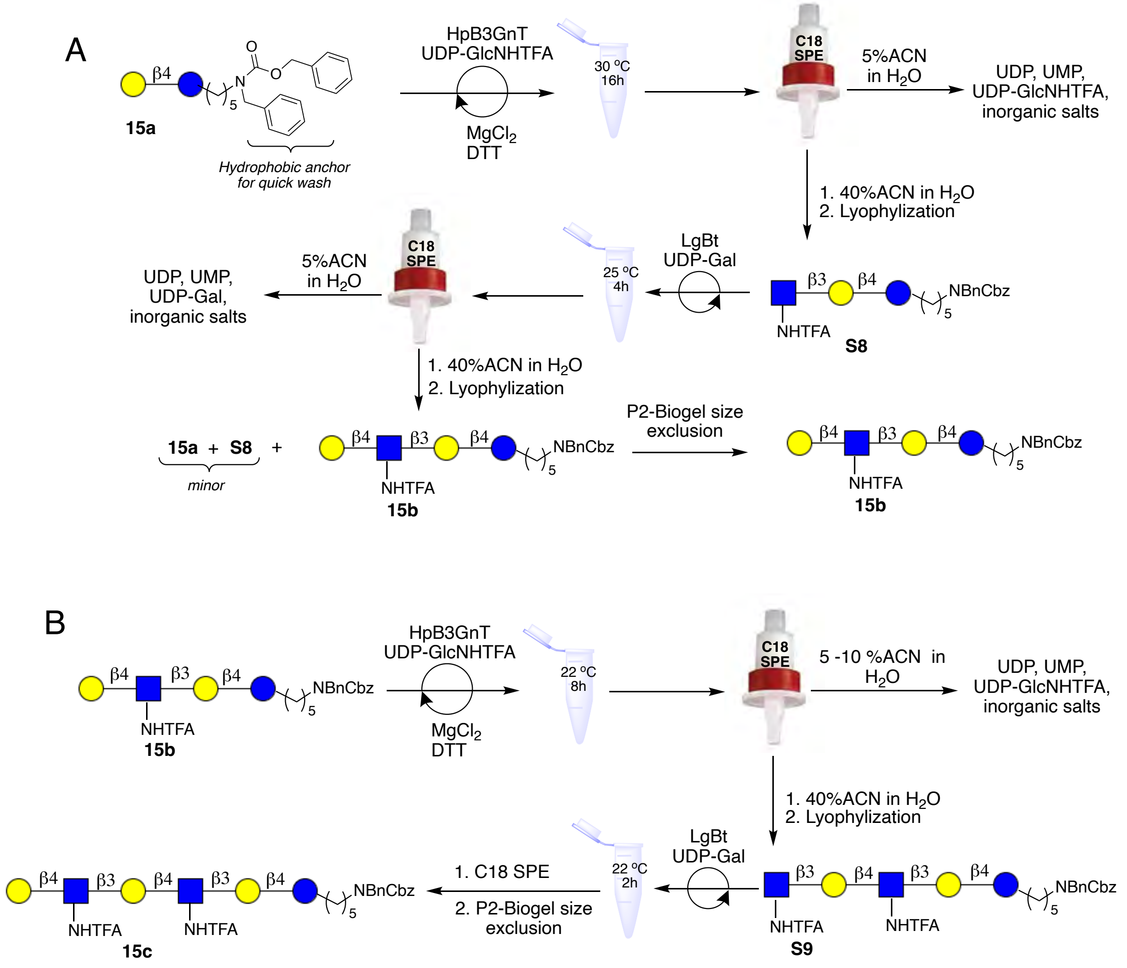


**Figure S7.** Schematic visualization of enzymatic flowchart for the installation of one (A) and two (B) lactosamine moiety.

**General protocol for the installation of unit using UDP-GlcNHTFA and UDP-Gal in combination with HpB3GnT and LgtB, respectively**

UDP-GlcNHTFA (1.5 eq) and acceptor **(15a, 15b)** were dissolved in Tris buffer (pH = 7.8, 100 mM) containing 10 mM $MgCl_2$ and 1 mM DTT to give a final concentration of 10 mM of acceptor. To this solution, Hpb3GlcNAcT (1%, w/w) and CIAP (10 mU) were added and the reaction mixture was incubated at 30 °C overnight in an incubator with shaking. MALDI-TOF MS was used to monitor the progress of the reaction and in case of incomplete conversion an extra portion of UDP-GlcNHTFA and B3GnT were added. Upon completion of the enzymatic transformation, the reaction mixture was centrifuged and supernatant was decanted using a 1 mL syringe and passed through Sep-Pak C18-reverse phase cartridge by first washing with 5% aqueous acetonitrile to remove inorganic salts and nucleotide bases, followed by elution with 40% aqueous acetonitrile to obtain the desired intermediate compounds. Fractions containing product were pooled and freeze-dried. The resulting residue was redissolved in water to which Tris buffer (100 mM, pH = 7.5), UDP-Gal (1.5 eq assuming quantitative transfer of GlcNHTFA), $MgCl_2$ (10 mM), CIAP (0.1 U) and NmLgtB enzyme were added. The reaction mixture was incubated at 25 $^oC$ with an occasional shaking. After several hours, MALDI-TOF MS analysis indicated the completion of galactose transfer. After centrifugation, the mixture was passed through a Sep-Pak C18 cartridge and washed with low concentration of acetonitrile and eluted with 40% ACN. Fractions containing product were combined and lyophilized. The crude product was applied to a Biogel-P-2 size exclusion column and eluted with 10 mM aqueous $NH_4HCO_3$. Product containing fractions were analyzed by MALDI-TOF MS, concentrated, lyophilized and used in the subsequent steps. Titled compounds were analyzed by NMR and MS.

***N*-(Benzyl)-benzyloxycarbonyl-5-aminopentyl *β*-D-galactopyranosyl-(1→4)-*β*-D-glucopyranoside (15a).**

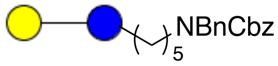


Compound **1** was synthesized in a multi-gram scale as described previously (2). MALDI-TOFMS *m/z* $C_{32}H_{45}NO_{13}$ $(M+Na)^+$ calcd 674.2789, found 674.5286.

$^1$H NMR (600 MHz, $D_2O$) δ: (non carbohydrate): 7.32 – 6.88 (m, 10H, Ar-H), 4.98 (d, $J$ = 91.9 Hz, 2H, *C*$H_2$ Cbz), 4.35 – 4.26 (2H, -C*H*$_2$NPh) 3.65 (-OCH*H* linker), 3.40 (-OC*H*H linker), 3.08 (t, $J$ = 7.4 Hz, 2H, -$CH_2$ linker), 1.41 – 1.29 (m, 4H, -$CH_2$ linker), 1.11 – 1.03 (m, 2H, -$CH_2$ linker).

$^{13}$C NMR (151 MHz, $D_2O$): $\delta$ (non carbohydrate): 129.8 – 126.5 (C Ar), 70.1 (-O*C*$H_2$ linker), 67.3 (*C*$H_2$ Cbz), 50.0 (-*C*$H_2$NPh), 47.2 (-*C*$H_2$ linker), 28.6 (-*C*$H_2$ linker), 27.1 (-*C*$H_2$ linker), 22.5 (-*C*$H_2$ linker).

|  | H1 | H2 | H3 | H4 | H5 | H6 |
|---|---|---|---|---|---|---|
| Glc | 4.28 | 3.22 | 3.56 | 3.57 | 3.43 | 3.84 – 3.73 |
| Gal | 4.37<br>d, $J$ = 7.6 Hz | 3.49,<br>t, $J$ = 8.7 Hz | 3.59 | 3.86,<br>d, $J$ = 2.7 Hz | 3.65 | 3.72 – 3.68 |

|  | C1 | C2 | C3 | C4 | C5 | C6 |
|---|---|---|---|---|---|---|
| Glc | 101.98 | 73.04 | 74.41 | 78.32 | 74.74 | 60.04 |
| Gal | 102.99 | 70.58 | 72.52 | 68.48 | 75.18 | 61.05 |

***N*-(Benzyl)-benzyloxycarbonyl-5-aminopentyl *β*-D-galactopyranosyl)-(1→4)-(2-trifluoroacetamido-2-deoxy-*β*-D-glucopyranosyl)-(1→3)-(*β*-D-galactopyranosyl)-(1→4)-*β*-D-glucopyranoside (15b).**

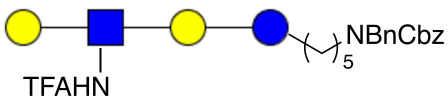


The general protocol for LacNHTFA installation was followed to obtain compound **15b** (480 mg, 88%) from **15a** (327 mg, 0.5 mmol) using UDP-GlcNHTFA (500 mg, 0.74 mmol) and UDP-Gal (440 mg, 0.75 mmol). MALDI-TOF MS *m/z* $C_{46}H_{65}F_3N_2O_{23}$ $(M+Na)^+$ 1093.3828 found 1093.5448.

$^1$H NMR (600 MHz, $D_2O$) δ: (non carbohydrate): 7.44 – 7.14 (m, 10H, Ar-H), 5.07 (d, $J$ = 29.1 Hz, 2H, $CH_2$, Cbz), 4.49 - 4.41 (2H, $CH_2$, -NC$\underline{H_2}$Ph), 3.73 (-OC$\underline{H}$H linker), 3.52 (-OCH$\underline{H}$ linker), 3.21 (m, -$CH_2$ linker), 1.54 – 1.40 (m, 4H, -$CH_2$ linker), 1.25 – 1.15 (m, 2H, -$CH_2$ linker).

$^{13}$C NMR (151 MHz, $D_2O$): $\delta$ (non carbohydrate): 22.5 (-$\underline{C}H_2$ linker), 27.2 (-$\underline{C}H_2$ linker), 28.9 (-$\underline{C}H_2$ linker), 47.0 (-$\underline{C}H_2$, linker), 50.4 (-N$\underline{C}H_2$Ph), 67.3 (-$\underline{C}H_2$, Cbz), 69.6 (-O$\underline{C}H_2$, linker), 125.9 – 129.7 (C Ar).

|  | H1 | H2 | H3 | H4 | H5 | H6 |
|---|---|---|---|---|---|---|
| Glc | 4.31 | 3.17 | 3.52 | 3.52 | 3.45 | 3.77 |
| Gal | 4.33<br>d, $J$ = 7.6 Hz | 3.48 | 3.63 | 4.09<br>d, $J$ = 3.2 Hz | na | 3.71 – 3.62 |
| GlcNAc | 4.73 | 3.79 | 3.51 | 3.68 | 3.60 | 3.87 – 3.82 |
| Gal(2) | 4.36<br>d, $J$ = 8.5 Hz | 3.44 | 3.56 | 3.84<br>d, $J$ = 3.1 Hz | na | 3.71 – 3.62 |

| | C1 | C2 | C3 | C4 | C5 | C6 |
|---|---|---|---|---|---|---|
| Glc | 102.4 | 72.9 | 74.6 | 78.8 | 74.8 | 59.7 |
| Gal | 102.8 | 69.7 | 82.7 | 68.3 | na | 60.8 |
| GlcNAc | 101.8 | 56.0 | 74.8 | 77.8 | 75.2 | 60.8 |
| Gal(2) | 103.1 | 71.2 | 72.4 | 68.5 | na | 3.90 – 3.62 |

**N-(Benzyl)-benzyloxycarbonyl-5-aminopentyl β-D-galactopyranosyl-(1→4)-2-trifluoroacetamido-2-deoxy-β-D-glucopyranosyl-(1→3)-β-D-galactopyranosyl-(1→4)-2-trifluoroacetamido-2-deoxy-β-D-glucopyranosyl-(1→3)-β-D-galactopyranosyl-(1→4)-β-D-glucopyranoside (15c).**

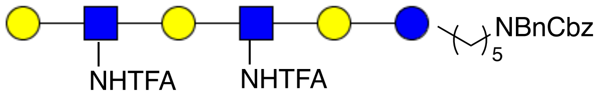


Compound **15b** (200 mg, 0.19 mmol) was reacted with UDP-GlcNHTFA (191 mg, 0.28 mmol) and UDP-Gal (164 mg, 0.28 mmol) according to the general protocol for LacNHTFA installation to yield compound **15c** (250 mg, 88%). MALDI-TOF MS *m/z* $C_{60}H_{85}F_6N_3O_{33}$ $(M+Na)^+$ 1512.4867 found 1512.6962, $(M+K)^+$ 1528.4607 found 1528.6309.

$^1$H NMR (600 MHz, $D_2O$) δ: (non carbohydrate): 7.30 – 6.88 (m, 10H, Ar-H), 6.80 (d, *J* = 9.8 Hz, 1H, -N<u>H</u>TFA), 6.74 (d, J = 9.0 Hz, 1H, -N<u>H</u>TFA), 4.93 (d, J = 92.5 Hz, 2H, $CH_2$, Cbz), 4.32 - 4.30 (2H, $CH_2$, NBn), 3.64 (-OC<u>*H*</u>H linker), 3.44 (-OCH<u>*H*</u> linker), 2.66 (t, J = 7.8 Hz, 2H, -$CH_2$, linker), 1.45 – 1.32 (m, 4H, -$CH_2$, linker), 1.14 – 1.05 (m, 2H, -$CH_2$ linker).

$^{13}$C NMR (151 MHz, $D_2O$): $\delta$ (non carbohydrate): 168.0 (-NH<u>*C*</u>O$CF_3$), 127.2 – 125.3 ($C_{Ar}$) 70.7 (-O$CH_2$ linker), 67.1 (<u>C</u>$H_2$ Cbz), 48.9 ($CH_2$ NBn ), 46.9 (-$CH_2$ linker), 29.1 (-$CH_2$ linker), 27.2 (-$CH_2$ linker), 23.2 (-$CH_2$ linker).

| | H1 | H2 | H3 | H4 | H5 | H6 |
|---|---|---|---|---|---|---|
| Glc | 4.26 | 3.13 | 3.49 | 3.48 | 3.50 | 3.80 – 3.71 |
| Gal | 4.35<br>d, *J* = 8.1 Hz | 3.46 | 3.60 | 4.04,<br>d, *J* = 2.8 Hz | na | 3.68 – 3.57 |
| GlcNAc | 4.68 | 3.74 | 3.57 | 3.61 | 3.47 | 3.80 – 3.71 |
| Gal(2) | 4.32<br>d, *J* = 8.4 Hz | 3.42 | 3.69 | 3.96<br>d, *J* = 2.5 Hz | na | 3.68 – 3.57 |
| GlcNAc2 | 4.68 | 3.74 | 3.58 | 3.61 | 3.47 | 3.80 – 3.71 |
| Gal (3) | 4.29<br>d, *J* = 8.1 Hz | 3.25 | 3.63 | 3.78 | na | 3.68 – 3.57 |

| | H1 | H2 | H3 | H4 | H5 | H6 |
|---|---|---|---|---|---|---|
| Glc | 102.1 | 72.3 | 74.7 | 78.2 | 74.4 | 60.2 |
| Gal | 102.8 | 69.7 | 82.4 | 68.1 | na | 60.8 |
| GlcNAc | 101.7 | 55.6 | 74.9 | 78.1 | 74.8 | 59.8 |
| Gal(2) | 102.7 | 71.3 | 72.9 | 74.4 | na | 60.8 |
| GlcNAc2 | 101.7 | 55.6 | 74.9 | 78.1 | 74.8 | 59.8 |
| Gal (3) | 102.5 | 71.2 | 72.3 | 68.4 | na | 60.8 |

#### 4. Chemical modification of enzymatically assembled di- and oligosaccharides

**General protocol for TFA removal.** Compounds **15b** and **15c** were dissolved in water and the pH of the resulting solutions were adjusted to 10. The reaction mixtures was incubated at 37 °C and progress of reactions were monitored by MALDI-TOF MS. Upon completion of the deacylation, the pH of the reaction was brought to neutral by dropwise addition of 1M HCl solution and then the samples were freeze-dried. Crude products were applied to a Biogel-P-2 size exclusion column and eluted with 100 mM aqueous $NH_4HCO_3$. Carbohydrate containing fractions were identified by MALDI-TOF MS and fractions containing products were pooled and lyophilized.

**General procedure for azido-transfer reaction.** To a solution of compound **19b** and **19c** in water, $K_2CO_3$ (10 eq), imidazole-1-sulfonyl azide hydrogen chloride (10 eq) (prepared following the protocol (3), and cat. amount of $CuSO_4{\cdot}5H_2O$ were added and the reaction mixture was incubated at 37 °C for 2 h. The conversion was monitored by MALDI-TOF MS and in case starting material was remaining an additional portion of diazo-transfer reagent was added followed by adjustment of pH to 8. The reaction mixture was further incubated until full conversion was observed by mass spectrometry and TLC analyses. Insoluble solid particles were removed by centrifugation and the supernatant was lyophilized. The resulting material was dissolved in water and applied onto benchtop C18 beads, which were washed with 10% acetonitrile in water followed by elution with 50% aqueous acetonitrile. Fractions containing products were collected and freeze-dried using $H_2O$:$^t$BuOH (3:1, v/v) as solvent mixture.

**General protocol for tin mediated allylation.** Compounds **15a**, **20b-c** and $Bu_2SnO$ (1.5 eq) were suspended in dry methanol and heated under reflux at 80 °C for 2 h, after which the suspension turned clear. The reaction mixture was stirred for a further 20 min under refluxing conditions after which, it was allowed cool to room temperature. Methanol was evaporated under reduced pressure, and the resulting residue was coevaporated from toluene to remove the residual methanol and then dried *in-vacuo* for 2 h. The residue was taken up in dry DMF (50 mL) and allyl bromide (3 eq) and TBAI (0.4 eq) were added and the reaction mixture was stirred at room temperature for 18 h. The progress of the reaction was monitored by TLC. In case of slow conversion 1 more eq of allyl bromide and 0.2 eq of TBAI were added. Upon completion, the reaction mixture was concentrated under reduced pressure, the residue dissolved up in minimum amount of DCM:MeOH (1/1, v/v), dry loaded on a silica gel column and chromatographed using a gradient of EtOAc:MeOH (100% → 60%). Fractions containing product were collected, concentrated under reduced pressure and the product characterized by MS and NMR.

**General protocol for benzylation.** Compounds **21a-c** were dissolved in DMF and the resulting solution cooled in an ice-bath. NaH (1.5 eq per each hydroxyl) and BnBr (2 eq per each hydroxyl) were added and the reaction mixture was stirred for 30 min at 0 °C and then 2 h at room temperature. The progress of the reactions were monitored by TLC and when completed, the reaction was quenched by the addition of ice cold methanol. The solvents were evaporated under reduced pressure and the residue were dissolved in DCM, subsequently washed with 1M aqueous HCl, sat. $NaHCO_3$ solution and brine. The organic layer was separated, dried ($Na_2SO_4$) and concentrated under reduced pressure. The residue was applied to a column silica gel and eluted with a gradient of petroleum ether: EtOAc = 20:1 → 6:1. Fractions containing product were concentrated under reduced pressure and used in the next step without further characterization.

**General procedure for conversion of azide into NHTCA.** Intermediate compounds obtained after benzylation of **21b-c** were dissolved in 90% aq. THF and to this solution 1M solution of $PMe_3$ in THF was added untill a final 5-fold excess of azido glycan (caution: no base was used for the reduction to avoid the *N*-benzylation with traces of remaining benzyl bromide from the previous reaction). The reaction mixtures stirred for 2 h at room temperature, after which TLC indicated full consumption of the starting material. The solvents were evaporated under reduced pressure and the residue was coevaporated twice from toluene to remove trace amounts of moisture and trimethyl phosphine. The residue was further dried in high *vacuo* for 2 h. The solid residue was dissolved in DCM, triethylamine (2 eq to starting material) was added followed by the addition of trichloroacetyl chloride (1.5 eq per each $NH_2$). TLC monitoring indicated the formation of trichloroacetamide product within 15 min. The reaction mixture was further stirred for 30 min, quenched with sat. solution of aqueous $NaHCO_3$, and washed with 1M HCl, brine. Organic phases were collected, dried ($Na_2SO_4$) and concentrated *in vacuo*. The residue was applied to a silica gel column chromatography and eluted with Tol:EtOAc=9:1, v:v.

**General procedure for allyl ether removal.** To solutions of compounds **22a-c** in MeOH:DCM = 4:1, v:v, a catalytic amount of $PdCl_2$ (0.2 eq) was added and the solution was stirring at room temperature. The progress of deallylation was monitored by TLC and when no more starting material was observed, the reactions were concentrated by rotary evaporation under reduced pressure. The remaining dark-brown oily crude product was applied to silicagel column chromatography and eluted with Tol:EtOAc=8:1, v:v eluent to afford the required compound.

**N-(Benzyl)-benzyloxycarbonyl-5-aminopentyl β-D-galactopyranosyl)-(1→4)-(2-amino-2-deoxy-β-D-glucopyranosyl)-(1→3)-(β-D-galactopyranosyl)-(1→4)-β-D-glucopyranoside (19b).**

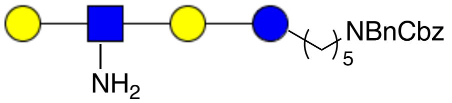


Compound **15b** (250 mg, 0.23 mmol) was subjected to deprotection according to general procedure to obtain 190 mg of compound **19b** (84%). MALDI-TOF MS *m/z* $C_{44}H_{66}N_2O_{22}$ $(M+Na)^+$ 997.4005 found 997.4405.

$^1$H NMR (600 MHz, $D_2O$) δ: (non carbohydrate): 7.41 – 7.02 (m, 10H, Ar-H), 5.08 (d, $J$ = 34.7 Hz, 2H, $CH_2$ Cbz), 4.44 (d, $J$ = 21.2 Hz, 2H, $CH_2$ NBn), 3.75 (-OC*__H__*H linker), 3.50 (-OCH*__H__*$_2$ linker), 3.23 (t, $J$ = 7.4 Hz, 2H, -$CH_2$ linker), 1.51 – 1.42 (m, 4H, -$CH_2$ linker), 1.41 – 1.34 (m, 2H, -$CH_2$ linker).

$^{13}$C NMR (151 MHz, $D_2O$): $\delta$ (non carbohydrate): 129.0 – 126.0 (C Ar) 70.3 (-$OCH_2$ linker), 67.9

(*C*$H_2$ Cbz), 50.2 (*C*$H_2$ NBn), 47.2 (-$CH_2$ linker), 28.5 (-$CH_2$ linker), 26.6 (-$CH_2$ linker), 22.1 (-$CH_2$ linker).

Carbohydrate region:

| | H1 | H2 | H3 | H4 | H5 | H6 |
|---|---|---|---|---|---|---|
| Glc | 4.32 | 3.20 | 3.54 | 3.54 | 3.45 | 3.87 – 3.65 |
| Gal | 4.41<br>d, *J* = 7.9 Hz | 3.56 | 3.71 | 3.84,<br>d, *J* = 3.3 Hz | 3.66 | 3.87 – 3.65 |
| GlcNAc | 4.54<br>d, *J* = 8.0 Hz | 2.66<br>d, J = 9.1 Hz | 3.49 | 3.57 | 3.49 | 3.87 – 3.65 |
| Gal(2) | 4.37<br>d, *J* = 8.4 Hz | 3.47 | 3.57 | 4.10<br>d, *J* = 2.8 Hz | 3.60 | 3.87 – 3.65 |

| | C1 | C2 | C3 | C4 | C5 | C6 |
|---|---|---|---|---|---|---|
| Glc | 102.6 | 73.4 | 75.0 | 74.4 | 71.3 | 59.0 – 61.0 |
| Gal | 102.6 | 74.9 | 82.5 | 68.1 | 75.3 | 59.0 – 61.0 |
| GlcNAc | 104.9 | 56.5 | 74.8 | 72.7 | na | 59.0 – 61.0 |
| Gal(2) | 103.5 | 78.7 | 78.7 | 68.6 | 69.8 | 59.0 – 61.0 |

**N-(Benzyl)-benzyloxycarbonyl-5-aminopentyl β-D-galactopyranosyl-(1→4)-2-amino-2-deoxy-β-D-glucopyranosyl-(1→3)-β-D-galactopyranosyl-(1→4)-2-amino-2-deoxy-β-D-glucopyranosyl-(1→3)-β-D-galactopyranosyl-(1→4)-β-D-glucopyranoside (19c).**

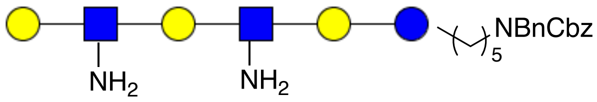


Compound **15c** (200 mg, 0.13 mmol) was subjected to TFA removal according to the general procedure to yield 140 mg (82%) of **19c**. MALDI-TOF MS *m/z* $C_{56}H_{87}N_3O_{31}$ $(M+Na)^+$ calculated 1321.2918, found 1321.0382.

$^1$H NMR (600 MHz, $D_2O$) δ: (non carbohydrate): 7.42 – 7.14 (m, 10H, Ar-H), 5.09 (d, *J* = 43.8 Hz, 2H, $CH_2$ Cbz ), 4.46 (m, 2H, -$NCH_2Ph$), 3.72 (-OC*H*H linker), 3.56 (-OCH*H* linker), 3.24 (t, *J* = 7.4 Hz, 2H, -$CH_2$ linker), 1.58 – 1.42 (m, 4H, -$CH_2$ linker), 1.26 – 1.16 (m, 2H, -$CH_2$ linker).

$^{13}$C NMR (151 MHz, $D_2O$): *δ* (non carbohydrate): 129.0 – 126.0 (C Ar), 70.2 (-O*C*$H_2$ linker), 67.3 (*C*$H_2$ Cbz), 50.1 (-$NCH_2Ph$), 47.2 (-*C*$H_2$ linker), 28.6 (-*C*$H_2$ linker), 26.3 (-$CH_2$ linker), 22.3 (-$CH_2$ linker).

Carbohydrate region:

| | H1 | H2 | H3 | H4 | H5 | H6 |
|---|---|---|---|---|---|---|
| Glc | 4.33 | 3.20 | 3.55 | 3.55 | 3.48 | 3.78<br>3.73 |
| Gal | 4.39<br>d, $J$ = 8.2 Hz | 3.46 | 3.78 | 3.78,<br>d, $J$ = 2.9 Hz | na | 3.74 – 3.66 |
| GlcNAc | 4.79<br>d, $J$ = 8.4 Hz | 2.94 | 3.67 | 3.67 | 3.55 | 3.89<br>3.78 |
| Gal(2) | 4.42<br>d, $J$ = 8.5 Hz | 3.63 | 3.78 | 4.06<br>d, $J$ = 2.9 Hz | na | 3.74 – 3.66 |
| GlcNAc2 | 4.80<br>d, $J$ = 8.2Hz | 2.94 | 3.67 | 3.64 | 3.55 | 3.89<br>3.78 |
| Gal(3) | 4.42<br>d, $J$ = 7.8 Hz | 3.63 | na | 3.90,<br>d, $J$ = 3.1 Hz | na | 3.74 – 3.66 |

| | C1 | C2 | C3 | C4 | C5 | C6 |
|---|---|---|---|---|---|---|
| Glc | 102.1 | 72.8 | 75.0 | 75.0 | 74.8 | 59.0 – 61.0 |
| Gal | 103.1 | 71.2 | 78.7 | 68.1 | na | 59.0 – 61.0 |
| GlcNAc | 101.7 | 55.9 | 75.4 | 74.4 | 78.3 | 59.0 – 61.0 |
| Gal(2) | 102.6 | 70.8 | 82.5 | 68.3 | na | 59.0 – 61.0 |
| GlcNAc (2) | 101.7 | 55.9 | 75.4 | 74.4 | 78.3 | 59.0 – 61.0 |
| Gal(3) | 102.6 | 70.7 | 82.5 | 68.3 | na | 59.0 – 61.0 |

**N-(Benzyl)-benzyloxycarbonyl-5-aminopentyl β-D-galactopyranosyl)-(1→4)-(2-azido-2-deoxy-β-D-glucopyranosyl)-(1→3)-(β-D-galactopyranosyl)-(1→4)-β-D-glucopyranoside (20b).**

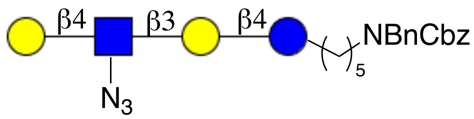


Compound **19b** (180 mg, 0.18 mmol) was converted into **20b** (166 mg) in 90% yield according to the **g**eneral protocol for azido transfer. MALDI-TOF MS *m/z* $C_{44}H_{64}N_4O_{22}$ $(M+Na)^+$ calculated 1023.3910, found 1023.4641.

$^1$H NMR (600 MHz, $D_2O$) δ: (non carbohydrate): 7.36 – 7.11 (m, 10H, Ar-H), 5.04 (d, J = 29.2 Hz, 2H, $CH_2$ Cbz ), 4.42 (d, J = 27.3 Hz, 2H, -$CH_2$NPh), 3.72 (-OC*H*H linker), 3.56 (-OCH*H* linker), 3.20 (m, 2H, -$CH_2$ linker), 1.45 – 1.39 (m, 4H, -$CH_2$ linker), 1.19 – 1.12 (m, 2H, -$CH_2$ linker).

$^{13}C$ NMR (151 MHz, $D_2O$): $\delta$ (non carbohydrate): 129.6 – 125.9 (C Ar) 67.66 ($\underline{C}H_2$ Cbz ), 50.2 (-$\underline{C}H_2$ NPh), 47.3 (-$\underline{C}H_2$ linker), 28.4 (-$\underline{C}H_2$ linker), 26.8 (-$\underline{C}H_2$ linker), 22.3 (-$\underline{C}H_2$ linker ).

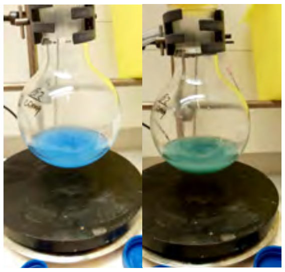

| | H1 | H2 | H3 | H4 | H5 | H6 |
|---|---|---|---|---|---|---|
| Glc | 4.23 | 3.15 | 3.50 | 3.50 | 3.48 | 3.82<br>3.70 |
| Gal | 4.33<br>d, $J$ = 7.7 Hz | 3.40 | 3.73 | 4.03<br>d, $J$ = 2.8 Hz | 3.66 | 3.88 – 3.55 |
| GlcNAc | 4.67 | 3.29<br>dd, $J_1$ = 8.5 Hz, $J_2$ = 10.7 Hz | 3.49 | 3.60 | 3.54 | 3.87 – 3.79 |
| Gal(2) | 4.34<br>d, $J$ = 7.3 Hz | 3.60 | 3.54 | 3.79<br>d, $J$ = 2.7 Hz | na | 3.88– 3.55 |

| | C1 | C2 | C3 | C4 | C5 | C6 |
|---|---|---|---|---|---|---|
| Glc | 101.8 | 72.7 | 74.3 | 78.3 | 72.3 | 59.7 |
| Gal | 102.4 | 71.0 | 81.9 | 68.5 | na | 60.7 |
| GlcNAc | 102.9 | 65.2 | 74.6 | 77.8 | 72.2 | 59.8 |
| Gal(2) | 102.8 | 70.2 | na | 68.4 | na | 60.9 |

**N-(Benzyl)-benzyloxycarbonyl-5-aminopentyl β-D-galactopyranosyl-(1→4)-2-azido-2-deoxy-β-D-glucopyranosyl-(1→3)-β-D-galactopyranosyl-(1→4)-2-azido-2-deoxy-β-D-glucopyranosyl-(1→3)-β-D-galactopyranosyl-(1→4)-β-D-glucopyranoside (20c).**

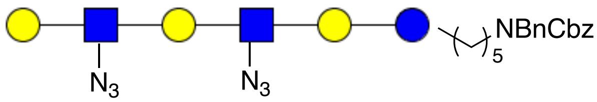


Compound **20c** (101 mg) was obtained from starting material, **19c** (130 mg, 0.1 mmol) following the general protocol for azido transfer. MALDI-TOF MS $m/z$ $C_{56}H_{83}N_7O_{31}$ $(M+Na)^+$ calculated 1372.5031, found 1372.6177.

$^1H$ NMR (600 MHz, $D_2O$) δ: (non carbohydrate): 7.40 – 7.15 (m, 10H, Ar-H), 5.10 (d, $J$ = 44.3

Hz, 2H, $CH_2$ Cbz), 4.44 (m, 2H, -$NCH_2$Ph), 3.72 (-OC*H*H linker), 3.53 (-OCH*H* linker), 3.26 (m -$CH_2$ linker), 1.54 – 1.44 (m, 4H, -$CH_2$ linker), 1.26 – 1.18 (m, 2H, -$CH_2$ linker).

$^{13}$C NMR (151 MHz, $D_2O$) $\delta$: (non carbohydrate): 129.6 – 126.2 (C Ar), 70.2 (-O*C*$H_2$ linker), 67.5 (*C*$H_2$ Cbz), 50.5 (-$NCH_2$Ph), 47.4 (-*C*$H_2$ linker), 28.2 (-*C*$H_2$ linker ), 26.9 (-*C*$H_2$ linker), 22.3 (-*C*$H_2$ linker).

| | H1 | H2 | H3 | H4 | H5 | H6 |
|---|---|---|---|---|---|---|
| Glc | 4.37 | 3.21 | 3.55 | na | na | 3.75 |
| Gal | 4.40 | 3.64 | 3.78 | 4.09 | na | 3.74 – 3.66 |
| GlcNAc | 4.72 | 3.34 | 3.66 | na | na | 3.89<br>3.76 |
| Gal(2) | 4.40 | 3.64 | 3.78 | 4.09 | na | 3.74 – 3.66 |
| GlcNAc(2) | 4.72 | 3.34 | 3.66 | na | na | 3.89<br>3.76 |
| Gal(3) | 4.39 | 3.45 | na | 3.86 | na | 3.74 – 3.66 |

| | C1 | C2 | C3 | C4 | C5 | C6 |
|---|---|---|---|---|---|---|
| Glc | 101.9 | 72.7 | na | na | na | 60.0 |
| Gal | 102.9 | 70.3 | 82.1 | 68.5 | na | 60.8 |
| GlcNAc | 102.7 | 65.4 | na | na | na | 59.8 |
| Gal(2) | 102.8 | 70.3 | 82.1 | 68.5 | na | 60.8 |
| GlcNAc(2) | 102.7 | 65.4 | na | na | na | 60.8 |
| Gal(3) | 102.8 | 70.9 | na | 68.6 | na | 60.8 |

***N*-(Benzyl)-benzyloxycarbonyl-5-aminopentyl 3-*O*-allyl-*β*-D-galactopyranosyl-(1→4)-*β*-D-glucopyranoside (21a).**

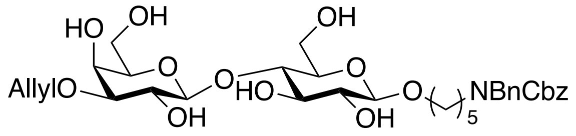


Compound **21a** (160 mg, 75%) was synthesized according to the general allylation procedure starting form **15a** (200 mg, 0.31 mmol), $Bu_2SnO$ (114 mg, 0.46 mmol), allyl bromide (80 $\mu$l, 0.92 mmol) and TBAI (61 mg, 0.16 mmol). $R_f$ = 0.6 (100% EtOAc). MALDI-TOF MS *m/z* $C_{35}H_{49}NO_{13}$ $(M+Na)^+$ calculated 714.3102, found 714.4697.

$^1$H NMR (600 MHz, $CD_3OD$-$d_4$) δ 7.36–7.01 (m, 10H, Ar-H), 5.89 (m, 1H, $CH_2$=*CH*-$CH_2$-), 5.21

(m, 1H, C*H*H=CH-CH2-), 5.06 (m, 3H, CH*H*=CH-CH2-, -CH2 Cbz), 4.42 (s, 2H, -NCH2Ph), 4.28 (d, *J* = 7.8 Hz, 1H, H-1 Gal), 4.19 – 4.11 (m, 2H, H-1 Glc, CH2=CH-C*H*H-), 4.02 (dd, $J_1$ = 5.1 Hz, $J_2$ = 12.5 Hz, 1H, CH2=CH-CH*H*-), 3.90 (d, *J* = 3.0 Hz, 1H, H-4 Gal), 3.80 – 3.65 (m, 4H, -CH*H* linker, H-6 Glc, H-6$_a$ Gal), 3.60 (dd, $J_1$ = 4.9 Hz, J2 = 11.5 Hz, 1H, H-6 Gal), 3.53 (t, *J* = 8.6 Hz, 1H, H-2 Gal), 3.48 – 3.38 (m, 3H, H-4 Glc, H-5 Gal, H-3 Glc), 3.49 (m, 1H, -CH*H* linker), 3.28 (m, 1H, H-5 Glc), 3.22 (d, *J* = 3.1 Hz, 1H, H-3 Gal), 3.17 – 3.09 (m, 3H, H-2 Glc, -C*H*2 linker), 1.53 – 1.37 (m, 4H, -CH2 linker), 1.26 – 1.16 (m, 2H, -CH2 linker).

$^{13}$C NMR (151 MHz, $CD_3OD$-d4): $\delta$ 22.9 (-*C*H2 linker), 27.26 (-*C*H2 linker), 28.9 (-*C*H2 linker), 46.4 (-*C*H2 linker), 50.1 (-*C*H2 -NCH2Ph), 60.7 (C6 Glc), 61.0 (C6 Gal), 65.9 (C4 Gal), 67.0 (-*C*H2 Cbz), 69.3 (-*C*H2 linker), 70.3 (-*C*H2 allyl), 70.5 (C2 Gal), 73.3 (C2 Glc), 75.0 (C5 Glc), 75.1 (C3 Glc), 75.5 (C5 Gal), 79.5 (C4 Glc), 80.1 (C3 Gal), 102.9 (C1 Glc), 103.6 (C1 Gal), 116.0 (*CH2*=CH-CH2-), 126.45, 126.9, 127.3, 127.5, 127.6, 127.7, 127.1, 127.2 (CAr), 135.2 (CH2=*CH*-CH2-).

***N*-(Benzyl)-benzyloxycarbonyl-5-aminopentyl 3-O-allyl-*β*-D-galactopyranosyl)-(1→4)-(2-azido-2-deoxy-*β*-D-glucopyranosyl)-(1→3)-(*β*-D-galactopyranosyl)-(1→4)-*β*-D-glucopyranoside (21b).**

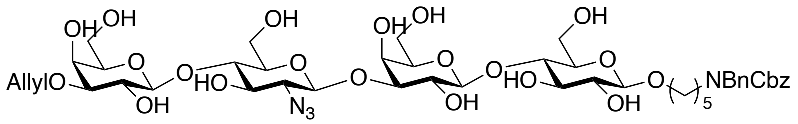


Compound **21b** (114 mg, 68%) was synthesized according to the general allylation procedure starting from **20b** (160 mg, 0.16 mmol), $Bu_2SnO$ (60 mg, 0.24 mmol), allyl bromide (42 $\mu$l, 0.48 mmol) and TBAI (32 mg, 0.08 mmol). $R_f$ = 0.5 (EtOAc:MeOH = 4:1). MALDI-TOF MS *m/z* $C_{47}H_{68}N_4O_{22}$ (M+2Na-$N_2$)$^+$ calculated 1058.4059, found 1057.8085.

$^1$H NMR (600 MHz, $D_2O$) δ 7.41–6.98 (m, 11H, Ar-H), 5.85 (m, 1H, CH2=*CH*-CH2-), 5.24 (d, J = 16.8 Hz, 1H, C*H*H=CH-CH2-), 5.15 (d, J = 10.5 Hz, 1H, CH*H*=CH-CH2-), 5.05 (d, J = 29.0 Hz, 2H, -CH2 Cbz), 4.67 (H-1, GlcNAc), 4.41 (m, 2H, -NCH2Ph), 4.39 – 4.24 (m, 3H, H-1 Gal-2, H-1 Gal-1, H-1 Glc), 4.13 – 3.96 (m, 4H, CH2=CH-C*H*2, H-4 Gal1-2), 3.87 – 3.80 (m, H-6 GlcNAc), 3.75 – 3.56 (m, H-6 Gal1-2, Glc, -OCH2 linker, H-2 Gal1, H-3 Gal1), 3.55 – 3.36 (m, H-2 Gal2,

H-3 Gal2, H-3 GlcNAc, H-4 GlcNAc, H-3 Glc), 3.31 (t, *J* = 9.1 Hz, 1H, H-2 GlcNAc) 3.23 – 3.13 (m, 3H, -NCH2 linker, H-2 Glc), 1.52 – 1.34 (m, 4H, -CH2 linker), 1.27 – 1.07 (m, 2H, -CH2 linker).

$^{13}$C NMR (151 MHz, $D_2O$): $\delta$ 22.2 (-*C*H2 linker), 27.1 (-*C*H2 linker), 28.2 (-*C*H2 linker), 47.2 (-*C*H2 linker), 50.4 (-*C*H2 NBn), 59.7 (C6 GlcNAc), 60.6 (C6 Gal1-2, C6 Glc), 64.6 (C4 Gal1-2), 65.2 (C2 GlcNAc), 67.8 (-*C*H2 Cbz), 70.2 (CH2=CH-*C*H2-), 70.4 (-*C*H2 linker), 72.6 (C2 Glc), 78.2 (C4 Glc), 78.4 (C3 Gal1), 79.5 (C3 Gal2), 101.9 (C1 Glc), 102.6 (C1 GlcNAc), 102.7 (C1 Gal1-2), 118.7 (*CH2*=CH-CH2-), 125.8 – 129.9 (C Ar), 133.7 (CH2=*CH*-CH2-).

**N-(Benzyl)-benzyloxycarbonyl-5-aminopentyl 3-O-allyl-β-D-galactopyranosyl-(1→4)-2-azido-2-deoxy-β-D-glucopyranosyl-(1→3)-β-D-galactopyranosyl-(1→4)-2-azido-2-deoxy-β-D-glucopyranosyl-(1→3)-β-D-galactopyranosyl-(1→4)-β-D-glucopyranoside (21c).**

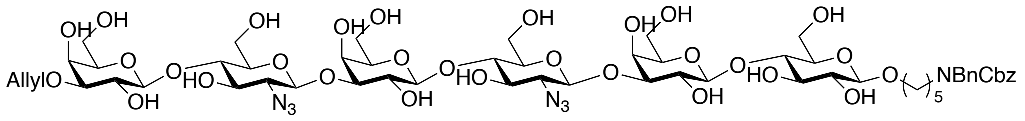


Compound **21c** (71.0 mg, 73%) was synthesized according to the general allylation procedure starting from **20c (**95 mg, 0.07 mmol), $Bu_2SnO$ (26 mg, 0.105 mmol), allyl bromide (25 $\mu$l, 0.21 mmol) and TBAI (32 mg, 0.08 mmol). $R_f$ = 0.2 (EtOAc:MeOH = 3:1). MALDI-TOF MS *m/z* $C_{59}H_{87}N_7O_{31}$ $(M+2Na-N_2)^+$ calculated 1407.5180, found 1407.0012.

$^1$H NMR (600 MHz, $D_2O$) δ 7.38–7.05 (m, 10H, Ar-H), 5.91 (m, 1H, CH2=*C**H***-CH2-), 5.30 (d, *J* = 17.3 Hz, 1H, CH*H*=CH-CH2-), 5.21 (d, *J* = 10.2 Hz, 1H, C*H*H=CH-CH2-), 5.04 (d, *J* = 54.4 Hz, 2H, CH2 Cbz), 4.74 (1H, H-1 GlcNAc1-2), 4.43 – 4.27 (m, 6H, -NCH2Ph, H-1 Gal1-3, H-1 Glc), 4.16 (dd, $J_1$ = 6.6 Hz, $J_2$ = 12.6 Hz, 1H, CH2=CH-CH*H*-), 4.11 (s, 1H, H-4 Gal2), 4.09 (s, 1H, H-4 Gal1), 4.04 (dd, $J_1$ = 5.6 Hz, $J_2$ = 12.3 Hz, 1H, CH2=CH-C*H*H-), 3.89 – 3.75 (m, 10H, H-4 Gal3, H-6 GlcNAc1-2, H-6 Glc, H-3 Gal1-2), 3.76 – 3.60 (m, 13H, H-6 Gal1-3, H-4 GlcNAc1-2, H-5 GlcNAc1-2, H-2 Gal1-2, H-3 Gal3), 3.60 – 3.41 (m, 5H, H-2 Gal3, H-3 Glc, H-4 Glc, H-3 GlcNAc1-2), 3.36 (m, 2H, H-2 GlcNAc1-2), 3.22 (m, 3H, H-2 Glc, -CH2 linker), 1.52 – 1.32 (m, 4H, -CH2 linker), 1.20 – 1.09 (m, 2H, -CH2 linker).

$^{13}C$ NMR (151 MHz, $D_2O$): δ 22.1 (-$\underline{C}H_2$ linker), 26.9 (-$\underline{C}H_2$ linker), 28.4 (-$\underline{C}H_2$ linker), 46.9 (-$\underline{C}H_2$ linker), 50.2 (-N$\underline{C}H_2$Ph), 59.7 (C6 Glc), 59.9 (C6 GlcNAc), 60.9 (C6 Gal1-3), 65.1 (C2 GlcNAc1-2), 67.6 (-$\underline{C}H_2$ Cbz), 68.1 (C4 Gal1-2), 68.4 (C4 Gal3), 69.8 (C2 Gal1-3), 70.1 ($CH_2$=CH-$\underline{C}H_2$), 72.4 (C5 Glc), 72.7 (C2 Glc), 74.3 (C3 Glc), 74.5 (C3 GlcNAc1-2), 78.0 (C4 GlcNAc1-2), 78.5 (C4 Glc), 79.4 (C3 Gal3), 81.8 (C3 Gal1-2), 102.2 (C1 Glc), 102.7 (C1 GlcNAc1-2), 102.9 (C1 Gal1-3), 118.7 ($\underline{C}H_2$=CH-$CH_2$-), 126.6 – 129.4 (C Ar), 133.9 ($CH_2$=$\underline{C}$H-$CH_2$-).

**N-(Benzyl)-benzyloxycarbonyl-5-aminopentyl 3-O-allyl-2,4,6-tri-O-benzyl-β-D-galactopyranosyl-(1→4)-2,3,6-tri-O-benzyl-β-D-glucopyranoside (22a).**

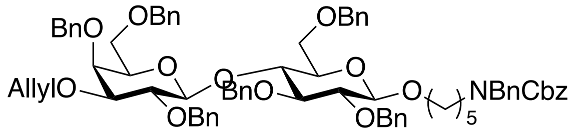


General procedure for *O*-benzylation was applied to convert **21a (**150 mg, 0.22 mol) using NaH (105.0 mg, 2.64 mmol) and BnBr (2.7 mL, 2.64 mmol) into **22a** in a quantitative yield. MALDI-TOF MS *m/z* $C_{77}H_{85}NO_{13}$ $(M+Na)^+$ calculated 1254.5919, found 1254.8246.

$^1H$ NMR (600 MHz, $CDCl_3$) δ 7.42–7.39 (m, 38H, Ar-H), 5.92 (m, 1H, $CH_2$=C$\underline{H}$-$CH_2$-), 5.33 (m, 1H, C$\underline{H}$H=CH-$CH_2$-), 5.20 – 5.13 (m, 3H, CH$\underline{H}$=CH-$CH_2$-, -$CH_2$ Cbz), 5.02 (d, $J$ = 1015 Hz, 1H, -OC$\underline{H}$HPh), 4.96 (d, $J$ = 11.8 Hz, 1H, -OCH$\underline{H}$Ph), 4.90 – 4.70 (m, 5H, -OC$\underline{H_2}$Ph), 4.57 – 4.45 (m, 4H, -OC$\underline{H_2}$Ph, -NC$\underline{H_2}$Ph), 4.44 (d, $J$ = 8.2 Hz, 1H, H-1, Gal), 4.41 – 4.31 (m, 3H, H-1 Glc, -OC$\underline{H_2}$Ph), 4.24 (d, $J$ = 11.8 Hz, 1H, -OC$\underline{H_2}$Ph), 4.20 – 4.12 (m, 2H, $CH_2$=CH-C$\underline{H_2}$-), 3.91 (t, $J$ = 9.8 Hz, 1H, H-4 Glc), 3.87 (d, $J$ = 2.7 Hz, 1H, H-4 Gal), 3.85 (m, 1H, -OC$\underline{H}$H linker), 3.77 (dd, $J_1$ = 4.2 Hz, $J_2$ = 10.8 Hz, 1H, H-6 Glc), 3.75 – 3.68 (m, 2H, H-6 Glc, H-2 Gal), 3.57 – 3.52 (m, 2H, H-3 Glc, H-6 Gal), 3.47 (m, 1H, -OCH$\underline{H}$ linker), 3.40 – 3.33 (m, 4H, H-2 Glc, H-5 Glc, H-5 Gal, H-6 Gal), 3.31 (dd, $J_1$ = 3.1 Hz, $J_2$ = 9.9 Hz, 1H, H-3 Gal), 3.26 – 3.13 (m, 2H, -NC$\underline{H_2}$ linker), 1.67 – 1.49 (m, 4H, -C$\underline{H_2}$$CH_2$C$\underline{H_2}$ linker), 1.26 – 1.16 (m, 2H, -$CH_2$C$\underline{H_2}$$CH_2$ linker).

$^{13}C$ NMR (151 MHz, $CDCl_3$): δ 23.5 (-$CH_2$$\underline{C}H_2$$CH_2$ linker), 27.7, 29.4 (-$\underline{C}H_2$$CH_2$$\underline{C}H_2$ linker), 46.7 (-N$\underline{C}H_2$ linker), 50.3 (-N$\underline{C}H_2$Ph), 67.2 (-$\underline{C}H_2$ Cbz), 68.2 (C6 Gal), 68.6 (C6 Glc), 69.8 (-$\underline{C}H_2$ linker), 71.2 ($CH_2$=CH-$\underline{C}H_2$-), 73.5 (C4 Gal), 72.9 – 75.3 (C5 Gal, C5 Glc, -OC$\underline{H_2}$Ph), 77.0 (C4

Glc), 79.9 (C2 Gal), 81.8 (C2 Glc), 82.6 (C3 Gal), 83.1 (C3 Glc), 102.7 (C1 Gal), 103.6 (C1 Glc), 116.4 (*C*$H_2$=CH-$CH_2$-), 127.0, 127.3, 127.4, 127.5, 127.7, 127.8, 127.91, 128.0, 128.1, 128.2, 128.3, 128.4, 128.5, 138.0, 138.1, 138.5, 138.7, 138.9, 139.0, 139.1 (C Ar), 135.2 ($CH_2$=*CH*-$CH_2$-), 156.5 (-C=O Cbz).

**N-(Benzyl)-benzyloxycarbonyl-5-aminopentyl 3-O-allyl-2,4-6-tri-O-benzyl-β-D-galactopyranosyl)-(1→4)-(2-azido-3,6-di-O-benzyl-2-deoxy-β-D-glucopyranosyl)-(1→3)-2,4-6-tri-O-benzyl-β-D-galactopyranosyl)-(1→4)-2,3,6-tri-O-benzyl-β-D-glucopyranoside (22b).**

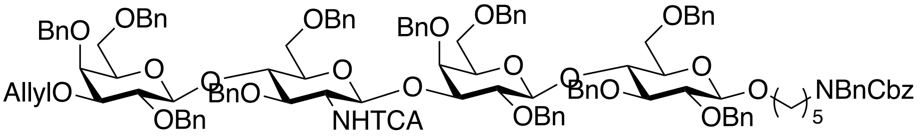


Compound **21b** (100 mg, 0.09 mmol) was benzylated using the general procedure in the presence of NaH (72 mg, 1.8 mmol) and benzyl bromide (181 $\mu$L), and then subjected to Staudinger azide reduction using a $PMe_3$ (450 $\mu$L) solution. The obtained amine intermediate was protected as a trichloroacetamide using 25 mg trichloroacetyl chloride following the general procedure for amine protection to furnish **22b** (135 mg, 70% over 3 steps). MALDI-TOF MS *m/z* $C_{126}H_{135}Cl_3N_2O_{23}$ $(M+Na)^+$ calculated 2171.8419, found 2174.1609.

$^1$H NMR (600 MHz, Chloroform-d) δ 7.38–7.03 (m, 65H, Ar-H), 6.61 (d, J = 7.8 Hz, 1H, -N*H*TCA), 5.93 (m, 1H, $CH_2$=*CH*-$CH_2$-), 5.33 (dd, $J_1$ = 1.7 Hz, $J_2$ = 17.30 Hz, 1H, CH*H*=CH-$CH_2$-), 5.18 (dd, $J_1$ = 1.5 Hz, $J_2$ = 10.5 Hz, 1H, C*H*H=CH-$CH_2$-), 5.14 (d, *J* = 12.2 Hz, 2H, $CH_2$ Cbz), 5.07 (d, *J* = 7.3 Hz, 1H, H-1 GlcNAc), 4.99 (d, *J* = 11.5 Hz, 1H, -OC$H_2$Ph), 4.96 – 4.91 (m, 3H, -OC*H*HPh, -OC$H_2$Ph), 4.85 (d, J = 11.8 Hz, 1H, -OC$H_2$Ph), 4.83 – 4.74 (m, 4H, -OC$H_2$Ph), 4.71 – 4.62 (m, 3H, -OC*H*HPh, -OC$H_2$Ph), 4.57 (d, *J* = 11.3 Hz, 1H, -OC*H*HPh,), 4.53 – 4.43 (m, 7H, -OC*H*HPh, -OC$H_2$Ph, -NC$H_2$Ph), 4.42 (d, *J* = 8.4 Hz, 1H, H-1 Gal2), 4.38 – 4.22 (m, 6H, -OC$H_2$Ph, H-1 Gal1, H-1 Glc), 4.21 – 4.13 (m, 4H, -OC$H_2$Ph, $CH_2$=CH-CH*H*-), 4.03 (t, *J* = 7.9 Hz, 1H, H-4 GlcNAc), 3.92 (d, *J* = 2.5 Hz, 1H, H-4 Gal1), 3.88 (t, *J* = 9.3 Hz, 1H, H-4 Glc), 3.85 (d, *J* = 3.1 Hz, 1H, H-4 Gal2), 3.81 – 3.74 (m, 4H, -$CH_2$ linker, H-$6_a$ GlcNAc, H-2 GlcNAc, H-3 GlcNAc), 3.74 – 3.63 (m, 5H, H-3 Gal1, H-2 Gal2, H-2 Gal1, H-$6_b$ GlcNAc, H-$6_a$ Glc), 3.57 (d, *J* = 10.1 Hz, 1H, H-$6_b$ Glc), 3.55 – 3.47 (m, 2H, H-$6_a$ Gal1, H-5 GlcNAc), 3.47 – 3.28 (m, 8H, H-3 Glc, H-2

Glc, H-3 Gal2, H-6 Gal2, H-5 Gal1, H-5 Gal2, -$CH_2$ linker), 3.26 (dd, $J_1$ = 5.0 Hz, $J_2$ = 9.2 Hz, 1H, H-$6_b$ Gal1), 3.12 (m, 2H, -$CH_2$ linker), 3.17 (m, 1H, H-5 Glc), 1.60 – 1.46 (m, 5H, -$CH_2$ linker), 1.36 – 1.18 (m, 2H, -$CH_2$ linker).

$^{13}$C NMR (151 MHz, $CDCl_3$): $\delta$ 23.4 (-$\underline{C}H_2$ linker), 27.8 (-$\underline{C}H_2$ linker), 29.7 (-$\underline{C}H_2$ linker), 46.7 (-$\underline{C}H_2$ linker), 50.3 (-N$\underline{C}H_2$Ph), 57.3 (C2 GlcNAc), 67.0 (-$\underline{C}H_2$ Cbz), 67.9 – 68.2, (C6 Gal1, C6 Glc, C6 GlcNAc, C6 Gal2, 69.8 (-$\underline{C}H_2$ linker), 71.6 (-$\underline{C}H_2$ allyl), 73.1, 73.2 2x(-(-O$\underline{C}H_2$Ph), 73.2, 73.3 (C5, Gal1 and Gal2), 73.4 (-O$\underline{C}H_2$Ph), 73.4 (-O$\underline{C}H_2$Ph), 73.6 (C4 Gal2), 73.9 (-O$\underline{C}H_2$Ph), 74.6, 74.7, 2x(-O$\underline{C}H_2$Ph), 74.1 (C3 Gal), 75.0 (C5 Glc), 75.3 (-O$\underline{C}H_2$Ph), 75.4 (C5 GlcNAc), 75.4 (-O$\underline{C}H_2$Ph), 76.3 (C4 Gal1), 76.3 (C4 Glc), 76.6 (C4 GlcNAc), 78. (C3 GlcNAc), 79.83 (C3 Gal), 79.9 (C2 Gal), 80.3 (C2 Gal), 81.7 (C3 Gal2), 82.2 (C2 Glc), 82.7 (C3 Glc), 100.3 (C1 GlcNAc), 102.6 (C1 Gal), 103.1 (C1 Gal), 103.6 (C1 Glc), 116.6 ($\underline{C}H_2$=CH-$CH_2$-), 127.1, 127.2, 127.3, 127.4, 127.5, 127.6, 127.7, 127.7, 127.8, 127.8, 127.9, 128.0, 128.1, 128.1, 128.2, 128.3, 128.4, 128.4, 128.4, 128.4, 128.4, 128.5 (C Ar), 134.9 ($CH_2$=$\underline{C}$H-$CH_2$-). 138.7 (-COO Cbz).

***N*-(Benzyl)-benzyloxycarbonyl-5-aminopentyl 3-O-allyl-2,4-6-tri-O-benzyl-*β*-D-galactopyranosyl)-(1→4)-(2-azido-3,6-di-O-benzyl-2-deoxy-*β*-D-glucopyranosyl)-(1→3)-(2,4-6-tri-O-benzyl-*β*-D-galactopyranosyl)-(1→4)-(2-azido-3,6-di-O-benzyl-2-deoxy-*β*-D-glucopyranosyl)-(1→3)-2,4-6-tri-O-benzyl-*β*-D-galactopyranosyl)-(1→4)-2,3,6-tri-O-benzyl-*β*-D-glucopyranoside (22c).**

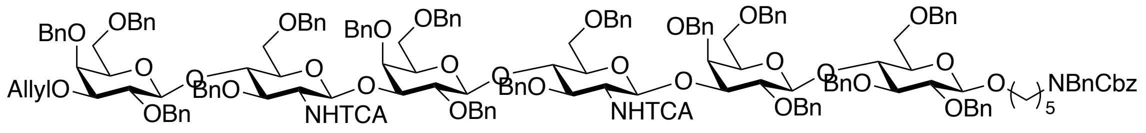


Compound **21c** (65 mg, 0.05 mmol) was benzylated using the general procedure in the presence of NaH (64 mg, 1.6 mmol) and benzyl bromide (160 $\mu$L), which was subsequently subjected to Staudinger azide reduction using a $PMe_3$ (250 $\mu$L) solution. The obtained amine intermediate was protected as a trichloroacetamide using trichloroacetyl chloride ( 27 mg, 0.15 mmol) following the general procedure for amine protection to furnish **22c** (104 mg, 68% over 3 steps). MALDI-TOF MS *m/z* $C_{175}H_{185}Cl_6N_3O_{33}$ $(M+Na)^+$ calculated 3089.0919, found 3093.3962.

[1]H NMR (600 MHz, Chloroform-d) δ 7.38–7.04 (m, 83H, Ar-H), 6.67 (d, *J* = 7.9 Hz, 1H, -N*H*TCA GlcNAc2), 6.59 (d, *J* = 8.4 Hz, 1H, -N*H*TCA GlcNAc1), 5.93 (m, 1H, $CH_2$=*CH*-$CH_2$-), 5.34 (dd, $J_1$ = 1.9 Hz, $J_2$ = 17.5 Hz, 1H, C*H*H=CH-$CH_2$-), 5.18 (dd, $J_1$ = 1.5Hz, $J_2$ = 10.6 Hz, 1H, CH*H*=CH-$CH_2$-), 5.14 (d, *J* = 12.0 Hz, 2H, $CH_2$ Cbz), 5.10 (d, *J* = 7.2 Hz, 1H, H-1 GlcNAc2), 5.01 (d, *J* = 7.3 Hz, 1H, H-1 GlcNAc1), 4.98 (d, *J* = 11.7 Hz, 1H, -OC*H₂*Ph), 4.97 (d, *J* = 11.5 Hz, 1H, -OC*H₂*Ph), 4.94 (d, *J* = 10.9 Hz, 2H, -OC*H₂*Ph), 4.86 (d, *J* = 10.8 Hz, 1H, -OC*H*HPh), 4.84 – 4.62 (m, 8H, -OC*H₂*Ph), 4.53 – 4.44 (m, 10H, -NC*H₂*Ph, -OC*H₂*Ph), 4.42 (d, *J* = 7.7 Hz, 1H, H-1 Gal3), 4.40 – 4.23 (m, 9H, H-1 Gal2, H-1 Gal, H-1 Glc, -OC*H₂*Ph), 4.21 – 4.09 (m, 4H, -OC*H₂*Ph, $CH_2$=CH-C*H₂*-), 3.99 (m, 2H, H-4 GlcNAc1,2), 3.91 (d, *J* = 2.7 Hz, 1H, H-4 Gal2), 3.89 (m, 1H, H-4 Glc), 3.87 (m, 1H, H-4 Gal), 3.85 (d, *J* = 2.6 Hz, 1H, H-4 Gal3), 3.84 – 3.80 (m, 2H, -$CH_2$ linker, H-6 GlcNAc), 3.80 – 3.75 (m, 3H, H-3 GlcNAc2, H-2 GlcNAc2, H-2 GlcNAc), 3.77 – 3.75 (m, 2H, H-3 Gal2, H-2 Gal2), 3.74 – 3.67 (m, 6H, H-3 GlcNAc1, H-2 Gal3, H-3 Gal1, H-2 Gal1, H-6 GlcNAc), 3.65 (dd, $J_1$ = 4.2 Hz, $J_2$ = 11.5 Hz, 1H, H-$6_a$ Glc), 3.60 (d, *J* = 10.3 Hz, 1H, H-$6_b$ Glc), 3.58 – 3.52 (m, 2H, H-6 GlcNAc), 3.54 (m, 1H, H-5 GlcNAc2), 3.50 – 3.33 (m, 9H, H-6 Gal2, H-3 Glc, H-6 Gal3, H-5 GlcNAc1, H-5 Gal3, H-5 Gal2, H-5 Gal1, -CH*H* linker), 3.37 – 3.31 (m, 2H, H-2 Glc, H-3 Gal3), 3.27 (m, 1H, H-6 Gal), 3.26 – 3.12 (m, 2H, -$CH_2$ linker), 3.17 (m, H, H-5 Glc), 1.63 – 1.45 (m, 5H, -$CH_2$ linker), 1.35 – 1.27 (m, 2H, -$CH_2$ linker).

[13]C NMR (151 MHz, $CDCl_3$): *δ* 23.4 (-*C*$H_2$ linker), 27.8 (-*C*$H_2$ linker), 29.6 (-*C*$H_2$ linker), 46.3 (-*C*$H_2$ linker), 50.4 (-N*C*$H_2$Ph), 57.6 (C2 GlcNAc), 57.7 (C2 GlcNAc2), 67.2 (-*C*$H_2$ Cbz), 67.7 – 68.3 (C6 Gal3, C6 Gal2, C6 GlcNAc2, C6 GlcNAc, C6 Gal, C6 Glc), 69.7 (-*C*$H_2$ linker), 71.6 (-*C*$H_2$ allyl), 73.1, 73.2 (-O*C*$H_2$Ph), 73.2 (C5 Gal1), 73.2 (C5 Gal2), 73.3, 73.4 (-O*C*$H_2$Ph), 73.5 (C5 Gal3), 73.5 (-O*C*$H_2$Ph), 73.5 (C4 Glc), 73.8, 74.04 (-O*C*$H_2$Ph), 74.7, 74.8, 74.9, 75.0 (-O*C*$H_2$Ph), 75.3 (C5 Glc), 75.4 (-O*C*$H_2$Ph), 75.4 (C5 GlcNAc, C5 GlcNAc), 75.6 (-O*C*$H_2$Ph), 76.0 (C4 Gal3), 76.2 (C4 Gal), 76.3 (C4 GlcNAc2, C4 GlcNAc1), 76.6 (C4 Gal2), 78.2 (C3 GlcNAc, C3 GlcNAc2), (C3 Gal2), 80.2 (C2 Gal1), 80.3 (C2 Gal2), 81.7 (C3 Gal3), 82.2 (C2 Glc), 82.7 (C3 Glc), 100.3 (C1 GlcNAc2), 100.0 (C1 GlcNAc1), 102.5 (C1 Gal1, 102.9 (C1 Gal2), 103.1 (C1 Gal3), 103.6 (C1 Glc), 115.6 (*CH₂*=CH-$CH_2$-), 127.1, 127.2, 127.3, 127.4, 127.5, 127.6, 127.7, 127.8, 127.9, 128.0, 128.1, 128.2, 128.3, 128.4, 128.5, 128.6 (C Ar), 134.9 ($CH_2$=*CH*-$CH_2$-), 137.9, 138.1, 138.2, 138.3, 138.3, 138.4, 138.7, 138.8, 138.9, 139.0, 139.1, 139.2 (C Ar), 139.0 (-COO Cbz), 161.6, 161.7 (-NH*TCA*).

**$N$-(Benzyl)-benzyloxycarbonyl-5-aminopentyl 2,4,6-tri-O-benzyl-$\beta$-D-galactopyranosyl-(1→4)-2,3,6-tri-O-benzyl-$\beta$-D-glucopyranoside (16a).**

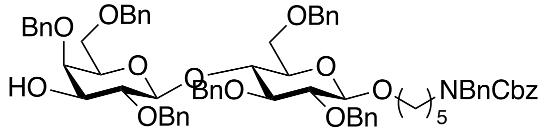


Compound **22a** (250 mg, 0.20 mmol) was treated with $PdCl_2$ (7 mg, 0.04 mmol) according to general procedure for allyl removal to yield **16a** (190 mg,79%). MALDI-TOF MS *m/z* $C_{74}H_{81}NO_{13}$ $(M+Na)^+$ calculated 1214.5606, found 1214.7406.

$^1$H NMR (600 MHz, $CDCl_3$) δ 7.40–7.09 (m, 40H, Ar-H), 5.14 (d, *J* = 14.2 Hz, 2H, $CH_2$ Cbz), 5.00 (d, *J* = 10.9 Hz, 1H, -OC*H*HPh), 4.85 (m, -OC*H*HPh), 4.80 (d, *J* = 11.8 Hz, 1H, -OCH*H*Ph), 4.77 – 4.65 (m, 4H, -OC*H*2Ph), 4.60 (d, *J* = 11.8 Hz, 1H, -OCH*H*Ph), 4.56 (d, *J* = 12.7 Hz, 1H, -OC*H*HPh), 4.45 (d, *J* = 16.6 Hz, 2H, -NC*H*2Ph), 4.43 – 4.36 (m, 3H, -OC*H*2Ph, H-1 Gal), 4.32 (m, 1H, H-1 Glc), 4.26 (d, $J_1$ = 11.4 Hz, 2H, -OCH*H*Ph), 3.94 (t, *J* = 9.0 Hz, 1H, H-4 Glc), 3.87 – 3.83 (m, 1H, -$CH_2$ linker), 3.83 (d, *J* = 2.7 Hz, 1H, H-4 Gal), 3.77 (dd, $J_1$ = 4.4 Hz, $J_2$ = 10.9 Hz, 1H, H-$6_a$ Gal), 3.73 (d, *J* = 9.1 Hz, 1H, H-$6_b$ Gal), 3.57 – 3.52 (m, 2H, H-$6_a$ Glc, H-3 Glc), 3.50 – 3.46 (m, 2H, H-3 Gal, H-2 Gal), 3.42 – 3.33 (m, 5H, H-5 Gal, H-2 Glc, H-5 Glc, H-$6_b$ Glc, -$CH_2$ linker), 3.26 – 3.12 (m, 2H, -$CH_2$ linker), 1.63 – 1.44 (m, 5H, -$CH_2$ linker), 1.39 – 1.26 (m, 2H, -$CH_2$ linker).

$^{13}$C NMR (151 MHz, $CDCl_3$): $\delta$ 23.4 (-*C*H2 linker), 27.7 (-*C*H2 linker), 29.5 (-*C*H2 linker), 46.3 (-*C*H2 linker), 50.2 (-NC*H*2Ph), 67.1 (-*C*H2 Cbz), 67.9 (C6 Glc), 68.4 (C6 Gal), 69.6 (-*C*H2 linker), 73.1 (-NC*H*2Ph), 73.2 (C5 Gal), 73.4 (-OC*H*2Ph), 74.1 (C3 Gal), 74.9 (-OC*H*2Ph), 75.0 (-OC*H*2Ph), 75.1 (-OC*H*2Ph), 75.3 (C5 Glc), 75.4 (-OC*H*2Ph), 75.9 (C4 Gal), 76.8 (C4 Glc), 80.7(C2 Gal), 81.7 (C2 Glc), 82.9 (C3 Glc), 102.7 (C1 Gal), 103.6 (C1 Glc), 127.2, 127.52, 127.54, 127.6, 127.6, 127.7, 127.7, 127.8, 127.8, 127.9, 128.1, 128.1, 128.3, 128.4, 128.4, 128.5, 128.5 (C Ar), 138.7 (-COO Cbz).

**N-(Benzyl)-benzyloxycarbonyl-5-aminopentyl 2,4-6-tri-O-benzyl-β-D-galactopyranosyl)-(1→4)-(2-azido-3,6-di-O-benzyl-2-deoxy-β-D-glucopyranosyl)-(1→3)- 2,4-6-tri-O-benzyl-β-D-galactopyranosyl)-(1→4)-2,3,6-tri-O-benzyl-β-D-glucopyranoside (16b).**

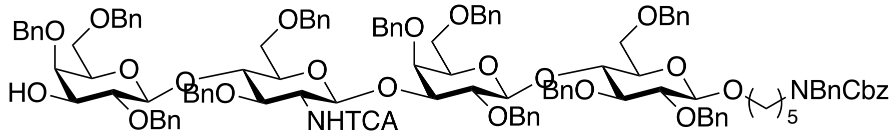


107 mg of compound **16b** was obtained from the reaction of 130 mg (0.06 mmol) of **22b** with $PdCl_2$ (2 mg, 0.012 mmol) according to general procedure for allyl removal. MALDI-TOF MS *m/z* $C_{123}H_{131}Cl_3N_2O_{23}$ $(M+Na)^+$ calculated 2131.8106, found 2134.2571.

$^1$H NMR (600 MHz, $CDCl_3$) δ 7.38–7.03 (m, 62H, Ar-H), 6.60 (d, *J* = 7.8 Hz, 1H, -N*H*TCA), 5.14 (d, *J* = 12.3 Hz, 2H, $CH_2$ Cbz), 5.09 (d, *J* = 7.4 Hz, 1H, H-1 GlcNAc), 5.00 (d, J = 11.2 Hz, 1H, -OCH*H*Ph), 4.96 – 4.92 (m, 2H, -$OCH_2$Ph), 4.85 (d, *J* = 11.8 Hz, 1H, -CH*H*Ph), 4.83 – 4.61 (m, 7H, -OCH*H*Ph, -$OCH_2$Ph), 4.57 (d, *J* = 11.3 Hz, 1H, -C*H*HPh), 4.54 – 4.46 (m, 5H, -OC*H*HPh, -$OCH_2$Ph), 4.44 (d, *J* = 16.6 Hz, 2H, -$NCH_2$Ph), 4.43 – 4.38 (m, 2H, -$OCH_2$Ph, H-1 Gal2), 4.36 – 4.28 (m, 4H, -C*H*HPh, -$OCH_2$Ph, H-1 Gal), 4.26 – 4.16 (m, 3H, H-1 Glc, -$OCH_2$Ph), 4.06 (t, *J* = 8.7 Hz, 1H, H-4 GlcNAc), 3.92 (d, *J* = 2.7 Hz, 1H, H-4 Gal), 3.89 (t, *J* = 9.4 Hz, 1H, H-4 Glc), 3.85 – 3.67 (m, 8H, -CH*H* linker , H-4 Gal2, H-6 GlcNAc, H-2 GlcNAc, H-3 GlcNAc, H-3 Gal, H-2 Gal), 3.65 (dd, $J_1$ = 4.2 Hz, $J_2$ = 11.3 Hz, 1H, H-$6_a$ Glc), 3.57 (d, *J* = 10.8 Hz, 1H, H-$6_b$ Glc), 3.55 – 3.30 (m, 12H, H-6 Gal, H-3 Glc, H-2 Glc, H-2 Gal, H-3 Gal2, H-6 Gal2, H-5 GlcNAc, H-5 Gal, H-5 Gal2, -$CH_2$ linker), 3.26 – 3.12 (m, 2H, -$CH_2$ linker), 3.17 (m, H, H-5 Glc), 1.63 – 1.43 (m, 5H, -$CH_2$ linker), 1.38 – 1.20 (m, 2H, -$CH_2$ linker).

$^{13}$C NMR (151 MHz, $CDCl_3$): *δ* 23.1 (-*C*$H_2$ linker), 27.3 (-*C*$H_2$ linker), 29.4 (-*C*$H_2$ linker), 46.9 (-*C*$H_2$ linker), 50.3 (-N*C*$H_2$Ph), 57.5 (C2 GlcNAc), 67.3 (-*C*$H_2$ Cbz), 67.1, (C6 Gal), 68.1 (C6 Glc), 68.1 (C6 GlcNAc), 68.3 (C6 Gal), 69.7 (-*C*$H_2$ linker), 73.1, 73.2 (-O*C*$H_2$Ph), 73.2 (C5 Gal), 73.3 73.9, 74.6, 74.8, 75.1 (-O*C*$H_2$Ph), 74.0 (C3 Gal), 75.2 (C5 Glc), 75.4 (C5 GlcNAc), 75.8 (C4 Gal2), 76.1 (C4 Gal), 76.2 (C4 Glc), 76.6 (C4 GlcNAc), 78.1 (C3 GlcNAc), 79.9 (C3 Gal), 80.2 (C2 Gal), 80.5 (C2 Gal2), 81.8 (C2 Glc), 82.8 (C3 Glc), 100.1 (C1 GlcNAc), 102.6 (C1 Gal), 103.1 (C1 Gal2), 103.5 (C1 Glc), 127.1, 127.2, 127.2, 127.2, 127.6, 127.6, 127.6, 127.7, 127.7, 127.7, 127.8, 127.90, 127.9, 128.0, 128.1, 128.1, 128.2, 128.3, 128.4, , 128.5, 128.6 ($C_{Ar}$), 138.7 (-COO Cbz).

***N*-(Benzyl)-benzyloxycarbonyl-5-aminopentyl 2,4-6-tri-O-benzyl-*β*-D-galactopyranosyl)-(1→4)-(2-azido-3,6-di-O-benzyl-2-deoxy-*β*-D-glucopyranosyl)-(1→3)- (2,4-6-tri-O-benzyl-*β*-D-galactopyranosyl)-(1→4)-(2-azido-3,6-di-O-benzyl-2-deoxy-*β*-D-glucopyranosyl)-(1→3)-2,4-6-tri-O-benzyl-*β*-D-galactopyranosyl)-(1→4)-2,3,6-tri-O-benzyl-*β*-D-glucopyranoside (16c).**

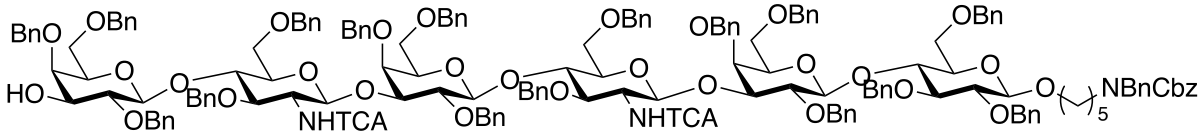

Compound **22c** (100 mg, 0.03 mmol) was treated with $PdCl_2$ (1.154 mg, 0.006 mmol) according yo the general procedure to yield **16c** (77 mg, 78%). MALDI-TOF MS *m/z* $C_{172}H_{181}Cl_6N_3O_{33}$ $(M+Na)^+$ calculated 3049.0606, found 3053.5083.

$^1$H NMR (600 MHz, Chloroform-d) δ 7.40–7.02 (m, 90H, Ar-H), 6.67 (d, *J* = 8.3 Hz, 1H, -N*H*TCA GlcNAc2), 6.58 (d, *J* = 7.6 Hz, 1H, -N*H*TCA GlcNAc1), 5.17 – 5.10 (m, 3H, H-1 GlcNAc2, $CH_2$ Cbz), 5.01 (d, *J* = 7.5 Hz, 1H, H-1 GlcNAc1), 5.00 (d, J = 11.2 Hz, 1H, -$OCH_2Ph$), 4.99 – 4.91 (m, 4H, -$OCH_2Ph$), 4.88 – 4.78 (m, 4H, -$OCH_2Ph$), 4.77 – 4.61 (m, 7H, -$OCH_2Ph$), 4.58 (d, *J* = 11.5 Hz, 1H, -OC*H*HPh), 4.55 – 4.37 (m, 11H, -OC*H*HPh, -$OCH_2Ph$, -$NCH_2Ph$, H-1 Gal3, H-1 Gal2), 4.34 – 4.28 (m, 5H, -$OCH_2Ph$, H-1 Gal), 4.28 – 4.20 (m, 3H, H-1 Glc, -$OCH_2Ph$), 4.18 (d, *J* = 12.0 Hz, 1H, -OC*H*HPh), 4.14 (d, *J* = 11.6 Hz, 1H, -OC*H*HPh), 4.07 (t, *J* = 8.5 Hz, 1H, H-4 GlcNAc2, 4.02 (t, *J* = 8.2 Hz, 1H, H-4 GlcNAc, 3.92 (d, *J* = 2.7 Hz, 1H, H-4 Gal2), 3.89 – 3.80 (m, 6H, H-4 Glc, H-4 Gal, H-4 Gal3, H-3 GlcNAc2, -OC*H*H linker, H-6 GlcNAc), 3.78 – 3.66 (m, 8H, H-2 GlcNAc2, H-2 GlcNAc, H-3 GlcNAc, H-3 Gal2, H-2 Gal2, H-3 Gal, H-2 Gal, H-6 GlcNAc), 3.63 (dd, $J_1$ = 3.3 Hz, $J_2$ = 11.0 Hz, 1H, H-$6_a$ Glc), 3.57 (d, *J* = 10.3 Hz, 1H, H-$6_b$ Glc), 3.57 – 3.29 (m, 15H, H-3 Gal2, H-2 Gal2, H-6 Gal2, H-5 GlcNAc2, H-3 Glc, H-2 Glc, H-6 Gal3, H-5 GlcNAc, H-5 Gal3, H-5 Gal2, H-5 Gal, -CH*H* linker, H-$6_a$ Gal2), 3.26 (m, 1H, H-$6_b$ Gal), 3.26 – 3.12 (m, 2H, -$CH_2$ linker), 3.17 (m, H, H-5 Glc), 1.63 – 1.43 (m, 5H, -$CH_2$ linker), 1.38 – 1.20 (m, 2H, -$CH_2$ linker).

$^{13}$C NMR (151 MHz, $CDCl_3$): *δ* 23.0 (-*C*$H_2$ linker), 27.8 (-*C*$H_2$ linker), 29.3 (-*C*$H_2$ linker), 46.9 (-*C*$H_2$ linker), 50.3 (-N*C*$H_2$Ph), 57.7 (C2 GlcNAc1,2), 67.1 (-*C*$H_2$ Cbz), 67.7 – 68.3 (C6 Gal3, C6 Gal2, C6 GlcNAc2, C6 GlcNAc, C6 Gal, C6 Glc), 69.6 (-*C*$H_2$ linker), 73.1, 73.2, 73.2, 73.3, 73.4,

73.4 (-O*C*$H_2$Ph), 73.4 (C5 Gal, C5 Gal2, C5 Gal3), 73.8 (C3 Gal3), 75.0 (C5 Glc), 75.0 (C5 GlcNAc, C5 GlcNAc2), 75.1, 75.2 (-O*C*$H_2$Ph), 75.3 (C4 Gal), 75.4 (C4 Gal2, C4 Gal3, C4 Glc), 76.1 (C4 GlcNAc), 76.3 (C4 GlcNAc2), 77.9 (C3 GlcNAc), 78.0 (C3 GlcNAc2), 79.8 (C3 Gal2), 79.9 (C2 Gal2), 80.2 (C3 Gal), 80.4 (C2 Gal), 80.6 (C2 Gal3), 81.3 (C2 Glc), 82.7 (C3 Glc), 100.2 (C1 GlcNAc, C1 GlcNAc2), 102.6 (C1 Gal), 103.0 (C1 Gal2), 103.1 (C1 Gal3), 103.6 (C1 Glc), 127.1, 127.2, 127.3, 127.4, 127.5, 127.6, 127.6, 127.7, 127.8, 127.8, 127.9, 127.9, 128.0, 128.0, 128.0, 128.15, 128.2, 128.3, 128.4, 128.4, 128.5, 128.6, 137.8, 137.9, 138.0, 138.2 ($C_{Ar}$), 138.7 (-COO Cbz), 161.6 (-NH*TCA*).

#### 5. Synthesis of target glycans with diverse terminal epitopes

**General procedure for glycosylation of lactosyl acceptors with glucuronic acid donor.** A mixture of acceptor and donor (1 eq and 2eq, respectively) were twice coevaporated from dry toluene and then dried in high *vacuo* for 2 h. It was taken up in DCM to a final concentration of donor and acceptor of 40 mM and 20 mM, respectively. To this solution 4Å molecular sieves was added and the mixture stirred for 2 h at room temperature. The solution was cooled to -40 ℃ , and then a solution of TMSOTf in DCM (3 eq in total) was added dropwise and the reaction mixture left stirring allowing the temperature gradually to rise to -20 ℃. The progress of the glycosylation was monitored by TLC and if necessary an additional portions of donor was added to drive the reaction to completion. When TLC analysis showed almost no acceptor remaining, triethylamine was added to quench the reaction. The mixture was filtered through a pad of celite and washed with saturated aqueous $NaHCO_3$. The aqueous phase was decanted and the DCM layer was dried ($Na_2SO_4$) and concentrated to dryness. The residue was dissolved in a minimum amount of toluene/Acetone (1/1, v/v) and applied to SX-1 size exclusion chromatography to separate the product from unreacted starting materials. The products was characterized by MALDI-TOF MS and NMR.

**General procedure for saponification and intralactone formation reactions.** To a solution of compounds **19a-c** in dioxane was added 1M aqueous KOH until the pH reached 11 and the mixture was stirred vigorously at 40 ℃. The progress of the reaction was monitored by TLC until disappearance of starting material. The reaction mixture was concentrated under reduced pressure and the residue redissolved in methanol. The solution was stirred for 18 h until all benzoyl ester were removed. The pH of the solution was adjusted to neutral with 1M HCl and then concentrated under reduced pressure. The residue was taken up in chloroform and washed with sat. aqueous $NaHCO_3$. The procedure was repeated until the chloroform layer was clear. The aqueous phase was decanted and the organic layer was dried ($Na_2SO_4$), concentrated under reduced pressure and the residue dried *in vacuo* before subjected to the next reaction.

The crude glucuronyl carboxylates were dissolved in $Ac_2O$ and heated to 85 ℃ for 2 h, after which TLC analysis indicated the formation of a 3,6-lactone. The reaction mixture was cooled to room temperature, after which pyridine, acetic anhydride and catalytic amount of DMAP were added. The reaction mixture stirred for 18 h and the progress of the reaction was monitored by MALDI-TOF MS. The mixture was evaporated to dryness under reduced pressure, twice coevaporated form toluene to ensure the complete removal of acetic anhydride and acetic acid. The crude product was dissolved in a minimum amount of toluene, and applied on silica-gel column chromatography eluting with a gradient of EtOAc in toluene (5% → 20%). Fractions containing product were collected, concentrated under reduced pressure and used in the next step after full characterized by NMR and MS.

**General protocol for methanolysis of lactones.** To 100 mM solution of anhydrous NaOAc in dry methanol, 3Å MS were added and the mixture was stirred for 2 h under an inert atmosphere. Compounds **20a-c** were dissolved in dry chloroform (1 mL) and the methanolic solution of sodium acetate (6 mL) was added and the reaction mixture was stirred for 2 h or until the starting material was fully consumed. After completion of the reaction, the solution was buffered by the addition of 50 mL glacial acetic acid and the solvent was evaporated under a stream of $N_2$. The crude product was taken up in toluene and transferred to Eppendorf tube and solid particles removed by centrifugation. The supernatant was applied to a silica-gel column and elutied with 20% EtOAc in toluene to afford the product.

**General protocol for *O*-sulfation.** Compounds **21a-c** were dissolved in pyridine to which $SO_3Py$ (20 eq) was added. The reaction mixture stirred until TLC indicated complete consumption of the alcohol, after which methanol was added to quench unreacted pyridinium sulfur trioxide. The mixture was concentrated under reduced pressure at room temperature. The residue was dissolved in methanol and then passed through Dowex $Na^+$ ion exchange resin to convert the pyridinium form of sulfates to more stable sodium salts.

**General procedure for global deprotection.** Compounds **21a-c** and **22a-c** were dissolved in 1:1 mixture of dioxane and water to which Degussa type $Pd(OH)_2/C$ catalyst was added along with 50 mL of acetic acid. The reaction mixture was stirred overnight under $H_2$ atmosphere. In case of incomplete hydrogenation, additional catalyst was added and the reaction mixture was stirred for a further 18 h under an atmosphere of $H_2$. The formation of desired products was confirmed with ESI-TOF MS in negative mode. The catalyst was filtered off over a pad of compressed celite, the filter washed with water and the supernatant lyophilized. The resulting white fluffy material were dissolved in water and the pH was adjusted to 8. The solution was incubated at 37 ℃ to hydrolyze acetyl and methyl esters. The reaction mixture was freeze-dried and the residue applied to P2 biogel size exclusion column chromatography using aqueous 10 mM $NH_4HCO_3$ as the eluent. Fractions containing product were were collected and lyophilized.

***N*-(Benzyl)-benzyloxycarbonyl-5-aminopentyl (methyl 2,3-di-O-benzoyl-4-O-benzyl-*β*-D-glucopyranosyluronate) -(1→3)-2,4,6-tri-O-benzyl-*β*-D-galactopyranosyl-(1→4)-2,3,6-tri-O-benzyl-*β*-D-glucopyranoside (23a).**

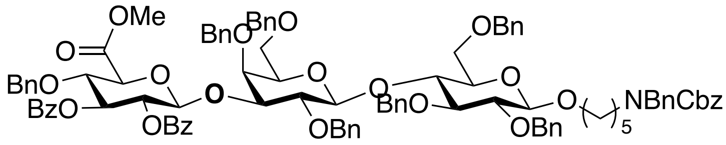


Compound **15a** (121 mg, 0.1 mmol) was reacted with **17** (140 mg 0.2 mmol) in the presence of TMSOTf (66 mg) to yield 23a (152 mg, 91%). MALDI-TOF MS *m/z* $C_{102}H_{105}NO_{21}$ $(M+Na)^+$ calculated 1702.7077, found 1703.1537.

$^1$H NMR (600 MHz, $CDCl_3$) δ 7.99 (m, 2H, Bz), 7.74 (m, 2H, Bz), 7.48 (m, 1H, Bz), 7.39–7.01 (m, 47H, Ar-H), 5.70 (t, $J$ = 9.4 Hz, 1H, H-3 GlcA), 5.52 (dd, $J_1$ = $J_2$ = 7.9 Hz, 1H, H-2 GlcA), 5.20 (d, $J$ =7.8 Hz, 1H, H-1 GlcA), 5.14 (d, $J$ = 12.1 Hz, 2H, $CH_2$ Cbz), 5.05 (d, $J$ = 11.1 Hz, 1H, -OCH*H*Ph), 4.88 (d, $J$ = 10.3 Hz, 1H, -OC*H*HPh), 4.81 (m, 1H, -OCH*H*Ph), 4.67 (d, $J$ = 11.12 Hz,

1H, -OC*H*HPh), 4.59 – 4.55 (m, 2H, -OCH2Ph), 4.54 – 4.50 (m, 3H, -OCH2Ph, -OC*H*HPh), 4.48 – 4.41 (m, 3H, -NC*H*HPh, -OC*H*HPh), 4.35 – 4.16 (m, 7H, -OCH2Ph, -OC*H*HPh, H-1 Gal, H-1 Glc, H-4 GlcA), 4.13 (d, $J$ = 9.5 Hz, 1H, H-5 GlcA), 3.90 (d, $J$ = 2.7 Hz, 1H, H-4 Gal), 3.8 (t, $J$ = 9.0 Hz, 1H, H-4 Glc), 3.81 (m, 1H, -OC*H*HP), 3.76 (s, 3H, -COO*Me*), 3.70 (dd, $J_1$ = 2.7 Hz, $J_2$ = 9.6 Hz, 1H, H-3 Gal), 3.65 (dd, $J_1$ = 4.5 Hz, $J_2$ = 10.9 Hz, 1H, H-6a Glc), 3.53 – 3.47 (m, 3H, H-6a Gal, H-6b Glc, H-2 Gal), 3.44 – 3.29 (m, 5H, H-5 Gal, H-3 Glc, H-2 Glc, H-6b Glc, -CH*H* linker), 3.20 (m, 2H, -CH2 linker), 3.16 (m, 1H, H-5 Glc), 1.59 – 1.43 (m, 4H, -CH2 linker), 1.34 – 1.22 (m, 2H, -CH2 linker).

$^{13}$C NMR (151 MHz, CDCl3): $\delta$ 23.3 (-*C*H2 linker), 27.5 (-*C*H2 linker), 29.4 (-*C*H2 linker), 46.8 (-*C*H2 linker), 50.6 (-N*C*H2Ph), 52.7 (-COO*Me*), 66.9 (-*C*H2 Cbz), 67.9 (C6 Gal), 68.2 (C6 Glc), 69.7 (-*C*H2 linker), 71.9 (C2 GlcA), 73.1 (C5 Gal), 73.2 (-O*C*H2Ph), 73.5 (-O*C*H2Ph), 74.3 (C3 GlcA), 74.5 (C5 GlcA), 74.7, 74.8, 74.9 (-O*C*H2Ph), 75.0 (C5 Glc), 75.1 (-O*C*H2Ph), 75.4 (-O*C*H2Ph), 76.1 (C4 Gal), 76.3 (C4 Glc), 77.6 (C4 GlcA), 80.1 (C2 Gal), 80.2 (C3 Gal), 81.7 (C2 Glc), 82.8 (C3 Glc), 101.5 (C1 GlcA), 102.6 (C1 Gal), 103.5 (C1 Glc), 127.0, 127.2, 127.3, 127.5, 127.6, 127.8, 127.9, 128.0, 128.1, 128.2, 128.2, 128.3, 128.3, 128.4, 128.5, 129.02, 129.2, 129.7, 129.7, 133.0, 133.2, 136.9, 138.2, 138.3, 138.4, 138.7, 139.0 ($C_{Ar}$), 139.3 (-COO Cbz), 164.9 (-OO*C* Bz), 165.2 (-OO*C* Bz), 169.1 (-$C6_{GlcA}$).

***N*-(Benzyl)-benzyloxycarbonyl-5-aminopentyl (methyl 2,3-di-O-benzoyl-4-O-benzyl-*β*-D-glucopyranosyluronate)-(1→3)-2,4-6-tri-O-benzyl-*β*-D-galactopyranosyl)-(1→4)-(2-azido-3,6-di-O-benzyl-2-deoxy-*β*-D-glucopyranosyl)-(1→3)-2,4-6-tri-O-benzyl-*β*-D-galactopyranosyl)-(1→4)-2,3,6-tri-O-benzyl-*β*-D-glucopyranoside (23b).**

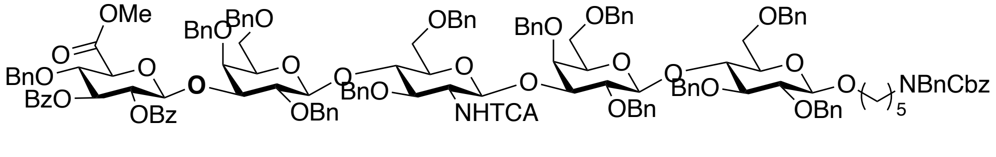


Compound **16b** (90 mg, 0.04 mmol) was reacted with donor **17** (59 mg, 0.08 mmol) followed the general protocol for glycosylations to give **23b** (92 mg, 84%). MALDI-TOF MS *m/z* $C_{151}H_{155}Cl_3N_2O_{31}$ $(M+Na)^+$ calculated 2619. 9577 found 2622.4424.

$^{1}$H NMR (600 MHz, $CDCl_3$) δ 7.89 (d, *J* = 8.1Hz, 2H, Bz), 7.76 (d, *J* = 8.1 Hz, 2H, Bz), 7.48 (t, *J* = 7.3 Hz, 1H, Bz), 7.39 (t, *J* = 7.7 Hz, 1H, Bz), 7.37–7.01 (m, 96H, Ar-H), 6.60 (d, *J* = 6.5 Hz, 1H, -N*H*TCA), 5.72 (t, *J* = 9.6 Hz, 1H, H-3 GlcA), 5.53 (dd, $J_1 = J_2$ = 7.7 Hz, 1H, H-2 GlcA), 5.20 (d, *J* =8.0 Hz, 1H, H-1 GlcA), 5.14 (d, *J* = 12.8 Hz, 2H, $CH_2$ Cbz), 5.06 (d, *J* = 11.8 Hz, 1H, -OCH*H*Ph), 4.99 (d, *J* = 8.4 Hz, 1H, H-1 GlcNAc), 4.97 – 4.91 (m, 2H, -$OCH_2$Ph), 4.82 (d, *J* = 11.8 Hz, 1H, -OCH*H*Ph), 4.77 – 4.41 (m, 16H, -$OCH_2$Ph, -$NCH_2$Ph), 4.38 (d, *J* = 10.1 Hz, 1H, -OCH*H*Ph), 4.35 – 4.10 (m, 15H, -$OCH_2$Ph, H-1 Gal1-2, -OCH*H*Ph, H-1 Glc, H-5 GlcA, H-4 GlcA), 3.97 (t, *J* = 8.7 Hz, 1H, H-4 GlcNAc), 3.94 – 3.81 (m, 4H, H-4 Gal1-2, H-4 Glc, -$CH_2$ linker), 3.76 (-COO*Me*), 3.74 – 3.62 (m, 8H, H-6 GlcNAc, H-2 GlcNAc H-3 GlcNAc, H-3 Gal1-2, H-2 Gal1), 3.58 – 3.52 (m, 3H, H-2 Gal2, H-6 Gal2), 3.48 (t, *J* = 7.9 Hz, 1H, H-$6_a$ Gal1), 3.43 (t, *J* = 8.8 Hz, 1H, H-3 Glc), 3.42 – 3.29 (m, 8H, H-$6_b$ Gal1, H-2 Glc, H-5 GlcNAc, H-5 Gal1,2, -CH*H*- linker, H-$6_a$ Glc), 3.22 – 3.12 (m, 4H, H-$6_b$ Glc, H-5 Glc, -$CH_2$ linker), 1.62 – 1.46 (m, 5H, -$CH_2$ linker), 1.33 – 1.19 (m, 2H, -$CH_2$ linker).

$^{13}$C NMR (151 MHz, $CDCl_3$): *δ* 23.4 ( -$CH_2$ linker), 27.8 ( -$CH_2$ linker), 29.5 (-*C*$H_2$ linker), 46.5 (-*C*$H_2$ linker), 50.9 (-$NCH_2$Ph), 52.7 (-COO*Me*), 57.8 (C2 GlcNAc), 67.2 (-*C*$H_2$ Cbz), 68.2 (C6 Gal1,2, C6 GlcNAc), 68.2 (C6 Glc), 69.5 (-*C*$H_2$), 71.9 (C2 GlcA), 73.1, (-O*C*$H_2$Ph), 73.2 (C5 Gal1,2), 73.2, 73.3, 73.4, 73.9, (-*C*$H_2$Ph), 74.2 (C3 GlcA), 74.5 (C5 GlcA), 74.8, 74.9, 75.00 (-*C*$H_2$Ph), 75.0 (C5 Glc), 75.1 (-*C*$H_2$Ph), 75.2 (C5 GlcNAc), 76.0 (C4 Gal1-2), 76.2 (C4 Glc), 76.3 (C4 GlcNAc), 77.6 (C4 GlcA), 78.1 (C3 GlcNAc), 79.8 – 80.3 (C3 Gal1-2, C2 Gal1-2), 81.8 (C2 Glc), 82.7 (C3 Glc), 100.1 (C1 GlcNAc), 101.6 (C1 GlcA), 102.5 (C1 Gal), 102.8 (C1 Gal2), 103.6 (C1 Glc), 127.1, 127.2, 127.3, 127.4, 127.5, 127.6, 127.7, 127.8, 127.8, 127.9, 127.9, 1278.0, 128.1, 128.2, 128.3, 128.4, 128.5, 129.0, 129.1, 129.2, 129.7, 136.9, 138.0, 138.1, 138.2, 138.3, 138.7, 138.8, 139.0 (C Ar), 139.2 (-COO Cbz), 161.6 (-NHTCA), 165.1 (-OO*C* Bz), 165.5 (-OO*C* Bz), 169.1 (-C6 GlcA).

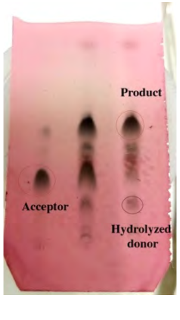

**N-(Benzyl)-benzyloxycarbonyl-5-aminopentyl (methyl 2,3-di-O-benzoyl-4-O-benzyl-β-D-glucopyranosyluronate)-(1→3)-2,4-6-tri-O-benzyl-β-D-galactopyranosyl)-(1→4)-(2-azido-3,6-di-O-benzyl-2-deoxy-β-D-glucopyranosyl)-(1→3)- (2,4-6-tri-O-benzyl-β-D-galactopyranosyl)-(1→4)-(2-azido-3,6-di-O-benzyl-2-deoxy-β-D-glucopyranosyl)-(1→3)-2,4-6-tri-O-benzyl-β-D-galactopyranosyl)-(1→4)-2,3,6-tri-O-benzyl-β-D-glucopyranoside (23c).**

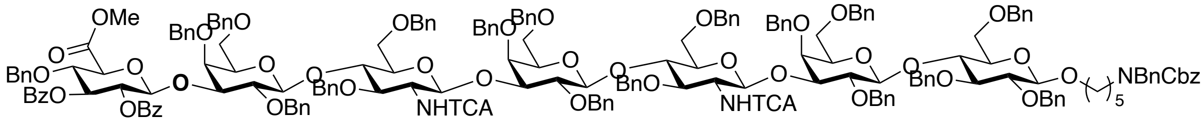


Compound **16c** (60 mg, 0.02 mmol) was reacted with donor **17** (27 mg, 0.039 mmol) according to general glycosylation protocol to give **23c** (62 mg, 89%). MALDI-TOF MS *m/z* $C_{200}H_{205}Cl_6N_3O_{41}$ $(M+K)^+$ calculated 3553.1817, found 3556.4220.

$^1$H NMR (600 MHz, Chloroform-d) δ 7.89 (d, *J* = 7.6 Hz, 2H, Bz), 7.77 (d, *J* = 7.6 Hz, 2H, Bz), 7.48 (t, *J* = 7.5 Hz, 1H, Bz), 7.39 (t, *J* = 7.6 Hz, 1H, Bz), 7.37–7.01 (m, 70H, Ar-H), 6.61 (d, *J* = 7.6 Hz, 1H, -N*H*TCA), 6.58 (d, *J* = 7.6 Hz, 1H, -N*H*TCA), 5.72 (t, *J* = 9.3 Hz, 1H, H-3 GlcA), 5.53 (dd, $J_1$ = $J_2$ = 7.8 Hz, 1H, H-2 GlcA), 5.20 (d, *J* = 7.6 Hz, 1H, H-1 GlcA), 5.14 (d, *J* = 13.0 Hz, 2H, $CH_2$ Cbz), 5.05 (d, *J* = 11.8 Hz, 1H, -OCH*H*Ph), 5.02 (d, *J* = 7.6 Hz, 1H, H-1 GlcNAc2), 5.00 (d, *J* = 7.8 Hz, 1H, H-1 GlcNAc1), 4.98 – 4.91 (m, 2H, -$OCH_2$Ph), 4.87 – 4.72 (d, *J* = 11.8 Hz, 4H, -$OCH_2$Ph), 4.71 – 4.61 (m, 3H, -$CH_2$Ph), 4.57 (d, *J* = 10.8 Hz, 1H, -OCH*H*Ph), 4.54 – 4.37 (m, 8H, -$NCH_2$Ph, -OCH*H*Ph, -$OCH_2$Ph), 4.36 (d, *J* = 7.4 Hz, 1H, H-1 GlcNAc3), 4.35 – 4.16 (m, 8H, -OCH*H*Ph, -$OCH_2$Ph, H-1 Gal1-2, H-1 Glc, H-4 GlcA), 4.16 – 4.11 (m, 2H, H-5 GlcA, -OCH*H*Ph), 4.03 – 3.95 (t, *J* = 8.6 Hz, 2H, H-4 GlcNAc1-2), 3.94 – 3.84 (m, 4H, H-4 Gal1-3, H-4 Glc, -CH*H* linker), 3.76 (-COOMe GlcA), 3.76 – 3.66 (m, 8H, H-6 GlcNAc, H-2 GlcNAc1,2, H-3 GlcNAc1,2, H-3 Gal1-3, H-2 Gal2-3), 3.62 – 3.52 (m, 3H, H-2 Gal2, H-6 Gal3), 3.48 (t, *J* = 8.5 Hz, 1H, H-$6_a$ Gal1), 3.44 (t, *J* = 9.1 Hz, 1H, H-3 Glc), 3.42 – 3.29 (m, 8H, H-$6_b$ Gal1, H-2 Glc, H-5 GlcNAc1-2, H-5 Gal1-3, -C*H*H linker, H-$6_a$ Glc), 3.27 – 3.12 (m, 4H, H-$6_b$ Glc, H-5 Glc, -$CH_2$ linker), 1.68 – 1.44 (m, 5H, -$CH_2$ linker), 1.33 – 1.23 (m, 2H, -$CH_2$ linker).

$^{13}$C NMR (151 MHz, $CDCl_3$): *δ* 23.5 ( -$CH_2$ linker), 27.6 ( -$CH_2$ linker), 29.4 (-*C*$H_2$ linker), 46.6 (-*C*$H_2$ linker), 50.5 (-N*C*$H_2$Ph), 52.4 (-COO*Me*), 57.8 (C2 GlcNAc), 67.1 (-*C*$H_2$ Cbz), 68.1 (C6 Gal1-3, C6 GlcNAc1,2), 68.2 (C6 Glc), 69.7 (-*C*$H_2$ linker), 71.9 (C2 GlcA), 73.1, 73.2 (-O*C*$H_2$Ph),

73.38 (C5 Gal1-3), 73.8, 73.9 (-O*C*H$_2$Ph), 74.1 (C3 GlcA), 74.4 (C5 GlcA), 74.8, 74.9, 75.0 (-O*C*H$_2$Ph), 75.1 (C5 Glc), 75.20 (C5 GlcNAc1-2), 76.1 (C4 GlcNAc1-2), 76.2 (C4 Glc, C4 Gal1-3), 77.6 (C4 GlcA), 78.2 (C3 GlcNAc1-2), 79.8 – 80.3 (C3 Gal1-3, C2 Gal1-3), 81.7 (C2 Glc), 82.6 (C3 Glc), 100.1 (C1 GlcNAc1,2), 101.6 (C1 GlcA), 102.6 (C1 Gal3), 102.8 (C1 Gal1,2), 103.5 (C1 Glc), 127.1, 127.2, 127.3, 127.4, 127.5, 127.6, 127.7, 127.8, 127.9, 128.0, 128.1, 128.2, 128.3, 128.4, 128.5, 129.7, 133.1, 133.3, 137.9. 138.0, 138.1, 138.2, 138.7, 138.8, 139.0 ($C_{Ar}$), 139.1 (-COO Cbz), 162.0 (-NHTCA), 165.2 (-OO*C*$_{Bz}$), 165.5 (-OO*C*$_{Bz}$), 168.5 (-C6$_{GlcA}$).

**N-(Benzyl)-benzyloxycarbonyl-5-aminopentyl (2-O-acetyl-4-O-benzyl-β-D-glucopyranosylurono-6,3-lactone)-(1→3)-2,4,6-tri-O-benzyl-β-D-galactopyranosyl-(1→4)-2,3,6-tri-O-benzyl-β-D-glucopyranoside (24a)**

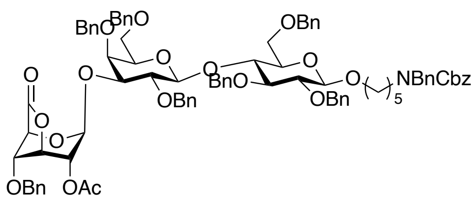


Compound **23a** (110 mg, 0.065 mmol) was subjected to lactonization using the general procedure to yield **24a** (95 mg, 97%). MALDI-TOF MS *m/z* $C_{88}H_{95}NO_{19}$ $(M+Na)^+$ calculated 1504.6396, found 1505.0507.

$^1$H NMR (600 MHz, CDCl$_3$) δ, 7.43–7.01 (m, 56H, Ar-H), 5.57 (s, 1H, H-1 GlcA), 5.18 – 5.12 (m, 3H, CH$_2$ Cbz, H-3 GlcA), 5.00 (d, J = 11.6 Hz, 1H, -OCH*H*Ph), 4.94 (d, J = 11.9 Hz, 1H, -OC*H*HPh), 4.84 – 4.79 (m, 3H, -OC*H*HPh, H-2 GlcA, -OCH*H*Ph), 4.69 (d, J = 11.6 Hz, 1H, -OCH*H*Ph), 4.68 – 4.63 (m, 3H, -OCH*H*Ph, -OCH$_2$Ph), 4.61 – 4.53 (m, 2H, -OCH*H*Ph, -OC*H*HPh), 4.54 – 4.50 (m, 3H, -OCH$_2$Ph, -OCH*H*Ph), 4.49 (d, J = 13.4 Hz, 1H, -OC*H*HPh), 4.45 (d, J = 14.3 Hz, 2H, -NCH$_2$Ph), 4.41 (d, J = 7.4 Hz, 1H, H-1 Gal), 4.35 (m, 2H, -OCH$_2$Ph), 4.29 – 4.17 (H-1 Glc, H-5 GlcA, -OCH$_2$Ph), 4.02 (t, *J* = 4.4 Hz, 1H, H-4 GlcA), 3.93 (t, *J* = 9.3 Hz, 1H, H-4 Glc), 3.89 (d, J = 2.4 Hz, 1H, H-4 Gal), 3.87 – 3.79 (m, 3H, H-3 Gal, H-2 Gal, -CH*H* linker), 3.71 (dd, $J_1$ = 10.7 Hz, $J_2$ = 4.1 Hz, 1H, H-6$_a$ Glc), 3.56 (d, $J_1$ = 10.3 Hz, 1H, H-6$_b$ Glc), 3.53 – 3.46 (m, 2H, H-6a Gal, H-3 Glc), 3.46 – 3.40 (m, 2H, H-5 Gal, -C*H*H linker), 3.37 – 3.29 (m, 2H, H-2 Glc, H-6$_b$ Gal), 3.24 – 3.10 (m, 3H, H-5 Glc, -CH$_2$ linker), 1.59 (s, 3H, C*H*$_3$COO-), 1.59 – 1.44 (m, 4H, -CH$_2$ linker), 1.34 – 1.22 (m, 2H, -CH$_2$ linker).

$^{13}$C NMR (151 MHz, CDCl$_3$): *δ* 20.0 (*C*H$_3$COO-), 23.4 (-*C*H$_2$ linker), 27.7 (-*C*H$_2$ linker), 29.6 (-

*C*$H_2$ linker), 46.5 (-*C*$H_2$ linker), 50.7 (-N*C*$H_2$Ph), 67.1 (-*C*$H_2$ Cbz), 68.1 (C6 Glc), 68.1 (C6 Gal), 68.2 (C5 GlcA), 69.5 (C3 GlcA), 69.6 (-*C*$H_2$ linker), 71.6 (-O*C*$H_2$Ph ), 72.1 (C2 GlcA), 73.1 (C4 GlcA), 73.2 (-O*C*$H_2$Ph ), 73.4 (-O*C*$H_2$Ph ), 73.6 (C5 Gal), 73.7 (-O*C*$H_2$Ph ), 74.1 (-O*C*$H_2$Ph ), 74.5 (-O*C*$H_2$Ph ), 74.9 (-O*C*$H_2$Ph ), 74.9 (C5 Glc), 75.4 (-O*C*$H_2$Ph ), 75.4 (-O*C*$H_2$Ph ), 75.5 (C4 Gal), 76.5 (C4 Glc), 77.6 (C3 Gal), 80.6 (C2 Gal), 81.6 (C2 Glc), 82.8 (C3 Glc), 100.2 (C1 GlcA), 102.7 (C1 Gal), 103.5 (C1 Glc), 126.5, 127.1, 127.4, 127.5, 127.6, 127.7, 127.8, 127.9, 128.0, 128.1, 128.2, 128.2, 128.3, 128.4, 128.5, 128.6, 129.1, 129.2, 129.6, 138.3, 138.7, 139.1 (C Ar), 139.7 (-COO Cbz), 169.4 (C6 GlcA), 170.9 (-*C*OO$CH_3$).

***N*-(Benzyl)-benzyloxycarbonyl-5-aminopentyl (2-O-acetyl-4-O-benzyl-*β*-D-glucopyranosylurono-6-3-lactone)-(1→3)-2,4-6-tri-O-benzyl-*β*-D-galactopyranosyl)-(1→4)-(2-azido-3,6-di-O-benzyl-2-deoxy-*β*-D-glucopyranosyl)-(1→3)-2,4-6-tri-O-benzyl-*β*-D-galactopyranosyl)-(1→4)-2,3,6-tri-O-benzyl-*β*-D-glucopyranoside (24b).**

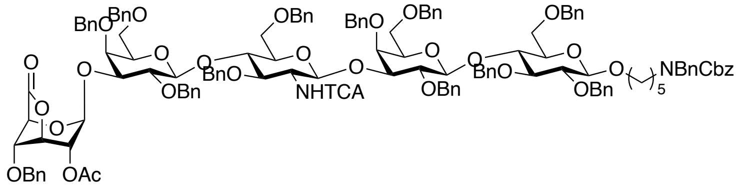


Compound **23b** (80 mg, 0.03 mmol) was subjected to lactonization using the general procedure to yield **24b** (60 mg (82%). MALDI-TOF MS *m/z* $C_{138}H_{145}Cl_3N_2O_{29}$ $(M+Na)^+$ calculated 2425. 0028, found 2424.2439.

$^1$H NMR (600 MHz, $CDCl_3$) δ 7.41–7.04 (m, 86H, Ar-H), 6.56 (d, $J$ = 7.6 Hz, 1H, -N*H*TCA), 5.57 (s, 1H, H-1 GlcA), 5.18 – 5.12 (m, 3H, $CH_2$ Cbz, H-2 GlcA), 5.03 (d, $J$ = 7.2 Hz, 1H, H-1 GlcNAc), 4.99 – 4.91 (m, 4H, -O$CH_2$Ph), 4.88 – 4.78 (m, 3H, H-3 GlcA, -O$CH_2$Ph), 4.74 (d, $J$ = 11.8 Hz, 1H, -OC*H*HPh), 4.71 – 4.60 (m, 6H, -O$CH_2$Ph), 4.53 – 4.41 (m, 9H, -O$CH_2$Ph, H-1 Gal2, -N$CH_2$Ph), 4.35 – 4.12 (m, 11H, -O$CH_2$Ph, H-1 Gal1, H-1 Glc, H-5 GlcA), 4.08 – 4.00 (m, 2H, H-4 GlcA, H-4 GlcNAc), 3.91 – 3.81 (m, 6H, H-4 Gal1-2, H-4 Glc, H-3 Gal1, H-2 Gal2, -OC*H*H linker), 3.76– 3.69 (m, 4H, H-6 GlcNAc, H-2 GlcNAc, H-3 GlcNAc), 3.69 – 3.65 (m, 2H, H-2 Gal1, H-3 Gal1), 3.65 – 3.52 (m, 2H, H-6 Gal2), 3.49 (t, $J$ = 7.7 Hz, 1H, H-$6_a$ Gal1), 3.46 – 3.40 (m, 3H, H-$6_a$ Glc, H-5 Gal2, H-3 Glc), 3.39 – 3.29 (m, 5H, H-$6_b$ Gal, H-2 Glc, H-5 GlcNAc, H-5 Gal, -OCH*H* linker), 3.26 (dd, $J_1$ = 4.8 Hz, $J_2$ = 3.9 Hz, 1H, H-$6_b$ Glc), 3.22 – 3.12 (m, 3H, H-5 Glc, -N$CH_2$ linker), 1.61 – 1.42 (m, 5H, -$CH_2$ linker), 1.33 – 1.18 (m, 2H, -$CH_2$ linker).

$^{13}$C NMR (151 MHz, $CDCl_3$): $\delta$ 23.4 (-*C*$H_2$ linker), 27.7 (-*C*$H_2$ linker), 29.6 (-*C*$H_2$ linker), 46.6 (-*C*$H_2$ linker), 50.38 (-N*C*$H_2$Ph), 57.8 (C2 GlcNAc), 67.1 (-*C*$H_2$ Cbz), 67.8 (C6 GlcNAc), 68.2 (C6 Gal1-2, C6 Glc), 68.3 (C5 GlcA), 69.5 (C3 GlcA), 69.7 (-*C*$H_2$ linker), 71.8 (-O*C*$H_2$Ph), 72.0 (C2 GlcA), 73.0 (C4 GlcA), 73.1 (-O*C*$H_2$Ph), 73.2 (C5 Gal), 73.3, 73.4 (-O*C*$H_2$Ph), 73.8 (C5 Gal2), 74.6, 74.9 (-O*C*$H_2$Ph), 75.0 (C5 Glc), 75.2 (C5 GlcNAc), 75.9 (C4 Glc), 76.3 (C4 Gal1-2), 76.4 (C4 GlcNAc), 76.4 (C3 Gal2), 78.1 (C3 GlcNAc), 79.4 (C3 Gal), 80.3 (C2 Gal2), 81.8 (C2 Glc), 82.7 (C3 Glc), 100.1 (C1 GlcNAc), 100.3 (C1 GlcA), 102.5 (C1 Gal), 103.0 (C1 Gal2), 103.6 (C1 Glc), 126.5, 127.1, 127.2, 127.3, 127.4, 127.5, 127.6, 127.6, 127.7, 127.8, 127.9, 128.0, 128.1, 128.2, 128.3, 128.3, 128.3, 128.4, 128.5, 128.6, 137.93. 138.1, 138.6, 138.7, 138.8, 139.0, 139.30 (C Ar), 139.5 (-COO Cbz), 161.6 (-NHTCA), 169.2 (-C6$_{GlcA}$), 169.4 (-OAc).

**_N_-(Benzyl)-benzyloxycarbonyl-5-aminopentyl (2-O-acetyl-4-O-benzyl-*β*-D-glucopyranosylurono-6-3-lactone)-(1→3)-2,4-6-tri-O-benzyl-*β*-D-galactopyranosyl)-(1→4)-(2-azido-3,6-di-O-benzyl-2-deoxy-*β*-D-glucopyranosyl)-(1→3)- (2,4-6-tri-O-benzyl-*β*-D-galactopyranosyl)-(1→4)-(2-azido-3,6-di-O-benzyl-2-deoxy-*β*-D-glucopyranosyl)-(1→3)-2,4-6-tri-O-benzyl-*β*-D-galactopyranosyl)-(1→4)-2,3,6-tri-O-benzyl-*β*-D-glucopyranoside (24c).**

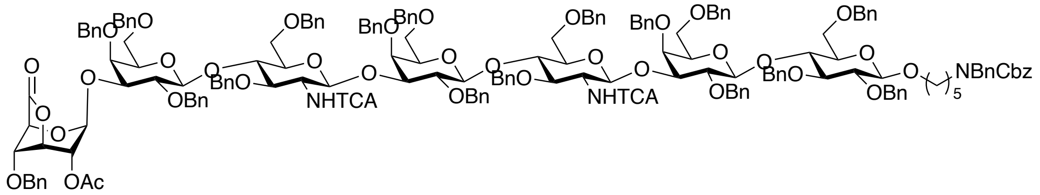


Compound **23c** (60 mg, 0.017 mmol) of was subjected to lactonization using the general procedure to yield **24c** (45 mg, 80%). MALDI-TOF MS *m/z* $C_{187}H_{195}Cl_6N_3O_{39}$ $(M+Na)^+$ calculated 3344.2888, found 3343.6135.

$^1$H NMR (600 MHz, $CDCl_3$) δ 7.45–6.98 (m, 116H, Ar-H), 6.62 (d, *J* = 8.5 Hz, 1H, -N*H*TCA$_{(2)}$), 6.56 (d, *J* = 8.7 Hz, 1H, -N*H*TCA1), 5.57 (s, 1H, H-1 GlcA), 5.18 – 5.12 (m, 3H, $CH_2$ Cbz, H-2 GlcA), 5.06 (d, *J* = 7.1 Hz, 1H, H-1 GlcNAc2), 4.99 (d, *J* = 8.3 Hz, 1H, H-1 GlcNAc1), 4.99 – 4.90 (m, 6H, -O$CH_2$Ph), 4.88 – 4.78 (m, 3H, H-3 GlcA, -O$CH_2$Ph), 4.79 (m, 2H, -O$CH_2$Ph), 4.74 (d, *J* = 11.6 Hz, 1H, -OCH*H*Ph), 4.71 – 4.61 (m, 8H, -O*C*$H_2$Ph), 4.53 – 4.44 (m, 12H, -O*C*$H_2$Ph,

H-1 Gal3, -NCH$_2$Ph), 4.36 – 4.21 (m, 11H, -O*C*H$_2$Ph, H-1 Gal2, H-1 Gal1, H-1 Glc), 4.22 – 4.10 (m, 5H, H-5 GlcA, -O*C*H$_2$Ph), 4.08 – 3.96 (m, 3H, H-4 GlcA, H-4 GlcNAc2, H-4 GlcNAc1), 3.92 – 3.85 (m, 6H, H-4 Gal1-3, H-4 Glc, H-3 Gal3, H-2 Gal2), 3.83 – 3.79 (m, 2H, H-2 Gal3, -CH*H* linker), 3.78– 3.62 (m, 14H, H-6 GlcNAc1-2, H-2 GlcNAc1-2, H-3 GlcNAc1-2, H-2 Gal1-2, H-3 Gal1,2), 3.62 – 3.51 (m, 4H, H-6 Gal2,3), 3.49 (t, $J$ = 7.9 Hz, 1H, H-6$_a$ Gal1), 3.46 – 3.29 (m, 12H, H-6$_b$ Gal1, H-5 Gal3, H-3 Glc, H-2 Glc, H-5 GlcNAc1-2, H-5 Gal1-2, -CH*H* linker), 3.26 (m, 2H H-6 Glc), 3.23 – 3.10 (m, 3H, H-5 Glc, -CH$_2$ linker), 1.64 – 1.21 (m, 6H, -CH$_2$ linker).

$^{13}$C NMR (151 MHz, CDCl$_3$): $\delta$ 23.5 (-*C*H$_2$ linker,), 27.48 (-*C*H$_2$ linker) 29.50 (-*C*H$_2$ linker), 46.4 (-*C*H$_2$ linker), 50.4 (-N*C*H$_2$Ph), 57.7 (C2 GlcNAc), 67.0 (-*C*H$_2$ Cbz), 67.9 (C6 GlcNAc1-2), 68.2 (C6 Gal1-3, C6 Glc), 68.3 (C5 GlcA), 69.5 (C3 GlcA), 69.6 (-*C*H$_2$ linker), 71.8, (-O*C*H$_2$Ph), 71.9 (C2 GlcA), 72.9 (C4 GlcA), 73.1 (-O*C*H$_2$Ph), 73.3 (C5 Gal1), 73.3, 73.4 (-O*C*H$_2$Ph), 73.5 (C5 Gal2), 73.8 (C5 Gal3), 74.5, 74.9 (-O*C*H$_2$Ph), 75.0 (C5 Glc), 75.3 (C5 GlcNAc), 75.5 (C4 Glc), 75.5 (C4 Gal1-3), 76.0 (C4 GlcNAc1), 76.3 (C4 GlcNAc2), 77.3 (C3 Gal3), 78.0 (C3 GlcNAc1-2), 79.6 (C3 Gal2), 79.8 (C3 Gal1), 80.1 (C2 Gal1), 80.3 (C2 Gal2), 80.5 (C2 Gal3), 81.7 (C2 Glc), 82.7 (C3 Glc), 100.1 (C1 GlcNAc2), 100.2 (C1 GlcA), 100.2 (C1 GlcNAc1), 102.5 (C1 Gal1), 102.6 (C1 Gal2), 103.1 (C1 Gal3), 103.4 (C1 Glc), 126.5, 127.2, 127.2, 127.4, 127.5, 127.60, 127.7, 127.8, 127.9, 128.0, 128.1, 128.2, 128.3, 128.4, 128.5, 128.6, 136.6, 137.9. 138.0, 138.1, 138.2, 138.3, 138.6, 138.7, 139.0, 139.2, 139.3 (C Ar), 139.5 (-COO Cbz), 161.6 (-NHTCA$_{(1)}$), 161.7 (-NHTCA2), 169.3 (-OAc), 170.7 (-C6 GlcA).

***N*-(Benzyl)-benzyloxycarbonyl-5-aminopentyl (methyl 2-O-acetyl-4-O-benzyl-*β*-D-glucopyranosyluronate) -(1→3)-2,4,6-tri-O-benzyl-*β*-D-galactopyranosyl-(1→4)-2,3,6-tri-O-benzyl-*β*-D-glucopyranoside (18a).**

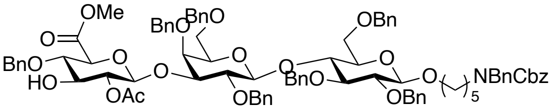


Compound **18a** (90 mg, 97%) was obtained from **24a** (90 mg, 0.061 mmol) according to the general procedure. MALDI-TOF MS *m/z* $C_{90}H_{99}NO_{20}$ (M+Na)$^+$ calculated 1537.7588, found 1537.1222.

$^1$H NMR (600 MHz, CDCl$_3$) δ 7.39–7.04 (m, 48H, Ar-H), 5.14 (d, J = 12.8 Hz, 2H, CH$_2$ Cbz),

4.97 (d, $J$ = 9.2 Hz, 1H, -OCH*H*Ph), 4.92 - 4.86 (m, 3H, -OCH*H*Ph, H-1GlcA, H-2 GlcA), 4.82 (m, 1H, -OCH*H*Ph), 4.77 – 4.59 (m, 4H, $-OCH_2Ph$, -OC*H*HPh), 4.54 (d, $J$ = 12.0 Hz, 1H, -$OCH_2Ph$), 4.49 (d, $J$ = 11.3 Hz, 1H, -OCH*H*Ph), 4.45 (d, $J$ = 16.9 Hz, 2H, $-NCH_2Ph$), 4.36 – 4.25 (m, 3H, $-OCH_2Ph$, H-1 Gal, H-1 Glc), 4.18 (d, $J_1$ = 11.4 Hz, 2H, -OCH*H*Ph), 3.93 – 3.88 (m, 2H, H-4 Glc, H-5 GlcA), 3.87 – 3.83 (m, 1H, -C*H*H linker), 3.83 (d, $J$ = 2.7 Hz, 1H, H-4 Gal), 3.81 (t, $J$ = 8.9 Hz, 1H, H-4 GlcA), 3.73 (s, 3H, -COO*Me*), 3.72 – 3.66 (m, 2H, H-3 GlcA, H-6a Gal), 3.66 – 3.60 (m, 2H, H-3 Gal, H-2 Gal), 3.59 (d, $J$ = 10.0 Hz, 1H, H-6b Gal), 3.52 – 3.46 (m, 2H, H-6a Glc, H-3 Glc), 3.42 (m, 1H, -CH*H* linker), 3.39 – 3.32 (m, 3H, H-5 Gal, H-2 Glc, H-6b Glc), 3.27 (dd, $J_1$ = 2.7 Hz, $J_2$ = 9.6 Hz, 1H, H-5 Glc), 3.26 – 3.12 (m, 2H, $-CH_2$ linker), 1.92 (s, 3H, -OAc), 1.63 – 1.42 (m, 4H, $-CH_2$ linker), 1.35 – 1.26 (m, 2H, $-CH_2$ linker).

$^{13}C$ NMR (151 MHz, $CDCl_3$): $\delta$ 20.8 (*C*$H_3$COO-), 23.6 (-*C*$H_2$ linker), 27.7 (-*C*$H_2$ linker) 29.6 (-*C*$H_2$ linker), 46.3 (-*C*$H_2$ linker), 50.2 ($-NCH_2Ph$), 52.6 (-OMe GlcA), 67.1 (-*C*$H_2$ Cbz), 68.2 (C6 Glc), 68.3 (C6 Gal), 69.5 (-*C*$H_2$ linker), 73.1 (C5 Gal), 73.2 ($-OCH_2Ph$), 73.4 ($-OCH_2Ph$), 74.1 (C2 GlcA), 74.2 (C5 GlcA), 74.8 ($-OCH_2Ph$), 74.9 ($-OCH_2Ph$), 75.0 ($-OCH_2Ph$), 75.1 (C5 Glc), 75.1 ($-OCH_2Ph$), 75.4 (C3 GlcA), 75.4 (C5 Glc), 75.5 ($-OCH_2Ph$), 76.2 (C4 Gal), 76.5 (C4 Glc), 79.8 (C4 GlcA), 80.2 (C2 Gal), 80.3 (C3 Gal), 81.7 (C2 Glc), 82.9 (C3 Glc), 101.3 (C1 GlcA), 102.5 (C1 Gal), 103.6 (C1 Glc), 127.1, 127.3, 127.5, 127.6, 127.8, 127.9, 128.1, 128.2, 128.3, 128.4, 128.5, 138.1, 138.4, 138.7 (C Ar), 139.2 (-COO Cbz), 168.5 (-*C*OOMe GlcA), 170.9 (-*C*OO$CH_3$ OAc).

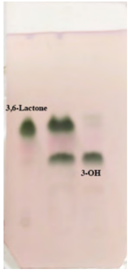

***N*-(Benzyl)-benzyloxycarbonyl-5-aminopentyl (methyl 2-O-acetyl-4-O-benzyl-*β*-D-glucopyranosyluronate)-(1→3)-2,4-6-tri-O-benzyl-*β*-D-galactopyranosyl)-(1→4)-(2-azido-3,6-di-O-benzyl-2-deoxy-*β*-D-glucopyranosyl)-(1→3)-2,4-6-tri-O-benzyl-*β*-D-galactopyranosyl)-(1→4)-2,3,6-tri-O-benzyl-*β*-D-glucopyranoside (18b).**

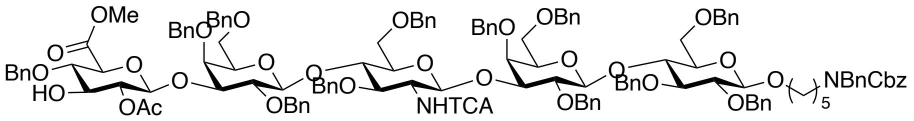


Compound **18b** (49 mg, 0.018 mmol) was obtained from **24b** (55 mg, 90%) according to the general procedure. MALDI-TOF MS *m/z* $C_{139}H_{149}Cl_3N_2O_{30}$ $(M+Na)^+$ calculated 2457. 0448, found 2456.3970.

$^1$H NMR (600 MHz, $CDCl_3$) δ 7.38–7.04 (m, 75H, Ar-H), 6.6 (d, *J* = 8.1 Hz, 1H, -*NH*TCA), 5.14 (d, *J* = 12.7 Hz, 2H, $CH_2$ Cbz), 5.1 (d, *J* = 7.5 Hz, 1H, H-1 GlcNAc), 4.9 (d, *J* = 9.2 Hz, 1H, -OC*H*HPh), 4.92 - 4.86 (m, 5H, -OC*H*HPh, -$OCH_2Ph$, H-1 GlcA, H-2 GlcA), 4.8 (m, 4H, -$OCH_2Ph$), 4.70 – 4.60 (m, 5H, -$OCH_2Ph$, -OC*H*HPh), 4.51 – 4.41 (m, 7H, -OC*H*HPh, -$OCH_2Ph$, -$NCH_2Ph$), 4.38 (d, *J* = 7.1 Hz, 1H, H-1Gal2), 4.35 (d, J = 7.2 Hz, 1H, H-1 Gal1), 4.33 – 4.29 (m, 3H, -OC*H*HPh, -$OCH_2Ph$), 4.27 – 4.22 (m, 2H, H-1 Glc, OC*H*HPh), 4.19 (d, *J* = 11.7 Hz, 1H, OCH*H*Ph), 4.13 (d, J = 11.8 Hz, 1H, OCH*H*Ph), 4.03 (t, *J* = 8.8 Hz, 1H, H-4 GlcNAc), 3.92 – 3.83 (m, 4H, H-4 Glc, H-5 GlcA, H-4 Gal1-2), 3.87 – 3.83 (m, 1H, -OC*H*H linker), 3.82 (t, *J* = 8.7 Hz, 1H, H-4 GlcA), 3.78 – 3.60 (m, 13H, H-3 GlcNAc, H-6a GlcNAc, H-2 GlcNAc, H-3 GlcA, H-3 Gal1-2, H-2 Gal1-2, H-6 Gal1-2, -COO*Me*), 3.56 (d, *J* = 10.1 Hz, 1H, H-6b Gal), 3.50 (m, 1H, H-6a Glc), 3.46 – 3.42 (m, H-3 Glc, H-5 GlcNAc), 3.42 – 3.30 (m, 6H, -OC*H*H linker, H-6b Gal, H-5 Gal1-2, H-2 Glc), 3.28 – 3.10 (m, 4H, H-6b Glc, H-5 Glc, -$CH_2$ linker), 1.90 (s, 3H, -OAc), 1.64 – 1.42 (m, 4H, -$CH_2$ linker), 1.34 – 1.19 (m, 2H, -$CH_2$ linker).

$^{13}$C NMR (151 MHz, $CDCl_3$): *δ* 21.0 (-COO*C*$H_3$), 23.1 (-*C*$H_2$ linker), 27.8 (-*C*$H_2$ linker), 29.5 (-*C*$H_2$ linker), 46.7 (-*C*$H_2$ linker), 50.4 (-N*C*$H_2$Ph), 52.4 (-OMe GlcA), 57.8 (C2 GlcNAc), 67.1 (-*C*$H_2$ Cbz), 67.9 (C6 GlcNAc), 68.2 (C6 Glc, C6 Gal1-2), 69.6 (-*C*$H_2$ linker), 73.1 (-O*C*$H_2$Ph), 73.2 (C5 Gal1,2), 73.3 (-O*C*$H_2$Ph), 73.4 (-O*C*$H_2$Ph), 73.8 (C2 GlcA), 74.1 (C5 GlcA), 74.9 (-O*C*$H_2$Ph), 75.0 (-O*C*$H_2$Ph), 75.0 (C5 Glc), 75.1 (-O*C*$H_2$Ph), 75.3 (C5 GlcNAc), 75.4 (C3 GlcA), 75.5 (-O*C*$H_2$Ph), 76.2 (C4 Gal1-2), 76.3 (C4 Glc), 76.5 (C4 GlcNAc), 78.1 (C3 GlcNAc), 79.7 (C4 GlcA), 79.9 (C3 Gal1-2), 80.0 (C3 Gal1-2), 80.3 (C2 Gal1-2), 80.3 (C2 Gal1-2), 81.7 (C2 Glc),

82.7 (C3 Glc), 101.1 (C1 GlcNAc), 101.3 (C1 GlcA), 102.5 (C1 Gal1), 102.7 (C1 Gal2), 103.6 (C1 Glc), 127.4, 127.5, 127.6, 127.7, 127.8, 127.9, 128.0, 128.1, 128.2, 128.3, 128.4, 128.55, 128.6 137.6, 137.9, 138.0, 138.2, 138.3, 138.7, 138.9 (C Ar), 139.0 (-COO Cbz), 161.5 (-NHTCA), 168.5 (-<u>C</u>OOMe GlcA), 170.5 (-<u>C</u>OOCH$_3$ OAc).

***N*-(Benzyl)-benzyloxycarbonyl-5-aminopentyl (methyl 2-O-acetyl-4-O-benzyl-*β*-D-glucopyranosyluronate)-(1→3)-2,4-6-tri-O-benzyl-*β*-D-galactopyranosyl)-(1→4)-(2-azido-3,6-di-O-benzyl-2-deoxy-*β*-D-glucopyranosyl)-(1→3)-(2,4-6-tri-O-benzyl-*β*-D-galactopyranosyl)-(1→4)-(2-azido-3,6-di-O-benzyl-2-deoxy-*β*-D-glucopyranosyl)-(1→3)-2,4-6-tri-O-benzyl-*β*-D-galactopyranosyl)-(1→4)-2,3,6-tri-O-benzyl-*β*-D-glucopyranoside (18c).**

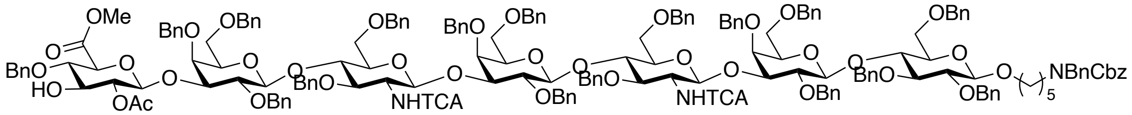


Compound **18c** (40 mg, 0.011 mmol) was obtained from **24c** (33 mg, 0.009 mmol, 84%) according to general procedure. MALDI-TOF MS *m/z* $C_{188}H_{199}Cl_6N_3O_{40}$ $(M+Na)^+$ calculated 3376.3308, found 3374.7163.

$^1$H NMR (600 MHz, CDCl$_3$) δ 7.37–7.04 (m, 107H, Ar-H), 6.65 (d, *J* = 8.4 Hz, 1H, -*<u>NH</u>*TCA$_2$), 6.56 (d, *J* = 8.4 Hz, 1H, -*<u>NH</u>*TCA$_1$), 5.14 (d, *J* = 12.2 Hz, 2H, CH$_2$ Cbz), 5.11 (d, *J* = 7.9 Hz, 1H, H-1 GlcNAc2), 5.01 (d, *J* = 7.4 Hz, 1H, H-1 GlcNAc1), 4.99 – 4.90 (m, 6H, -OCH$_2$Ph, H-1 GlcA, H-2 GlcA), 4.97 (d, *J* = 10.4 Hz, 1H, -OC*<u>H</u>*HPh), 4.84 – 4.62 (m, 11H, -O*<u>C</u>*H$_2$Ph, -OC*<u>H</u>*HPh), 4.53 – 4.41 (m, 10H, -OCH$_2$Ph, -NCH$_2$Ph), 4.38 (d, *J* = 7.2 Hz, 2H, H-1 Gal2-3), 4.35 – 4.29 (m, 5H, H-1 Gal1, -OCH$_2$Ph), 4.25 (m, 3H, H-1 Glc, -OCH$_2$Ph), 4.19 (d, *J* = 11.9 Hz, 1H, -OC*<u>H</u>*HPh), 4.16 – 4.12 (m, 2H, -OCH$_2$Ph), 4.02 (m, 2H, H-4 GlcNAc1-2), 3.92 – 3.85 (m, 5H, H-4 Glc, H-5 GlcA, H-4 Gal1-3), 3.87 – 3.79 (m, 3H, -OC*<u>H</u>*H linker, H-3 GlcNAc2, H-4 GlcA), 3.78 – 3.58 (m, 18H, H-3 GlcNAc1, H-6a GlcNAc1-2, H-2 GlcNAc1-2, H-3 GlcA, H-3 Gal1-3, H-2 Gal1-3, H-6 Gal1-2, -COO*<u>Me</u>*), 3.56 (d, *J* = 10.0 Hz, 1H, H-6b Gal), 3.50 (m, 1H, H-6a Glc), 3.46 – 3.42 (m, H-3 Glc, H-5 GlcNAc), 3.42 – 3.30 (m, 6H, -OCH*<u>H</u>* linker, H-6b Gal, H-5 Gal1-2, H-2 Glc), 3.28 – 3.10 (m, 4H, H-6b Glc, H-5 Glc, -CH$_2$ linker), 1.90 (s, 3H, -OAc), 1.64 – 1.42 (m, 4H, -CH$_2$ linker), 1.34 – 1.19 (m, 2H, -CH$_2$ linker).

$^{13}C$ NMR (151 MHz, $CDCl_3$): $\delta$ 21.0 (-COO$\underline{C}H_3$), 23.5 (-$\underline{C}H_2$ linker), 27.9 (-$\underline{C}H_2$ linker), 29.4 (-$\underline{C}H_2$ linker), 46.5 (-$\underline{C}H_2$ linker), 50.3 (-$NCH_2Ph$), 52.4 (-OMe GlcA), 57.9 (C2 GlcNAc), 67.0 (-$\underline{C}H_2$ Cbz), 67.8 (C6 GlcNAc), 68.0 (C6 Glc), 68.3 (C6 Gal1-2), 69.7 (-$\underline{C}H_2$ linker), 72.9 (-O$\underline{C}H_2$Ph), 73.2 (C5 Gal1-2), 73.3 (-O$\underline{C}H_2$Ph), 73.4 (-O$\underline{C}H_2$Ph), 73.9 (C2 GlcA), 74.0 (C5 GlcA), 74.3 (-O$\underline{C}H_2$Ph), 75.0 (-O$\underline{C}H_2$Ph), 75.2 (C5 Glc), 75.1 (-O$\underline{C}H_2$Ph), 75.3 (C5 GlcNAc1-2), 75.5 (C3 GlcA), 76.3 (C4 Gal1-2, C4 Glc), 76.4 (C4 GlcNAc), 77.7 (C3 GlcNAc), 79.9 (C4 GlcA), 80.5 (C2 Gal1-3), 81.8 (C2 Glc), 82.9 (C3 Glc), 100.2 (C1 GlcNAc), 100.3 (C1 GlcNAc), 101.2 (C1 GlcA), 102.5 (C1 Gal1), 102.8 (C1 Gal2-3), 103.5 (C1 Glc), 127.4, 127.5, 127.5, 127.6, 127.7, 127.9, 127.9, 128.0, 128.1, 128.2, 128.3, 128.4, 128.5, 128.6, 137.7, 137.9, 138.0, 138.2, 138.3, 138.7, 138.9 (C Ar), 139.0 (-COO Cbz), 161.5 (-NHTCA), 168.5 (-$\underline{C}$OOMe GlcA), 170.5 (-$\underline{C}OOCH_3$ OAc).

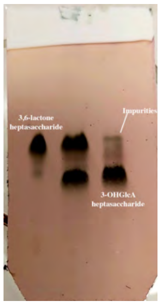


**5-aminopentyl 3-O-sulfo-*β*-D-glucopyranosyluronate-(1→3)-*β*-D-galactopyranosyl-(1→4)-*β*-D-glucopyranoside (1).**

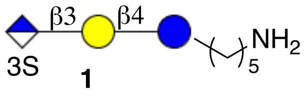

**1**

Compound **1** (6 mg, 66%) of was obtained from **18a** (20 mg, 0.0135 mmol) using the general protocols for *O*-sulfation, hydrogenation and saponification. ESI TOF-MS *m/z* $C_{23}H_{41}NO_{20}S$ $(M - H)^-$ calculated 682.6205, found 682.3445.

$^1H$ NMR (600 MHz, $D_2O$) δ: (non carbohydrate): 3.87 (-$OCH_2$ linker), 2.94 (t, $J$ = 7.5 Hz, 2H, -$CH_2$ linker), 1.69 – 1.56 (m, 4H, -$CH_2$ linker), 1.43 – 1.35 (m, 2H, -$CH_2$ linker).

$^{13}C$ NMR (151 MHz, $D_2O$): $\delta$ (non carbohydrate): 70.1 (-O$\underline{C}H_2$ linker), 39.6 (-$\underline{C}H_2$ linker), 28.2 (-

*C*H2 linker), 26.1 (-*C*H2 linker), 22.3 (-*C*H2 linker).

| | H1 | H2 | H3 | H4 | H5 | H*H*-6 | *H*H-6 |
|---|---|---|---|---|---|---|---|
| Glc | 4.42 d, $J = 7.8$ Hz | 3.25 t, $J = 8.3$ Hz | 3.59 | 3.60 | 3.67 | 3.92, dd, $J_1 = 2.1$ Hz, $J_2 = 12.5$ Hz, | 3.74 |
| Gal | 4.45 d, $J = 8.0$ Hz | 3.64 | 3.77 | 4.12 d, $J = 2.4$ Hz | 3.52 | 3.70 | 3.68 |
| GlcA | 4.70 d, $J = 8.5$ Hz | 3.53 | 4.27 t, $J = 9.3$ Hz | 3.63 | 3.72 | | |

| | C1 | C2 | C3 | C4 | C5 | C6 |
|---|---|---|---|---|---|---|
| Glc | 102.1 | 72.8 | 74.6 | 78.3 | 75.2 | 60.1 |
| Gal | 102.5 | 70.2 | 82.3 | 68.1 | 71.9 | 60.8 |
| GlcA | 103.2 | 74.7 | 83.8 | 70.2 | 76.1 | 175.3 |

**5-aminopentyl 3-O-sulfo-*β*-*D*-glucopyranosyluronate-(1→3)-*β*-D-galactopyranosyl)-(1→4)-(2-acetamido-2-deoxy-*β*-D-glucopyranosyl-(1→3)-*β*-D-galactopyranosyl-(1→4)-*β*-D-glucopyranoside (2).**

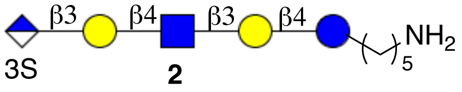


Compound **2** (4 mg, 0.005 mmol, 76%) was obtained from **18b** (13 mg) using the general protocols for *O*-sulfation, hydrogenation and saponification.

ESI TOF-MS *m/z* $C_{37}H_{64}N_2O_{30}S$ (M - H)$^-$ calculated 1047.3192, found 1047.5192.

$^1$H NMR (600 MHz, $D_2O$) δ: (non carbohydrate): 3.85 (-OCH*H* linker), 3.60 (-OC*H*H linker), 2.93 (t, $J = 7.5$ Hz, 2H, -CH2 linker), 1.95 (s, 3H, -NHAc), 1.65 – 1.56 (m, 4H, -CH2 linker), 1.34 (m, 2H, -CH2 linker).

$^{13}$C NMR (151 MHz, $D_2O$): *δ* (non carbohydrate): 70.27 (-OCH2 linker), 39.36 (-CH2 linker), 28.53 (-CH2 linker), 26.43 (-CH2 linker), 22.53 (-NHAc), 22.35 (-CH2 linker).

|  | H1 | H2 | H3 | H4 | H5 | H6 |
|---|---|---|---|---|---|---|
| Glc | 4.40 *J* = 8.0 Hz | 3.22 | 3.56 | 3.57 | 3.63 | 3.89 – 3.67 |
| Gal | 4.35 *J* = 7.6 Hz | 3.50 | 3.64 | 4.07 d, *J* = 2.5 Hz | n/a | 3.89 – 3.67 |
| GlcNAc | 4.62 *J* = 8.5 Hz | 3.72 | 3.64 | 3.67 | n/a | 3.89 – 3.67 |
| Gal | 4.46 *J* = 8.0 Hz | 3.66 | 3.75 | 4.10 d, *J* = 2.8 Hz | n/a | 3.89 – 3.67 |
| GlcA | 4.68 | 3.50 | 4.25 t, *J* = 9.4 Hz | 3.63 | 3.71 | 3.89 – 3.67 |

|  | C1 | C2 | C3 | C4 | C5 | C6 |
|---|---|---|---|---|---|---|
| Glc | 101.95 | 72.84 | 74.76 | n/a | n/a | 59.56 – 62.01 |
| Gal | 103.12 | 78.37 | 82.21 | 68.65 | n/a | 59.56 – 62.01 |
| GlcNAc | 102.94 | 55.37 | 74.93 | n/a | n/a | 59.56 – 62.01 |
| Gal | 102.77 | 78.19 | 82.39 | 68.30 | n/a | 59.56 – 62.01 |
| GlcA | 103.68 | 75.11 | 84.02 | 75.28 | 76.39 | 59.56 – 62.01 |

**5-aminopentyl 3-O-sulfo-*β*-D-glucopyranosyluronate-(1→3)-*β*-D-galactopyranosyl-(1→4)-(2-acetamido-2-deoxy-*β*-D-glucopyranosyl-(1→3)-*β*-D-galactopyranosyl-(1→4)-(2-acetamido-2-deoxy-*β*-D-glucopyranosyl-(1→3)-*β*-D-galactopyranosyl-(1→4)-*β*-D-glucopyranoside (3).**

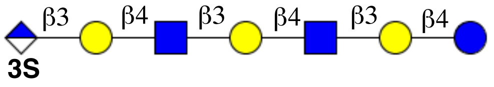


Compound **5** (5 mg, 84%) was obtained from **18c** (15 mg, 0.0042 mmol) using the general protocols for *O*-sulfation, hydrogenation and saponification. ESI TOF-MS *m/z* $C_{51}H_{86}N_3O_{40}S$ (M - 2H)$^{2-}$ calculated 705.7220, found 705.8901.

$^1$H NMR (600 MHz, $D_2O$) δ: (non carbohydrate): 3.85 (-$OCH_2$ linker), 2.93 (t, *J* = 7.7 Hz, 2H, -$CH_2$ linker), 1.95 (s, 6H, 2x-NHAc), 1.65 – 1.55 (m, -$CH_2$ linker), 1.41 – 1.34 (m, 2H, -$CH_2$ linker).

$^{13}$C NMR (151 MHz, $D_2O$): δ (non carbohydrate): 70.2 (-O$\underline{C}H_2$ linker), 39.2 (-$\underline{C}H_2$ linker), 28.2 (-$\underline{C}H_2$ linker), 26.4 (-$\underline{C}H_2$ linker), 22.2 ($\underline{C}H_3$CO-), 22.2 (-$\underline{C}H_2$ linker).

Carbohydrate region:

| | H1 | H2 | H3 | H4 | H5 | H6 |
|---|---|---|---|---|---|---|
| Glc | 4.41 d, $J$ = 8.4 Hz | 3.21, t, $J$ = 8.9 Hz | 3.56 | 3.57 | na | 3.87 3.77 |
| Gal | 4.35 d, $J$ = 8.0 Hz | 3.51 | 3.64 | 4.07, d, $J$ = 2.6 Hz | na | 3.70 – 3.53 |
| GlcNAc | 4.62 d, $J$ = 8.1 Hz | 3.72 | 3.65 | 3.67 | na | 3.89 |
| Gal2 | 4.39 d, $J$ = 8.5 Hz | 3.52 | 3.65 | 4.08, d, $J$ = 2.6 Hz | na | 3.70 – 3.53 |
| GlcNAc2 | 4.62 d, $J$ = 8.1 Hz | 3.72 | 3.65 | 3.67 | na | 3.89 |
| Gal3 | 4.46, d, $J$ = 7.9 Hz, | 3.63 | 3.75 | 4.11, d, $J$ = 3.0 Hz | na | 3.70 – 3.53 |
| GlcA | 4.68 | 3.54 | 4.24 t, $J$ = 9.2 Hz | 3.61 | 3.71 | |

| | C1 | C2 | C3 | C4 | C5 | C6 |
|---|---|---|---|---|---|---|
| Glc | 101.9 | 72.8 | 74.7 | 78.2 | na | 59.9 |
| Gal | 102.9 | 69.8 | 82.1 | 68.4 | na | 60.9 |
| GlcNAc | 102.8 | 55.2 | 75.1 | 78.1 | na | 59.8 |
| Gal2 | 102.8 | 69.8 | 82.1 | 68.4 | na | 60.8 |
| GlcNAc2 | 102.8 | 55.2 | 75.1 | 78.1 | na | 59.8 |
| Gal3 | 102.4 | 70.2 | 82.4 | 68.4 | na | 60.8 |
| GlcA | 103.2 | 71.2 | 83.9 | 75.1 | 76.2 | |

**5-aminopentyl *β*-D-glucopyranosyluronate-(1→3)-*β*-D-galactopyranosyl-(1→4)-*β*-D-glucopyranoside (4).**

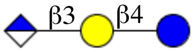


Compound **4 (**3 mg, 73%) of was obtained from **18a** (10 mg, 0.007 mmol) using the general protocols for hydrogenation and saponification. ESI TOF-MS *m/z* $C_{23}H_{41}NO_{17}$ $(M - H)^-$ calculated 602.2302, found 602.3908.

$^1$H NMR (600 MHz, $D_2O$) δ: (non carbohydrate): 3.85 (-OCH*H* linker), 3.60 (-OC*H*H linker), 2.93 (t, $J$ = 7.5 Hz, 2H, -$CH_2$ linker), 1.69 – 1.56 (m, 4H, -$CH_2$ linker), 1.43 - 1.34 (m, 2H, -$CH_2$ linker).

$^{13}$C NMR (151 MHz, $D_2O$): $\delta$ (non carbohydrate): 70.1 (-O$\underline{C}$H$_2$ linker), 39.4 (-$\underline{C}$H$_2$ linker), 26.2 (-$\underline{C}$H$_2$ linker), 25.8 (-$\underline{C}$H$_2$ linker), 22.1 (-$\underline{C}$H$_2$ linker).

| | **H1** | **H2** | **H3** | **H4** | **H5** | **H6a** | **H6b** |
|---|---|---|---|---|---|---|---|
| Glc | 4.41<br>d, *J* = 8.0 Hz | 3.23<br>t, *J* = 8.3 Hz | 3.56 | 3.57 | 3.64 | 3.90 | 3.73 |
| Gal | 4.44<br>d, *J* = 8.4 Hz | 3.63 | 3.73 | 4.11<br>d, *J* = 3.0 Hz | 3.51 | 3.67 | 3.64 |
| GlcA | 4.59<br>d, *J* = 7.8 Hz | 3.33<br>t, *J* = 8.1 Hz | 3.44 | 3.44 | 3.65 | | |

| | **C1** | **C2** | **C3** | **C4** | **C5** | **C6** |
|---|---|---|---|---|---|---|
| Glc | 102.5 | 72.9 | 74.6 | 78.3 | 75.1 | 60.1 |
| Gal | 102.6 | 76.7 | 82.3 | 68.1 | 74.8 | 60.3 |
| GlcA | 103.6 | 73.2 | 75.4 | 71.9 | 75.7 | 175.5 |

**5-aminopentyl *β-D*-glucopyranosyluronate-(1→3)-*β*-D-galactopyranosyl)-(1→4)-(2-acetamido-2-deoxy-*β*-D-glucopyranosyl-(1→3)-*β*-D-galactopyranosyl-(1→4)-*β*-D-glucopyranoside (5).**

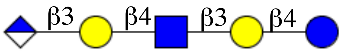


compound **5** (3 mg, 62%) was obtained from **18b** (13 mg, 0.005 mmol) using the general protocols for hydrogenation and saponification. ESI TOF-MS *m/z* $C_{37}H_{64}N_2O27$ (M - H)$^-$ calculated 967.3624, found 967.5421.

$^1$H NMR (600 MHz, $D_2O$) δ: (non carbohydrate): 3.85 (-OCH$\underline{H}$ linker), 3.60 (-OC$\underline{H}$H linker), 2.93 (t, *J* = 7.5 Hz, 2H, -CH$_2$ linker), 1.95 (s, 3H, -NHAc), 1.65 – 1.56 (m, 4H, -CH$_2$ linker), 1.34 (m, 2H, -CH$_2$ linker).

$^{13}$C NMR (151 MHz, $D_2O$): $\delta$ (non carbohydrate): 70.3 (-O$\underline{C}$H$_2$ linker), 39.4 (-$\underline{C}$H$_2$ linker), 28.5 (-$\underline{C}$H$_2$ linker), 26.4 (-$\underline{C}$H$_2$ linker), 22.4 (-$\underline{C}$H$_2$ linker), 22.53 (-NHAc).

|  | H1 | H2 | H3 | H4 | H5 | H6 |
|---|---|---|---|---|---|---|
| Glc | 4.40<br>d, $J$ = 8.0 Hz | 3.22<br>t, $J$ = 8.4 Hz | 3.56 | 3.64 | na | 3.77 – 3.65 |
| Gal | 4.35<br>d, $J$ = 7.8 Hz | 3.51 | 3.63 | 4.08<br>d, J = 2.9 Hz | na | 3.77 – 3.65 |
| GlcNAc | 4.62<br>d, $J$ = 8.3 Hz | 3.72 | 3.54 | 3.66 | na | 3.88 |
| Gal2 | 4.45<br>d, J = 7.8 Hz | 3.61 | 3.73 | 4.11<br>d, J = 3.0 Hz | na | 3.77 – 3.65 |
| GlcA | 4.60<br>d, J = 7.9 Hz | 3.33 | 3.44 | 3.43 | na | - |

**5-aminopentyl *β*-D-glucopyranosyluronate-(1→3)-*β*-D-galactopyranosyl-(1→4)-(2-acetamido-2-deoxy-*β*-D-glucopyranosyl-(1→3)-*β*-D-galactopyranosyl-(1→4)-(2-acetamido -2-deoxy-*β*-D-glucopyranosyl-(1→3)-*β*-D-galactopyranosyl-(1→4)-*β*-D-glucopyranoside (6).**

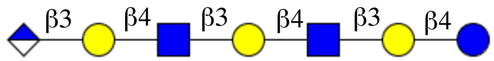


Compound **6** (3 mg, 54%) was obtained from **18c (**15 mg, 0.0042 mmol) using the general protocols for hydrogenation and saponification. ESI TOF-MS *m/z* $C_{51}H_{87}N_3O_{37}$ $(M - H)^-$ calculated 1332.4946, found 1332.7584.

$^1$H NMR (600 MHz, $D_2O$) δ: (non carbohydrate): 3.85 (-$OCH_2$ linker), 2.92 (t, $J$ = 7.5 Hz, 2H, -$CH_2$ linker), 1.95 (s, 6H, 2x-NHAc), 1.64 – 1.56 (m, -$CH_2$ linker), 1.41 – 1.33 (m, 2H, -$CH_2$ linker).

$^{13}$C NMR (151 MHz, $D_2O$): $\delta$ (non carbohydrate): 70.0 (-O$\underline{C}H_2$ linker), 39.3 (-$\underline{C}H_2$ linker), 28.2 (-$\underline{C}H_2$ linker), 26.3 (-$\underline{C}H_2$ linker), 21.9 ($\underline{C}H_3$CO-), 22.1 (-$\underline{C}H_2$ linker).

|  | H1 | H2 | H3 | H4 | H5 | H6 |
|---|---|---|---|---|---|---|
| Glc | 4.38<br>d, $J$ = 8.7 Hz | 3.22, t,<br>$J$ = 7.7 Hz | 3.55 | 3.57 | 3.63 | 3.82<br>3.77 |
| Gal | 4.35<br>d, $J$ = 7.6 Hz | 3.62 | 3.64 | 4.01 – 3.99 | na | 3.67 |
| GlcNAc | 4.62<br>d, $J$ = 8.5 Hz | 3.73 | 3.50 | 3.67 | 3.64 | 3.86 |
| Gal2 | 4.35<br>d, $J$ = 7.1 Hz | 3.54 | 3.57 | 4.01 – 3.99 | na | 3.67 |
| GlcNAc2 | 4.62<br>d, $J$ = 8.5 Hz | 3.73 | 3.50 | 3.67 | 3.64 | 3.86 |

| | | | | | | |
|---|---|---|---|---|---|---|
| Gal3 | 4.45, d, $J$ = 7.1 Hz, | 3.62 | 3.74 | 4.11, d, $J$ = 3.4 Hz | na | 3.67 |
| GlcA | 4.06 d, $J$ = 8.0 Hz, | 3.33, t, J = 8.2 Hz | 3.43 | 3.44 | 3.64 | |

| | C1 | C2 | C3 | C4 | C5 | C6 |
|---|---|---|---|---|---|---|
| Glc | 101.8 | 72.9 | 74.6 | 78.6 | 74.9 | 59.7 |
| Gal | 102.9 | 74.9 | 82.3 | 68.2 | na | 60.9 |
| GlcNAc | 102.6 | 55.1 | 75.0 | 78.2 | 75.4 | 59.8 |
| Gal2 | 102.8 | 74.4 | 82.3 | 68.2 | na | 60.9 |
| GlcNAc2 | 102.6 | 55.1 | 75.0 | 78.2 | 75.4 | 59.8 |
| Gal3 | 102.4 | 74.9 | 82.4 | 68.4 | na | 60.9 |
| GlcA | 103.4 | 73.3 | 71.9 | 75.4 | 74.9 | - |

**5-aminopentyl 3-O-sulfo-*β-D*-glucopyranosyl-(1→3)-*β*-D-galactopyranosyl)-(1→4)-2-acetamido-2-deoxy-*β*-D-glucopyranosyl-(1→3)-*β*-D-galactopyranosyl-(1→4)-*β*-D-glucopyranoside (7).**

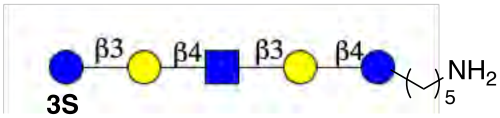


Compound **18b** (10 mg, 0.0037 mmol) was sulfated and hydrogenated according to the general protocol for *O*-sulfation and hydrogenation protocols. The intermediate was treated with $NaBH_4$ (2.6 mg, 0.07 mmol) to reduce methyl glucuronate to glucose. After completion of the reduction, excess of $NaBH_4$ was quenched with AcOH and the solid material were removed by centrifugation after which the supernatant was freeze-dried. The residue was applied to Biogel-P-2 size exclusion column chromatography and eluted with 100 mM $NH_4HCO_3$ solution to obtain **7** (2 mg 52% yield over 3 steps). ESI TOF-MS *m/z* $C_{37}H_{66}N_2O_{29}S$ $(M - H)^-$ calculated 1033.3399, found 1033.5229.

$^1$H NMR (600 MHz, $D_2O$) δ: (non carbohydrate): 3.85 (-OCH<u>*H*</u> linker), 3.60 (-OC<u>*H*</u>H linker), 2.93 (t, $J$ = 7.5 Hz, 2H, $-CH_2$ linker), 1.95 (s, 3H, -NHAc), 1.65 – 1.56 (m, 4H, $-CH_2$ linker), 1.34 (m, 2H, $-CH_2$ linker).

$^{13}$C NMR (151 MHz, $D_2O$) δ: (non carbohydrate): 70.3 (-O<u>*C*</u>H2 linker), 39.4 (-<u>*C*</u>H2 linker), 28.5 (-<u>*C*</u>H2 linker), 26.4 (-<u>*C*</u>H2 linker), 22.5 (-<u>*C*</u>H3CO), 22.35 (-<u>*C*</u>H2 linker).

| | H1 | H2 | H3 | H4 | H5 | H6 |
|---|---|---|---|---|---|---|
| Glc | 4.41<br>d, $J$ = 8.2 Hz | 3.23,<br>t, $J$ = 8.6 Hz | 3.57 | 3.54 | 3.66 | 3.78<br>3.72 |
| Gal | 4.36<br>d, $J$ = 7.9 Hz | 3.51 | 3.65 | 4.09<br>d, $J$ = 3.4 Hz | 3.52 –<br>3.57 | 3.71 –<br>3.64 |
| GlcNAc | 4.64<br>d, $J$ = 8.6 Hz | 3.74 | 3.68 | 3.69 | 3.48 | 3.90 |
| Gal2 | 4.47<br>d, $J$ = 7.8 Hz | 3.65 | 3.78 | 4.11<br>d, $J$ = 3.4 Hz | 3.52 –<br>3.57 | 3.71 –<br>3.64 |
| Glc2 | 4.71 | 3.45 | 4.26<br>t, $J$ = 8.9 Hz | 3.58 | 3.65 | 3.82<br>3.73 |

| | C1 | C2 | C3 | C4 | C5 | C6 |
|---|---|---|---|---|---|---|
| Glc | 102.1 | 72.8 | 74.5 | 78.3 | 72.2 | 59.8 |
| Gal | 102.7 | 74.8 | 82.2 | 68.3 | 68.1 – 70.0 | 60.3 – 61.7 |
| GlcNAc | 102.7 | 55.1 | 74.9 | 77.9 | 72.1 | 59.9 |
| Gal2 | 102.8 | 75.0 | 82.0 | 68.3 | 68.1 – 70.0 | 60.3 – 61.7 |
| Glc2 | 103.5 | 75.0 | 84.1 | 78.5 | 74.8 | 60.5 |

**5-aminopentyl *β-D*-glucopyranosyl-(1→3)-*β*-D-galactopyranosyl)-(1→4)-(2-acetamido-2-deoxy-*β*-D-glucopyranosyl-(1→3)-*β*-D-galactopyranosyl-(1→4)-*β*-D-glucopyranoside (8).**

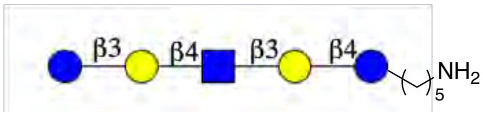


Compound **18b** (10 mg, 0.0037 mmol) was subjected to hydrogenolysis and $NaBH_4$ (2.6 mg, 0.07 mmol) reduction. After reduction of the glucuronate to glucose, the remaining reducing agent was quenched with AcOH and the solids removed. After freeze drying solid, the residue was applied to Biogel P-2 size exclusion column and eluted with 100 mM $NH_4HCO_3$ solution to obtain **7** (2.0 mg 57% overall yield). MALDI-TOF MS *m/z* $C_{37}H_{66}N_2O_{26}$ $(M+Na)^+$ calculated 977.3801, found 977.4213.

$^1$H NMR (600 MHz, $D_2O$) δ: (non carbohydrate): 3.80 (-$OCH_2$ linker), 2.87 (t, $J$ = 6.9 Hz, 2H, -$CH_2$ linker), 1.90 (s, 3H, -NHAc), 1.60 – 1.50 (m, 4H, -$CH_2$ linker), 1.36 – 1.27 (m, 2H, -$CH_2$ linker).

$^{13}$C NMR (151 MHz, $D_2O$): $\delta$ (non carbohydrate): 174.1 ($CH_3\underline{C}O$-), 70.1 (-$O\underline{C}H_2$ linker), 39.4 (-

$CH_2$ linker), 28.2 (-$CH_2$ linker), 26.1 (-$CH_2$ linker), 22.2 (-NHAc), 22.1 (-$CH_2$ linker).

Carbohydrate region:

| | H1 | H2 | H3 | H4 | H5 | H6 |
|---|---|---|---|---|---|---|
| Glc | 4.35 d, $J$ = 8.6 Hz | 3.17 | 3.50 | 3.54 | 3.55 | 3.84 3.71 |
| Gal | 4.30 d, $J$ = 7.8 Hz | 3.46 | 3.59 | 4.02 | na | 3.65 – 3.58 |
| GlcNAc | 4.58 d, $J$ = 8.8 Hz | 3.68 | 3.62 | 3.61 | 3.48 | 3.84 3.72 |
| Gal2 | 4.40 d, $J$ = 7.3 Hz | 3.56 | 3.71 | 4.06 | na | 3.65 – 3.58 |
| Glc2 | 4.55 d, $J$ = 8.0 Hz | 3.30 | 3.39 | 3.60 | 3.60 | 3.82 3.73 |

| | C1 | C2 | C3 | C4 | C5 | C6 |
|---|---|---|---|---|---|---|
| Glc | 102.1 | 72.9 | 74.7 | 78.3 | na | 59.7 |
| Gal | 102.7 | 81.9 | 82.0 | 68.1 | na | 60.0 – 61.7 |
| GlcNAc | 102.7 | 55.4 | 75.4 | 78.0 | na | 59.8 |
| Gal2 | 102.5 | 82.1 | 82.2 | 69.2 | na | 60.0 – 61.7 |
| Glc2 | 103.5 | 75.7 | 75.3 | 78.3 | na | 60.3 |

**5-aminopentyl 3-O-sulfo-*β*-D-galactopyranosyl)-(1→4)-2-acetamido-2-deoxy-*β*-D-glucopyranosyl-(1→3)-*β*-D-galactopyranosyl-(1→4)-*β*-D-glucopyranoside (9).**

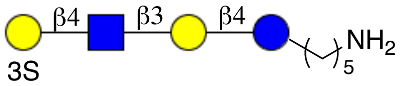


**16b** (5 mg, 0.002 mmol) was sulfated and hydrogenated according to general *O*-sulfation and hydrogenation protocol. After purification by Biogel-P-2 column chromatography compound **9** was obtained (1.5 mg, 83%). ESI TOF-MS *m/z* $C_{31}H_{55}N_2O_{24}S$ $(M - H)^-$ calculated 871.2871, found 871.4774.

$^1$H NMR (600 MHz, $D_2O$) δ: (non carbohydrate): 3.85 (-OC*H*H linker), 3.60 (-OCH*H* linker), 2.92 (t, J = 7.7 Hz, 2H, -$CH_2$ linker), 1.97 (s, 3H, -NHAc), 1.69 – 1.56 (m, 4H, -$CH_2$ linker), 1.43 – 1.35 (m, 2H, -$CH_2$ linker).

$^{13}$C NMR (151 MHz, $D_2O$): $\delta$ (non carbohydrate): 175.2 (-NH$C$O$CH_3$), 70.3 (-O$CH_2$ linker), 39.3

(-*C*$H_2$ linker), 26.6 (-*C*$H_2$ linker), 28.4 (-*C*$H_2$ linker), 22.3 (-NHCO*C*$H_3$), 21.9 (-*C*$H_2$ linker).

| | H1 | H2 | H3 | H4 | H5 | H6 |
|---|---|---|---|---|---|---|
| Glc | 4.42 d, $J$ = 8.5 Hz | 3.22, t, $J$ = 8.5 Hz | 3.54 | 3.55 | 3.63 | na |
| Gal | 4.36 d, $J$ = 7.8 Hz | 3.50 | 3.63 | 4.09 d, $J$ = 3.0 Hz | na | na |
| GlcNAc | 4.64 d, $J$ = 8.2 Hz | 3.72 | 3.68 | 3.70 | na | na |
| Gal | 4.53 d, $J$ = 7.7 Hz | 3.63 | 4.27 dd, $J_1$ = 3.6 Hz, $J_2$ = 9.9 Hz | 4.23 d, $J$ = 3.0 Hz | na | na |

| | C1 | C2 | C3 | C4 | C5 | C6 |
|---|---|---|---|---|---|---|
| Glc | 102.03 | 72.92 | 74.71 | 78.40 | 74.62 | 59.85 |
| Gal | 102.89 | 74.71 | 82.12 | 68.27 | 69.97 | 60.93 |
| GlcNAc | 102.78 | 55.20 | 75.08 | 78.11 | 72.47 | 59.93 |
| Gal2 | 102.42 | 75.08 | 80.10 | 66.74 | 68.98 | 60.93 |

**5-aminopentyl 3-O-sulfo-*β*-D-galactopyranosyl-(1→4)-2-acetamido-2-deoxy-*β*-D-glucopyranosyl-(1→3)-*β*-D-galactopyranosyl-(1→4)-2-acetamido-2-deoxy-*β*-D-glucopyranosyl-(1→3)-*β*-D-galactopyranosyl-(1→4)-*β*-D-glucopyranoside (10).**

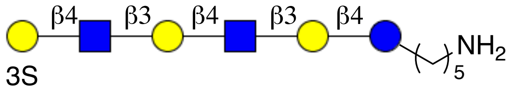


**16c** (5 mg, 0.0016 mmol) was sulfated and hydrogenated according to general *O*-sulfation and hydrogenation protocols. After purification over Biogel-P-2 column compound **10** was obtained (1.2 mg 63%). ESI TOF-MS *m/z* $C_{45}H_{78}N_3O_{34}S$ (M - H)$^-$ calculated 1236.4193, found 1236.6036.

$^1$H NMR (600 MHz, $D_2O$) δ: (non carbohydrate): 3.84 (-OCH*H* linker), 3.60 (-OC*H*H linker), 2.92 (t, J = 7.41 Hz, 2H, -N$CH_2$ linker), 1.96 (s, 6H, 2x-NHAc), 1.66 – 1.56 (m, 4H, -$CH_2$ linker), 1.41 – 1.34 (m, 2H, -$CH_2$ linker).

$^{13}$C NMR (151 MHz, C $D_2O$): *δ* (non carbohydrate): 174.89 (-NH*C*O$CH_3$), 70.06 (-O*C*$H_2$ linker), 39.31 (-N*C*$H_2$ linker), 28.31 (-*C*$H_2$ linker), 26.21 (-*C*$H_2$ linker), 22.11 (-NHCO*C*$H_3$), 22.18 (-*C*$H_2$ linker).

Carbohydrate region:

| | H1 | H2 | H3 | H4 | H5 | H6 |
|---|---|---|---|---|---|---|
| Glc | 4.42<br>d, $J$ = 8.3 Hz | 3.26,<br>t, $J$ = 8.3 Hz | 3.56 | 3.54 | na | na |
| Gal | 4.34<br>d, $J$ = 8.4 Hz | 3.51 | 3.64 | 4.07 | 3.49 | na |
| GlcNAc | 4.64<br>d, $J$ = 8.1 Hz | 3.72 | 3.69 | 3.67 | na | na |
| Gal2 | 4.40<br>d, $J$ = 8.4 Hz | 3.51 | 3.64 | 4.07 | 3.49 | na |
| GlcNAc2 | 4.64<br>d, $J$ = 8.1 Hz | 3.72 | 3.69 | 3.67 | na | na |
| Gal3 | 4.53<br>d, $J$ = 7.9 Hz | 3.62 | 4.25<br>dd, $J_1$ = 3.3 Hz,<br>$J_2$ = 9.9 Hz | 4.24<br>d, $J$ = 3.1 Hz | 3.60 | na |

| | C1 | C2 | C3 | C4 | C5 | C6 |
|---|---|---|---|---|---|---|
| Glc | 101.9 | 72.9 | 74.1 | 78.4 | - | 59.7 |
| Gal | 102.8 | 74.7 | 82.1 | 68.3 | 69.9 | 60.0 |
| GlcNAc | 102.7 | 55.1 | 74.9 | 78.2 | 72.2 | 59.8 |
| Gal2 | 102.6 | 74.7 | 82.1 | 68.3 | 69.9 | 60.9 |
| GlcNAc2 | 102.7 | 55.1 | 74.9 | 78.2 | 72.2 | 59.8 |
| Gal3 | 102.4 | 75.0 | 80.1 | 66.7 | 68.7 | 60.8 |

**5-aminopentyl *β*-D-galactopyranosyl-(1→4)-2-acetamido-2-deoxy-*β*-D-glucopyranosyl-(1→3)-*β*-D-galactopyranosyl-(1→4)-2-acetamido-2-deoxy-*β*-D-glucopyranosyl-(1→3)-*β*-D-galactopyranosyl-(1→4)-*β*-D-glucopyranoside (11).**

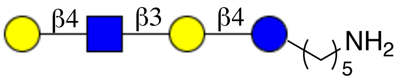


Compound **16b** (8.0 mg, 0.0037 mmol) was subjected to hydrogenolysis using the general protocol to provide **11** (2.5 mg, 83% yield). MALDI-TOF MS *m/z* $C_{31}H_{56}N_2O_{21}$ $(M+Na)^+$ calculated 815.3273, found 815.5653.

$^1$H NMR (600 MHz, $D_2O$) δ: (non carbohydrate): 3.81 (-OCH*H* linker), 3.56 (-OC*H*H linker), 2.87 (t, $J$ = 7.5 Hz, 2H, -$NCH_2$ linker), 1.97 (s, 3H, -NHAc), 1.59 – 1.50 (m, 4H, -$CH_2$ linker), 1.35 – 1.28 (m, 2H, -$CH_2$ linker).

$^{13}$C NMR (151 MHz, $D_2O$): $\delta$ (non carbohydrate): 174.6 (-NH<u>C</u>O$CH_3$), 70.2 (-O<u>C</u>$H_2$ linker), 39.4

(-N$\underline{C}$H$_2$ linker), 28.46 (-$\underline{C}$H$_2$ linker), 26.4 (-$\underline{C}$H$_2$ linker), 22.3 (-NHCO$\underline{CH_3}$), 21.1 (-$\underline{C}$H$_2$ linker).

| | H1 | H2 | H3 | H4 | H5 | H6 |
|---|---|---|---|---|---|---|
| Glc | 4.36<br>d, $J$ = 8.3 Hz | 3.18,<br>t, $J$ = 8.4 Hz | 3.55 | 3.51 | 3.47 | 3.86<br>3.72 |
| Gal | 4.35<br>d, $J$ = 7.8 Hz | 3.47 | 3.60 | 4.03<br>d, $J$ = 3.2 Hz | na | 3.68 – 3.58 |
| GlcNAc | 4.57<br>d, $J$ = 8.9 Hz | 3.68 | 3.60 | 3.60 | 3.44 | 3.86 – 3.81 |
| Gal2 | 4.31<br>d, $J$ = 7.9 Hz | 3.42 | 3.50 | 3.80 | na | 3.68 – 3.58 |

| | C1 | C2 | C3 | C4 | C5 | C6 |
|---|---|---|---|---|---|---|
| Glc | 102.2 | 72.9 | 74.5 | 78.1 | na | 59.7 |
| Gal | 102.5 | na | 82.1 | 68.2 | na | 60.0 – 64.0 |
| GlcNAc | 102.8 | 55.2 | 75.2 | 78.4 | na | 59.9 |
| Gal2 | 102.9 | na | 72.4 | 68.6 | na | 60.0 – 64.0 |

**5-aminopentyl *β*-D-galactopyranosyl-(1→4)-2-acetamido-2-deoxy-*β*-D-glucopyranosyl-(1→3)-*β*-D-galactopyranosyl-(1→4)-2-acetamido-2-deoxy-*β*-D-glucopyranosyl-(1→3)-*β*-D-galactopyranosyl-(1→4)-*β*-D-glucopyranoside (12).**

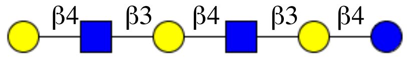


Compound **16c** (10 mg, 0.003 mmol) was subjected to hydrogenolysis using the general protocol to provide compound **12** (3 mg, 86%). MALDI-TOF MS *m/z* $C_{45}H_{79}N_3O_{31}$ $(M+Na)^+$ calculated 1180.4595, found 1180.8492.

$^1$H NMR (600 MHz, D$_2$O) δ: (non carbohydrate): 3.86 (-OCH$\underline{H}$ linker), 3.61 (-OC$\underline{H}$H linker), 2.93 (t, $J$ = 7.7 Hz, 2H, -NCH$_2$ linker), 1.95 (s, 6H, 2x-NHAc), 1.65 – 1.55 (m, 4H, -CH$_2$ linker), 1.42 – 1.33 (m, 2H, -CH$_2$ linker).

$^{13}$C NMR (151 MHz, D$_2$O): $\delta$ (non carbohydrate): 70.1 (-O$\underline{C}$H$_2$ linker), 39.4 (-N$\underline{C}$H$_2$ linker), 28.2 (-$\underline{C}$H$_2$ linker), 26.2 (-$\underline{C}$H$_2$ linker), 22.2 (-NHCO$\underline{CH_3}$), 22.1 (-$\underline{C}$H$_2$ linker).

Carbohydrate region:

| | H1 | H2 | H3 | H4 | H5 | H6 |
|---|---|---|---|---|---|---|
| Glc | 4.40<br>d, $J$ = 7.9 Hz | 3.21,<br>t, $J$ = 8.9 Hz | 3.56 | 3.54 | 3.46 | 3.76<br>3.88 |
| Gal | 4.41<br>d, $J$ = 7.5 Hz | 3.46 | 3.63 | 4.07 | na | 3.72 – 3.63 |
| GlcNAc | 4.62<br>d, $J$ = 8.4 Hz | 3.72 | 3.65 | 3.64 | 3.50 | 3.90 – 3.85 |
| Gal2 | 4.40<br>d, $J$ = 7.8 Hz | 3.51 | 3.63 | 4.07 | na | 3.72 – 3.63 |
| GlcNAc2 | 4.62<br>d, $J$ = 8.4 Hz | 3.72 | 3.65 | 3.64 | 3.50 | 3.90 – 3.85 |
| Gal3 | 4.35<br>d, $J$ = 7.9 Hz | 3.46 | 3.64 | 3.84 | na | 3.72 – 3.63 |

| | C1 | C2 | C3 | C4 | C5 | C6 |
|---|---|---|---|---|---|---|
| Glc | 101.2 | 72.9 | 74.7 | 78.4 | na | 59.9 |
| Gal | 102.6 | na | 82.1 | 68.3 | na | 59.8 – 61.8 |
| GlcNAc | 102.7 | 55.6 | 75.1 | 78.3 | na | 60.0 |
| Gal2 | 102.6 | na | 82.1 | 68.3 | na | 59.8 – 61.8 |
| GlcNAc2 | 102.7 | 55.6 | 75.1 | 78.3 | na | 60.0 |
| Gal3 | 103.0 | na | 72.1 | 68.5 | na | 59.8 – 61.8 |

**General procedure for the sialylation of Lacto-N-neotetraose and Lacto-N-neohexaose.** Compounds **11** and **12** and CMP-Neu5Ac (1.5 eq for each reaction) were dissolved in TrisHCl buffer (10 mM, pH 8.0) and $MgCl_2$ (20 mM), CIAP (10 U $\mu L^{-1}$) were added followed by the addition of PmST1 (1% w/w per acceptor). The reaction mixtures were incubated for 2-4 h and the progress of the reactions was monitored by TLC (EtOAc:MeOH:$H_2O$:AcOH = 4:2:1:0.1) and ESI-MS (negative mode). Upon completion of the reactions, they were stopped by the addition of MeOH and further incubated at 0 °C for 30 min and then concentrated under reduced pressure at 25 °C and applied on size-exclusion Biogel P-2 column and eluted with 10 mM $NH_4HCO_3$. Fractions containing product were detected by TLC staining and confirmed by LC-MS, after which were combined and lyophilized. To the precipitate after freeze-drying deionized water was added and freeze-dried to remove the residual $NH_4HCO_3$.

**5-aminopentyl α-5-acetamido-3,5-dideoxy-D-glycero-β-D-galacto-non-2-ulopyranosylate-(2→3)-*β*-D-galactopyranosyl-(1→4)-2-acetamido-2-deoxy-*β*-D-glucopyranosyl-(1→3)-*β*-D-galactopyranosyl-(1→4)-*β*-D-glucopyranoside (13).**

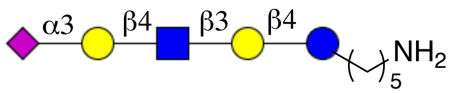


Compound **11** (1 mg, 0.0013 mmol) was sialylated to obtain 0.9 mg of **13** (69%). ESI-TOF MS *m/z* $C_{42}H_{73}N_3O_{29}$ $(M - H)^-$ calculated 1082.4257, found 1082.6159.

$^1$H NMR (600 MHz, $D_2O$) δ: (non carbohydrate): 3.80 (-OCH*H* linker), 3.55 (-OC*H*H linker), 2.86 (t, *J* = 7.8 Hz, 2H, -$NCH_2$ linker), 1.89 (s, 6H, -NHAc GlcNAc, -NHAc Neu5Ac), 1.59 – 1.50 (m, 4H, -$CH_2$ linker), 1.35 – 1.29 (m, 2H, -$CH_2$ linker).

$^{13}$C NMR (151 MHz, $D_2O$): *δ* (non carbohydrate): 174.5 (-*C*OOH), 174.8 (-NH*C*OCH$_3$), 70.1 (-O*C*H$_2$ linker), 39.5 (-N*C*H$_2$ linker), 28.4 (-*C*H$_2$ linker), 26.6 (-*C*H$_2$ linker), 22.1 (-*C*H$_2$ linker), 21.9 (-NHCO*CH*$_3$).

Carbohydrate region:

| | H1 | H2 | H3 | H4 | H5 | H6 |
|---|---|---|---|---|---|---|
| Glc | 4.34<br>d, *J* = 8.3 Hz | 3.17,<br>t, *J* = 9.0 Hz | 3.49 | 3.51 | na | na |
| Gal | 4.29<br>d, *J* = 7.4 Hz | 3.45 | 3.59 | 4.02<br>d, *J* = 2.9 Hz | na | na |
| GlcNAc | 4.56<br>d, *J* = 8.7 Hz | 3.68 | 3.58 | 3.61 | 3.45 | na |
| Gal2 | 4.42<br>d, *J* = 8.0 Hz | 3.44 | 3.98<br>dd, $J_1$ = 3.0 Hz,<br>$J_2$ = 10.2 Hz | 3.83 | na | na |

| | H3 | H4 | H5 | H6 | H7 | H8 | H9 |
|---|---|---|---|---|---|---|---|
| NeuNAc | 2.62 (eq)<br>dd, *J*= 4.7, 12.5 Hz<br>1.63 (ax)<br>t, *J* = 12.6 Hz | 3.56 | 3.70 | 3.40 | na | n/a | n/a |

| | C1 | C2 | C3 | C4 | C5 | C6 |
|---|---|---|---|---|---|---|
| Glc | 102.0 | 72.8 | 74.6 | na | na | na |
| Gal | 102.1 | 71.8 | 71.6 | 68.5 | na | na |
| GlcNAc | 102.8 | 55.1 | 75.3 | na | na | na |

| | | | | | | |
|---|---|---|---|---|---|---|
| Gal2 | 102.9 | 75.6 | 75.5 | 67.6 | na | na |

| | C1 | C2 | C3 | C4 | C5 | C6 | C7 | C8 | C9 |
|---|---|---|---|---|---|---|---|---|---|
| Neu5Ac | 174.0 | 99.7 | 39.6 | 68.6 | 51.6 | 72.4 | na | Na | na |

**5-aminopentyl α-5-acetamido-3,5-dideoxy-D-glycero-β-D-galacto-non-2-ulopyranosylate)-(2→3)-*β*-D-galactopyranosyl-(1→4)-2-acetamido-2-deoxy-*β*-D-glucopyranosyl-(1→3)-*β*-D-galactopyranosyl-(1→4)-2-acetamido-2-deoxy-*β*-D-glucopyranosyl-(1→3)-*β*-D-galactopyranosyl-(1→4)-*β*-D-glucopyranoside (14).**

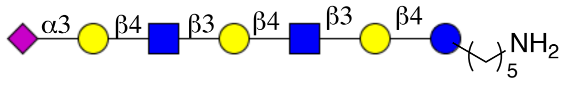


Compound **12** (1.0 mg 0.0008 mmol) was sialylated to obtain of **14** (1.1 mg 88%). ESI-TOF MS *m/z* $C_{56}H_{96}N_4O_{39}$ (M - H)$^-$ calculated 1447.5579, found 1447.7662.

$^1$H NMR (600 MHz, $D_2O$) δ: (non carbohydrate): 3.80 (-OCH*H* linker), 3.55 (-OC*H*H linker), 2.88 (t, *J* = 7.9 Hz, 2H, -$NCH_2$ linker), 1.89 (s, 9H, -NHAc GlcNAc1-2, -NHAc Neu5Ac), 1.59 – 1.50 (m, 4H, -$CH_2$ linker), 1.35 – 1.28 (m, 2H, -$CH_2$ linker).

$^{13}$C NMR (151 MHz, $D_2O$): *δ* (non carbohydrate): 174.5 (-*C*OOH), 174.8 (-NH*C*OCH$_3$), 70.1 (-O*C*H$_2$ linker), 39.3 (-N*C*H$_2$ linker), 28.1 (-*C*H$_2$ linker), 26.3 (-*C*H$_2$ linker), 22.1 (-*C*H$_2$ linker), 21.9 (-NHCO*C*H$_3$).

Carbohydrate region:

| | H1 | H2 | H3 | H4 | H5 | H6 |
|---|---|---|---|---|---|---|
| Glc | 4.36<br>d, *J* = 8.6 Hz | 3.16,<br>t, *J* = 9.5 Hz | 3.50 | 3.54 | na | 3.82<br>3.73 |
| Gal | 4.30<br>d, *J* = 7.7 Hz | 3.46 | 3.61 | 4.02 | na | na |
| GlcNAc | 4.56<br>d, *J* = 8.4 Hz | 3.67 | 3.59 | na | na | 3.83 |
| Gal2 | 4.33<br>d, *J* = 7.7 Hz | 3.45 | 3.59 | 4.02 | na | na |
| GlcNAc2 | 4.56<br>d, *J* = 8.4 Hz | 3.67 | 3.59 | na | na | 3.83 |

| | | | | | | |
|---|---|---|---|---|---|---|
| Gal3 | 4.42, d, $J$ = 8.0 Hz, | 3.44 | 3.98 dd, $J_1$ = 3.3 Hz, $J_2$ = 9.7 Hz | 3.82 | na | na |

| | H3 | H4 | H5 | H6 | H7 | H8 | H9 |
|---|---|---|---|---|---|---|---|
| Neu5Ac | 2.62 (eq) dd, $J$ = 4.2, 12.7 Hz 1.66 (ax) $J$ = 12.2 Hz | 3.55 | 3.71 | na | na | na | na |

| | C1 | C2 | C3 | C4 | C5 | C6 |
|---|---|---|---|---|---|---|
| Glc | 101.7 | 72.7 | na | na | na | 59.7 |
| Gal | 102.9 | na | na | 68.1 | na | 61.0 |
| GlcNAc | 102.7 | 55.3 | na | na | na | 59.8 |
| Gal2 | 102.4 | na | na | 68.1 | na | 61.0 |
| GlcNAc2 | 102.7 | 55.3 | na | na | na | 59.8 |
| Gal3 | 102.5 | na | 75.5 | 67.3 | na | 61.0 |

| | C1 | C2 | C3 | C4 | C5 | C6 | C7 | C8 | C9 |
|---|---|---|---|---|---|---|---|---|---|
| NeuNAc | 176.3 | 100.5 | 39.6 | 68.2 | 51.9 | na | na | na | na |

#### 6. Synthesis of building blocks

##### Chemoenzymatic synthesis of UDP-GlcNHTFA nucleotide donor

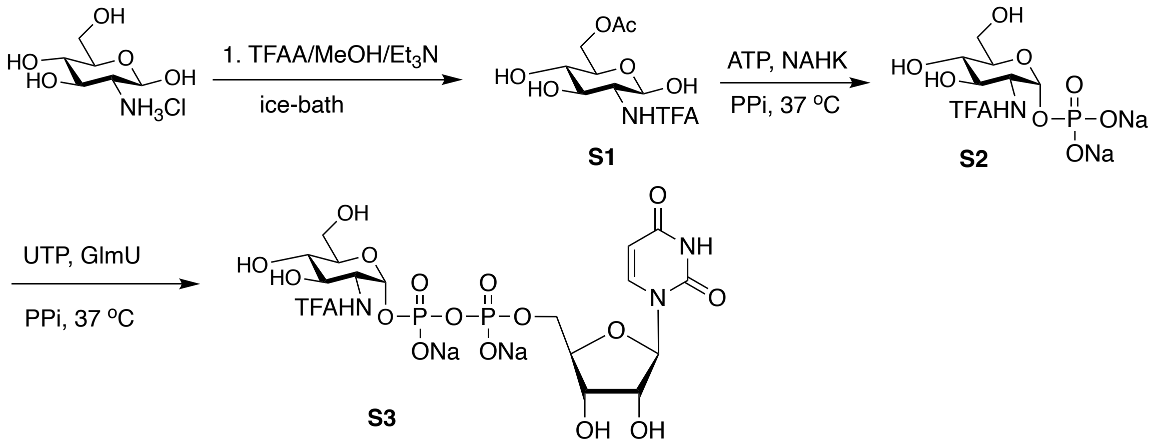


**Scheme S1.** Chemoenzymatic synthesis of UDP-GlcNHTFA.

2-Deoxy-2-amino-glucopyranosyl hydrochloride (4.3 g, 20 mmol) was suspended in methanol to which triethylamine (10 mL) and trifluoroacetic anhydride (10 mL) were added at 0 °C. The reaction was stirred at that temperature for 30 min at 0 °C and then at room temperature for a further 30 min. Upon completion, the reaction mixture was concentrated and residue was coevaporated with toluene and purified over silicagel column chromatography using EtOAc:MeOH = 4:1 as an eluent. Product containing fractions were pooled and concentrated by rotary evaporator and the obtained solid residue was dissolved in deionized water and lyophilized resulting in 6 g white fluffy N-trifluoroacetyl glucosamine were obtained.

S1 (1.0 g, 3.6 mmol) and ATP disodium salt (3.0 g, 5.4 mmol) were mixed Tris buffer (pH = 7.8, 100 mM) containing $MgCl_2$ (20 mM) to which Bifidobacterum longum N-Acetylhexosamine 1-kinase enzyme was added (10 mg, 1% w/w) along with inorganic pyrophosphotase (0.1 U). The reaction mixture was incubated at an incubator at 37 $^0$C with occasional shaking. The progress of anomeric phosphate formation was monitored by TLC until disappearance of the starting material. The reaction mixture was spin filtered using 10 MWCO spin filters and lyophilized and then used in the next step.

To the mixture of S2 and UTP (2.4 g, 5.3 mmol) in Tris buffer (pH = 8.0) N-acetyl hexosamine-1-phosphate uridyl transferase (PmGlmU, 5 – 10 mg) and inorganic pyrophosphatase (PpA, 0.1 U) were added and the reaction mixture was incubated in a shaker at 37 $^0$C for 2 h and the for another 18 h at room temperature. The progress of sugar-nucleotide formation was monitored by ESI-TOF. An additional portion of GlmU and PpA were added and the mixture was further incubated for 3 h at 37 $^0$C. Upon consumption of the starting material, the reaction mixture was treated with 1M $BaCl_2$ solution, vortexed and centrifuged to remove barium salts of monophosphate and diphosphate nucleotides and insoluble proteins. Supernatant was decanted and freeze dried. Crude UDP-GlcNHTFA was dissolved in minimum amount of water and applied on Biogel P-2 size exclusion chromatography for desalting. Fractions containing nucleotide sugar donors were combined and freeze-dried to yield UDP-GlcNHTFA (1.2 g, 48%) as a fluffy material. ESI TOF-MS *m/z* $C_{17}H_{24}F_3N_3O_{17}P_2$ $(M-H)^-$ calcd 660.0460 found 659.8330. $^1$H NMR characteristics are in accordance with the previously published literature (4).

**Synthesis of glucuronate phosphate donor**

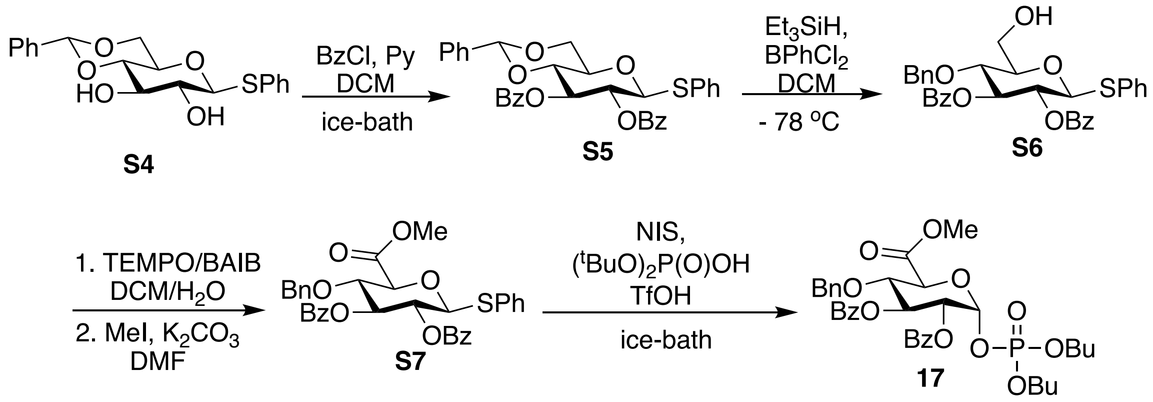


**Scheme S2.** Synthesis glucuronate phosphate donor (**17**).

**4,6-O-benzylidene-1-thio-*β*-D-glucopyranose (S4).**

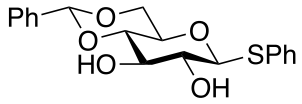

Peracetylated glucopyranose (19.5 g, 0.05 mol) was dissolved in dry DCM (100 mL) and the resulting solution cooled to 0 ℃. To this solution, PhSH (9 mL, 0.075 mol) and $BF_3Et_2O$ (10 mL, 0,075 mL) were added and the mixture left stirring for 30 min at 0 ℃ and then 2 h at room temperature. The progress of the reaction was monitored by TLC until complete consumption of the starting material, after which it was quenched by the addition of sat. $NaHCO_3$ solution. The organic phase was separated, dried ($Na_2SO_4$) and concentrated under reduced pressure. The residue was subjected to silica gel column chromatography using PE:EtOAc (10:1 ⟶ 1:1) as eluent furnished the required thioglycoside. The product was dissolved in MeOH and cat. amount of NaOMe solution was added until the pH of the solution reached 9. The reaction mixture was stirred for 2 h until TLC indicated complete deacetylation of the material. The reaction was quenched with Amberlyte $H^+$ and filtered. The filtrate was concentrated under reduced pressure and the residue twice coevaporated from toluene. The residue was further dried in *high-vacuo*. The solid residue was resuspended in acetonitrile (100 mL) and $PhCH(OMe)_2$ (10 mL) and cat. amount of p-toluenesulfonic acid (0.5 g) were added. The reaction mixture was stirred for 2 h at room temperature. Upon completion of the reaction the product started to precipitate. The reaction was quenched by triethyl amine and the precipitate was filtered off and washed with ice-cold diethyl ether (2x). The white precipitates were collected and dried in high vacuo to yield the titled product (15 g, 83% over 3 steps. MALDI

TOF-MS *m/z* $C_{19}H_{20}SO_5$ $(M+Na)^+$ calcd 383.0929 found 383.0500.

$^1$H NMR (600 MHz, $CDCl_3$) δ 7.55–7.21 (m, 10H, Ar-H), 7.91 – 7.88 (m, 2H, Ar-H), 5.51 (s, 1H, *CH*Ph), 4.63 (d, *J* = 9.81 Hz, 1H, H-1), 4.36 (dd, $J_1$ = 4.53 Hz, $J_2$ = 10.41 Hz, 1H, H-6a), 3.83 (t, *J* = 8.45 Hz, 1-H, H-4), 3.76 (m, 1H, H-6b), 3.53 – 3.48 (m, 2H, H-3, H-5), 3.46 (t, *J* = 9.36 Hz, 1H, H-2).

$^{13}$C NMR (151 MHz, $CDCl_3$): *δ* 68.5 (C6), 70.4 (C5), 72.9 (C2), 74.8 (C4), 80.3 (C3), 88.7 (C1), 102.0 (*CH*Ph), 126.0, 126.4, 128.3, 128.9, 129.0, 129.3, 129.69, 129.80, 132.9 ($C_{Ar}$).

**2,3-di-O-benzoyl-4,6-O-benzylidene-1-thio-*β*-D-glucopyranose (S5).**

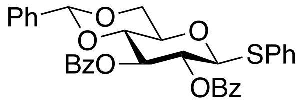


Compound S4 (14 g, 38.67 mmol) of was dissolved in DCM (100 mL) to which pyridine (20 mL) was added and the resulting solution was cooled to 0 ℃, followed by the addition of BzCl (20 mL, ~150 mmol). The reaction mixture was stirred at that temperature for 30 min when TLC indicated the formation of the product. The reaction mixture was diluted with DCM and washed with aqueous HCl (1M) and sat. $NaHCO_3$ solution, respectively. The organic layer was separated, dried (anhydrous $Na_2SO_4$) and concentrated *in vacuo*. The residue was purified by silica gel column chromatography to afford the titled compound (20 g, 91%). MALDI-TOF MS *m/z* $C_{33}H_{28}SO_7$ $(M+Na)^+$ calcd 591.1453 found 591.2877.

$^1$H NMR (600 MHz, $CDCl_3$) δ 7.98–7.91 (m, 4H, Ar-H), 7.55 – 7.28 (m, 16H, Ar-H), 5.79 (t, J = 9.67 Hz, 1H, H-3), 5.54 (s, 1H, *CH*Ph), 5.47 (t, J = 9.27 Hz, 1H, H-2), 5.03 (d, J = 9.81 Hz, 1H, H-1), 4.46 (dd, $J_1$ = 5.0 Hz, $J_2$ = 10.53 Hz, 1H, H-6a), 3.92 – 3.85 (m, 2H, H-4, H-6b), 3.76 (ddd, J = 4.87 Hz, 1H, H-5).

$^{13}$C NMR (151 MHz, $CDCl_3$): *δ* 68.7 (C6), 71.0 (C2), 71.1 (C5) 73.1 (C3), 79.0 (C4), 87.2 (C1), 101.3 (*CH*Ph), 126.0, 128.2, 128.3, 128.4, 129.0, 129.7, 129.9, 133.1, 133.4 ($C_{Ar}$), 165.2 (-$C_{Bz}$), 165.6 (-$C_{Bz}$).

**2,3-di-O-benzoyl-4-O-benzyl-1-thio-*β*-D-glucopyranose (S6).**

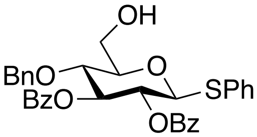


To a solution of S5 (10 g, 17.54 mmol) in dry DCM, 4Å MS was added and the resulting suspension was stirred for 2 h at room temperature. Next, the mixture was cooled to -78 ℃ and triethylsilane (5.2 mL, 35 mmol) was added followed by the addition of $PhBCl_2$ (3.5 mL, 26 mmol). After stirring for 30 min at -78℃, TLC indicated consumption of the starting material. The reaction was quenched by the addition of triethylamine and filtered over a pad of celite. The filtrate was washed with sat. $NaHCO_3$, dried ($Na_2SO_4$) and concentrated under reduced pressure. The crude product was chromatographed using PE:EtOAc = 4:1 as an eluent to furnish the title compound (9.0 g, 90%). MALDI TOF-MS *m/z* $C_{33}H_{30}SO_7$ $(M+Na)^+$ calcd 593.1610 found 593.3301.

$^1H$ NMR (600 MHz, $CDCl_3$) δ 7.95–7.87 (m, 4H, Ar-H), 7.54 – 7.10 (m, 16H, Ar-H), 5.74 (t, J = 9.56 Hz, 1H, H-3), 5.35 (t, J = 9.89 Hz, 1H, H-2), 4.97 (d, J = 10.15 Hz, 1H, H-1), 4.57 (s, 2H, $CH_2$Ph) 3.98 (m, 1H, H-6a), 3.93 (t, J = 9.6 Hz, 1H, H-4), 3.82 (m, 1H, H-6b), 3.63 (m, 1H, H-5).

$^{13}C$ NMR (151 MHz, $CDCl_3$): *δ* 61.8 (C6), 71.0 (C2), 75.0 (*C*$H_2$Ph), 75.4 (C4), 76.2 (C3), 79.5 (C5) 86.25 (C1), 128.0, 128.2, 128.3, 128.4, 129.0, 129.7, 129.9, 132.6, 133.2, 133.6 (C Ar), 165.3 (-*C* Bz), 165.7 (-*C* Bz).

**Methyl 2,3-di-O-benzoyl-4-O-benzyl-1-thio-*β*-D-glucopyranosyluronate (S7).**

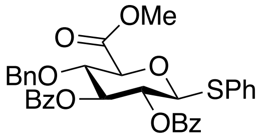


Compound **4** (6.0 g, 10 mmol) was dissolved in DCM (80 mL) and $H_2O$ (40 mL) was added to the mixture, after which it was cooled in an ice-bath. TEMPO (300 mg, 2 mmol) and bis(acetoxy)iodobenzene (10 g, 30 mmol) were added and the reaction mixture was stirred for 2 at room temperature when TLC (EtOAc:PE = 1:2, R*f* = 0) indicated the formation of the oxidation product. The reaction was quenched with 10% solution of $Na_2S_2O_3$ (40 mL) and sat. $NaHCO_3$ solution. The aqueous phase was separated and washed with EtOAc (2x), the combined organic fractions were dried ($Na_2SO_4$), filtered and concentrated and used in the following reaction without purification. Crude glucuronic acid was dissolved in DMF, $K_2CO_3$ (7.0 g, 50 mmol) and $CH_3I$ (14.0 g, 100 mmol) were added at 0 ℃. After 1 h of stirring at 0°C, TLC showed the complete conversion of the acid into the methyl ester.

The reaction was quenched by the addition of water and then concentrated under reduced pressure. The crude product was dissolved in EtOAc and washed with $H_2O$. The organic phase was separated and dried (anhydrous $Na_2SO_4$) and concentrated under reduced pressure. The residue was applied to silica gel column chromatography and eluted using PE:EtOAc = 100% → 75% affording methyl ester product (5 g, 83%). R*f* = 0.5 EtOAc:PE=1:4. MALDI-TOF MS *m/z* $C_{34}H_{30}SO_8$ $(M+Na)^+$ calcd 621.1559 found 621.2950.

$^1$H NMR (600 MHz, Chloroform-d) δ 7.95–7.87 (m, 4H, Ar-H), 7.54 – 7.03 (m, 16H, Ar-H), 5.72 (t, J = 8.8 Hz, 1H, H-3), 5.37 (t, J = 10.26 Hz, 1H, H-2), 4.97 (d, J = 10.2 Hz, 1H, H-1), 4.56 (d, J = 11.36 Hz, 1H, $CH_{2a}$Ph), 4.51 (d, J = 11.00 Hz, 1H, $CH_{2b}$Ph) 4.18 – 4.10 (m, 2H, H-4, H-5), 3.81 (s, 3H, -COO$CH_3$).

$^{13}$C NMR (151 MHz, $CDCl_3$): *δ* 52.46 ($CH_3$), 70.3 (C2), 74.6 (*C*$H_2$Ph), 75.3 (C3), 76.9 (C4), 77.8 (C5) 86.9 (C1), 127.9, 128.0, 128.3, 128.3, 128.4, 129.0, 129.2, 129.7, 129.9, 131.9, 132.9, 133.3, 138.9 (C Ar), 165.1 (-*C* Bz), 165.5 (-*C* Bz), 168.1 (-*C*OOMe).

**Methyl 2,3-di-O-benzoyl-4-O-benzyl-1-di-O-butylphosphatidyl-*β*-D-glucopyranosyluronate (17).**

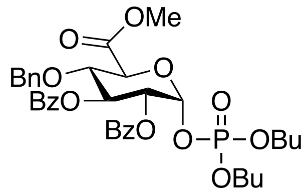


Compound **5** (1 g, 1.67 mmol) and $HOPO(OBu)_2$ (0.87 g, 4.2 mmol) were dissolved in dichloromethane (50 mL) and 4Å MS were added and the resulting suspension was stirred for 30 min at room temperature. The mixture was cooled to 0 ℃ and NIS (0.75 g, 3.33 mmol) was added followed by the addition of cat. amount of TfOH (25 mg, 0.167 mmol). The reaction mixture was stirred for 30 min at 0 ℃, after which TLC showed the full conversion of the starting material into phosphate donor. The mixture was diluted with DCM and 20 mL of 1:1 (v/v) $Na_2S_2O_3$ and $NaHCO_3$ solutions were used to quench the reaction. The organic phase was separated and dried ($Na_2SO_4$) and concentrated under reduced pressure. Flash silica gel column chromatography using PE:EtOAc = 100% →75% as eluent furnished the title compound in quantitative yield. R*f* = 0.5, PE:EtOAc = 1:1. MALDI-TOF MS *m/z* $C_{36}H_{43}PO_{12}$ $(M+Na)^+$ calcd 721.6915 found 721.4044.

$^1$H NMR (600 MHz, Chloroform-d) δ 7.95–7.90 (m, 4H, Ar-H), 7.54 – 7.48 (m, 2H, Ar-H), 7.40 – 7.33 (m, 4H, Ar-H), 7.18 – 7.14 (m, 3H, Ar-H), 7.11 – 7.05 (m, 2H, Ar-H), 5.72 (t, *J* = 9.7 Hz, 1H, H-3), 5.55 (t, *J* = 7.3 Hz, 1H, H-1), 5.50 (t, *J* = 8.93 Hz, 1H, H-2), 4.58 (d, J = 11.2 Hz, 1H, $CH_{2a}$Ph), 4.52 (d, *J* = 11.0 Hz, 1H, -OCH*H*Ph) 4.27 (d, *J* = 9.6 Hz, 1H, H-5), 4.18 (t, *J* = 9.3 Hz, 1H, H-4), 4.09 – 3.98 (m, 3H, $-OCH_2Ph$), 3.81 – 3.77 (m, 1H, $CH_2$ (OBu)), 3.76 (s, 3H, -COO$CH_3$). 3.73 – 3.67 (m, 1H, $CH_2$ OBu), 1.68 – 1.58 (m, 3H, $CH_2$ OBu), 1.44 – 1.23 (m, 5H, $CH_2$ OBu),1.07 – 1.00 (m, 2H, $CH_2$ OBu), 0.95 – 0.91 (m, 1H, $CH_2$ OBu), 0.90 (t, *J* = 7.4 Hz, C$H_3$OBu)), 0.69 (t, *J* = 7.3 Hz, $CH_2$ OBu).

$^{13}$C NMR (151 MHz, $CDCl_3$): $\delta$ 13.3, 13.4 ($\underline{C}H_3$ OBu), 18.2, 18.5 ($\underline{C}H_2$ OBu), 31.7, 32.0 ($\underline{C}H_2$ OBu). 52.7 ($CH_3$), 67.9, 68.2 ($\underline{C}H_2$ OBu),. 71.3 (C2), 73.5 (C3), 74.7 (C5), 74.8 (*C*$H_2$Ph), 77.2 (C4), 96.46 (C1), 128.0, 128.1, 128.3, 128.4, 128.8, 129.0, 129.8, 129.9, 131.9, 133.4, 133.5, 136.7 (C Ar), 165.0 (-*C* Bz), 165.2 (-*C* Bz), 167.9 (-*C*OOMe).

#### 7. Microarray

##### Serology serum samples

Ten patients with anti-MAG neuropathy and three healthy controls were used in this study. Patient samples were examined for anti-MAG IgM antibodies as part of standard diagnostic care by ELISA (Bühlmann Laboratories) according to the manufacturer's recommendations. Samples were collected after approval of the Institutional Review Board of the Erasmus MC (MEC-2016-606) and anonymized for further studies. Controls consisted of healthy Dutch blood bank donors. Sera were aliquoted and stored at -20 °C.

**Table S1.** Characteristics of MAG patients and anti-MAG ELISA results.

| | | | | Anti-MAG IgM ELISA | |
|---|---|---|---|---|---|
| SS# | Code | Gender | Age (years) | BTU* | Outcome |
| SS01 | MAG-01 | F | 75 | 11,524 | Positive |
| SS02 | MAG-02 | F | 77 | 29,191 | Positive |
| SS03 | MAG-03 | M | 57 | 6,420 | Weak positive |
| SS04 | MAG-04 | M | 72 | 5,310 | Weak positive |
| SS05 | MAG-05 | M | 54 | 50,851 | Positive |
| SS06 | MAG-06 | F | 73 | 40,085 | Positive |
| SS07 | MAG-07 | M | 71 | 5,587 | Weak positive |
| SS08 | MAG-08 | M | 72 | 63,706 | Positive |
| SS09 | MAG-09 | M | 69 | 59,507 | Positive |
| SS10 | MAG-10 | M | 66 | 28,869 | Positive |

*Anti-MAG IgM antibody titers (BTU: Bühlmann titer units determined by ELISA from Bühlmann Laboratories).

##### Glycan array printing

The synthetic glycans (100 μM in sodium phosphate (250 mM), pH 8.5 buffer) were printed as replicates of 6 on activated glass slides (Nexterion Slide H, Schott Inc) by piezoelectric non-contact printing (sciFLEXARRAYER S3, Scienion Inc) with a drop volume of ~400 pL and 1 drop per spot at 50% relative humidity. Each slide contained 24 subarrays (3x8). The slides were incubated overnight in a saturated NaCl chamber (providing a 75% relative humidity environment), after which the remaining activated esters were quenched with ethanolamine (50 mM) in TRIS (100 mM), pH 9.0. Slides were rinsed with DI water, dried by centrifugation, and stored in a desiccator at RT.

**Microarray binding assays and serum sample screening**

Sub-arrays were incubated with the protein of interest at the indicated concentrations in TSM binding buffer (20 mM Tris Cl, pH 7.4, 150 mM NaCl, 2 mM $CaCl_2$, 2 mM $MgCl_2$, 0.05% Tween, 1% BSA) for 1 h followed by washing. Wash steps involved 4 successive washes of the whole slides with TSM wash buffer (20 mM Tris Cl, pH 7.4, 150 mM NaCl, 2 mM $CaCl_2$, 2 mM $MgCl_2$, 0.05% Tween-20) - TSM buffer (20 mM Tris Cl, pH 7.4, 150 mM NaCl, 2 mM $CaCl_2$, 2 mM $MgCl_2$) - 2x deionized $H_2O$ with 5 min soak times. All incubation and wash steps were performed at RT.

Step-wise incubations and washes were performed as described above with biotinylated mouse monoclonal IgM anti-human CD57 (ProSpec ant-288) at 100 and 150 µg/mL and Streptavidin-AlexaFluor635 (5 µg/mL; ThermoFisher Scientific, S32364); mouse monoclonal IgG1 anti-human L2/HNK-1 carbohydrate (Abcam ab174437) at 3 and 10 µg/mL and a mixture of biotinylated goat anti-mouse IgG (5 µg/mL; Sigma-Aldrich B7264) and Streptavidin-AlexaFluor635 (5 µg/mL); recombinant human galectin-3 (R&D Systems 1154-GA) at 10, 30 µg/mL and a mixture of biotinylated goat anti-human galectin-3 (5 µg/mL; R&D Systems BAF1154) and Streptavidin-AlexaFluor 635 (5 µg/mL).

Human serum samples were provided randomly numbered and were blindly assayed on the microarray similarly as described above. Sub-arrays were incubated with human serum samples (1:250) in TSM binding buffer for 1 h. After washing steps, the sub-arrays were incubated with AlexaFluor647-labeled goat anti-human IgM (5 µg/mL; Jackson Immuno Research, 109-165-129) for 1 h.

Washed arrays were dried by centrifugation and immediately scanned for fluorescence on a GenePix 4000 B microarray scanner (Molecular Devices). The detection gain was adjusted to avoid saturation of the signal, whereby the same settings were used for each experiment to allow comparison between patient and control samples. The data were processed with GenePix Pro 7 software and further analyzed using our home written Microsoft Excel macro. After removal of the lowest and highest value of the six replicates, the mean fluorescent intensities (corrected for mean background) and standard deviations (SD) were calculated (n=4). Data were fitted using Prism software Version 8.3.0 (GraphPad Software, Inc). Bar graphs represent the mean ± SD for each compound. The highest possible protein concentration / serum dilution was employed at which good responsiveness was observed to achieve an appropriate dynamic range.

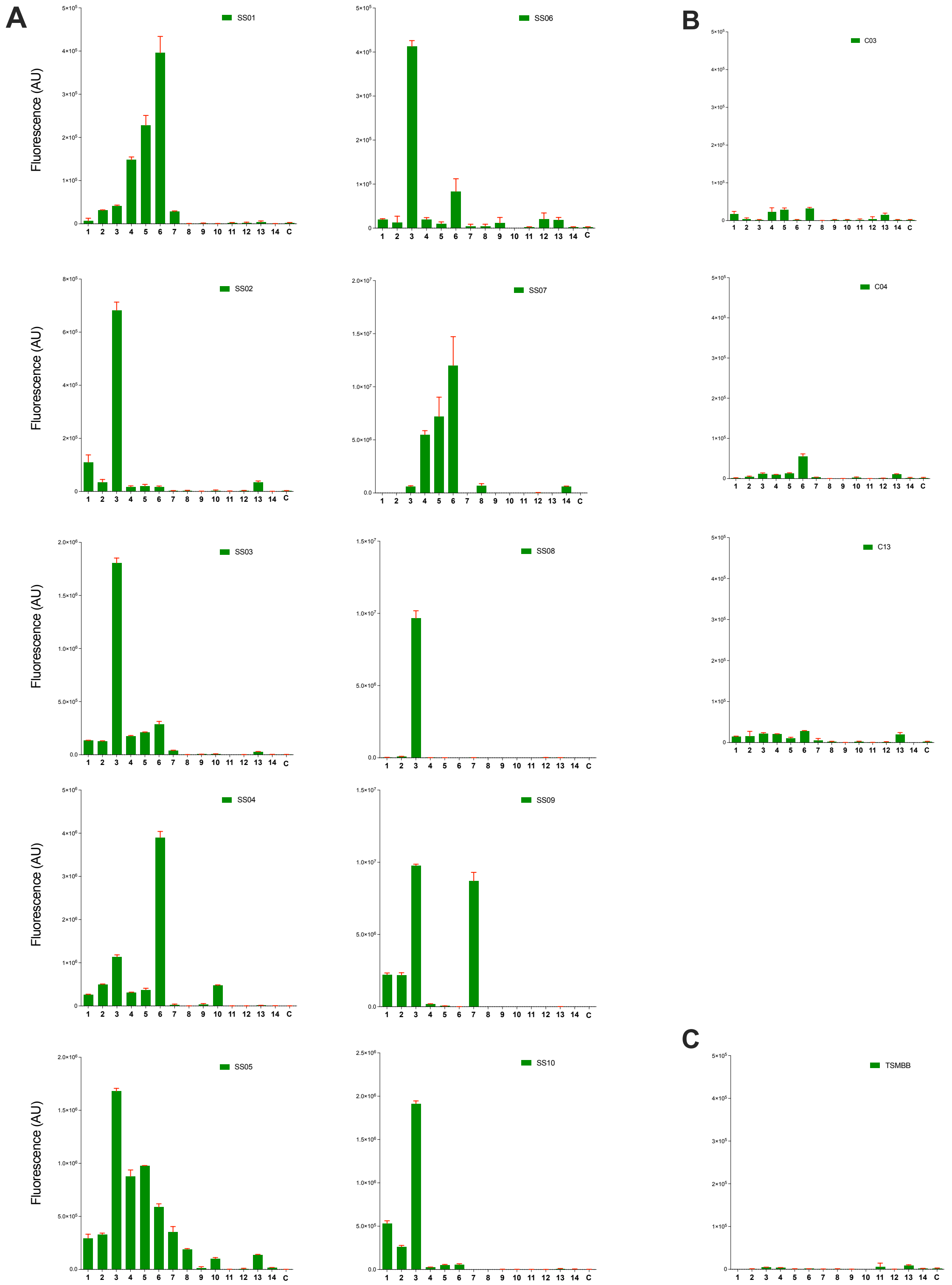


**Figure S8.** Microarray IgM results of the synthetic library with serum samples (1:250 diluted) of MAG patients (**A**), healthy donors (**B**) and TSM binding buffer (**C**). Bars represent the mean ± SD (n=4). C indicates buffer control. Assays are performed in the same session. MAG patient results are scaled to maximize the readout, while controls are shown with the same scale as the lowest patient. The IgM results of the MAG patients are also shown in Fig. 5 using a different presentation.

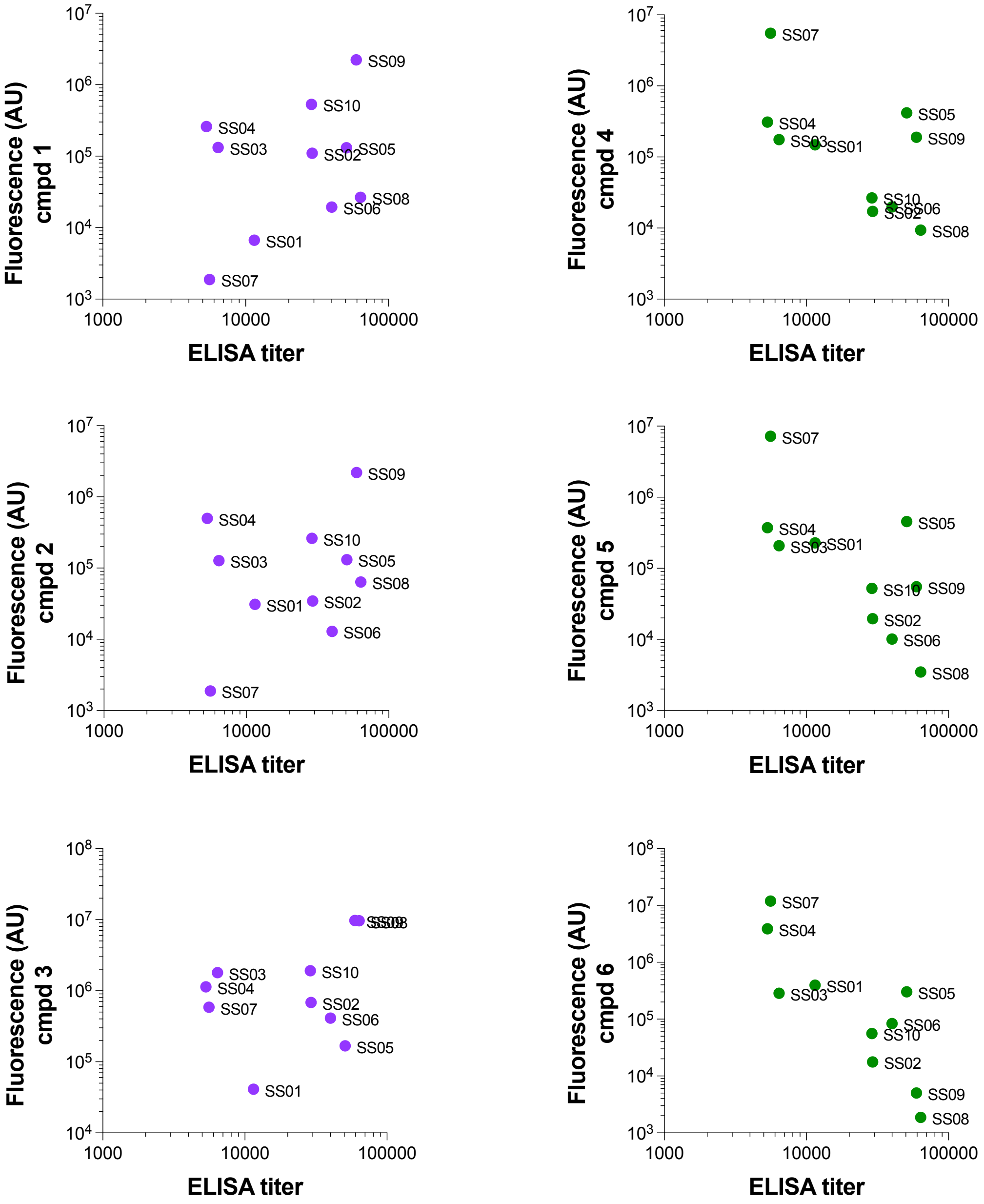


**Figure S9.** Correlation of MAG ELISA data *vs* microarray data for compounds 1-6. No statistically significant correlation was found (using Spearman correlation).


#### 9. Copies of NMR and MS spectra

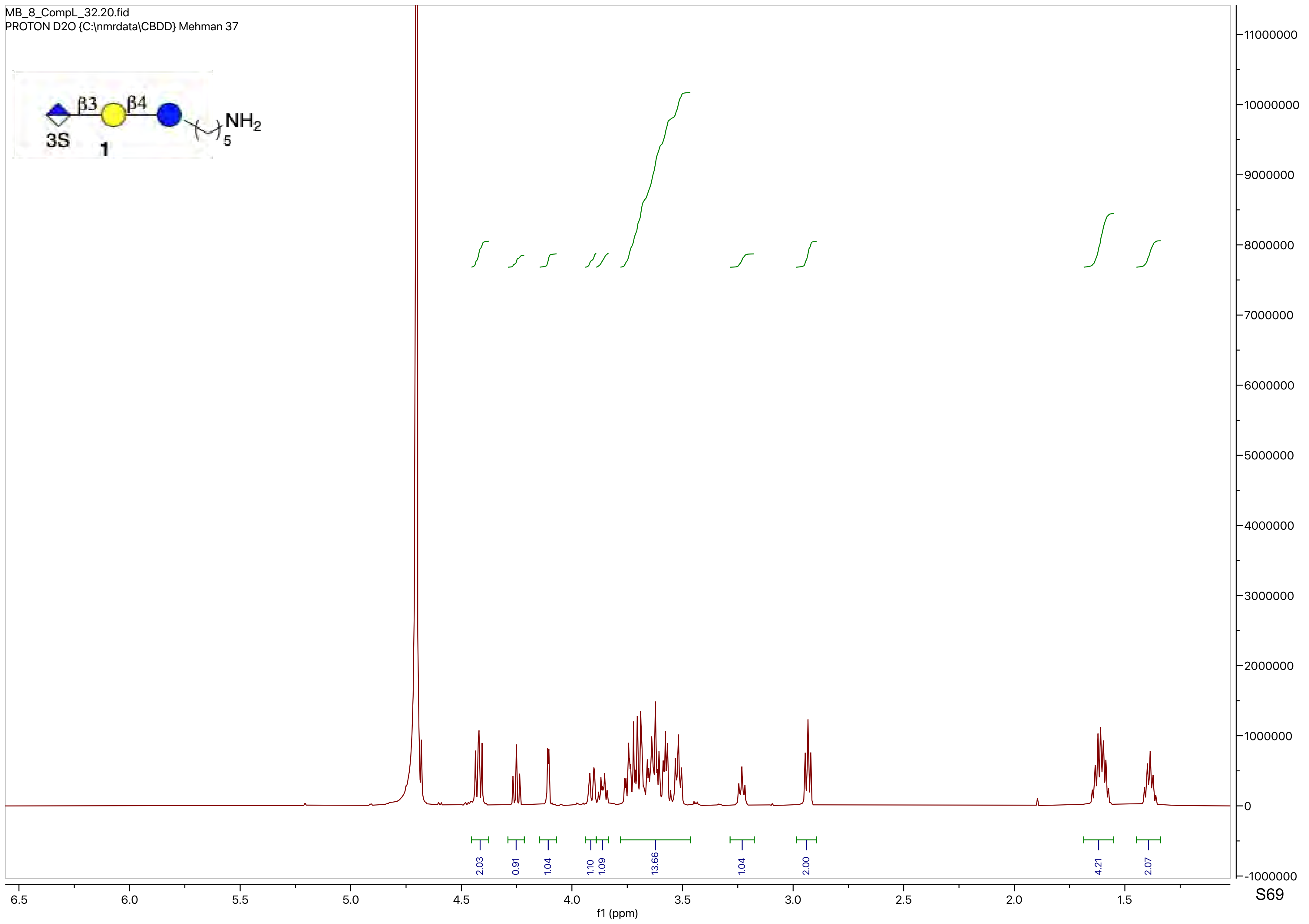
MB_8_CompL_32.20.fid
PROTON D2O {C:\nmrdata\CBDD} Mehman 37
β3
β4
NH2
3S
1
5
2.03
0.91
1.04
1.10
1.09
13.66
1.04
2.00
4.21
2.07
f1 (ppm)

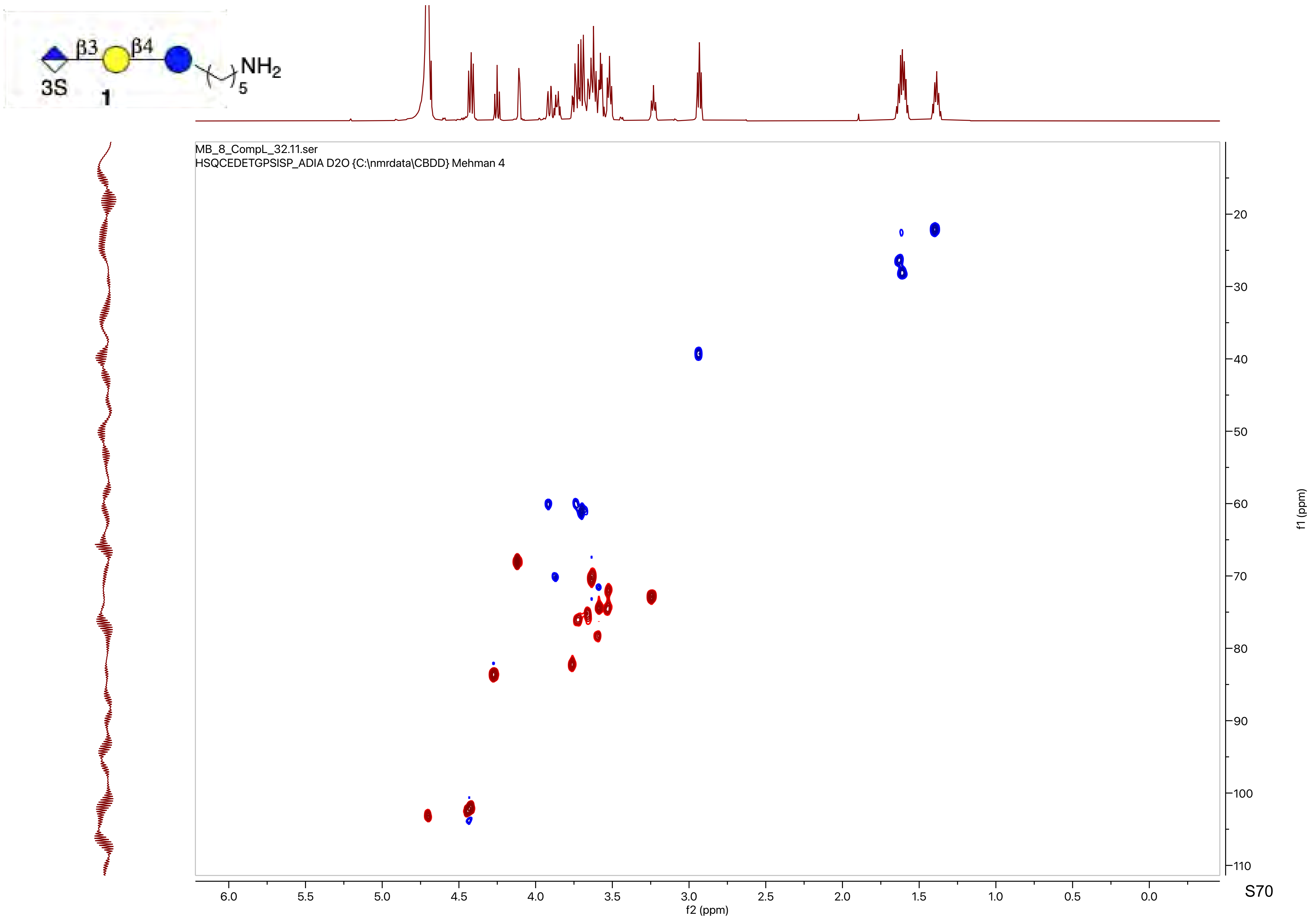

β3
β4
3S
1
NH2
5
MB_8_CompL_32.11.ser
HSQCEDETGPSISP_ADIA D2O {C:\nmrdata\CBDD} Mehman 4
f2 (ppm)
f1 (ppm)

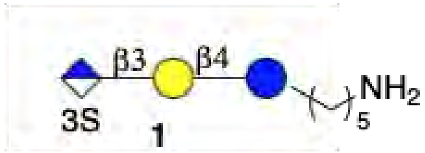
β3
β4
NH2
3S
1
5

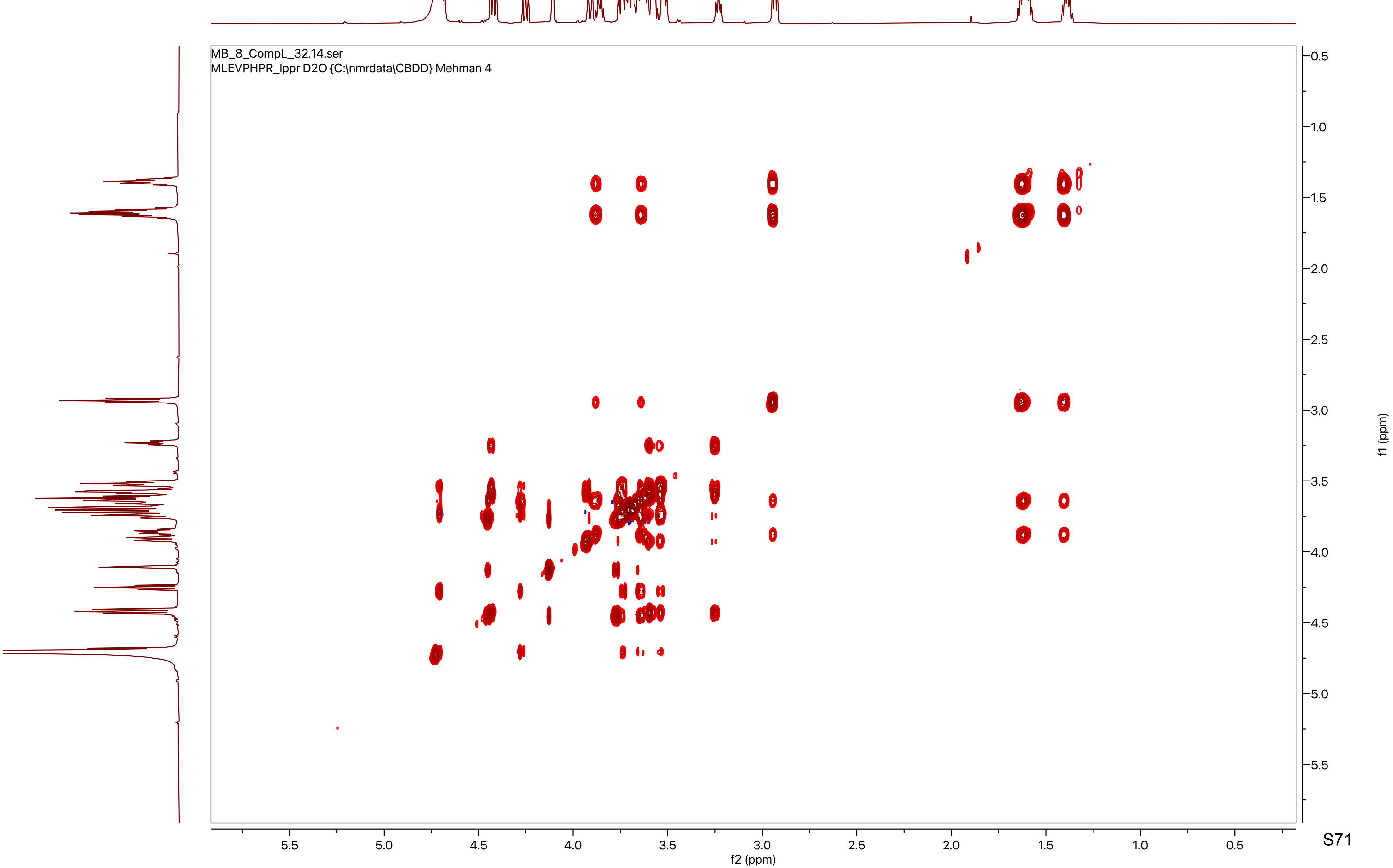
MB_8_CompL_32.14.ser
MLEVPHPR_lppr D2O {C:\nmrdata\CBDD} Mehman 4
f1 (ppm)
f2 (ppm)
5.5
5.0
4.5
4.0
3.5
3.0
2.5
2.0
1.5
1.0
0.5

MB_8_CompL_32.21.fid
WATER_D2O_Lowpowersupp D2O {C:\nmrdata\CBDD} Mehman 37

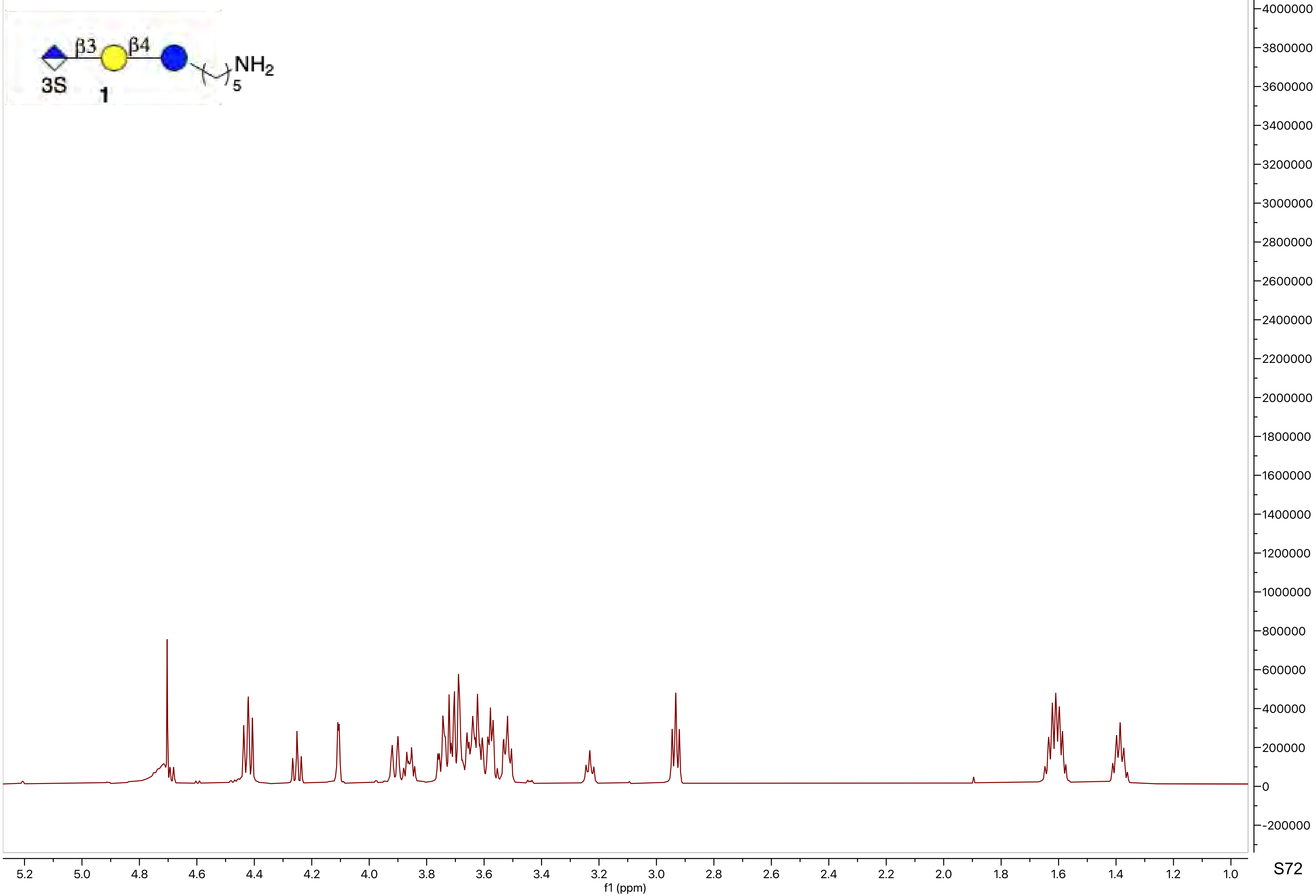

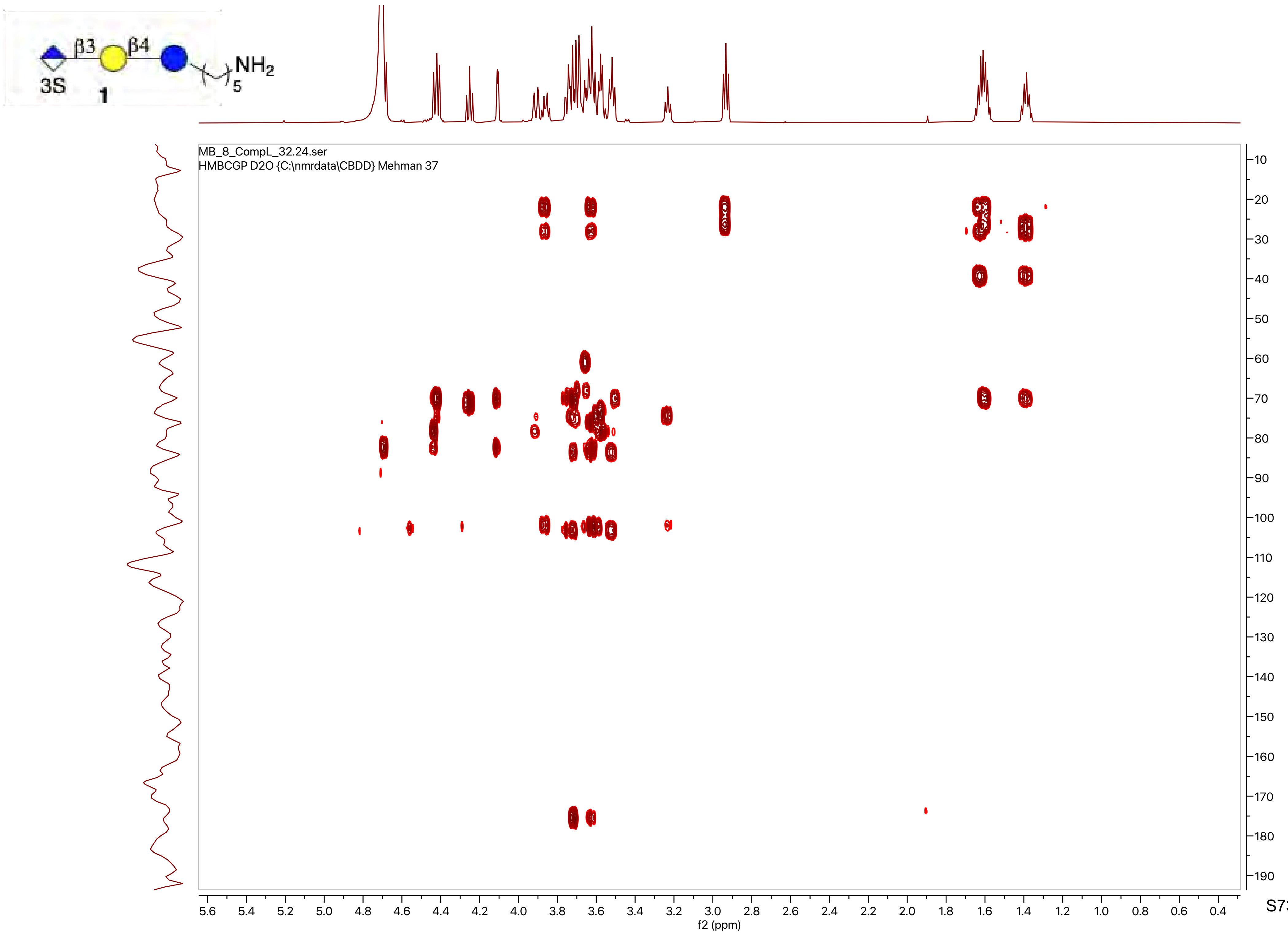

β3
β4
NH2
5
3S
1
MB_8_CompL_32.24.ser
HMBCGP D2O {C:\nmrdata\CBDD} Mehman 37
f2 (ppm)
f1 (ppm)

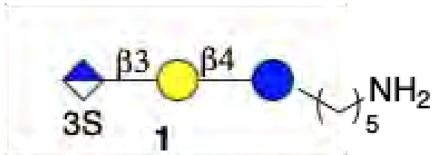
β3
β4
NH2
5
3S

1

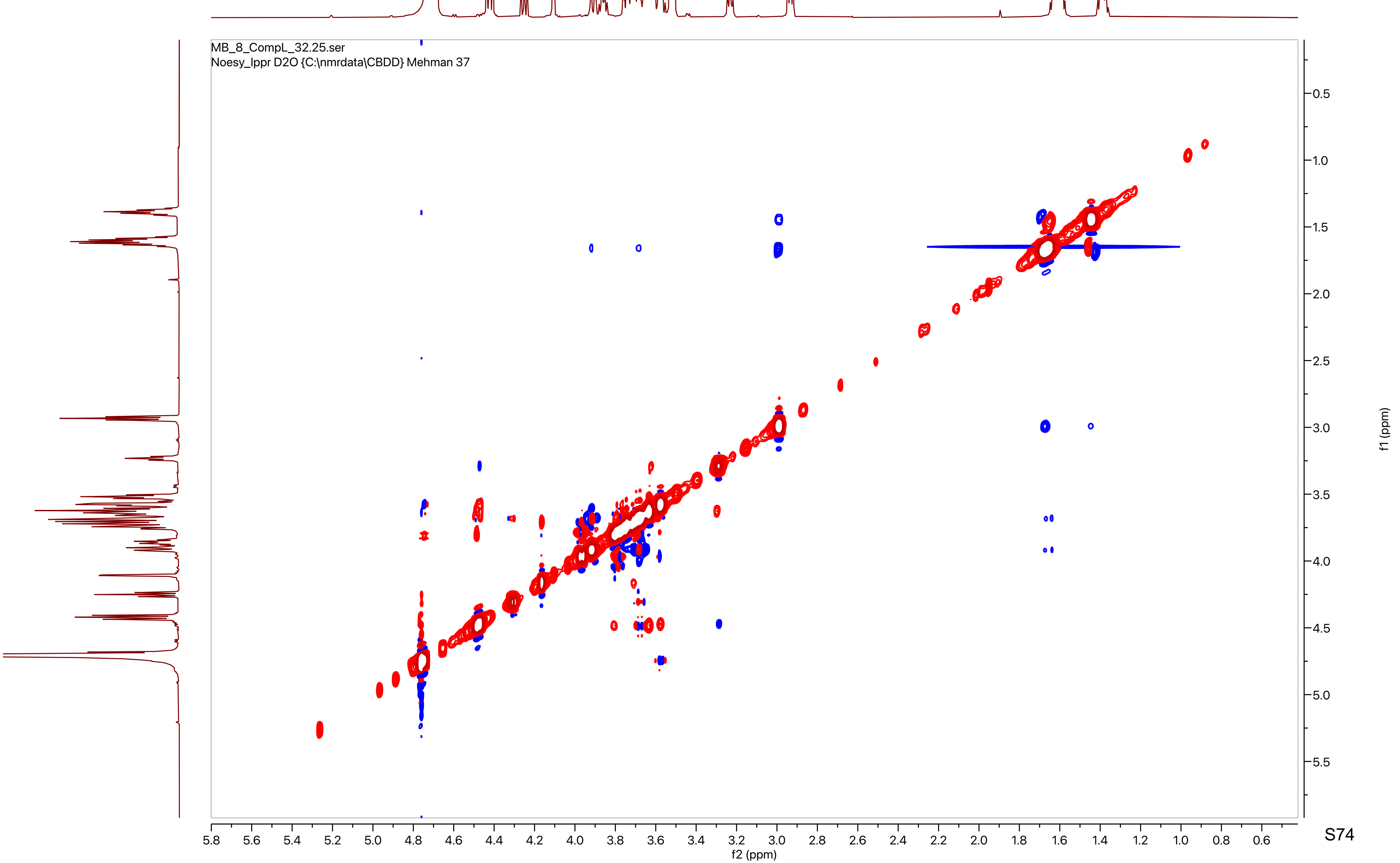
MB_8_CompL_32.25.ser
Noesy_lppr D2O {C:\nmrdata\CBDD} Mehman 37
f2 (ppm)
f1 (ppm)

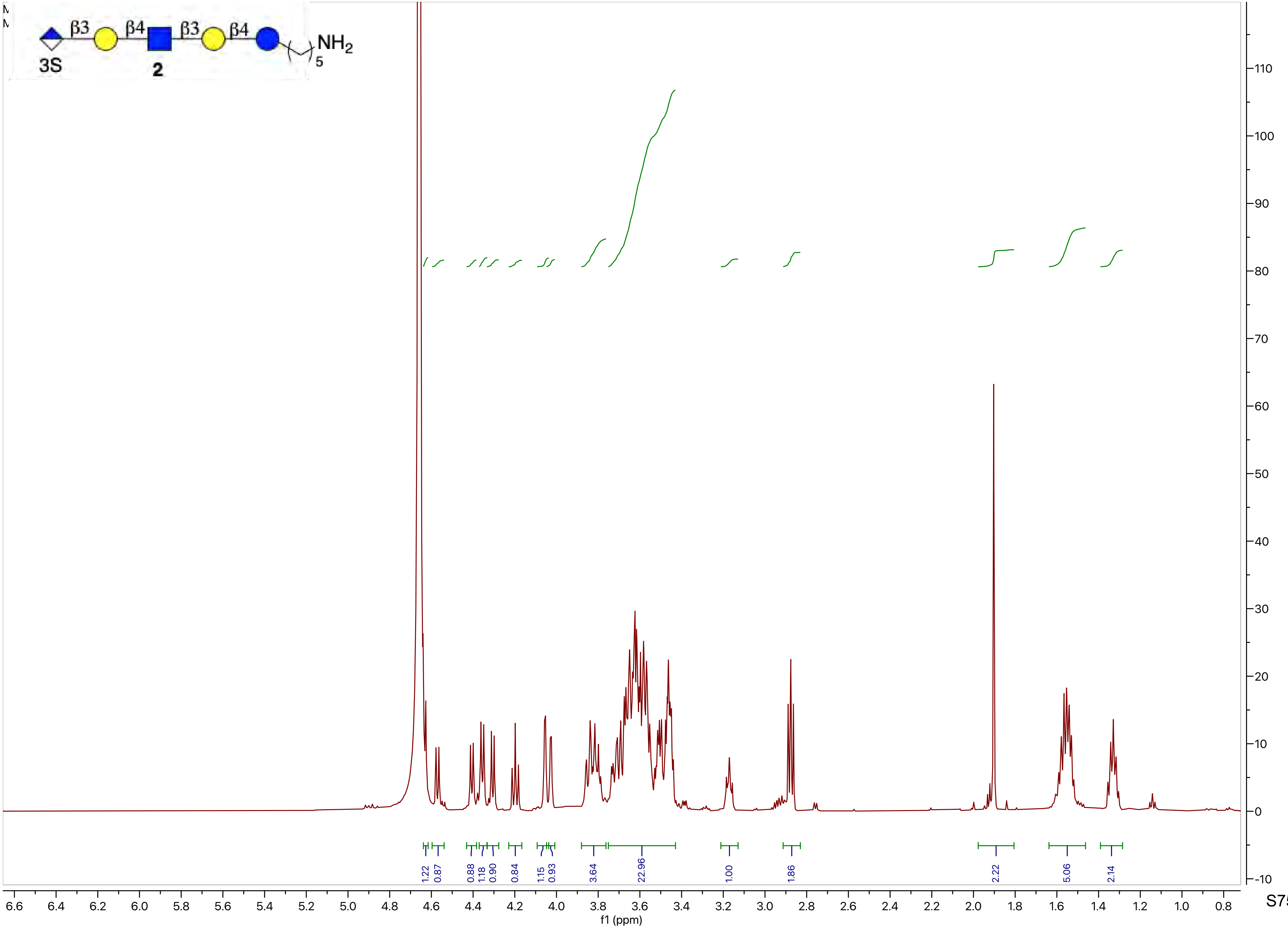
β3
β4
β3
β4
NH2
5
3S
2
f1 (ppm)
1.22
0.87
0.88
1.18
0.90
0.84
1.15
0.93
3.64
22.96
1.00
1.86
2.22
5.06
2.14

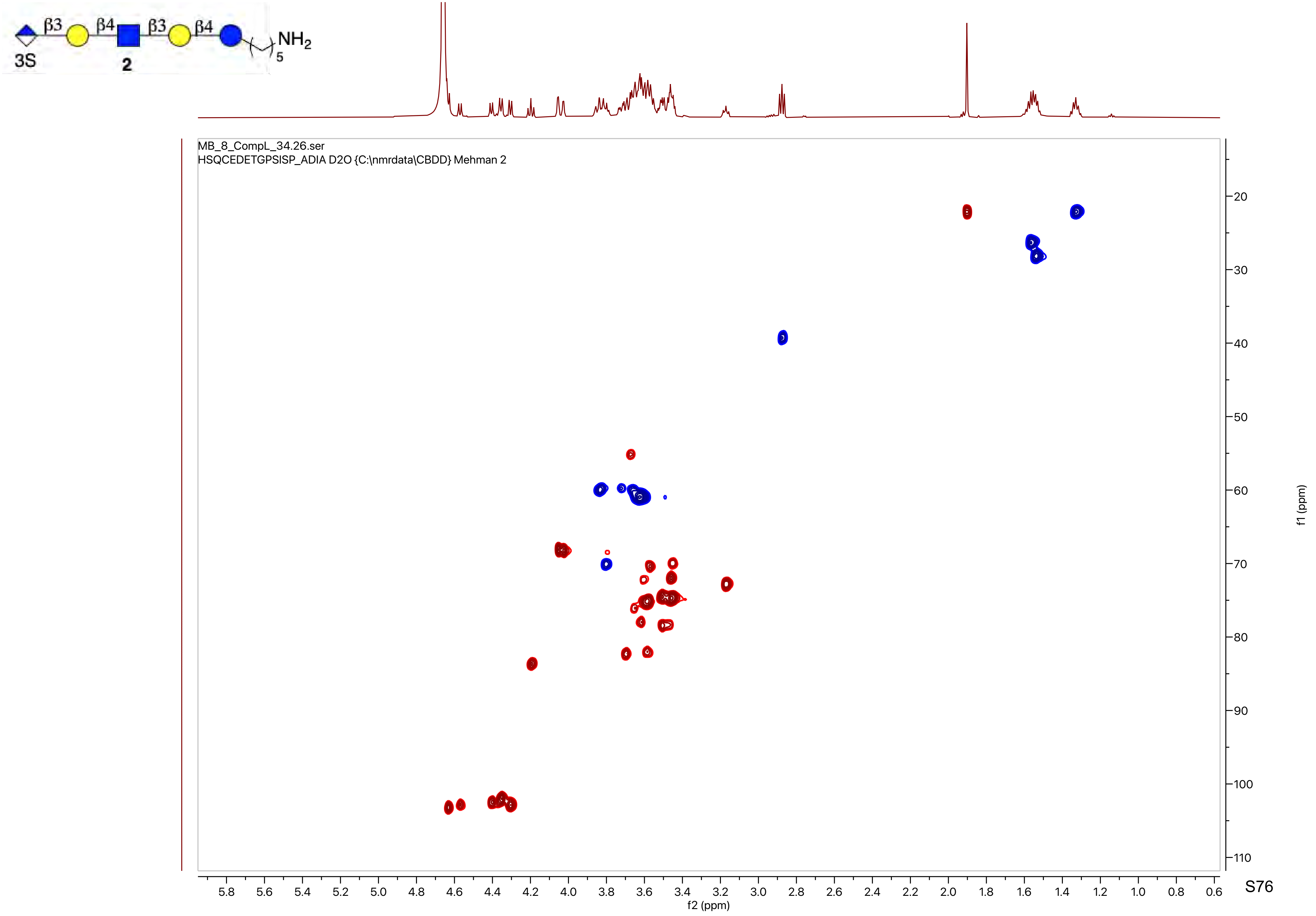
β3
β4
β3
β4
NH2
5
3S
2
MB_8_CompL_34.26.ser
HSQCEDETGPSISP_ADIA D2O {C:\nmrdata\CBDD} Mehman 2
f1 (ppm)
f2 (ppm)

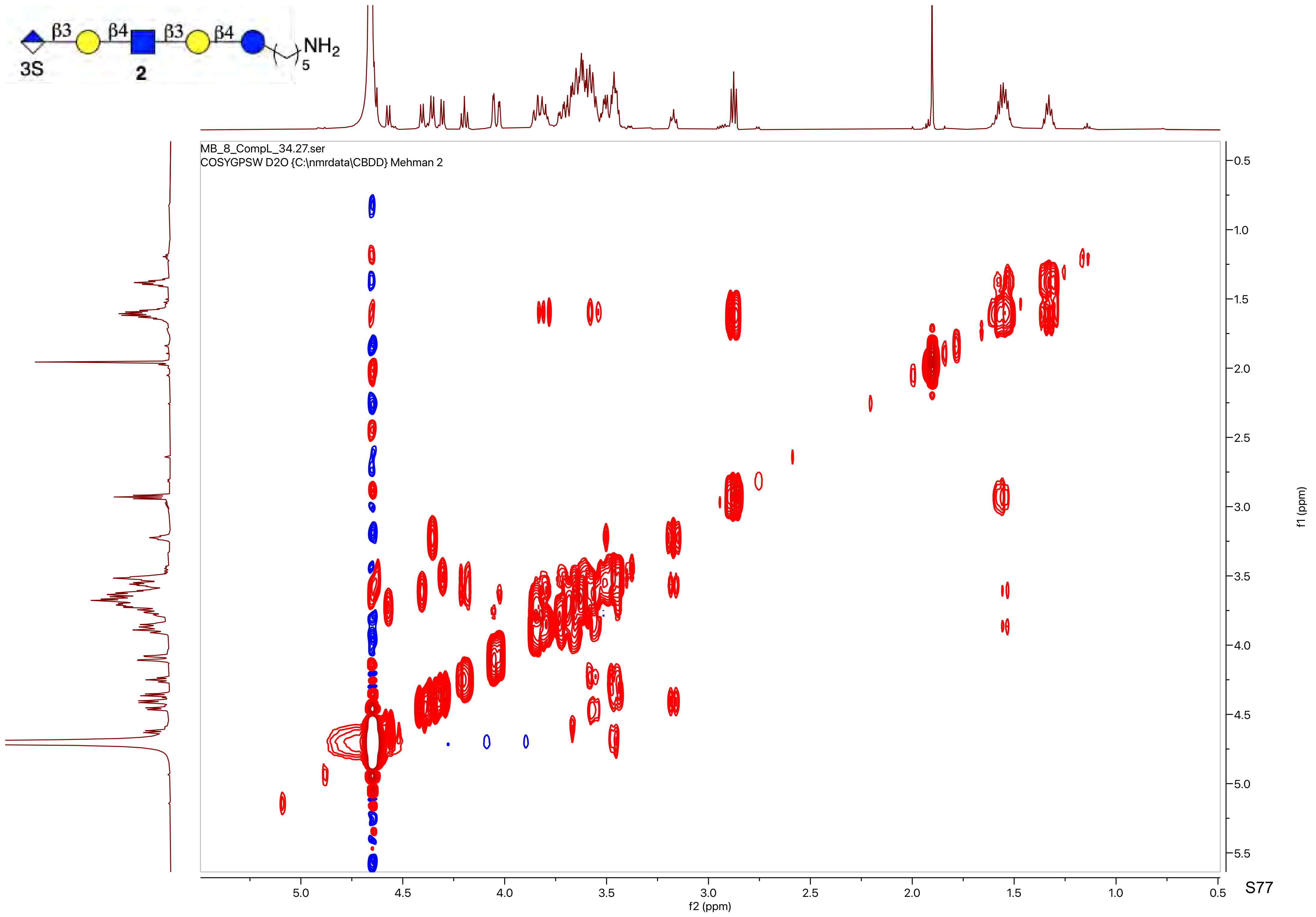

β3
β4
β3
β4
NH2
5
3S
2
MB_8_CompL_34.27.ser
COSYGPSW D2O {C:\nmrdata\CBDD} Mehman 2
f2 (ppm)
f1 (ppm)

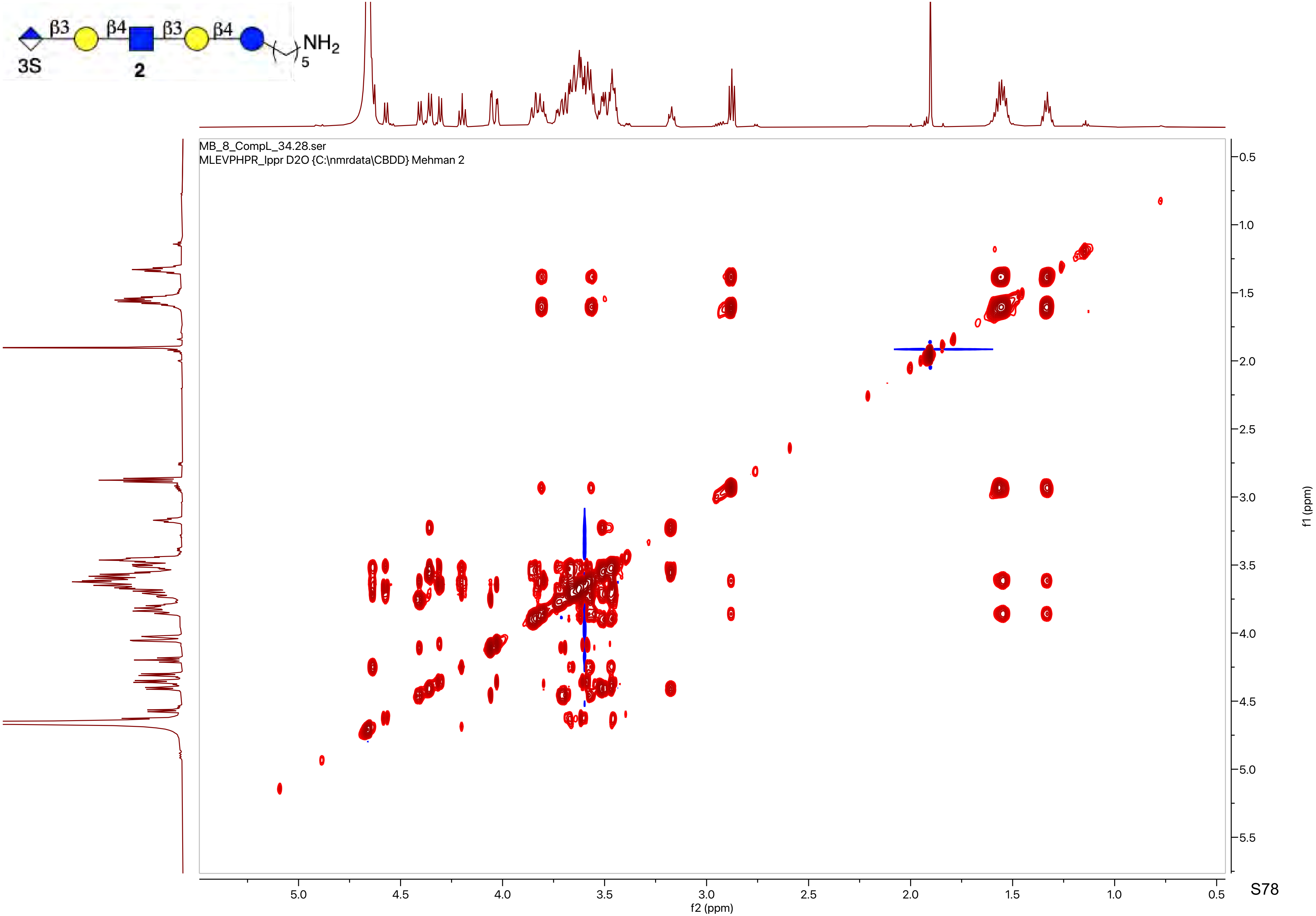
β3
β4
β3
β4
NH2
5
3S
2
MB_8_CompL_34.28.ser
MLEVPHPR_Ippr D2O {C:\nmrdata\CBDD} Mehman 2
f2 (ppm)
f1 (ppm)

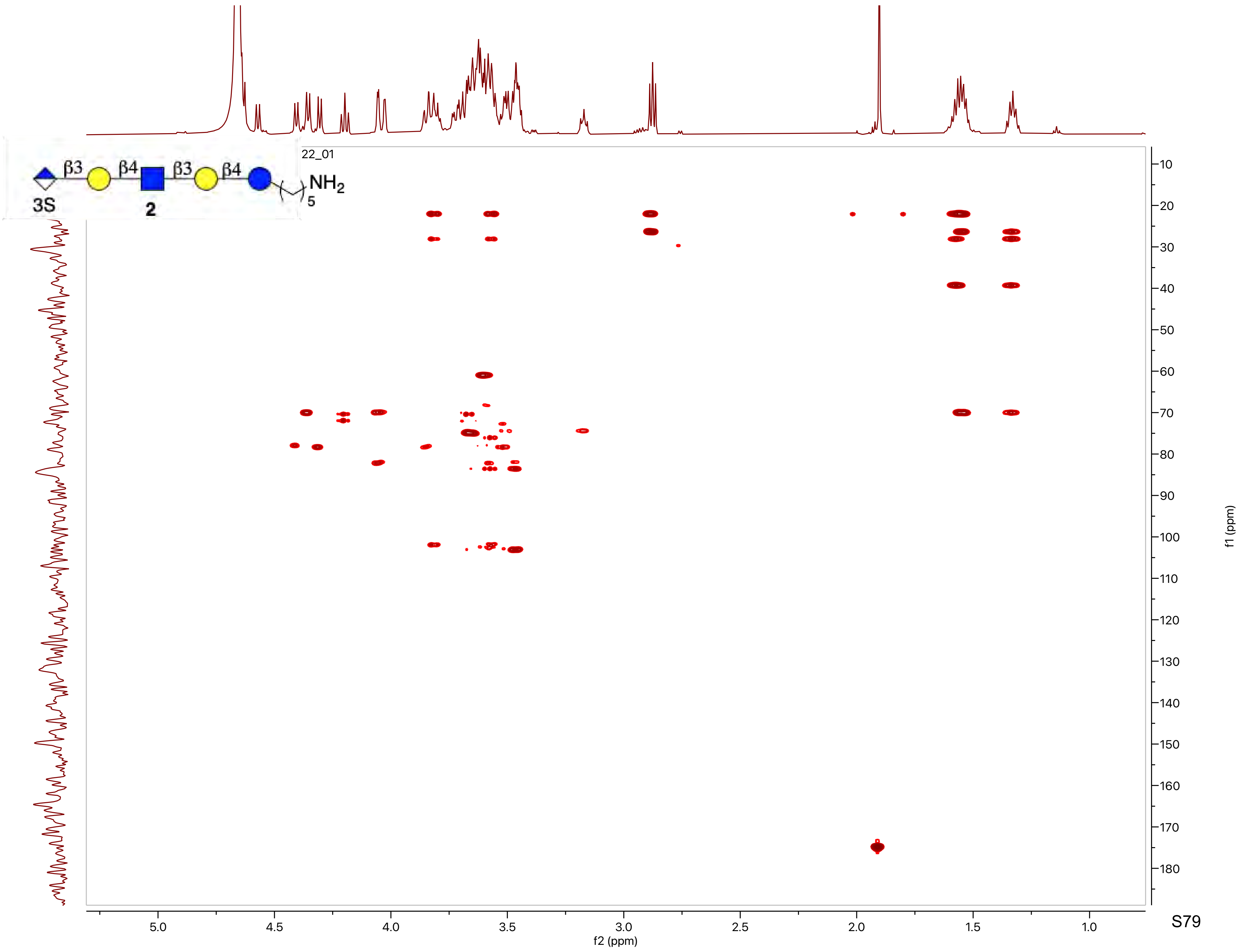

β3
β4
β3
β4
NH2
5
3S
2
22_01
f1 (ppm)
f2 (ppm)

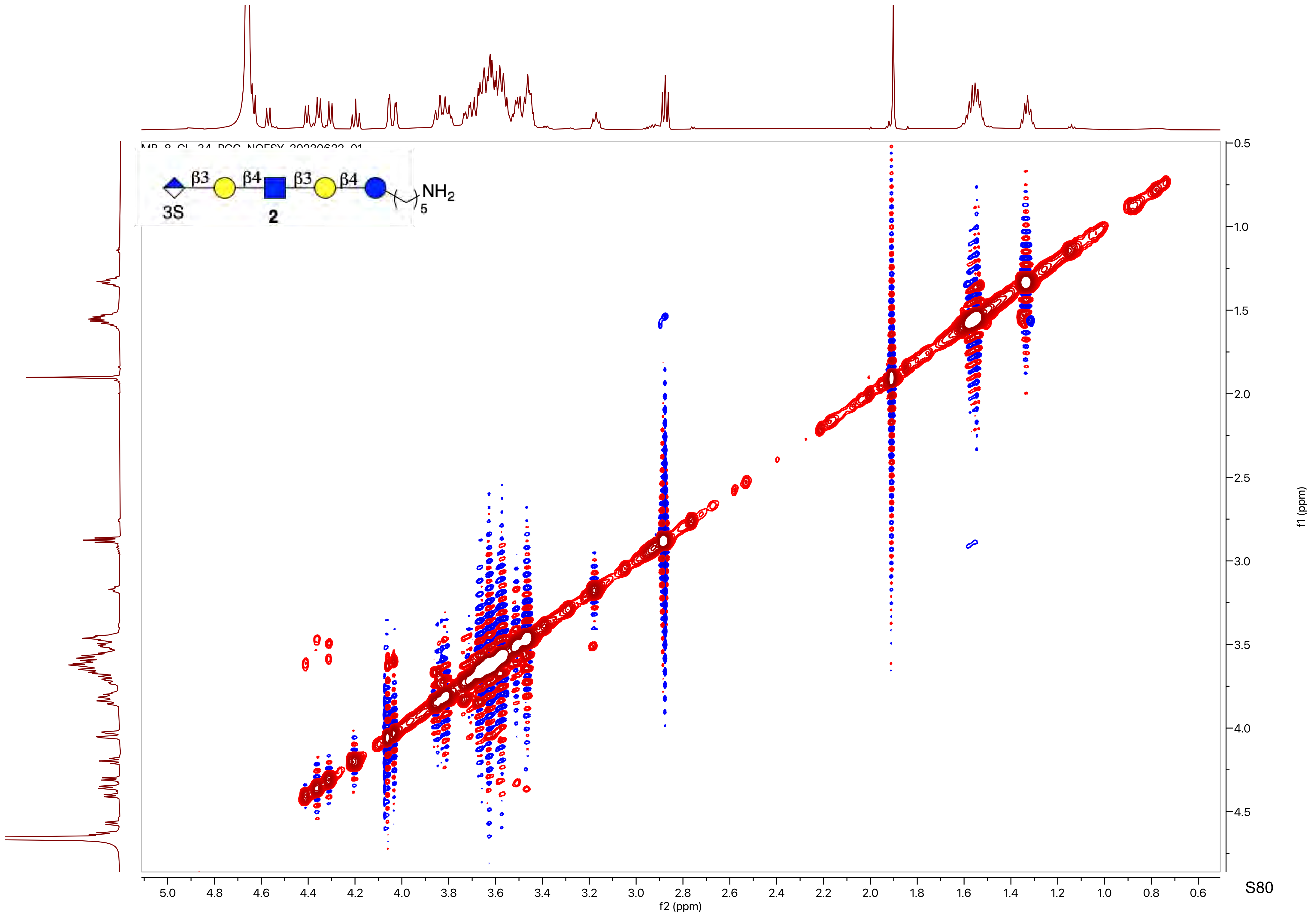
β3
β4
β3
β4
NH2
5
3S
2
f2 (ppm)
f1 (ppm)

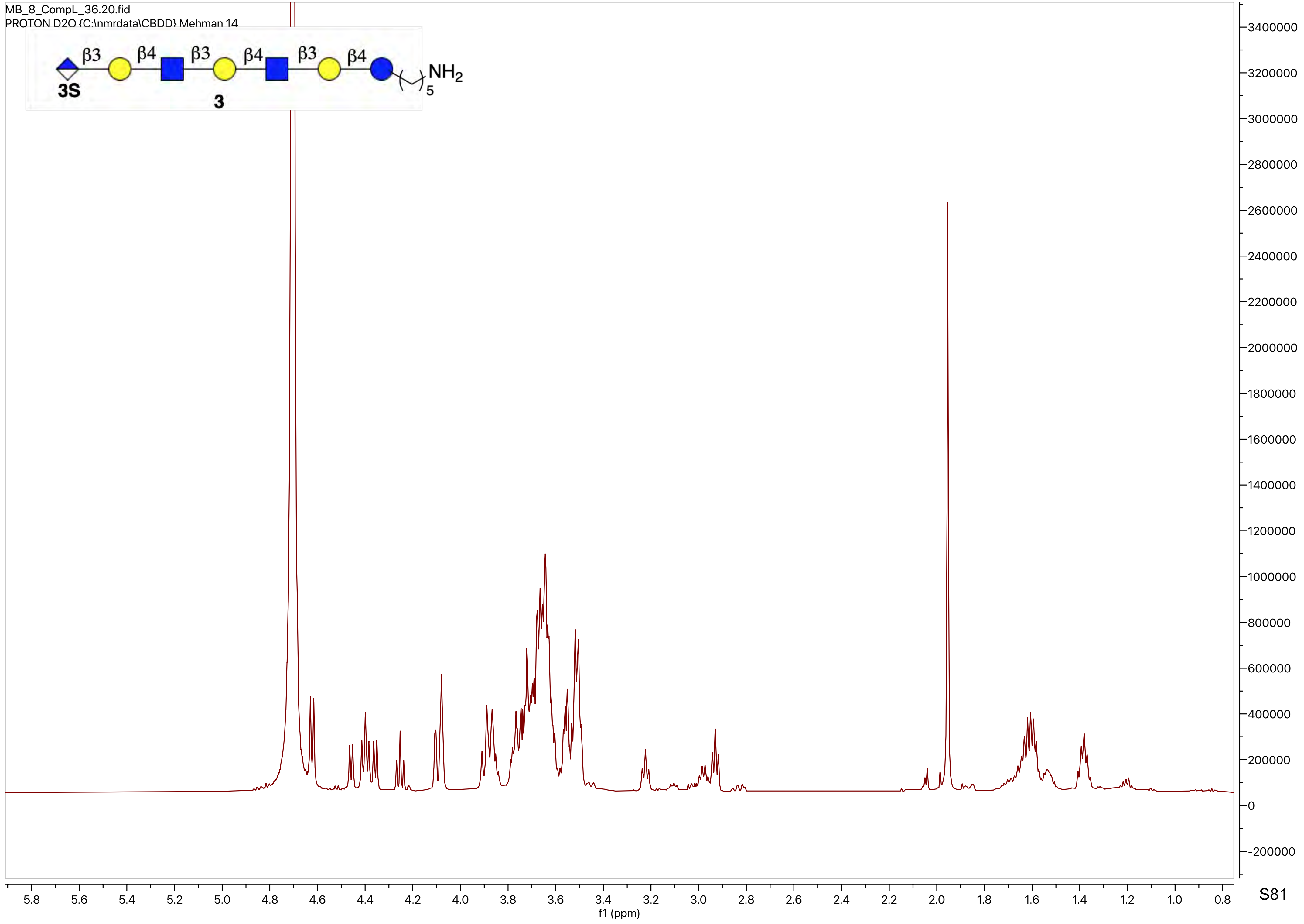
MB_8_CompL_36.20.fid
PROTON D2O {C:\nmrdata\CBDD} Mehman 14
β3
β4
β3
β4
β3
β4
NH2
5
3S
3
f1 (ppm)
5.8 5.6 5.4 5.2 5.0 4.8 4.6 4.4 4.2 4.0 3.8 3.6 3.4 3.2 3.0 2.8 2.6 2.4 2.2 2.0 1.8 1.6 1.4 1.2 1.0 0.8
3400000 3200000 3000000 2800000 2600000 2400000 2200000 2000000 1800000 1600000 1400000 1200000 1000000 800000 600000 400000 200000 0 -200000

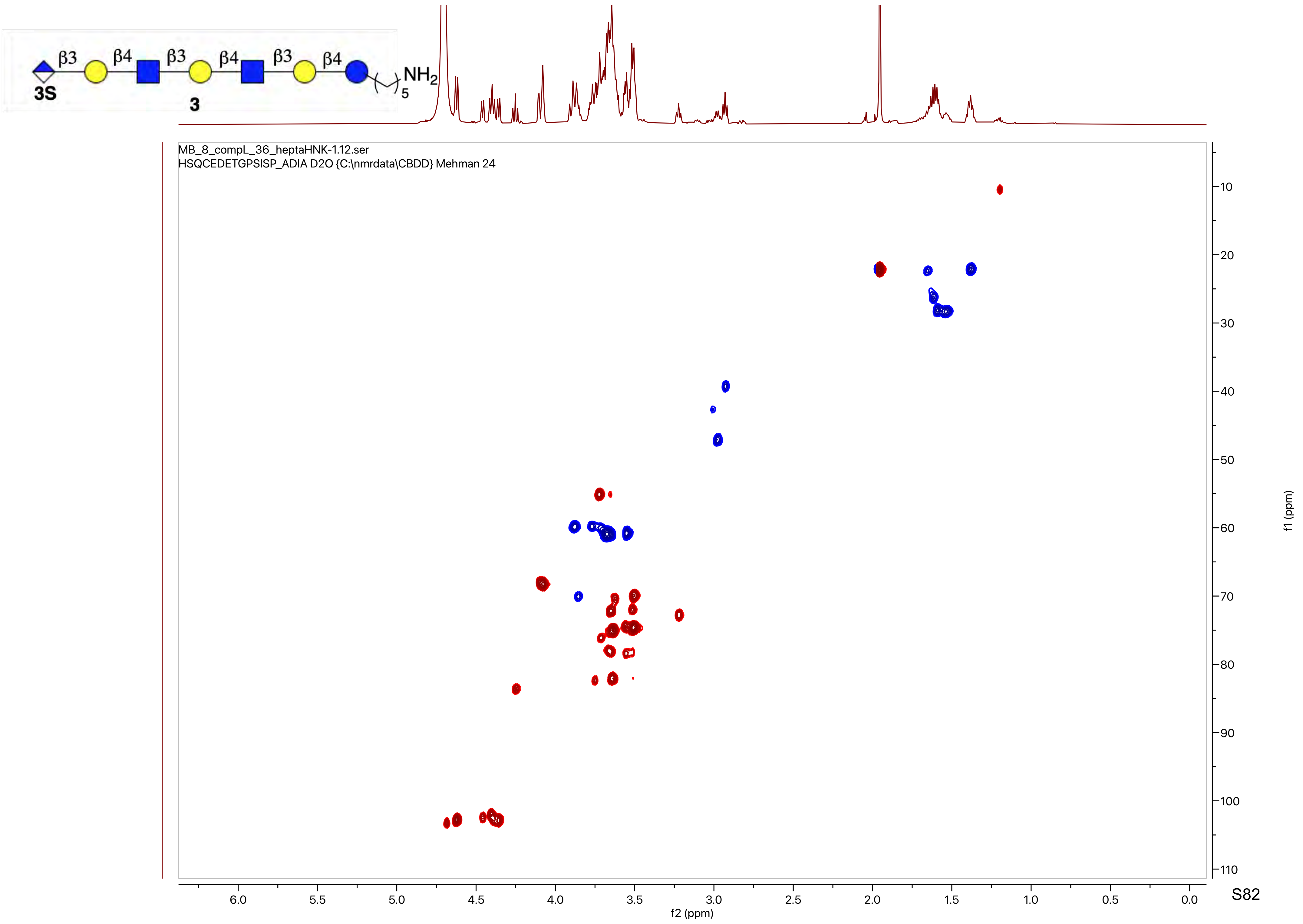

β3
β4
β3
β4
β3
β4
NH2
5
3S
3
MB_8_compL_36_heptaHNK-1.12.ser
HSQCEDETGPSISP_ADIA D2O {C:\nmrdata\CBDD} Mehman 24
f2 (ppm)
f1 (ppm)

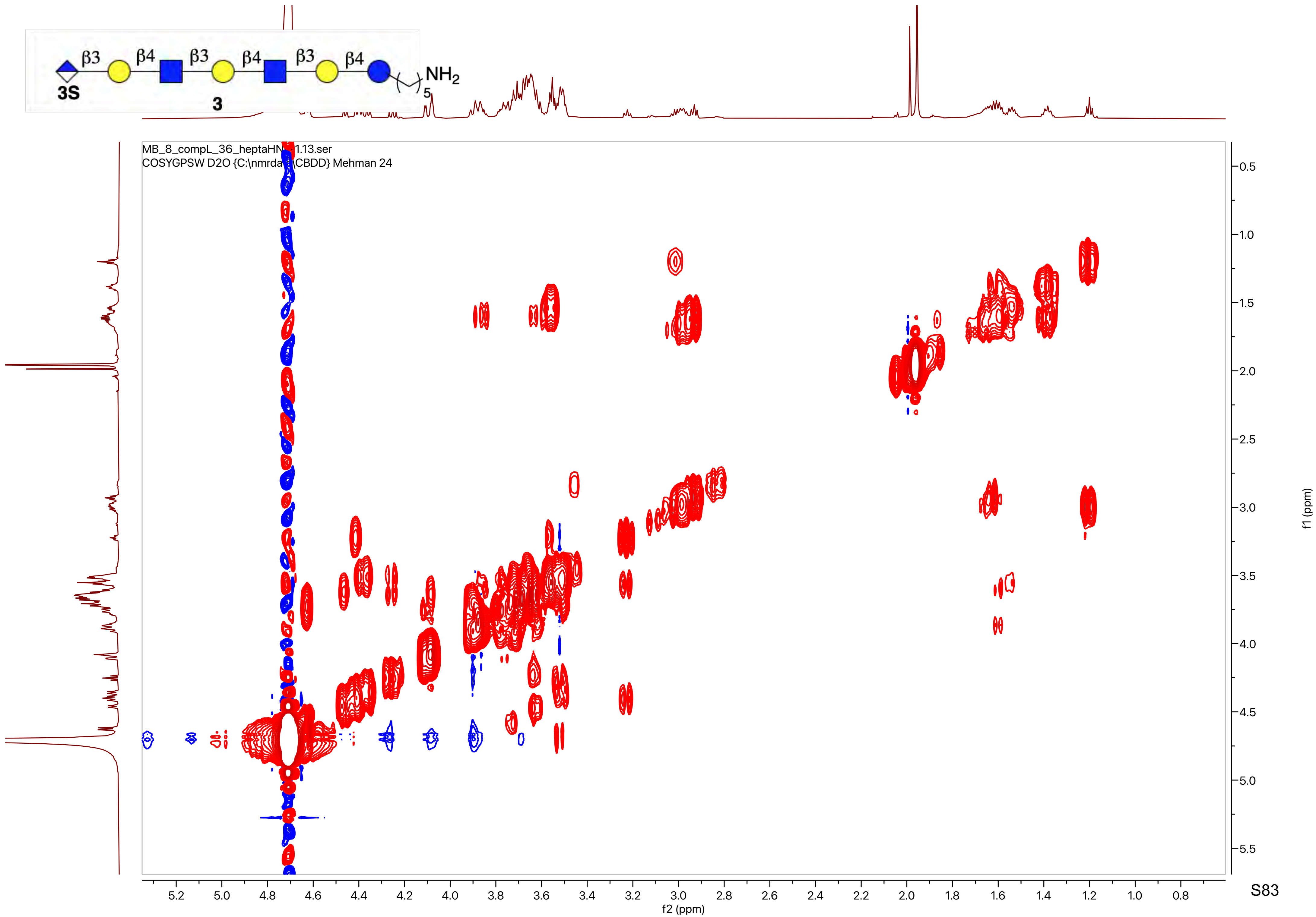
β3
β4
β3
β4
β3
β4
NH2
5
3S
3
MB_8_compL_36_heptaHN 1.13.ser
COSYGPSW D2O {C:\nmrda CBDD} Mehman 24
f2 (ppm)
f1 (ppm)

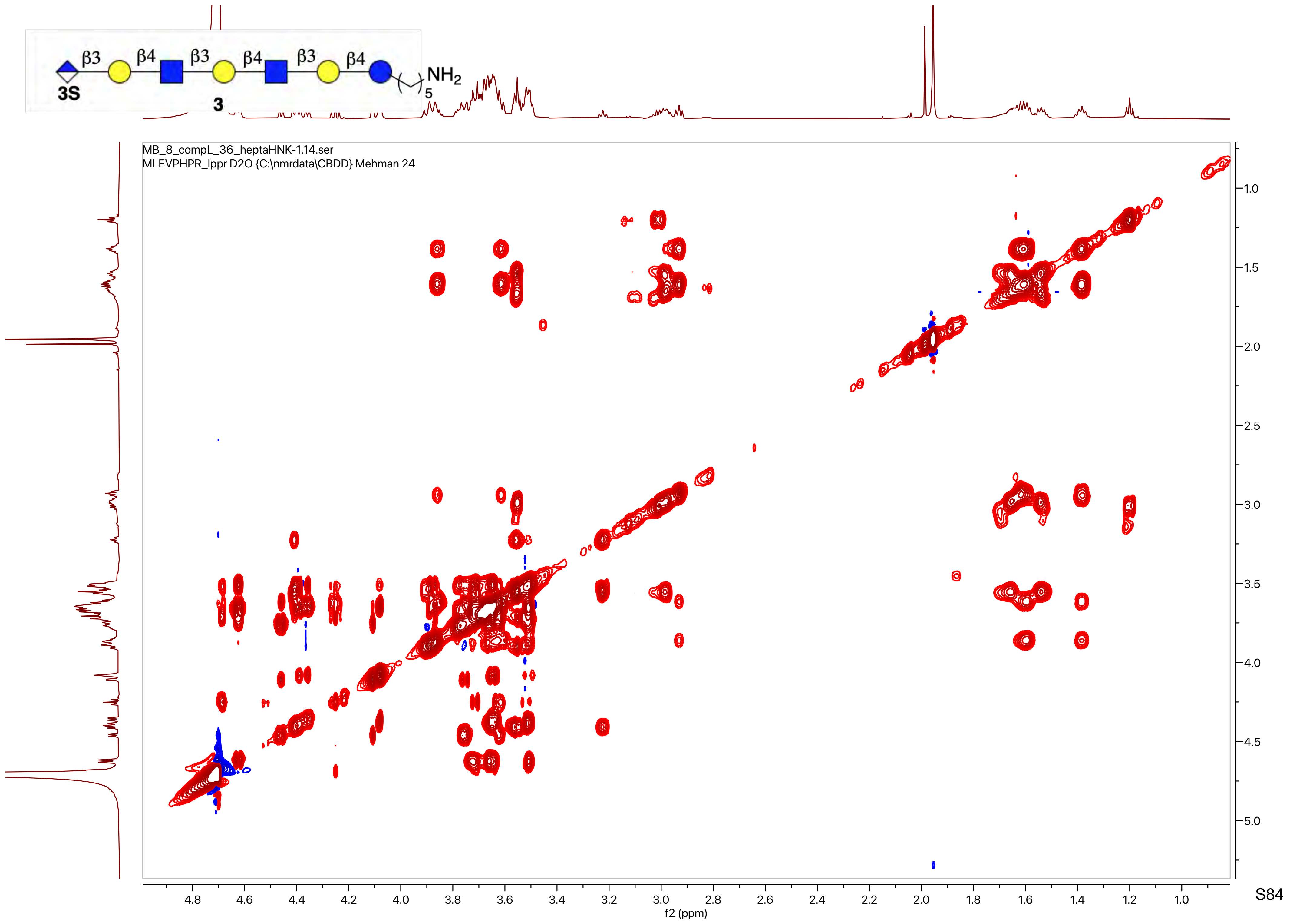
β3
β4
β3
β4
β3
β4
3S
3
NH2
5
MB_8_compL_36_heptaHNK-1.14.ser
MLEVPHPR_lppr D2O {C:\nmrdata\CBDD} Mehman 24
f2 (ppm)
f1 (ppm)

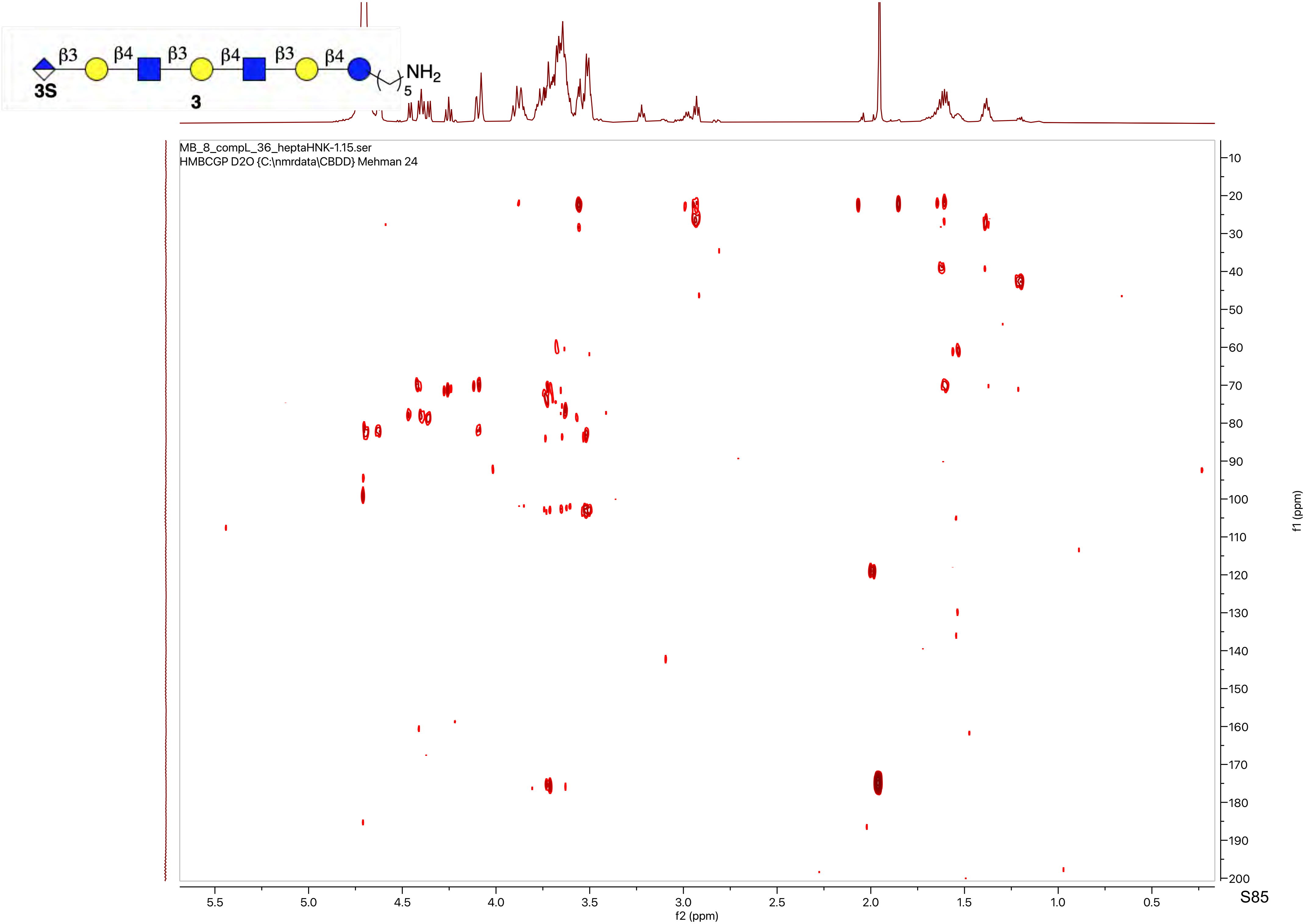
β3
β4
β3
β4
β3
β4
NH2
5
3S
3
MB_8_compL_36_heptaHNK-1.15.ser
HMBCGP D2O {C:\nmrdata\CBDD} Mehman 24
f2 (ppm)
f1 (ppm)

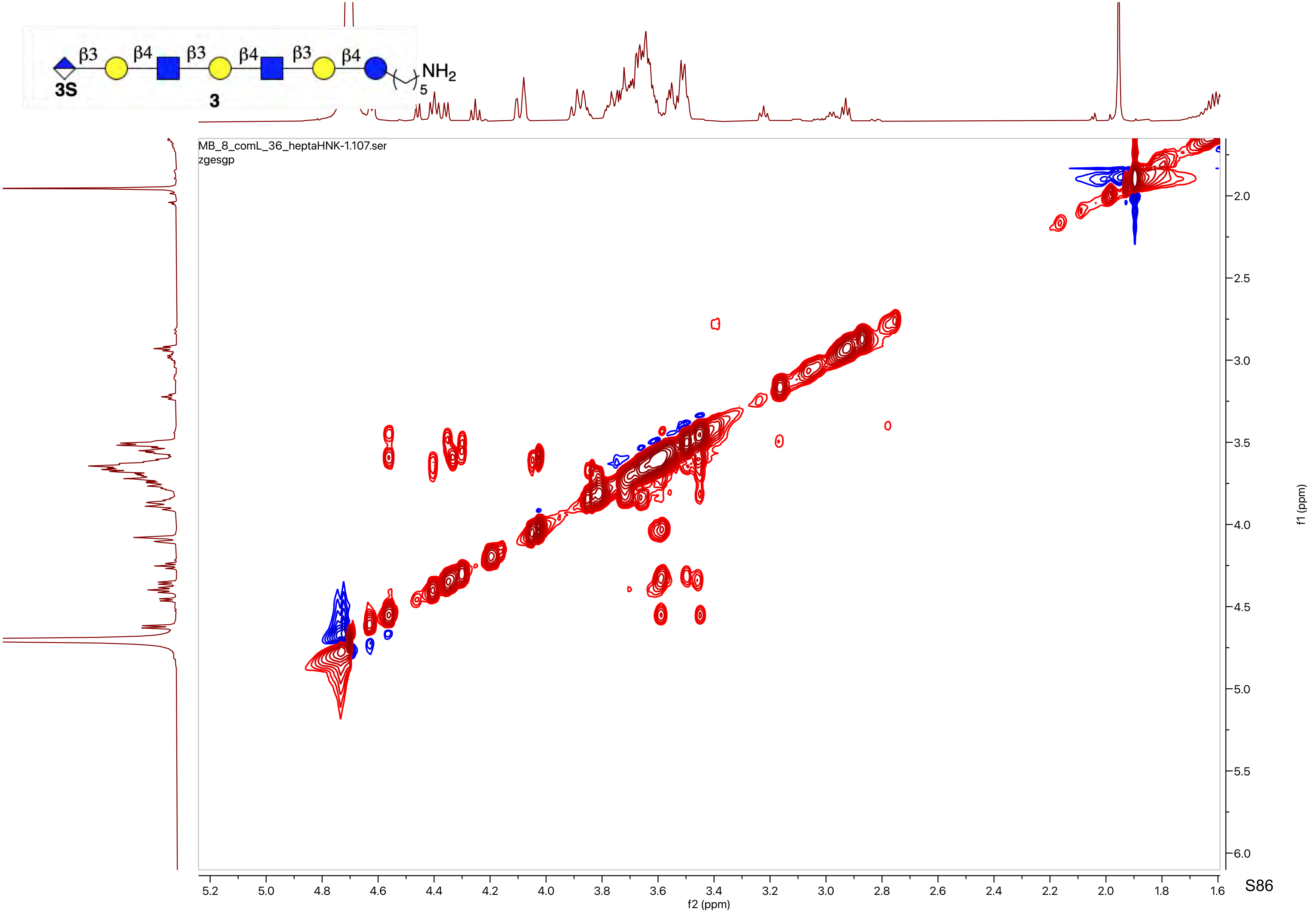
β3
β4
β3
β4
β3
β4
NH2
5
3S
3
MB_8_comL_36_heptaHNK-1.107.ser
zgesgp
f2 (ppm)
f1 (ppm)

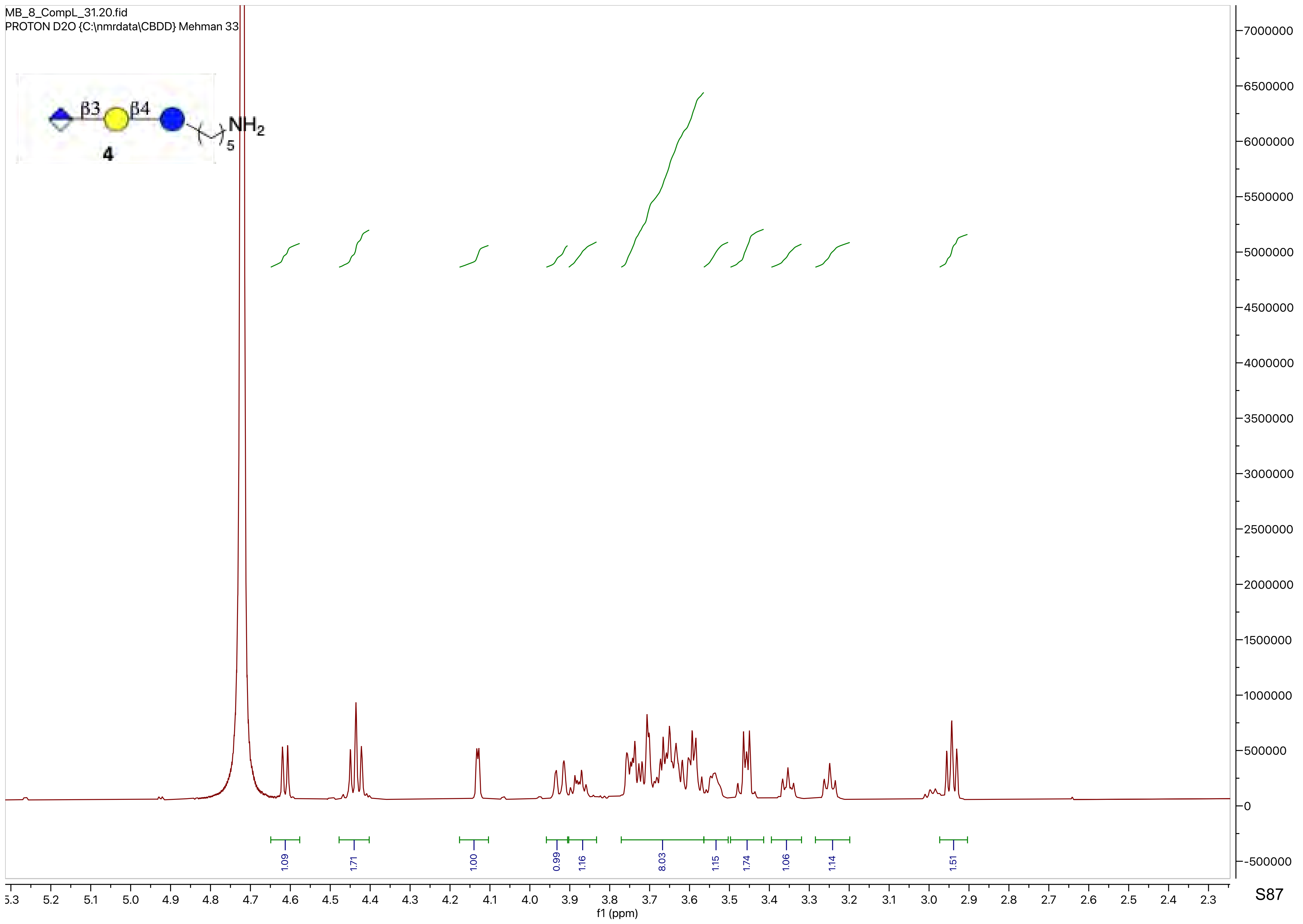
MB_8_CompL_31.20.fid
PROTON D2O {C:\nmrdata\CBDD} Mehman 33
β3
β4
NH2
5
4
1.09
1.71
1.00
0.99
1.16
8.03
1.15
1.74
1.06
1.14
1.51
f1 (ppm)

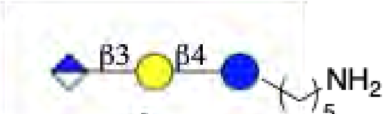
β3
β4
NH2
5

**4**

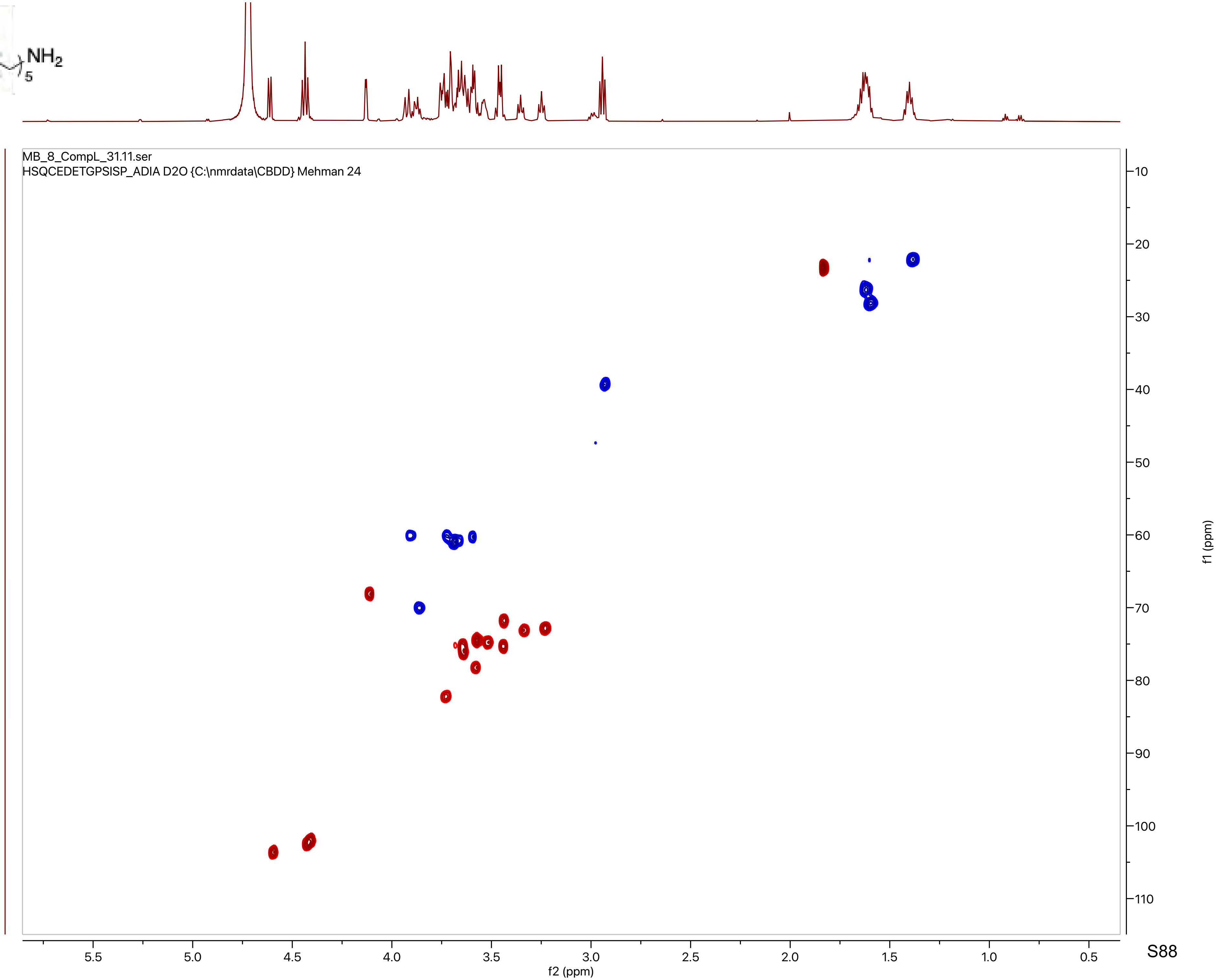
MB_8_CompL_31.11.ser
HSQCEDETGPSISP_ADIA D2O {C:\nmrdata\CBDD} Mehman 24
f2 (ppm)
f1 (ppm)

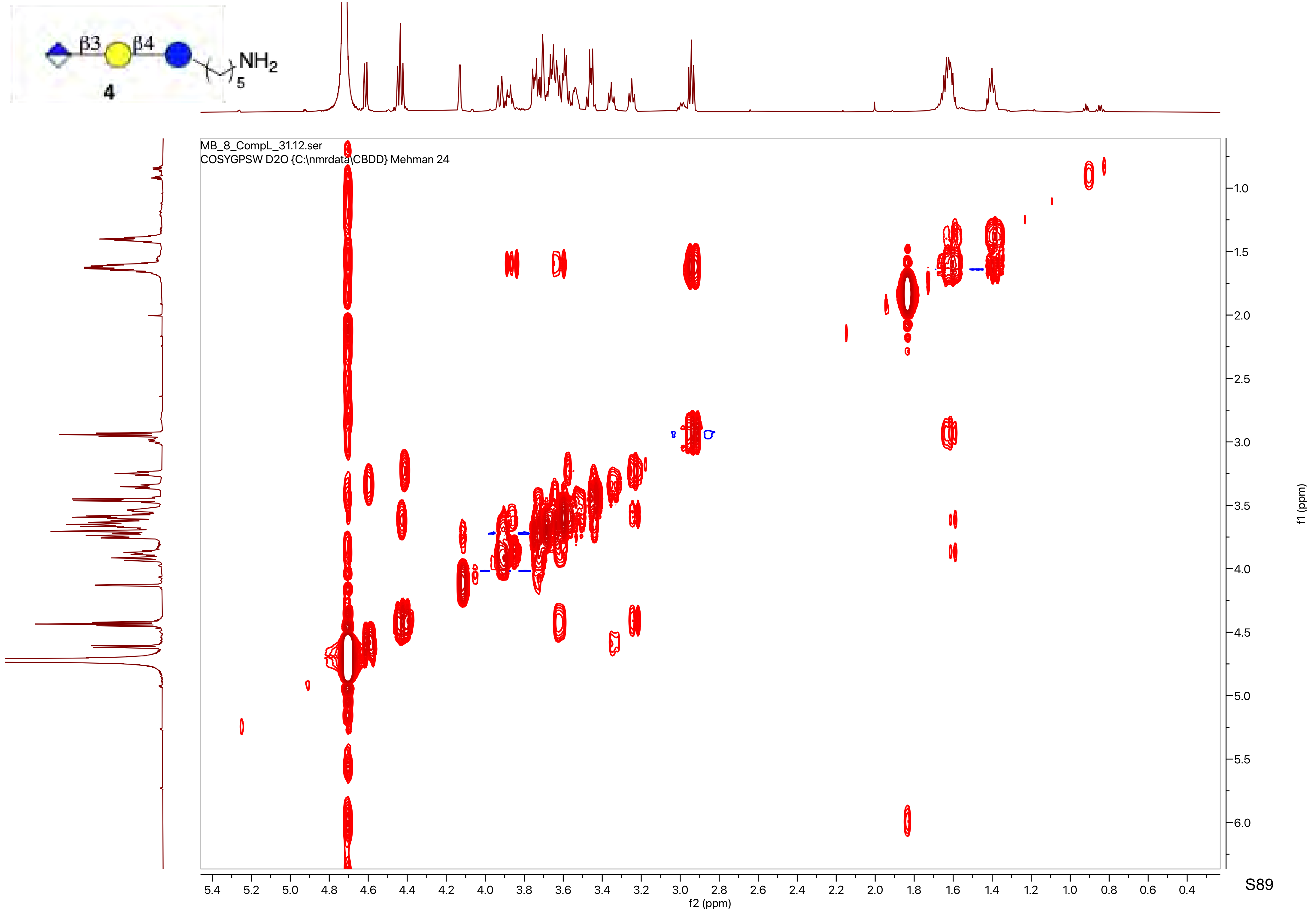
β3
β4
NH2
5
4
MB_8_CompL_31.12.ser
COSYGPSW D2O {C:\nmrdata\CBDD} Mehman 24
f2 (ppm)
f1 (ppm)

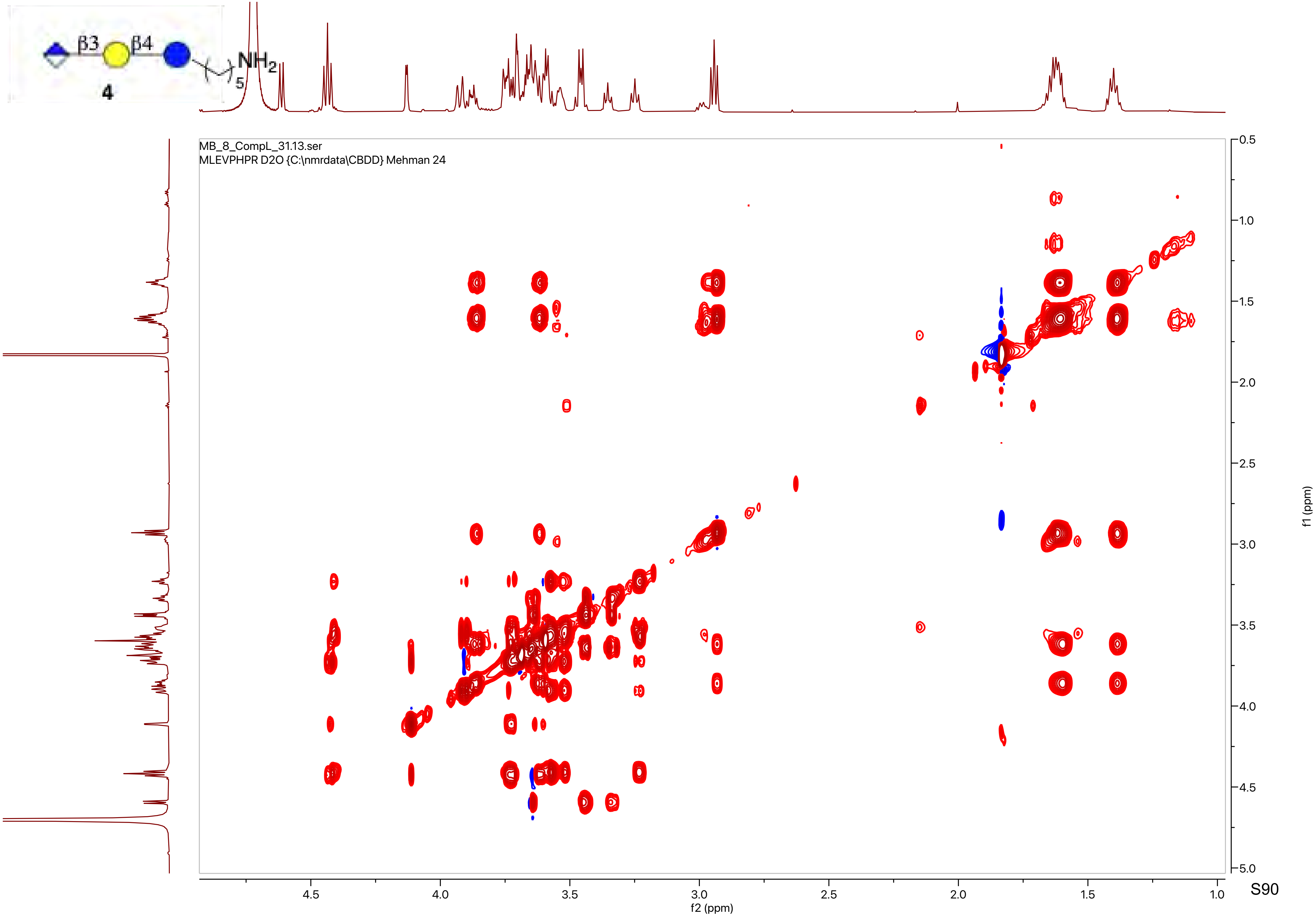
β3
β4
NH2
5
4
MB_8_CompL_31.13.ser
MLEVPHPR D2O {C:\nmrdata\CBDD} Mehman 24
f2 (ppm)
f1 (ppm)

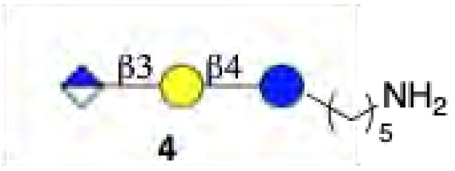
β3
β4
NH2
5
4

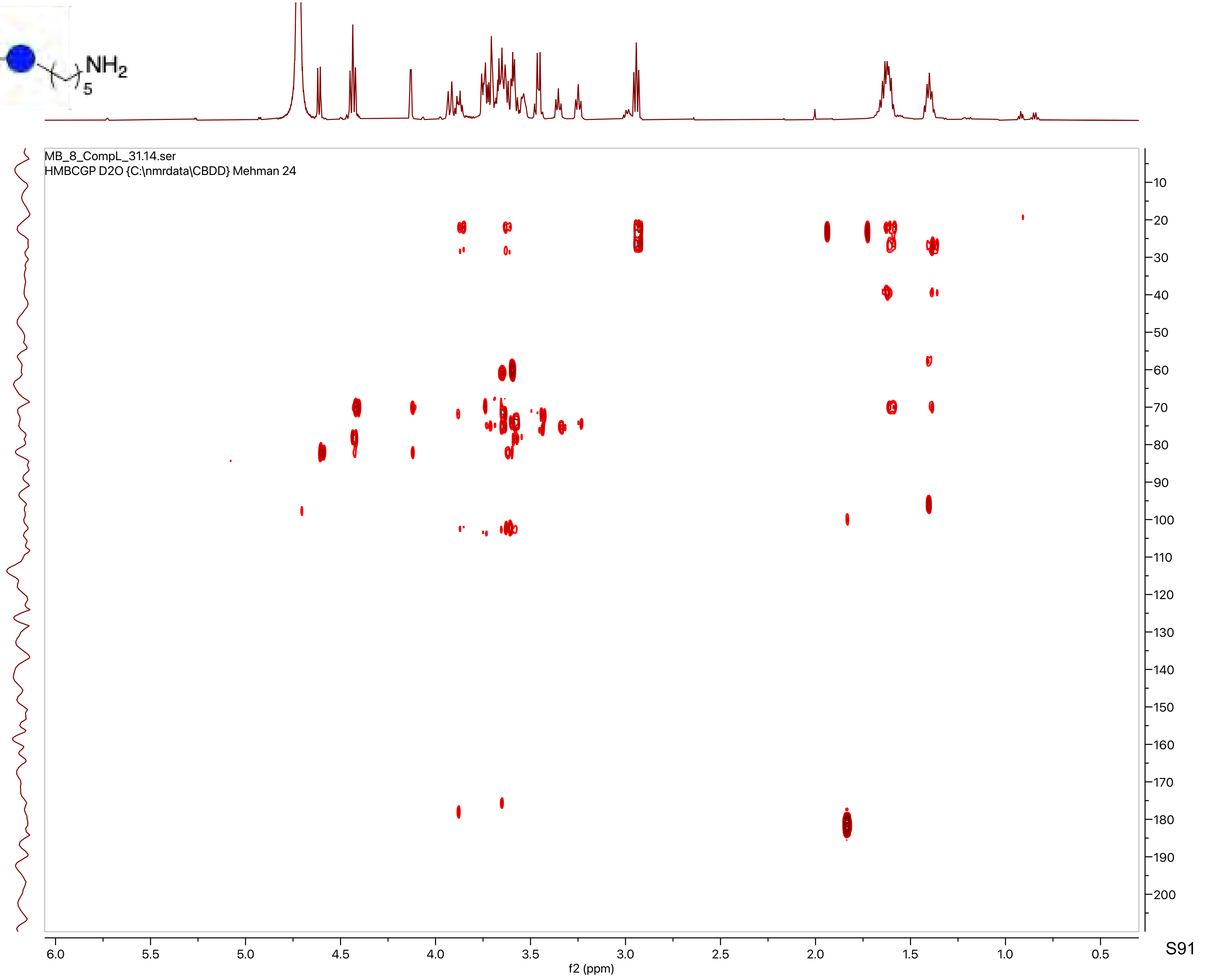
MB_8_CompL_31.14.ser
HMBCGP D2O {C:\nmrdata\CBDD} Mehman 24
f2 (ppm)
f1 (ppm)

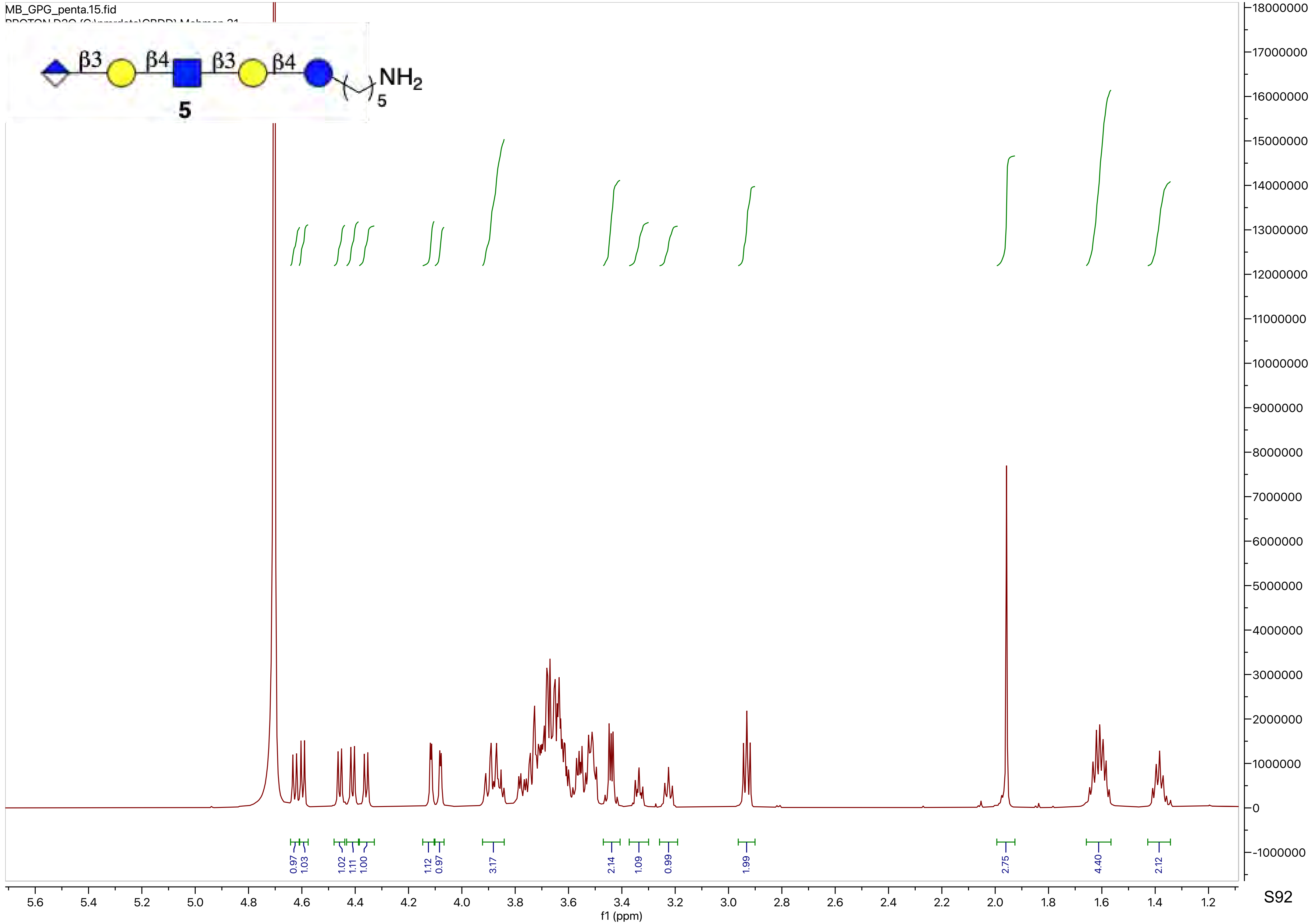
MB_GPG_penta.15.fid
β3
β4
β3
β4
NH2
5
5
0.97
1.03
1.02
1.11
1.00
1.12
0.97
3.17
2.14
1.09
0.99
1.99
2.75
4.40
2.12
f1 (ppm)

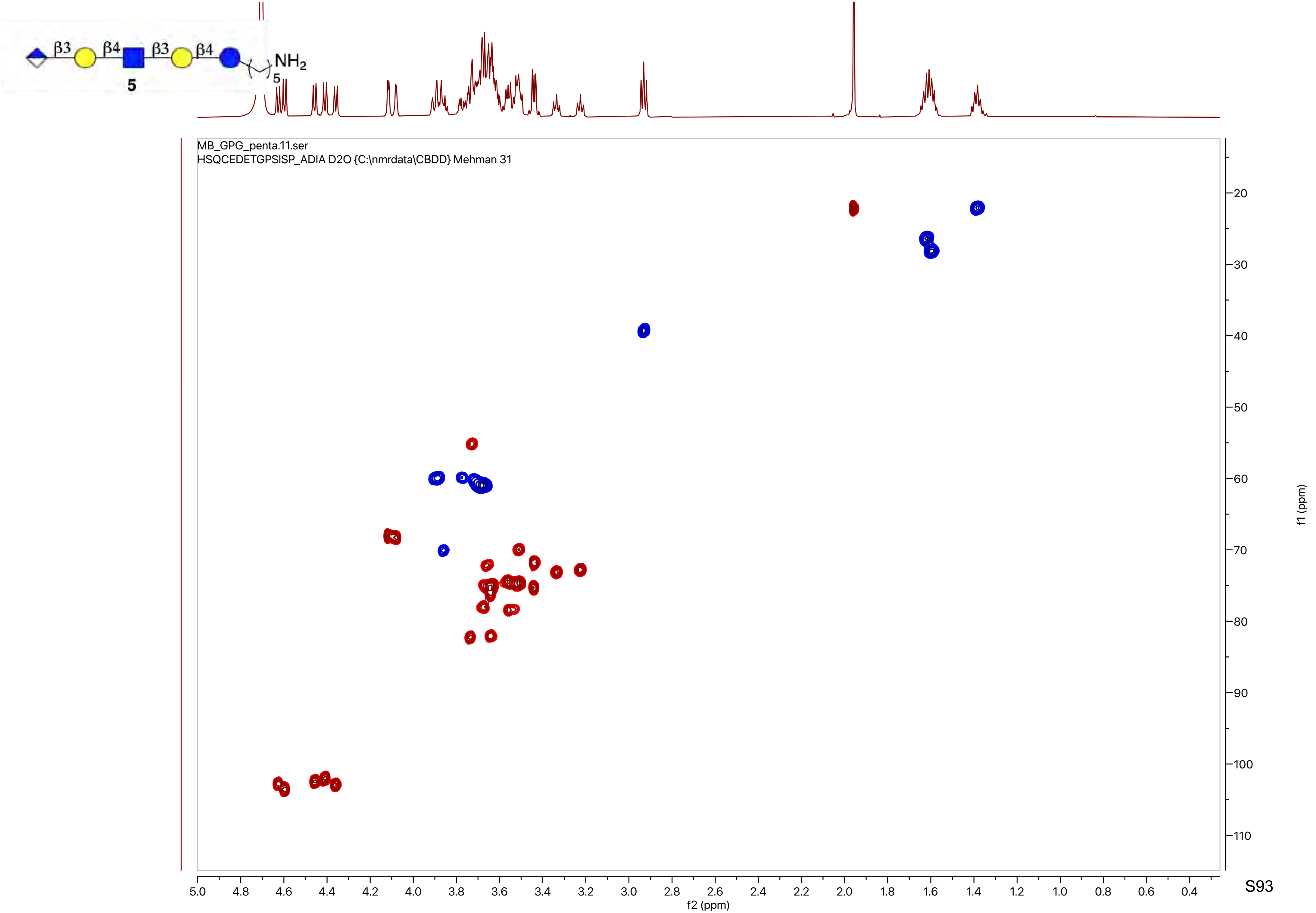

β3
β4
β3
β4
NH2
5
5
MB_GPG_penta.11.ser
HSQCEDETGPSISP_ADIA D2O {C:\nmrdata\CBDD} Mehman 31
f2 (ppm)
f1 (ppm)

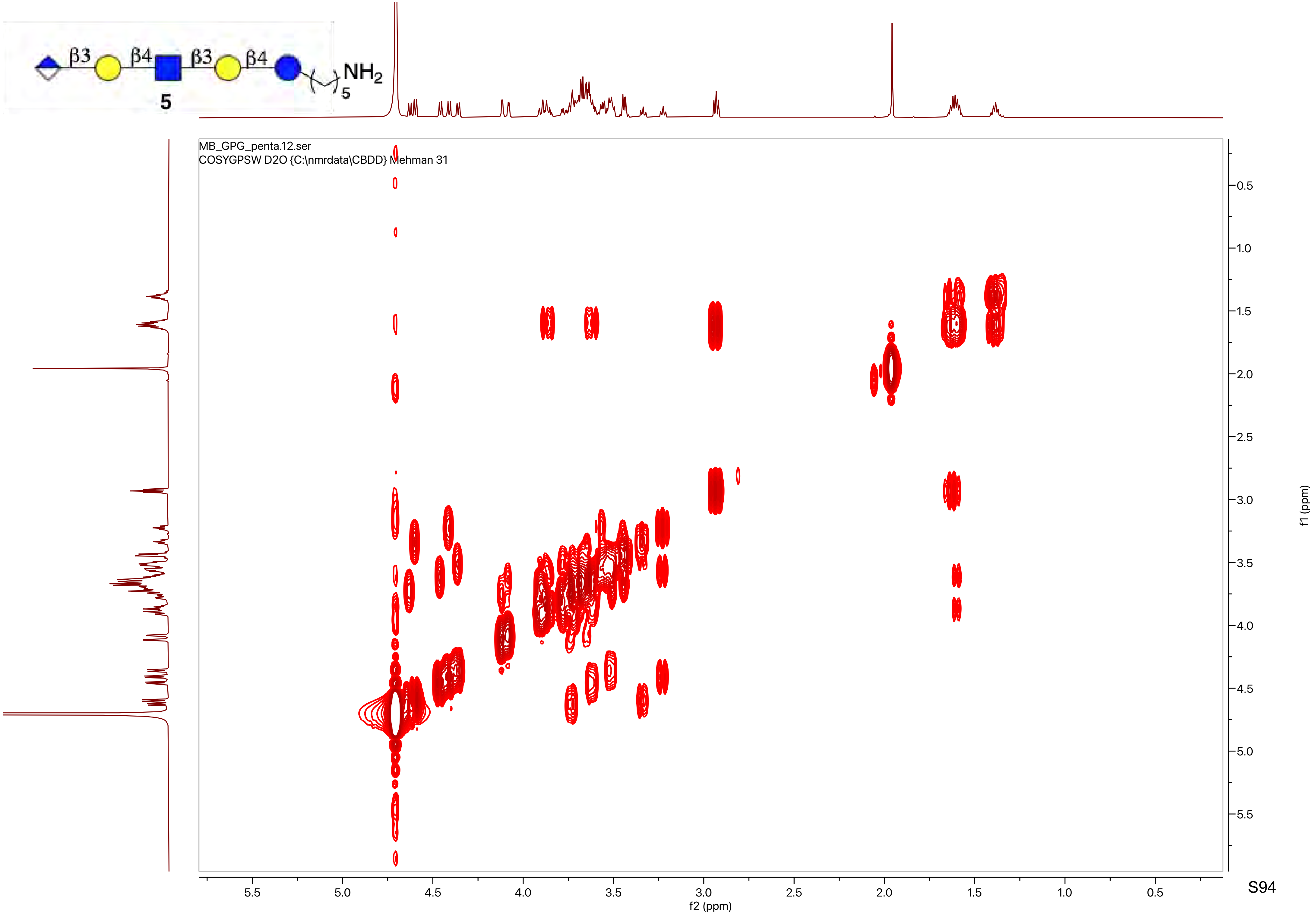

β3
β4
β3
β4
NH2
5
5
MB_GPG_penta.12.ser
COSYGPSW D2O {C:\nmrdata\CBDD} Mehman 31
f2 (ppm)
f1 (ppm)

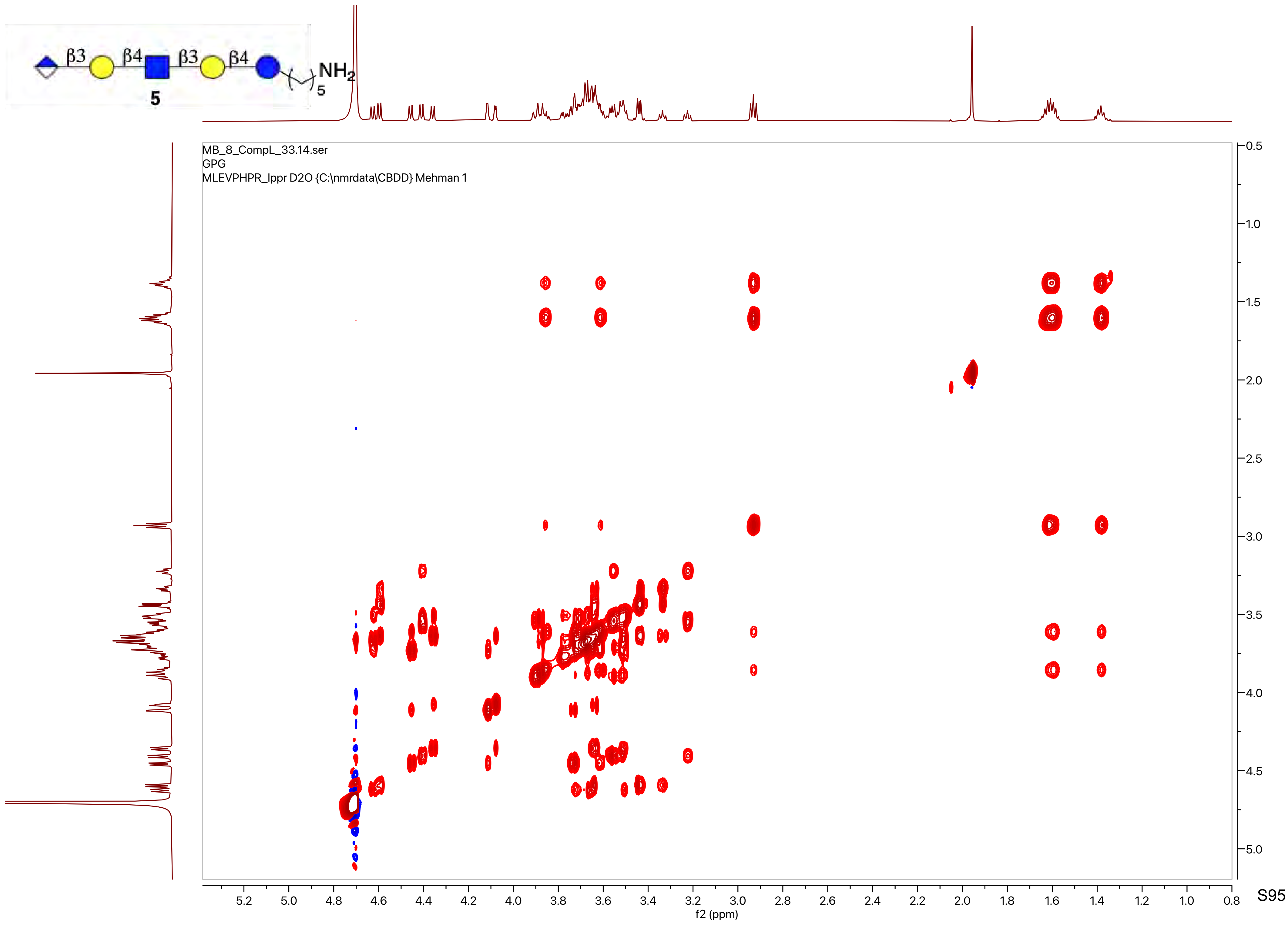
β3
β4
β3
β4
NH2
5
5
MB_8_CompL_33.14.ser
GPG
MLEVPHPR_lppr D2O {C:\nmrdata\CBDD} Mehman 1
f2 (ppm)
f1 (ppm)

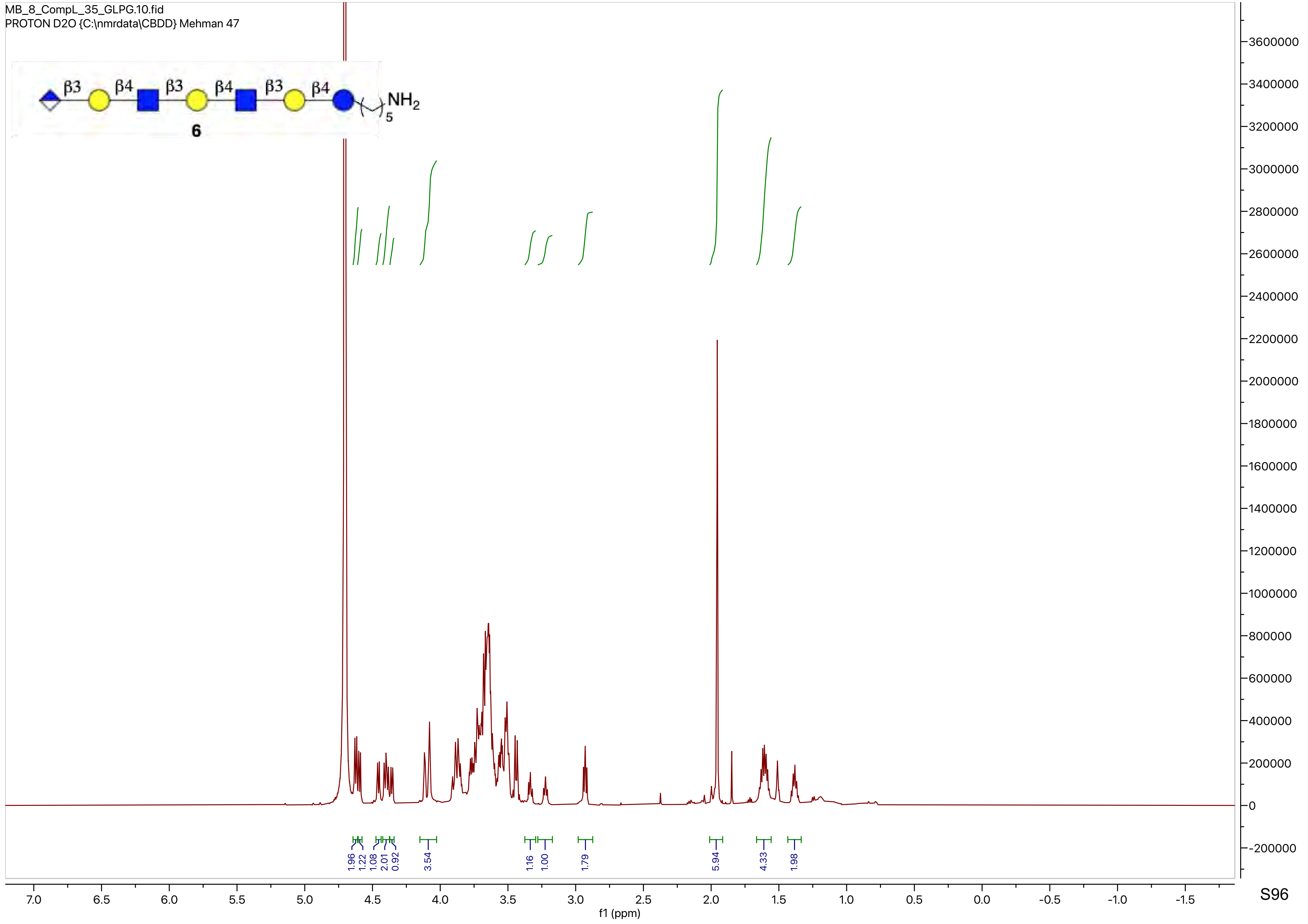
MB_8_CompL_35_GLPG.10.fid
PROTON D2O {C:\nmrdata\CBDD} Mehman 47
β3 β4 β3 β4 β3 β4
NH2
5
6
1.96 1.22 1.08 2.01 0.92 3.54 1.16 1.00 1.79 5.94 4.33 1.98
f1 (ppm)

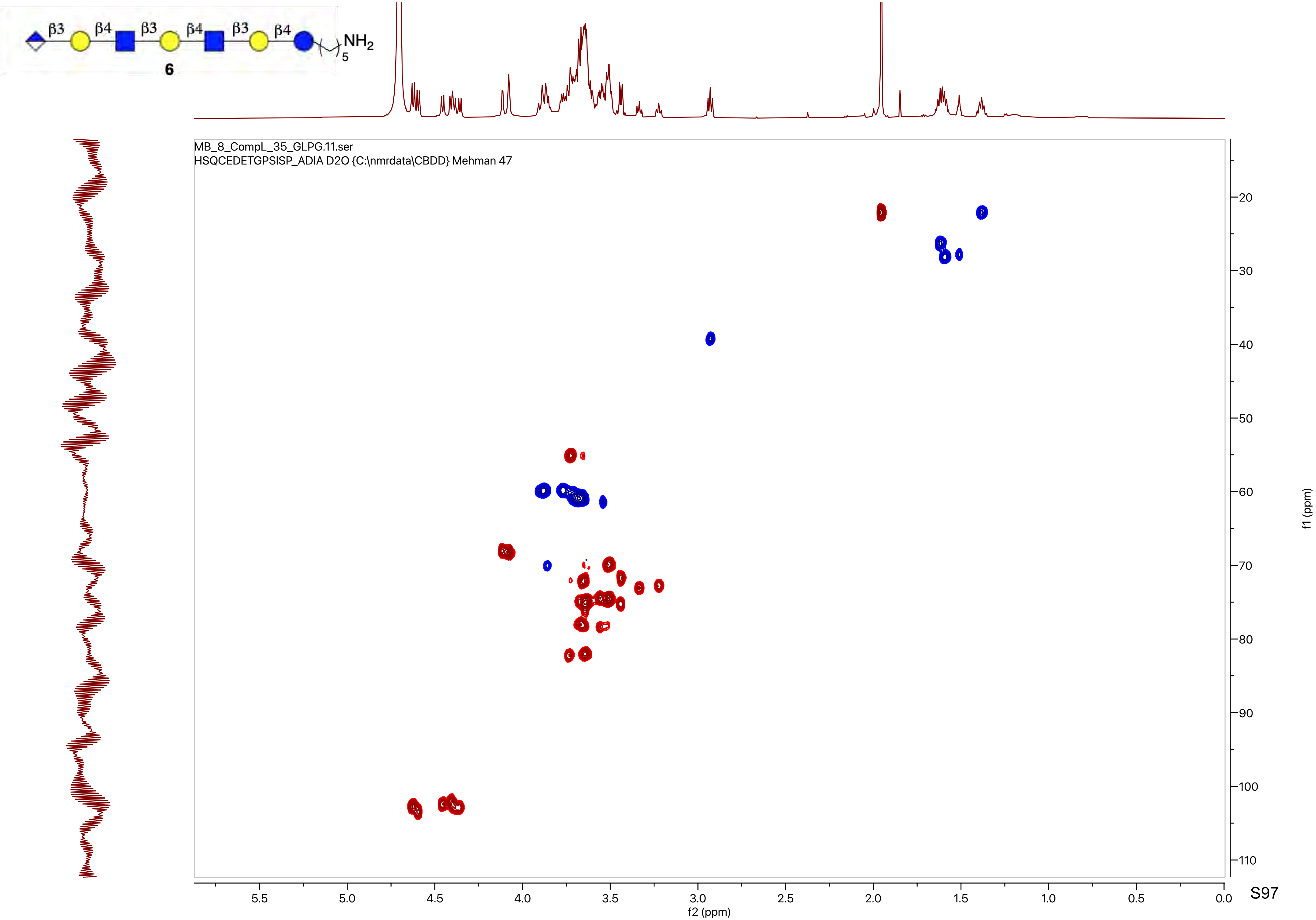
β3
β4
β3
β4
β3
β4
NH2
5
6
MB_8_CompL_35_GLPG.11.ser
HSQCEDETGPSISP_ADIA D2O {C:\nmrdata\CBDD} Mehman 47
f1 (ppm)
f2 (ppm)

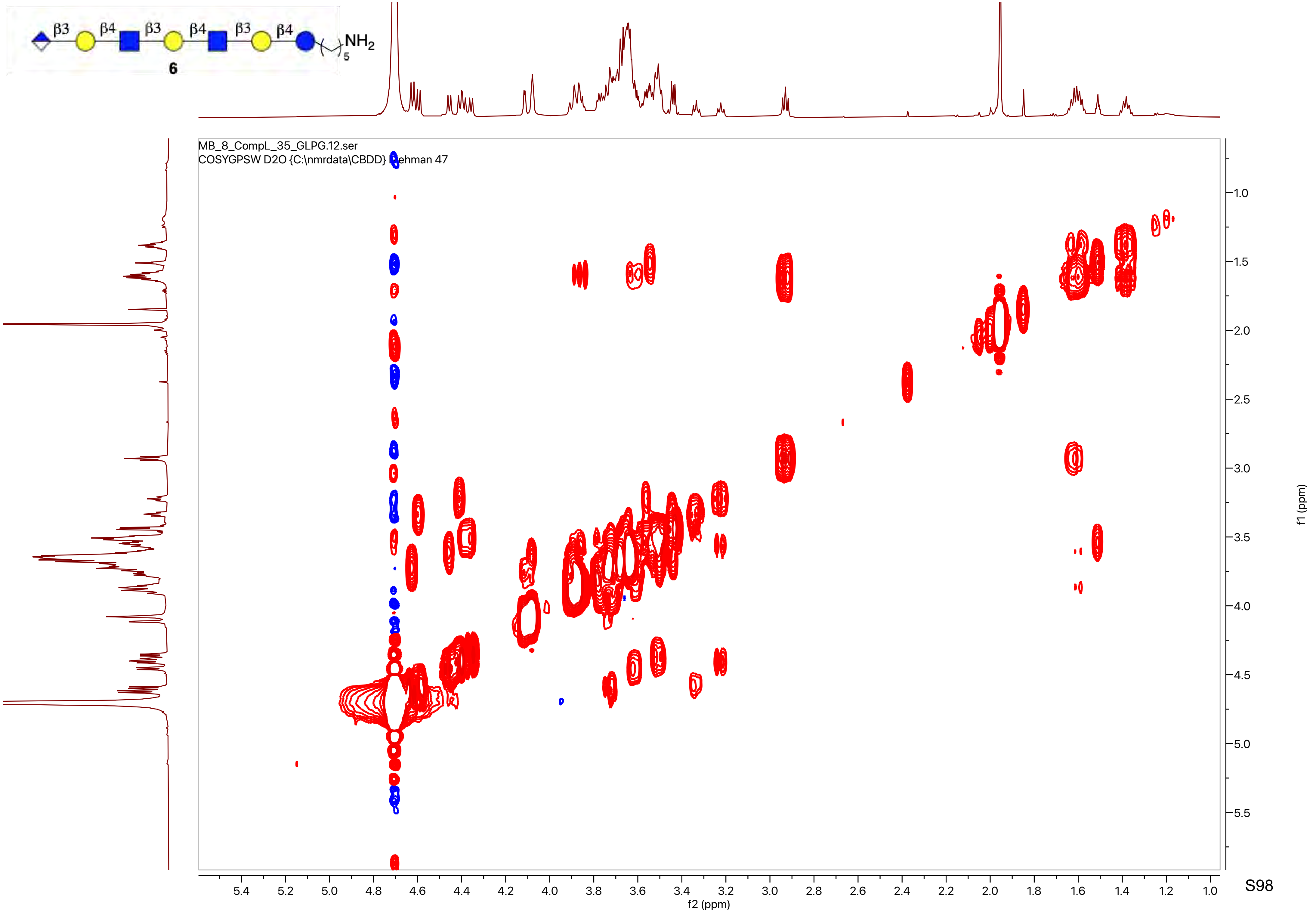
β3
β4
β3
β4
β3
β4
NH2
5
6
MB_8_CompL_35_GLPG.12.ser
COSYGPSW D2O {C:\nmrdata\CBDD} [illegible]ehman 47
f2 (ppm)
f1 (ppm)

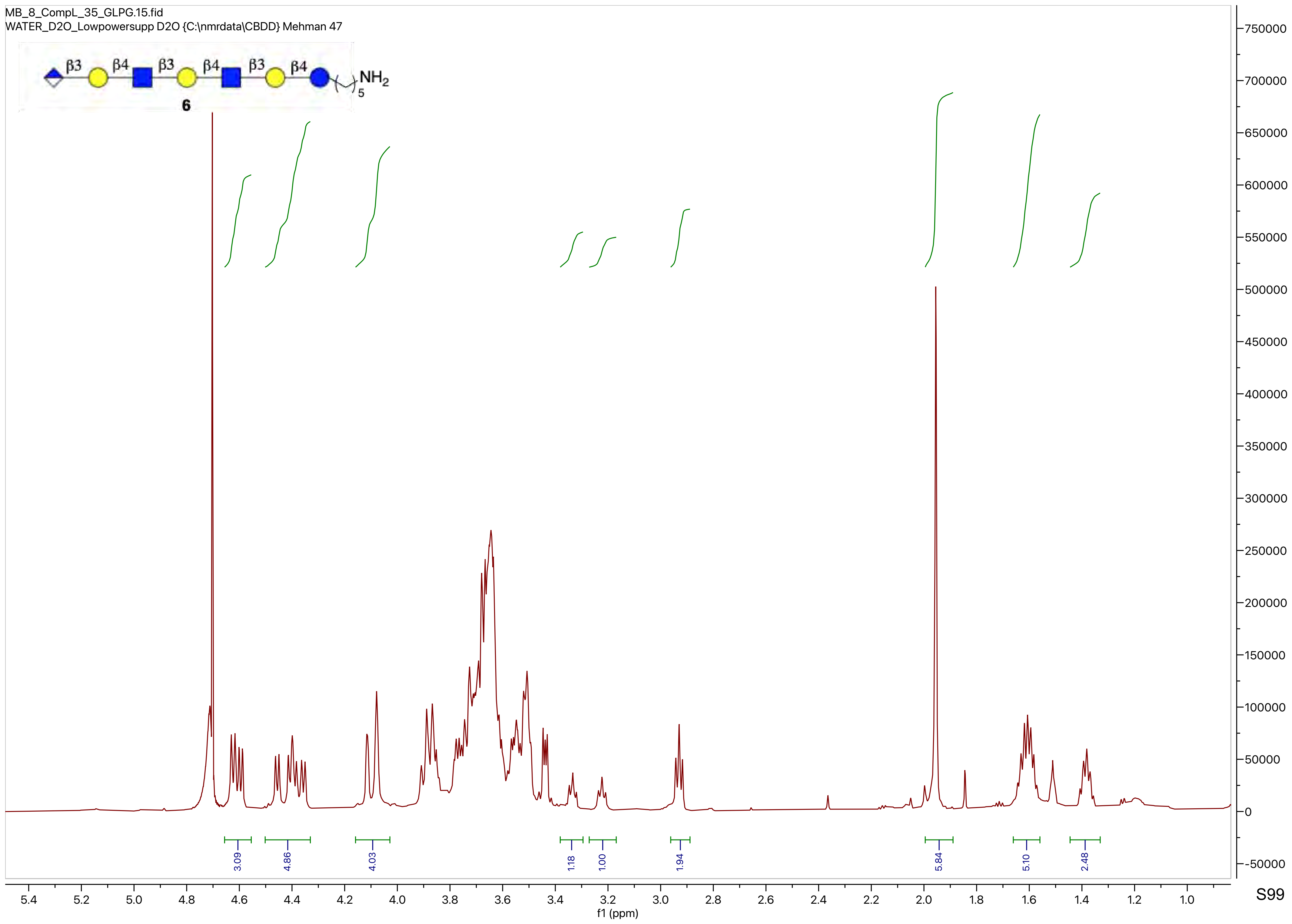

MB_8_CompL_35_GLPG.15.fid
WATER_D2O_Lowpowersupp D2O {C:\nmrdata\CBDD} Mehman 47
β3
β4
β3
β4
β3
β4
NH2
5
6
3.09
4.86
4.03
1.18
1.00
1.94
5.84
5.10
2.48
f1 (ppm)

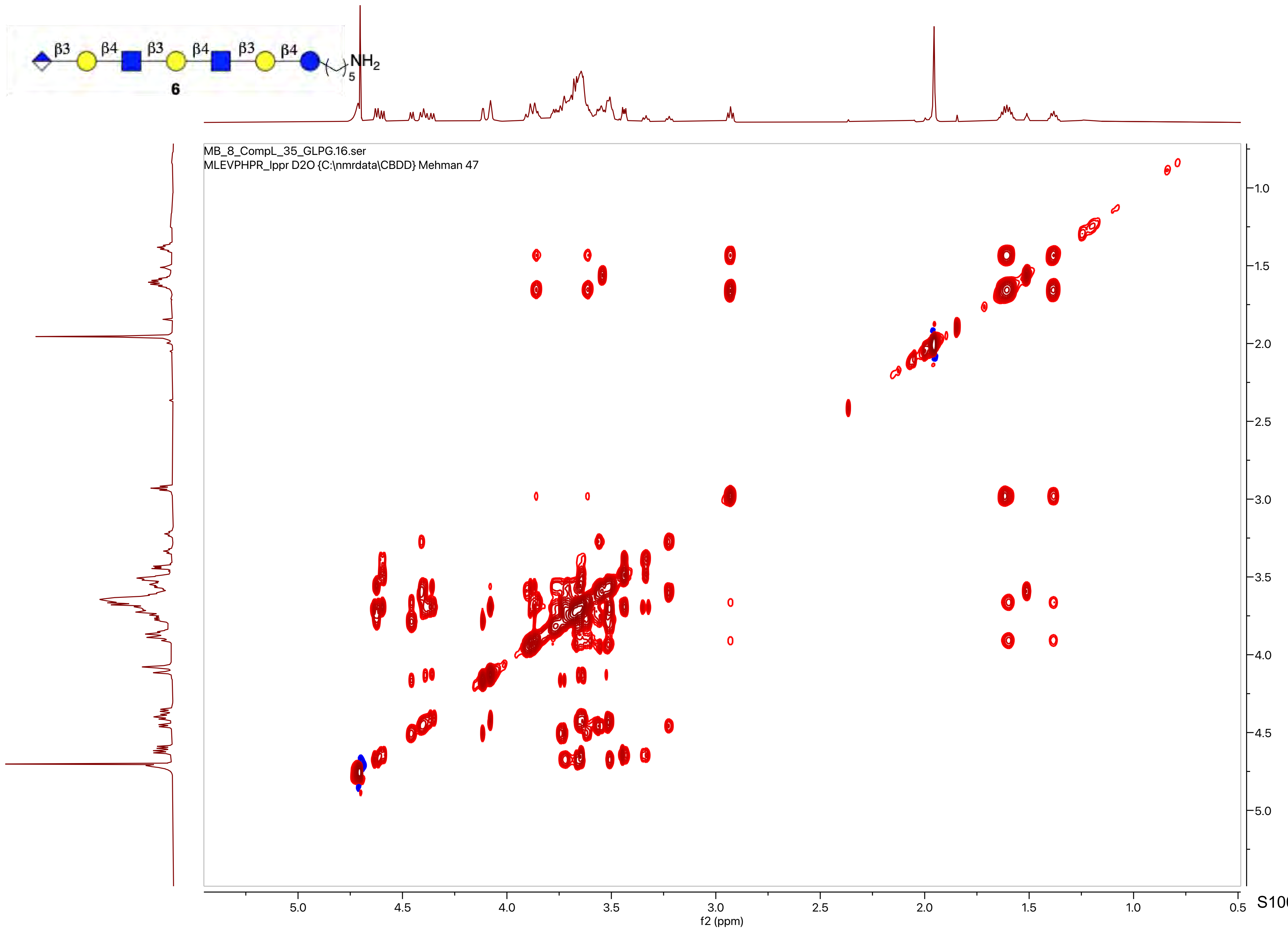
β3
β4
β3
β4
β3
β4
NH2
5
6
MB_8_CompL_35_GLPG.16.ser
MLEVPHPR_lppr D2O {C:\nmrdata\CBDD} Mehman 47
f2 (ppm)
f1 (ppm)

S100

MB_8_CompL_38_3SGlcParagloboside.15.fid

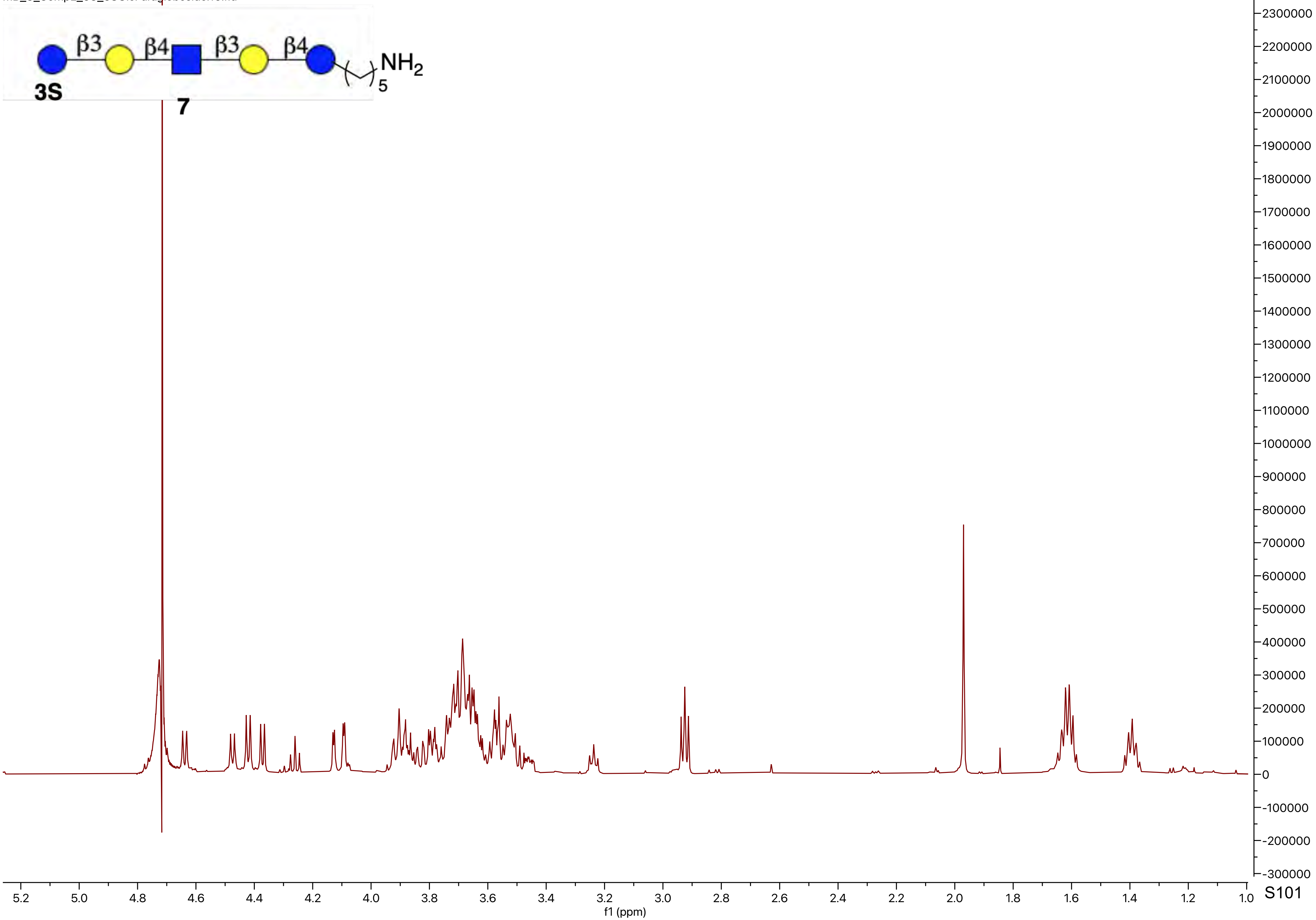

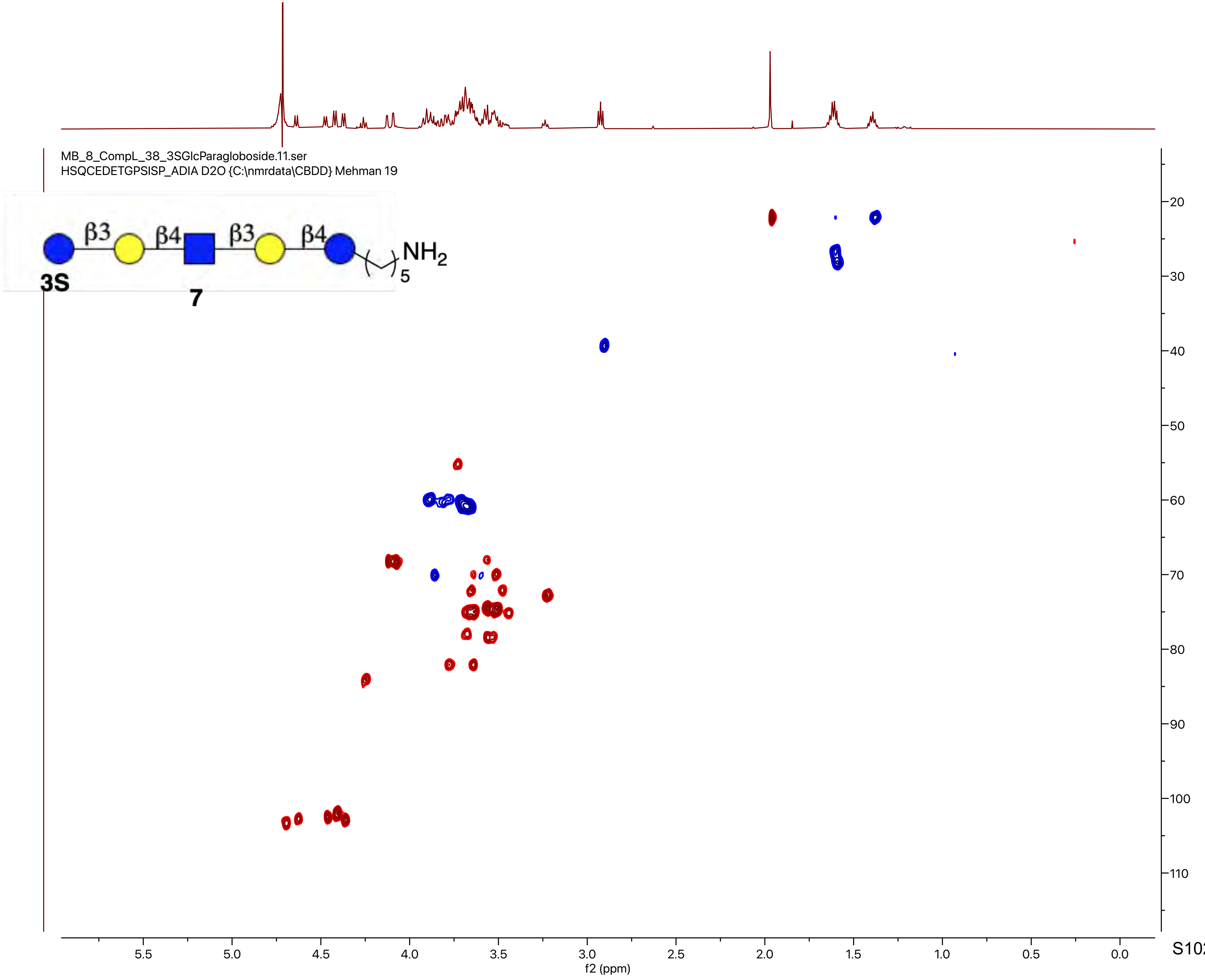

MB_8_CompL_38_3SGlcParagloboside.11.ser
HSQCEDETGPSISP_ADIA D2O {C:\nmrdata\CBDD} Mehman 19
β3
β4
β3
β4
NH2
5
3S
7
f2 (ppm)
f1 (ppm)

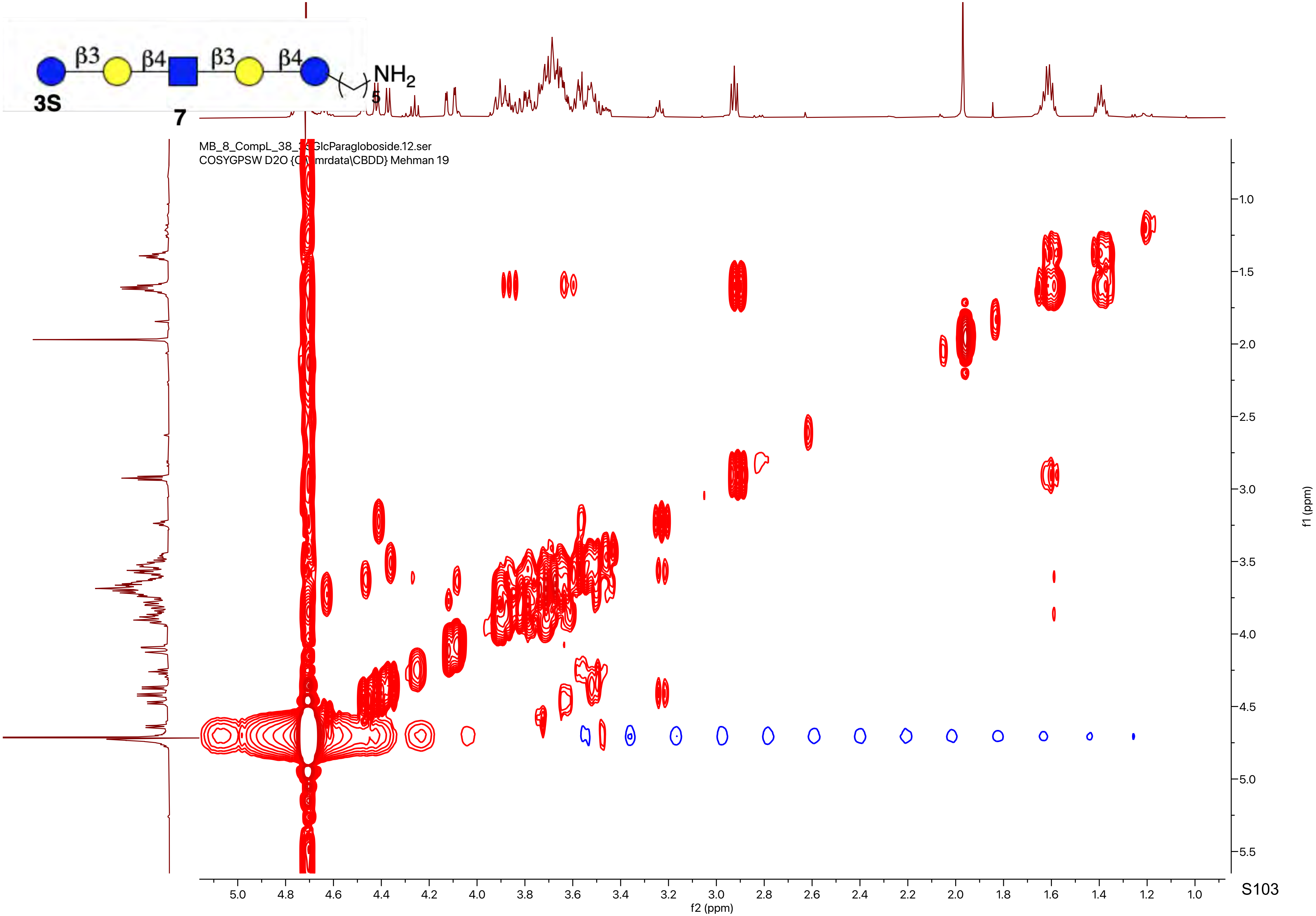

β3
β4
β3
β4
NH2
5
3S
7
GlcParagloboside.12.ser
COSYGPSW D2O
mrdata\CBDD} Mehman 19
f2 (ppm)
f1 (ppm)

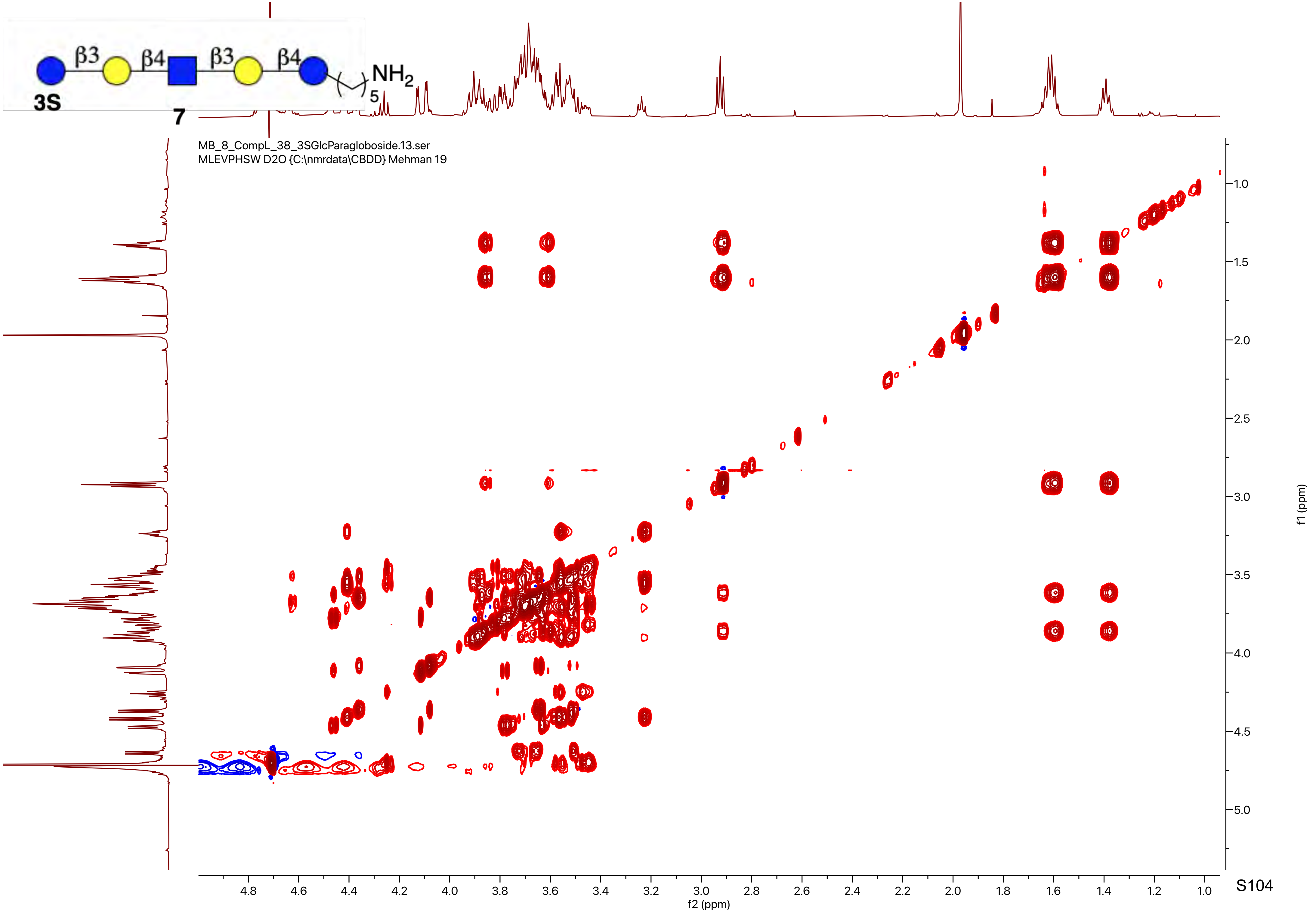
β3
β4
β3
β4
NH2
5
3S
7
MB_8_CompL_38_3SGlcParagloboside.13.ser
MLEVPHSW D2O {C:\nmrdata\CBDD} Mehman 19
f1 (ppm)
f2 (ppm)

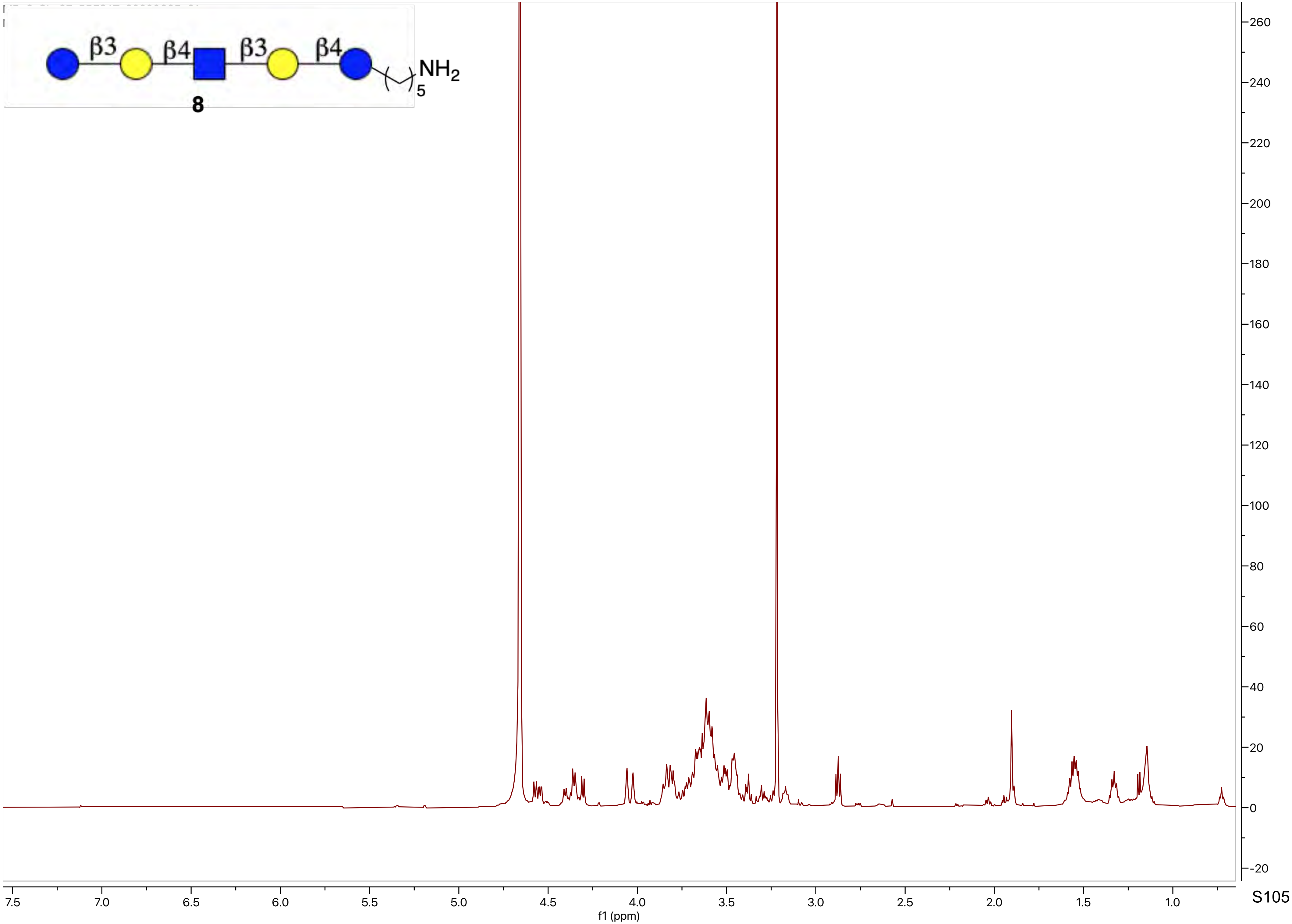
β3
β4
β3
β4
NH2
5
8
260
240
220
200
180
160
140
120
100
80
60
40
20
0
-20
7.5
7.0
6.5
6.0
5.5
5.0
4.5
4.0
3.5
3.0
2.5
2.0
1.5
1.0
f1 (ppm)

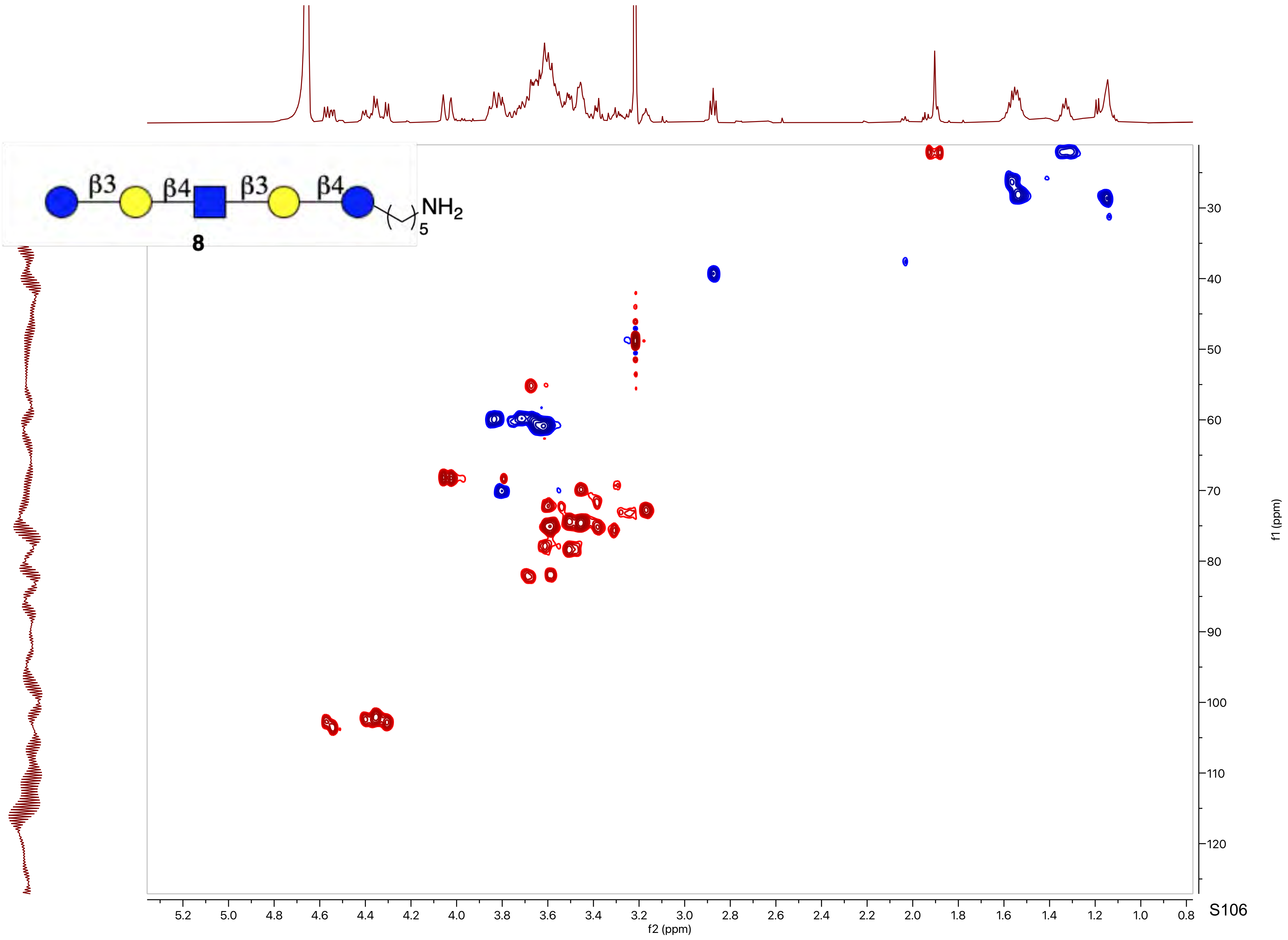
β3
β4
β3
β4
NH2
5
8
f1 (ppm)
f2 (ppm)
30
40
50
60
70
80
90
100
110
120
5.2
5.0
4.8
4.6
4.4
4.2
4.0
3.8
3.6
3.4
3.2
3.0
2.8
2.6
2.4
2.2
2.0
1.8
1.6
1.4
1.2
1.0
0.8

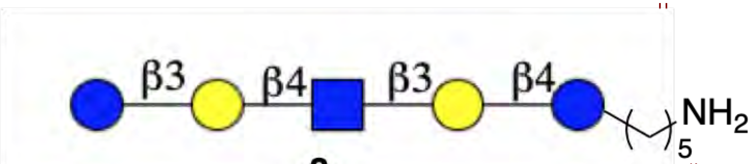
β3
β4
β3
β4
NH2
5

8

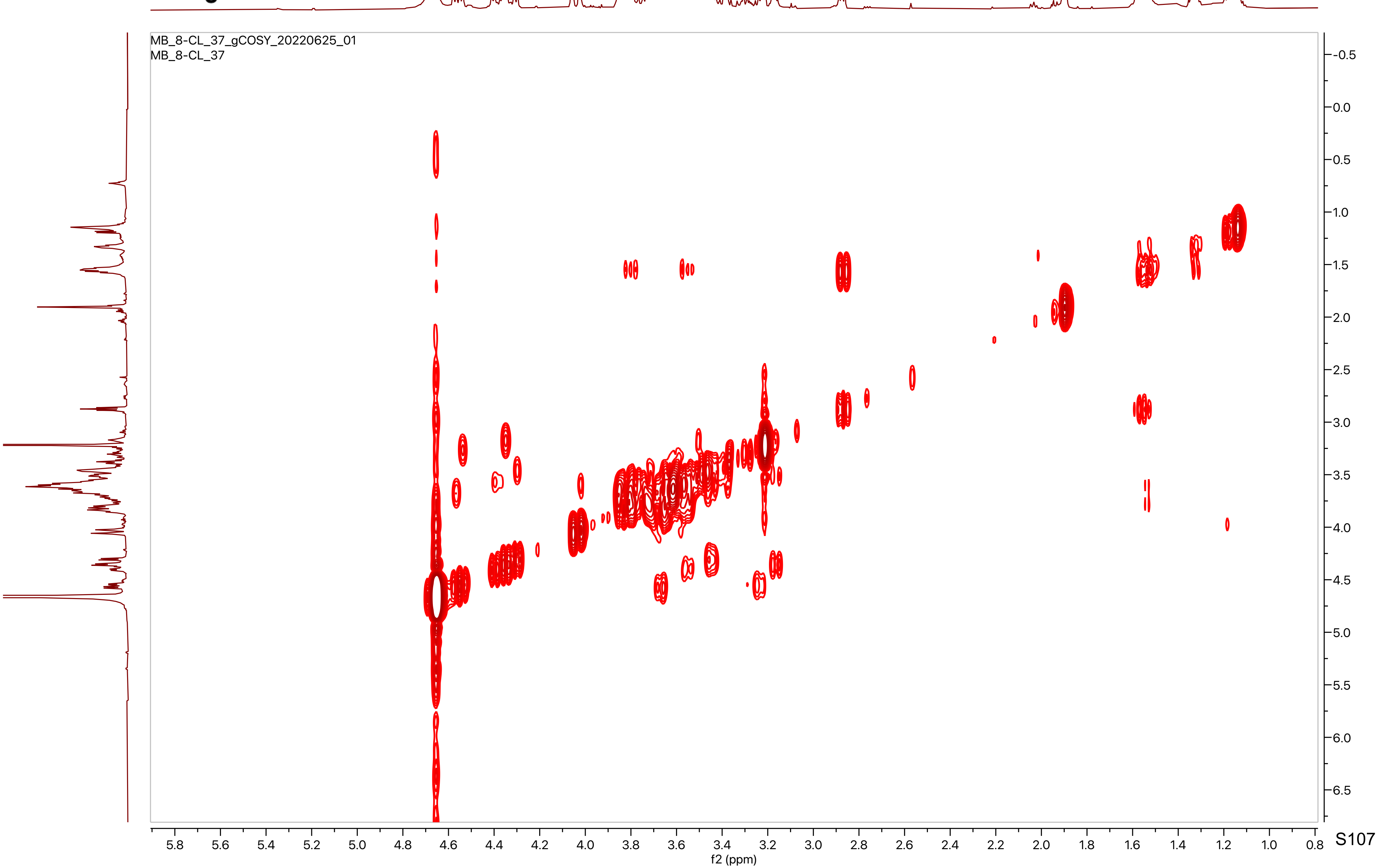
MB_8-CL_37_gCOSY_20220625_01
MB_8-CL_37
f2 (ppm)
f1 (ppm)

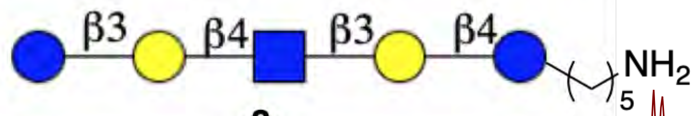
β3
β4
β3
β4
NH2
5
**8**

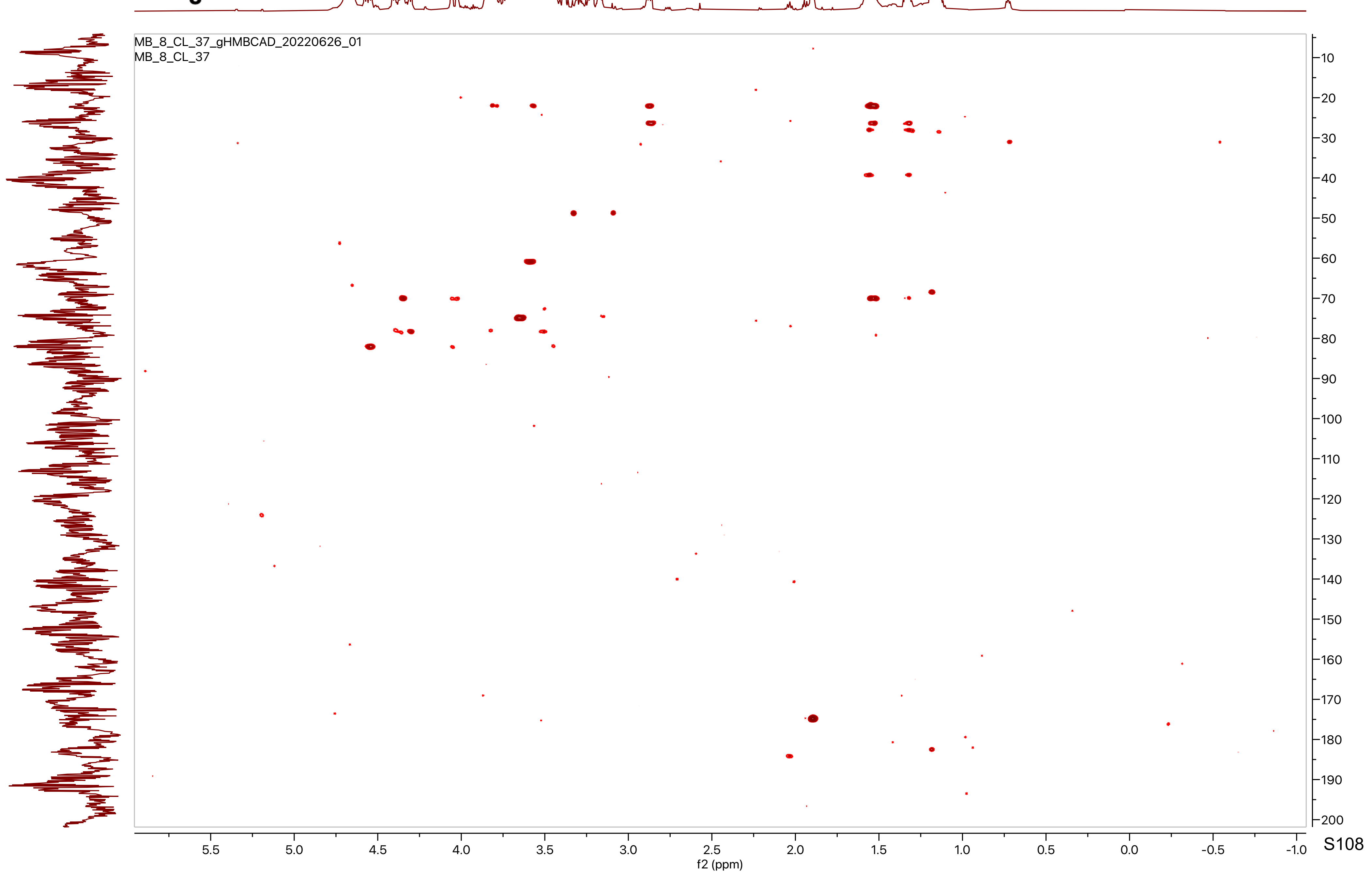
MB_8_CL_37_gHMBCAD_20220626_01
MB_8_CL_37
f2 (ppm)
f1 (ppm)

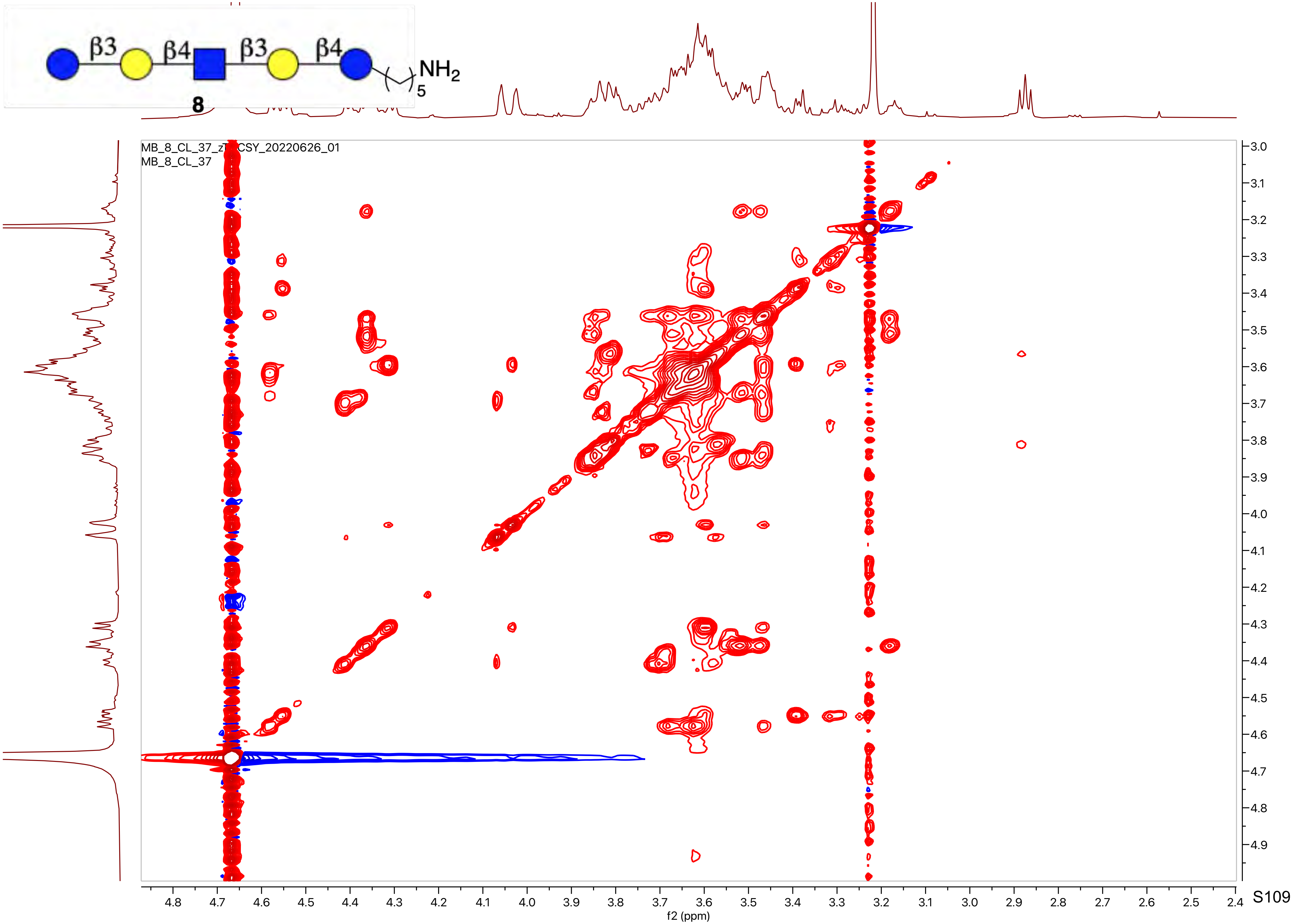
β3
β4
β3
β4
NH2
5
8
MB_8_CL_37_z[illegible]CSY_20220626_01
MB_8_CL_37
f2 (ppm)
f1 (ppm)

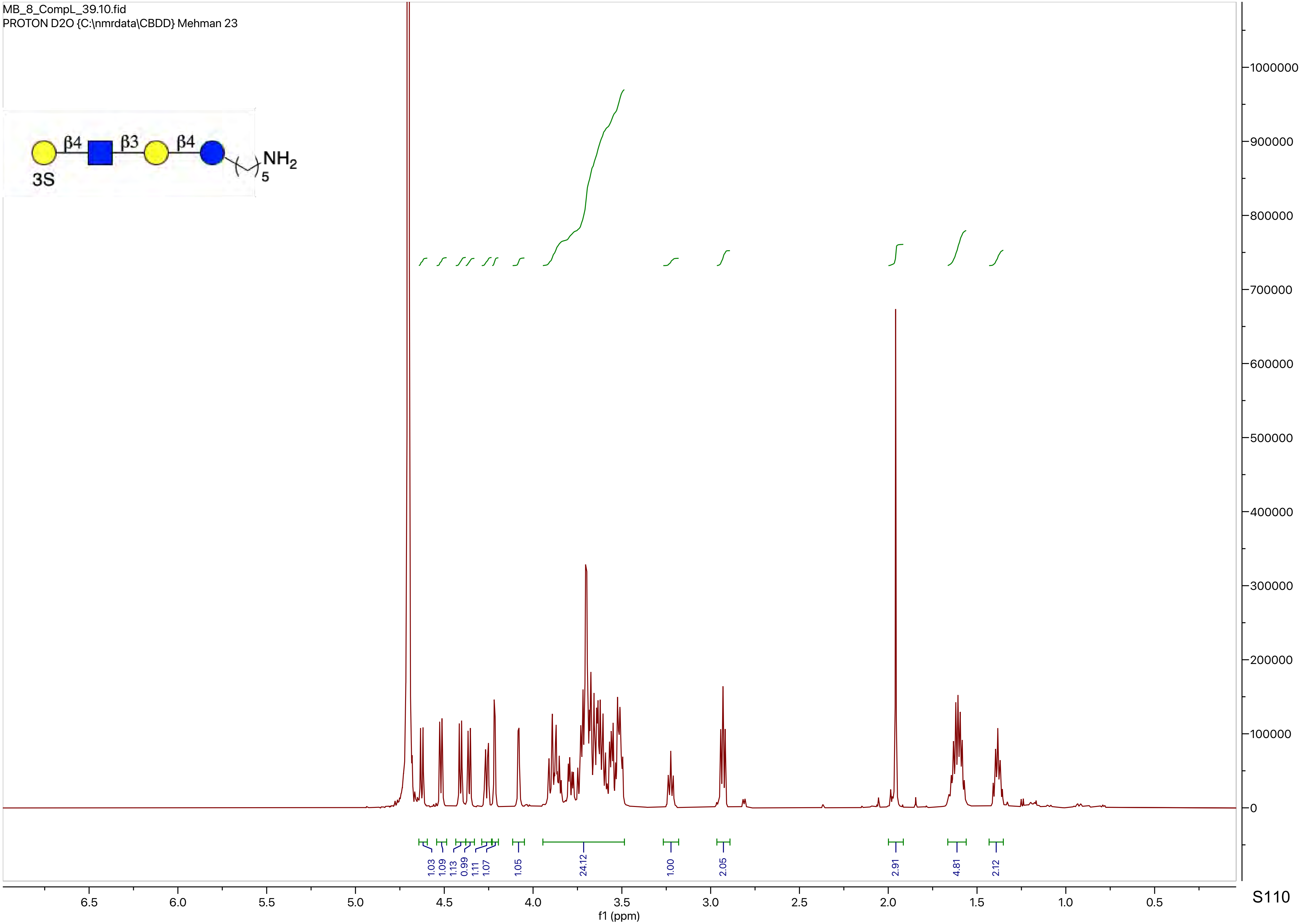
MB_8_CompL_39.10.fid
PROTON D2O {C:\nmrdata\CBDD} Mehman 23
β4
β3
β4
NH2
5
3S
1000000
900000
800000
700000
600000
500000
400000
300000
200000
100000
0
1.03
1.09
1.13
0.99
1.11
1.07
1.05
24.12
1.00
2.05
2.91
4.81
2.12
6.5
6.0
5.5
5.0
4.5
4.0
3.5
3.0
2.5
2.0
1.5
1.0
0.5
f1 (ppm)

MB_8_CompL_39.11.fid
WATER_D2O_Lowpowersupp D2O {C:\nmrdata\CBDD} Mehman 23

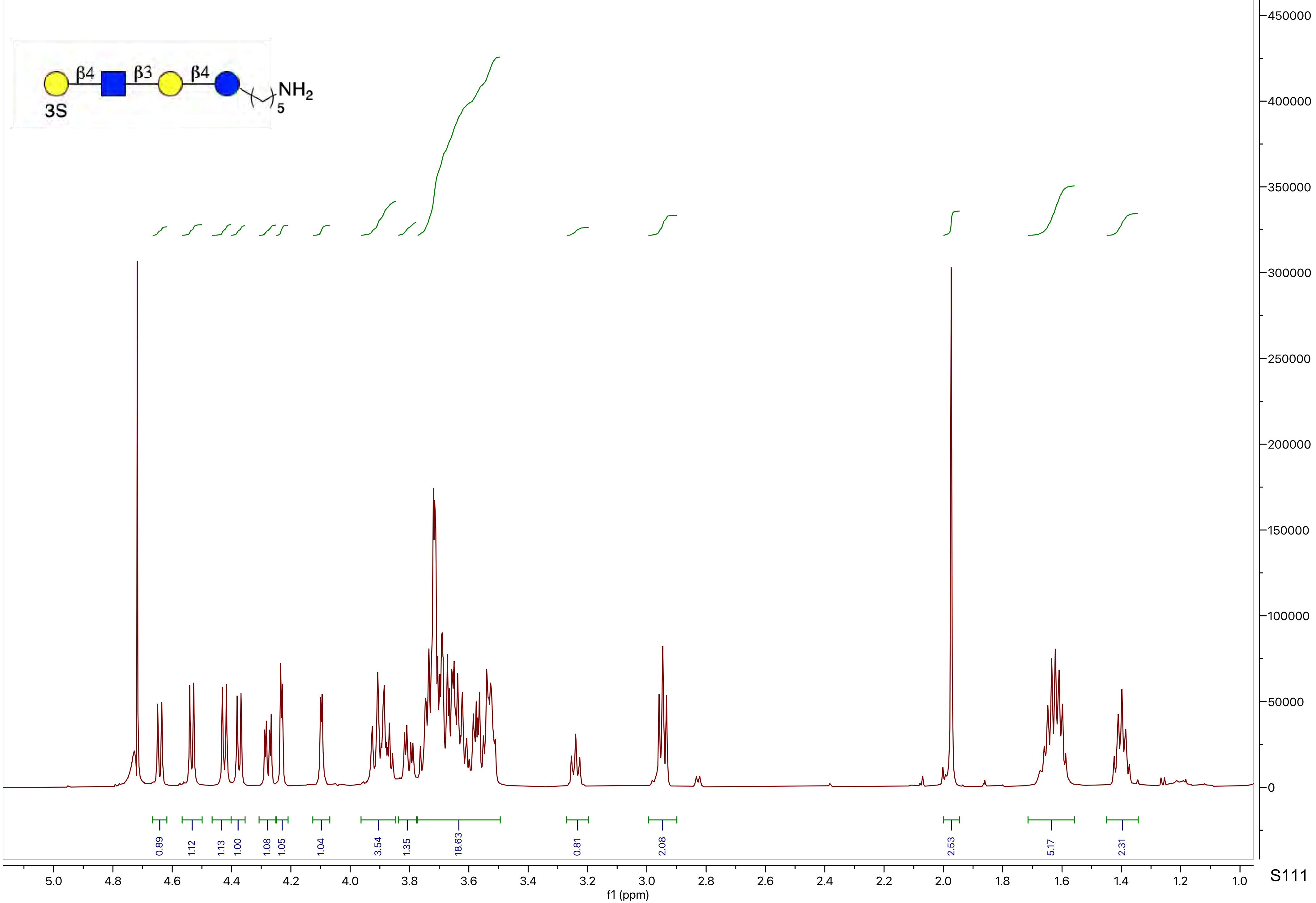

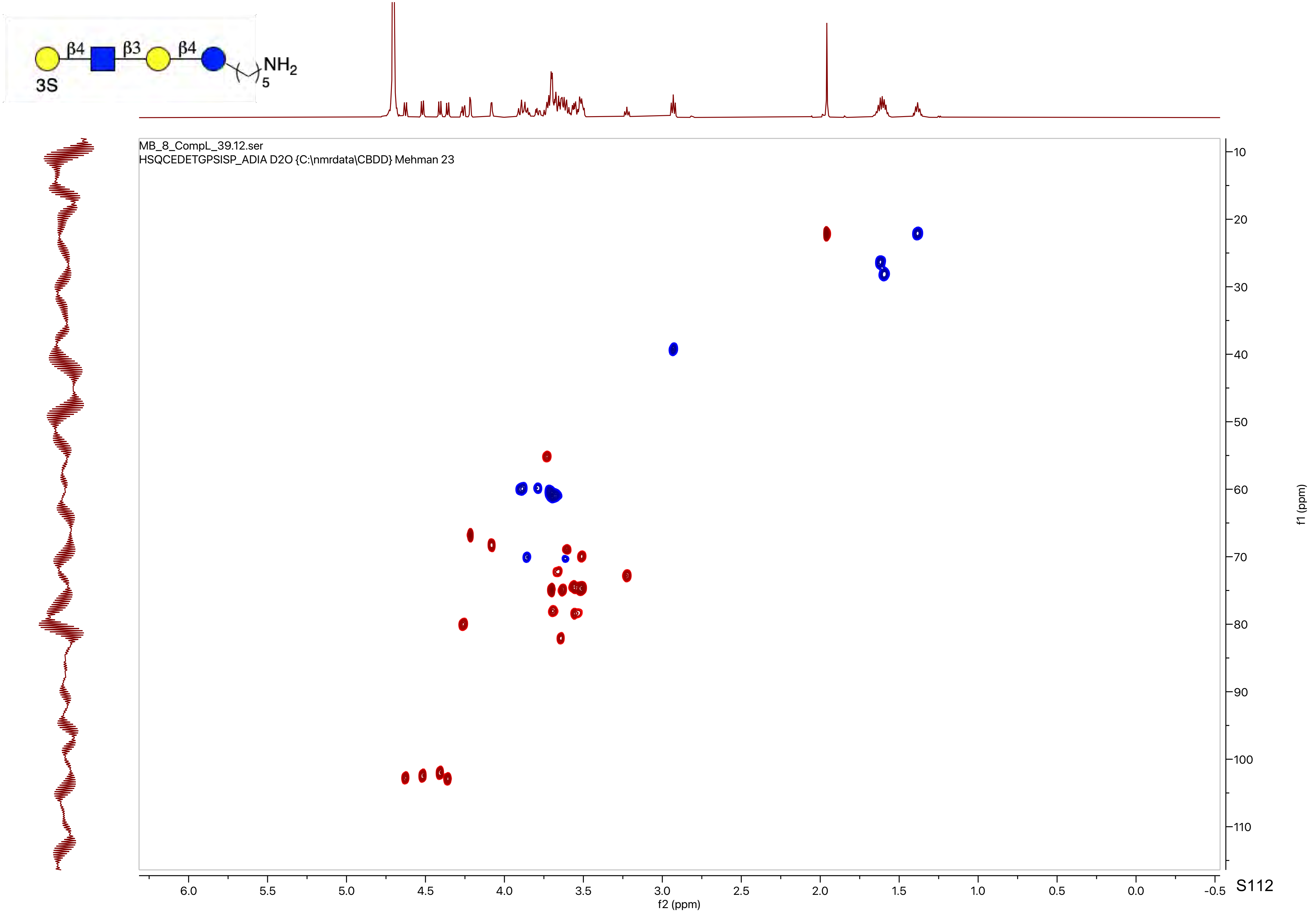

β4
β3
β4
3S
NH2
5
MB_8_CompL_39.12.ser
HSQCEDETGPSISP_ADIA D2O {C:\nmrdata\CBDD} Mehman 23
f2 (ppm)
f1 (ppm)

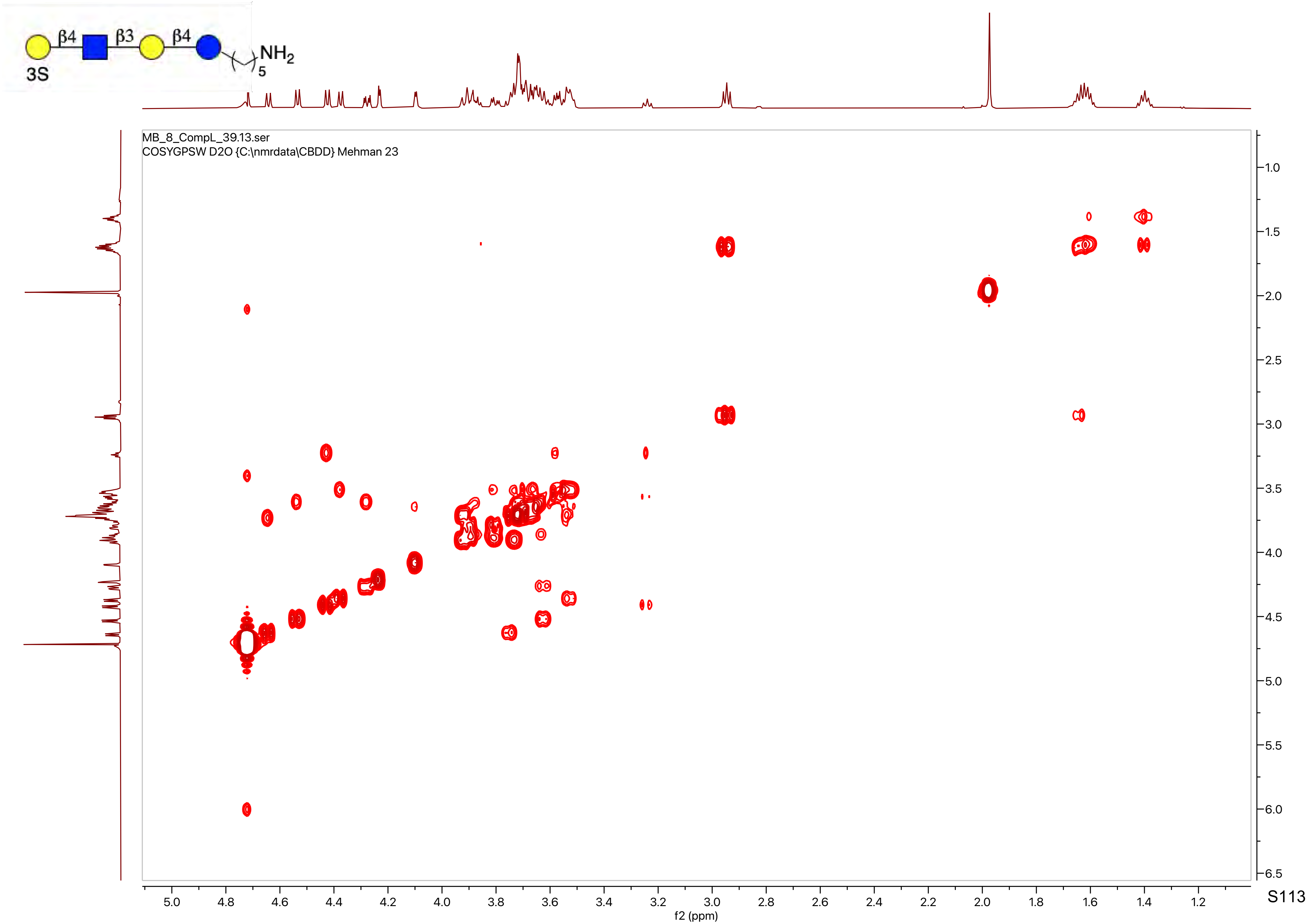
β4
β3
β4
NH2
5
3S
MB_8_CompL_39.13.ser
COSYGPSW D2O {C:\nmrdata\CBDD} Mehman 23
f2 (ppm)
f1 (ppm)

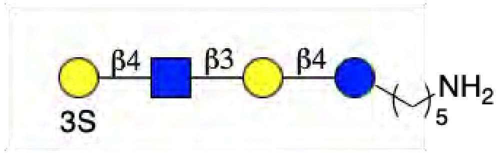
β4
β3
β4
NH2
5
3S

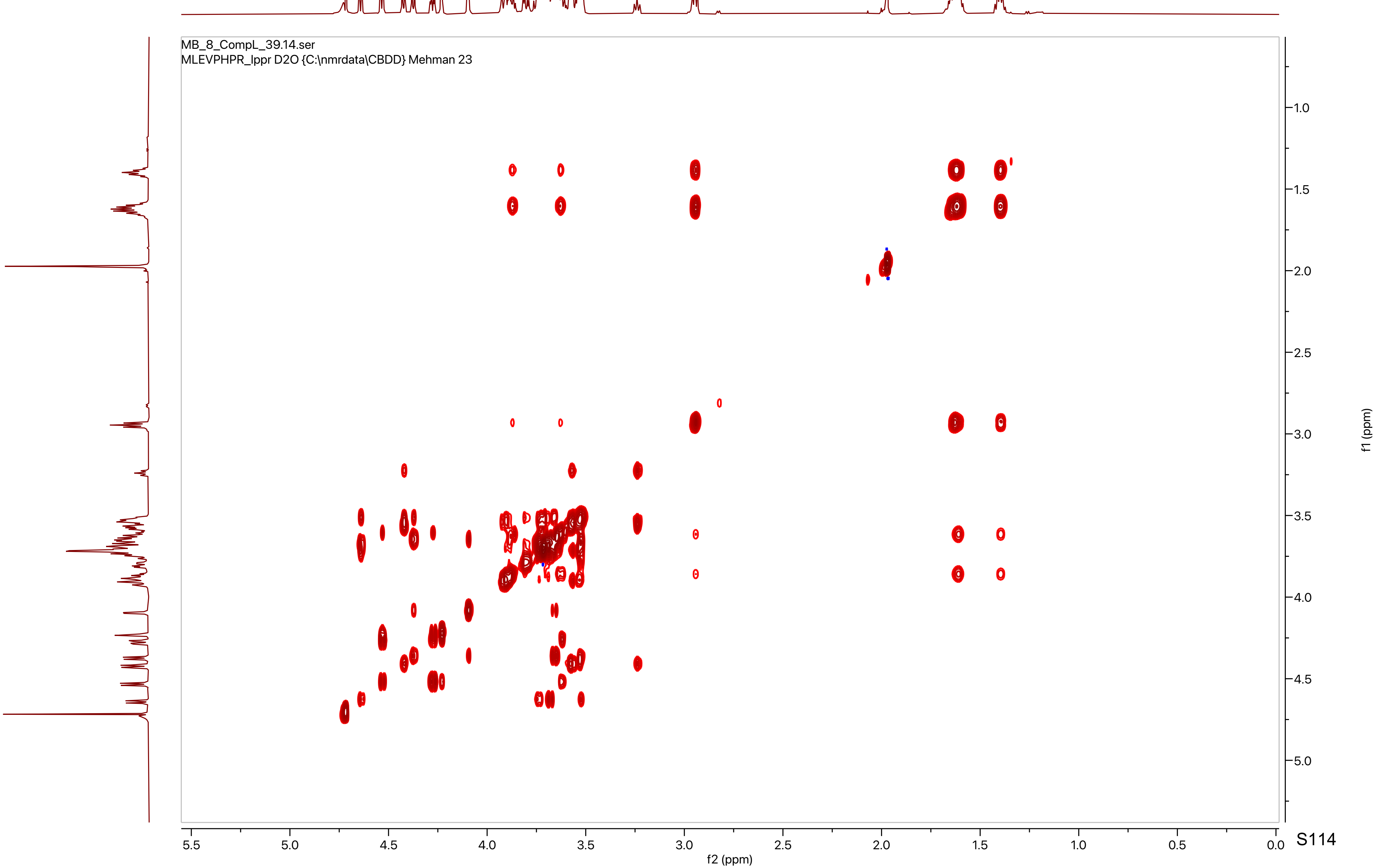
MB_8_CompL_39.14.ser
MLEVPHPR_lppr D2O {C:\nmrdata\CBDD} Mehman 23
f2 (ppm)
f1 (ppm)

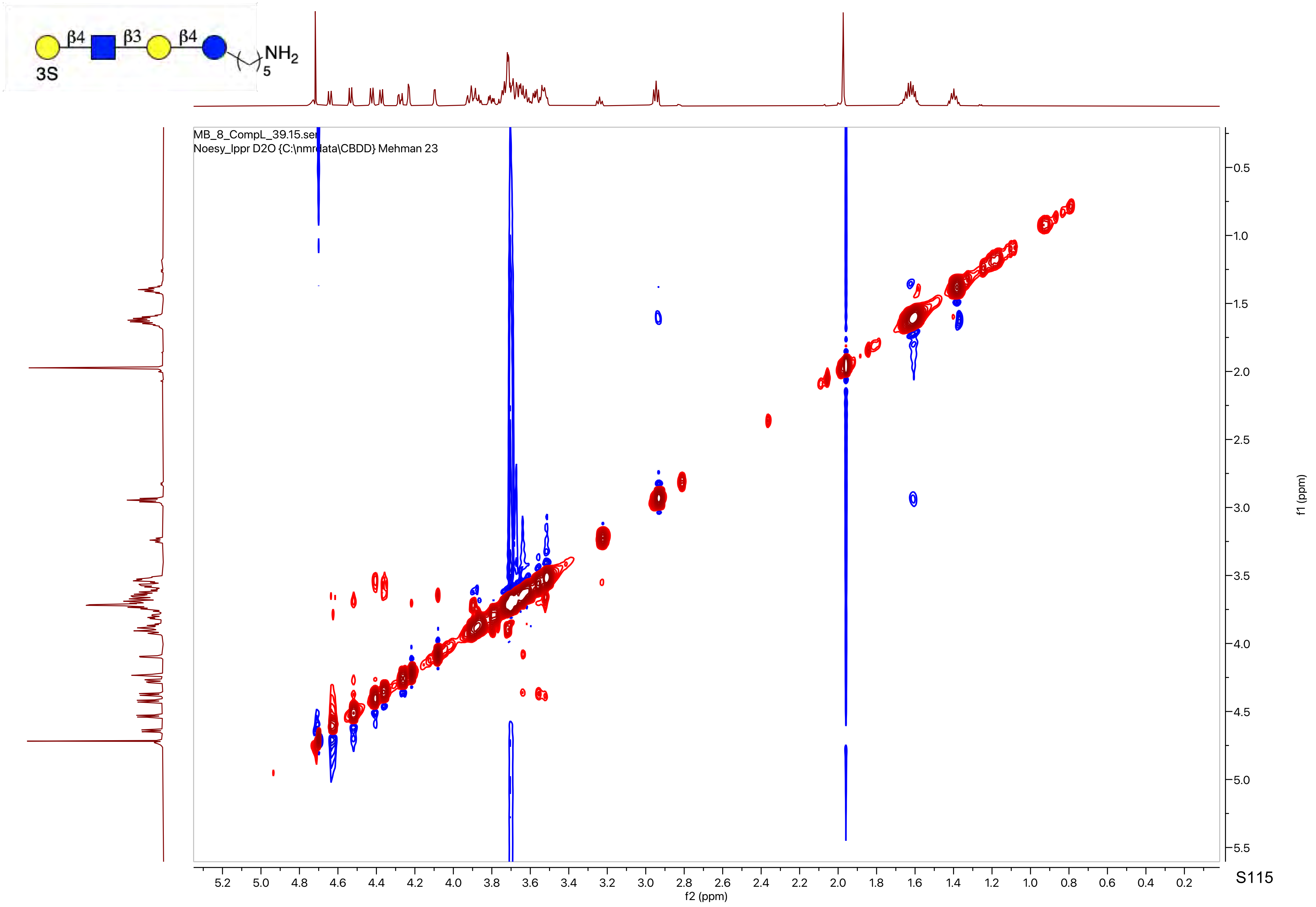
β4
β3
β4
3S
NH2
5
MB_8_CompL_39.15.ser
Noesy_lppr D2O {C:\nmrdata\CBDD} Mehman 23
f2 (ppm)
f1 (ppm)

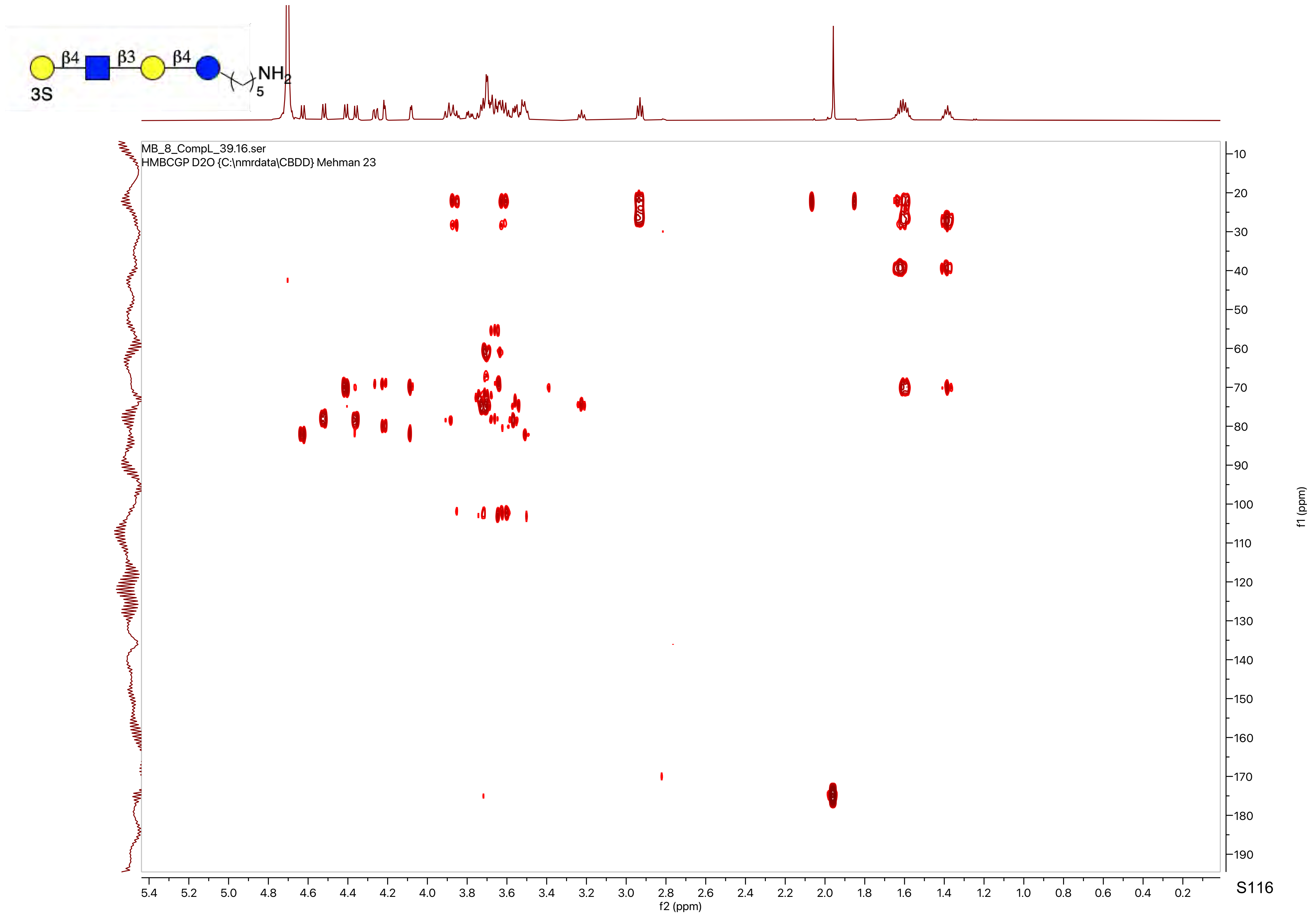
β4
β3
β4
NH2
5
3S
MB_8_CompL_39.16.ser
HMBCGP D2O {C:\nmrdata\CBDD} Mehman 23
f2 (ppm)
f1 (ppm)

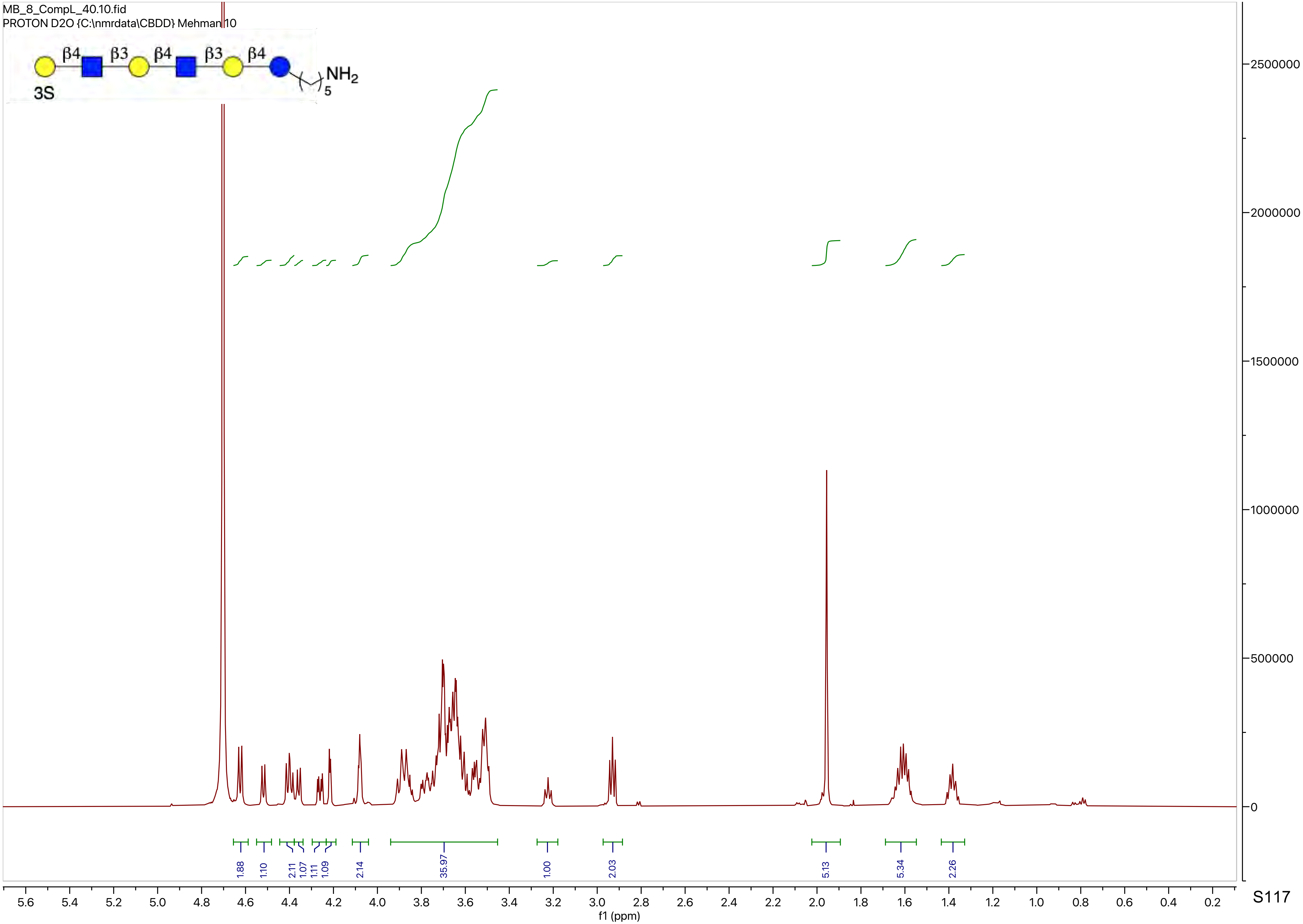

MB_8_CompL_40.10.fid
PROTON D2O {C:\nmrdata\CBDD} Mehman 10
β4 β3 β4 β3 β4
3S
NH2
5
1.88
1.10
2.11
1.07
1.11
1.09
2.14
35.97
1.00
2.03
5.13
5.34
2.26
2500000
2000000
1500000
1000000
500000
0
5.6 5.4 5.2 5.0 4.8 4.6 4.4 4.2 4.0 3.8 3.6 3.4 3.2 3.0 2.8 2.6 2.4 2.2 2.0 1.8 1.6 1.4 1.2 1.0 0.8 0.6 0.4 0.2
f1 (ppm)

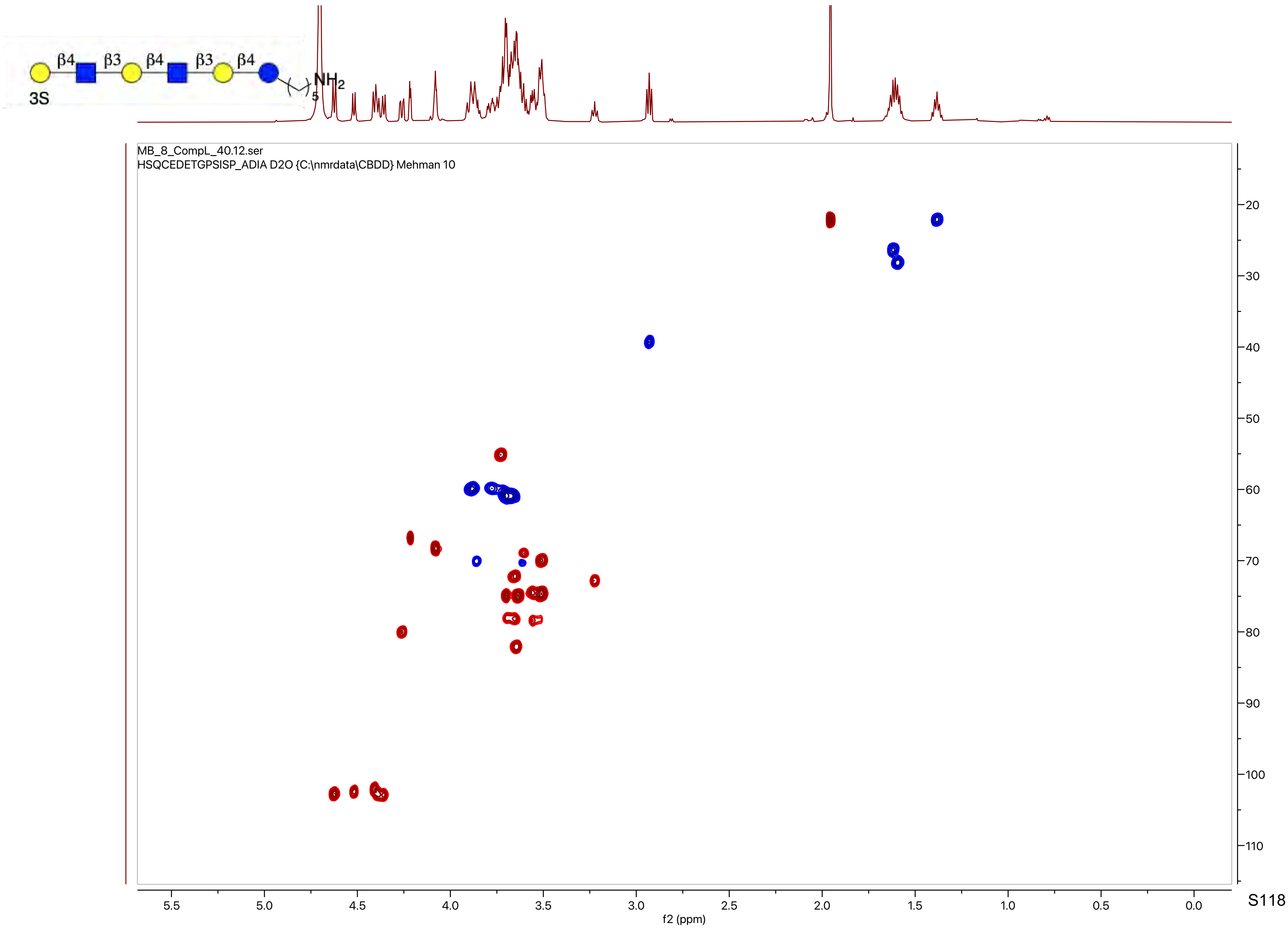
β4
β3
β4
β3
β4
NH2
5
3S
MB_8_CompL_40.12.ser
HSQCEDETGPSISP_ADIA D2O {C:\nmrdata\CBDD} Mehman 10
f2 (ppm)
f1 (ppm)

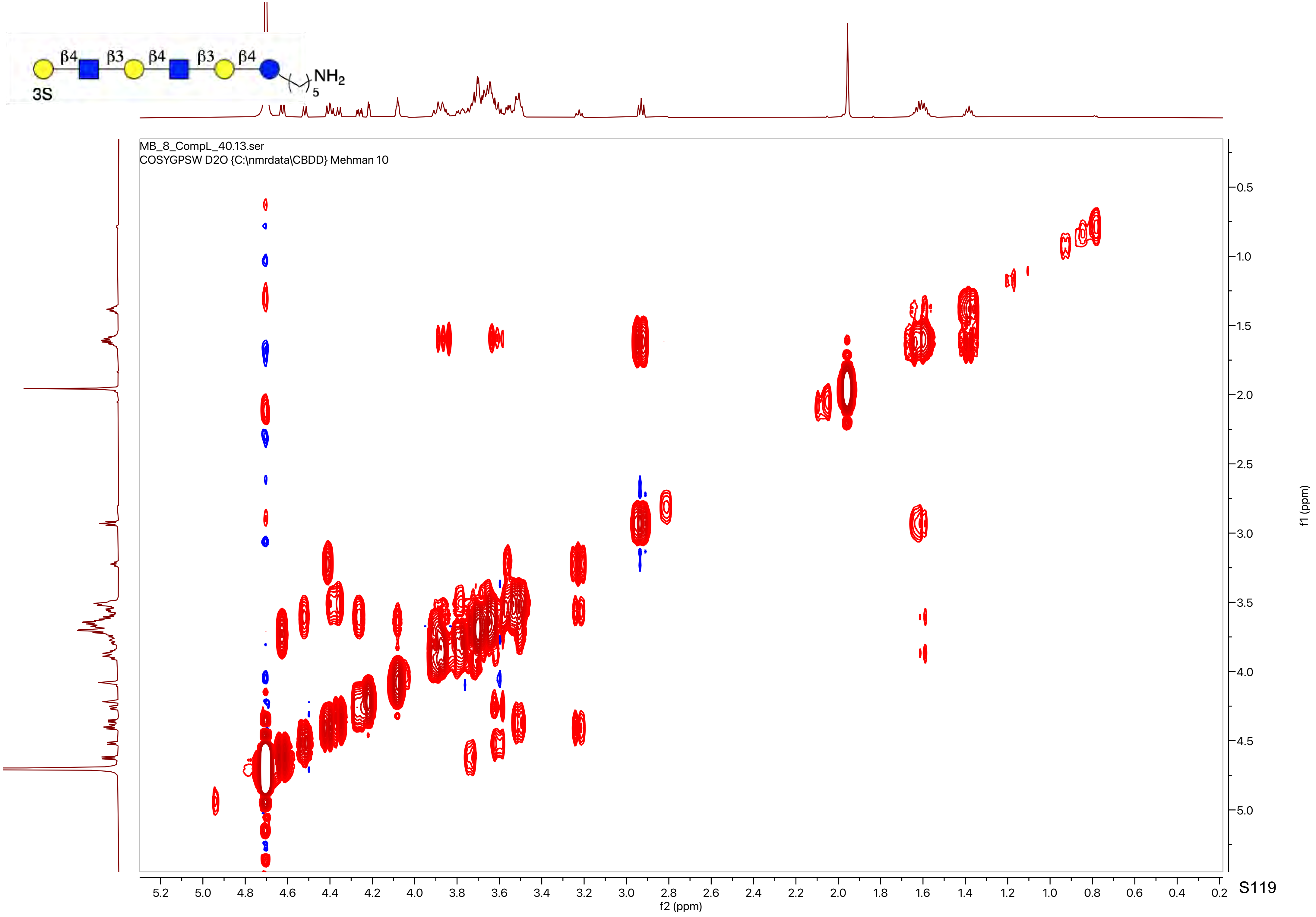

β4
β3
β4
β3
β4
$NH_2$
5
3S
MB_8_CompL_40.13.ser
COSYGPSW D2O {C:\nmrdata\CBDD} Mehman 10
f2 (ppm)
f1 (ppm)

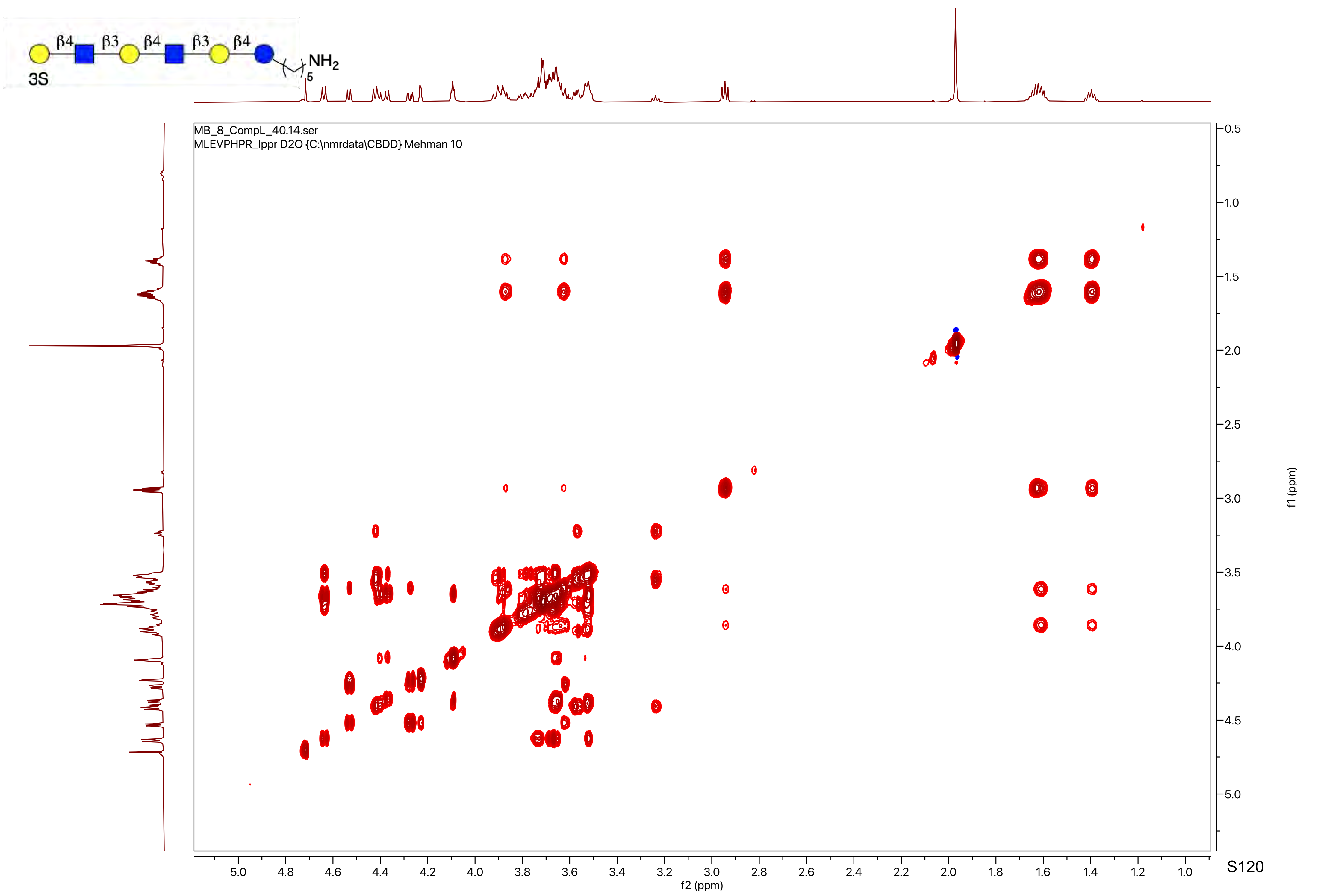

β4
β3
β4
β3
β4
NH2
5
3S
MB_8_CompL_40.14.ser
MLEVPHPR_lppr D2O {C:\nmrdata\CBDD} Mehman 10
f2 (ppm)
f1 (ppm)

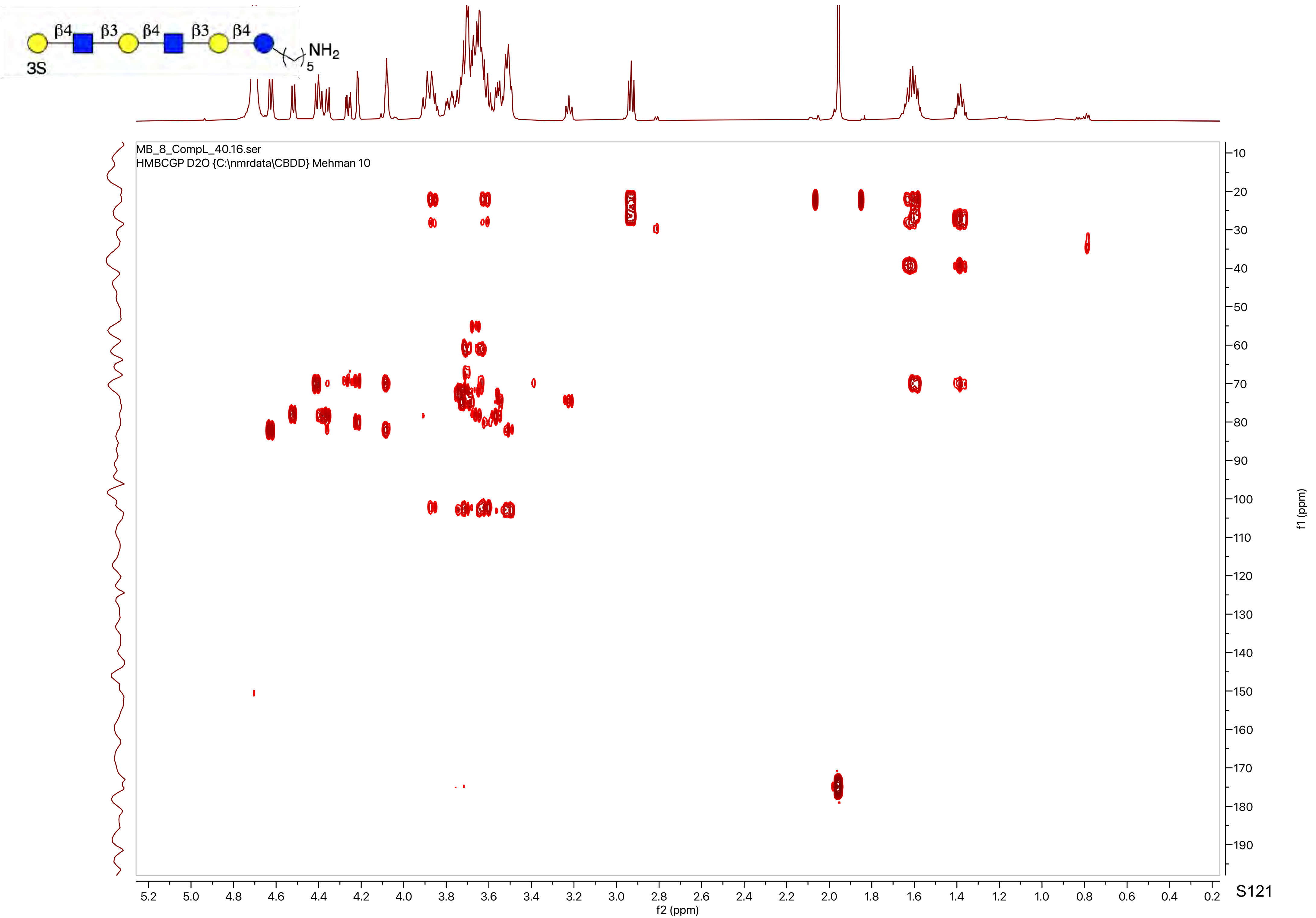

3S
β4
β3
β4
β3
β4
$NH_2$
5
MB_8_CompL_40.16.ser
HMBCGP D2O {C:\nmrdata\CBDD} Mehman 10
f2 (ppm)
f1 (ppm)

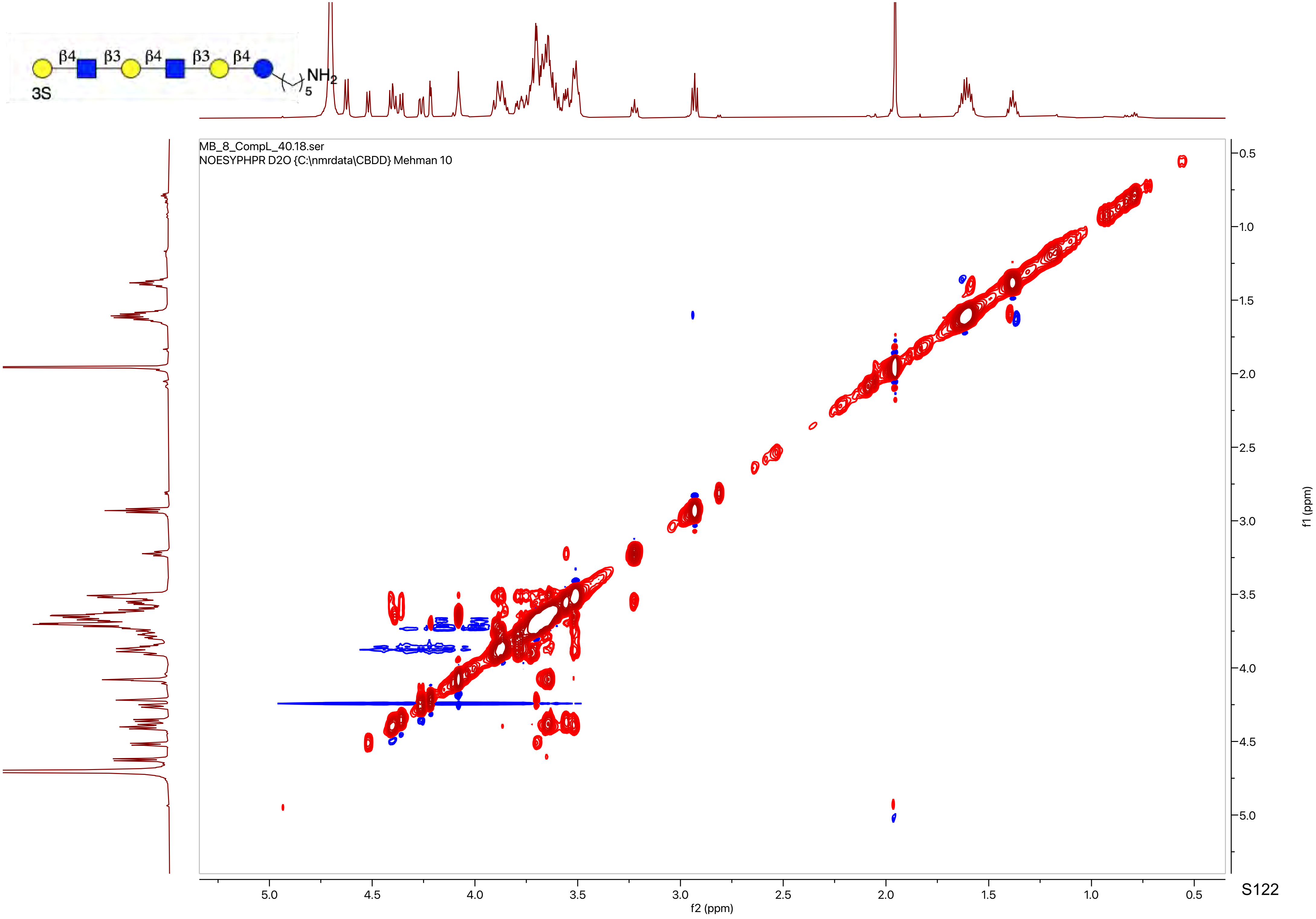

β4
β3
β4
β3
β4
3S
NH2
5
MB_8_CompL_40.18.ser
NOESYPHPR D2O {C:\nmrdata\CBDD} Mehman 10
f2 (ppm)
f1 (ppm)

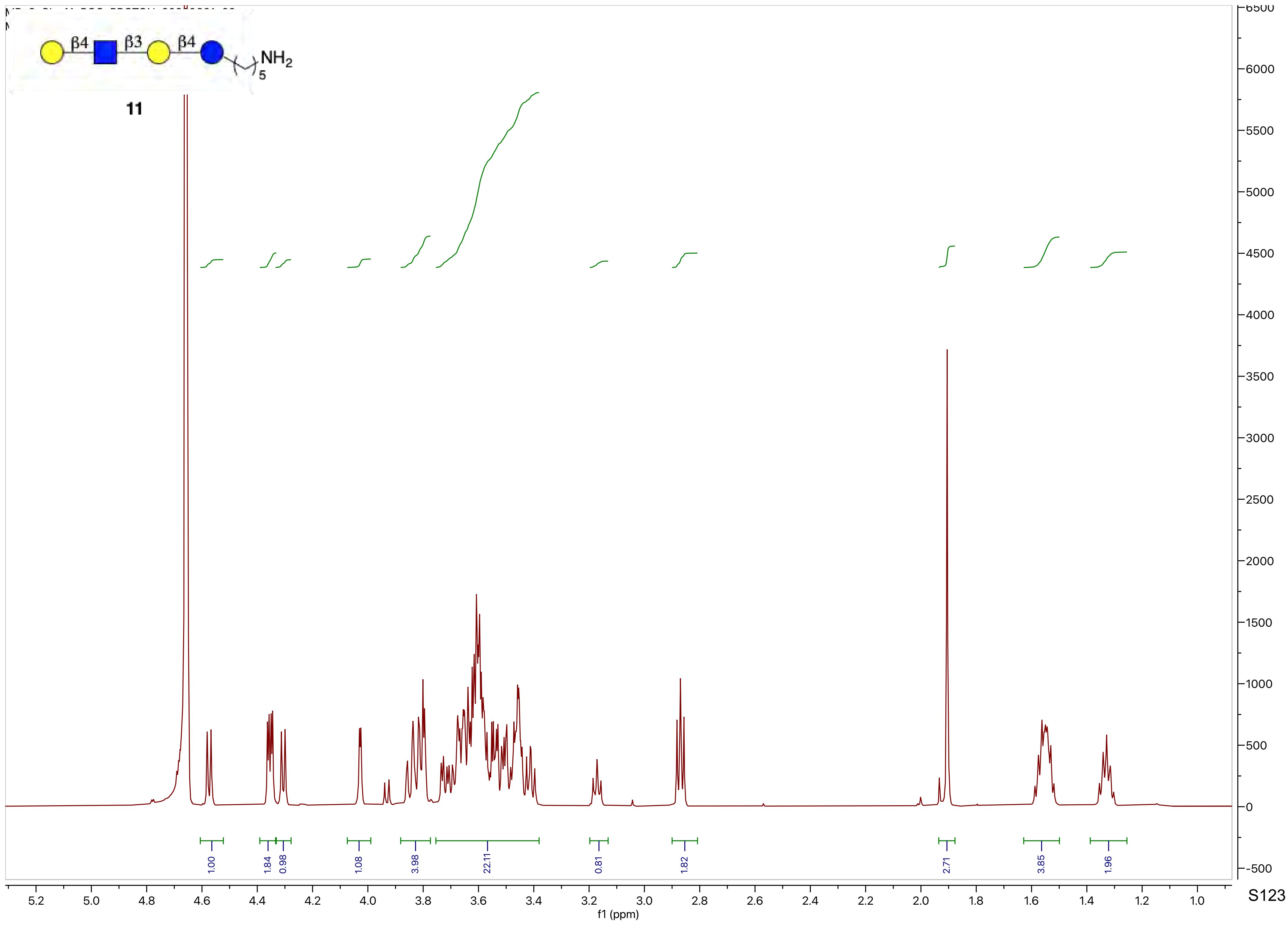
β4
β3
β4
NH2
5
11
1.00
1.84
0.98
1.08
3.98
22.11
0.81
1.82
2.71
3.85
1.96
f1 (ppm)

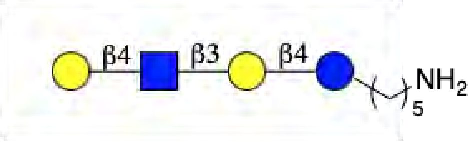
β4
β3
β4
NH2
5

**11**

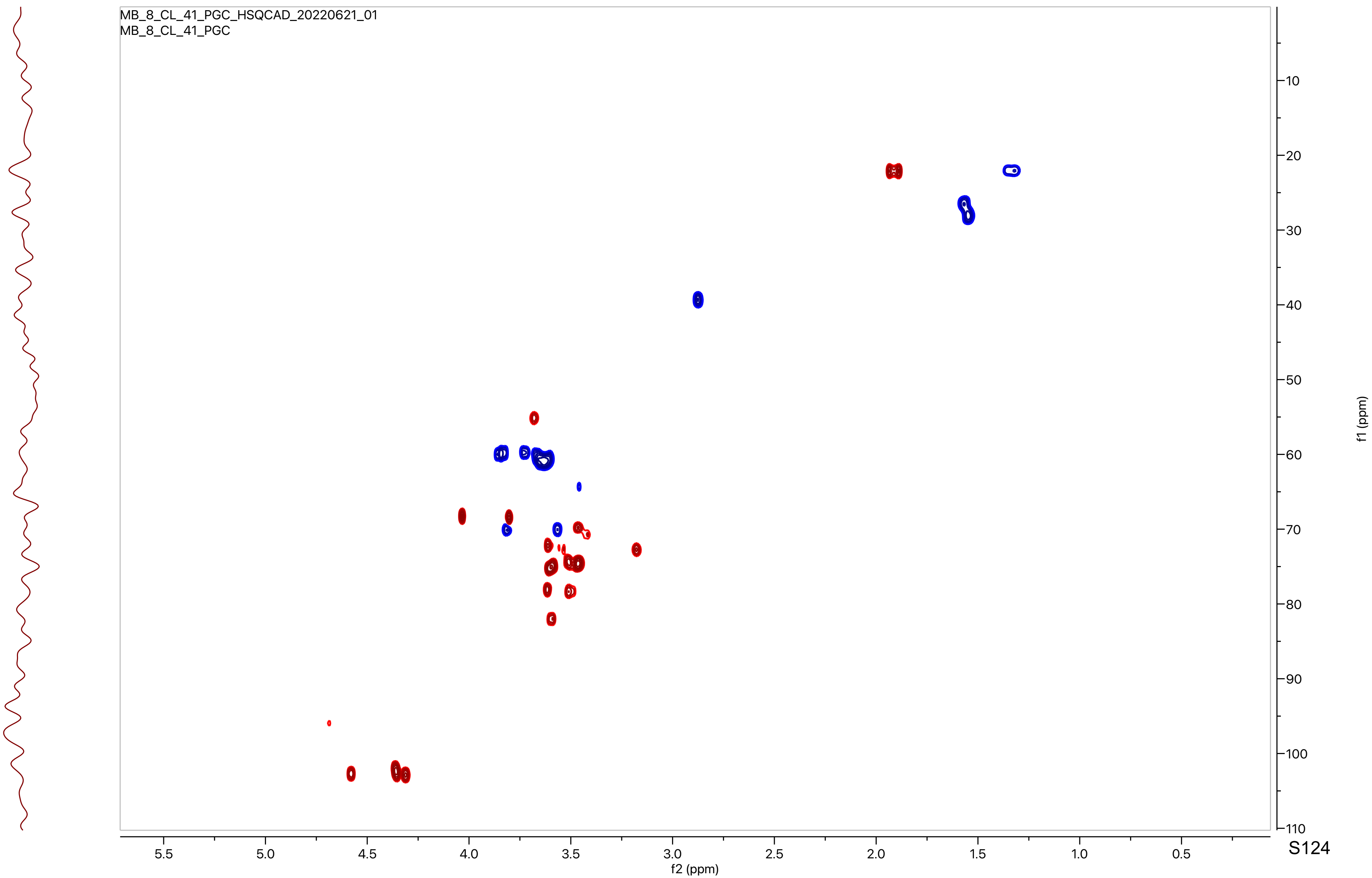
MB_8_CL_41_PGC_HSQCAD_20220621_01
MB_8_CL_41_PGC
f2 (ppm)
f1 (ppm)

**11**

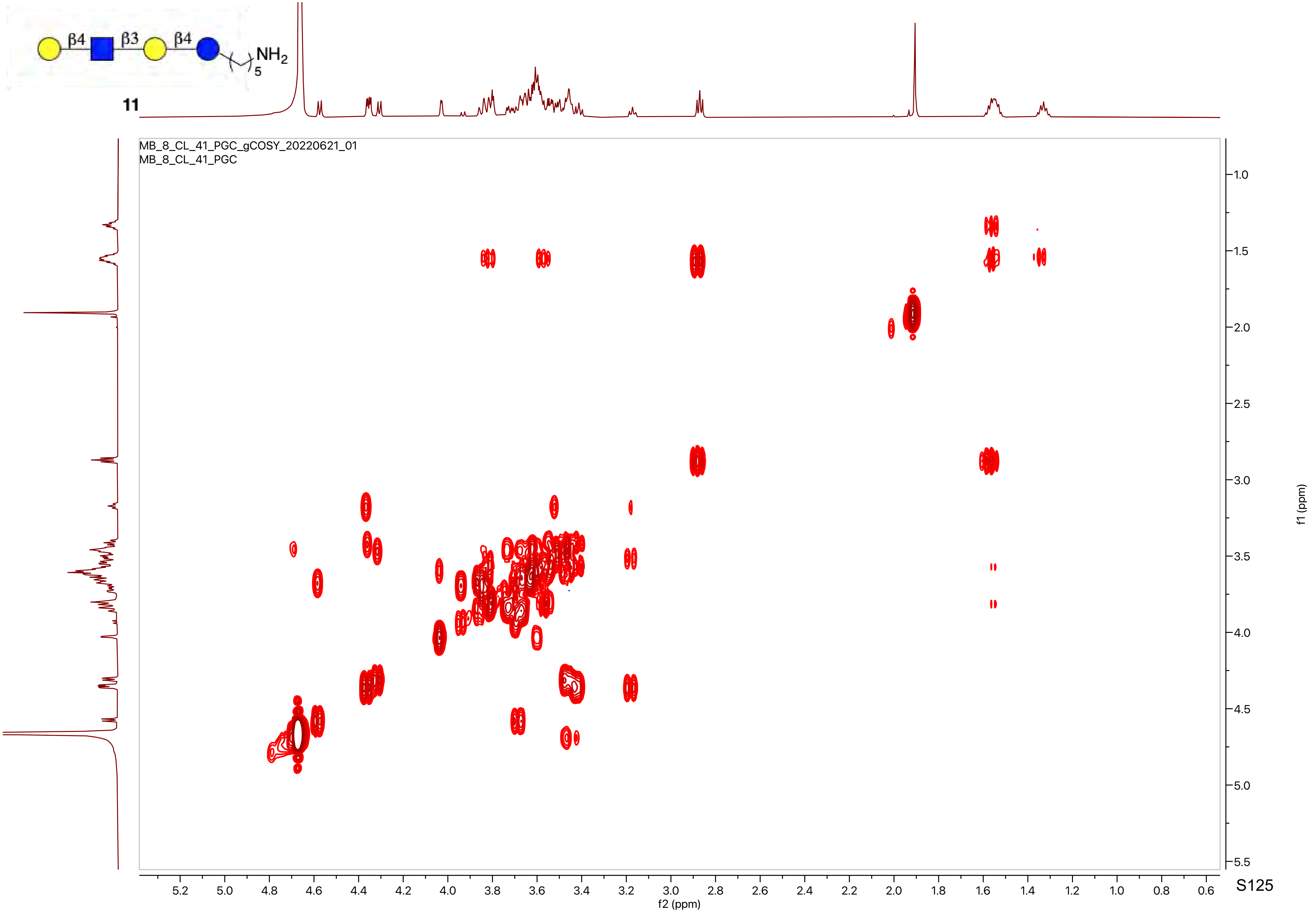

β4
β3
β4
NH2
5
MB_8_CL_41_PGC_gCOSY_20220621_01
MB_8_CL_41_PGC
f2 (ppm)
f1 (ppm)

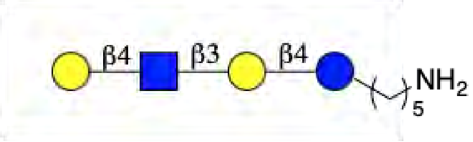
β4
β3
β4
NH2
5

**11**

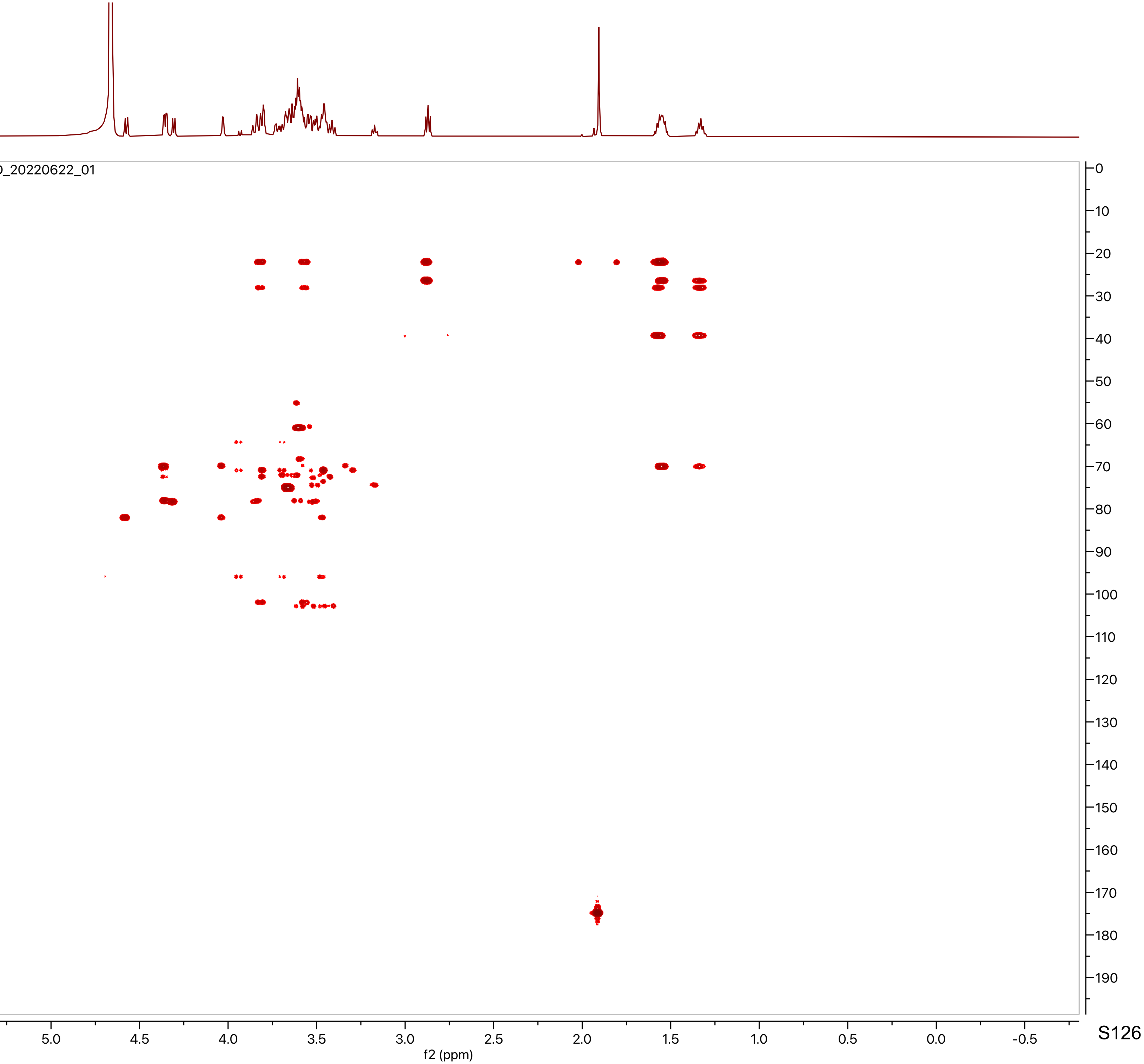
MB_8_CL_41_PGC_gHMBCAD_20220622_01
MB_8_CL_41_PGC
f2 (ppm)
f1 (ppm)

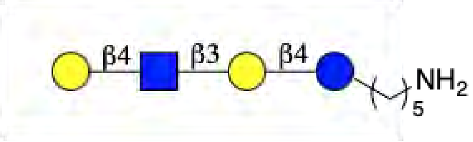
β4
β3
β4
NH2
5

**11**

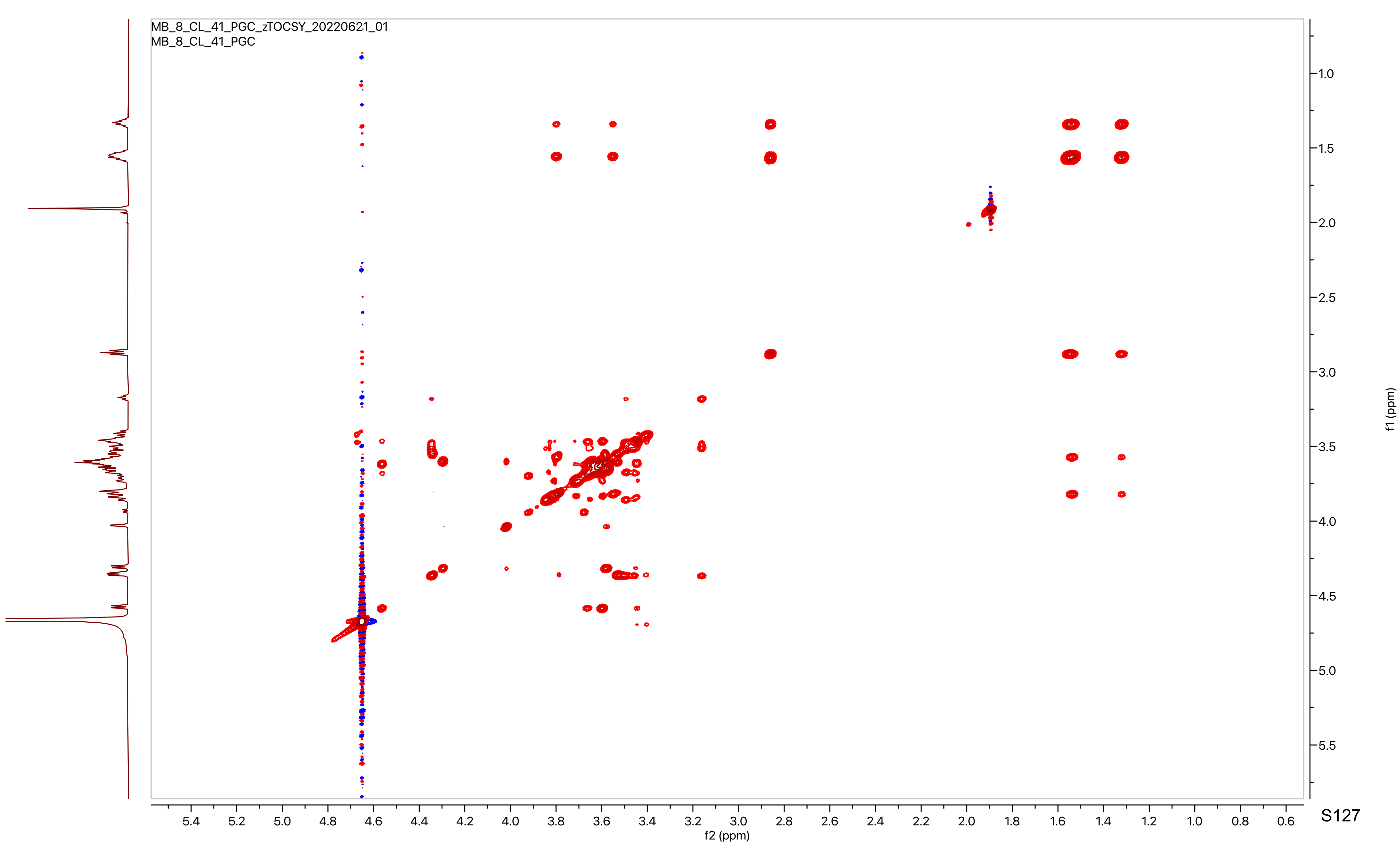
MB_8_CL_41_PGC_zTOCSY_20220621_01
MB_8_CL_41_PGC
f2 (ppm)
f1 (ppm)

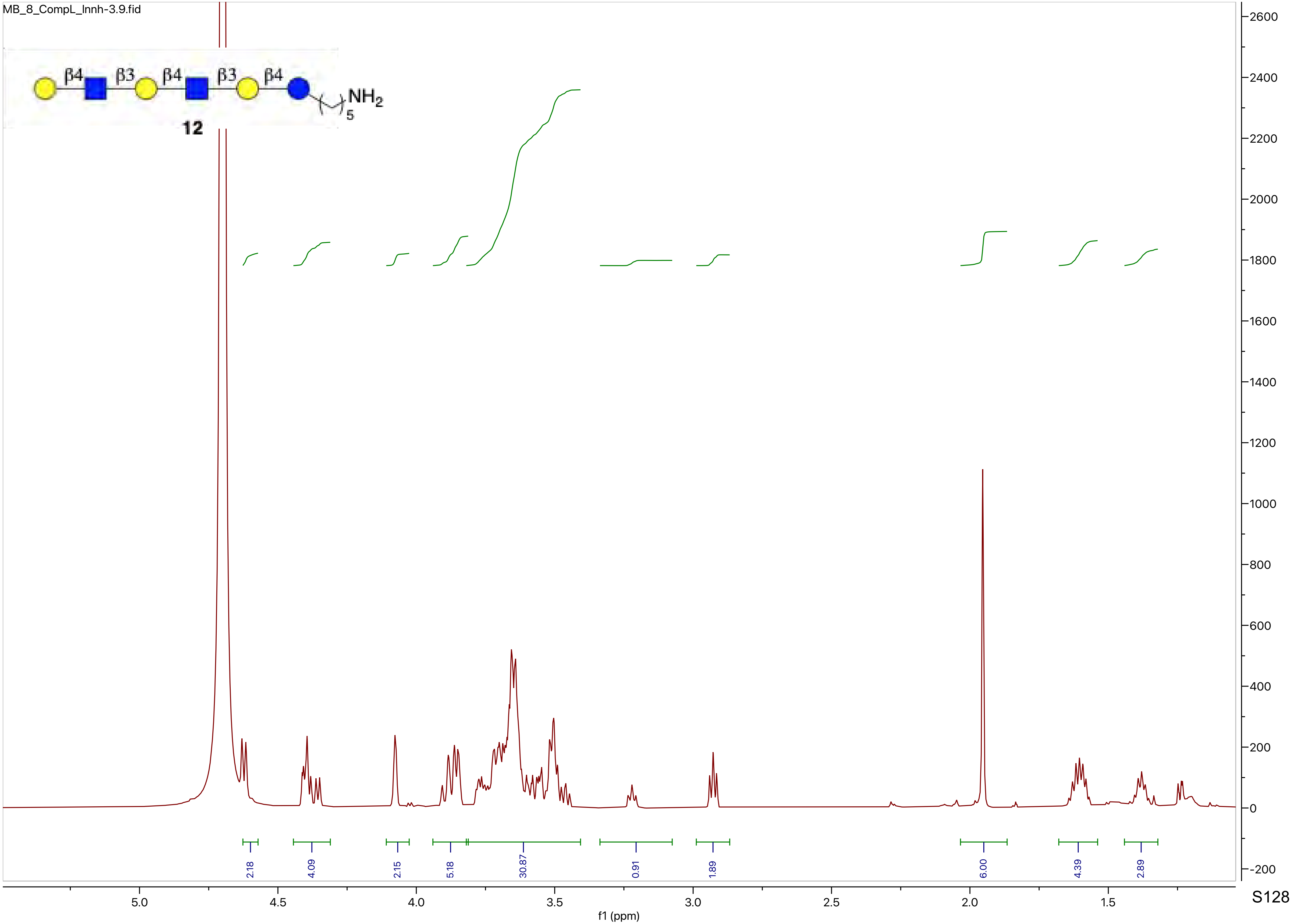
MB_8_CompL_lnnh-3.9.fid
β4
β3
β4
β3
β4
NH2
5
12
2.18
4.09
2.15
5.18
30.87
0.91
1.89
6.00
4.39
2.89
5.0
4.5
4.0
3.5
3.0
2.5
2.0
1.5
f1 (ppm)
2600
2400
2200
2000
1800
1600
1400
1200
1000
800
600
400
200
0
-200

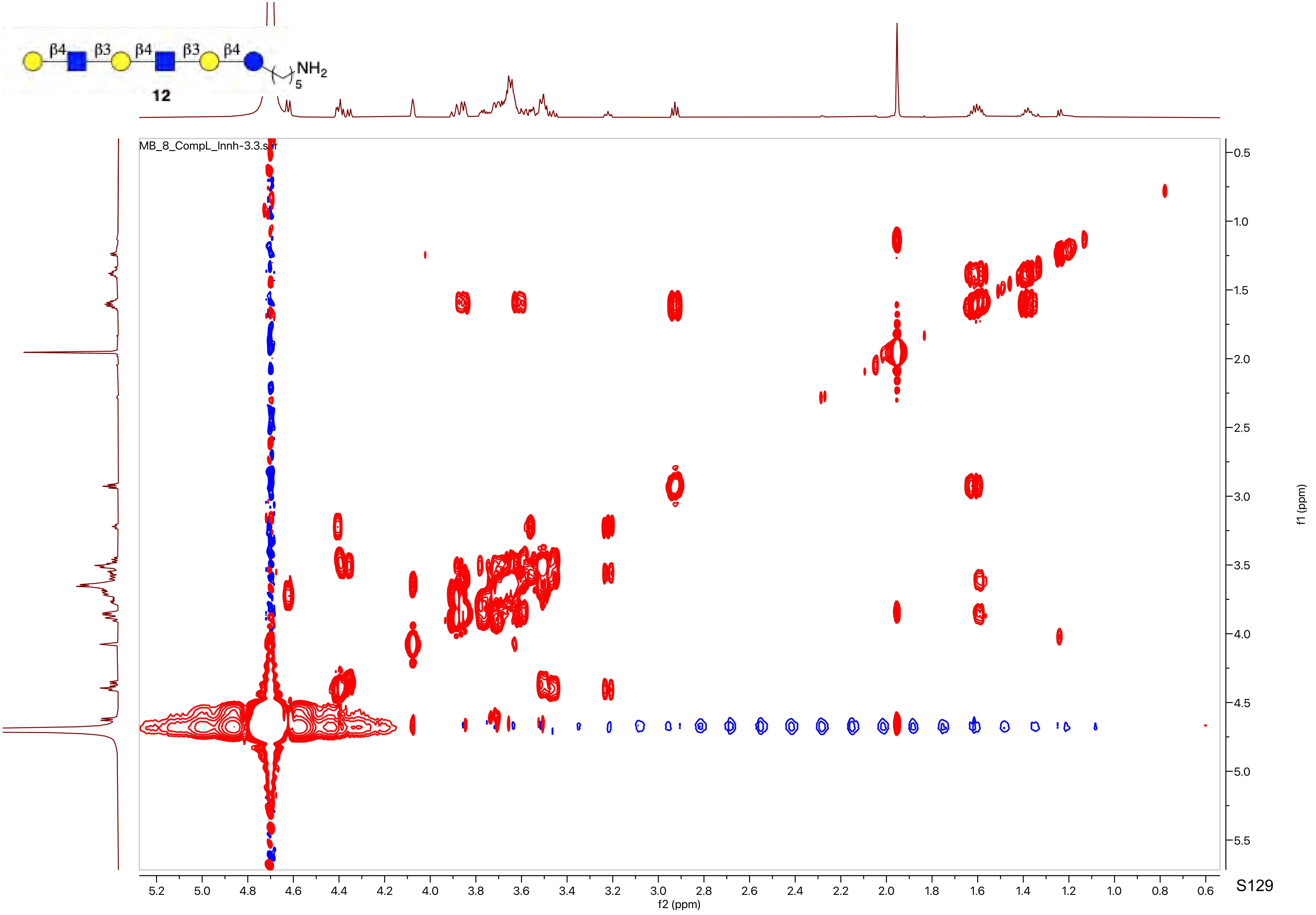
β4
β3
β4
β3
β4
NH2
5
12
MB_8_CompL_Innh-3.3.ser
f2 (ppm)
f1 (ppm)

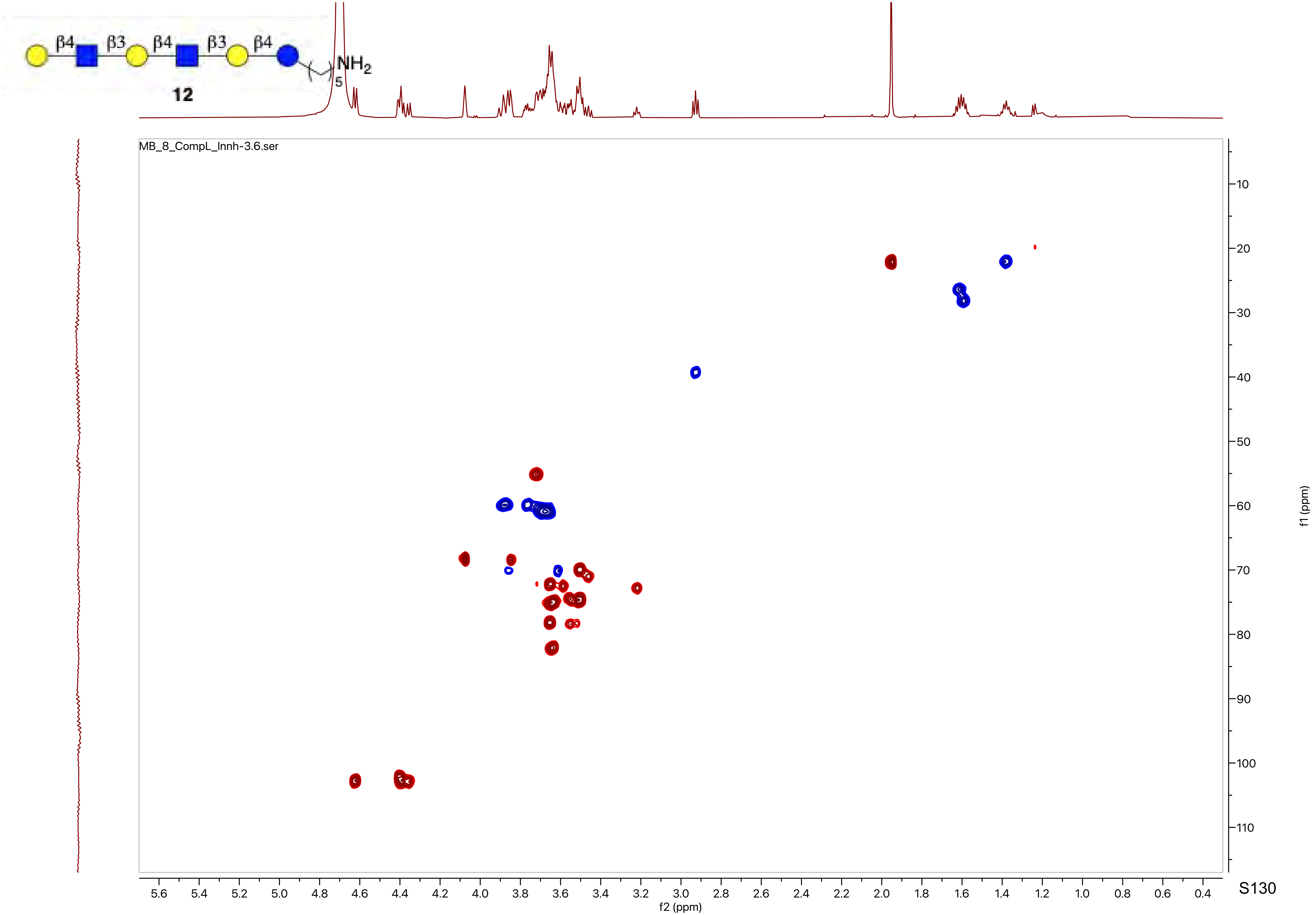
β4
β3
β4
β3
β4
NH2
5
12
MB_8_CompL_Innh-3.6.ser
f2 (ppm)
f1 (ppm)

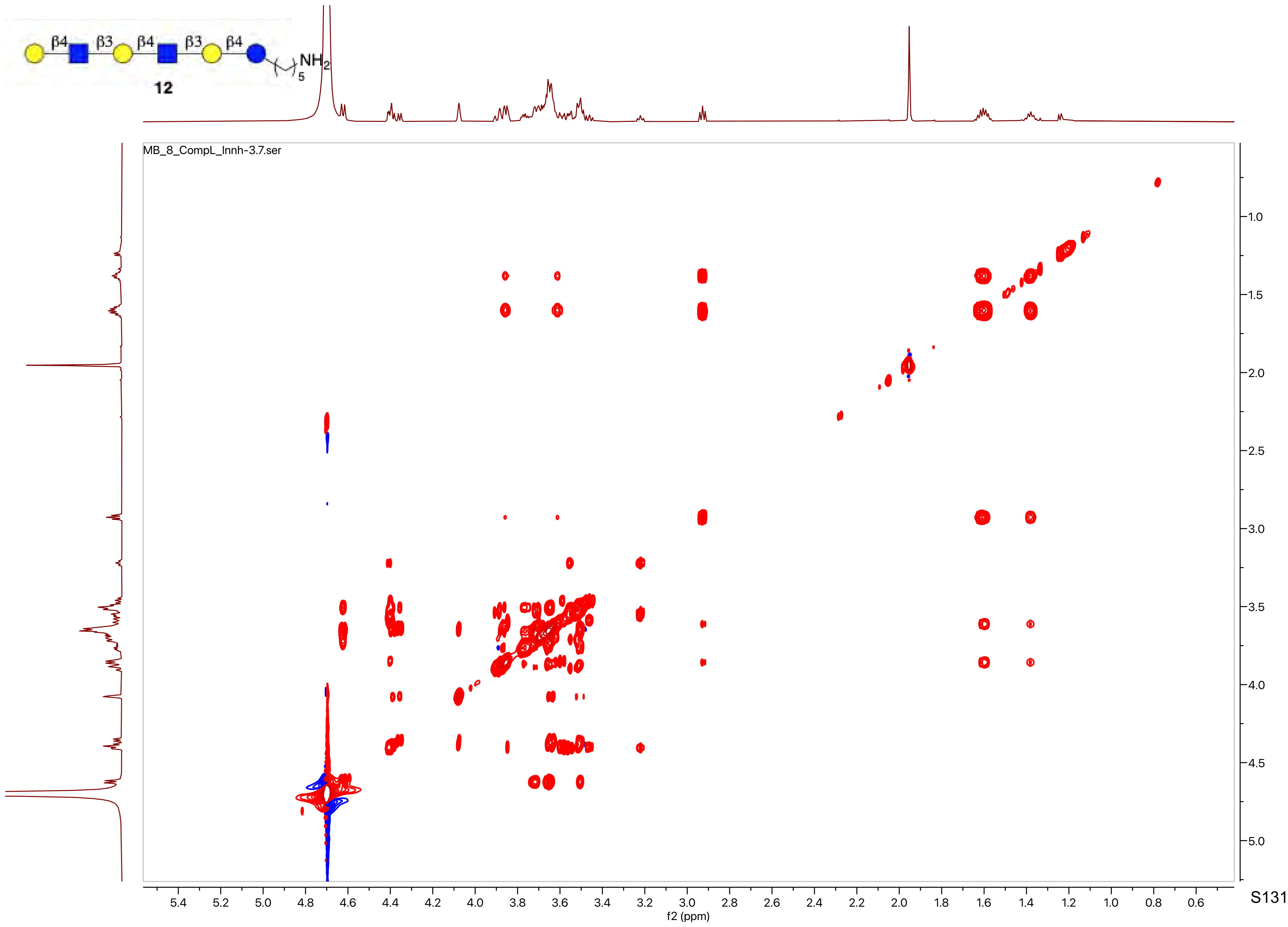

β4
β3
β4
β3
β4
NH2
5
12
MB_8_CompL_Innh-3.7.ser
f2 (ppm)
f1 (ppm)

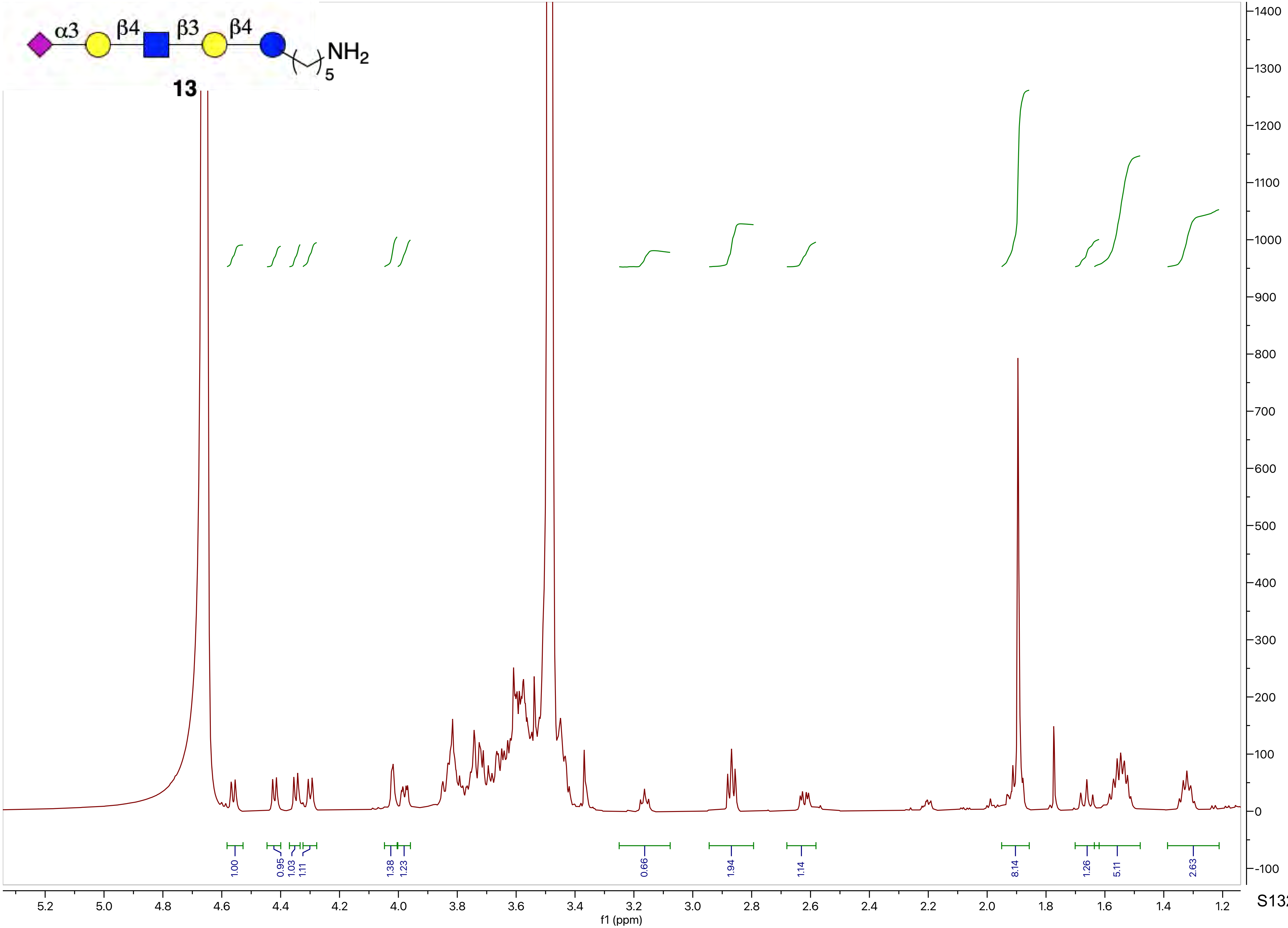

α3
β4
β3
β4
$NH_2$
5
13
f1 (ppm)

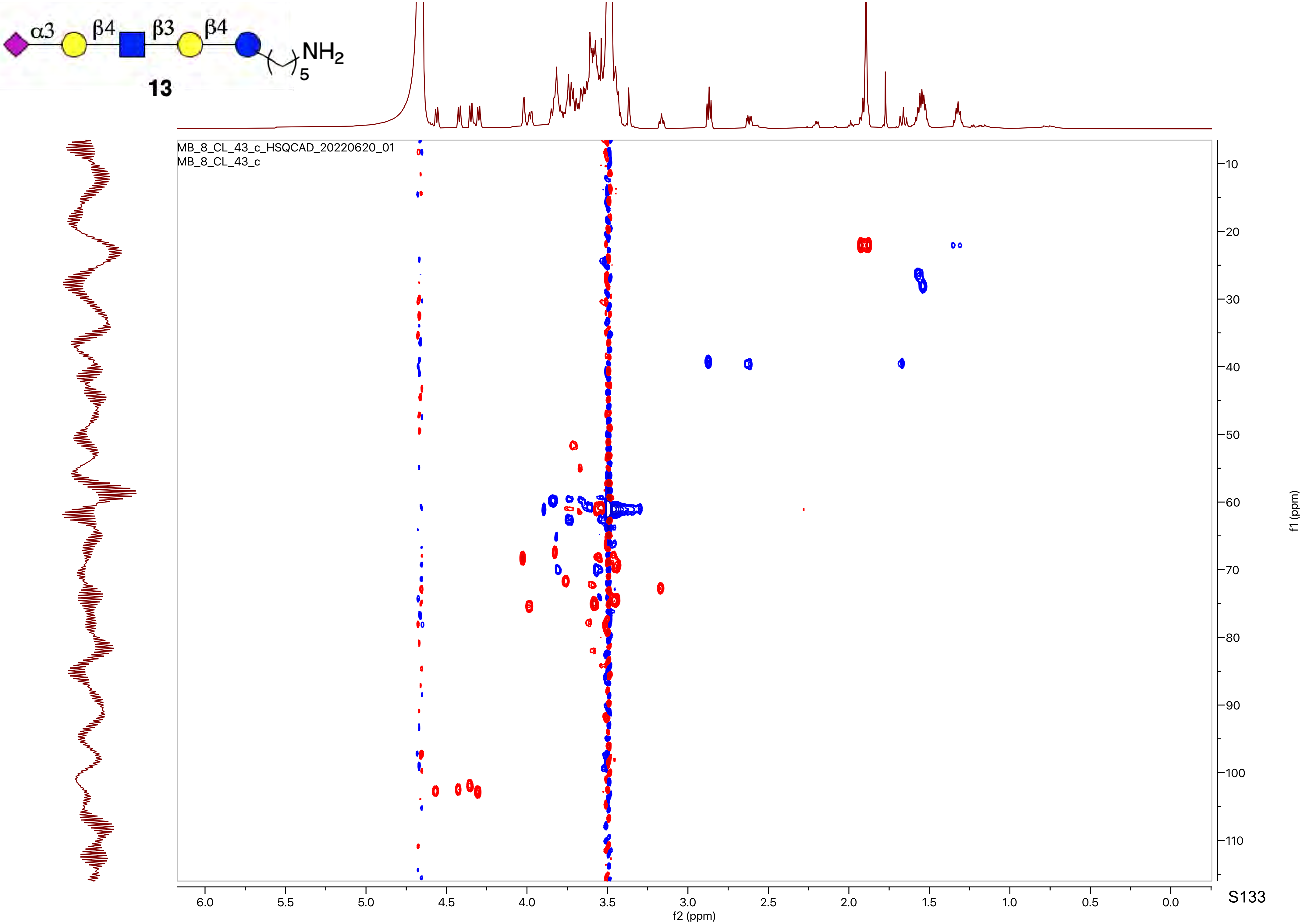
α3
β4
β3
β4
$NH_2$
5
MB_8_CL_43_c_HSQCAD_20220620_01
MB_8_CL_43_c
f2 (ppm)
f1 (ppm)

**13**

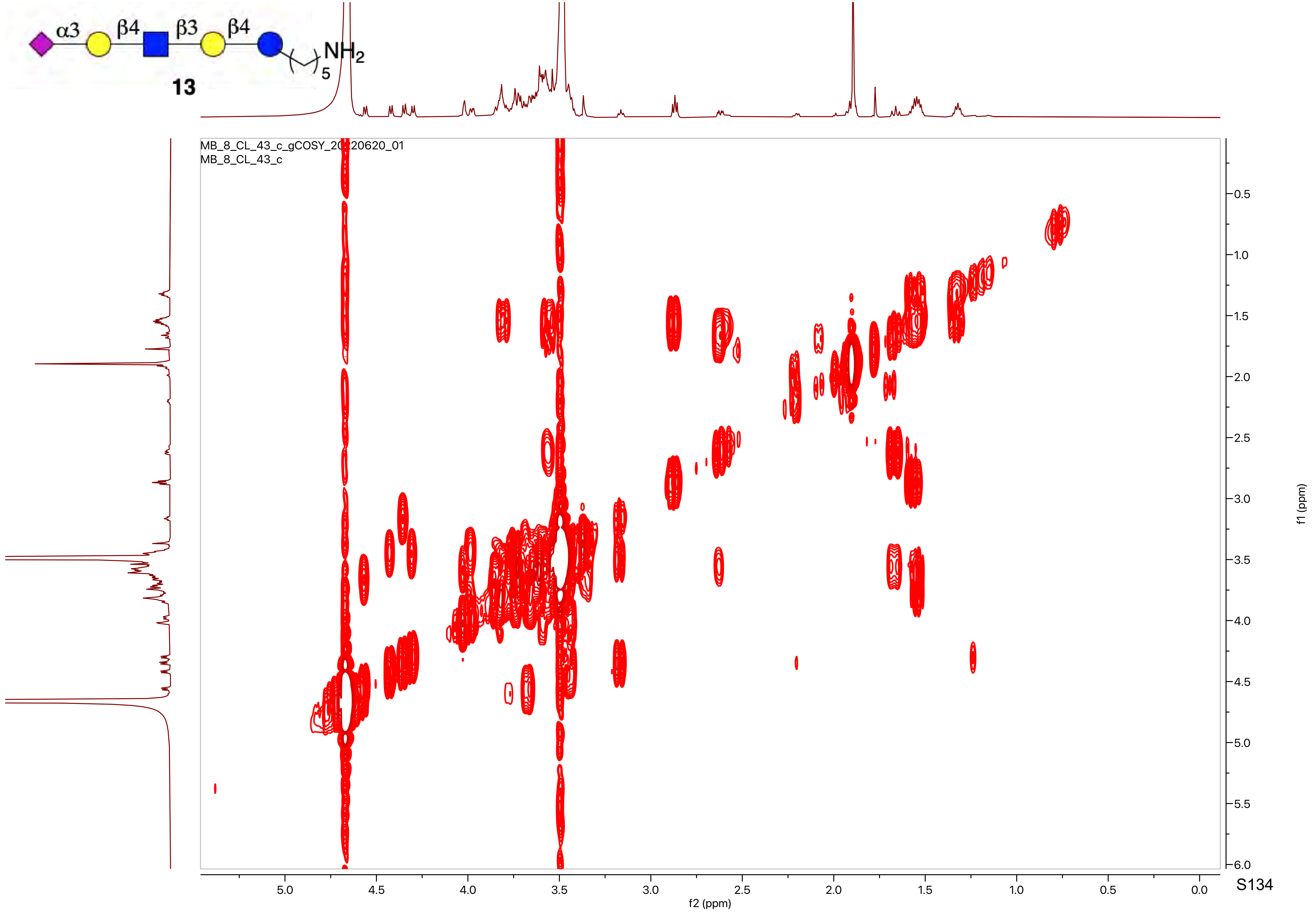
α3
β4
β3
β4
NH2
5
13
MB_8_CL_43_c_gCOSY_20220620_01
MB_8_CL_43_c
f2 (ppm)
f1 (ppm)

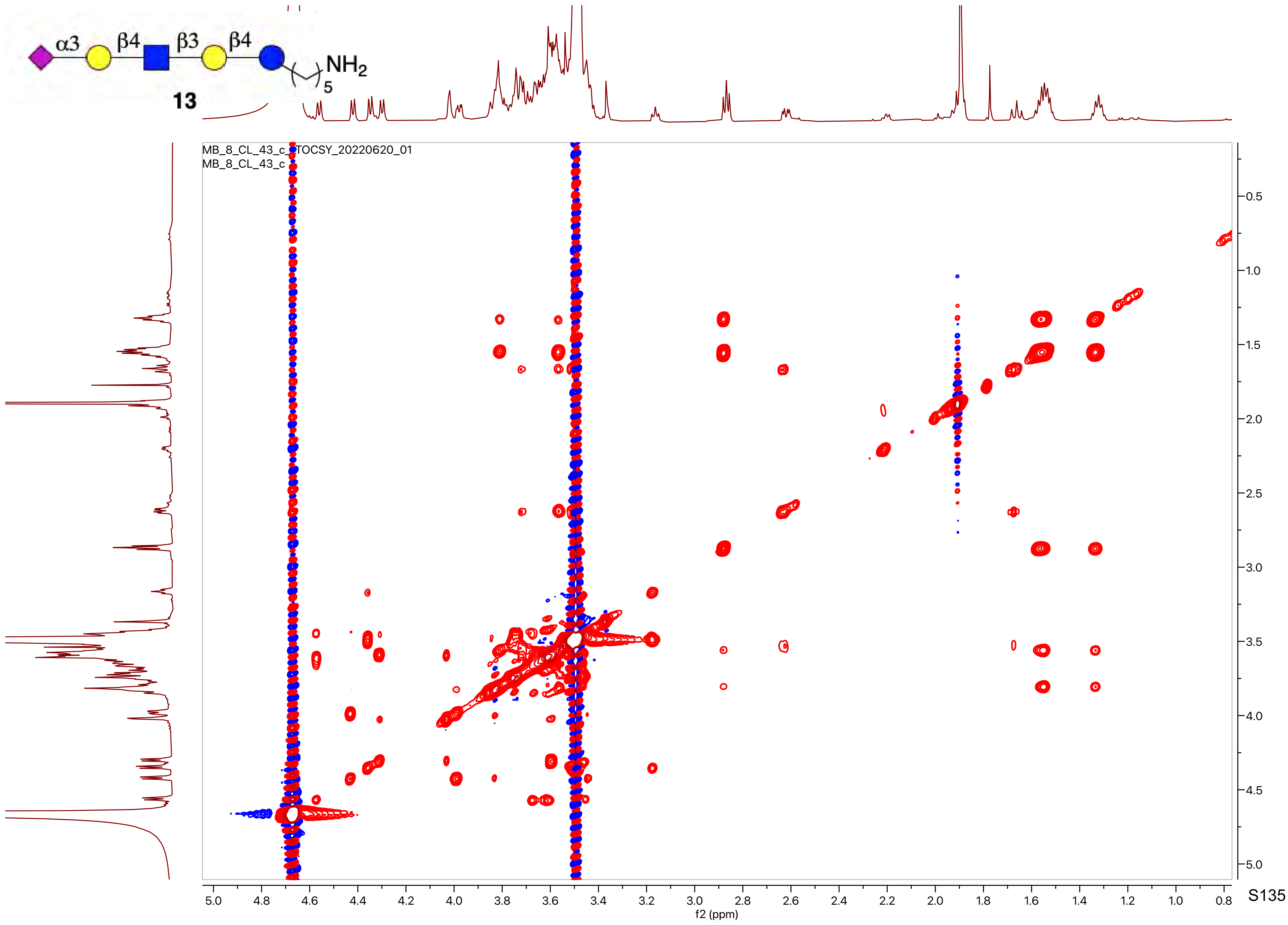

α3
β4
β3
β4
NH2
5
13
MB_8_CL_43_c_TOCSY_20220620_01
MB_8_CL_43_c
f2 (ppm)
f1 (ppm)

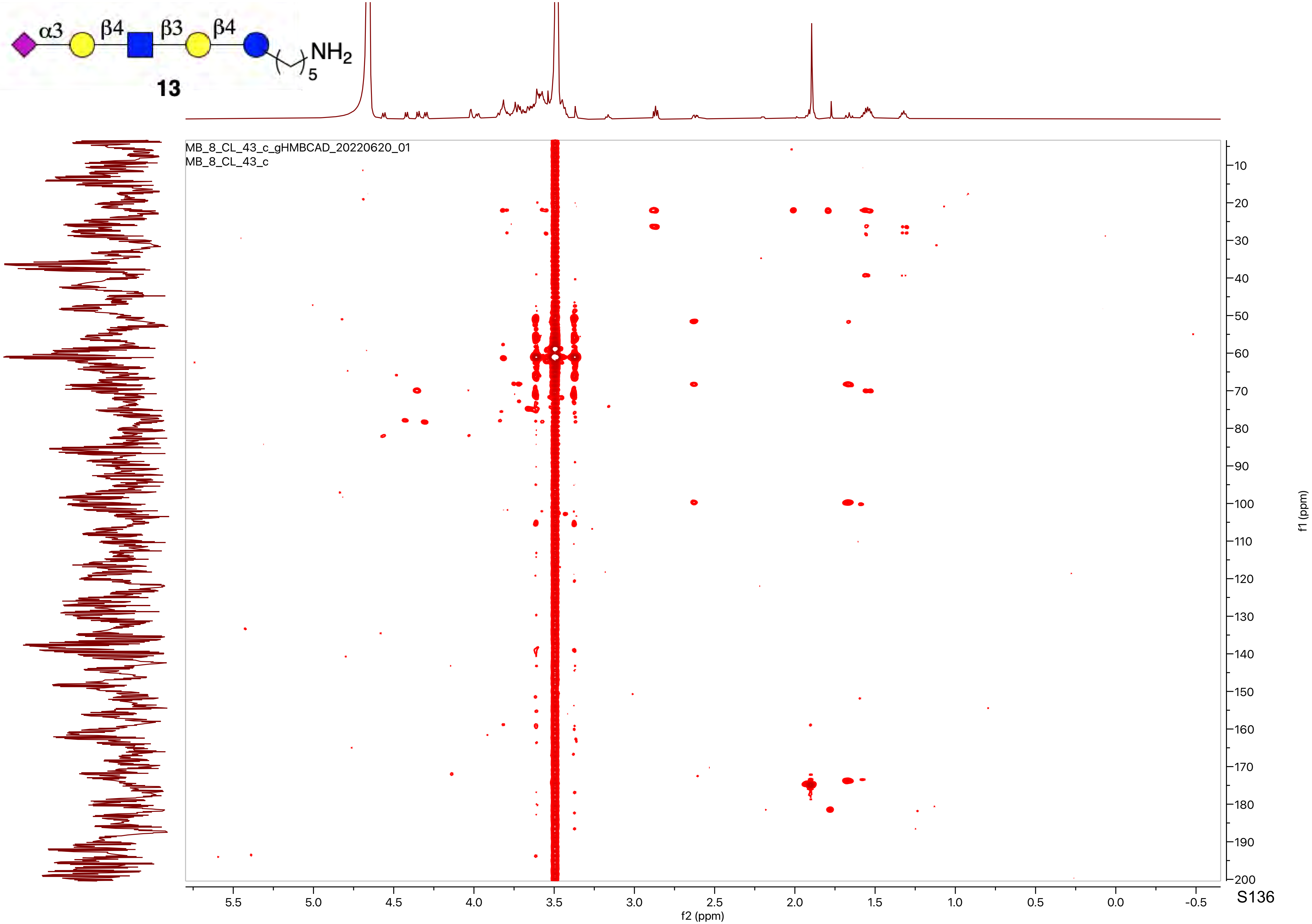
α3
β4
β3
β4
NH2
5
MB_8_CL_43_c_gHMBCAD_20220620_01
MB_8_CL_43_c
f2 (ppm)
f1 (ppm)

**13**

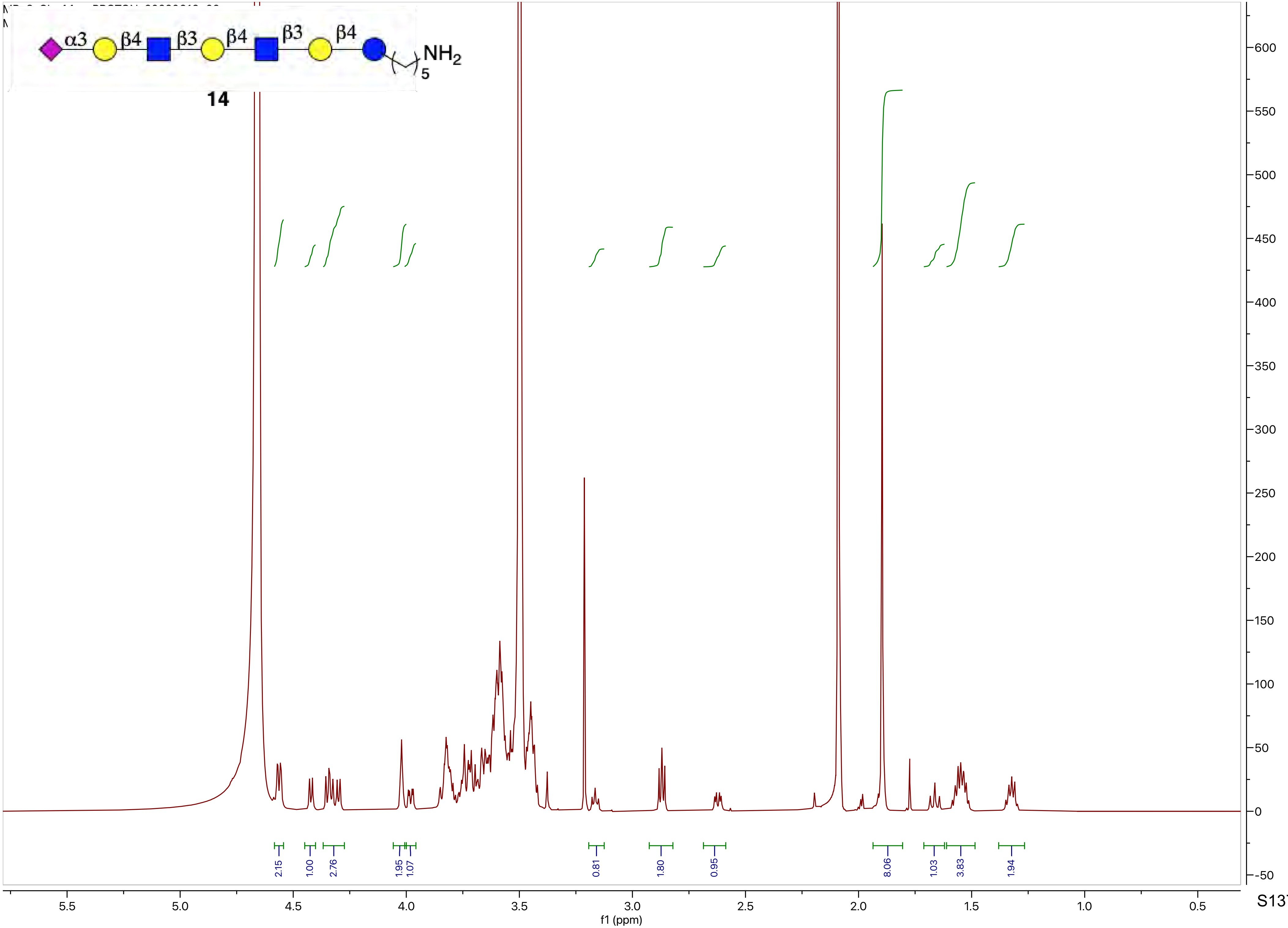
α3
β4
β3
β4
β3
β4
NH2
5
14
2.15
1.00
2.76
1.95
1.07
0.81
1.80
0.95
8.06
1.03
3.83
1.94
f1 (ppm)

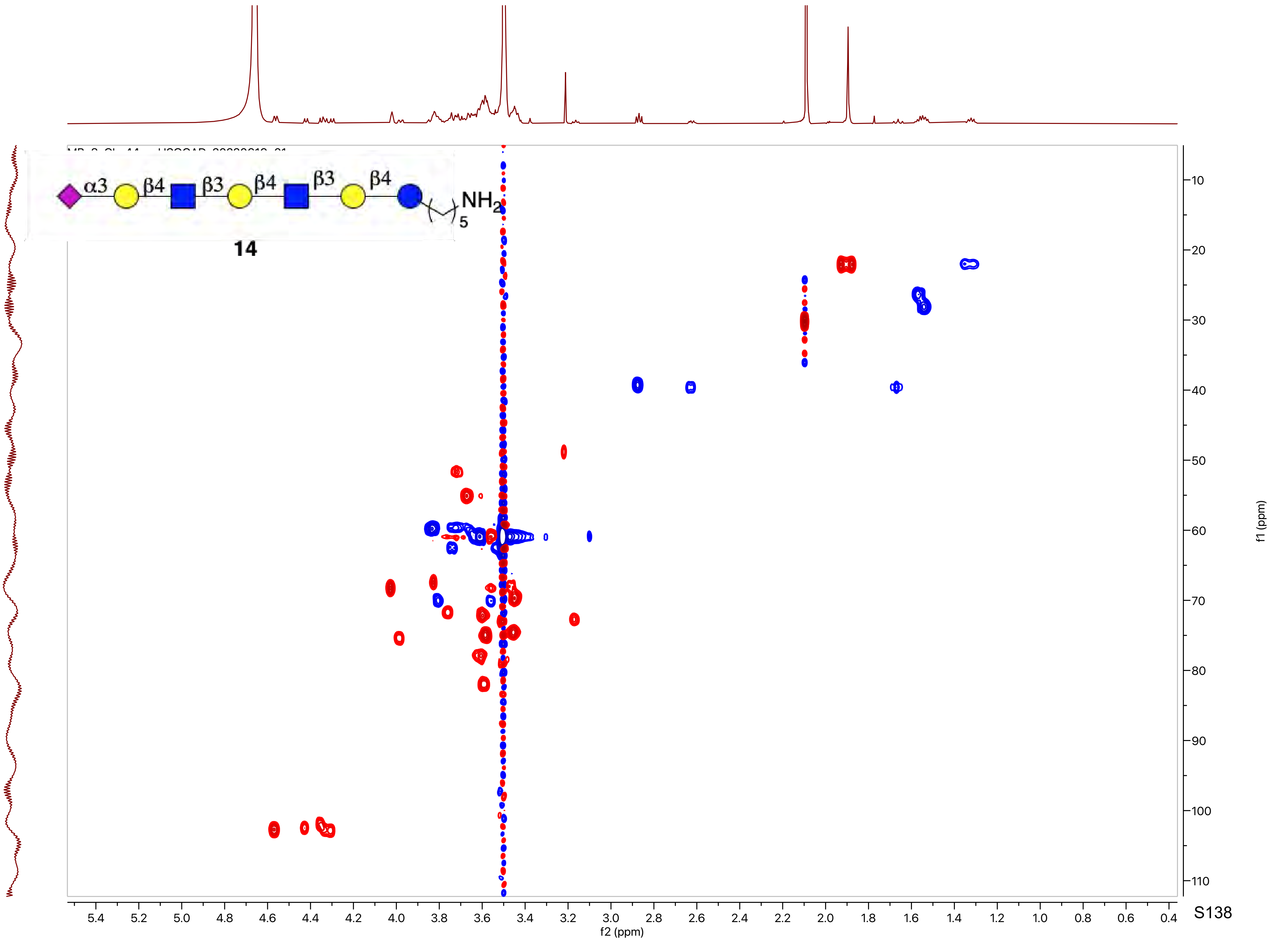
α3
β4
β3
β4
β3
β4
NH2
5
14
f2 (ppm)
f1 (ppm)

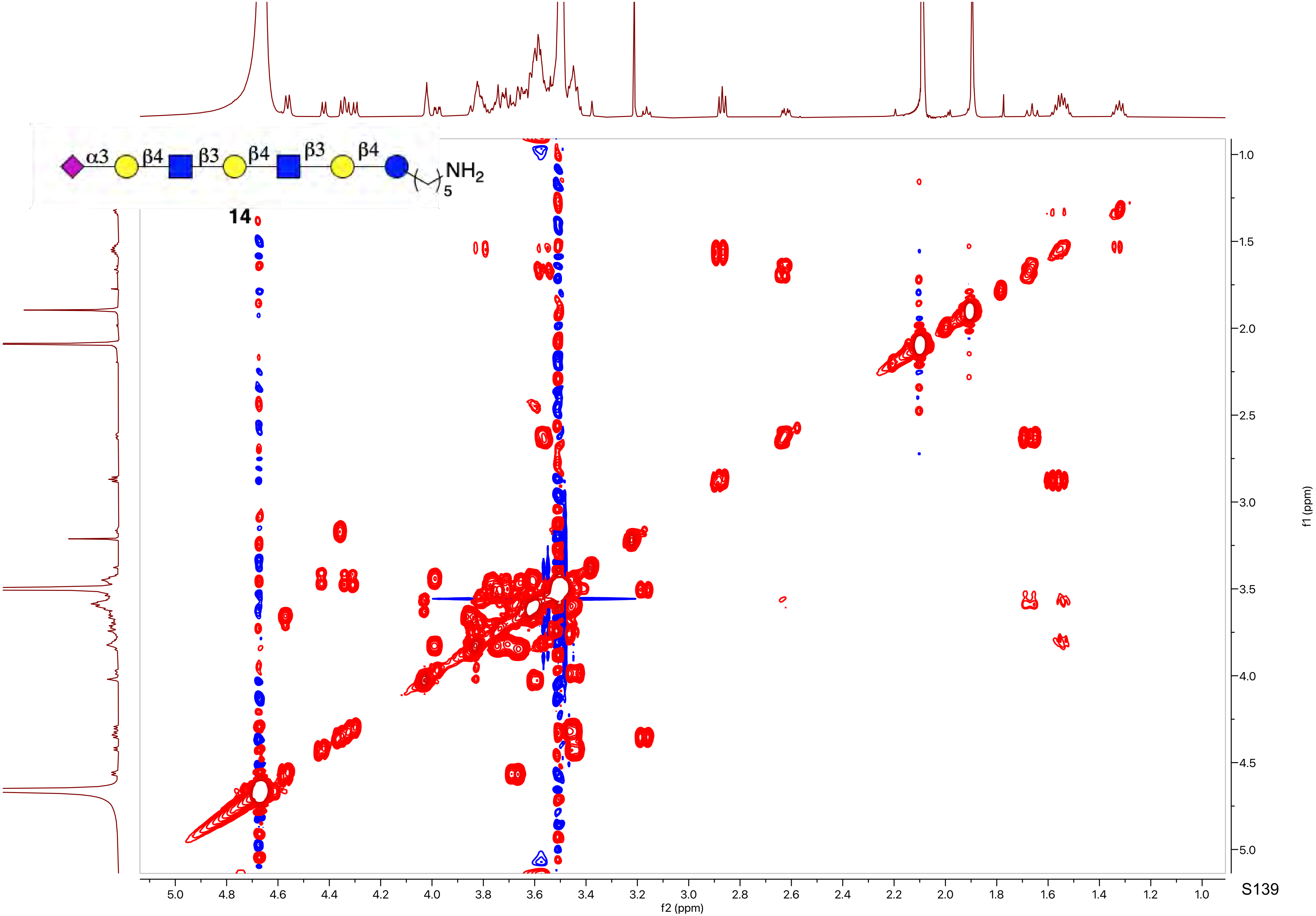

α3
β4
β3
β4
β3
β4
NH2
5
14
f2 (ppm)
f1 (ppm)

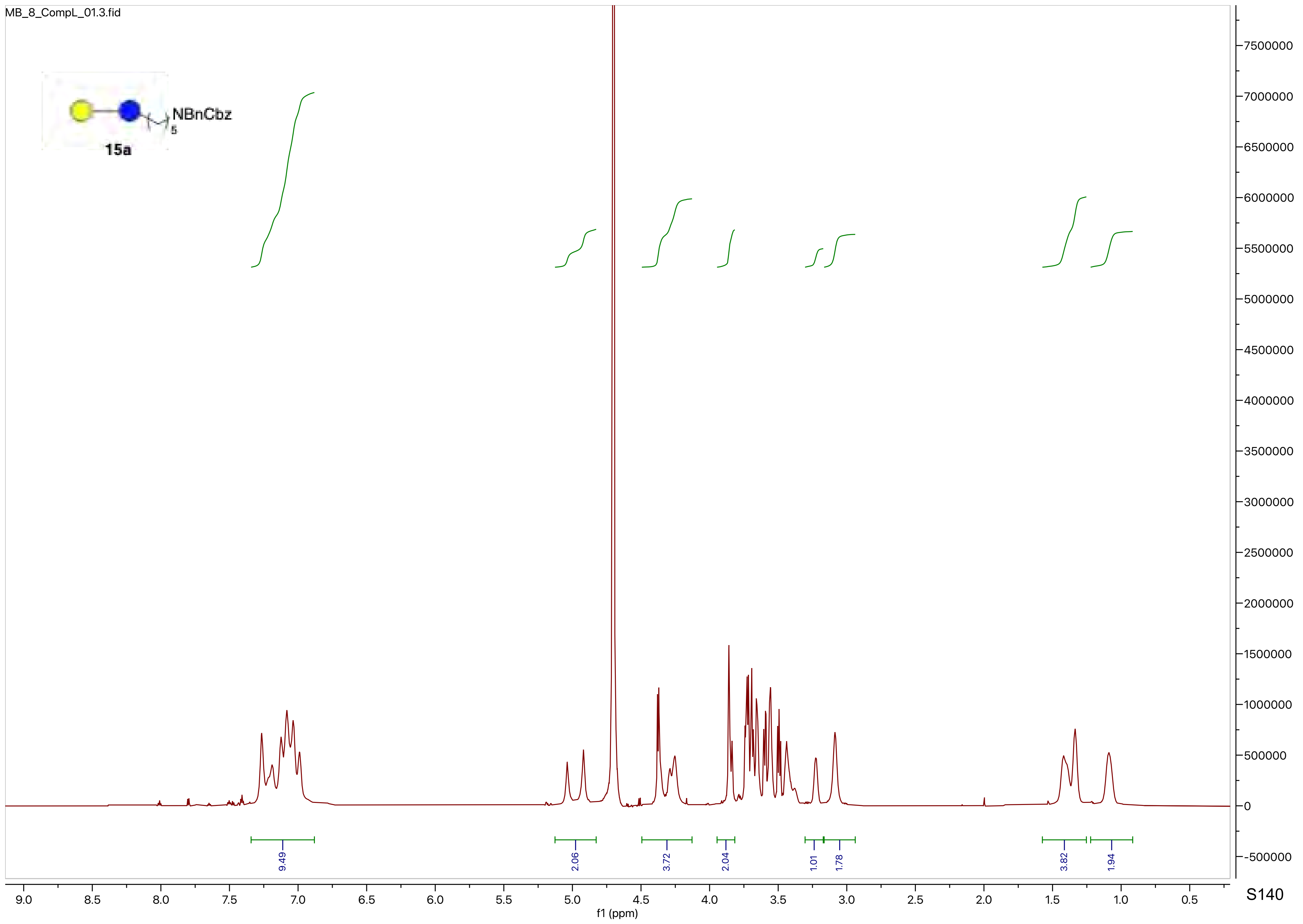

MB_8_CompL_01.3.fid
NBnCbz
5
15a
9.49
2.06
3.72
2.04
1.01
1.78
3.82
1.94
f1 (ppm)

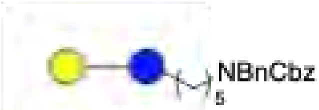
NBnCbz
5

15a

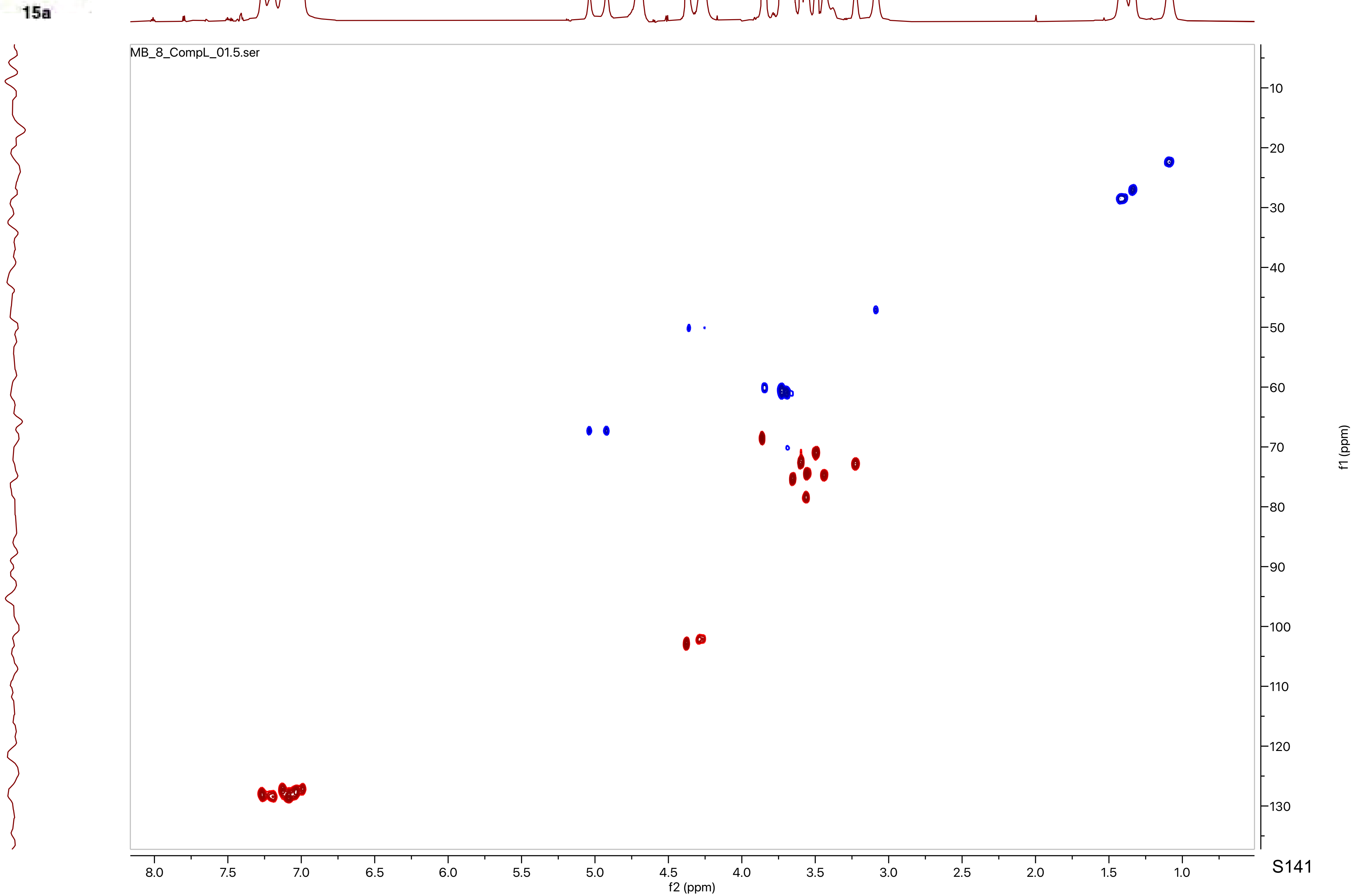
MB_8_CompL_01.5.ser
f2 (ppm)
f1 (ppm)
8.0
7.5
7.0
6.5
6.0
5.5
5.0
4.5
4.0
3.5
3.0
2.5
2.0
1.5
1.0
10
20
30
40
50
60
70
80
90
100
110
120
130

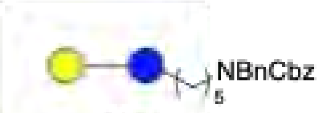

15a

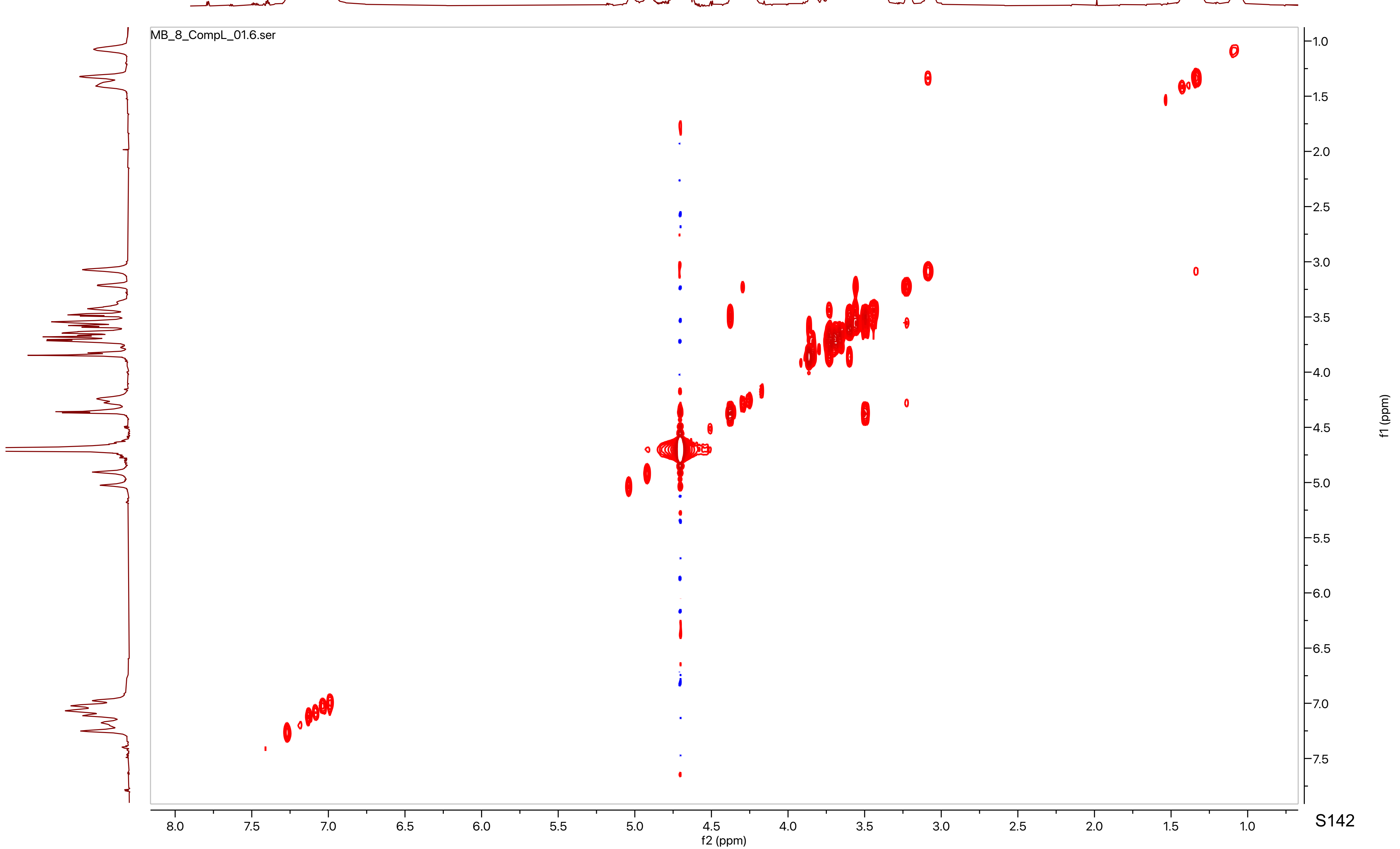

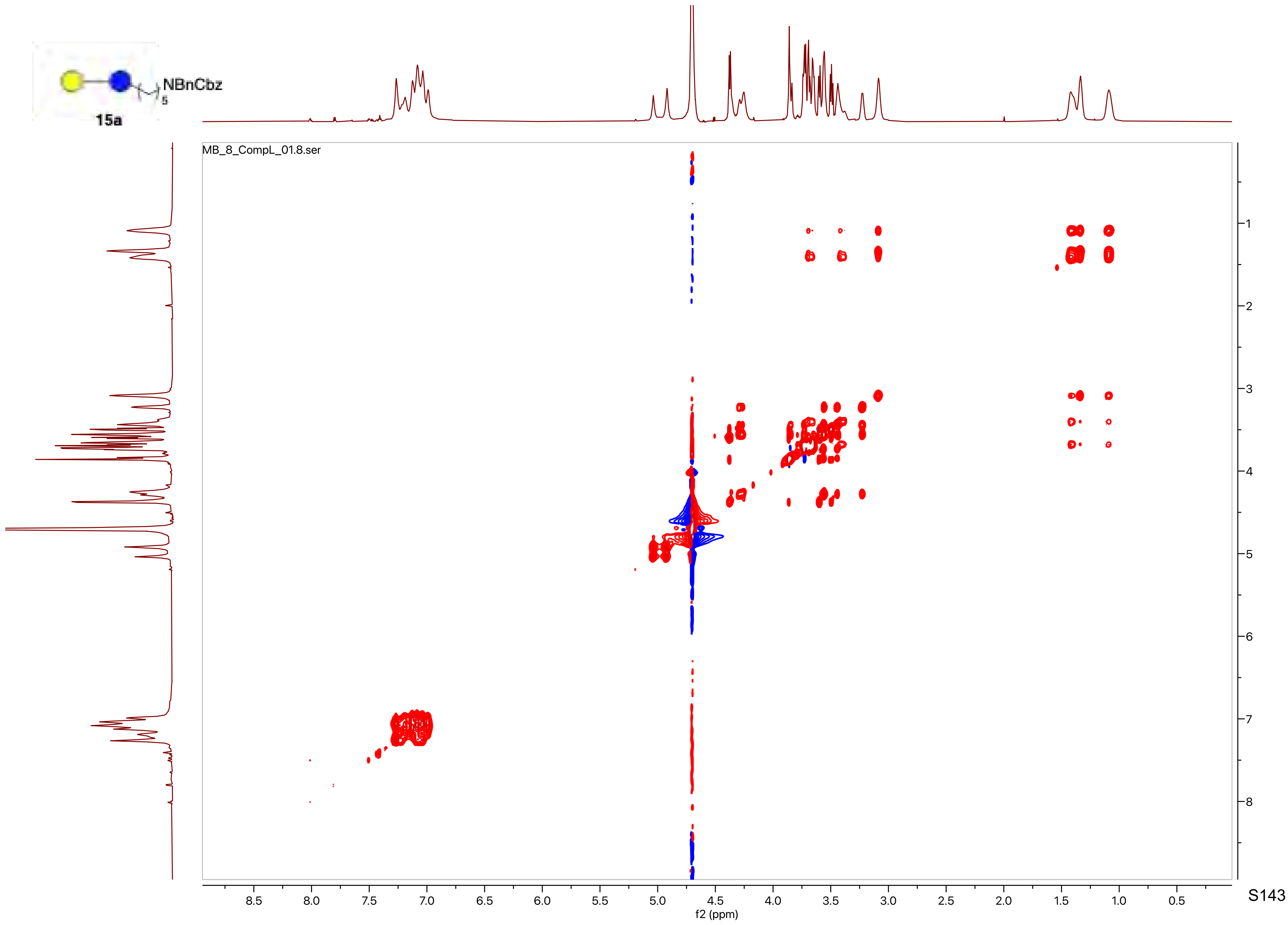
NBnCbz
5
15a
MB_8_CompL_01.8.ser
f2 (ppm)
f1 (ppm)

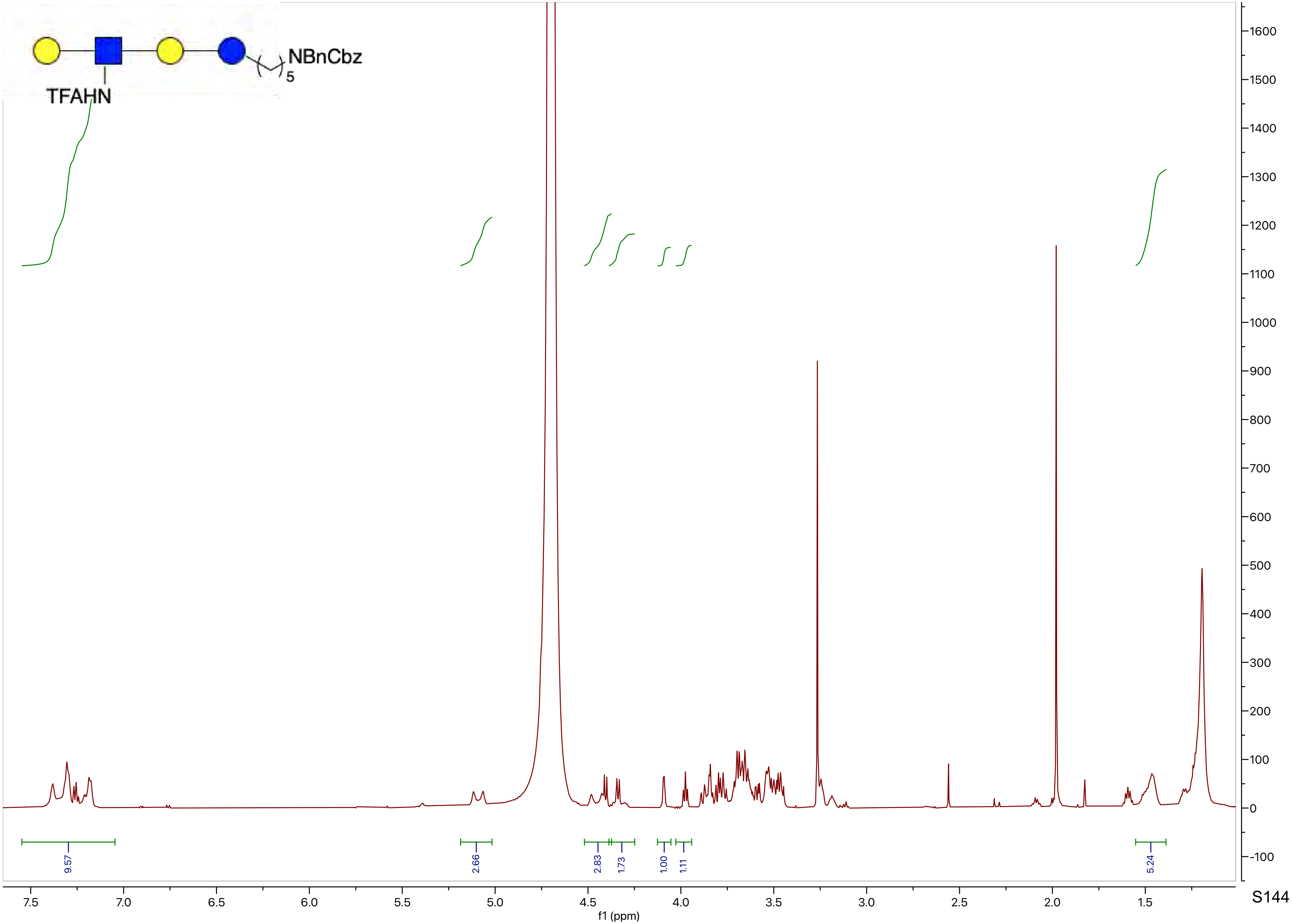
NBnCbz
5
TFAHN
9.57
2.66
2.83
1.73
1.00
1.11
5.24
f1 (ppm)

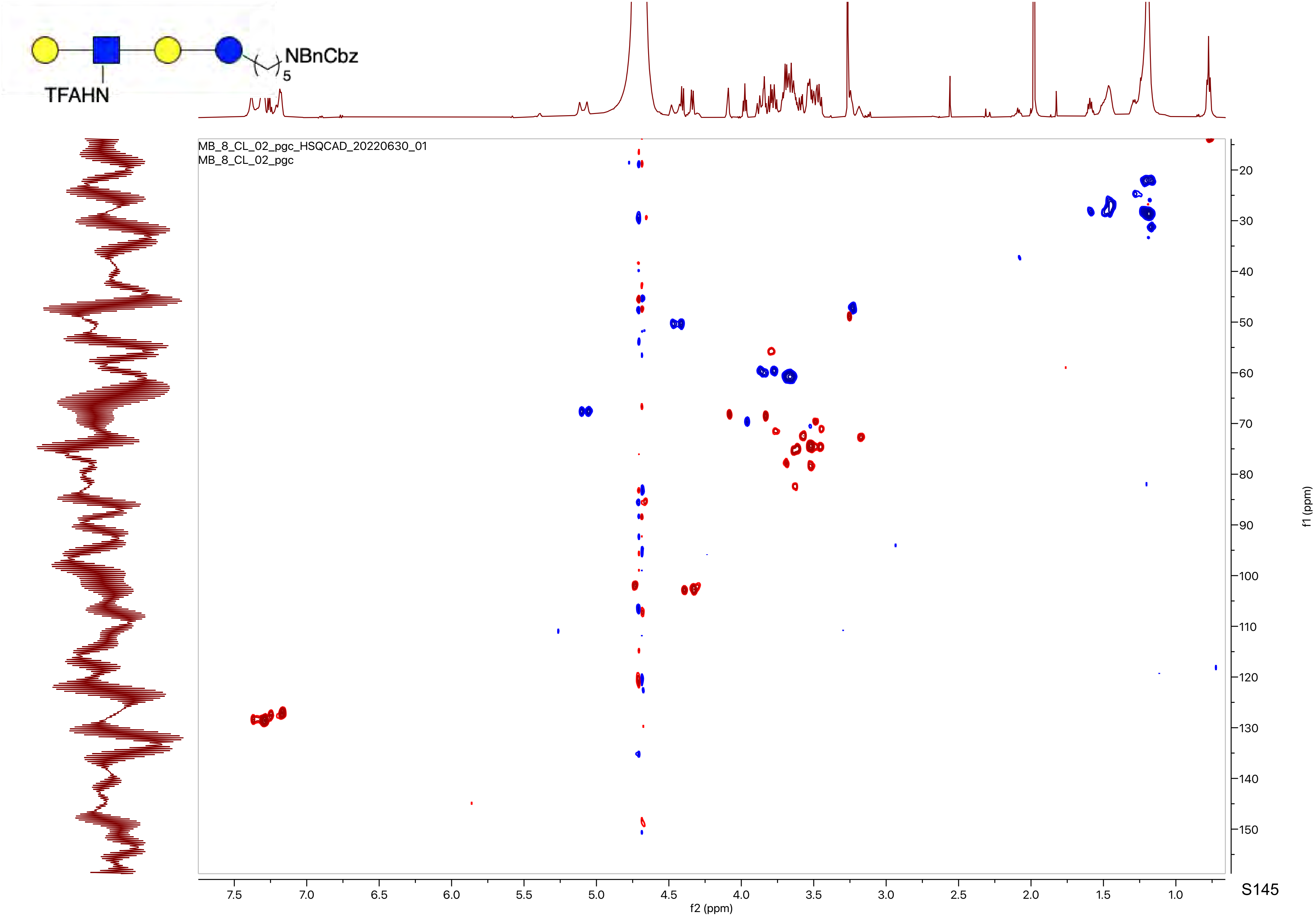
NBnCbz
5
TFAHN
MB_8_CL_02_pgc_HSQCAD_20220630_01
MB_8_CL_02_pgc
f2 (ppm)
f1 (ppm)

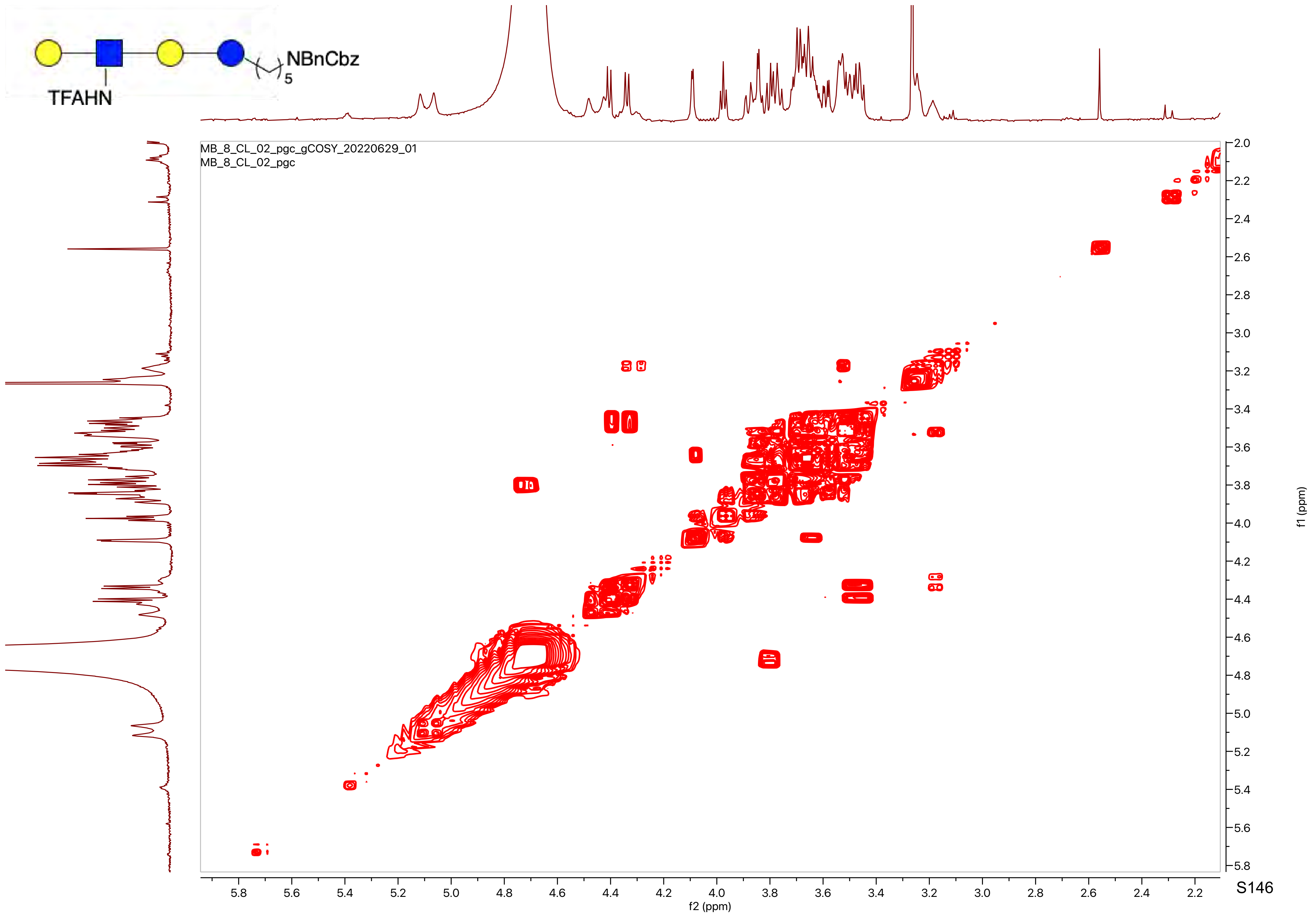
NBnCbz
5
TFAHN
MB_8_CL_02_pgc_gCOSY_20220629_01
MB_8_CL_02_pgc
f2 (ppm)
f1 (ppm)

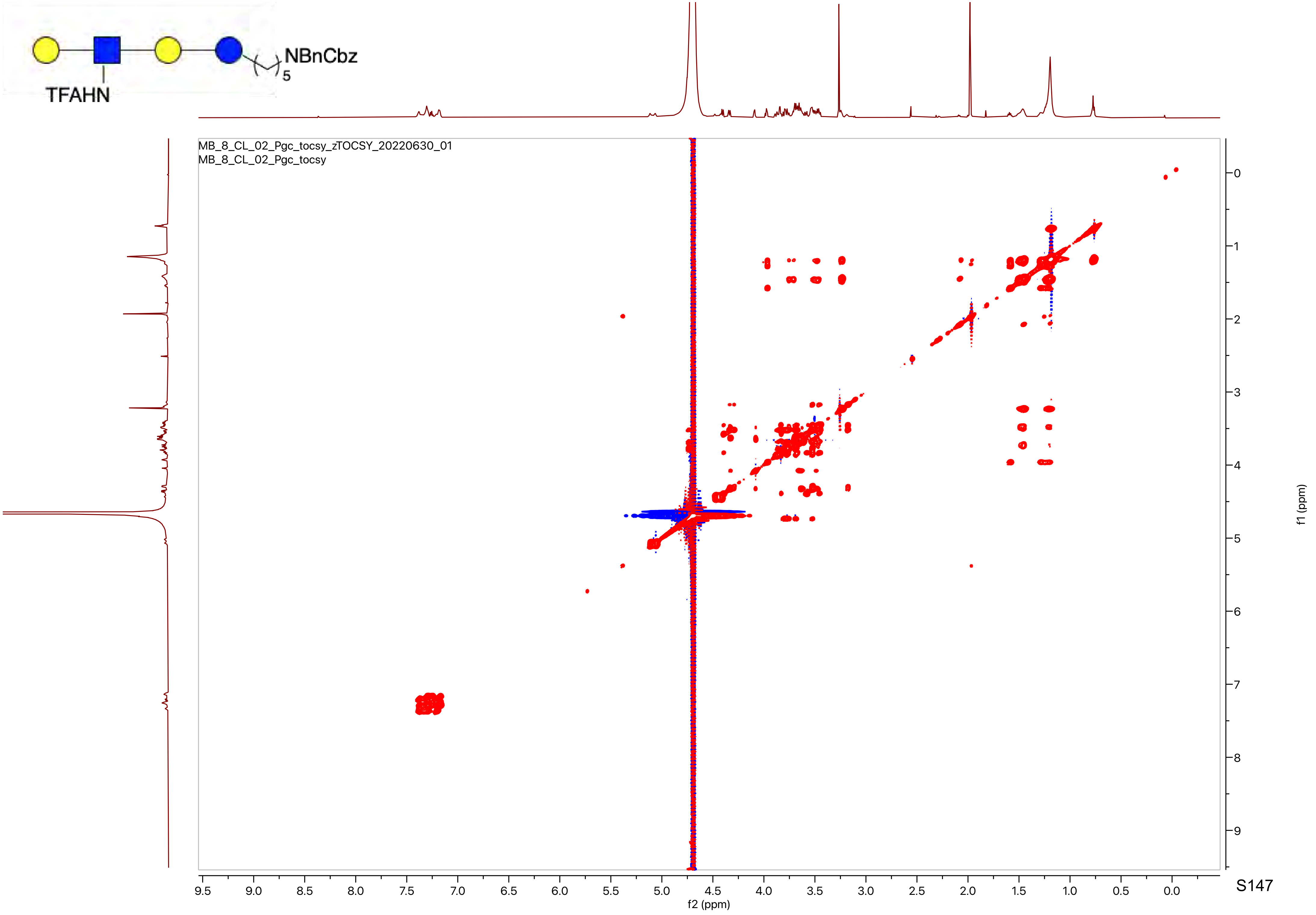

NBnCbz
5
TFAHN
MB_8_CL_02_Pgc_tocsy_zTOCSY_20220630_01
MB_8_CL_02_Pgc_tocsy
f2 (ppm)
f1 (ppm)

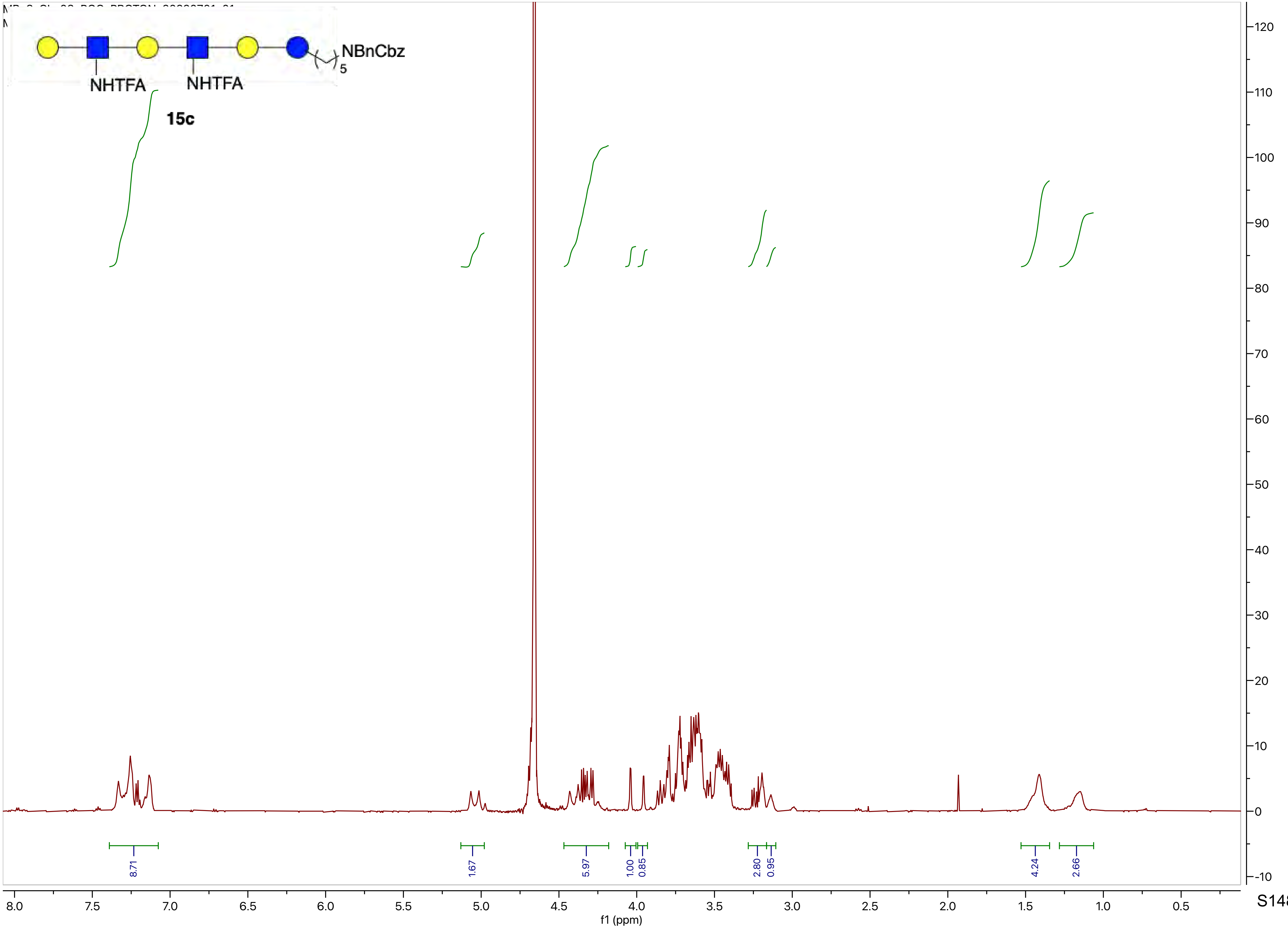
NBnCbz
5
NHTFA
NHTFA
15c
8.71
1.67
5.97
1.00
0.85
2.80
0.95
4.24
2.66
8.0
7.5
7.0
6.5
6.0
5.5
5.0
4.5
4.0
3.5
3.0
2.5
2.0
1.5
1.0
0.5
f1 (ppm)
120
110
100
90
80
70
60
50
40
30
20
10
0
-10

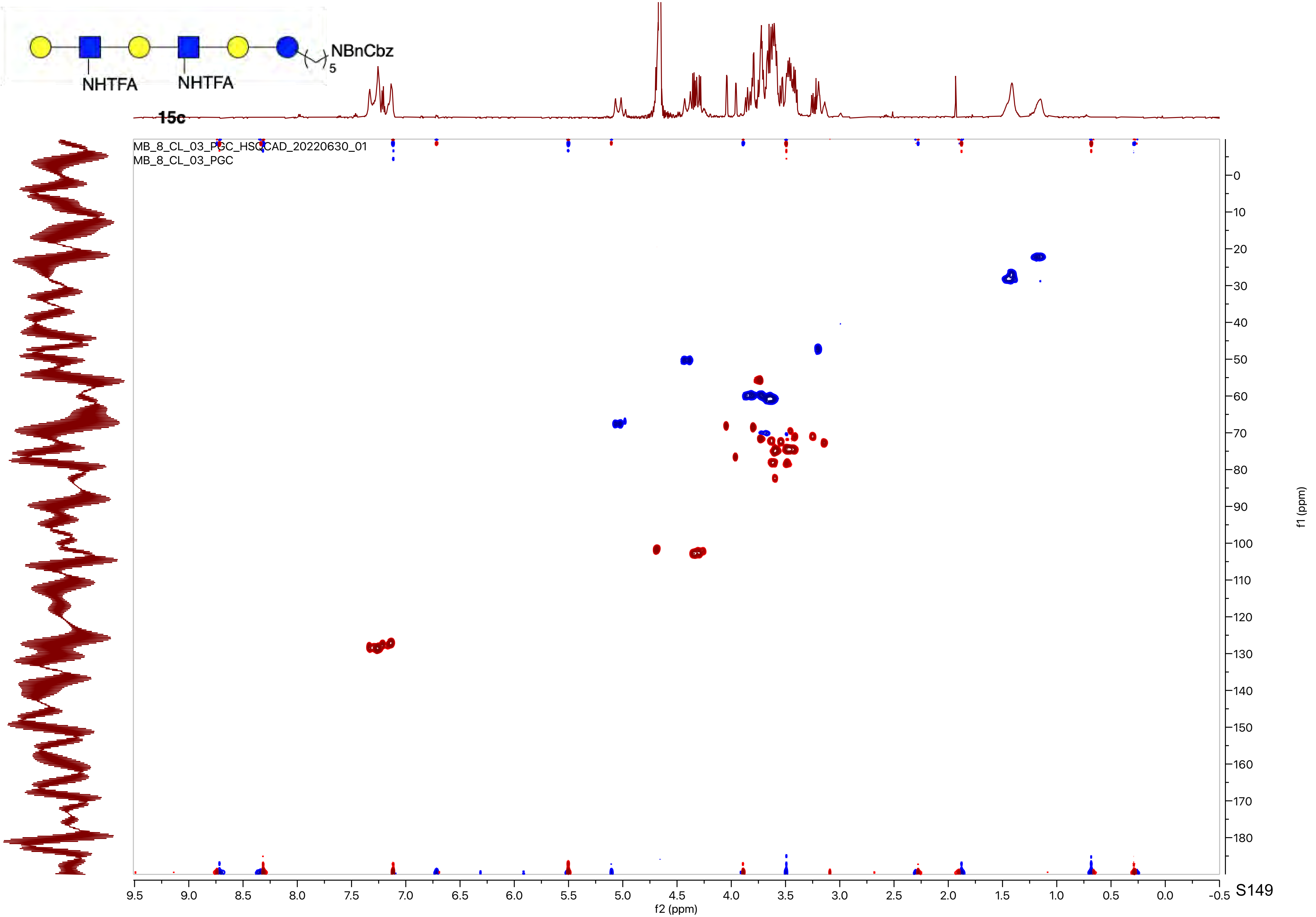

NBnCbz
5
NHTFA
NHTFA
15c
MB_8_CL_03_PGC_HSQCAD_20220630_01
MB_8_CL_03_PGC
f1 (ppm)
f2 (ppm)

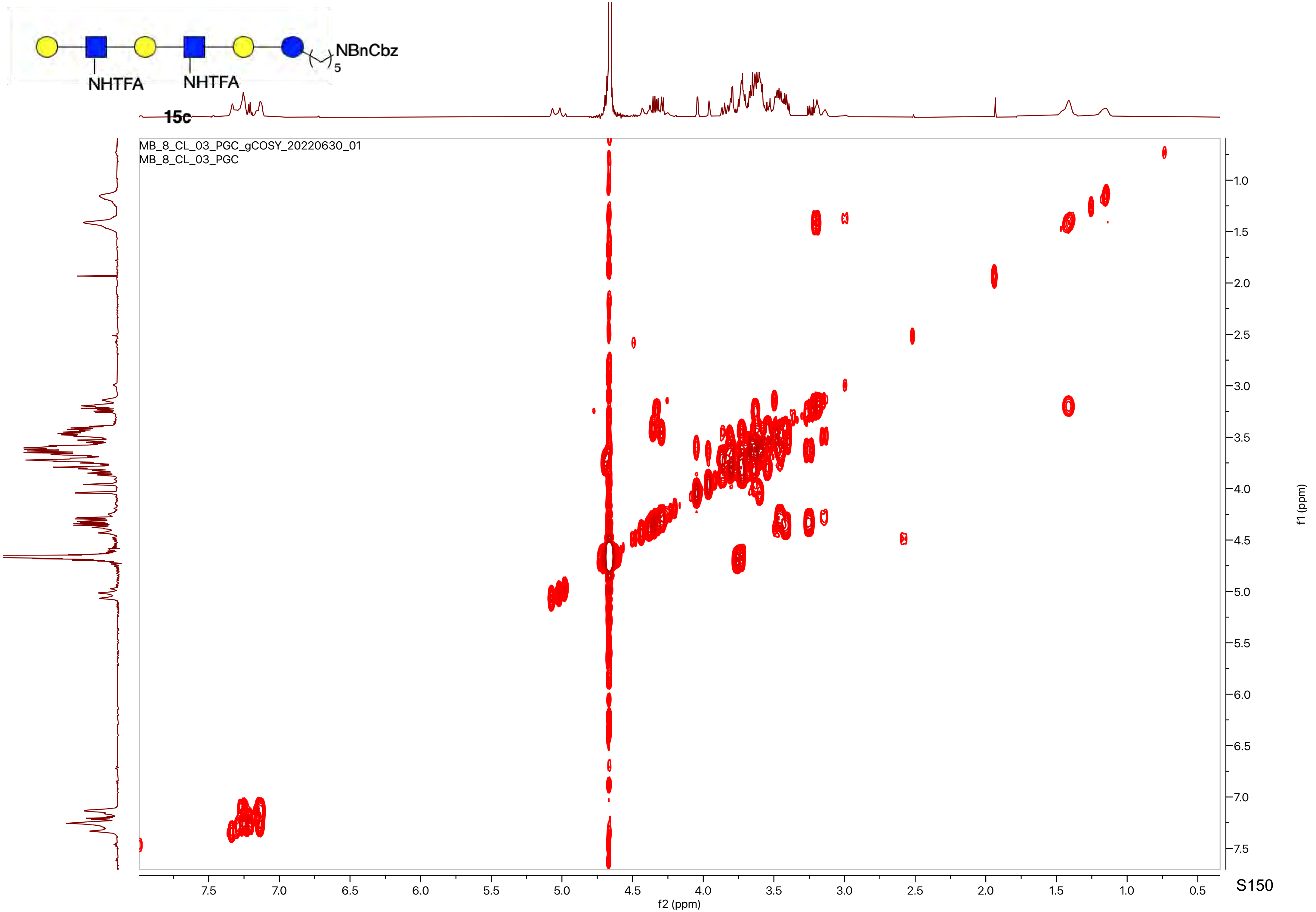
NBnCbz
5
NHTFA
NHTFA
15c
MB_8_CL_03_PGC_gCOSY_20220630_01
MB_8_CL_03_PGC
f2 (ppm)
f1 (ppm)

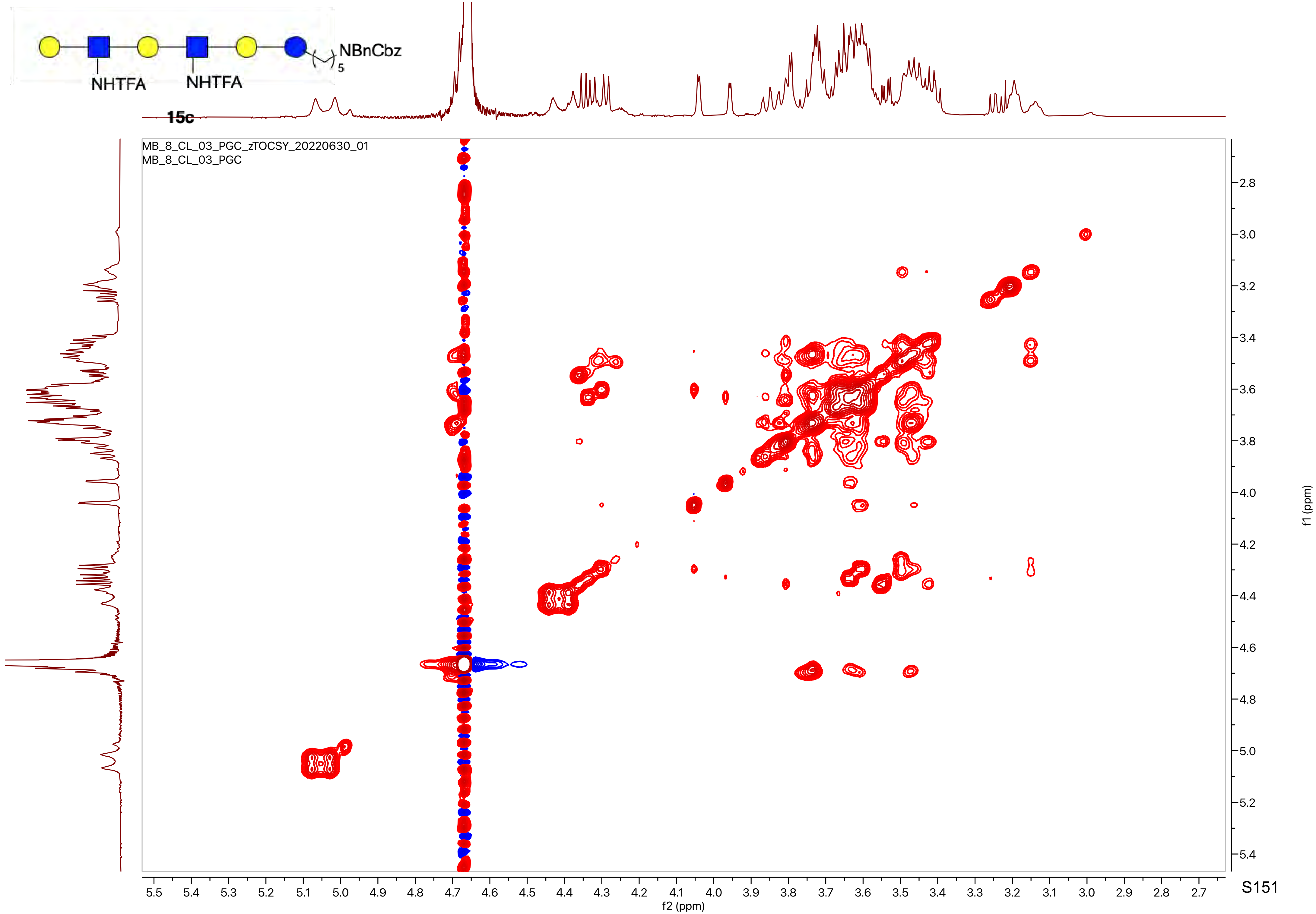
NBnCbz
5
NHTFA
NHTFA
15c
MB_8_CL_03_PGC_zTOCSY_20220630_01
MB_8_CL_03_PGC
f2 (ppm)
f1 (ppm)

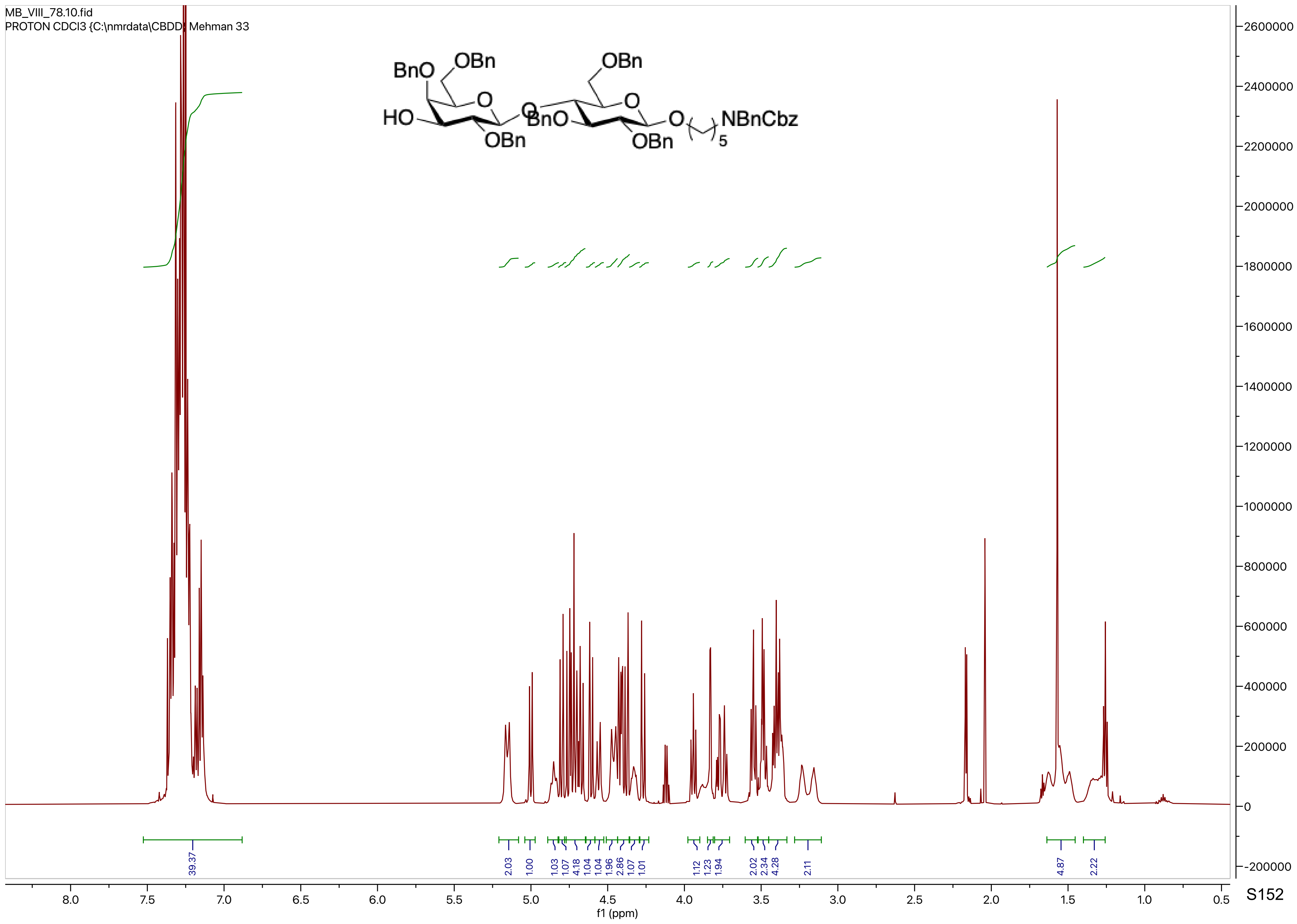
MB_VIII_78.10.fid
PROTON CDCl3 {C:\nmrdata\CBDD} Mehman 33
BnO
OBn
HO
O
OBn
BnO
NBnCbz
5
39.37
2.03
1.00
1.03
1.07
4.18
1.04
1.04
1.96
2.86
1.07
1.01
1.12
1.23
1.94
2.02
2.34
4.28
2.11
4.87
2.22
f1 (ppm)

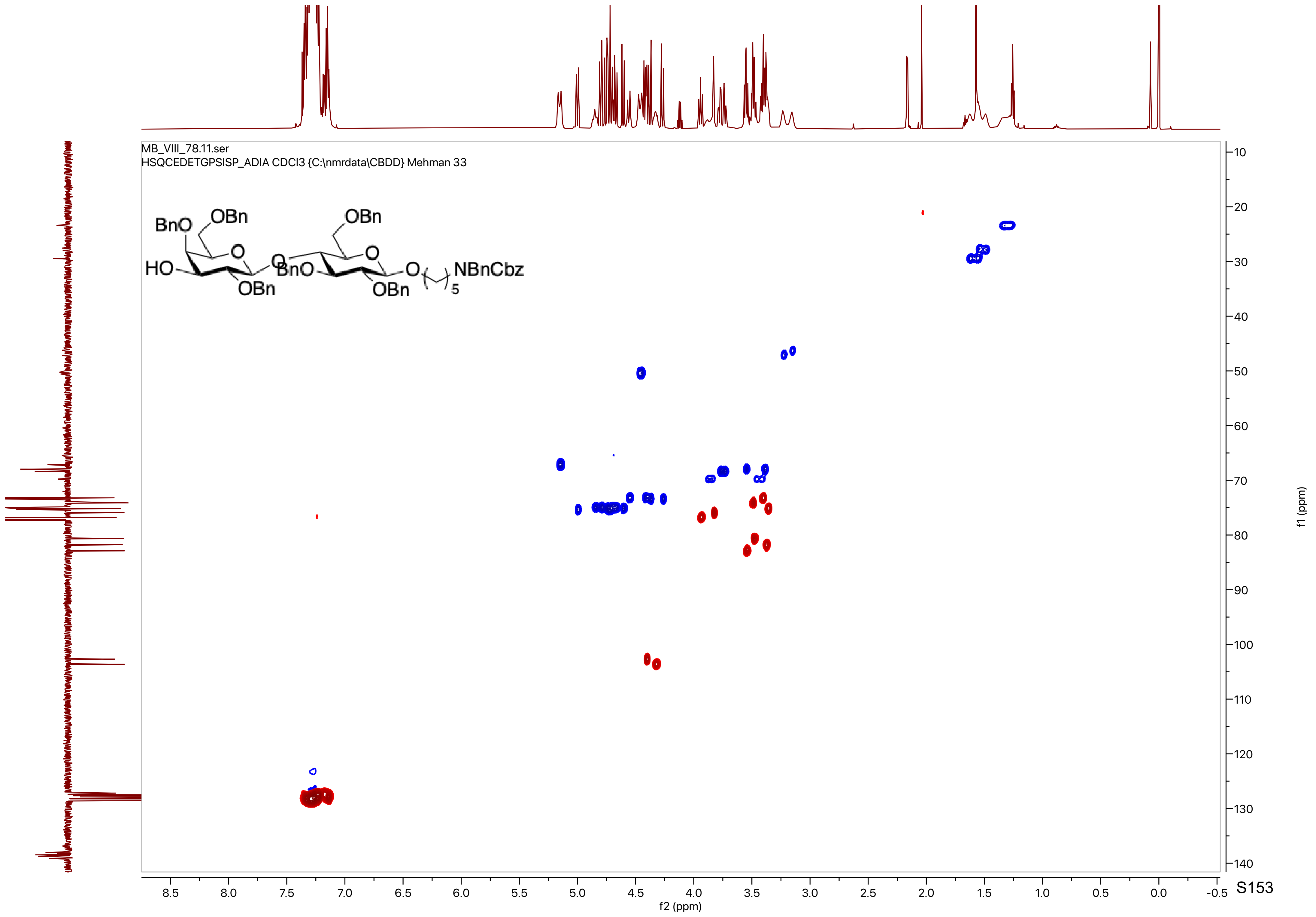

MB_VIII_78.11.ser
HSQCEDETGPSISP_ADIA CDCl3 {C:\nmrdata\CBDD} Mehman 33
BnO
OBn
O
HO
OBn
O
BnO
OBn
O
O
OBn
NBnCbz
5
f2 (ppm)
f1 (ppm)

MB_VIII_78.13.fid
C13APT CDCl3 {C:\nmrdata\CBDD} Mehman 33

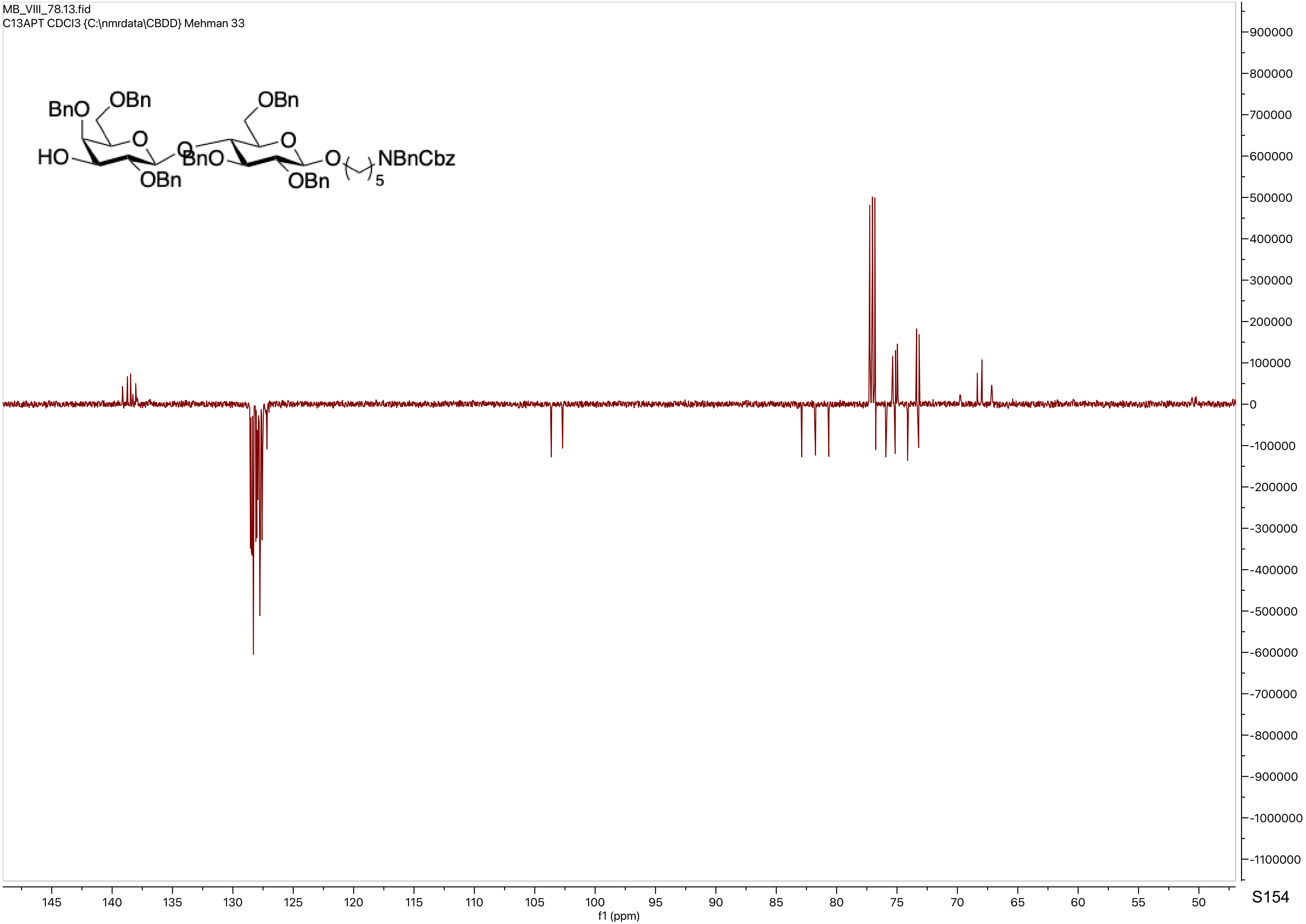

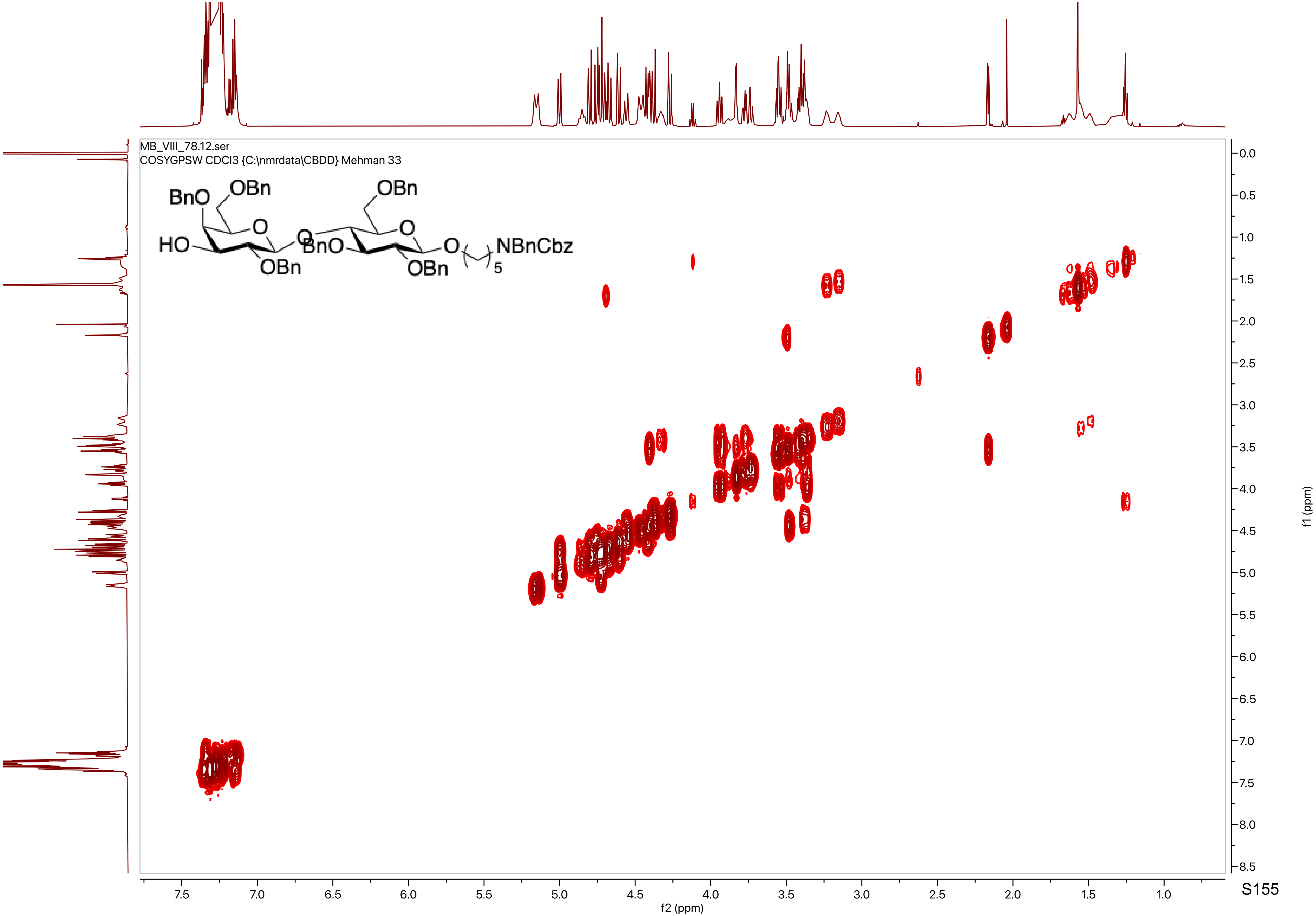
MB_VIII_78.12.ser
COSYGPSW CDCl3 {C:\nmrdata\CBDD} Mehman 33
BnO
OBn
O
HO
OBn
O
OBn
O
BnO
OBn
O
NBnCbz
5
f2 (ppm)
f1 (ppm)

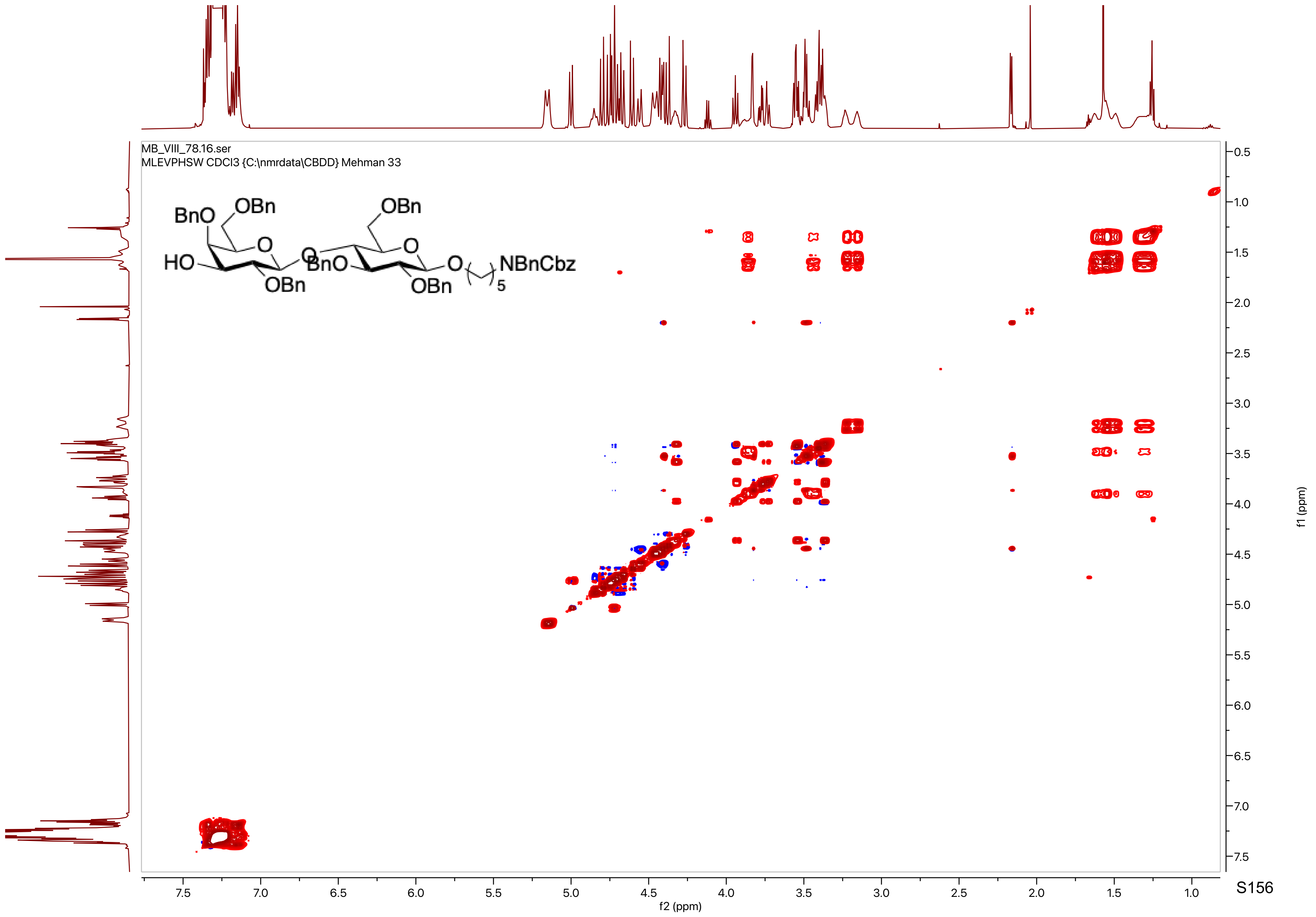
MB_VIII_78.16.ser
MLEVPHSW CDCl3 {C:\nmrdata\CBDD} Mehman 33
BnO
OBn
O
HO
OBn
O
BnO
OBn
O
O
NBnCbz
5
OBn
f2 (ppm)
f1 (ppm)

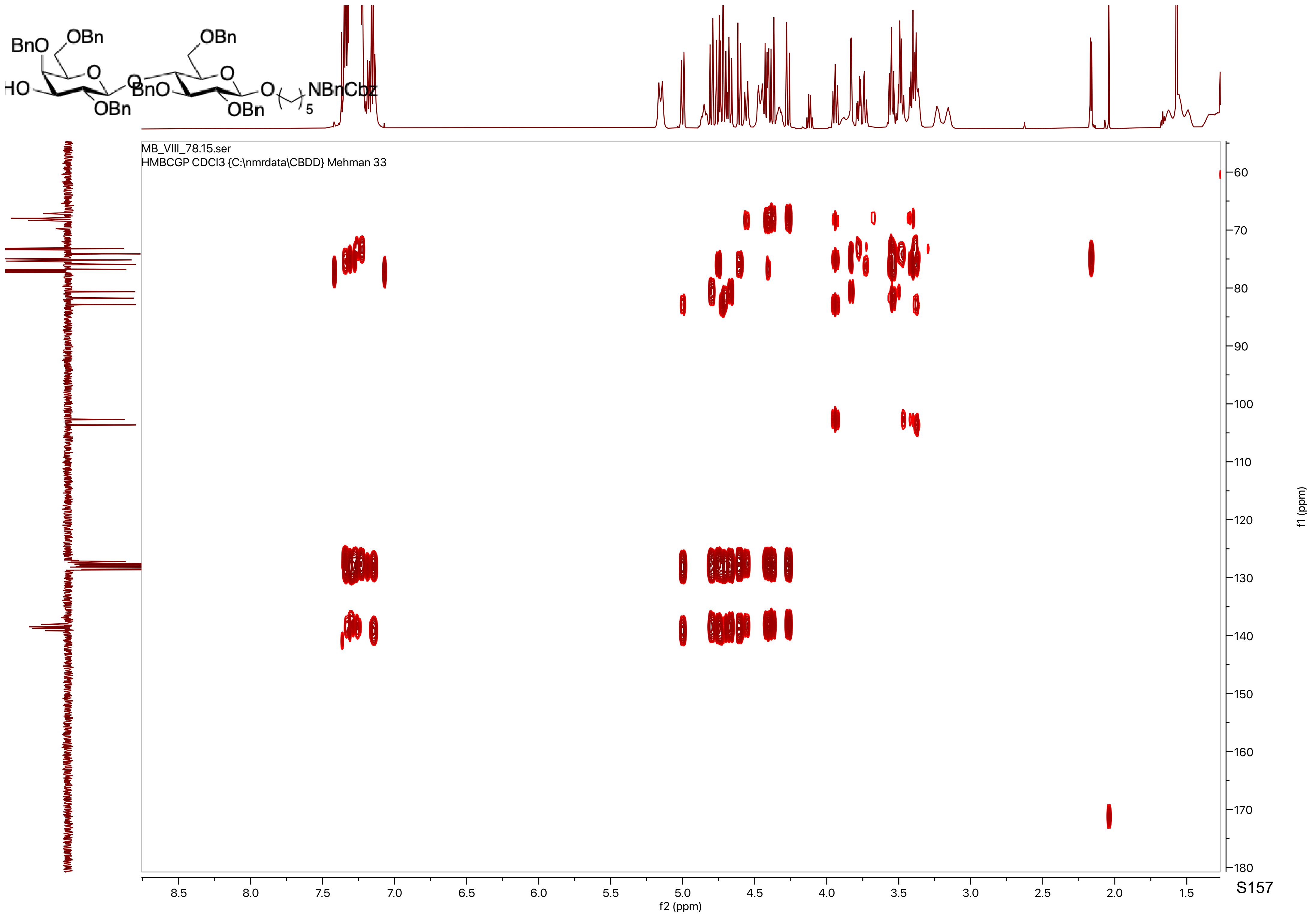
BnO
OBn
HO
O
O
OBn
BnO
O
OBn
O
NBnCbz
5
MB_VIII_78.15.ser
HMBCGP CDCl3 {C:\nmrdata\CBDD} Mehman 33
f2 (ppm)
f1 (ppm)

MB_VIII_80.10.fid
PROTON CDCl3 {C:\nmrdata\CBDD} Mehman 39

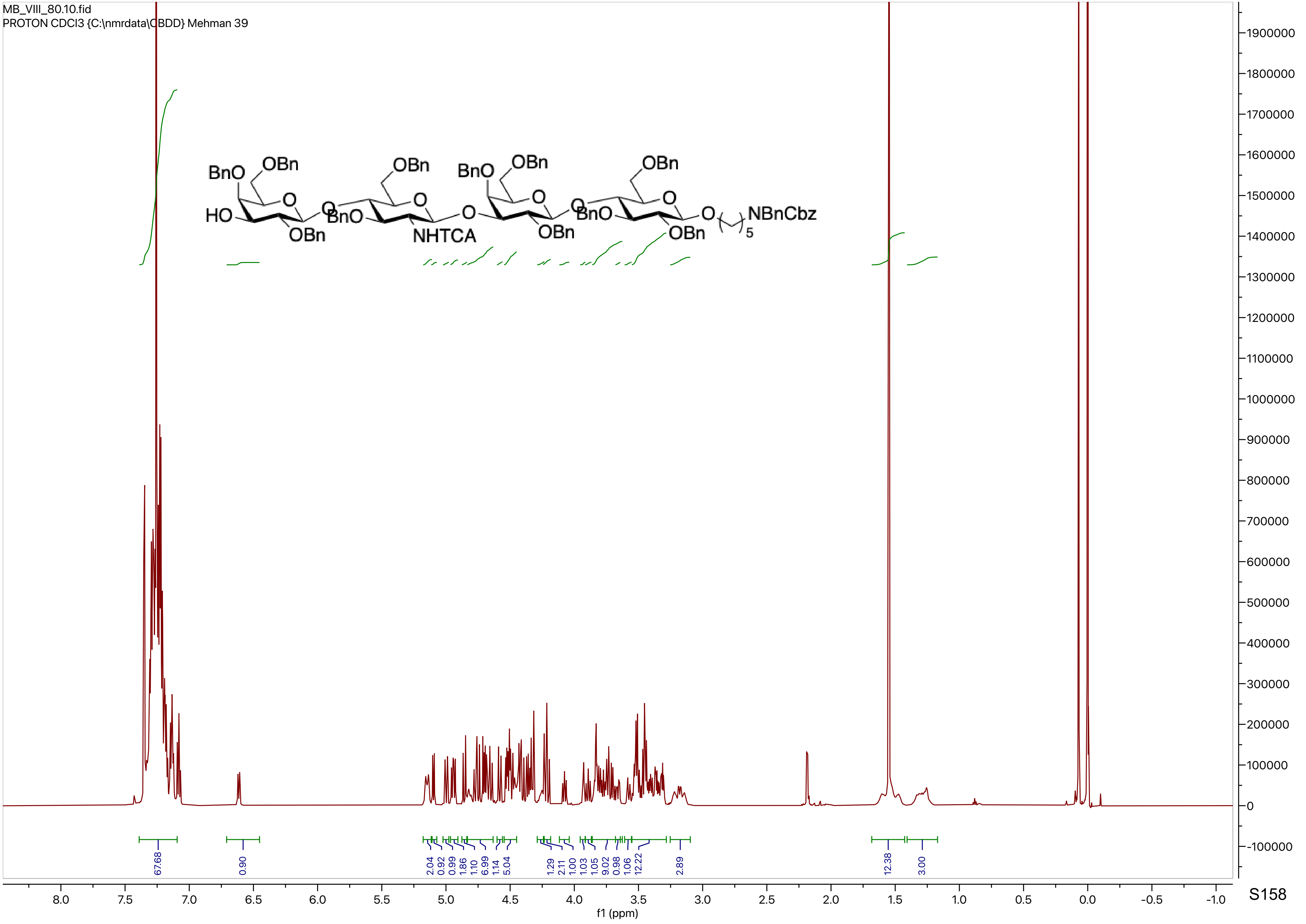

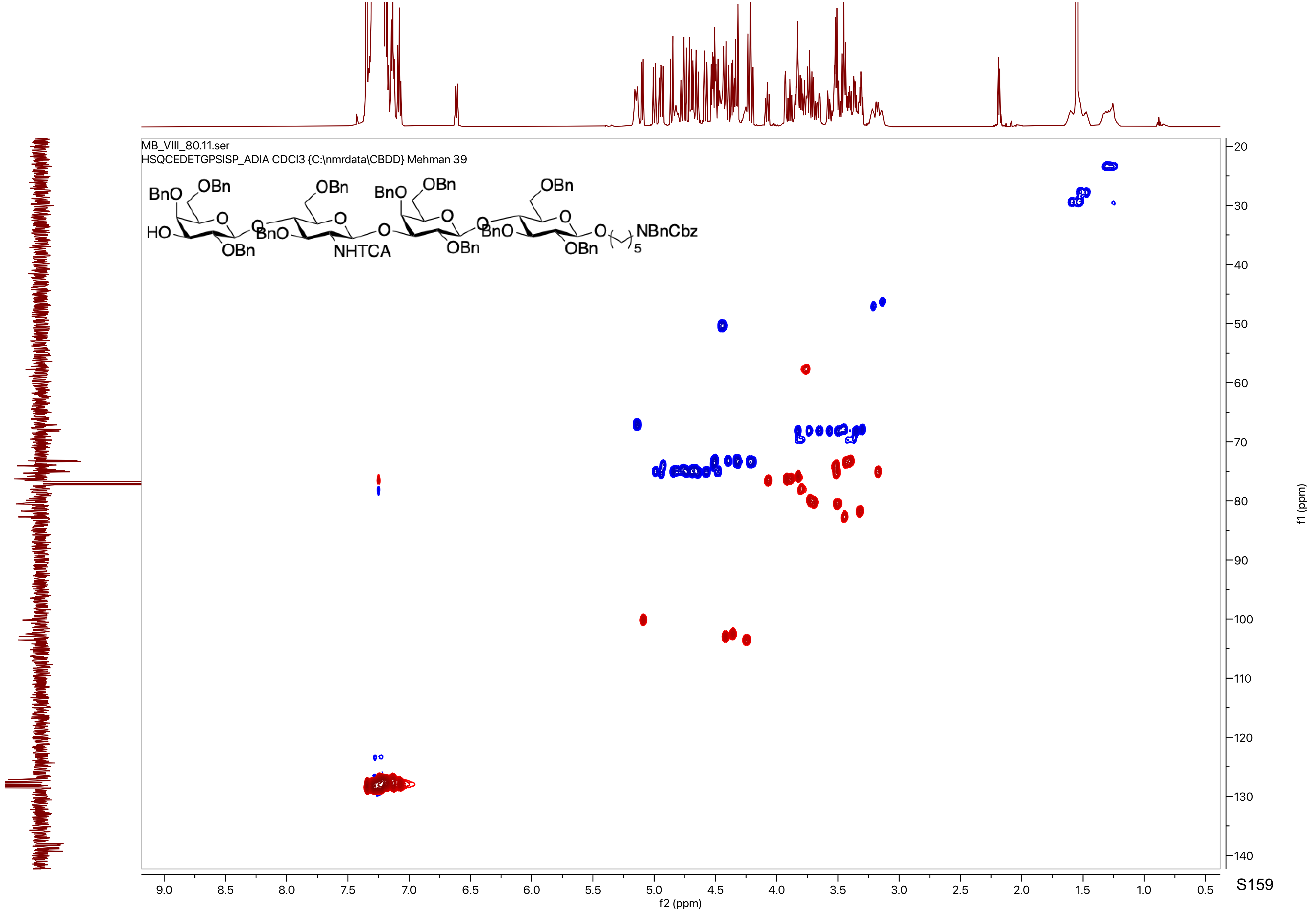
MB_VIII_80.11.ser
HSQCEDETGPSISP_ADIA CDCl3 {C:\nmrdata\CBDD} Mehman 39
BnO
OBn
HO
O
OBn
BnO
NHTCA
OBn
NBnCbz
5
f2 (ppm)
f1 (ppm)

MB_VIII_80.13.fid
C13APT CDCl3 {C:\nmrdata\CBDD} Mehman 39

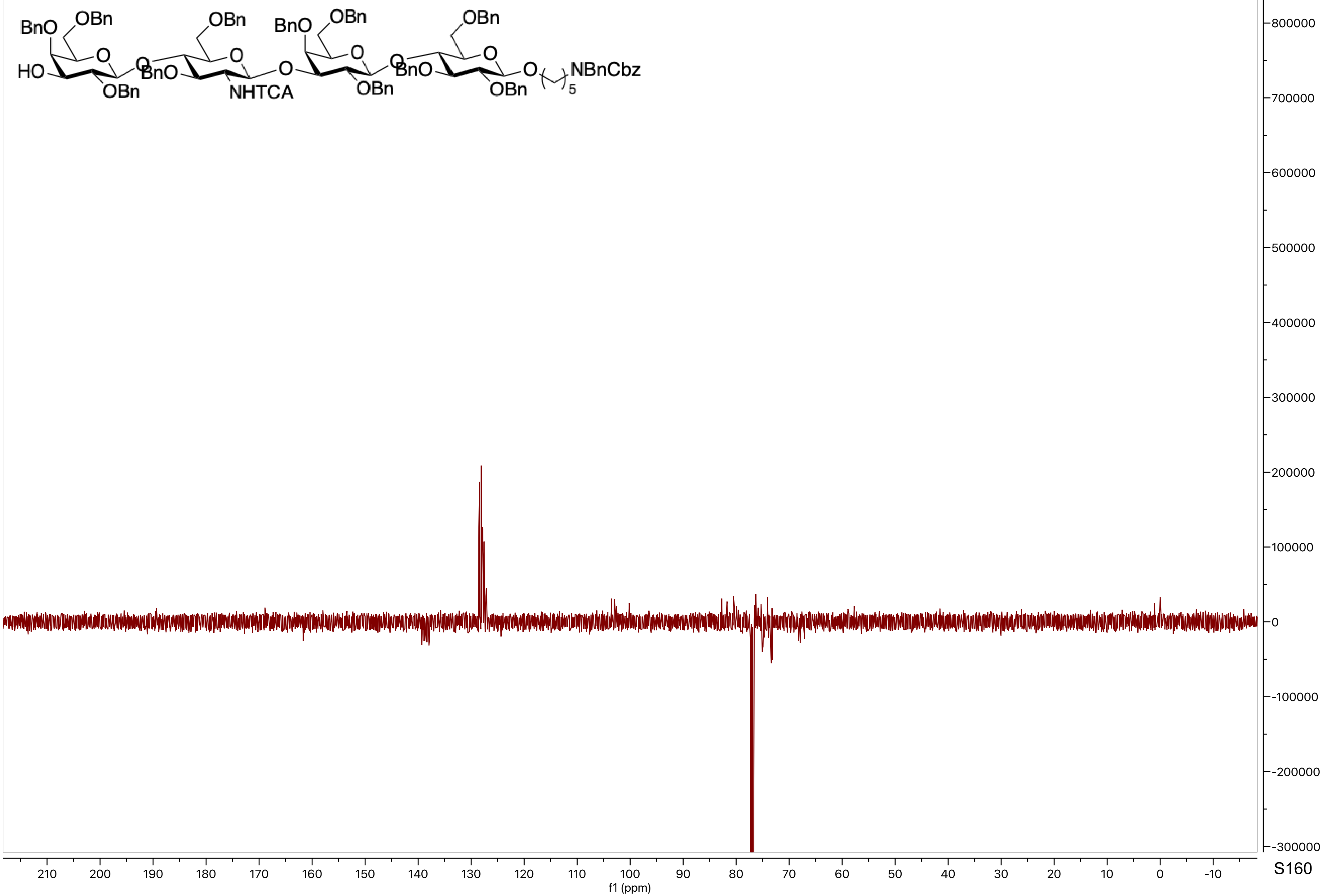

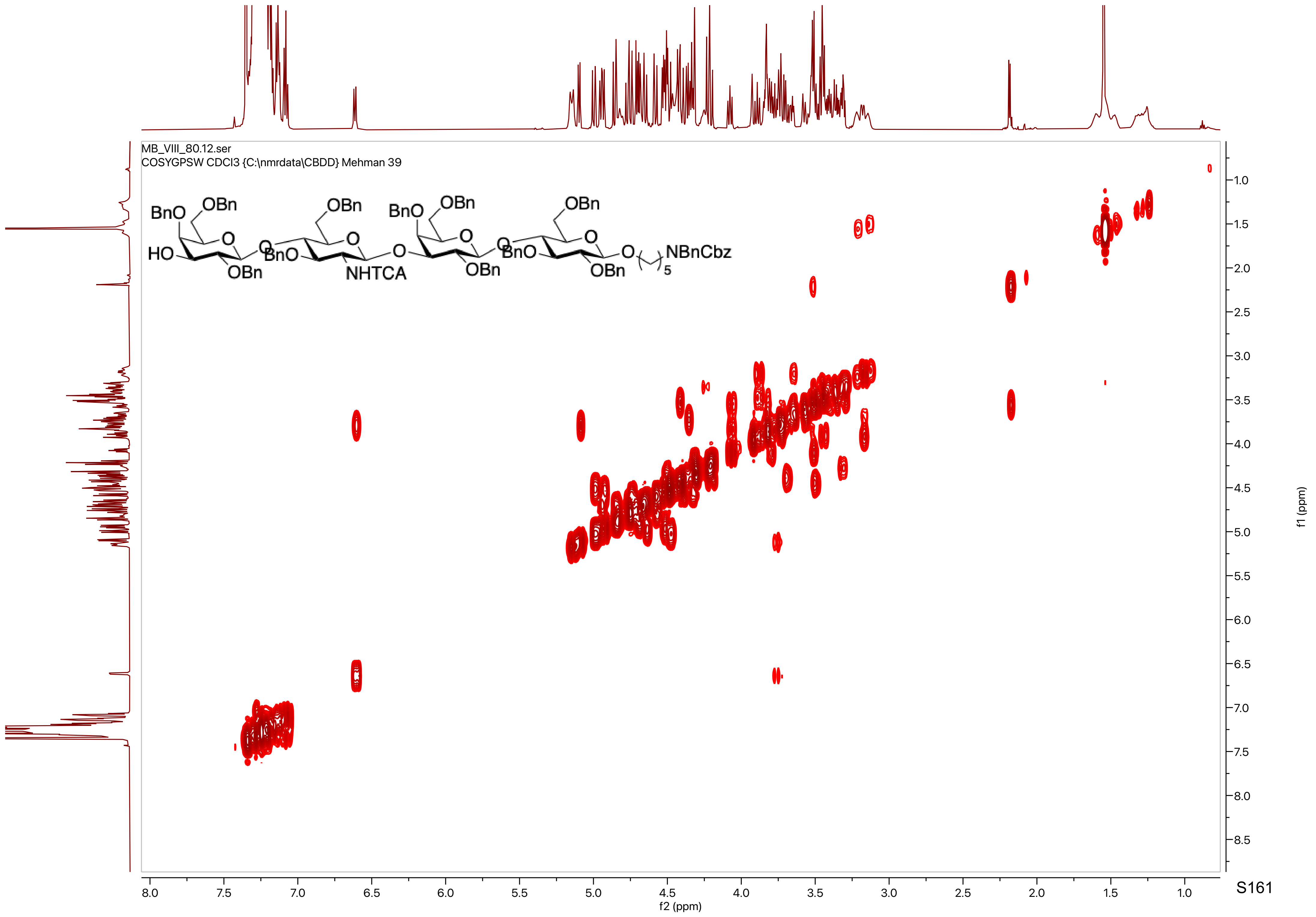

MB_VIII_80.12.ser
COSYGPSW CDCl3 {C:\nmrdata\CBDD} Mehman 39
BnO
OBn
HO
OBn
BnO
OBn
NHTCA
BnO
OBn
BnO
OBn
OBn
NBnCbz
5
f2 (ppm)
f1 (ppm)

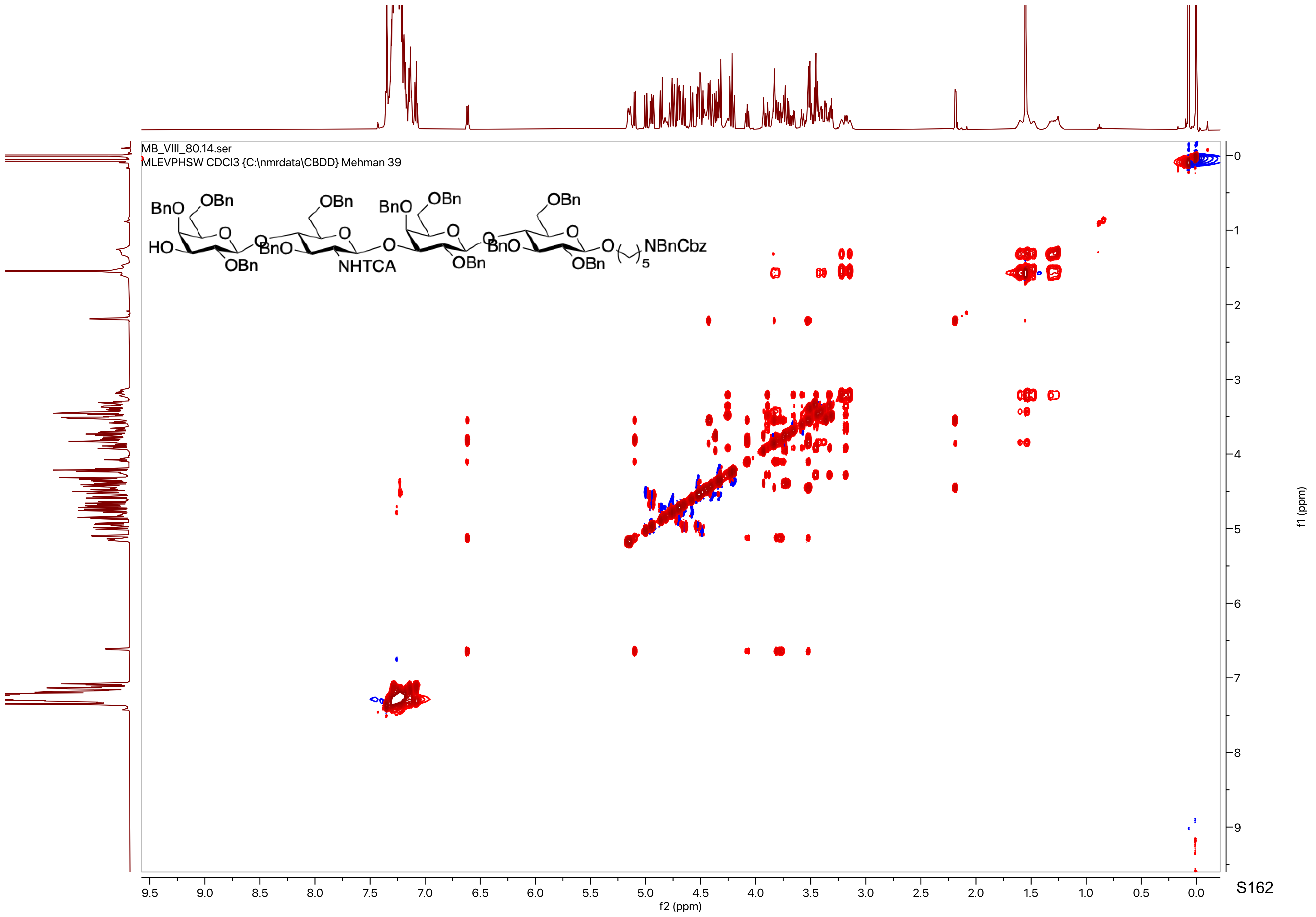

MB_VIII_80.14.ser
MLEVPHSW CDCl3 {C:\nmrdata\CBDD} Mehman 39
BnO
OBn
HO
O
OBn
BnO
NHTCA
NBnCbz
5
f2 (ppm)
f1 (ppm)

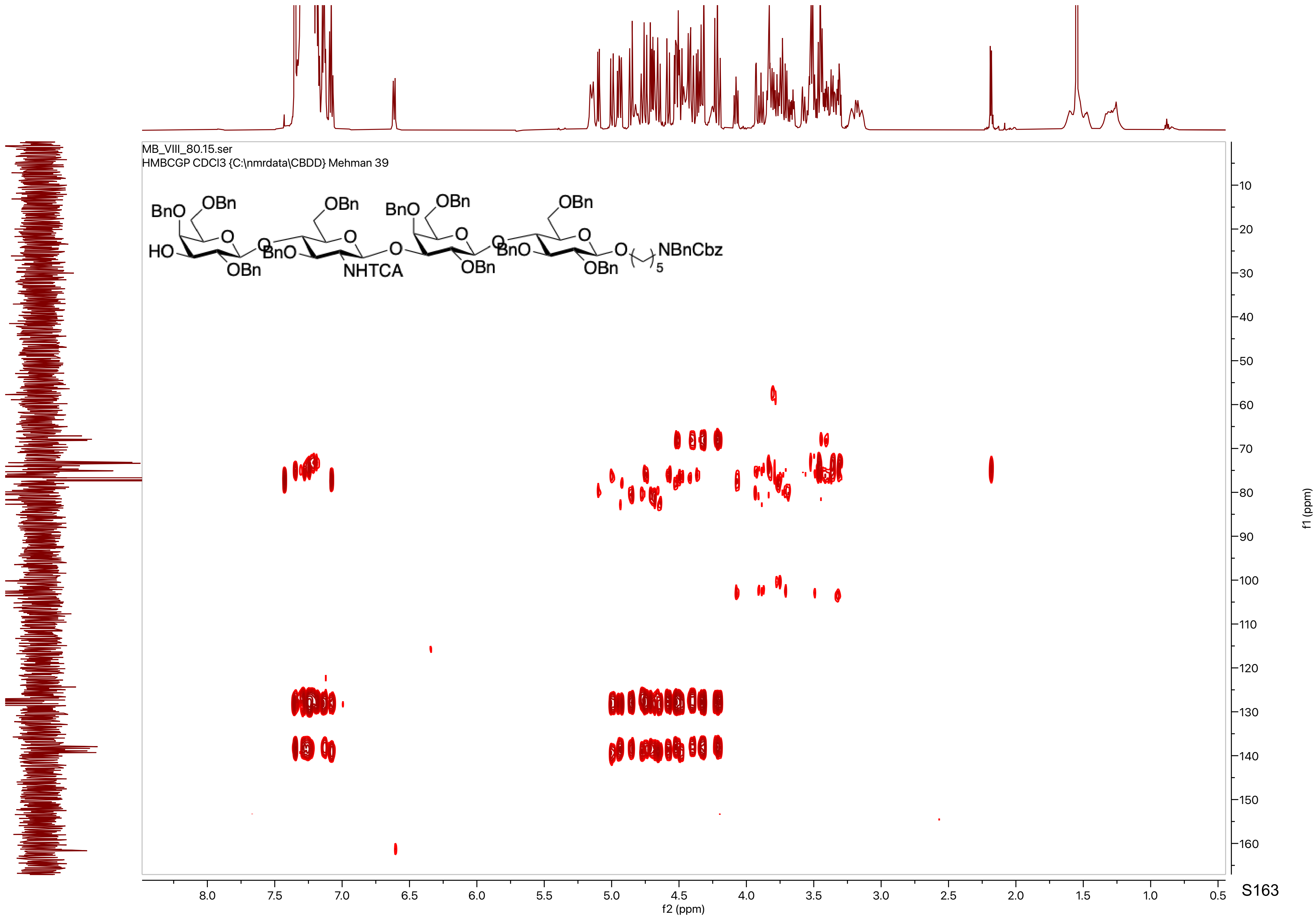
MB_VIII_80.15.ser
HMBCGP CDCl3 {C:\nmrdata\CBDD} Mehman 39
BnO
OBn
HO
OBn
BnO
OBn
NHTCA
BnO
OBn
OBn
BnO
OBn
OBn
NBnCbz
5
f2 (ppm)
f1 (ppm)

MB_VIII_82.10.fid
PROTON CDCl3 {C:\nmrdata\CBDD} Mehman 13

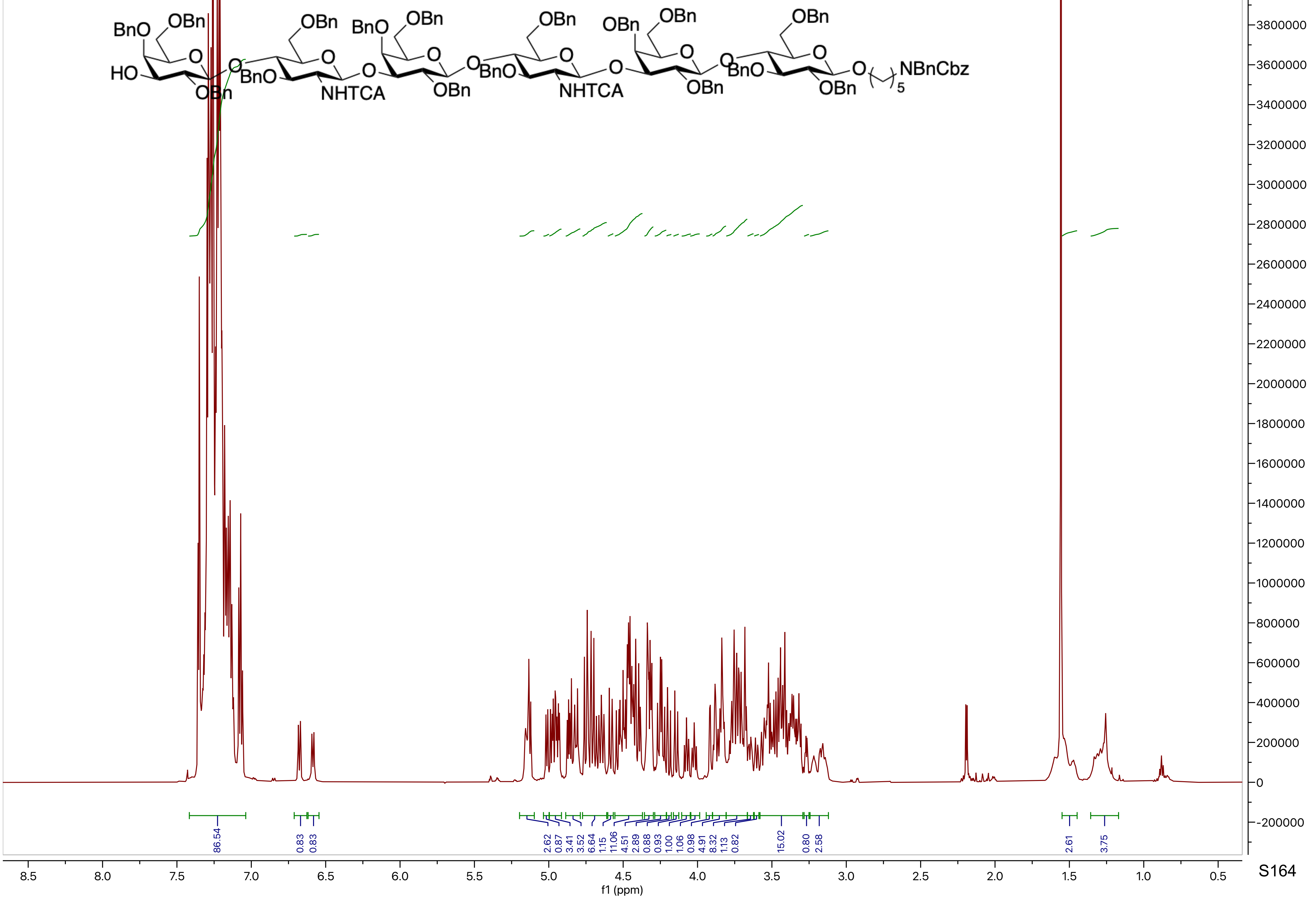

MB_VIII_82.15.fid
C13APT CDCl3 {C:\nmrdata\CBDD} Mehman 13

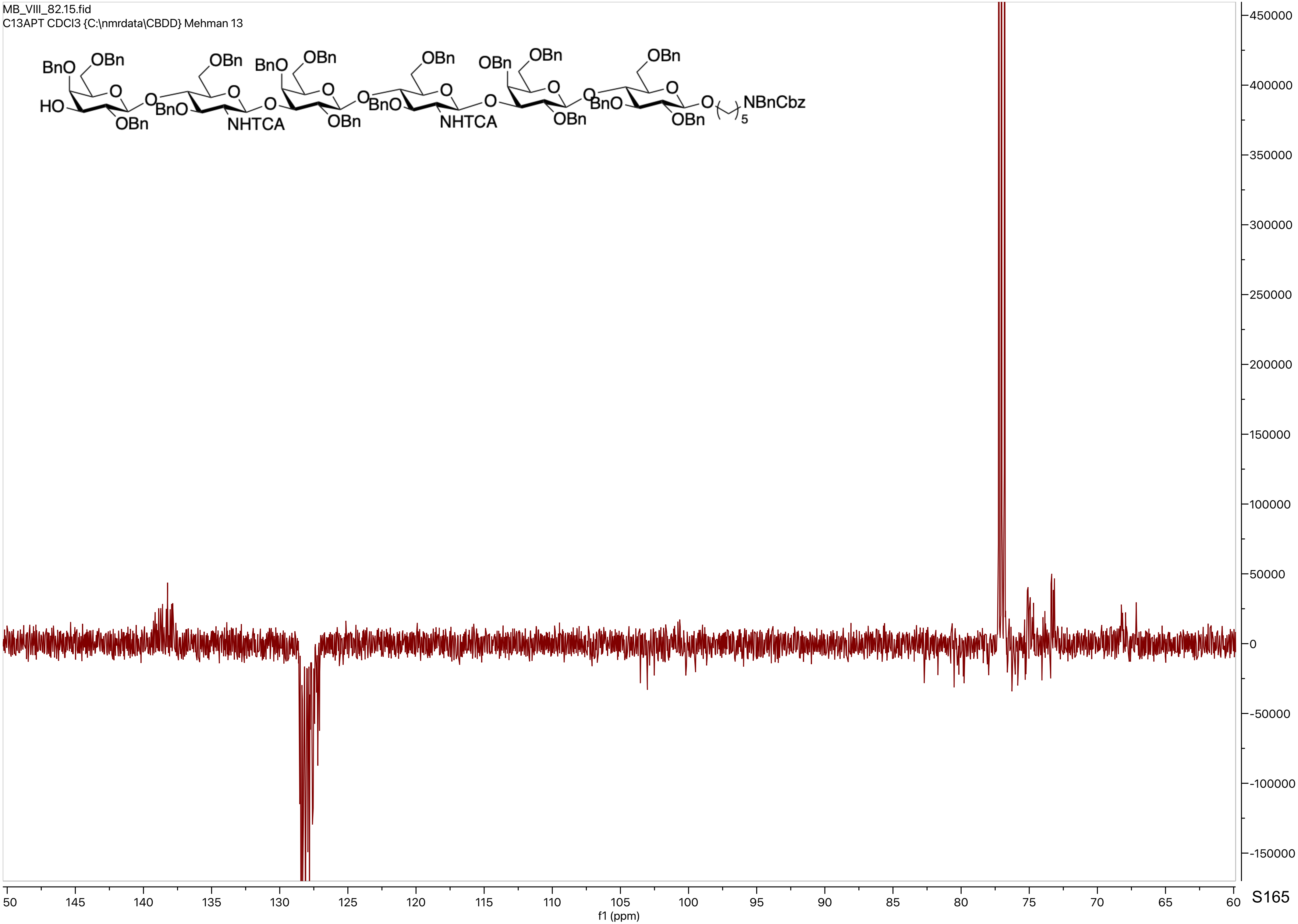

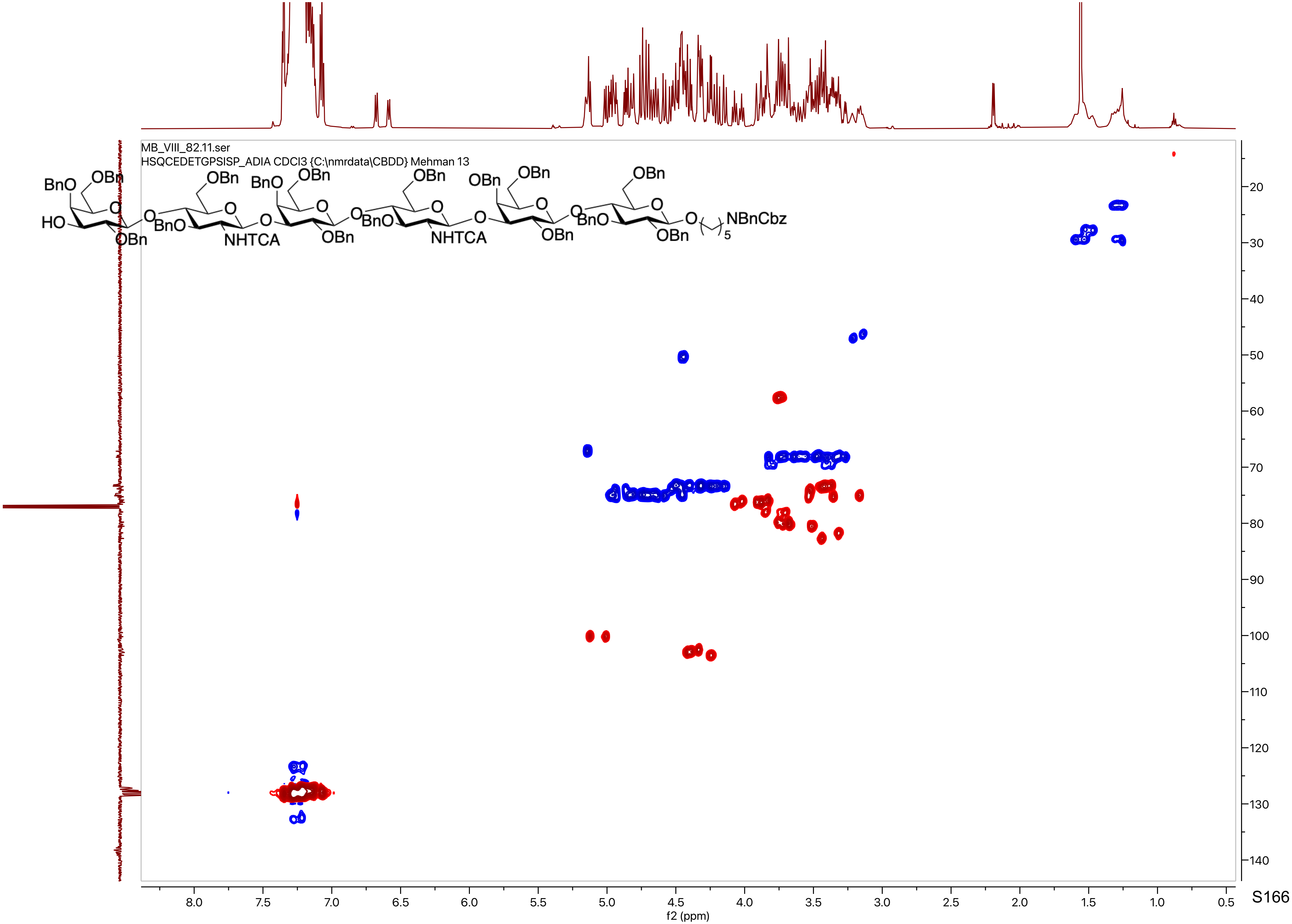
MB_VIII_82.11.ser
HSQCEDETGPSISP_ADIA CDCl3 {C:\nmrdata\CBDD} Mehman 13
f2 (ppm)
f1 (ppm)

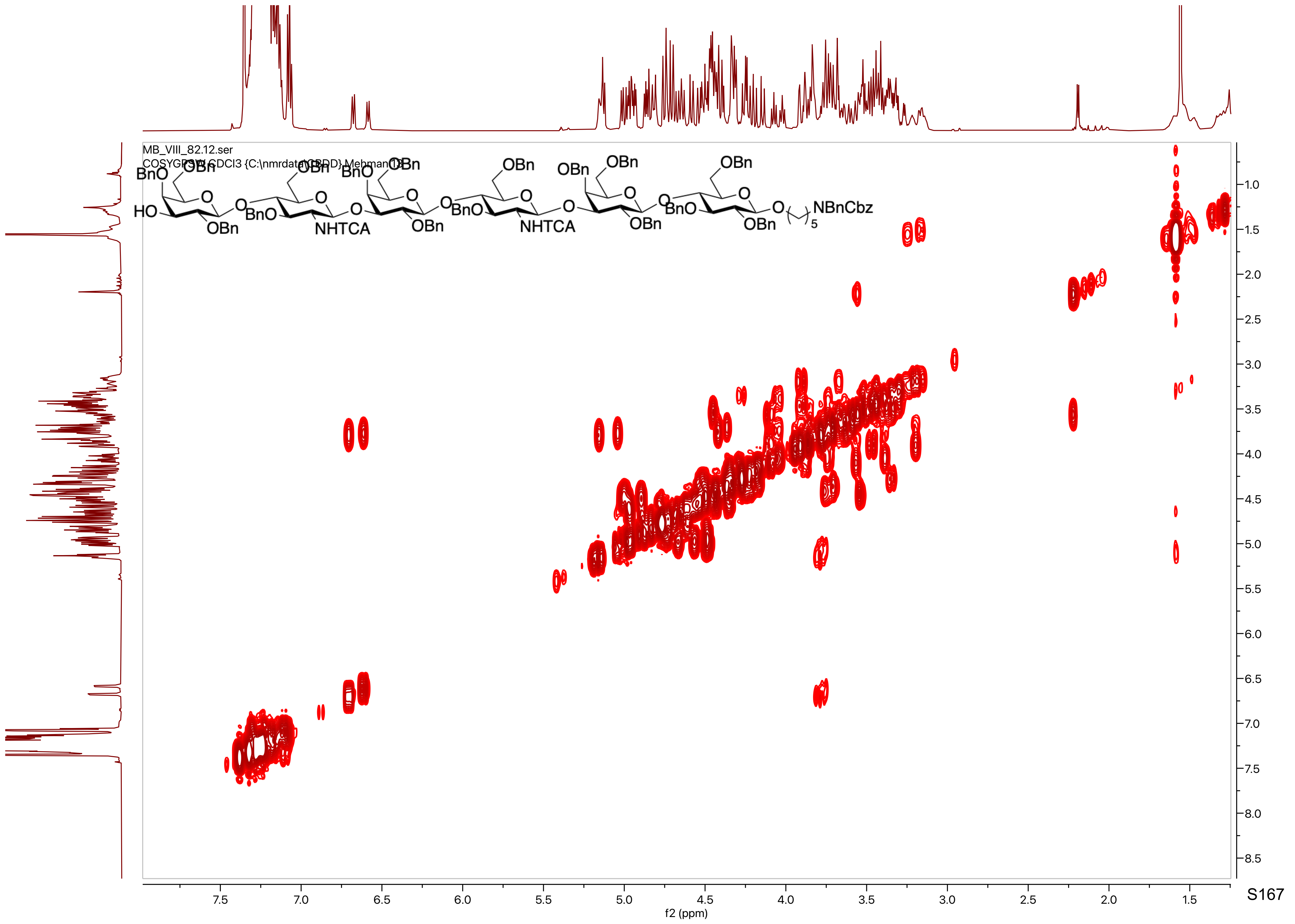
MB_VIII_82.12.ser
COSYGPSW CDCl3 {C:\nmrdata\CBDD} Mehman 13
BnO
OBn
HO
O
BnO
NHTCA
NBnCbz
f2 (ppm)
f1 (ppm)
7.5
7.0
6.5
6.0
5.5
5.0
4.5
4.0
3.5
3.0
2.5
2.0
1.5
1.0
8.0
8.5

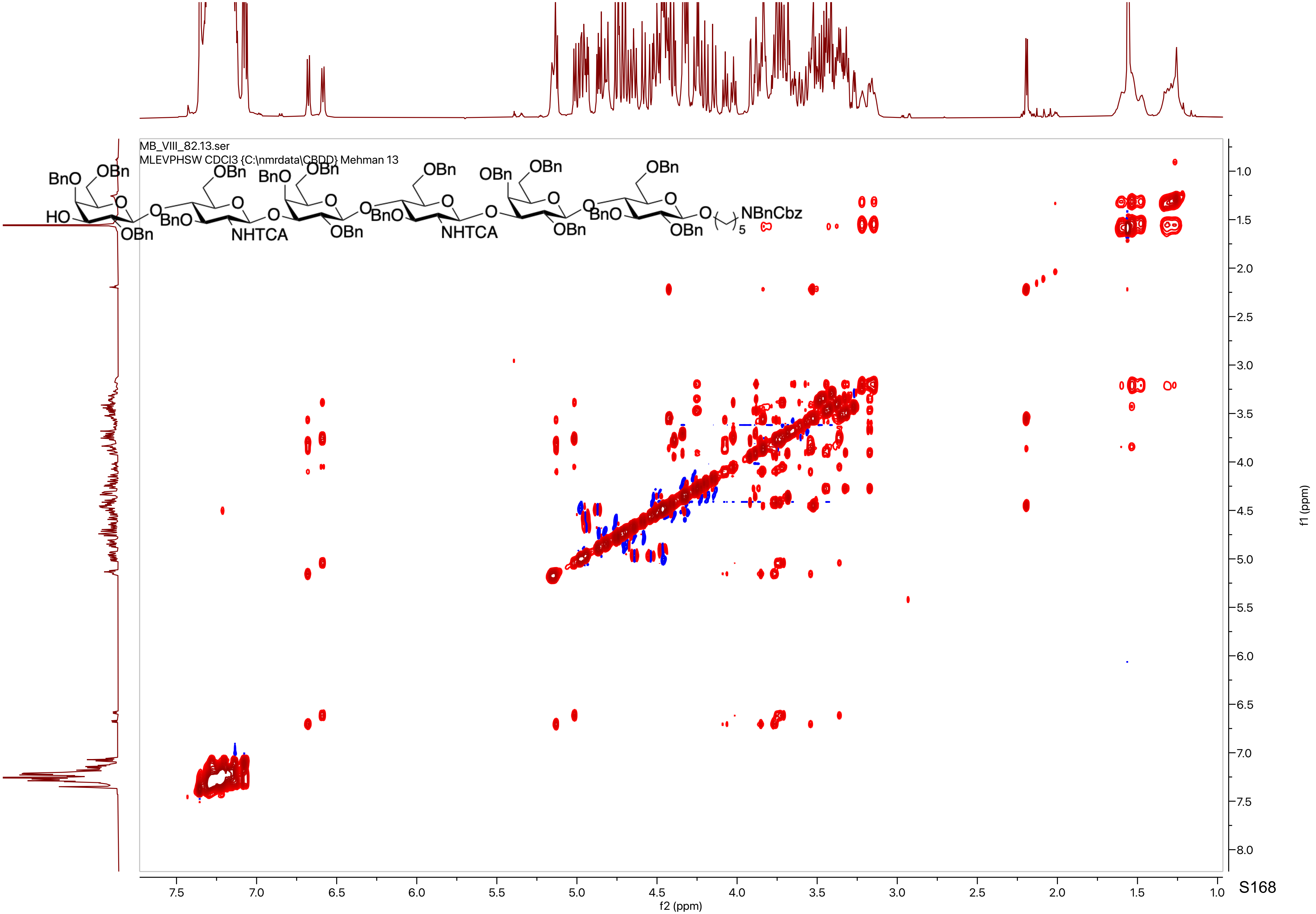

MB_VIII_82.13.ser
MLEVPHSW CDCl3 {C:\nmrdata\CBDD} Mehman 13
BnO
OBn
HO
OBn
BnO
NHTCA
NBnCbz
f2 (ppm)
f1 (ppm)

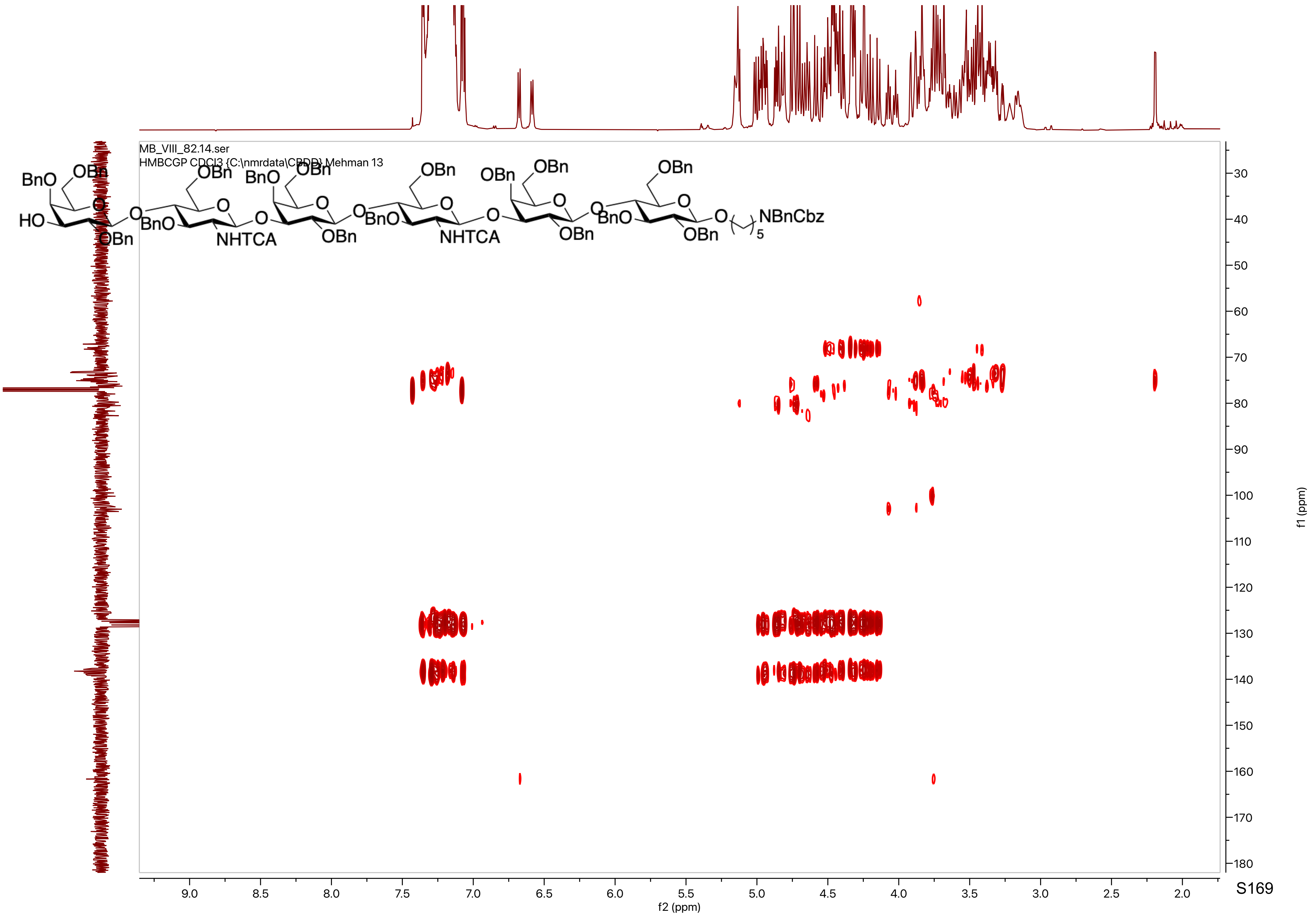
MB_VIII_82.14.ser
HMBCGP CDCl3 {C:\nmrdata\CBDD} Mehman 13
BnO
OBn
HO
O
OBn
BnO
NHTCA
OBn
NHTCA
NBnCbz
5
f2 (ppm)
f1 (ppm)
9.0
8.5
8.0
7.5
7.0
6.5
6.0
5.5
5.0
4.5
4.0
3.5
3.0
2.5
2.0
30
40
50
60
70
80
90
100
110
120
130
140
150
160
170
180

S169

MB_VIII_44.10.fid
PROTON CDCl3 {C:\nmrdata\[illegible]} Mehman 19

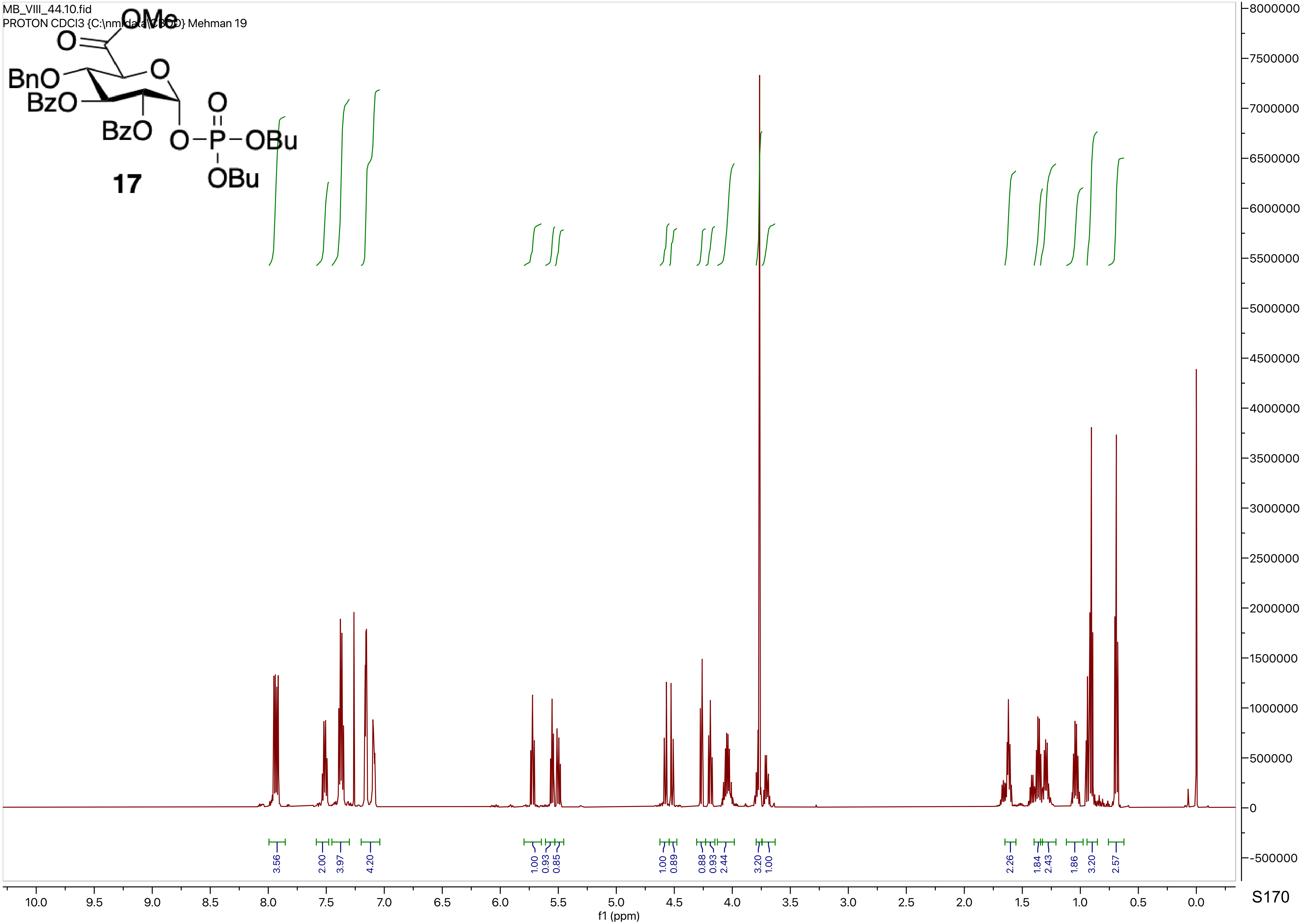

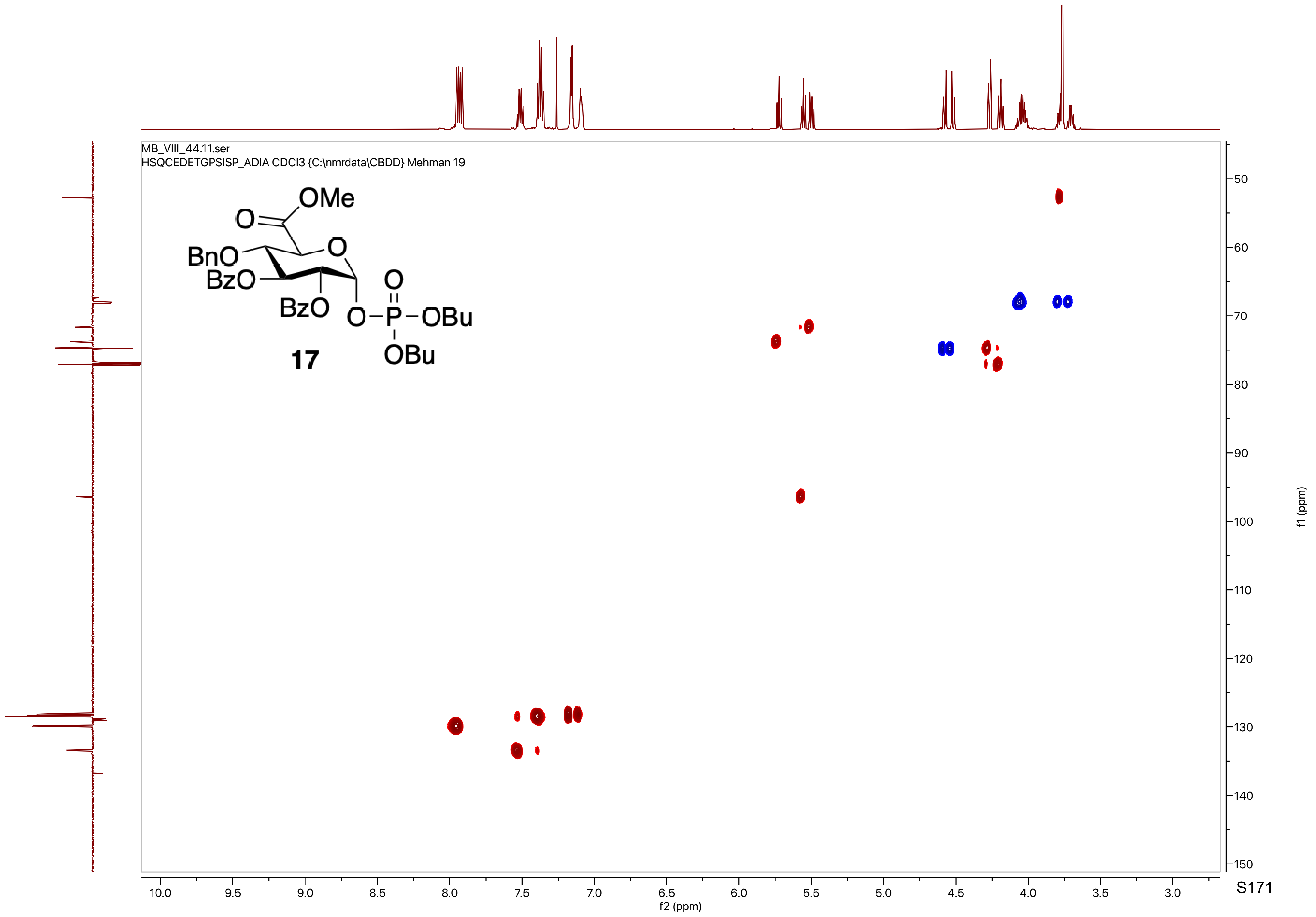

MB_VIII_44.11.ser
HSQCEDETGPSISP_ADIA CDCl3 {C:\nmrdata\CBDD} Mehman 19
OMe
O
BnO
BzO
BzO
O
O
O–P–OBu
OBu
17
f2 (ppm)
f1 (ppm)

MB_VIII_44.12.fid
C13APT CDCl3 {C:\nmrdata\CBDD} Mehman 19

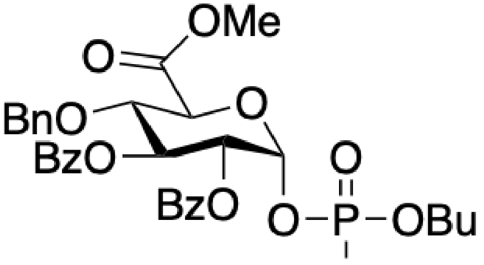


**17**

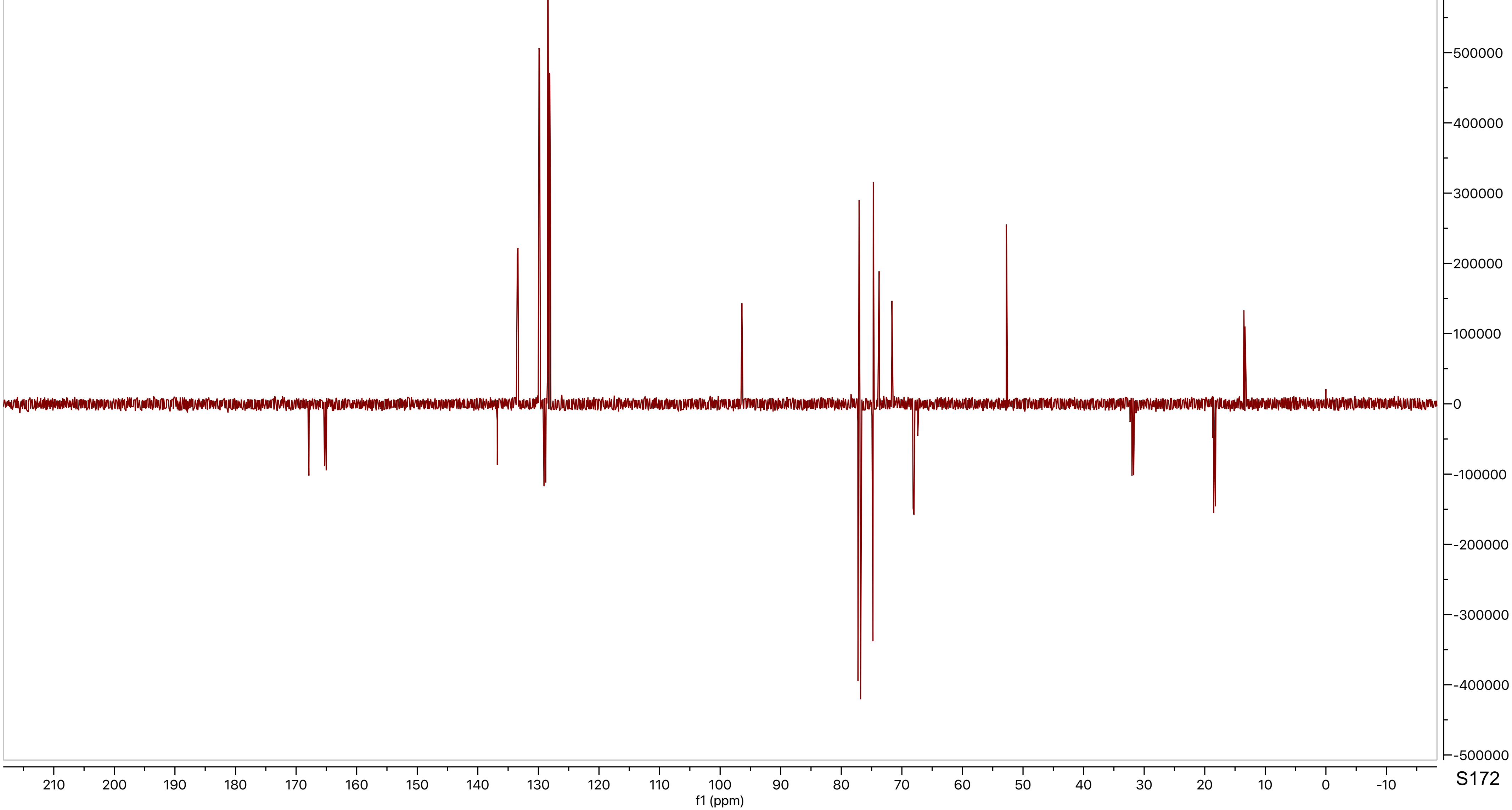

MB_VIII_44.13.ser
COSYGPSW CDCl3 {C:\nmrdata\CBDD} Mehman 19

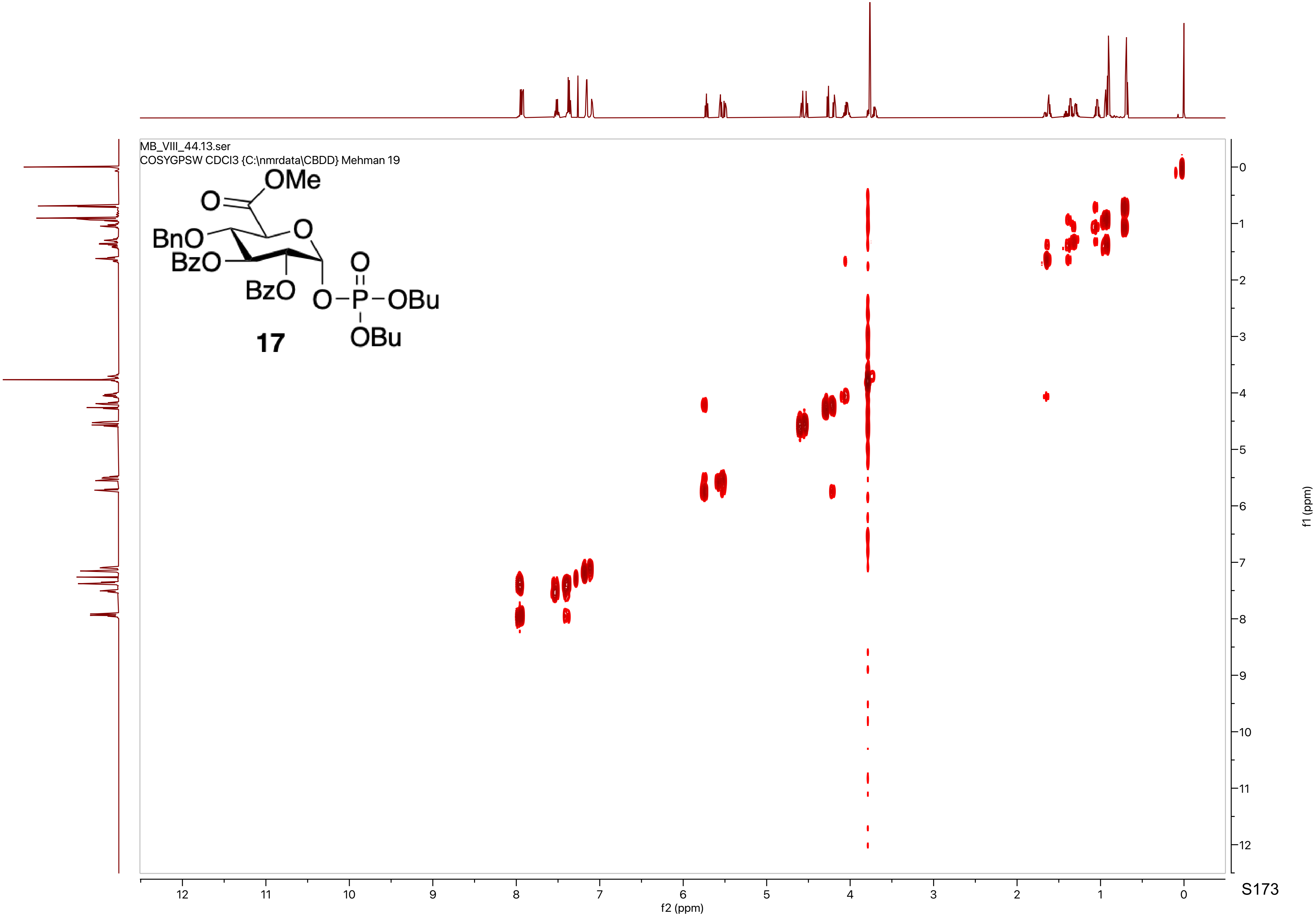

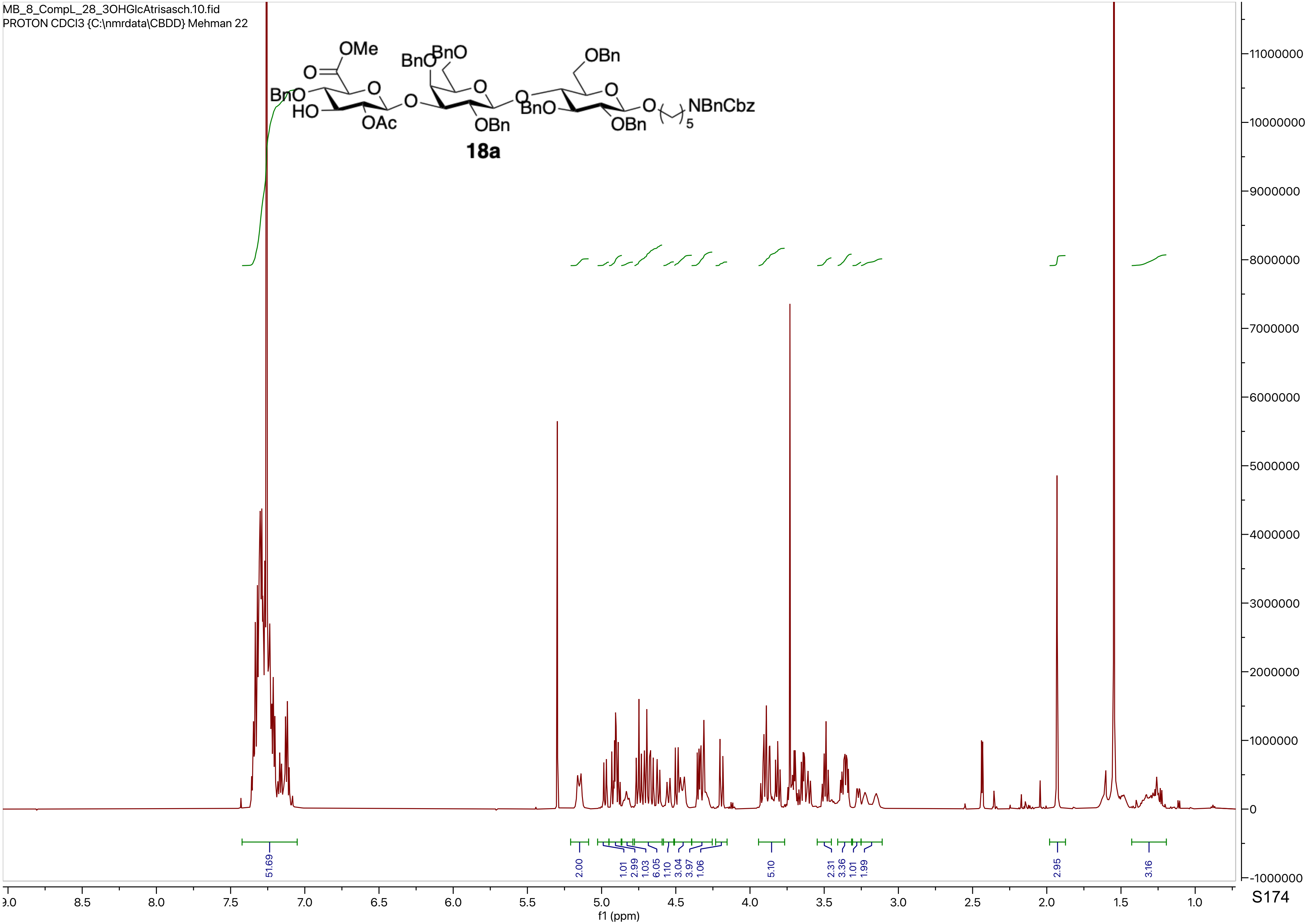
MB_8_CompL_28_3OHGlcAtrisasch.10.fid
PROTON CDCl3 {C:\nmrdata\CBDD} Mehman 22
OMe
O
BnO
HO
OAc
O
BnO
BnO
O
OBn
OBn
BnO
O
OBn
O
NBnCbz
5
18a
51.69
2.00
1.01
2.99
1.03
6.05
1.10
3.04
3.97
1.06
5.10
2.31
3.36
1.01
1.99
2.95
3.16
f1 (ppm)

MB_8_CompL_28_3OHGlcAtrisasch.15.fid
C13APT CDCl3 {C:\nmrdata\CBDD} Mehman 22

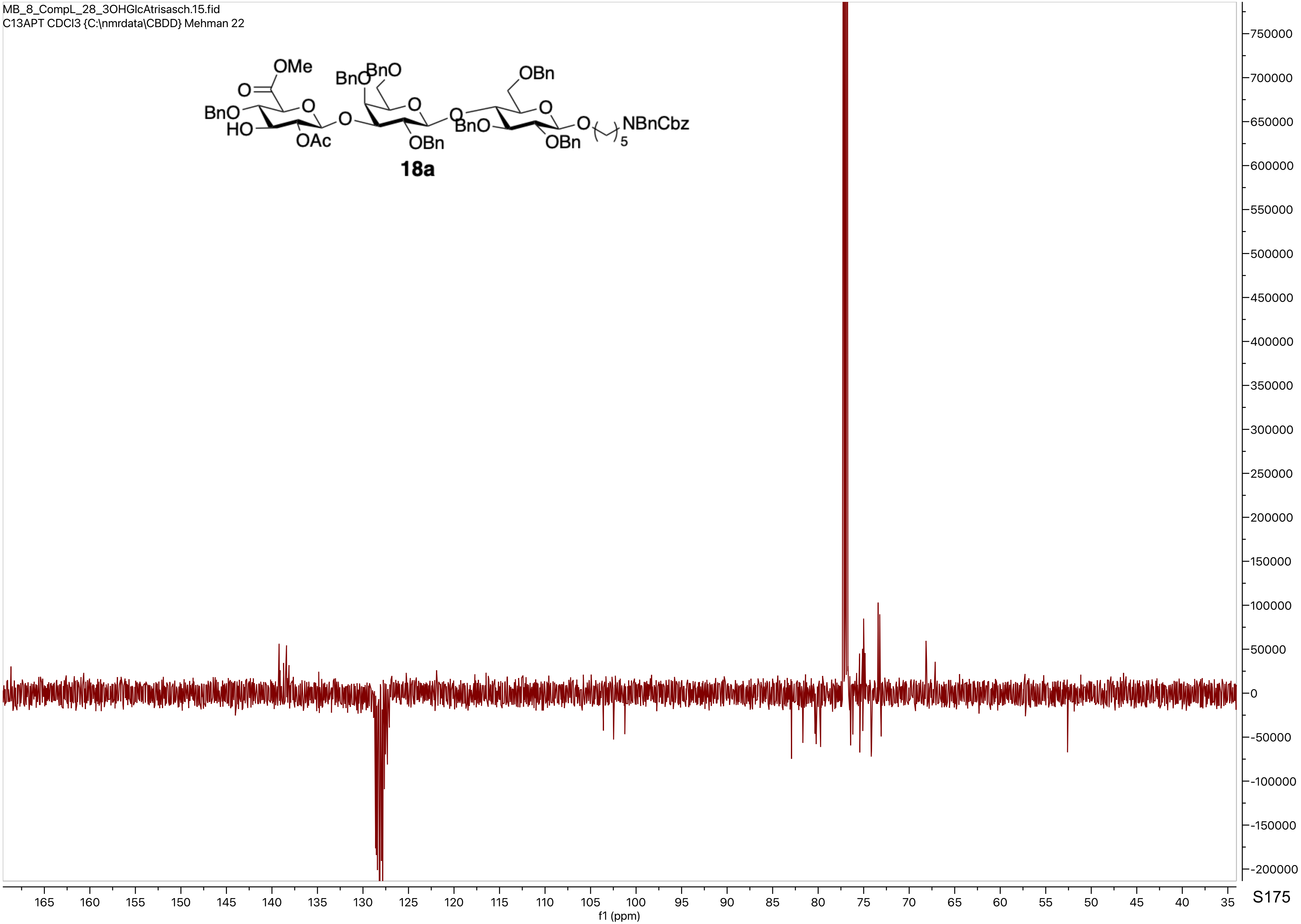

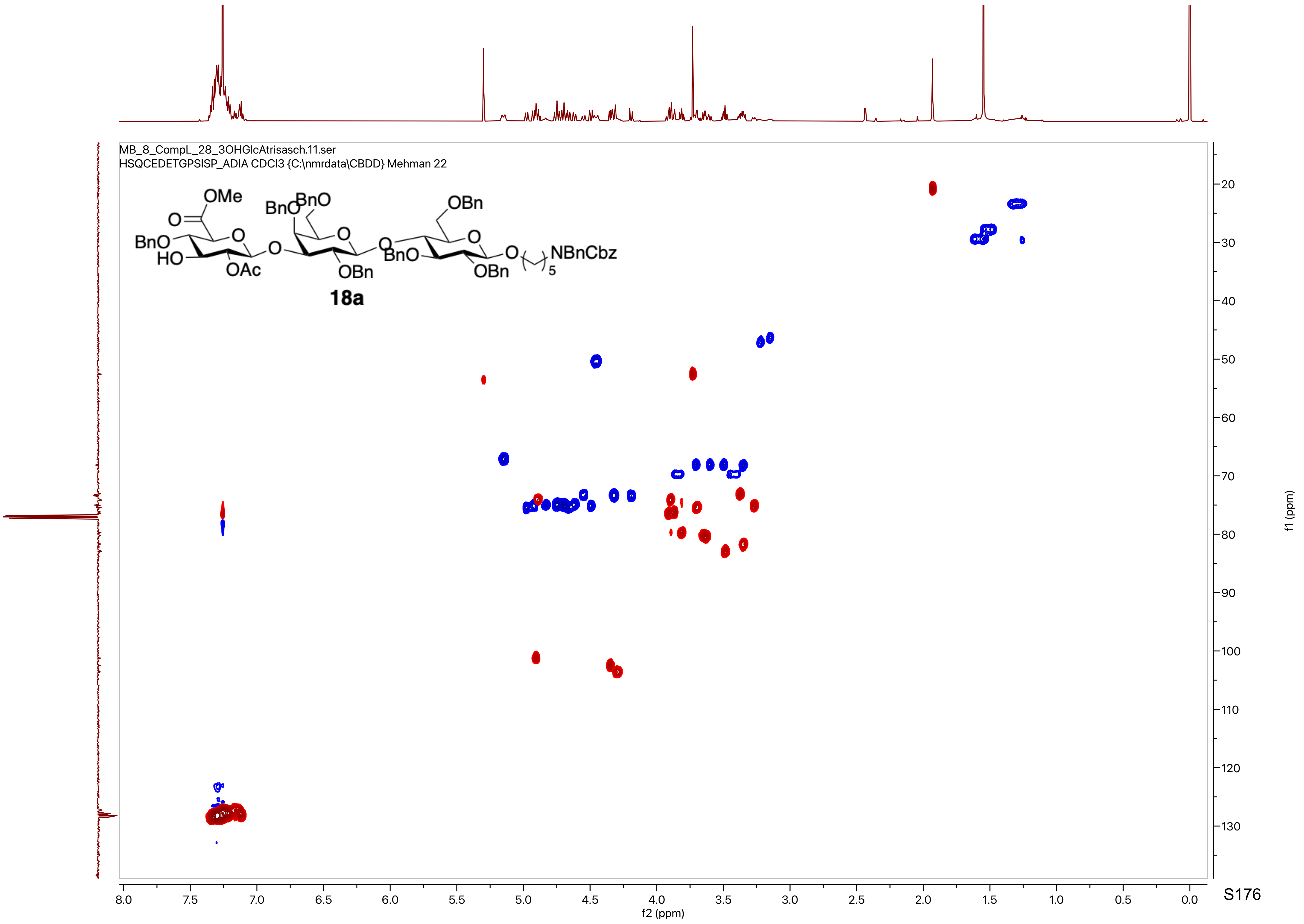
MB_8_CompL_28_3OHGlcAtrisasch.11.ser
HSQCEDETGPSISP_ADIA CDCl3 {C:\nmrdata\CBDD} Mehman 22
OMe
BnO
HO
OAc
OBn
NBnCbz
18a
f1 (ppm)
f2 (ppm)

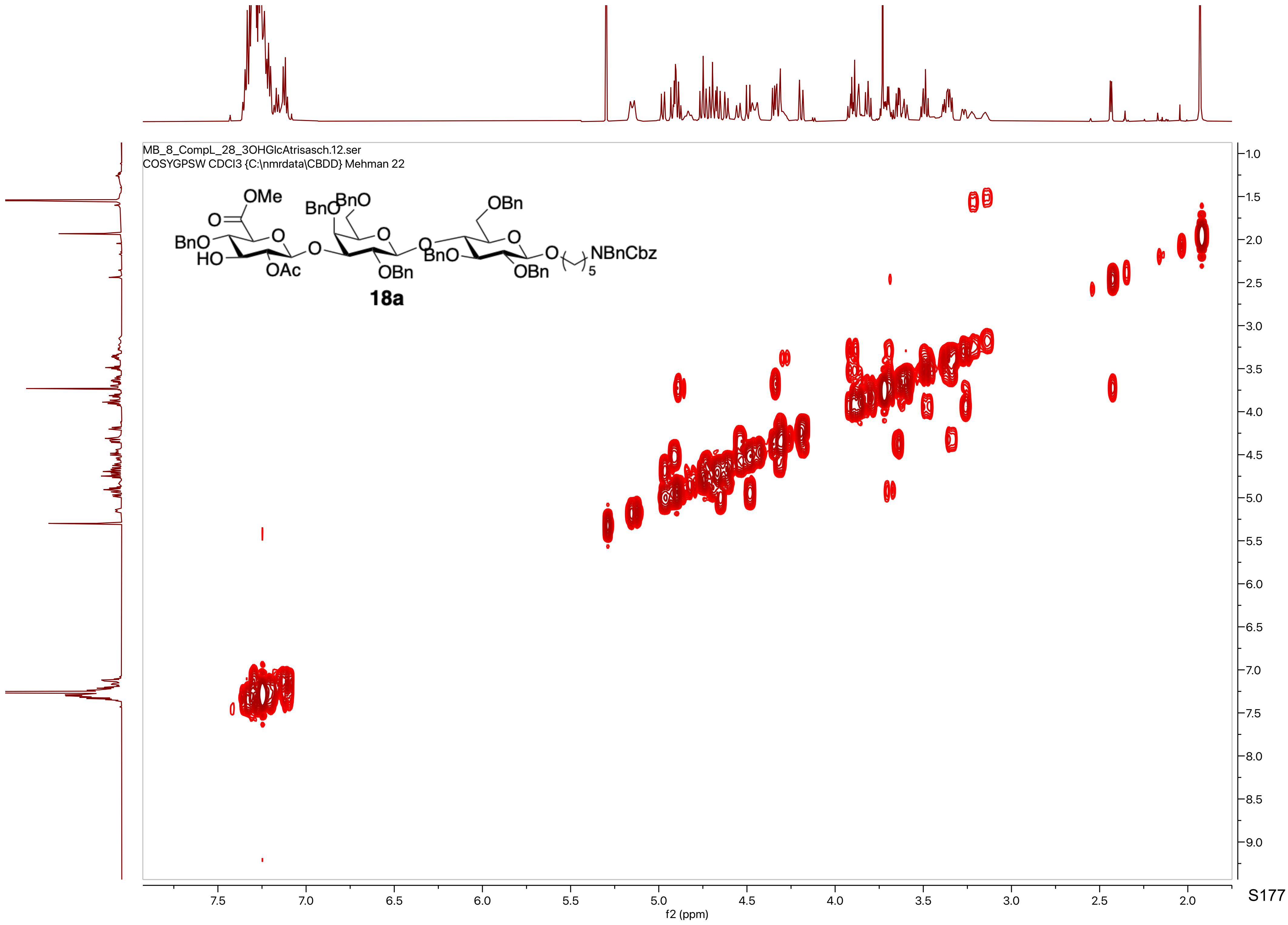

MB_8_CompL_28_3OHGlcAtrisasch.12.ser
COSYGPSW CDCl3 {C:\nmrdata\CBDD} Mehman 22
OMe
O
BnO
HO
OAc
O
BnO
BnO
OBn
BnO
OBn
OBn
O
NBnCbz
5
18a
f1 (ppm)
f2 (ppm)

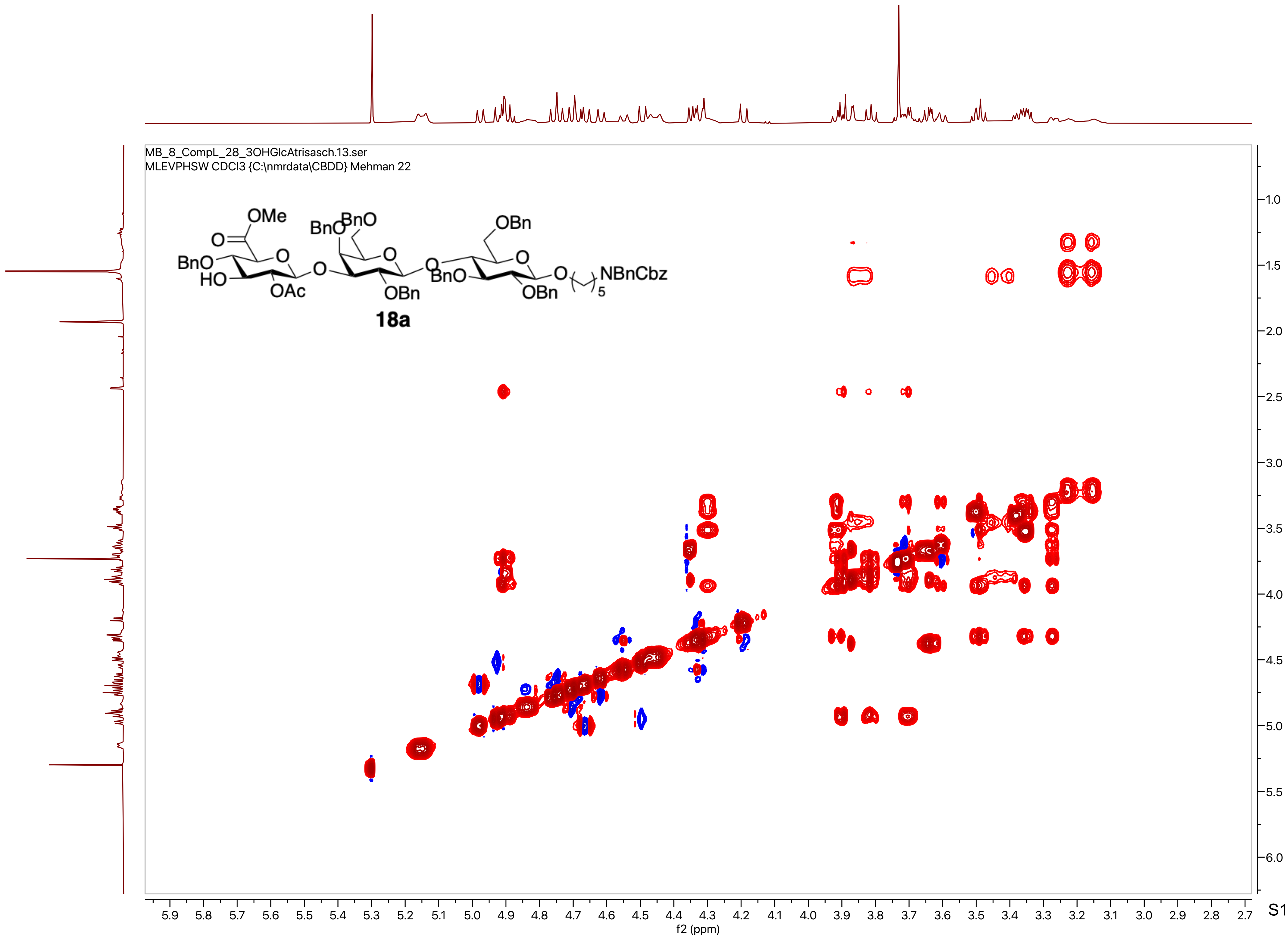
MB_8_CompL_28_3OHGlcAtrisasch.13.ser
MLEVPHSW CDCl3 {C:\nmrdata\CBDD} Mehman 22
OMe
O
BnO
HO
OAc
BnO
BnO
O
OBn
OBn
BnO
O
OBn
NBnCbz
5
18a
f2 (ppm)
f1 (ppm)

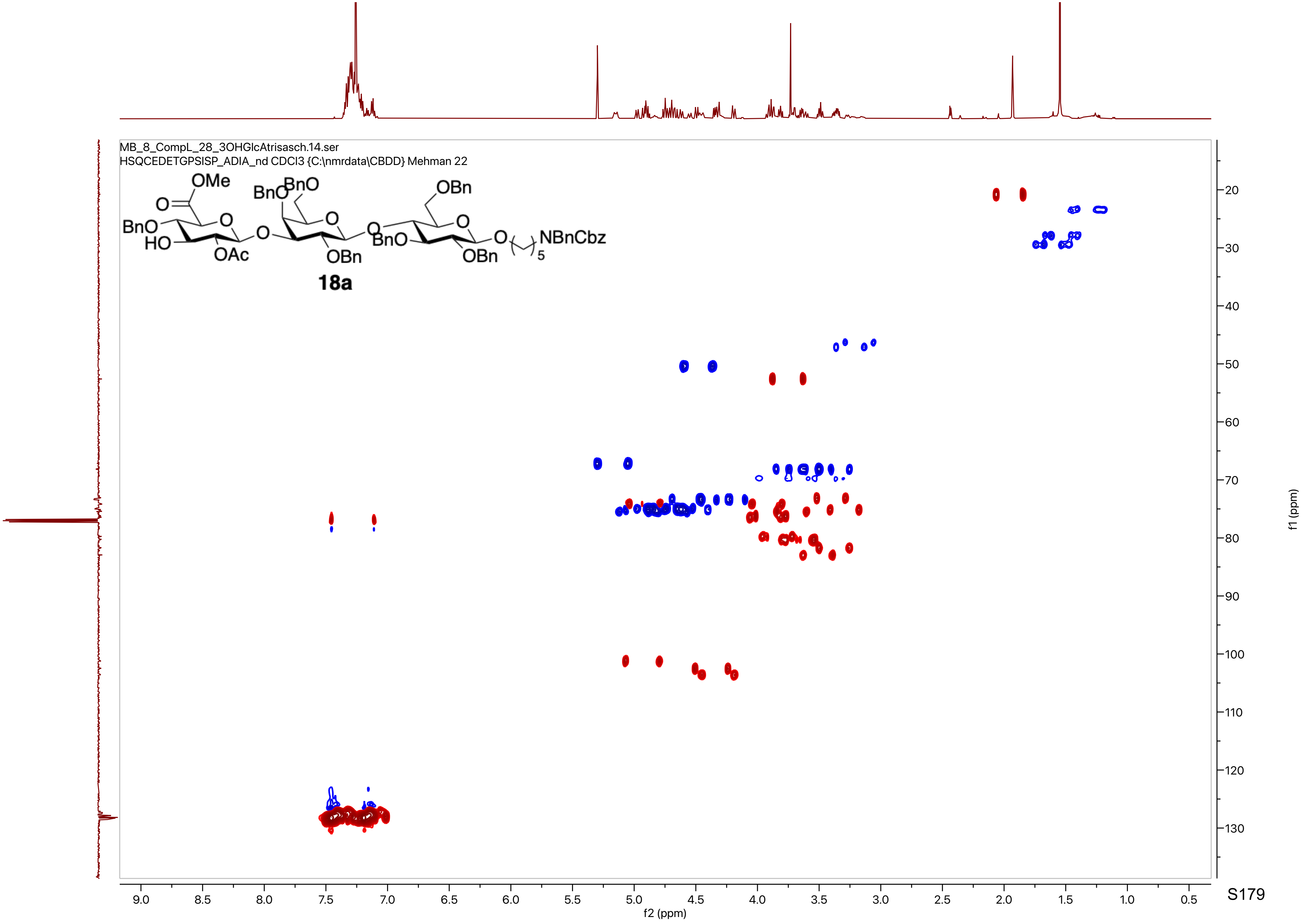
MB_8_CompL_28_3OHGlcAtrisasch.14.ser
HSQCEDETGPSISP_ADIA_nd CDCl3 {C:\nmrdata\CBDD} Mehman 22
OMe
BnO
HO
OAc
BnO
OBn
NBnCbz
18a
f1 (ppm)
f2 (ppm)

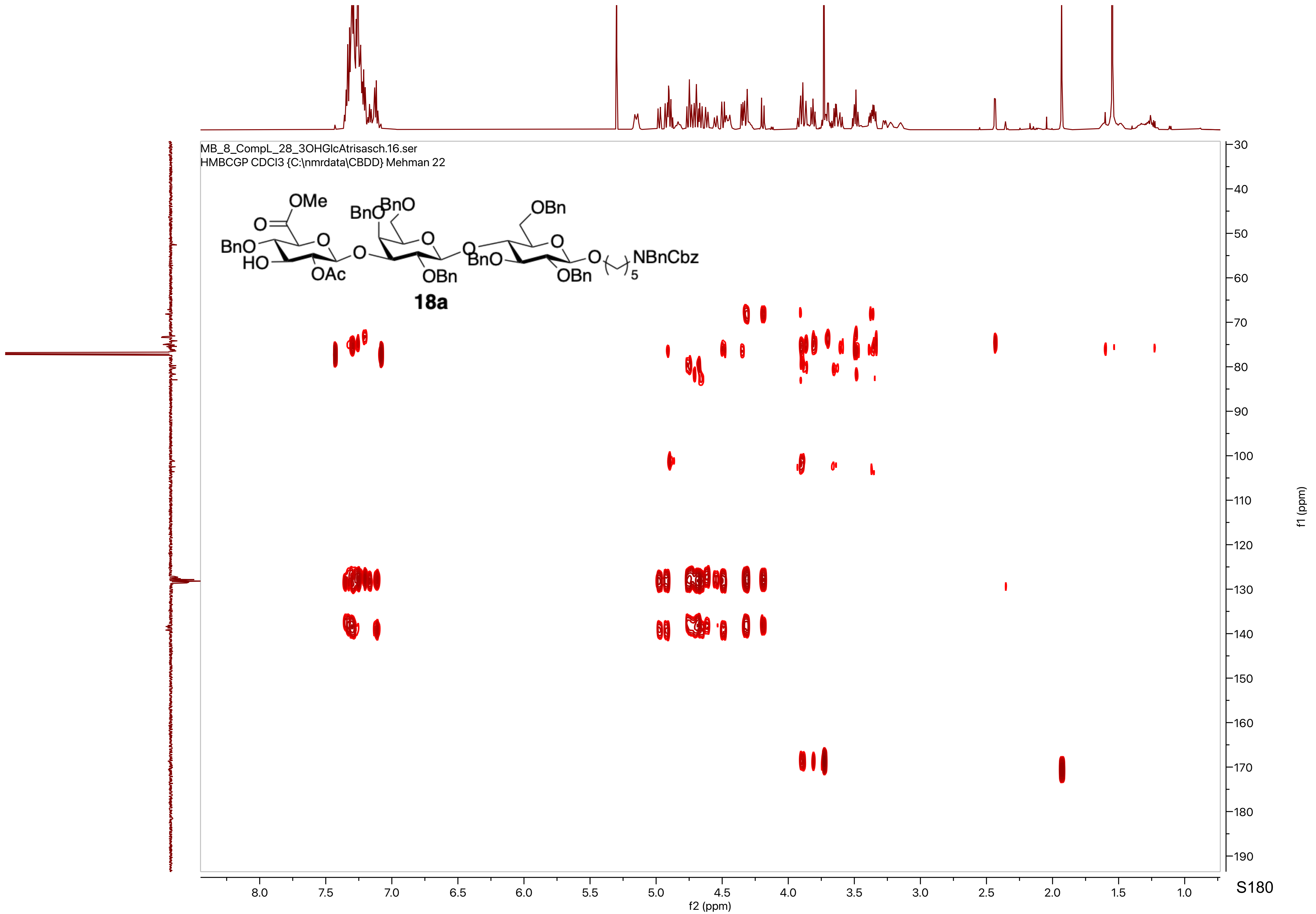

MB_8_CompL_28_3OHGlcAtrisasch.16.ser
HMBCGP CDCl3 {C:\nmrdata\CBDD} Mehman 22
OMe
O
BnO
HO
O
OAc
O
BnO
BnO
O
OBn
O
OBn
BnO
O
OBn
O
NBnCbz
5
18a
f2 (ppm)
f1 (ppm)

MB_8_CompL_29_3OH_penta.10.fid
PROTON CDCl3 {C:\nmrdata\CBDD} Mehman 35

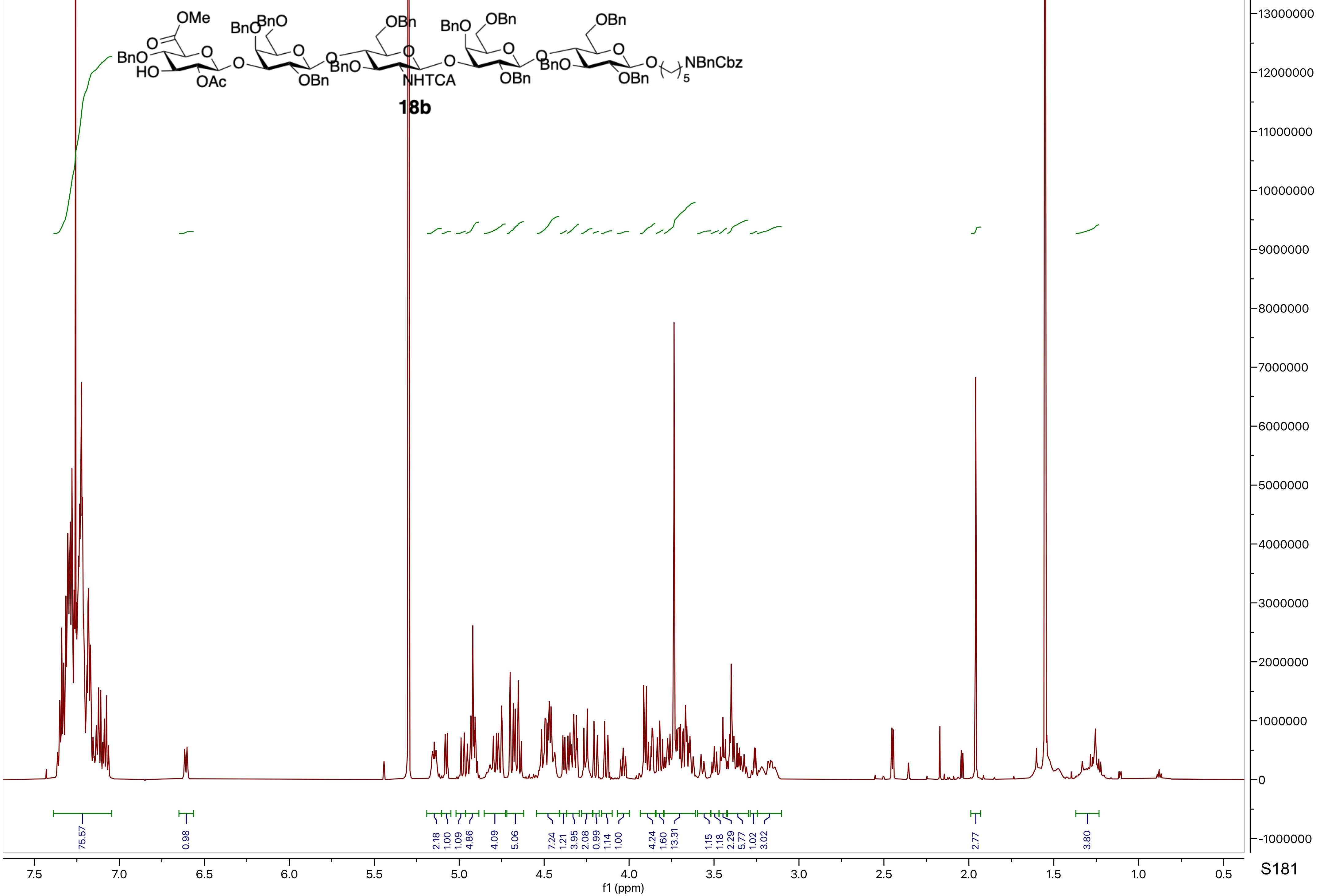

MB_8_CompL_29_3OH_penta.15.fid
C13APT CDCl3 {C:\nmrdata\CBDD} Mehman 35

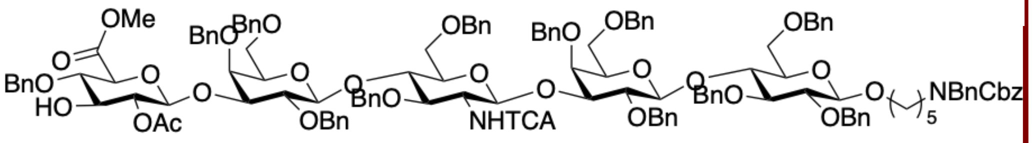

**18b**

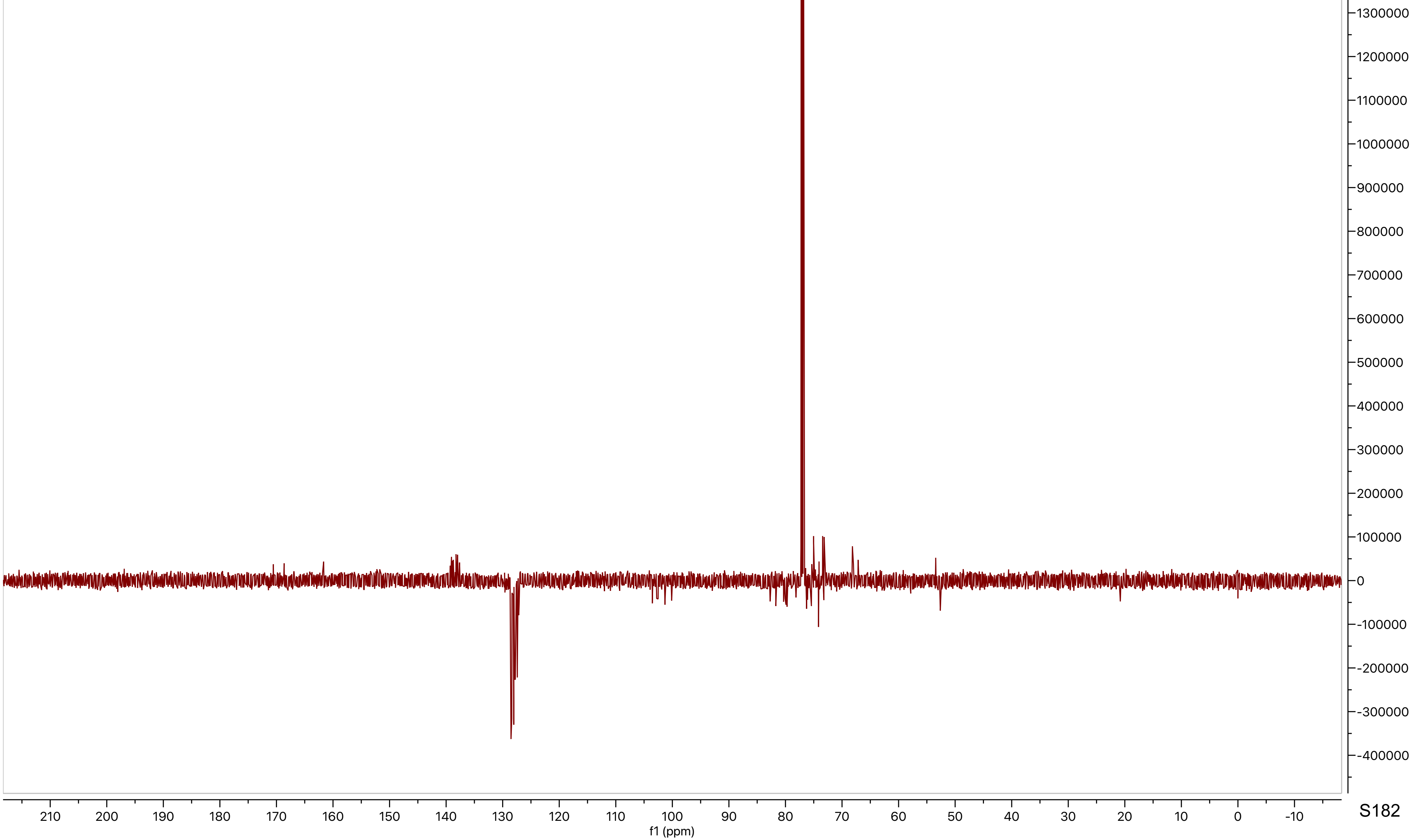

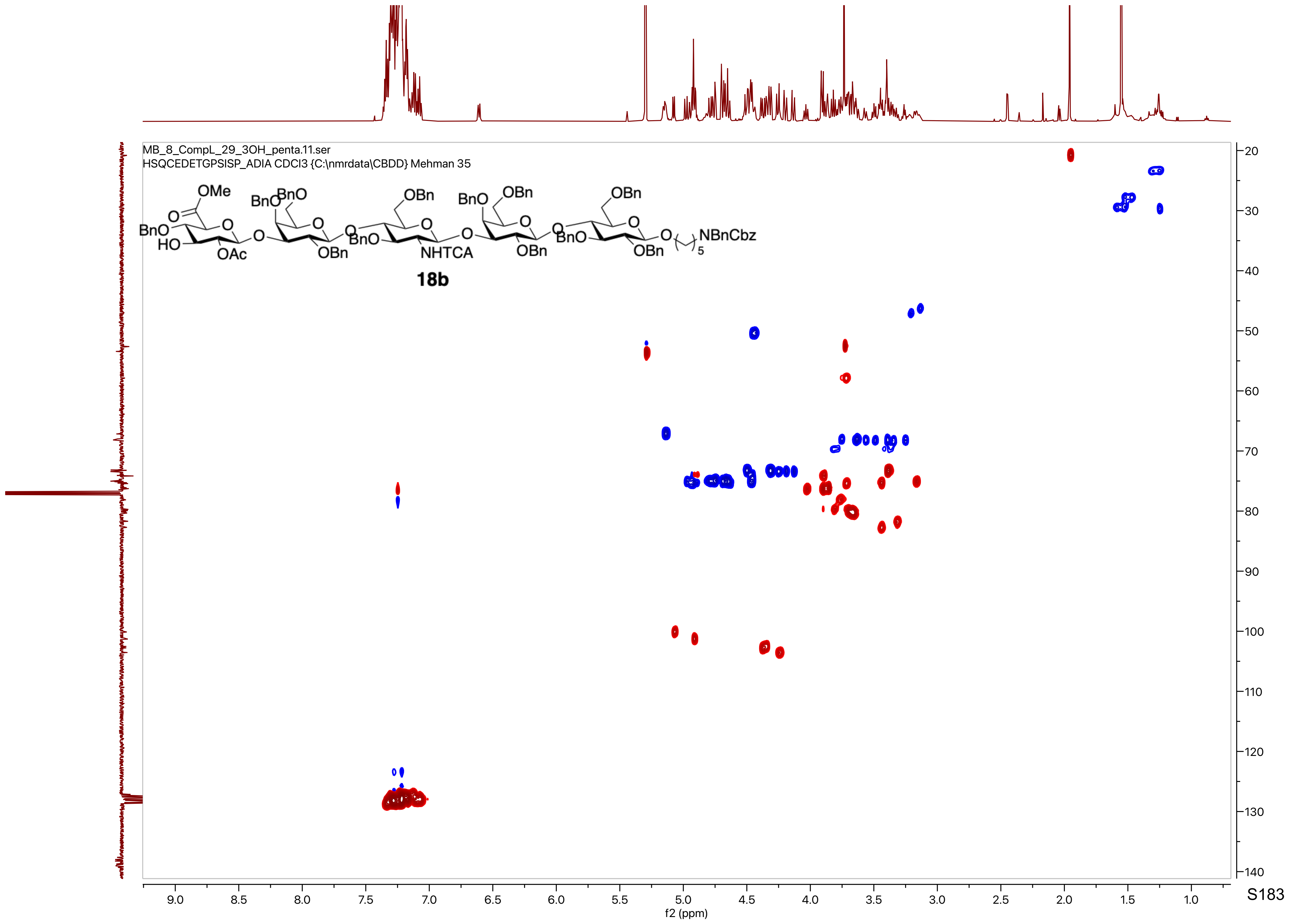
MB_8_CompL_29_3OH_penta.11.ser
HSQCEDETGPSISP_ADIA CDCl3 {C:\nmrdata\CBDD} Mehman 35
18b
f2 (ppm)
f1 (ppm)

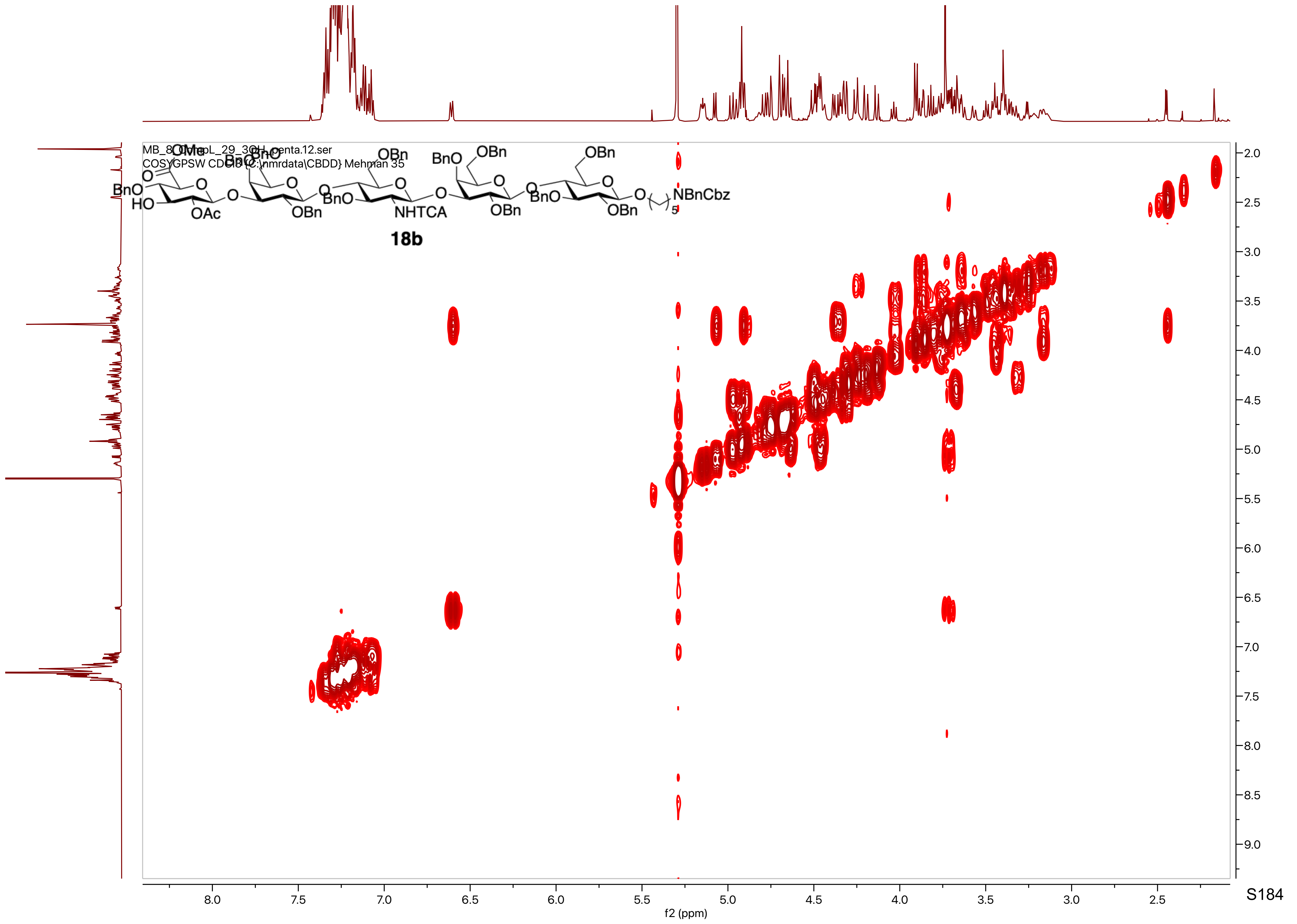
MB_8_OlympL_29_3OH_penta.12.ser
COSYGPSW CDCl3 {C:\nmrdata\CBDD} Mehman 35
18b
NBnCbz
NHTCA
f2 (ppm)
f1 (ppm)

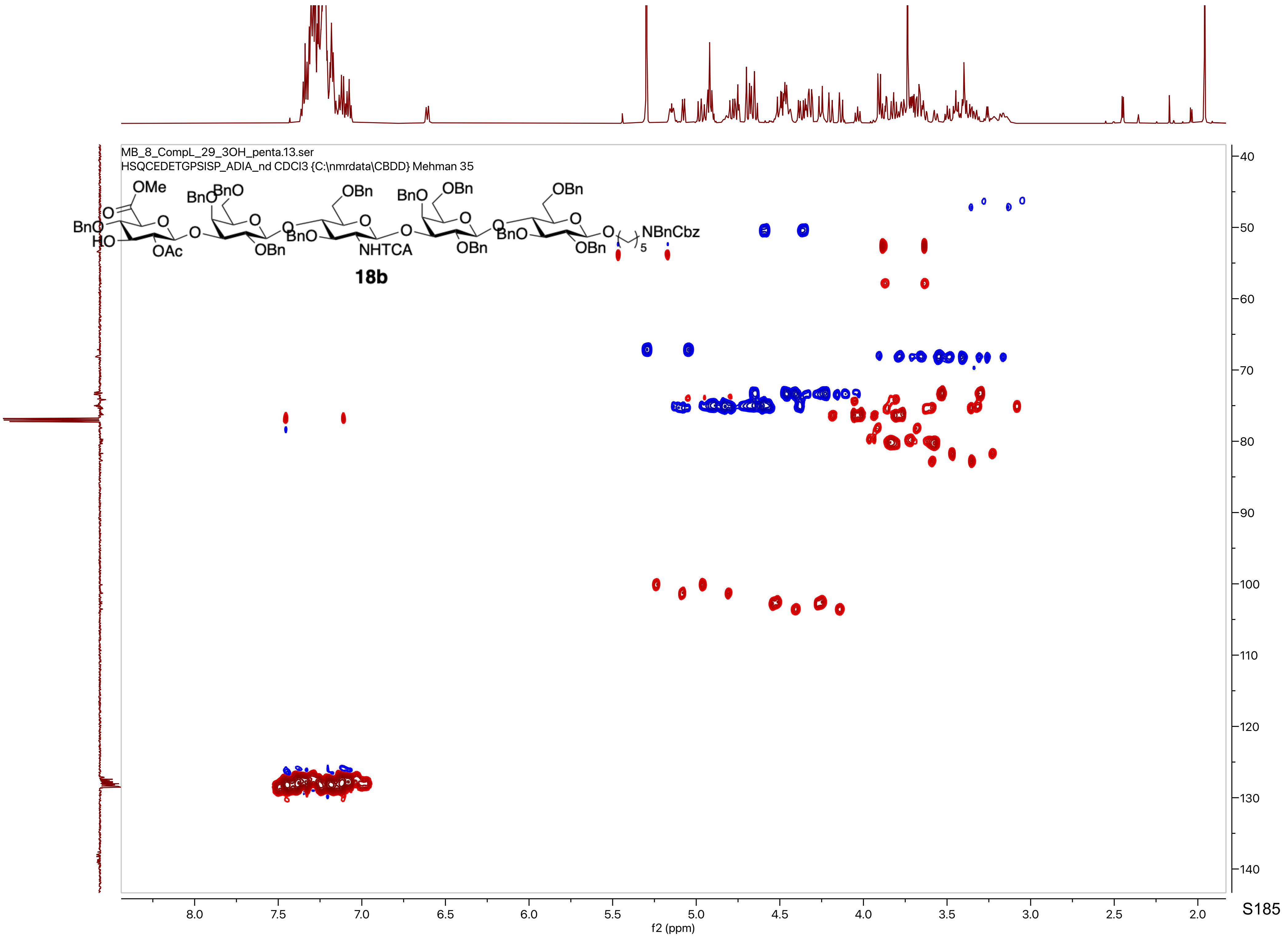
MB_8_CompL_29_3OH_penta.13.ser
HSQCEDETGPSISP_ADIA_nd CDCl3 {C:\nmrdata\CBDD} Mehman 35
18b
f2 (ppm)
f1 (ppm)

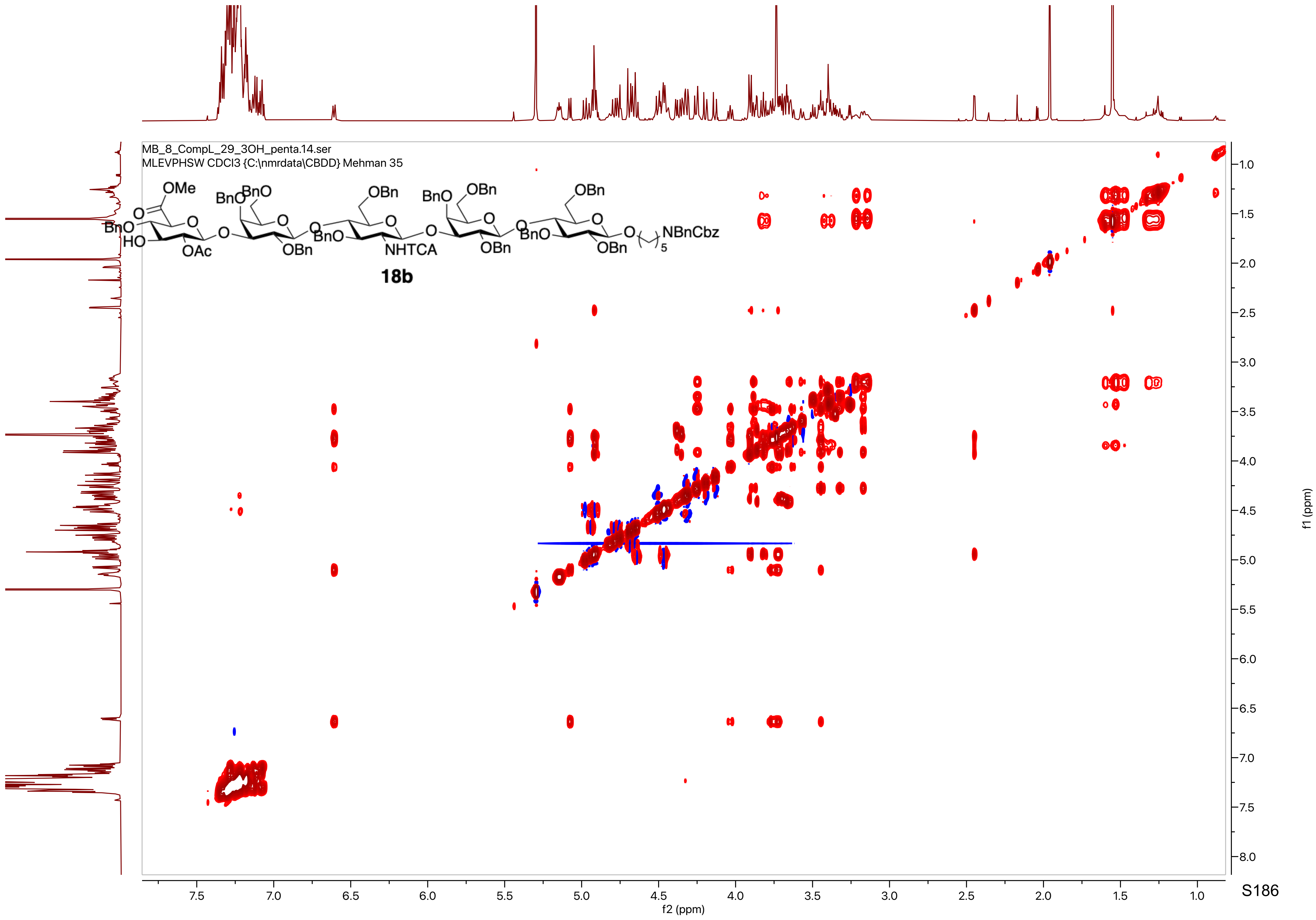
MB_8_CompL_29_3OH_penta.14.ser
MLEVPHSW CDCl3 {C:\nmrdata\CBDD} Mehman 35
OMe
O
BnO
HO
O
OAc
BnO
BnO
O
OBn
BnO
OBn
NHTCA
BnO
OBn
OBn
BnO
OBn
OBn
NBnCbz
5
18b
f2 (ppm)
f1 (ppm)

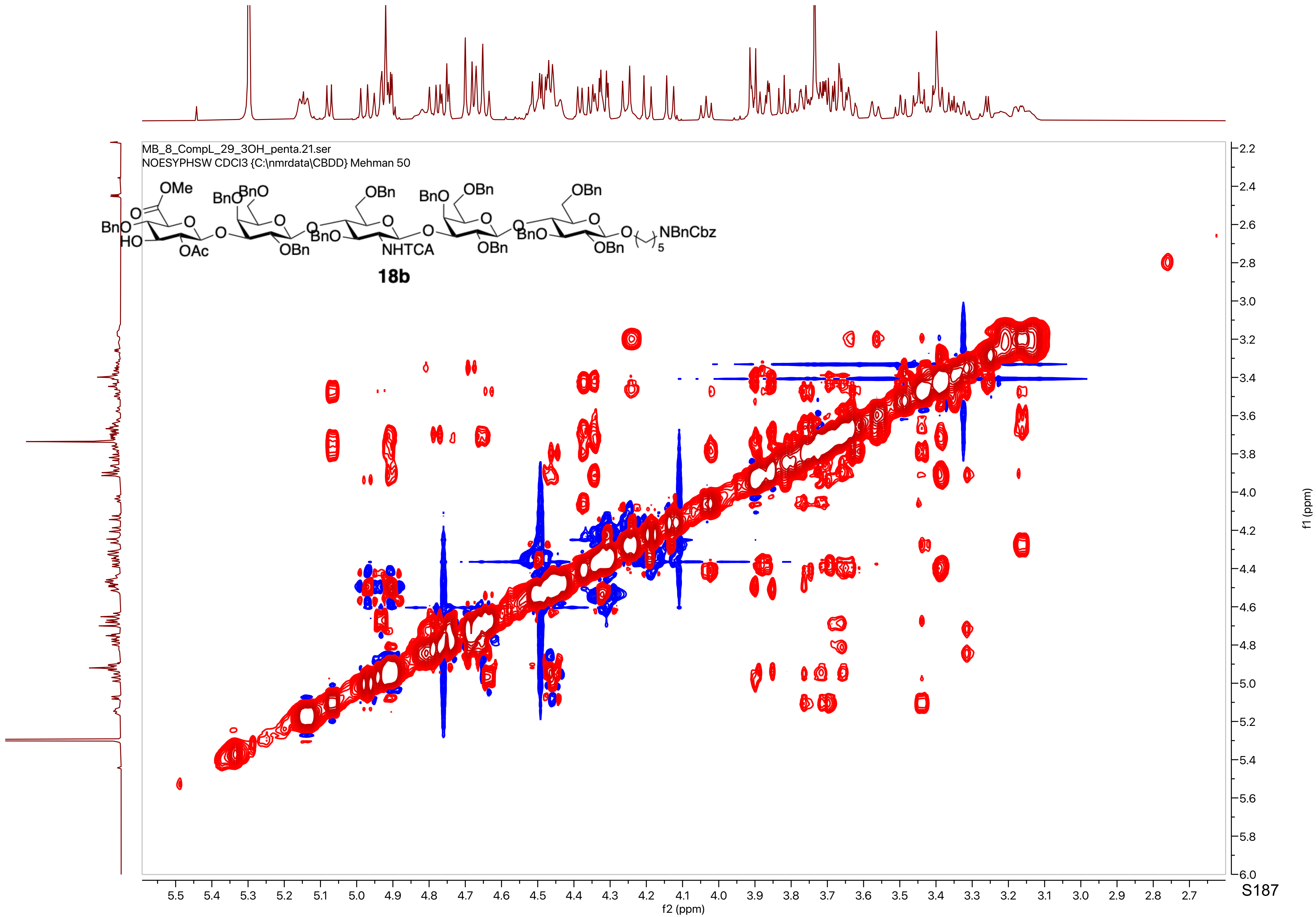
MB_8_CompL_29_3OH_penta.21.ser
NOESYPHSW CDCl3 {C:\nmrdata\CBDD} Mehman 50
OMe
BnO
BnO
BnO
OBn
BnO
OBn
OBn
HO
OAc
OBn
BnO
NHTCA
OBn
BnO
OBn
NBnCbz
5
18b
f2 (ppm)
f1 (ppm)

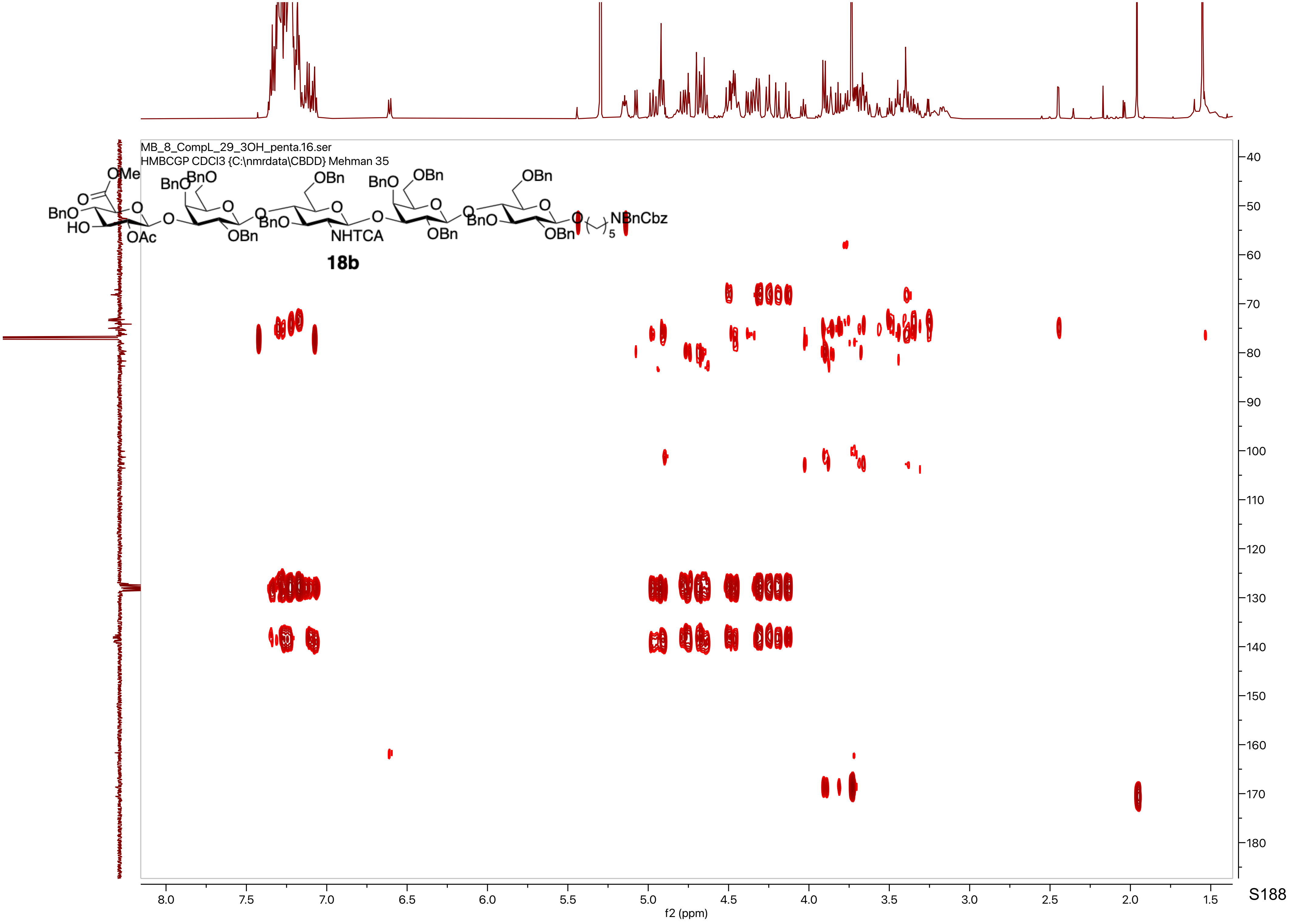

MB_8_CompL_29_3OH_penta.16.ser
HMBCGP CDCl3 {C:\nmrdata\CBDD} Mehman 35
OMe
BnO
HO
OAc
OBn
NHTCA
NBnCbz
18b
f2 (ppm)
f1 (ppm)

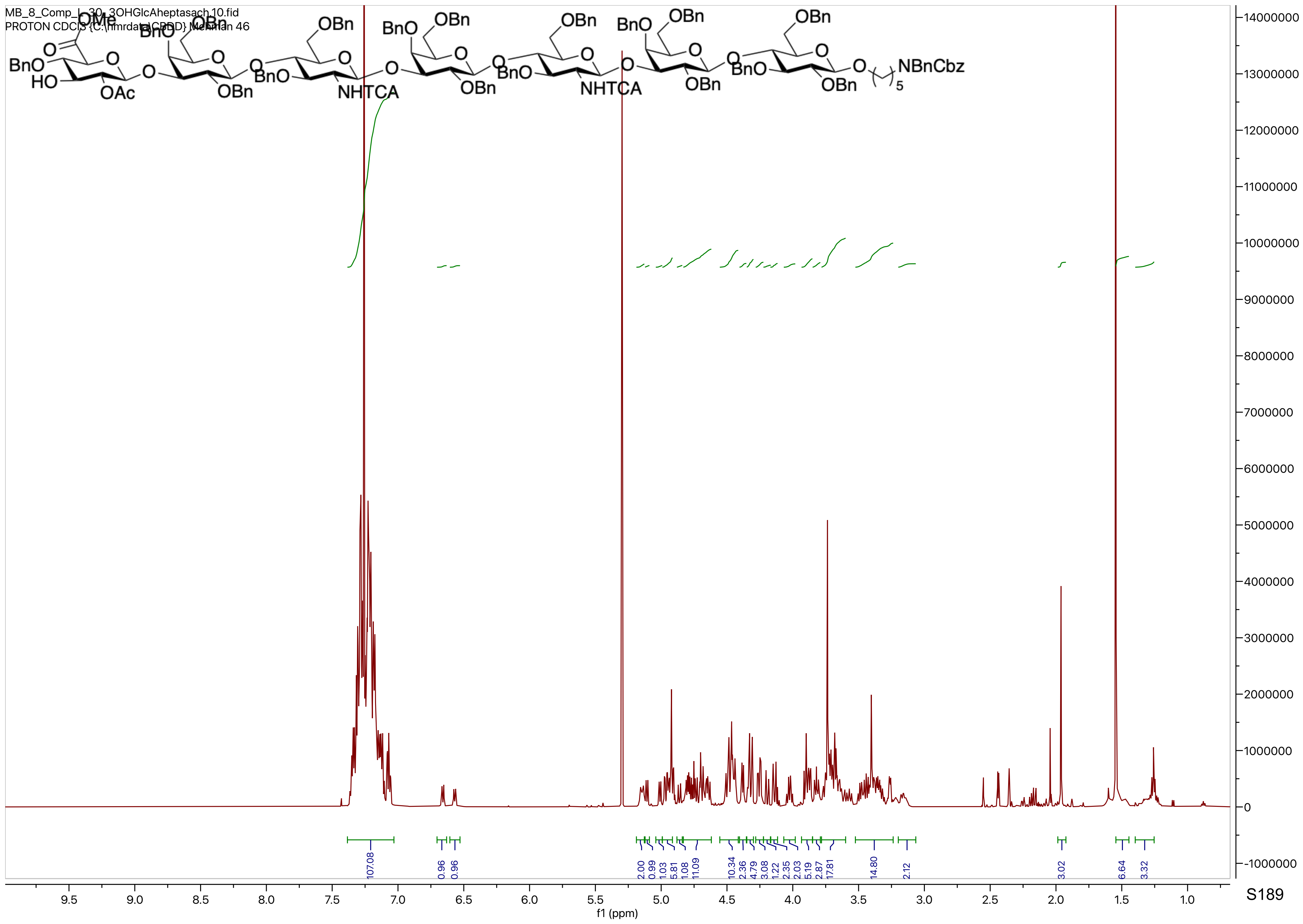
MB_8_Comp_L_30_3OHGlcAheptasach.10.fid
PROTON CDCl3 {C:\nmrdata\CBDD} Mehman 46
NHTCA
NBnCbz
f1 (ppm)

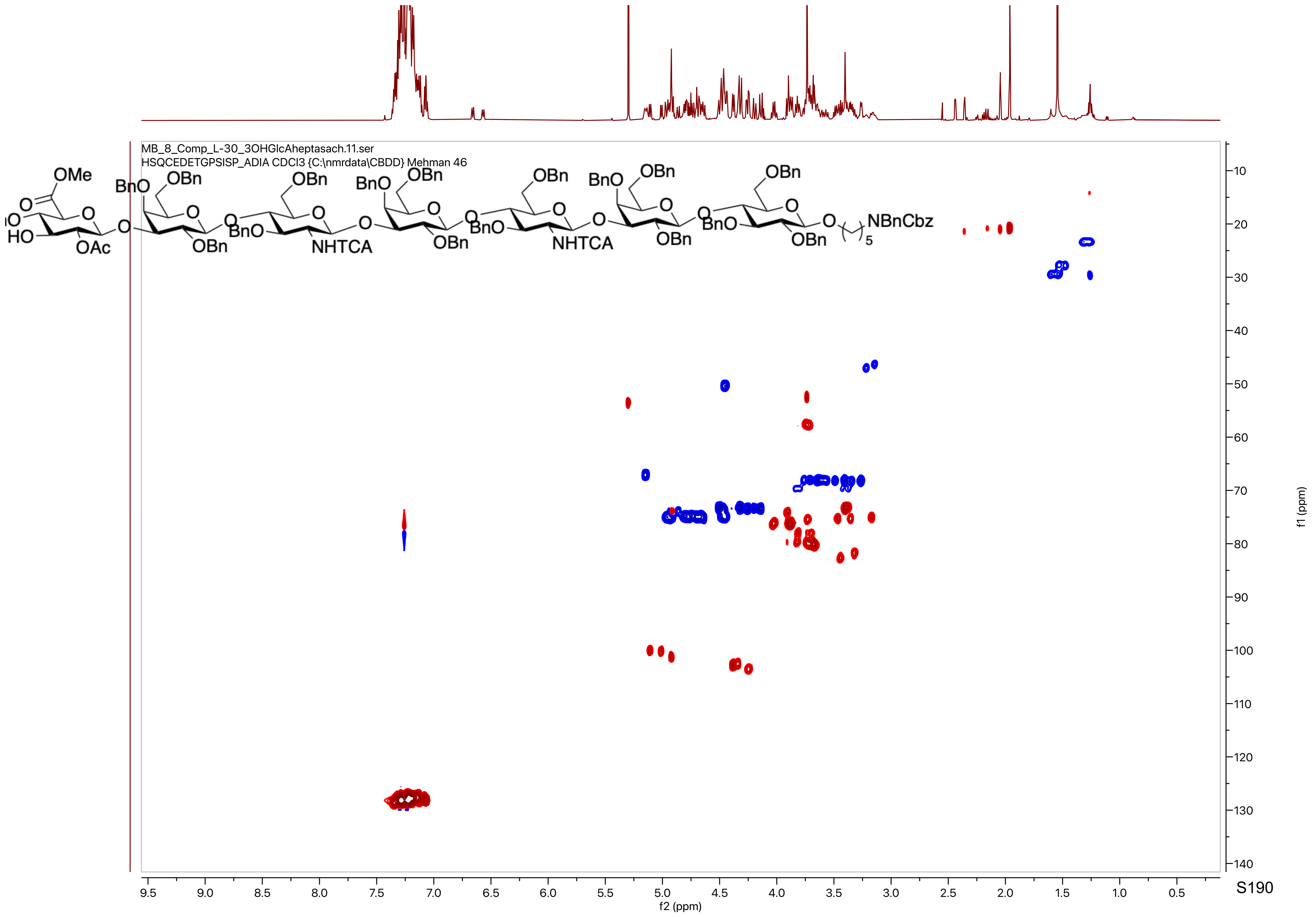

MB_8_Comp_L-30_3OHGlcAheptasach.11.ser
HSQCEDETGPSISP_ADIA CDCl3 {C:\nmrdata\CBDD} Mehman 46
OMe
OAc
NHTCA
NBnCbz
f2 (ppm)
f1 (ppm)

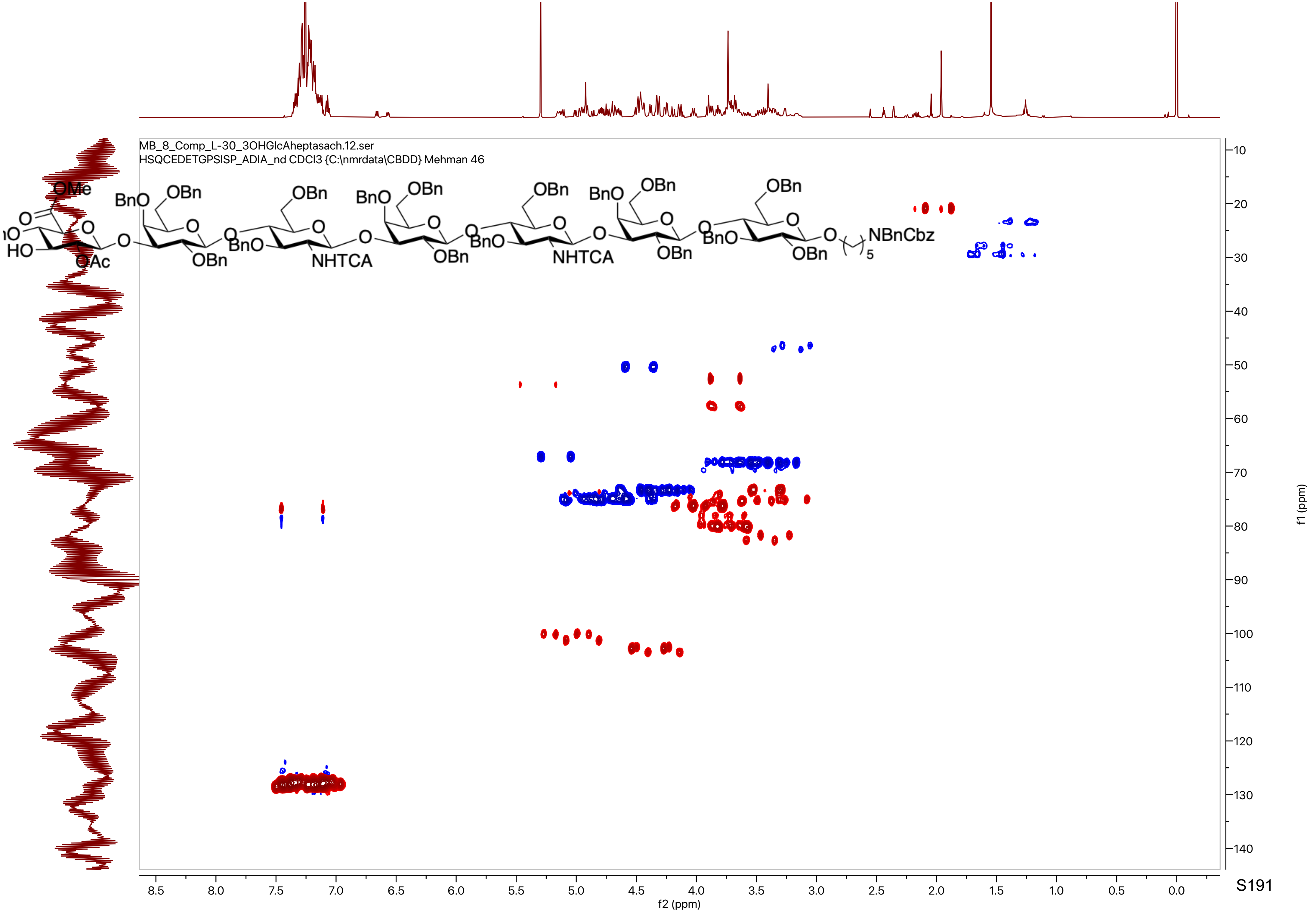

MB_8_Comp_L-30_3OHGlcAheptasach.12.ser
HSQCEDETGPSISP_ADIA_nd CDCl3 {C:\nmrdata\CBDD} Mehman 46
f2 (ppm)
f1 (ppm)

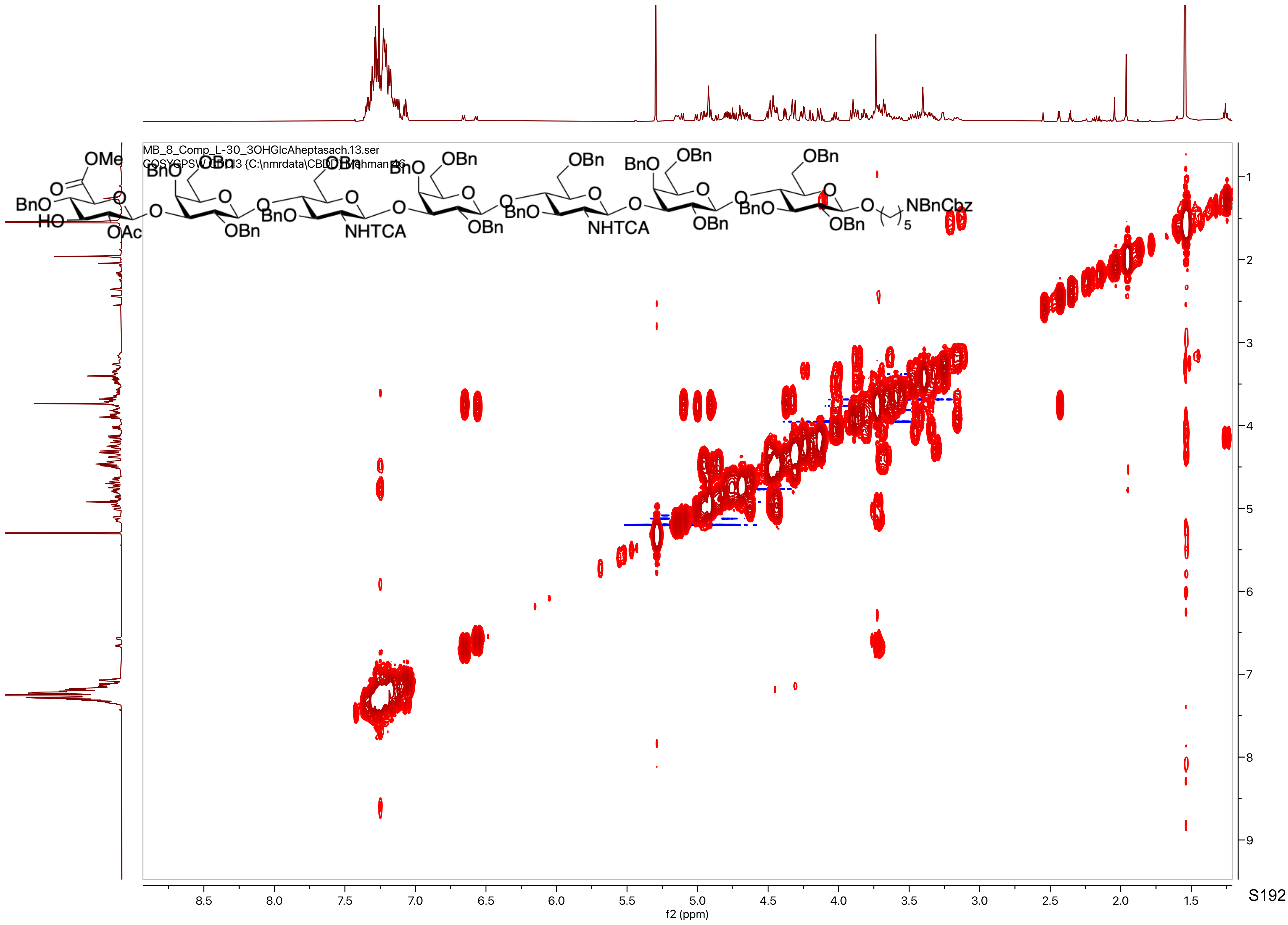

MB_8_Comp_L-30_3OHGlcAheptasach.13.ser
COSYGPSW CDCl3 {C:\nmrdata\CBDD} Rahman 46
OMe
BnO
HO
OAc
OBn
NHTCA
NBnCbz
f2 (ppm)
f1 (ppm)

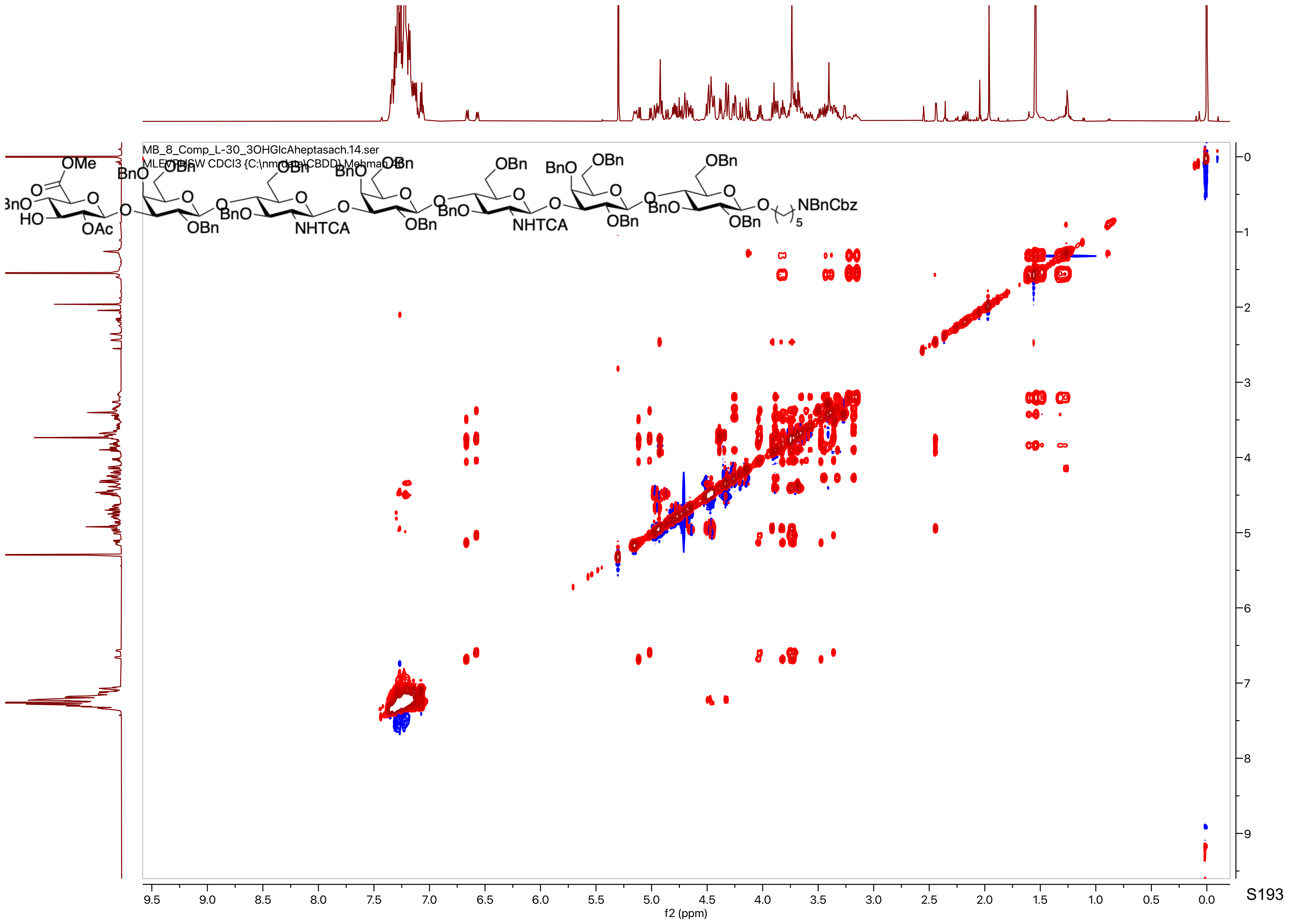
MB_8_Comp_L-30_3OHGlcAheptasach.14.ser
f2 (ppm)
f1 (ppm)

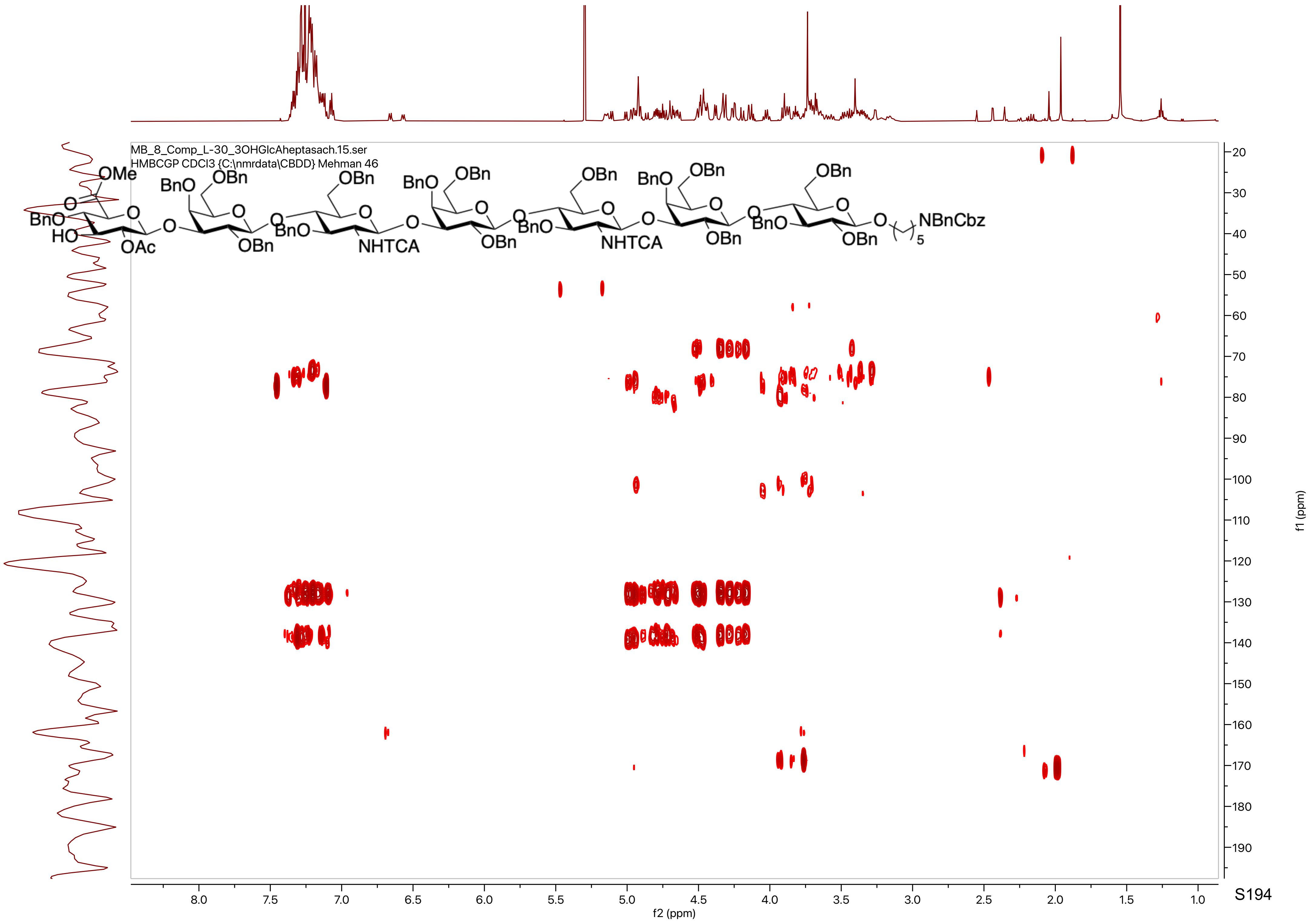
MB_8_Comp_L-30_3OHGlcAheptasach.15.ser
HMBCGP CDCl3 {C:\nmrdata\CBDD} Mehman 46
f1 (ppm)
f2 (ppm)

MB_8_comp_4.10.fid
PROTON D2O {C:\nmrdata\CBDD} Mehman 10

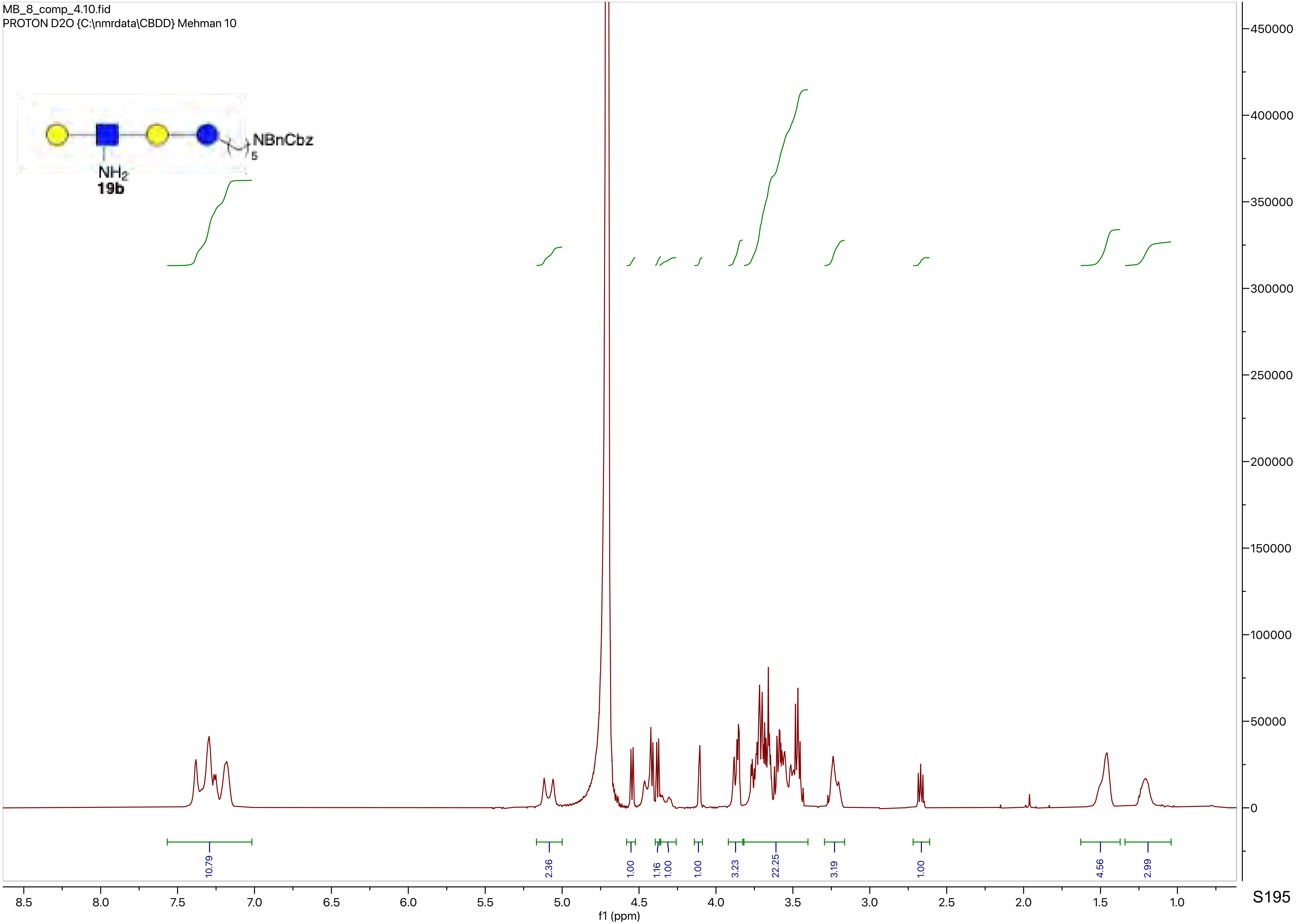

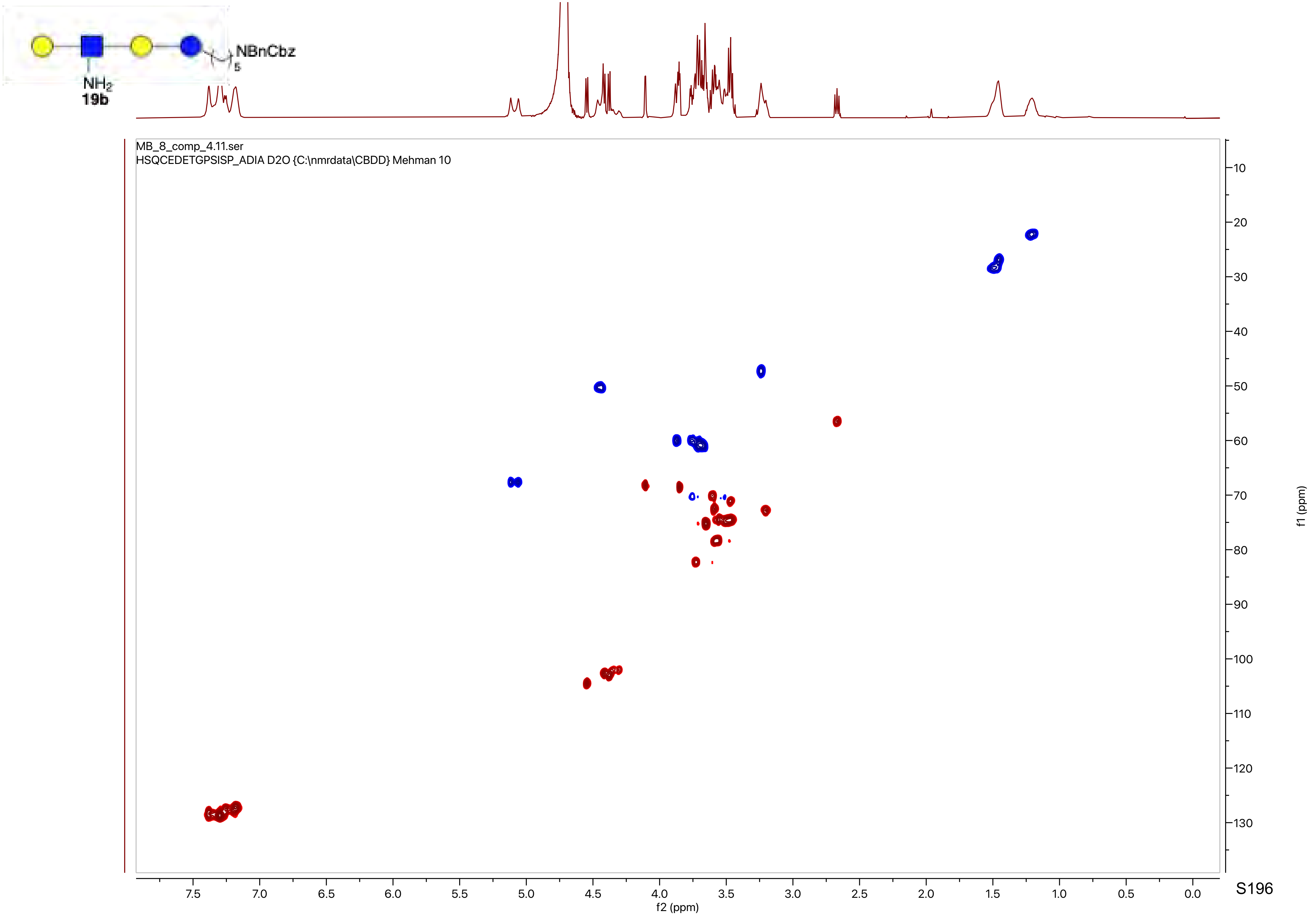

NBnCbz
5
NH2
19b
MB_8_comp_4.11.ser
HSQCEDETGPSISP_ADIA D2O {C:\nmrdata\CBDD} Mehman 10
f2 (ppm)
f1 (ppm)

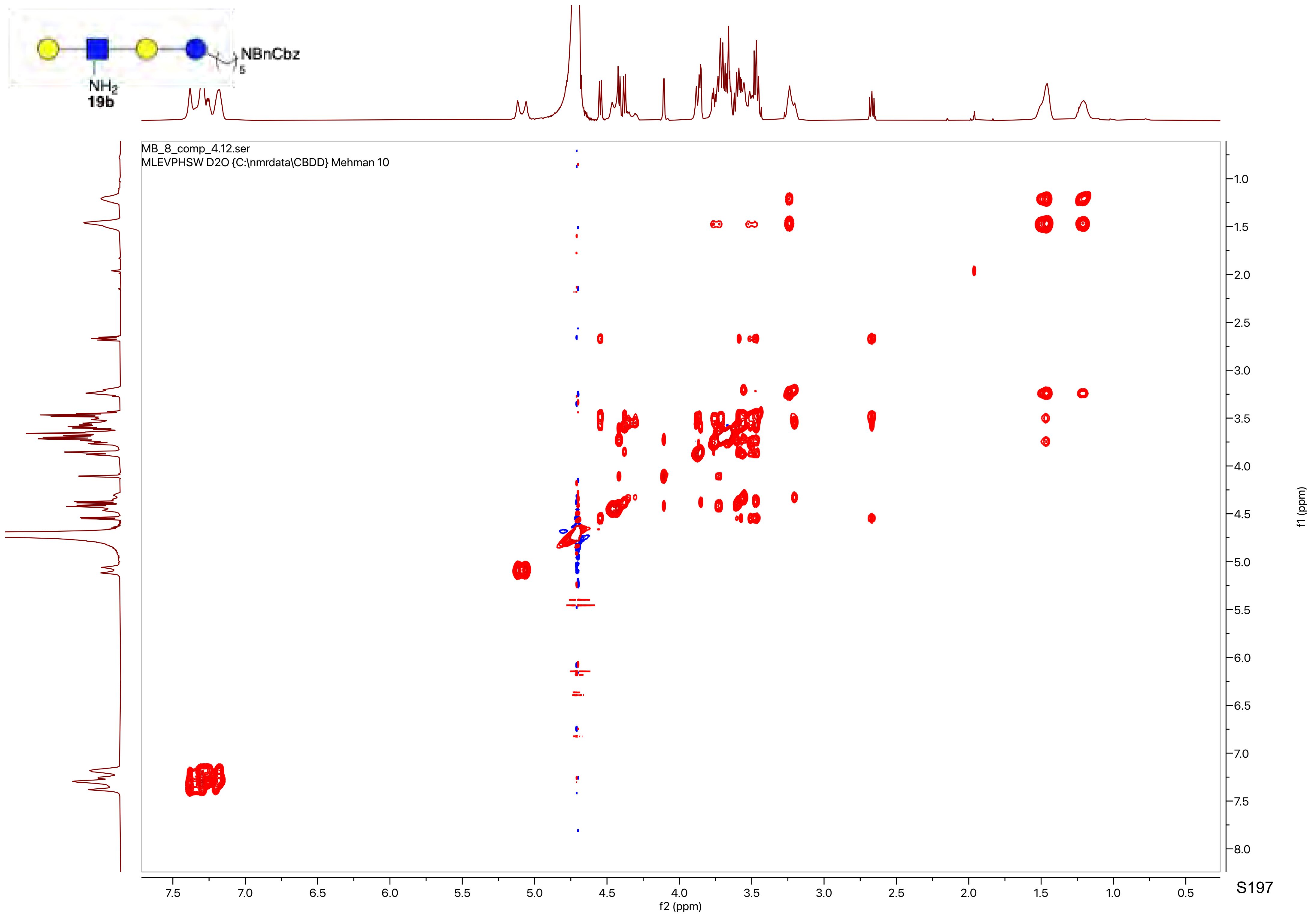
NBnCbz
5
NH2
19b
MB_8_comp_4.12.ser
MLEVPHSW D2O {C:\nmrdata\CBDD} Mehman 10
f2 (ppm)
f1 (ppm)

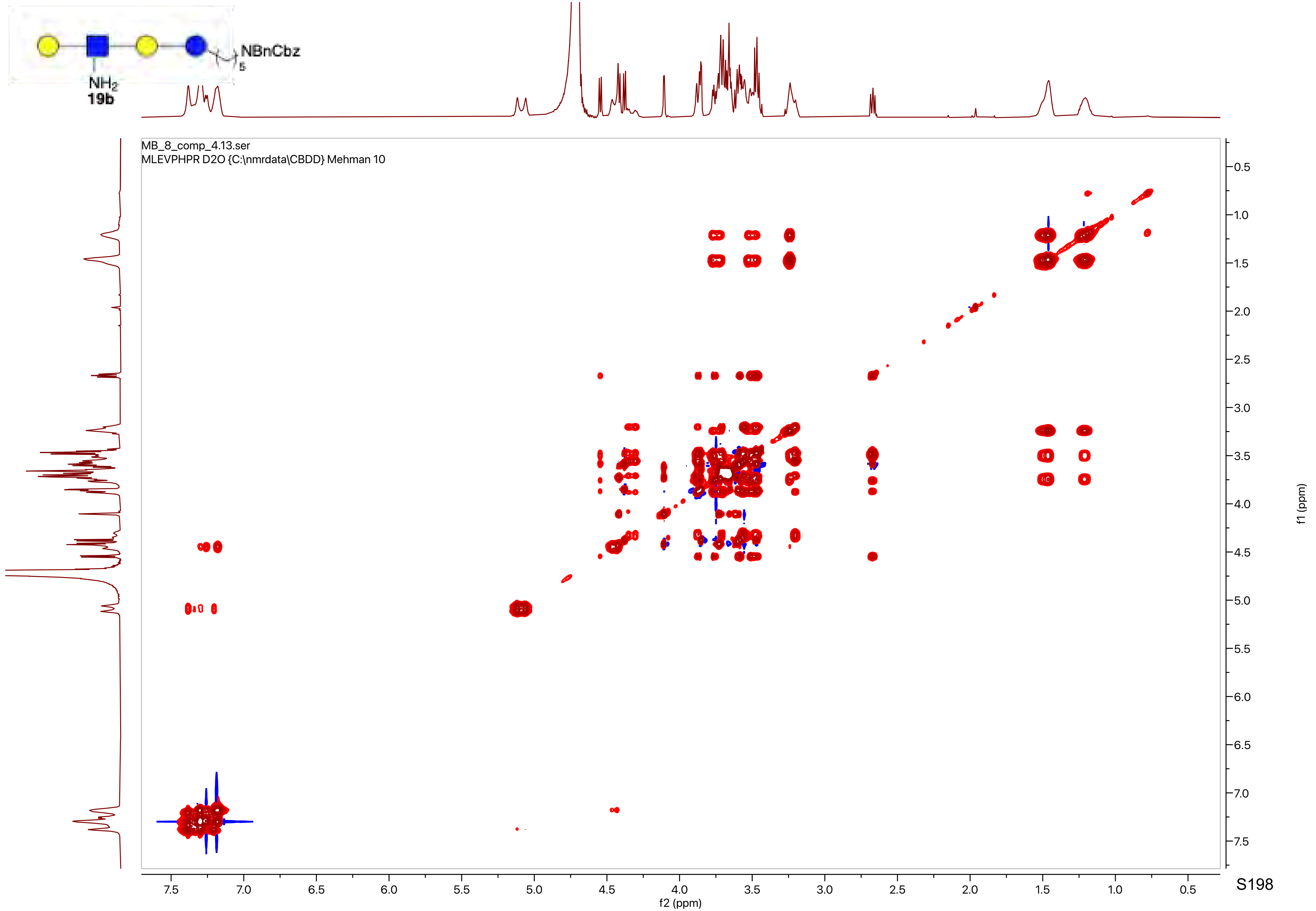

NBnCbz
5
NH2
19b
MB_8_comp_4.13.ser
MLEVPHPR D2O {C:\nmrdata\CBDD} Mehman 10
f2 (ppm)
f1 (ppm)

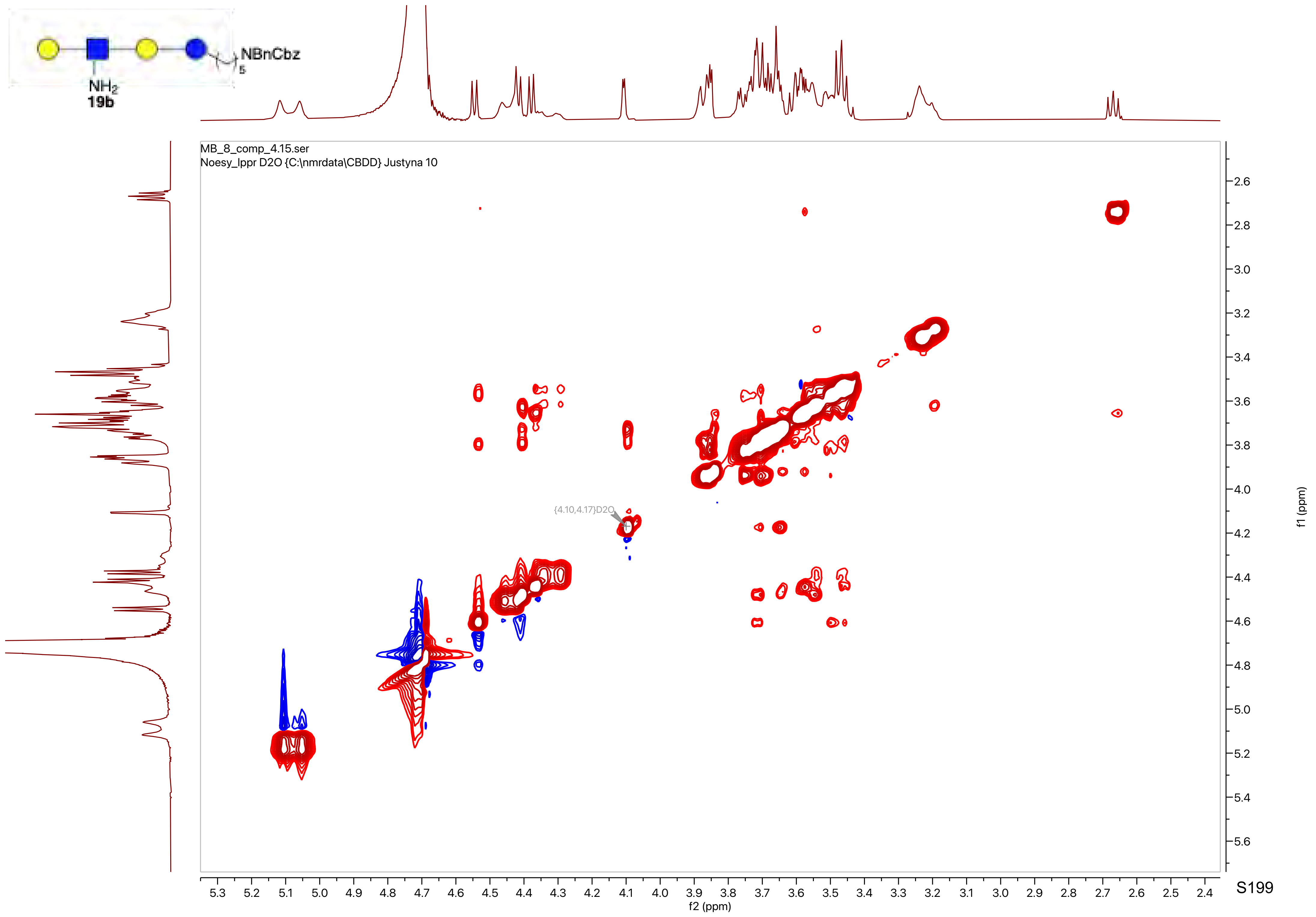

NBnCbz
5
NH2
19b
MB_8_comp_4.15.ser
Noesy_lppr D2O {C:\nmrdata\CBDD} Justyna 10
{4.10,4.17}D2O
f2 (ppm)
f1 (ppm)

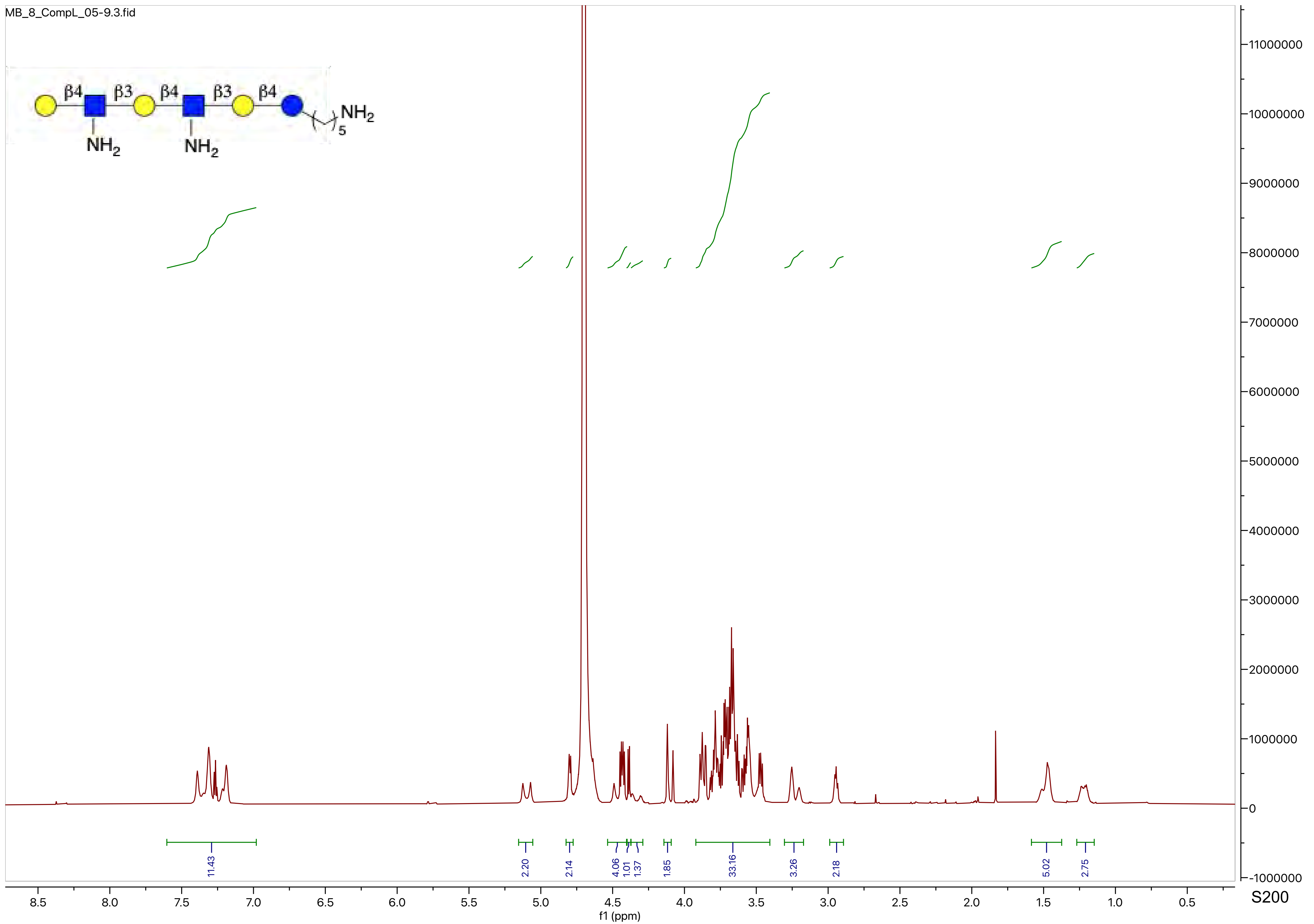
MB_8_CompL_05-9.3.fid
β4
β3
β4
β3
β4
NH2
NH2
NH2
5
11.43
2.20
2.14
4.06
1.01
1.37
1.85
33.16
3.26
2.18
5.02
2.75
8.5
8.0
7.5
7.0
6.5
6.0
5.5
5.0
4.5
4.0
3.5
3.0
2.5
2.0
1.5
1.0
0.5
f1 (ppm)
11000000
10000000
9000000
8000000
7000000
6000000
5000000
4000000
3000000
2000000
1000000
0
-1000000

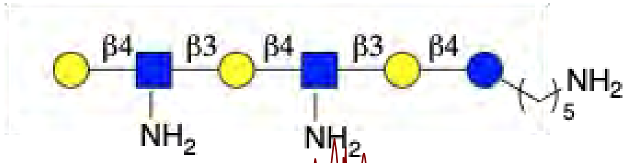
β4
β3
β4
β3
β4
NH2
NH2
NH2
5

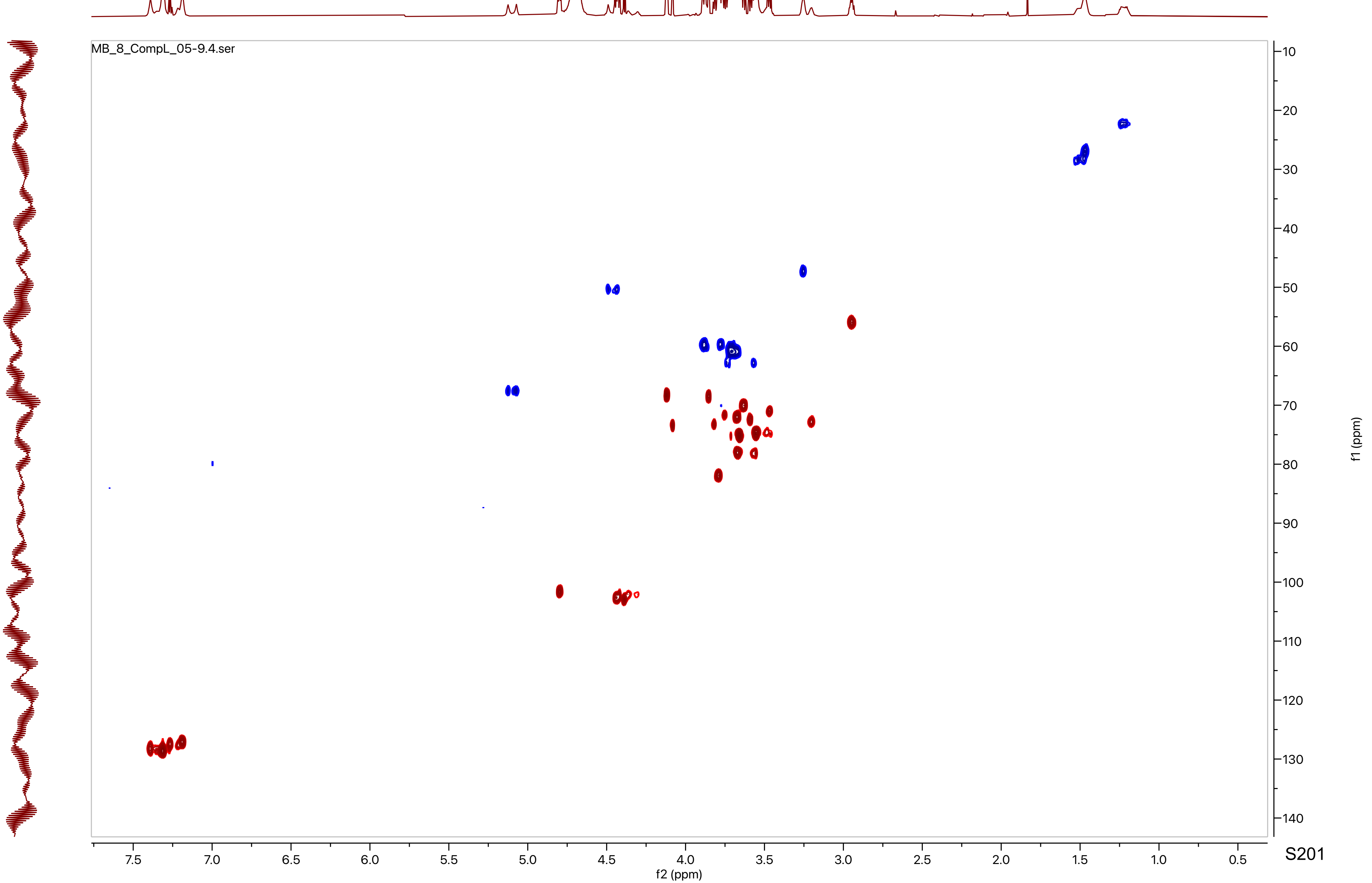
MB_8_CompL_05-9.4.ser
f2 (ppm)
f1 (ppm)

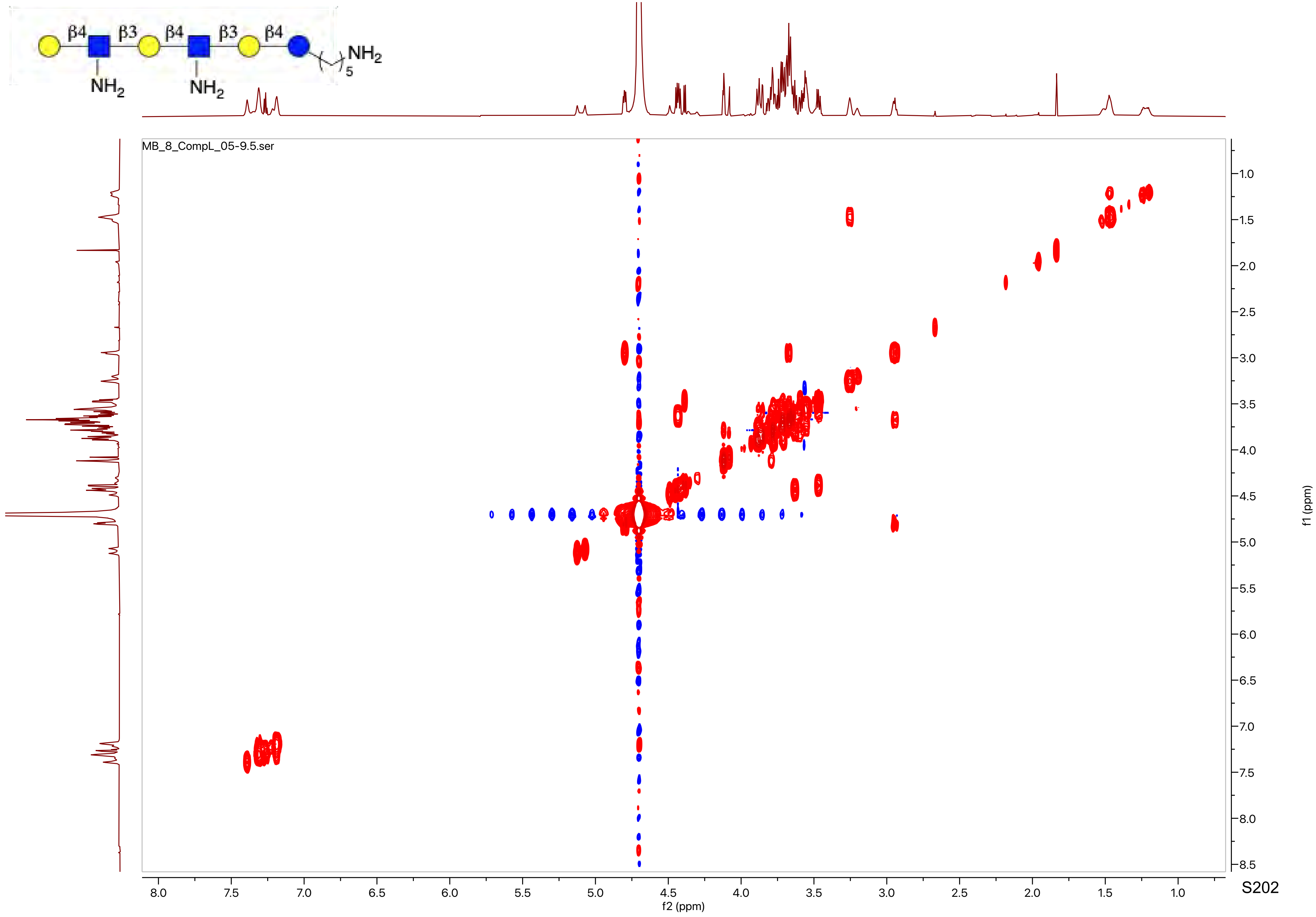

β4
β3
β4
β3
β4
NH2
5
NH2
NH2
MB_8_CompL_05-9.5.ser
f2 (ppm)
f1 (ppm)

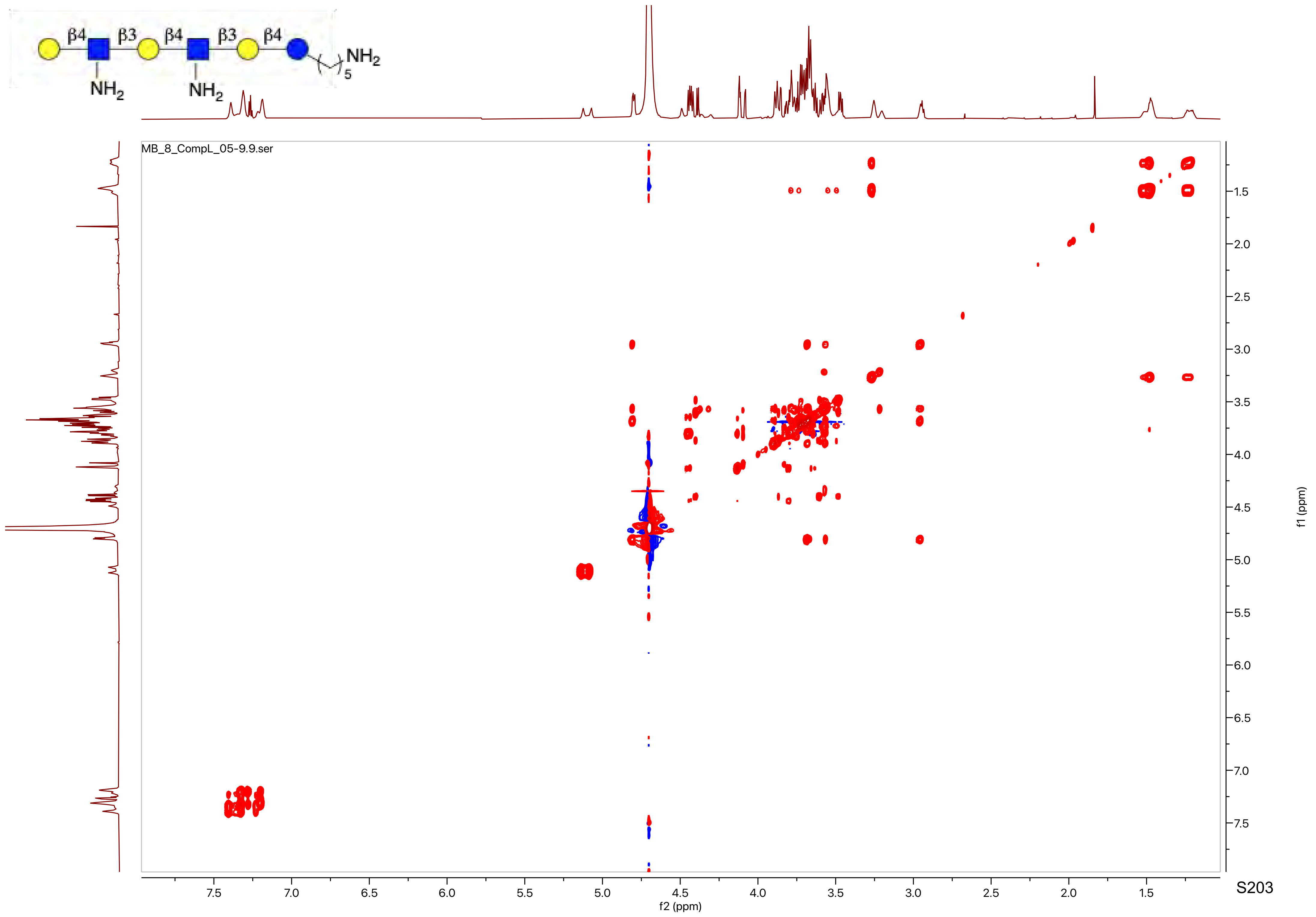
β4
β3
β4
β3
β4
NH2
5
NH2
NH2
MB_8_CompL_05-9.9.ser
f2 (ppm)
f1 (ppm)

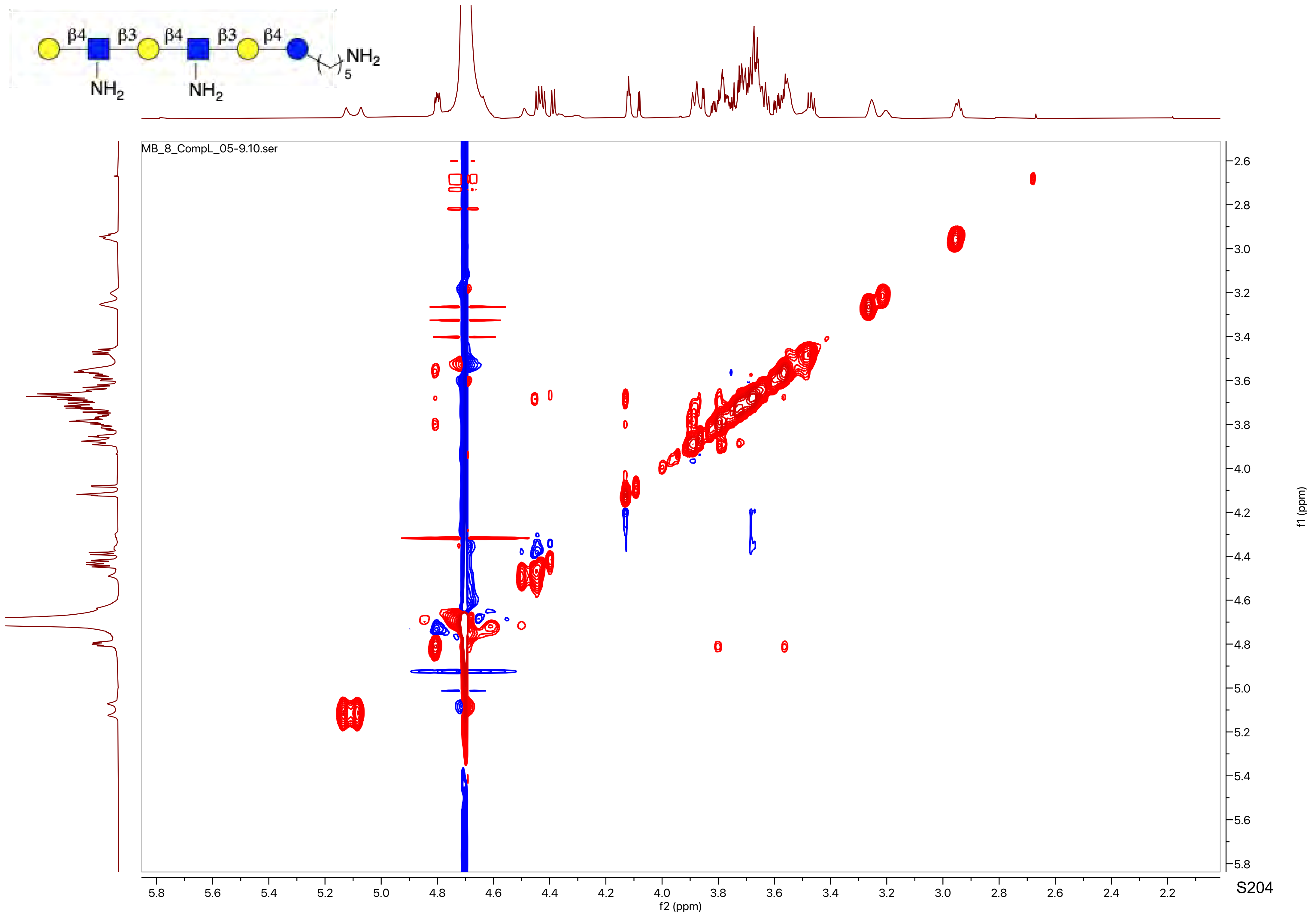
β4
β3
β4
β3
β4
NH2
5
NH2
NH2
MB_8_CompL_05-9.10.ser
f2 (ppm)
f1 (ppm)

MB_8_CL_06_C18_PROTON_20220704_02
MB_8_CL_06_C18

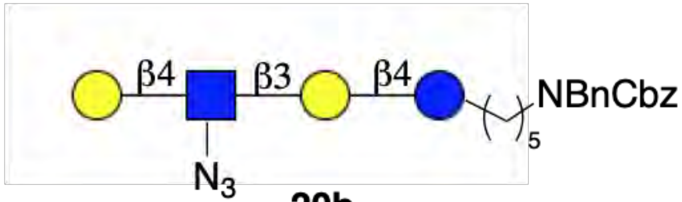


**20b**

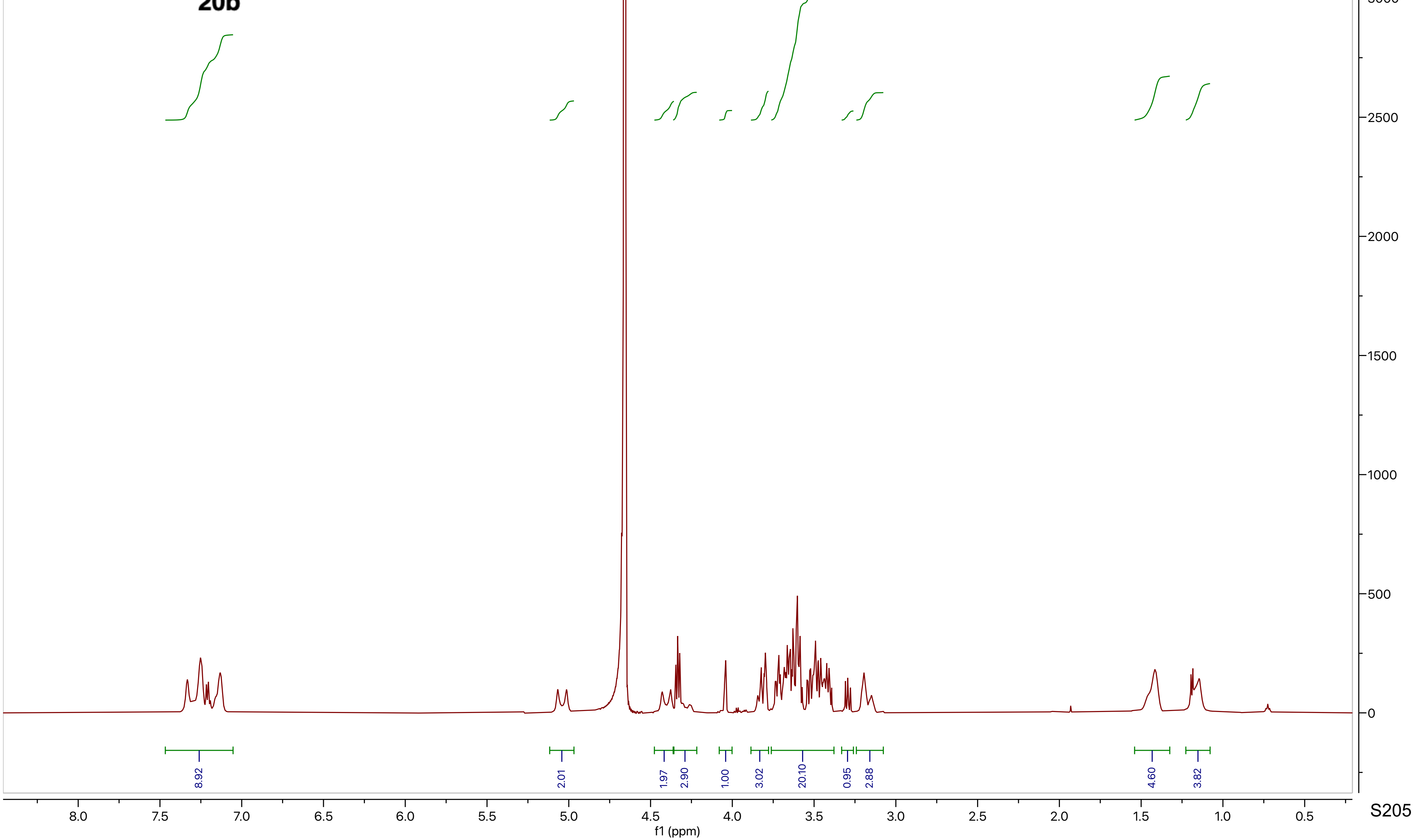

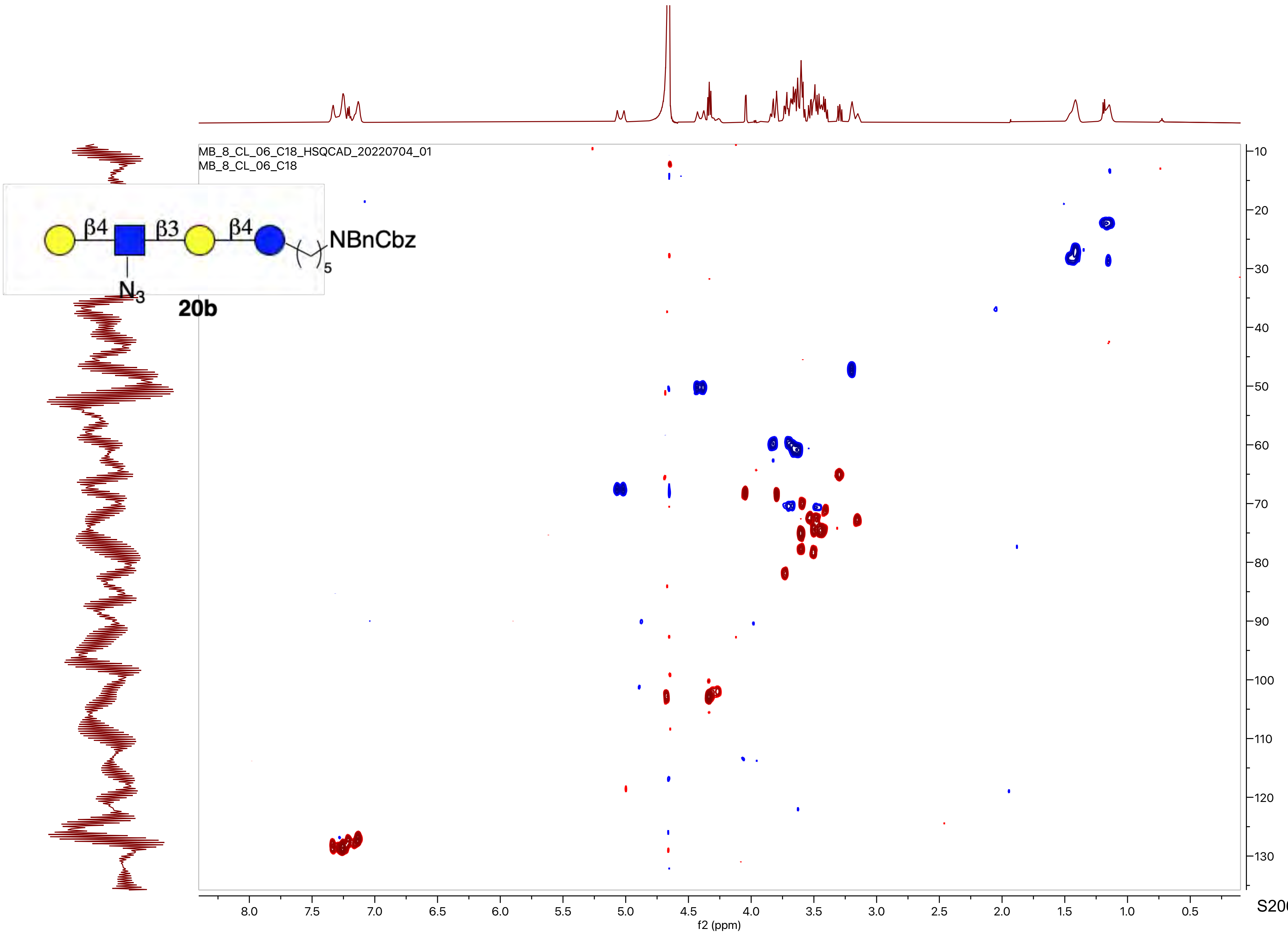

MB_8_CL_06_C18_HSQCAD_20220704_01
MB_8_CL_06_C18
β4
β3
β4
NBnCbz
5
N3
20b
f2 (ppm)
f1 (ppm)

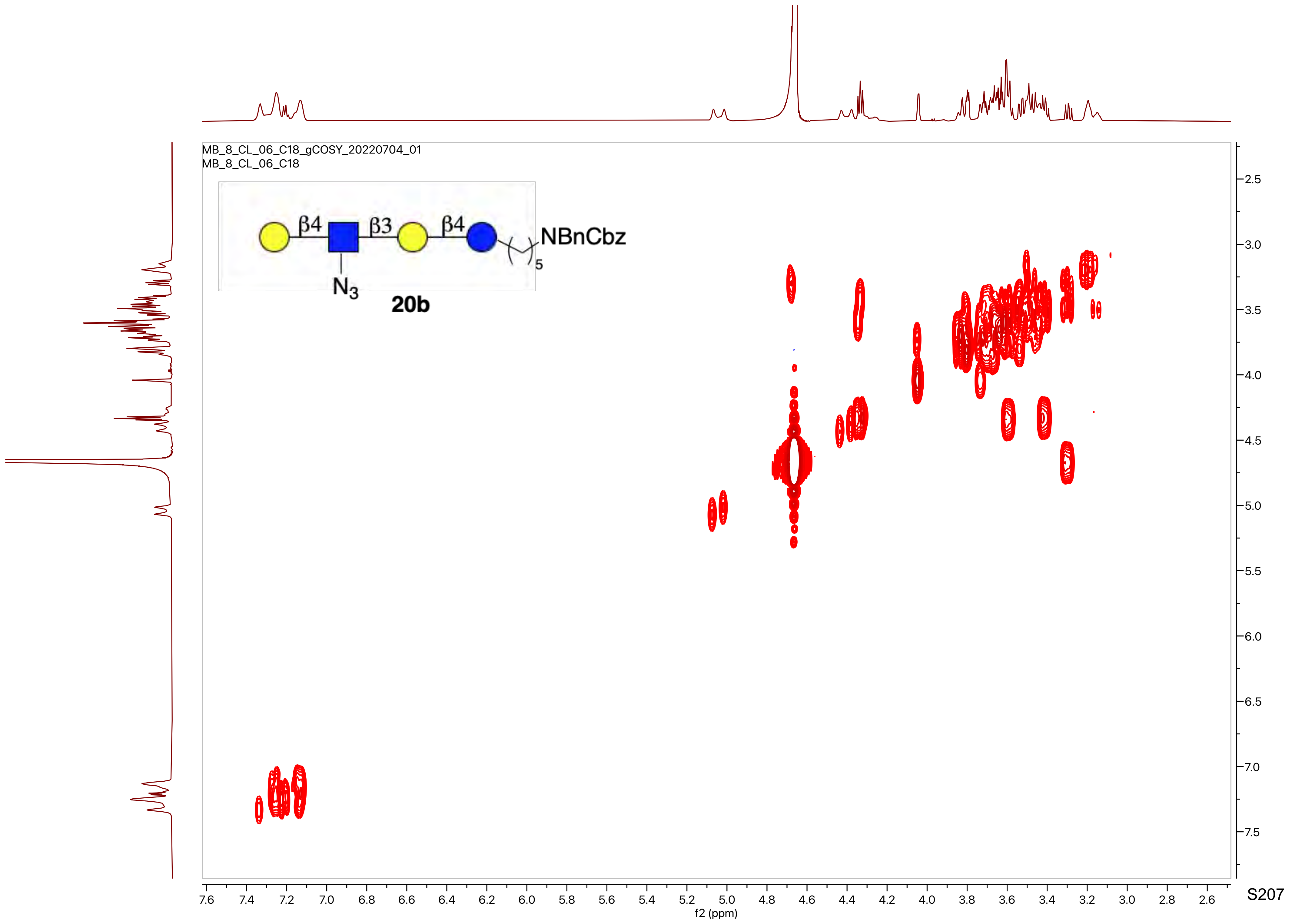

MB_8_CL_06_C18_gCOSY_20220704_01
MB_8_CL_06_C18
β4
β3
β4
NBnCbz
5
$N_3$
20b
f2 (ppm)
f1 (ppm)

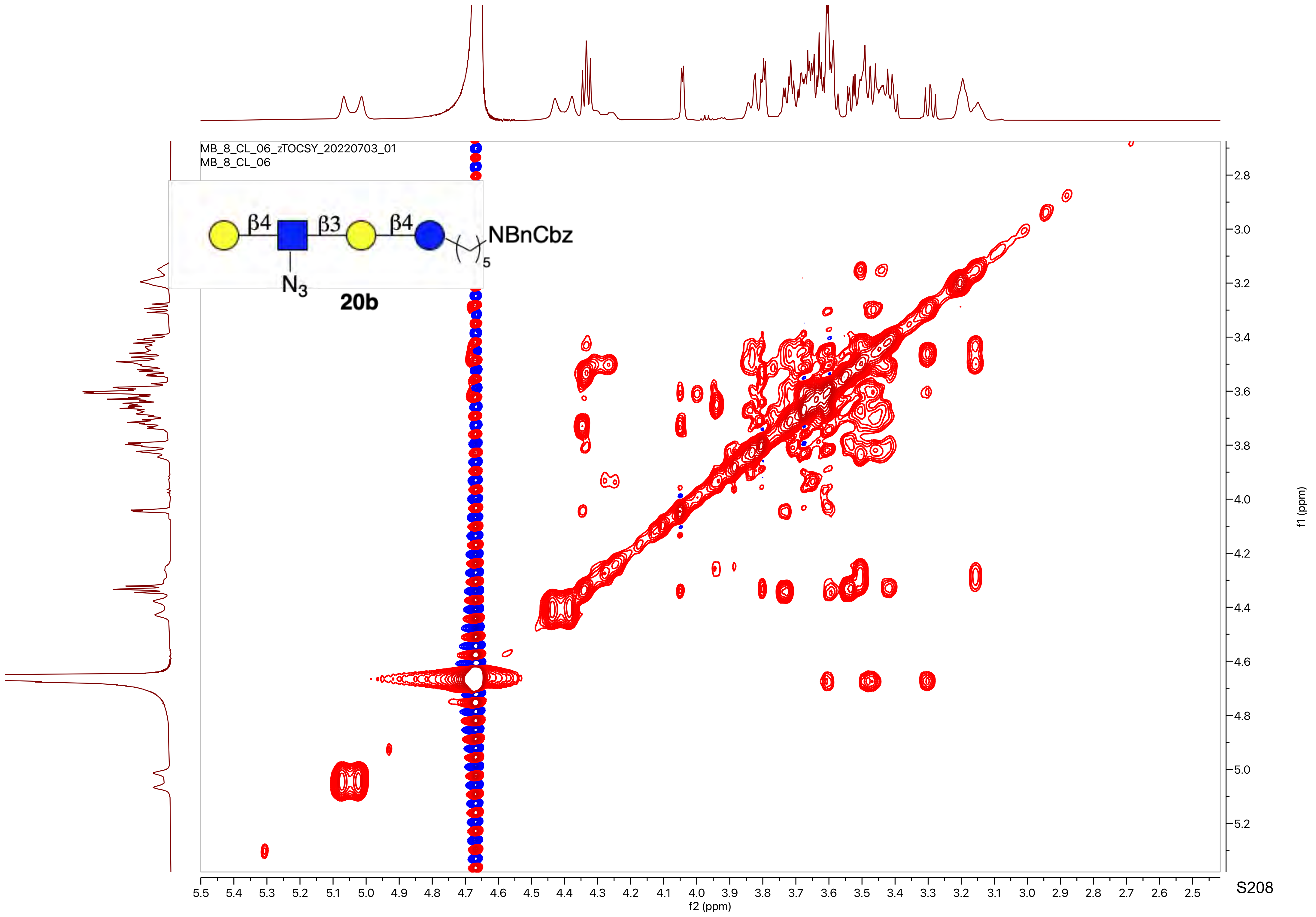

MB_8_CL_06_zTOCSY_20220703_01
MB_8_CL_06
β4
β3
β4
NBnCbz
5
N3
20b
f1 (ppm)
f2 (ppm)

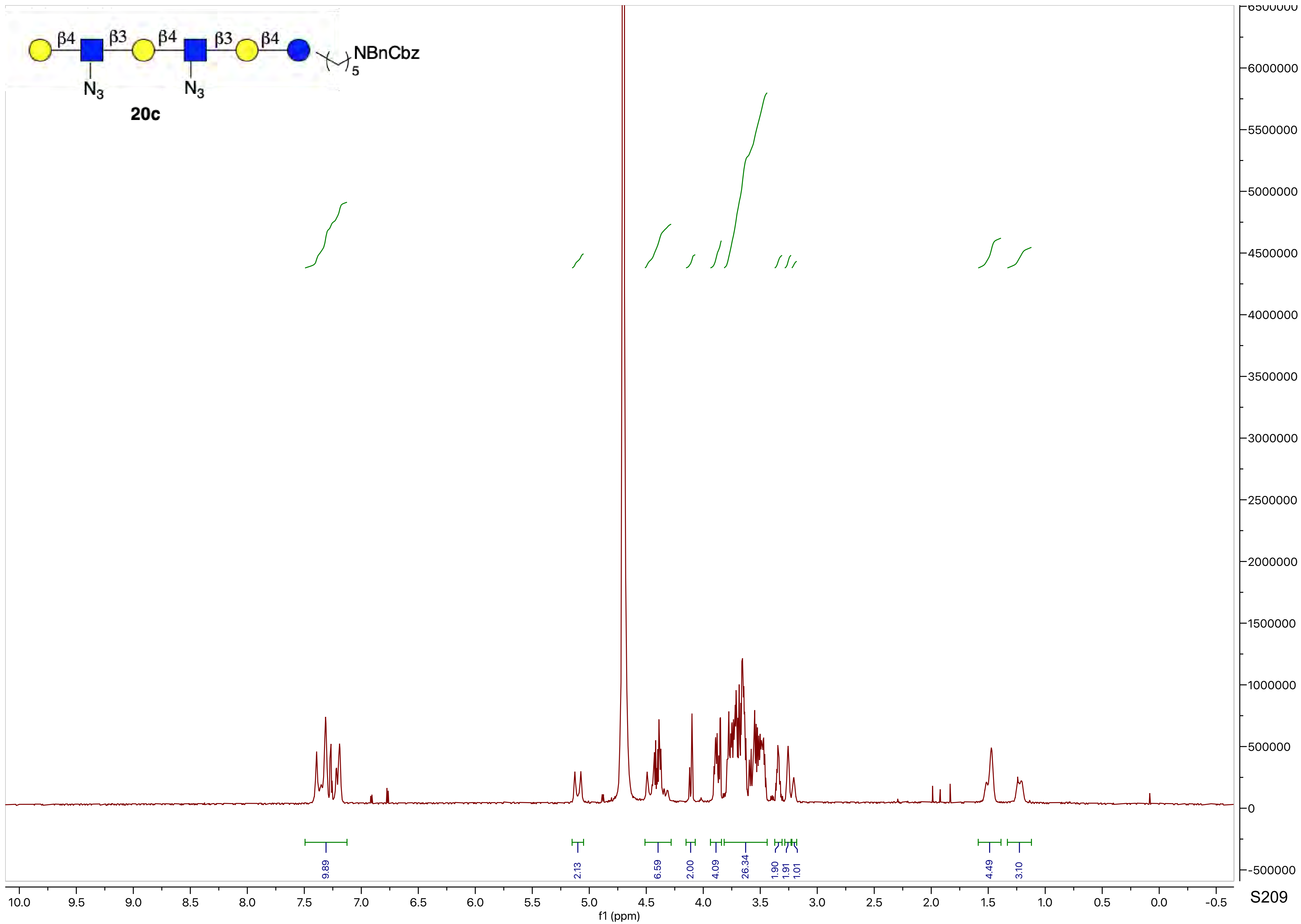

β4
β3
β4
β3
β4
NBnCbz
5
N3
N3
20c
9.89
2.13
6.59
2.00
4.09
26.34
1.90
1.91
1.01
4.49
3.10
10.0
9.5
9.0
8.5
8.0
7.5
7.0
6.5
6.0
5.5
5.0
4.5
4.0
3.5
3.0
2.5
2.0
1.5
1.0
0.5
0.0
-0.5
f1 (ppm)
6500000
6000000
5500000
5000000
4500000
4000000
3500000
3000000
2500000
2000000
1500000
1000000
500000
0
-500000

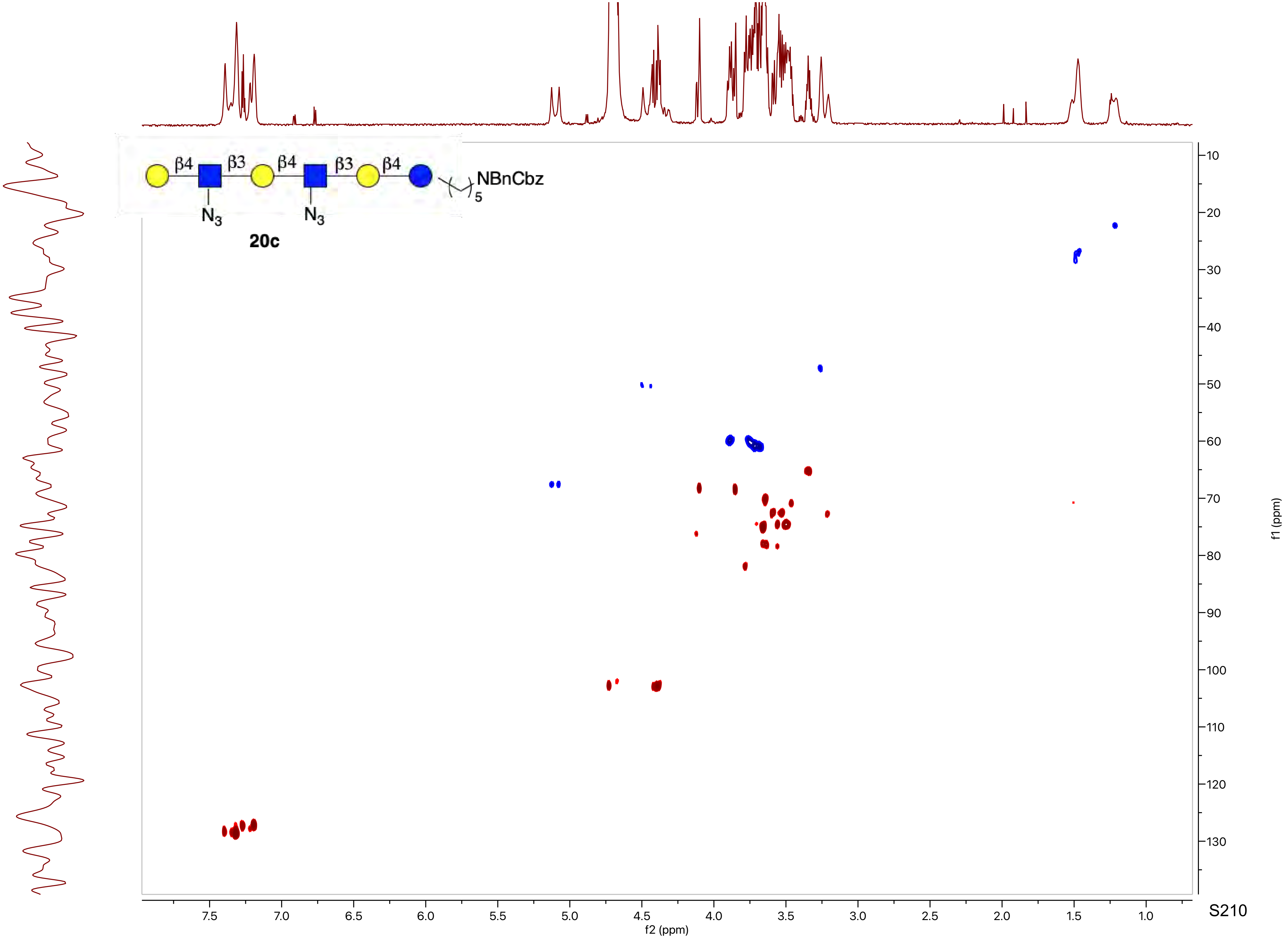
β4
β3
β4
β3
β4
N3
N3
NBnCbz
5
20c
f2 (ppm)
f1 (ppm)
7.5
7.0
6.5
6.0
5.5
5.0
4.5
4.0
3.5
3.0
2.5
2.0
1.5
1.0
10
20
30
40
50
60
70
80
90
100
110
120
130

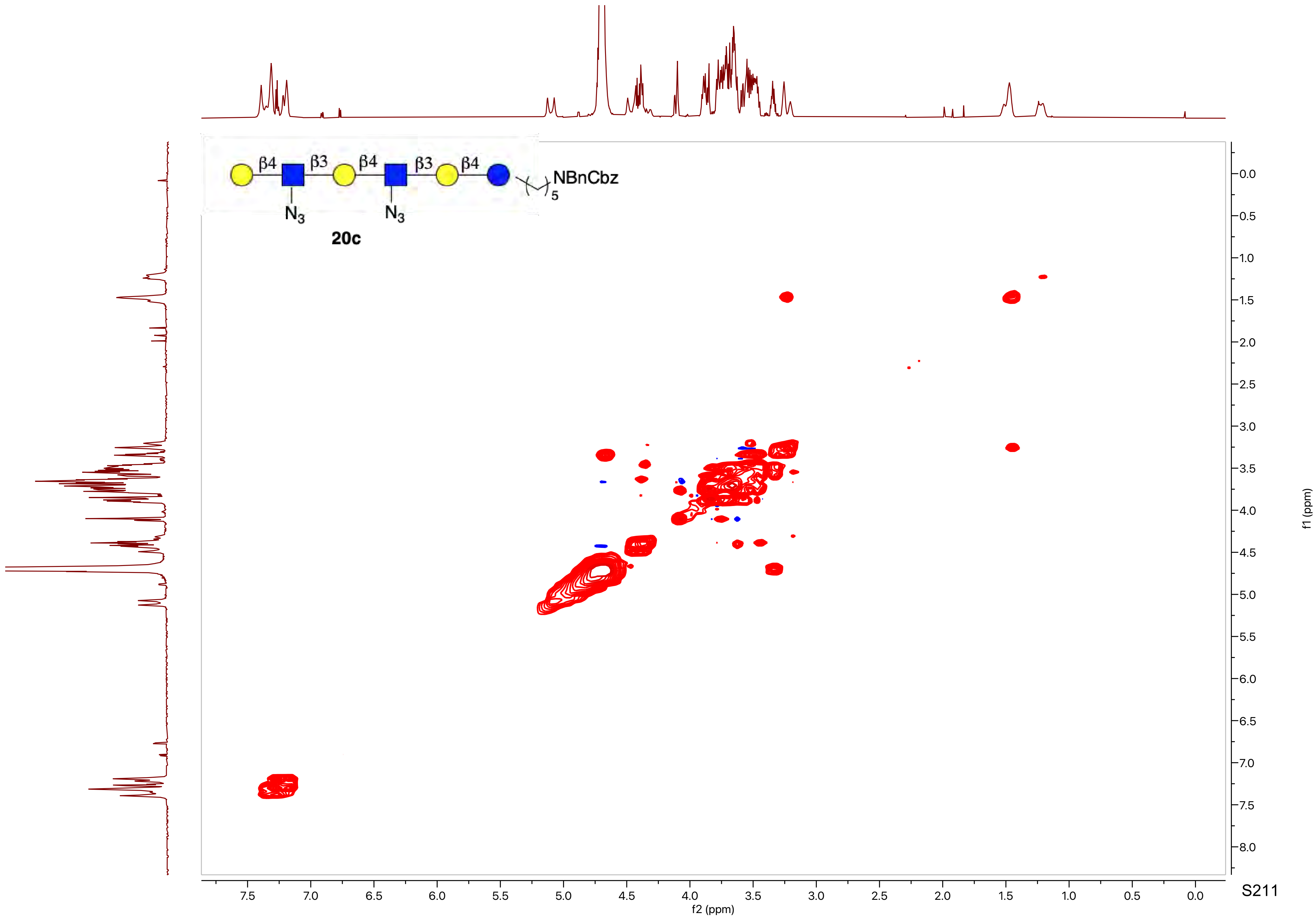
β4
β3
β4
β3
β4
N3
N3
NBnCbz
5
20c
f2 (ppm)
f1 (ppm)

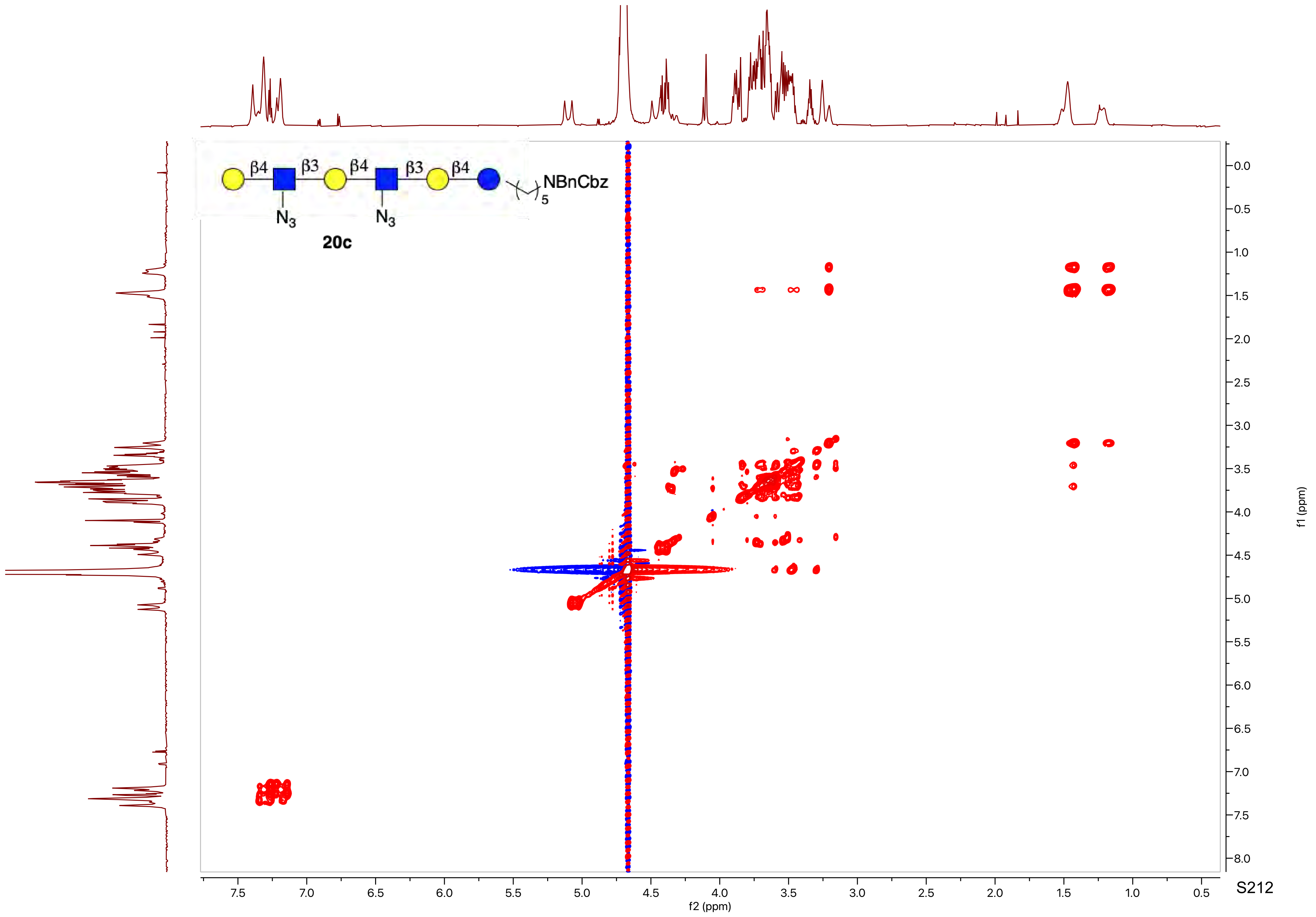
β4
β3
β4
β3
β4
N3
N3
NBnCbz
5
20c
f1 (ppm)
f2 (ppm)

MB_VIII_77.10.fid
PROTON MeOD {C:\nmrdata\CBDD} Mehman 16

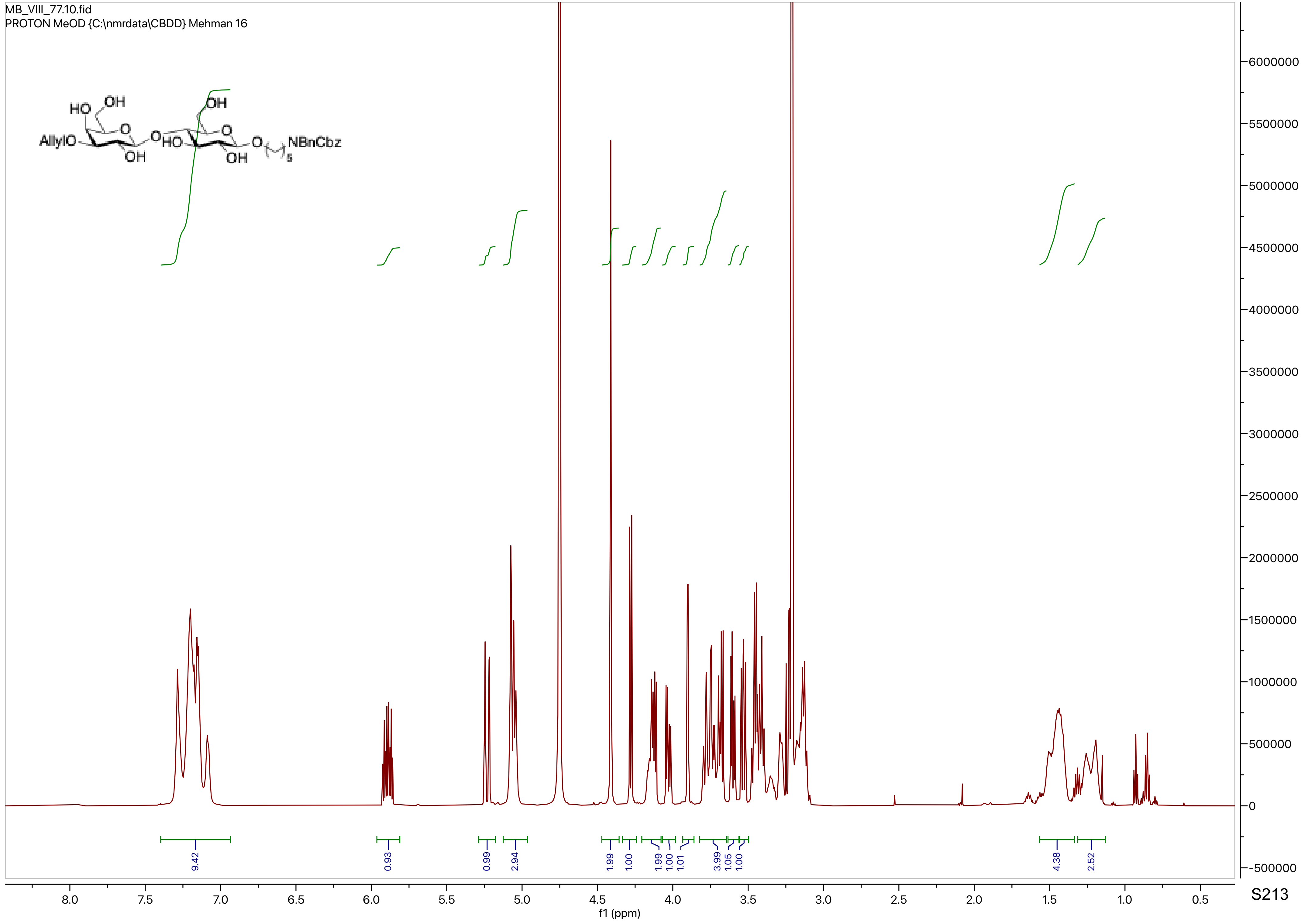

MB_VIII_77.12.fid
C13APT MeOD {C:\nmrdata\CBDD} Mehman 16

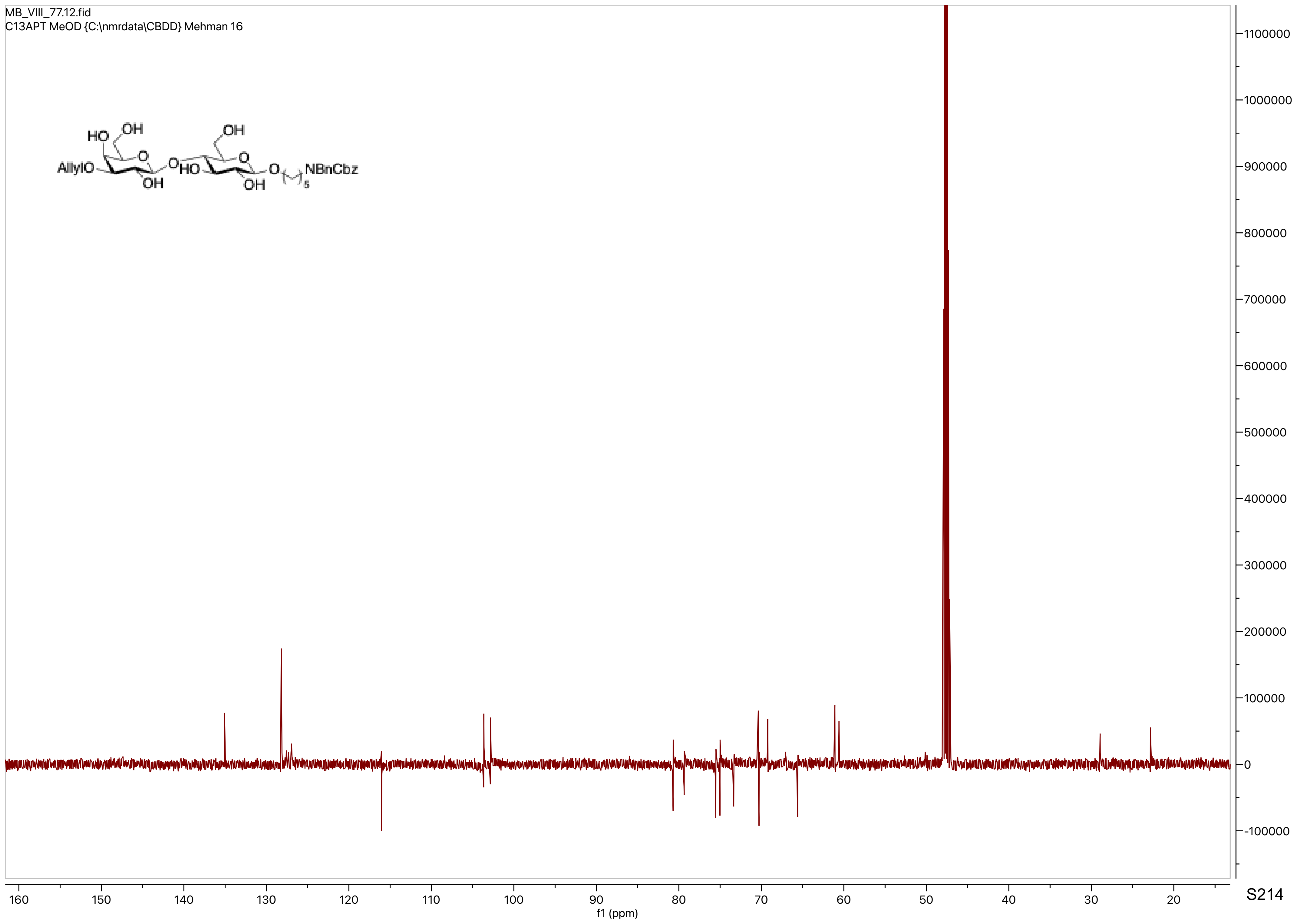

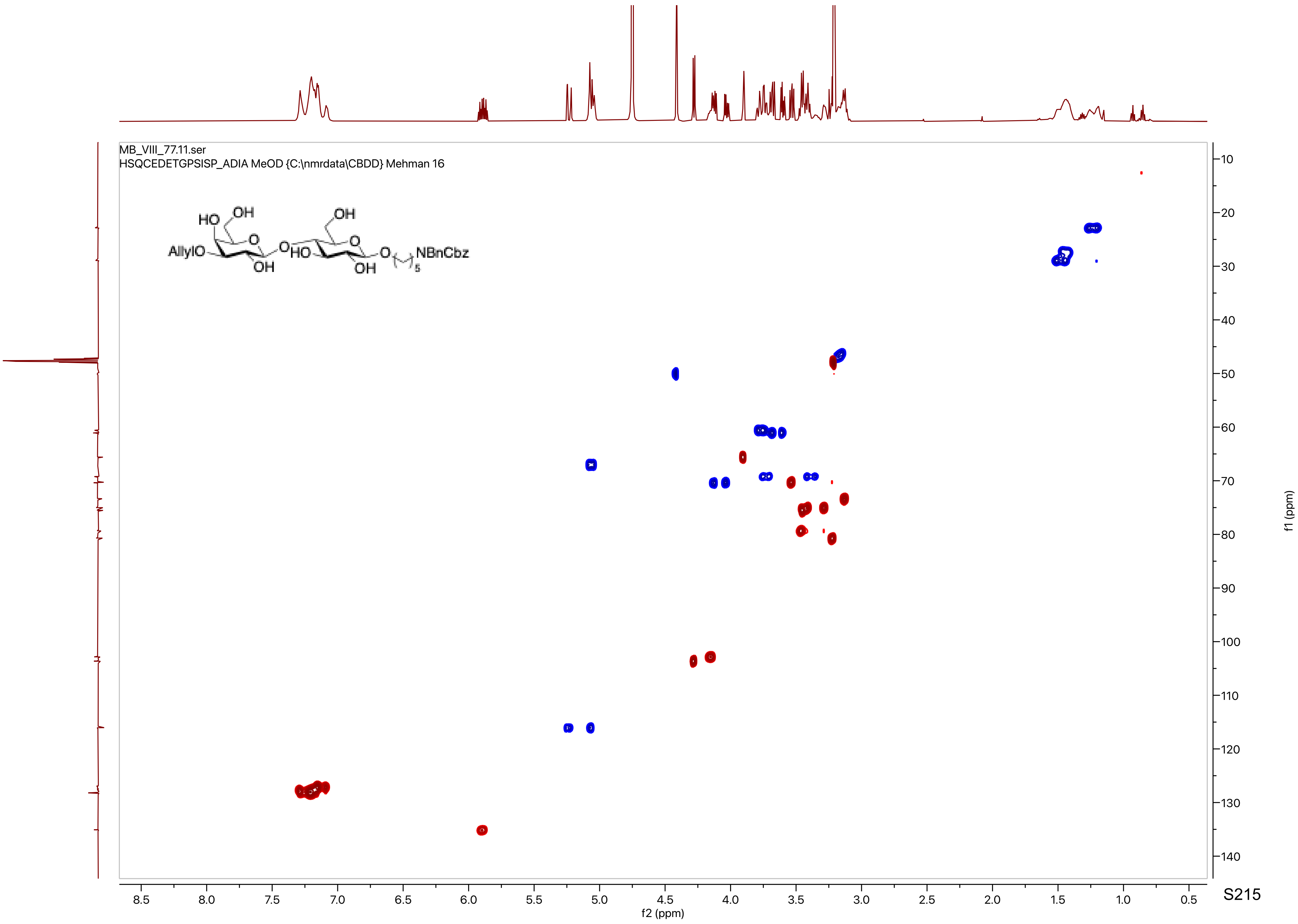

MB_VIII_77.11.ser
HSQCEDETGPSISP_ADIA MeOD {C:\nmrdata\CBDD} Mehman 16
AllylO
HO
OH
O
OH
HO
OH
O
O
OH
O
NBnCbz
5
f2 (ppm)
f1 (ppm)

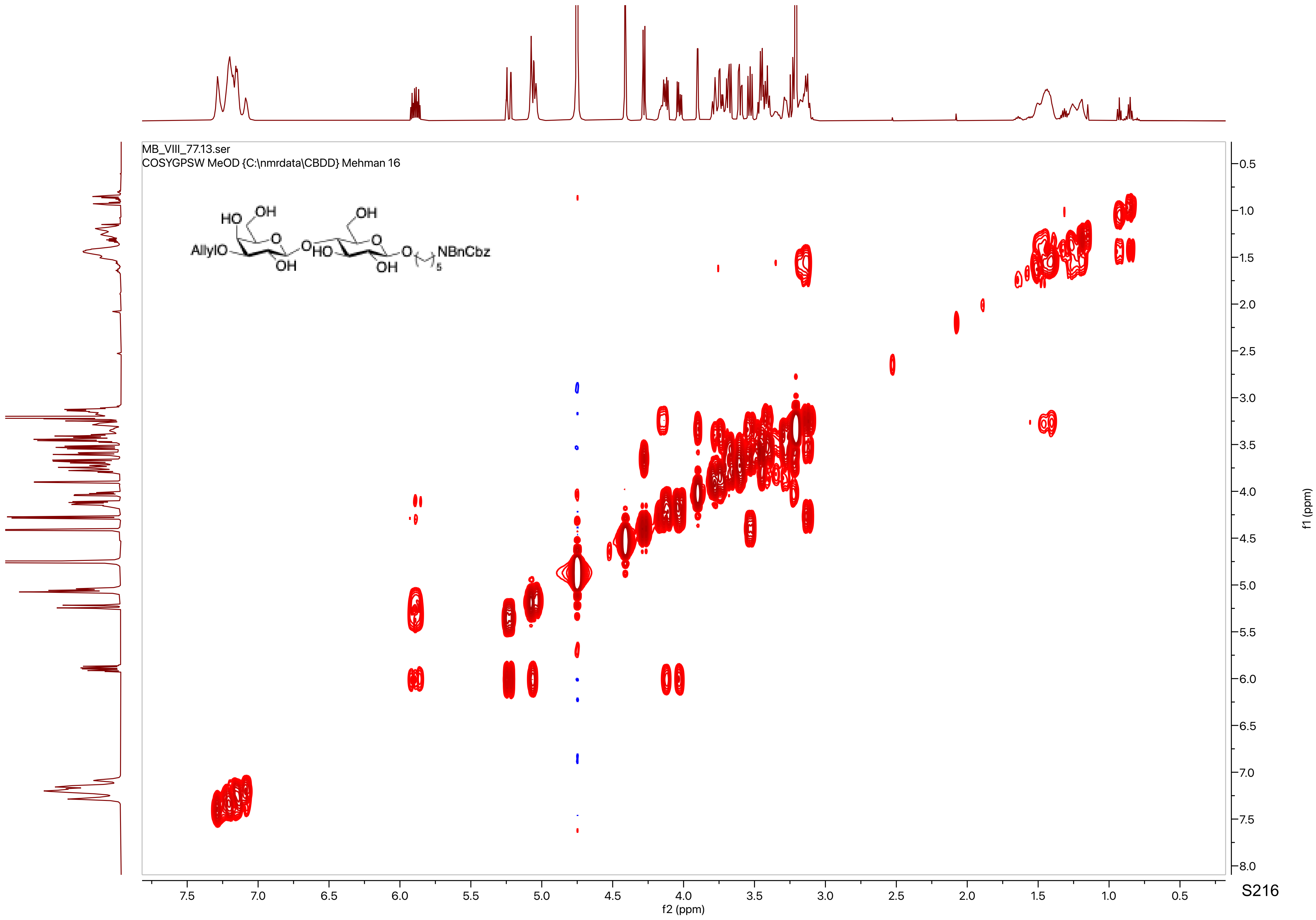
MB_VIII_77.13.ser
COSYGPSW MeOD {C:\nmrdata\CBDD} Mehman 16
HO
OH
OH
AllylO
O
O
O
HO
O
NBnCbz
OH
OH
5
f2 (ppm)
f1 (ppm)
7.5
7.0
6.5
6.0
5.5
5.0
4.5
4.0
3.5
3.0
2.5
2.0
1.5
1.0
0.5
8.0

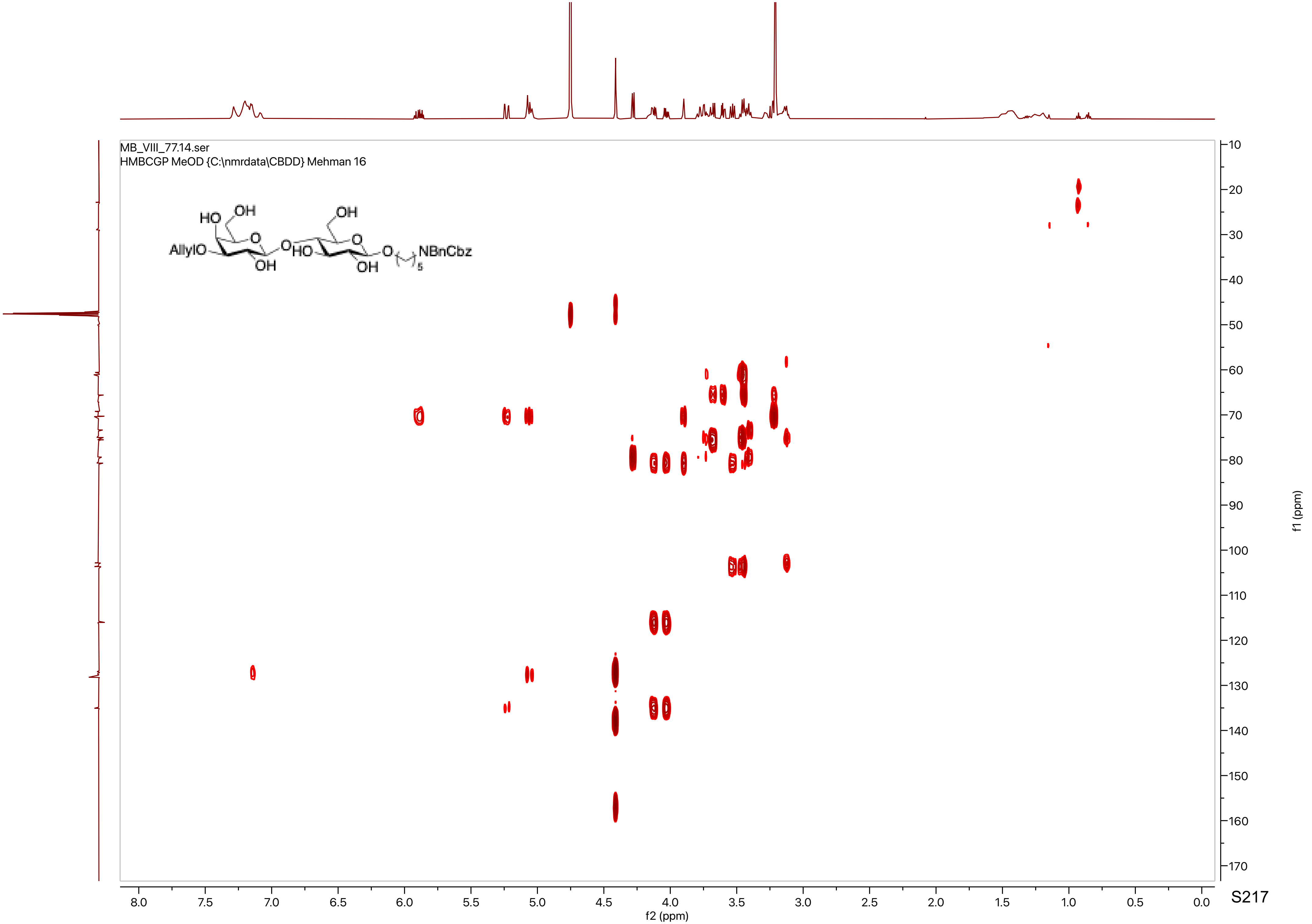

MB_VIII_77.14.ser
HMBCGP MeOD {C:\nmrdata\CBDD} Mehman 16
HO
OH
OH
AllylO
O
O
OH
HO
O
OH
O
NBnCbz
5
f2 (ppm)
f1 (ppm)

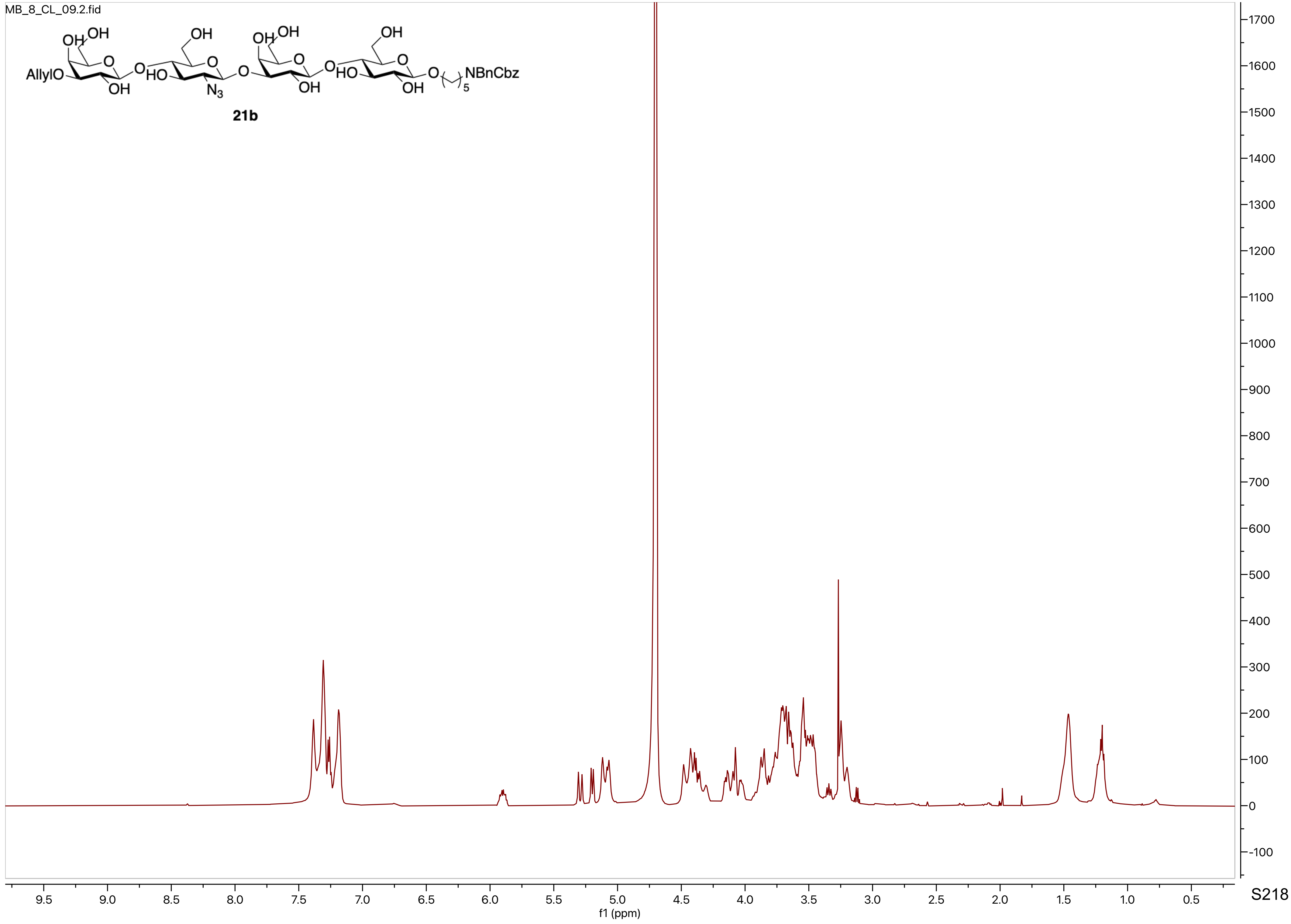
MB_8_CL_09.2.fid
AllylO
NBnCbz
21b
f1 (ppm)

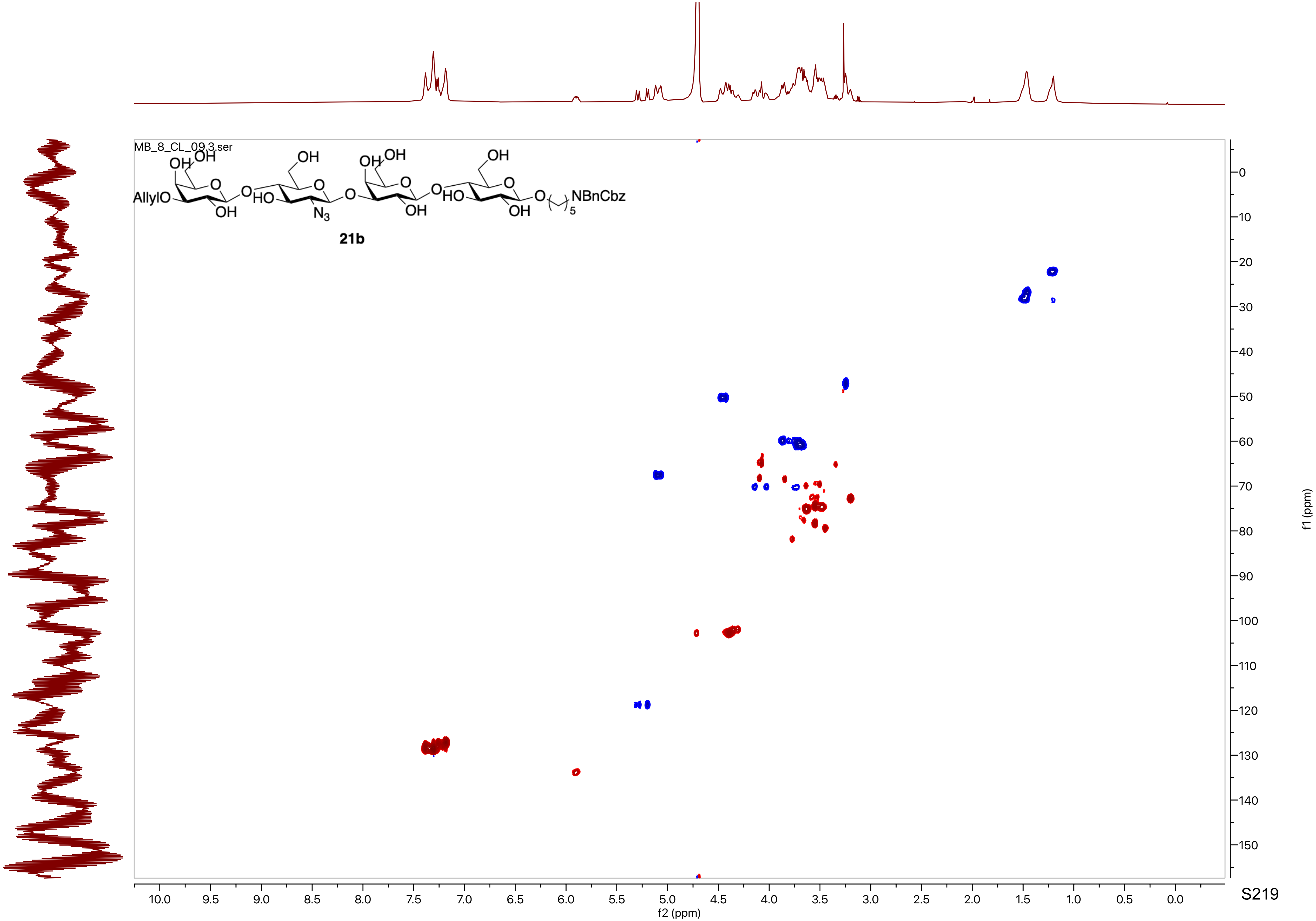

MB_8_CL_09.3.ser
AllylO
NBnCbz
21b
f2 (ppm)
f1 (ppm)

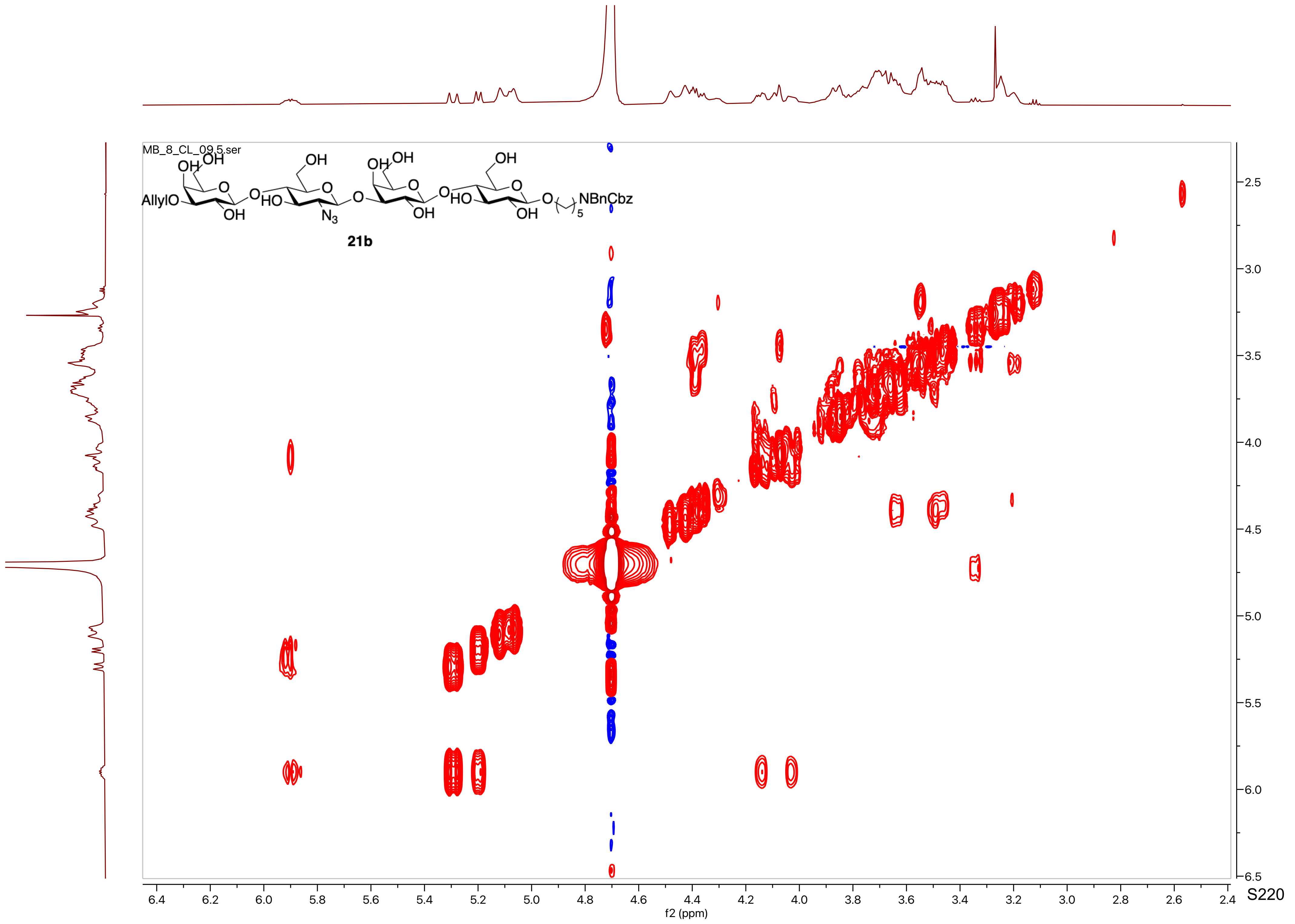

MB_8_CL_09.5.ser
AllylO
HO
N3
HO
NBnCbz
21b
f2 (ppm)
f1 (ppm)

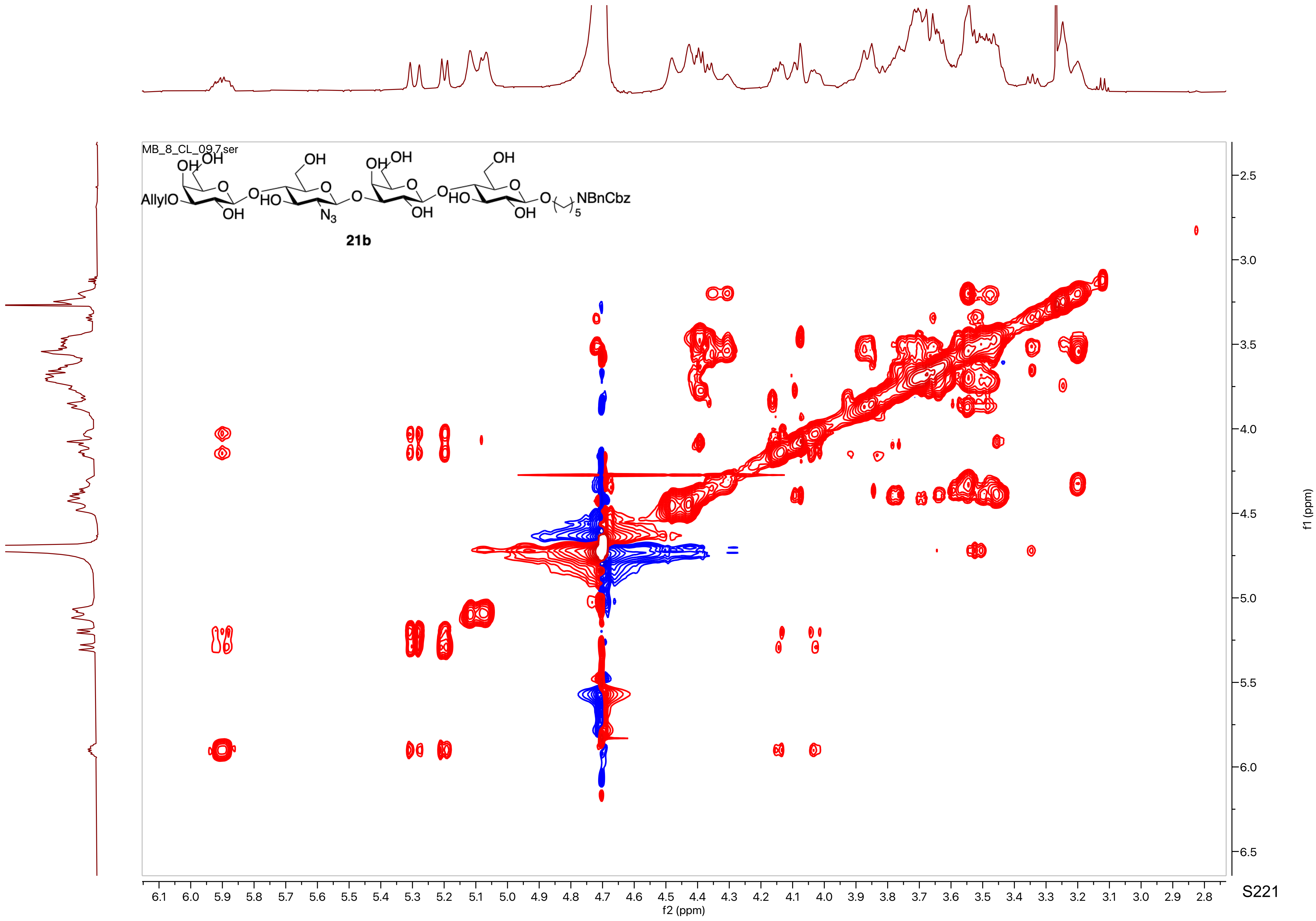
MB_8_CL_09.7.ser
AllylO
NBnCbz
21b
f2 (ppm)
f1 (ppm)

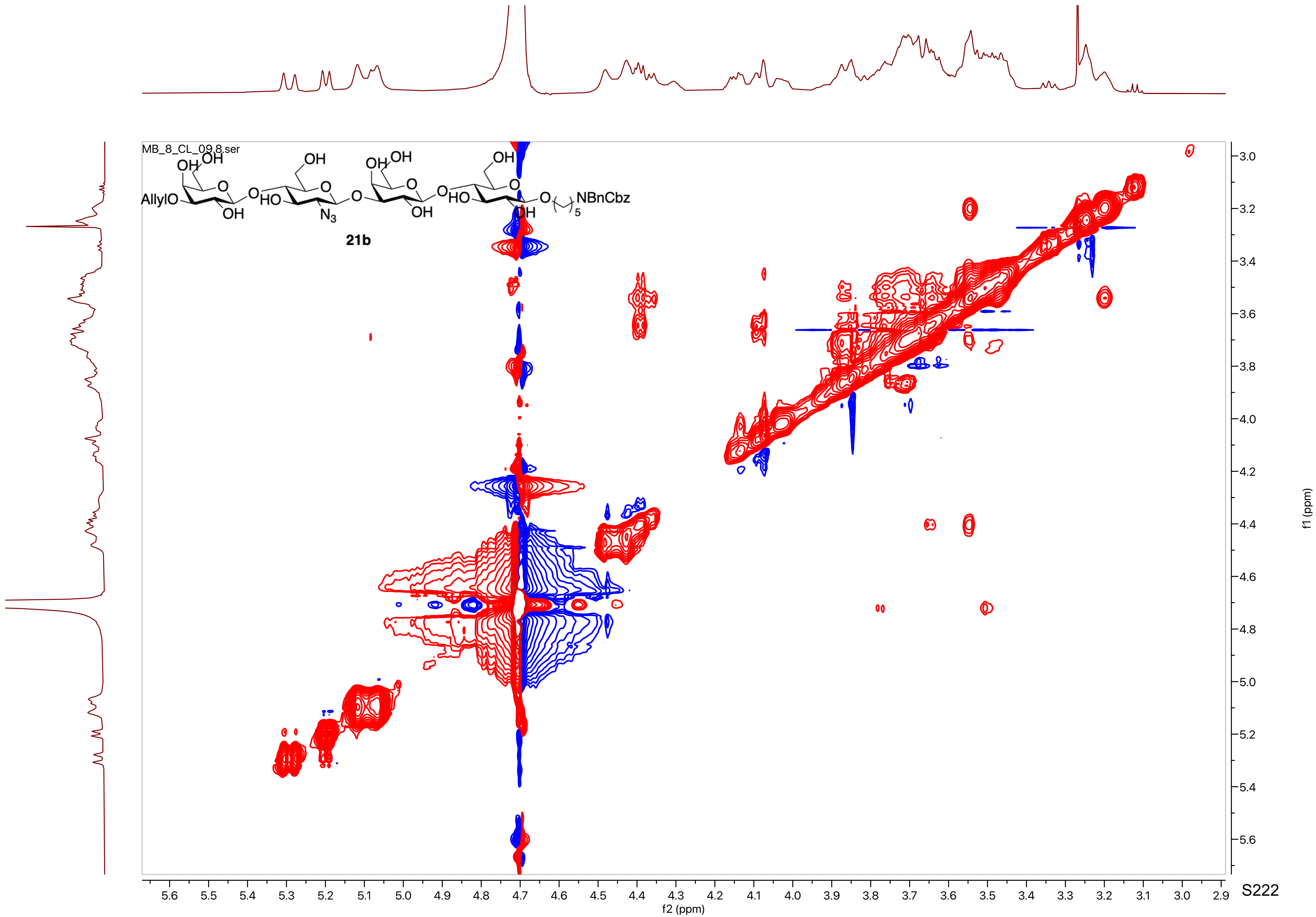
MB_8_CL_09.8.ser
AllylO
NBnCbz
21b
f2 (ppm)
f1 (ppm)

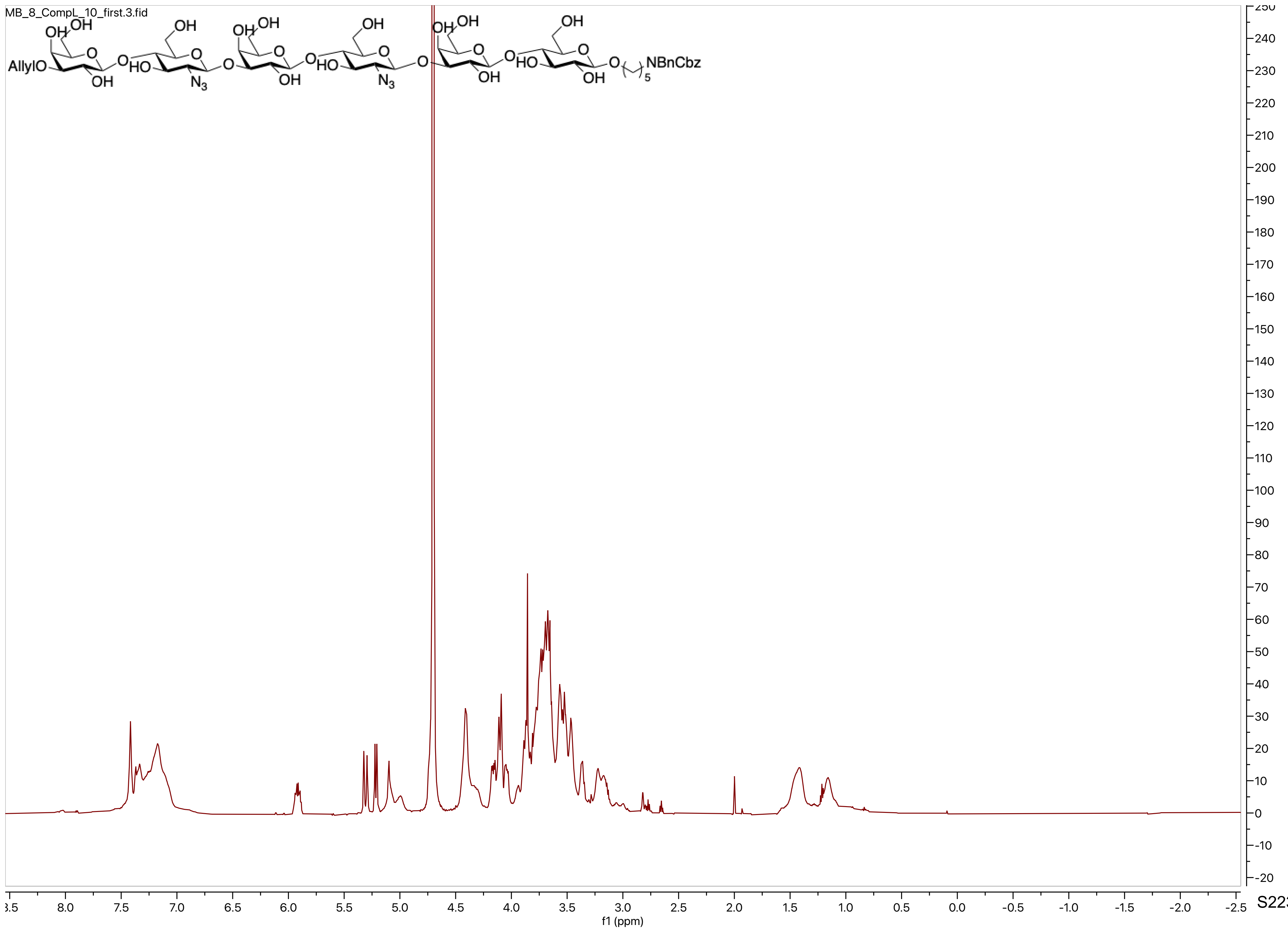
MB_8_CompL_10_first.3.fid
AllylO
NBnCbz
f1 (ppm)

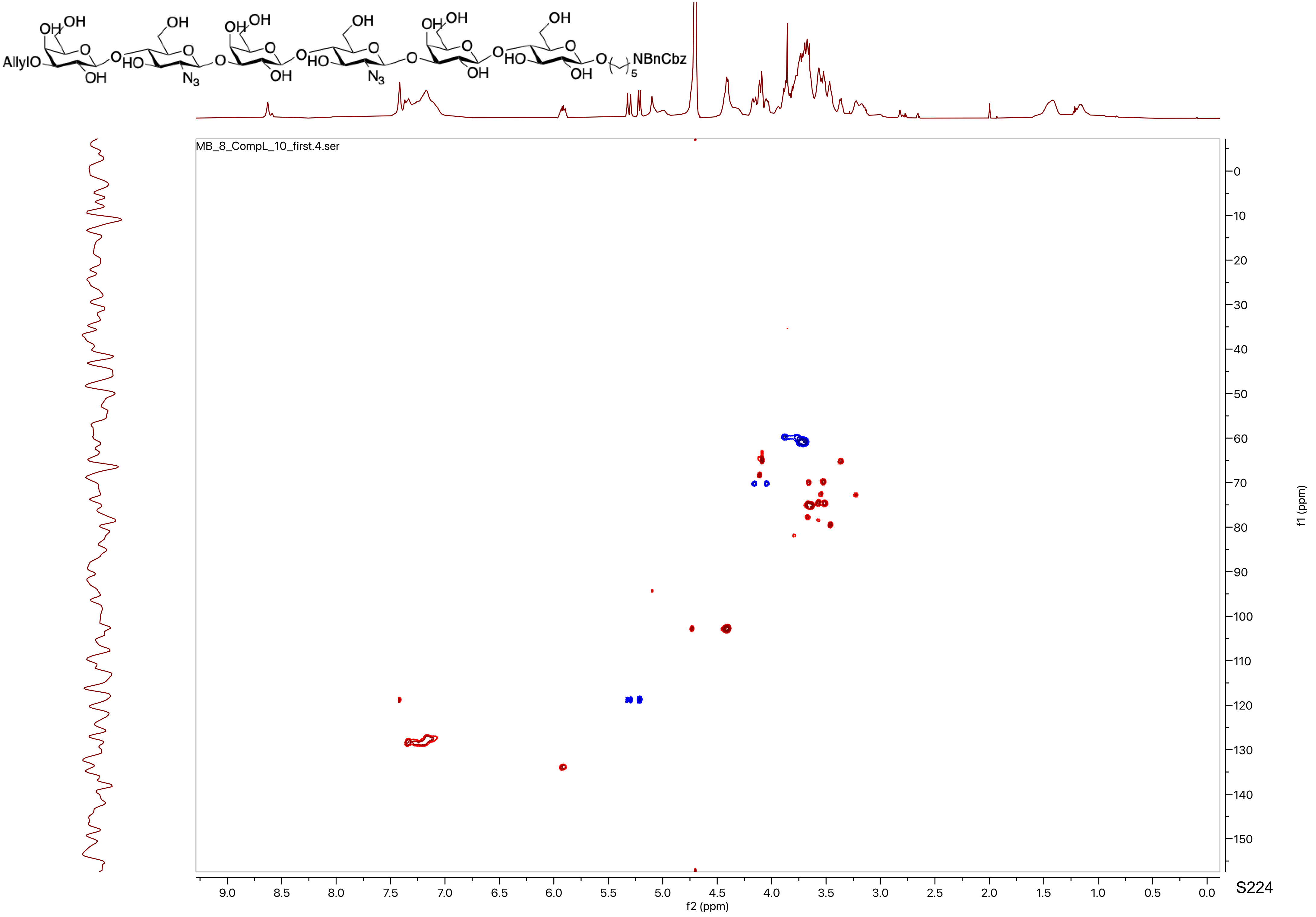

AllylO
NBnCbz
MB_8_CompL_10_first.4.ser
f2 (ppm)
f1 (ppm)

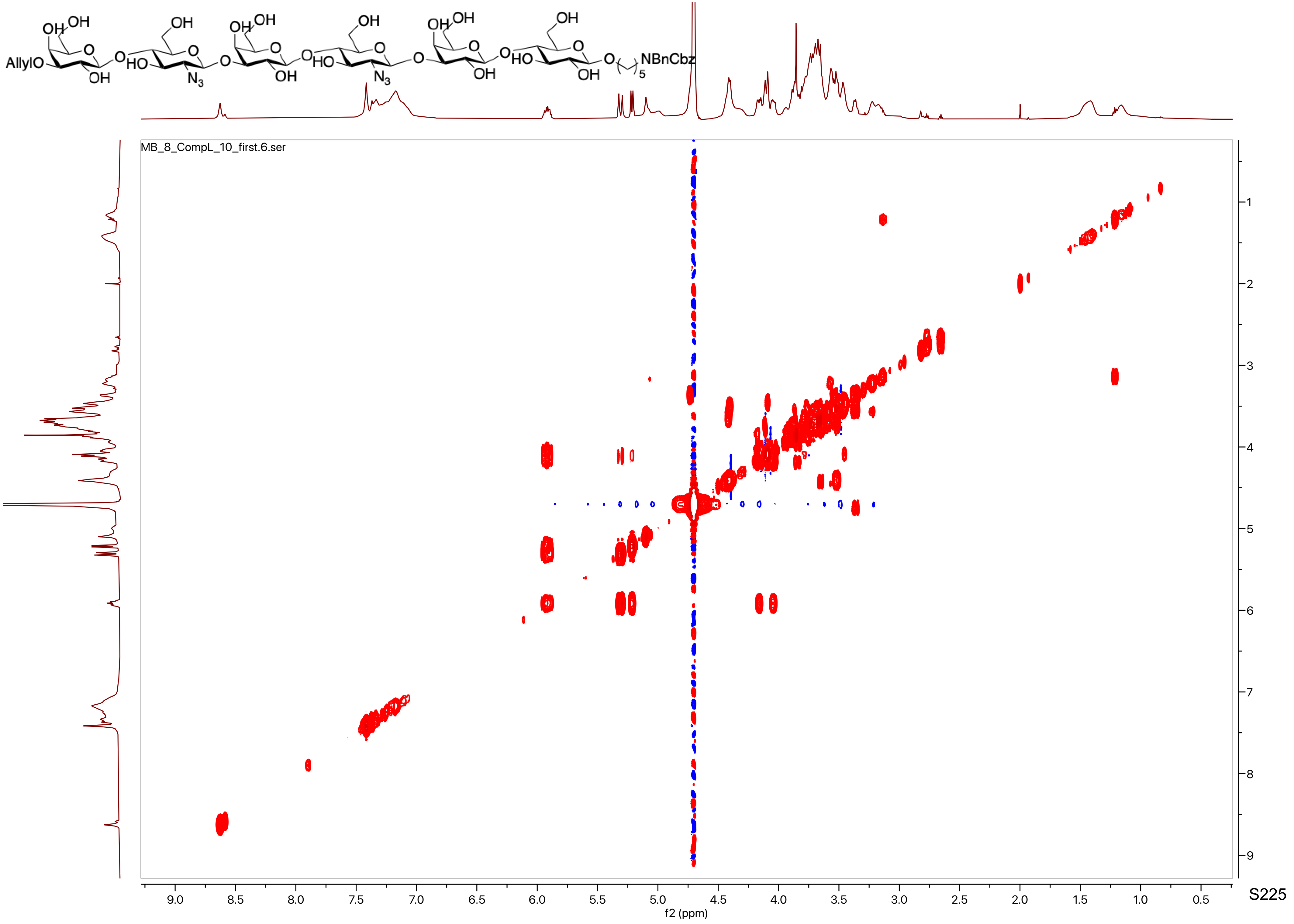

AllylO
NBnCbz
MB_8_CompL_10_first.6.ser
f2 (ppm)
f1 (ppm)

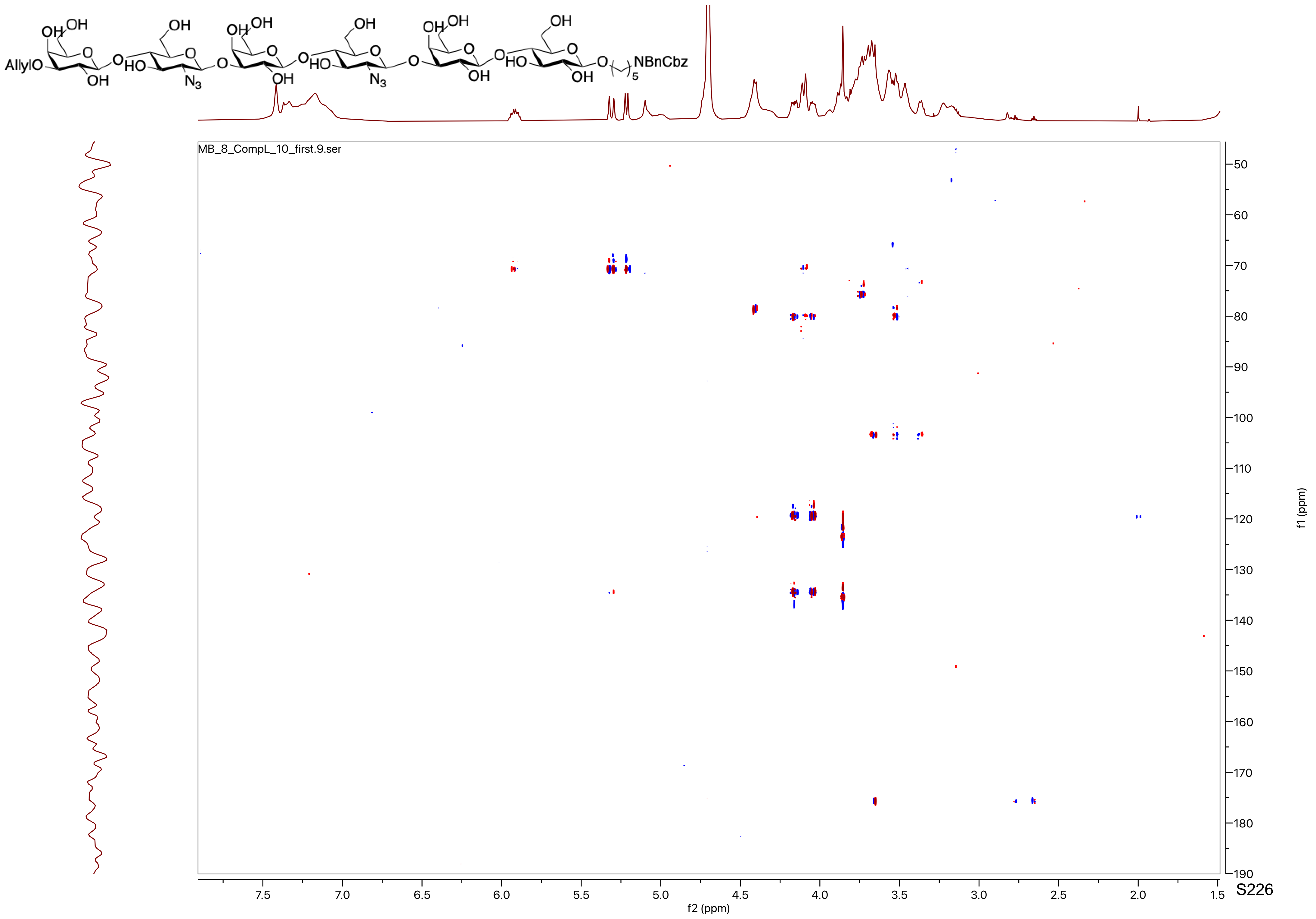

AllylO
NBnCbz
MB_8_CompL_10_first.9.ser
f2 (ppm)
f1 (ppm)

MB_8_CL_11_PROTON_20220627_01
MB_8_CL_11

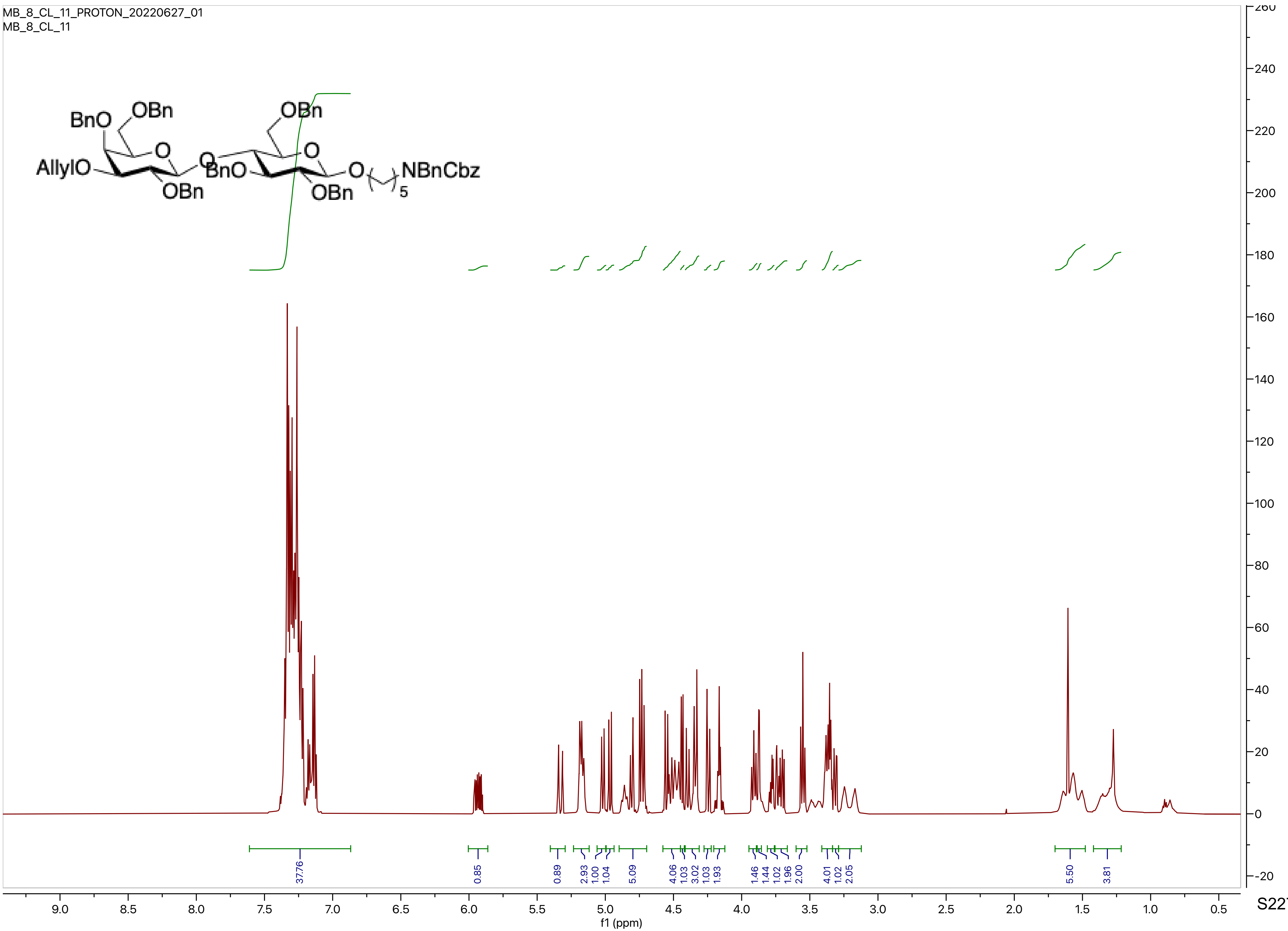

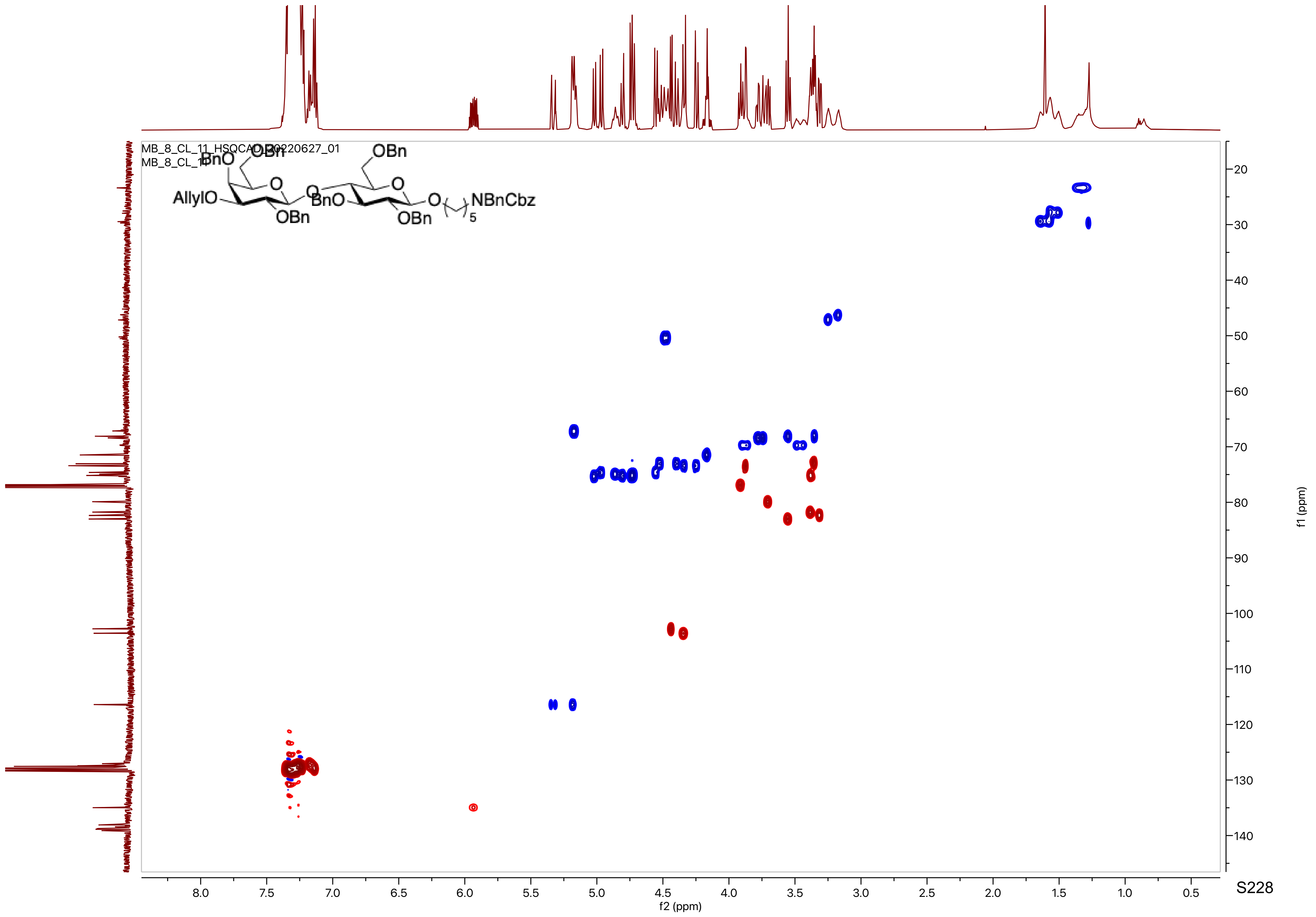
MB_8_CL_11_HSQCAD_20220627_01
MB_8_CL_11
BnO
OBn
OBn
AllylO
O
O
BnO
O
O
NBnCbz
5
OBn
OBn
f1 (ppm)
f2 (ppm)
20
30
40
50
60
70
80
90
100
110
120
130
140
8.0
7.5
7.0
6.5
6.0
5.5
5.0
4.5
4.0
3.5
3.0
2.5
2.0
1.5
1.0
0.5

MB_8_CL_11_CARBON_20220627_01
MB_8_CL_11

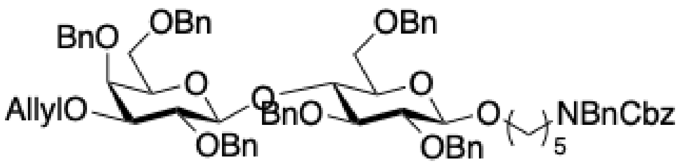

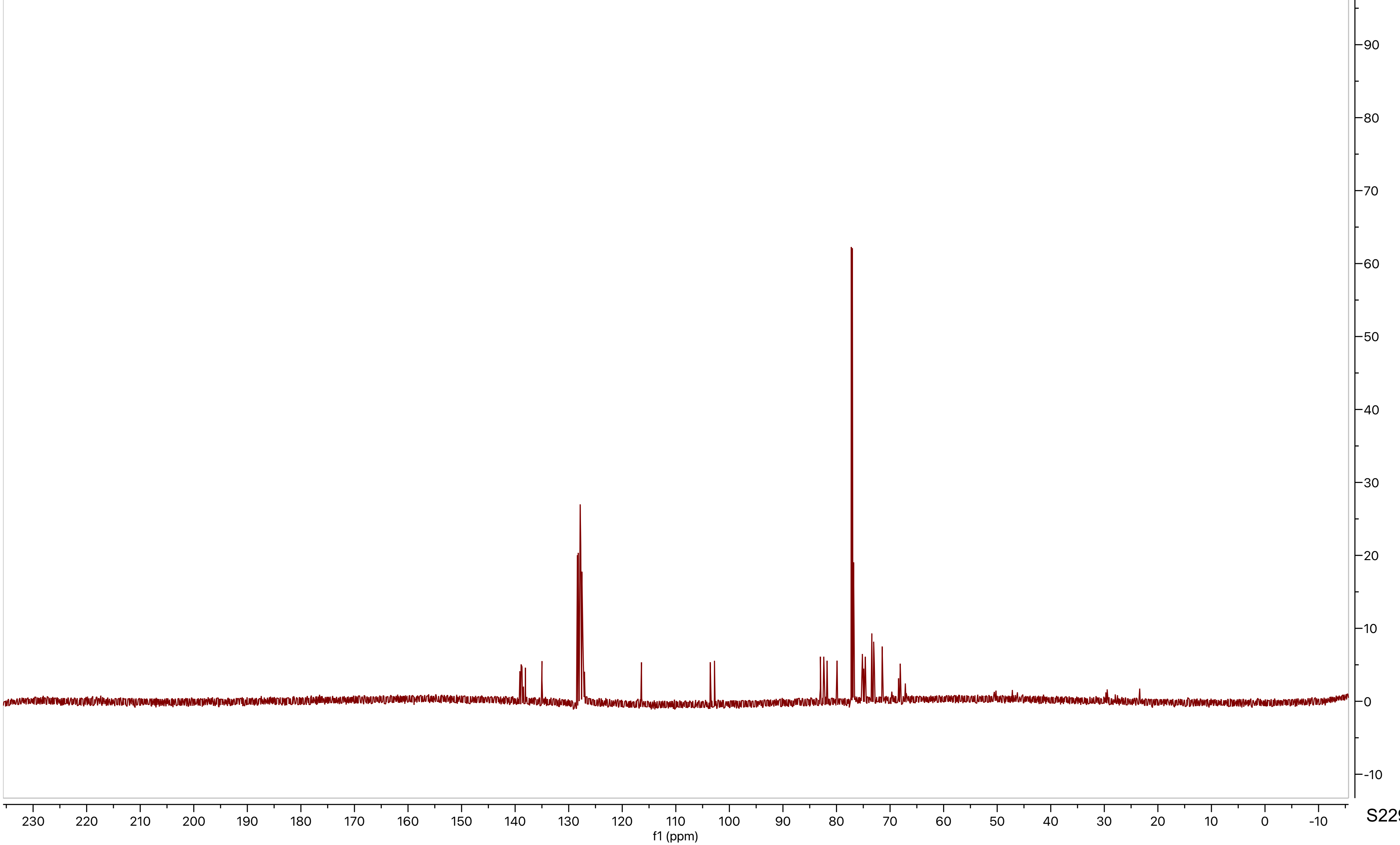

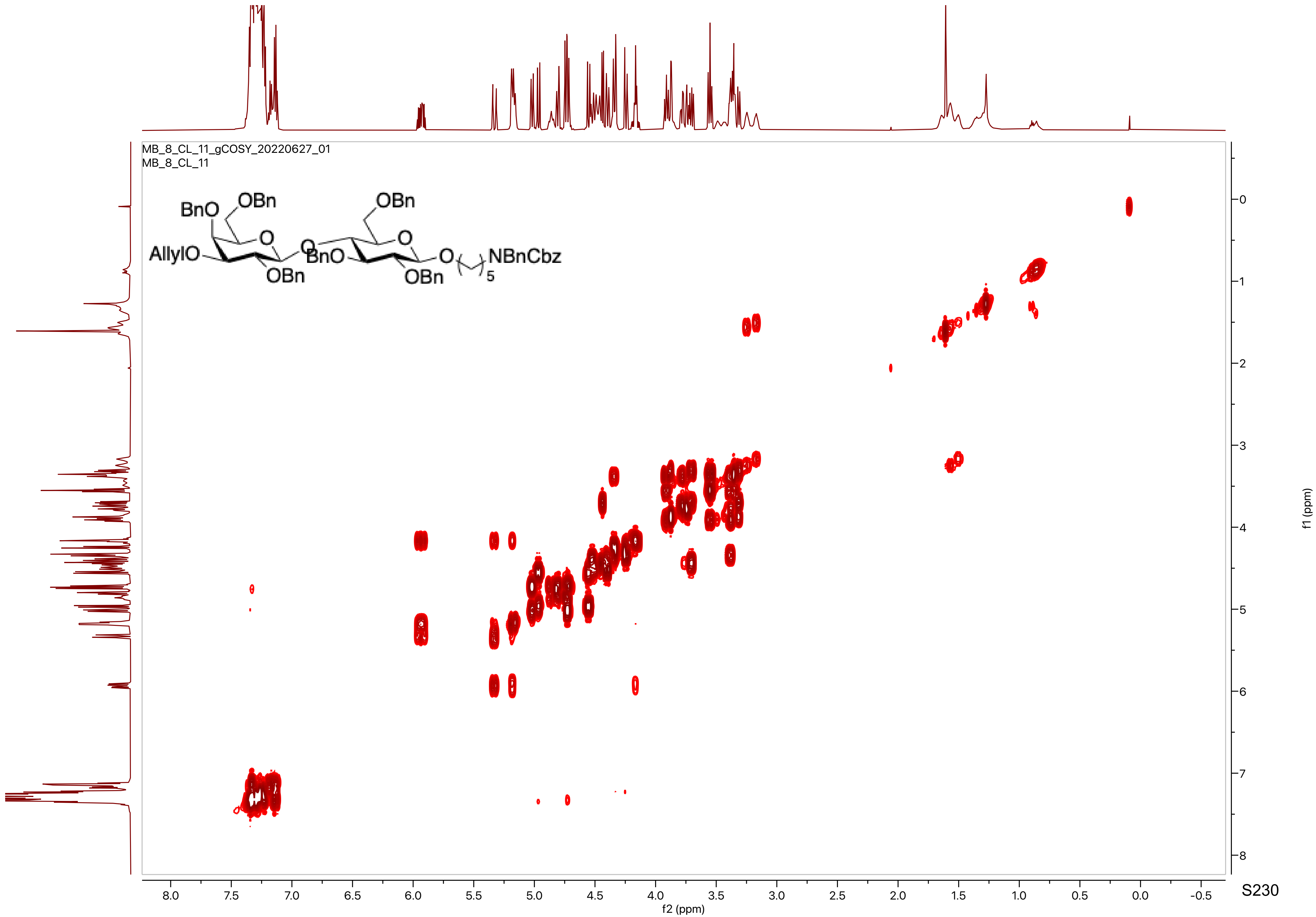
MB_8_CL_11_gCOSY_20220627_01
MB_8_CL_11
BnO
OBn
O
AllylO
OBn
O
BnO
OBn
O
O
OBn
NBnCbz
5
f2 (ppm)
f1 (ppm)

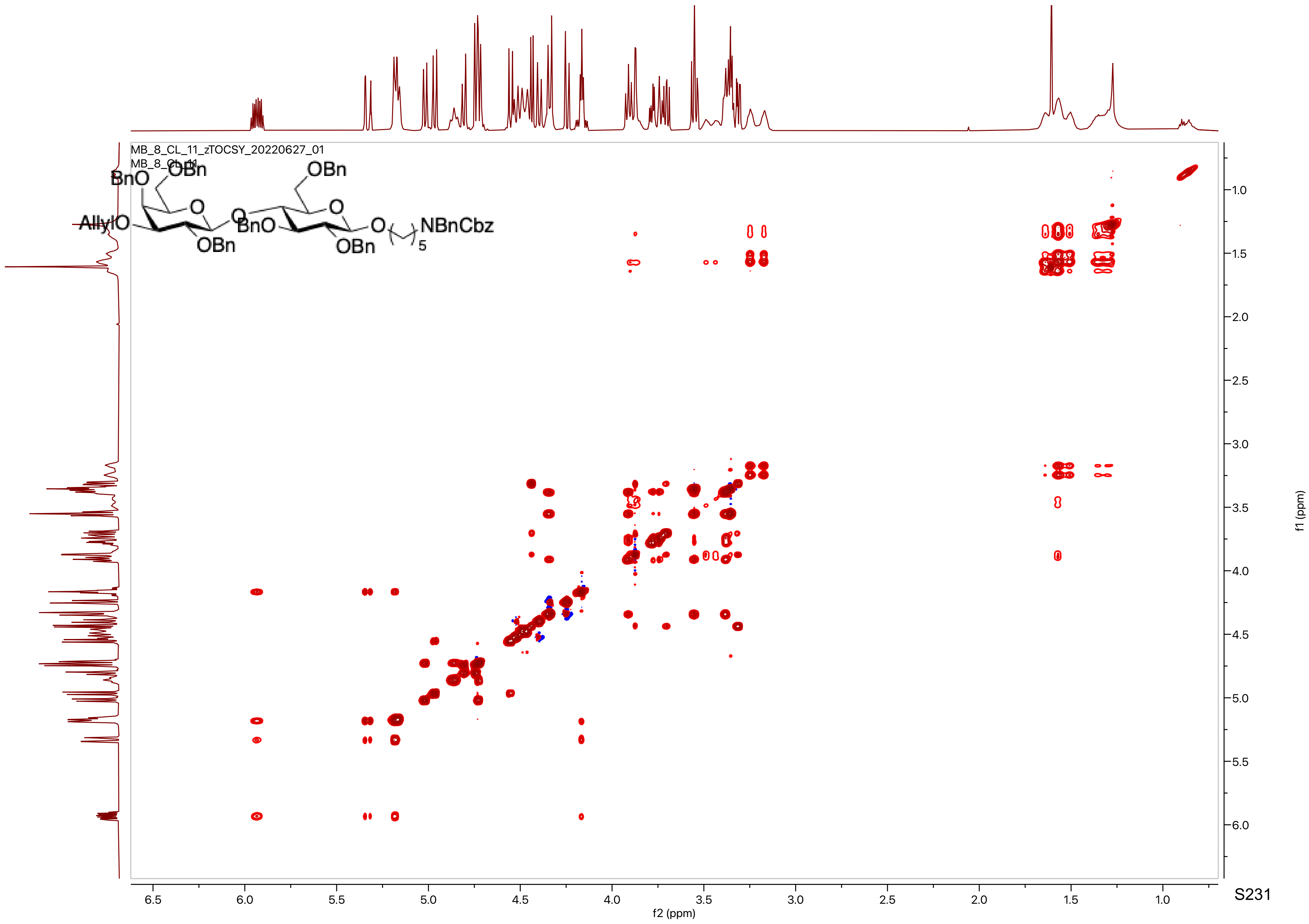
MB_8_CL_11_zTOCSY_20220627_01
MB_8_CL_11
f1 (ppm)
f2 (ppm)

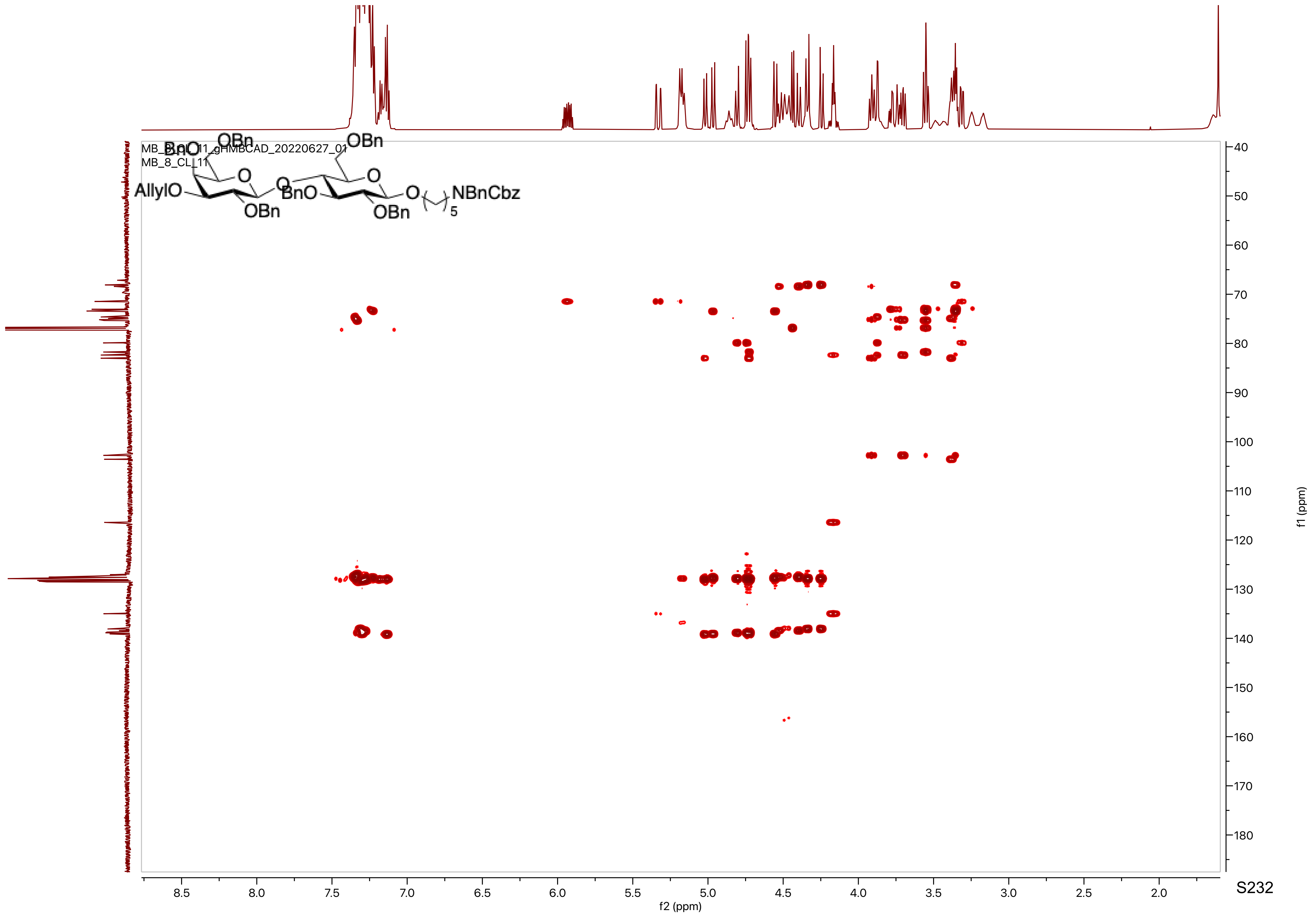

MB_8_CL_11_gHMBCAD_20220627_01
MB_8_CL_11
AllylO
BnO
OBn
O
BnO
OBn
O
O
OBn
OBn
O
NBnCbz
5
f2 (ppm)
f1 (ppm)

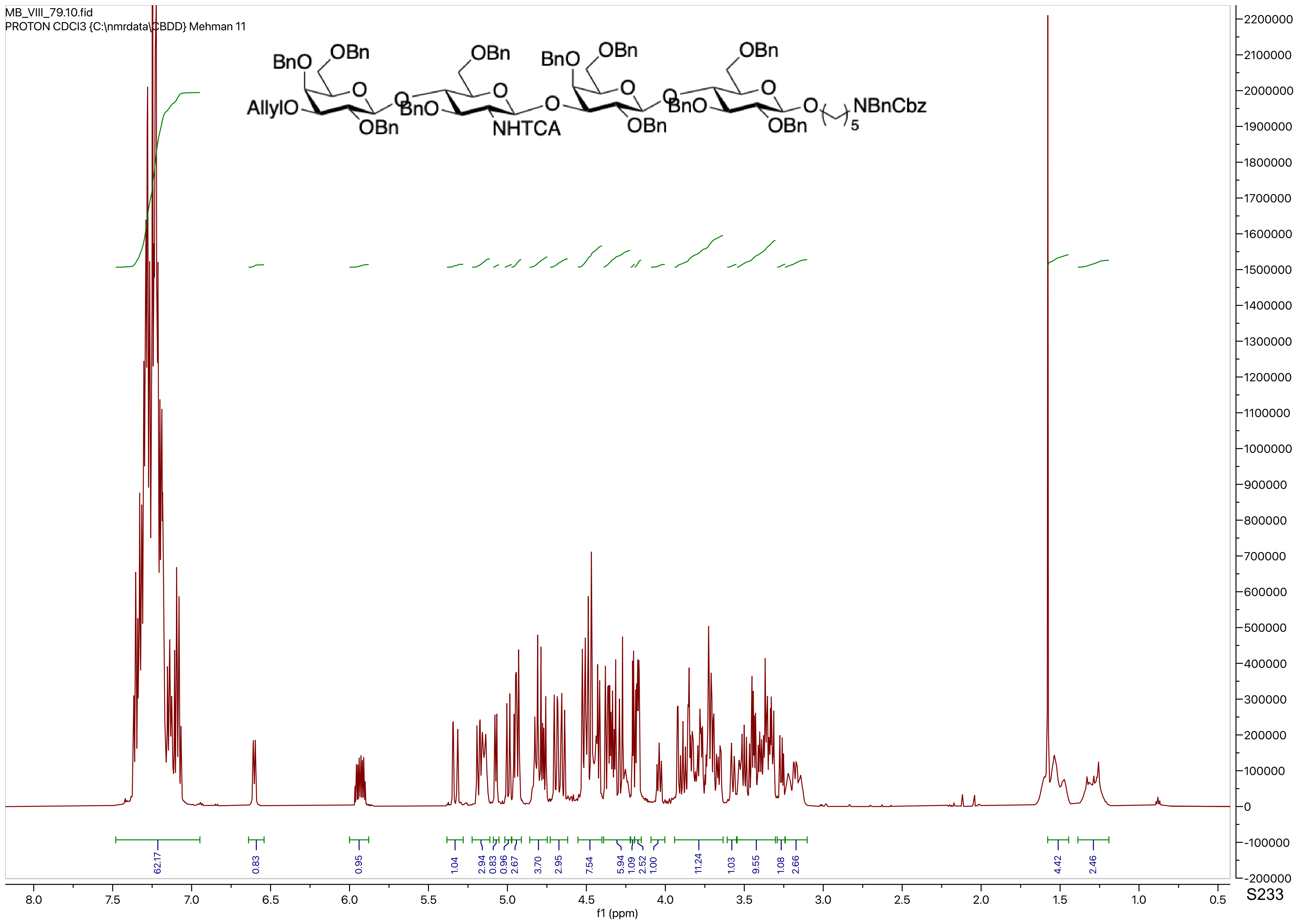

MB_VIII_79.10.fid
PROTON CDCl3 {C:\nmrdata\CBDD} Mehman 11
BnO
OBn
AllylO
OBn
BnO
NHTCA
OBn
BnO
OBn
OBn
BnO
OBn
NBnCbz
5
62.17
0.83
0.95
1.04
2.94
0.83
0.96
2.67
3.70
2.95
7.54
5.94
1.09
2.52
1.00
11.24
1.03
9.55
1.08
2.66
4.42
2.46
f1 (ppm)

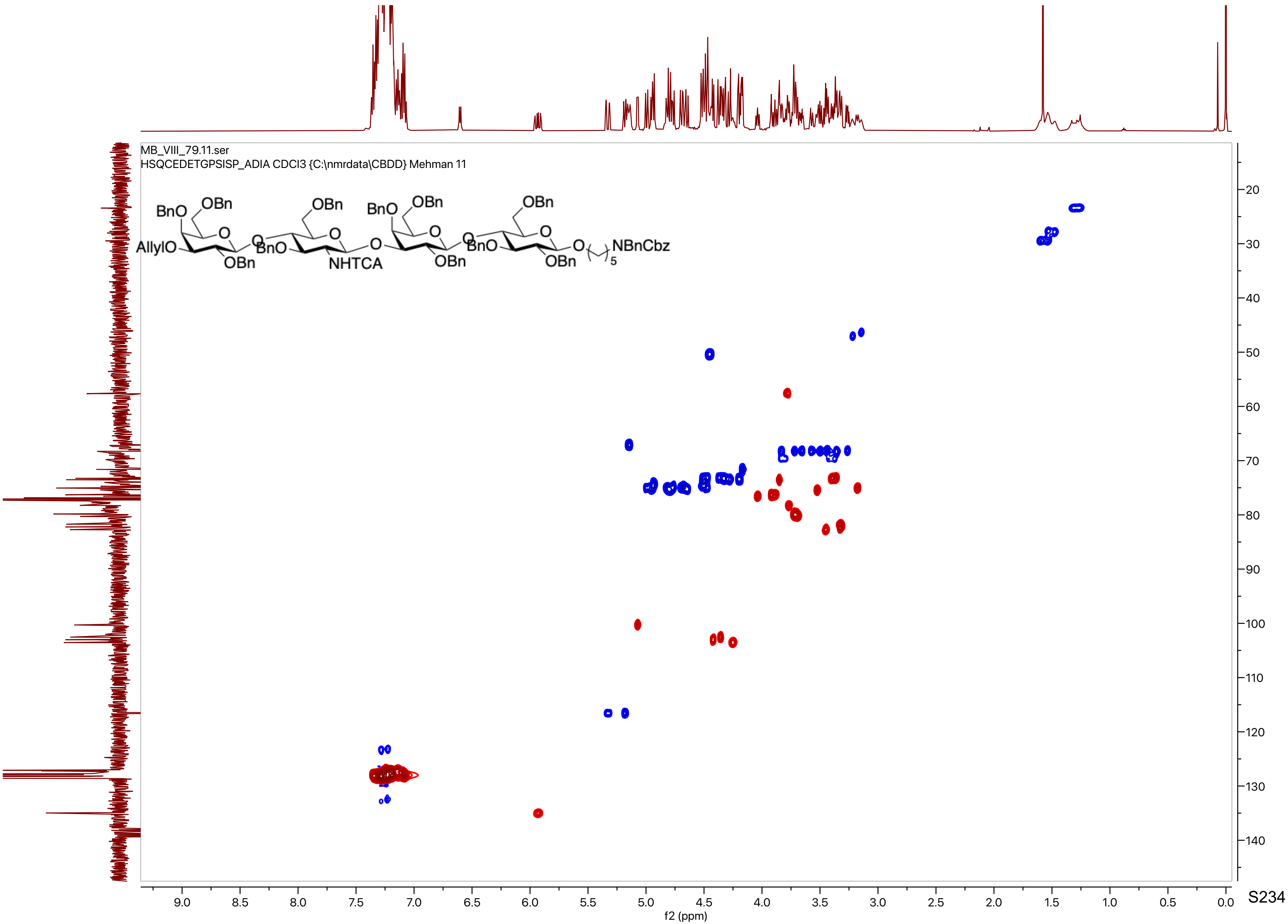

MB_VIII_79.11.ser
HSQCEDETGPSISP_ADIA CDCl3 {C:\nmrdata\CBDD} Mehman 11
BnO
OBn
AllylO
OBn
NHTCA
NBnCbz
f1 (ppm)
f2 (ppm)

MB_VIII_79.12.fid
C13APT CDCl3 {C:\nmrdata\CBDD} Mehman 11

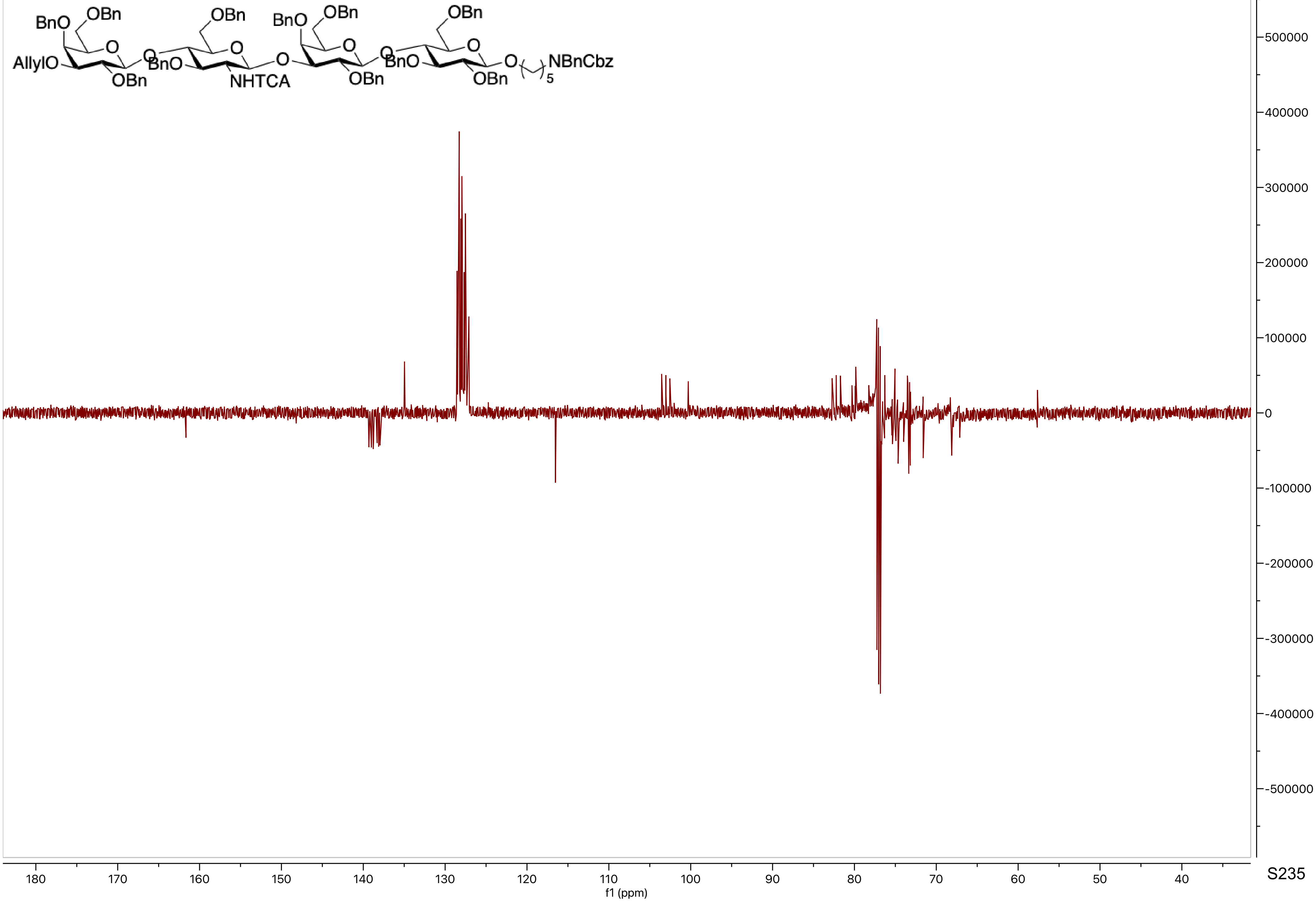

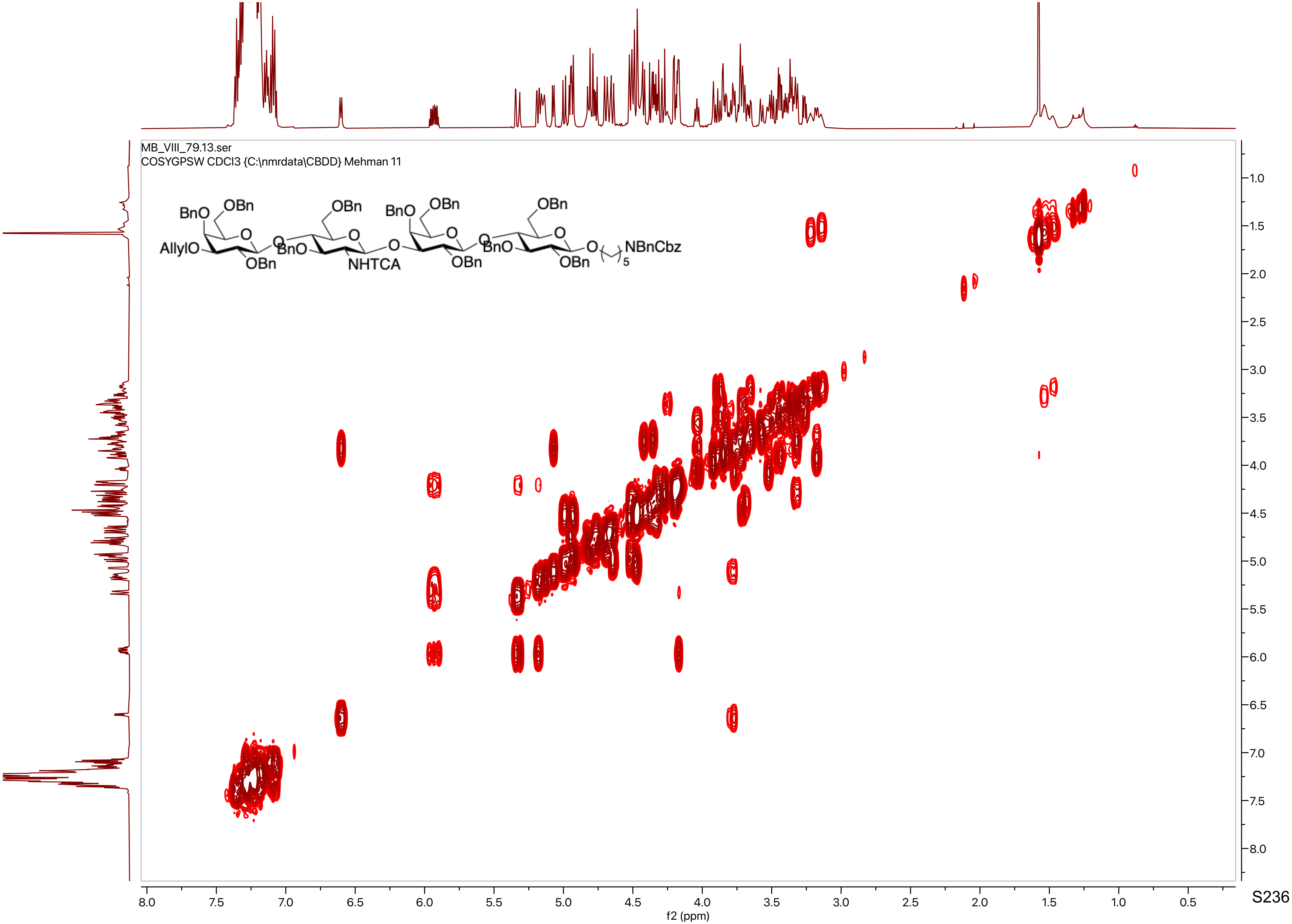
MB_VIII_79.13.ser
COSYGPSW CDCl3 {C:\nmrdata\CBDD} Mehman 11
AllylO
BnO
OBn
NHTCA
NBnCbz
f2 (ppm)
f1 (ppm)

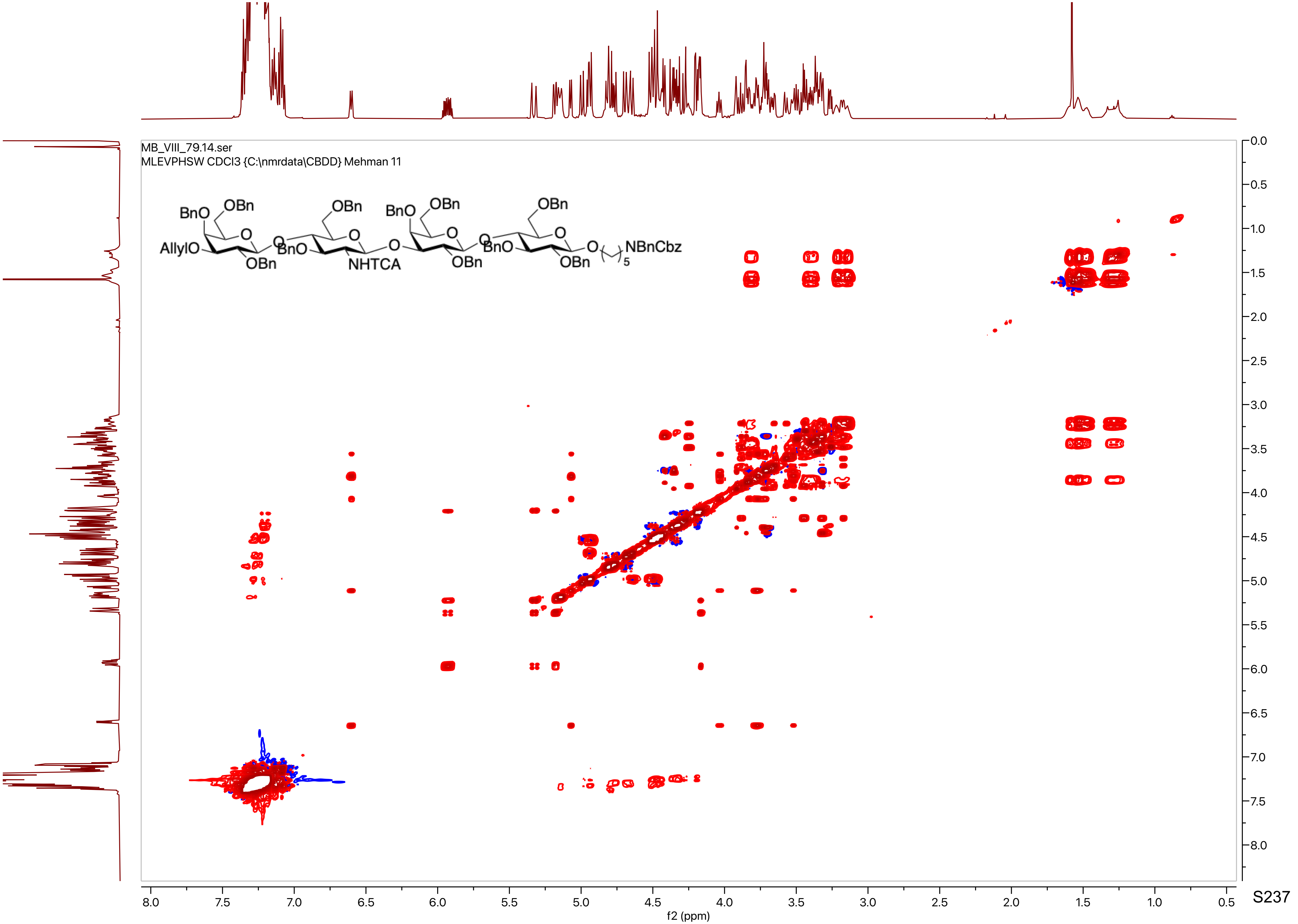

MB_VIII_79.14.ser
MLEVPHSW CDCl3 {C:\nmrdata\CBDD} Mehman 11
BnO
OBn
AllylO
OBn
NHTCA
NBnCbz
f2 (ppm)
f1 (ppm)

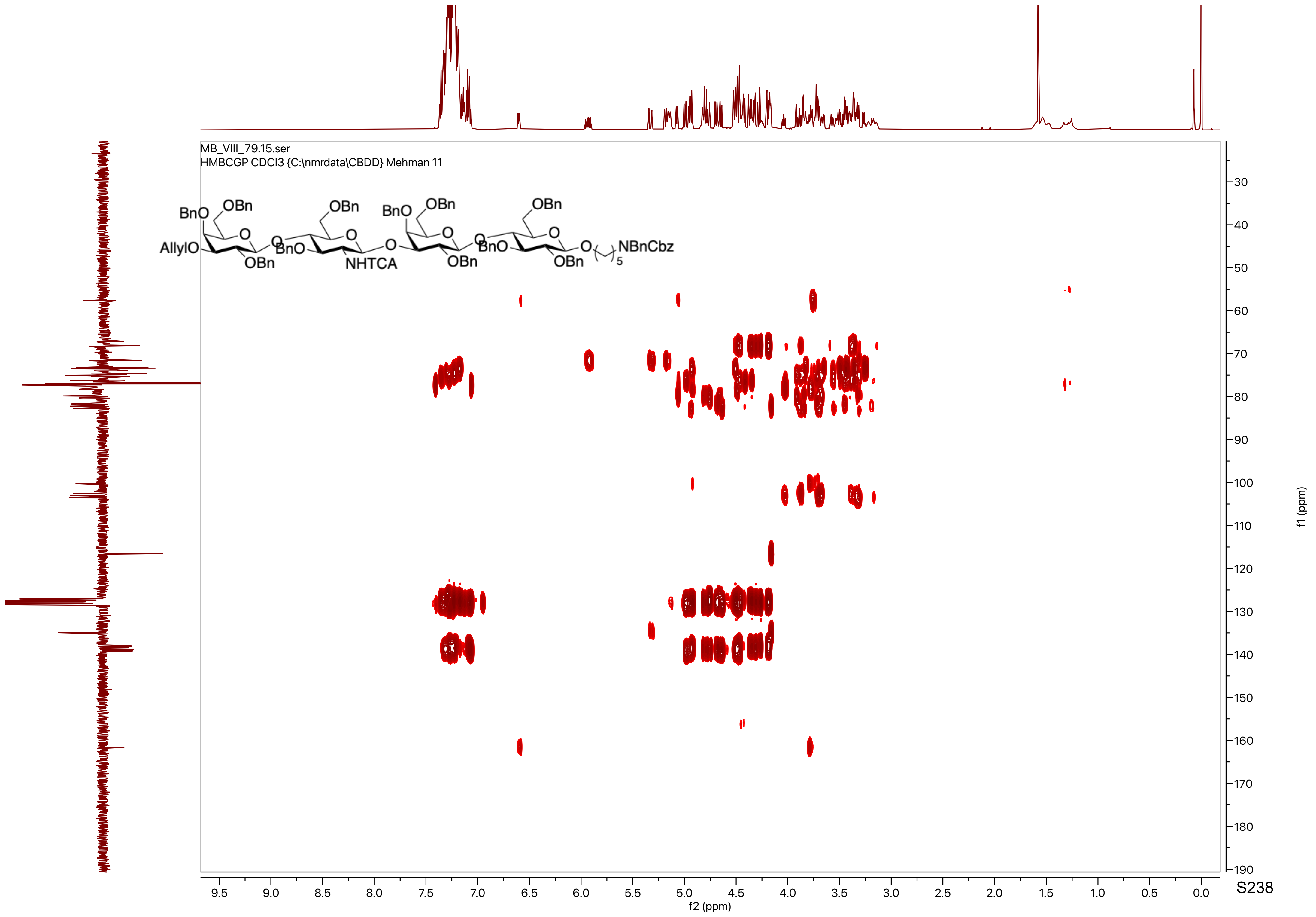
MB_VIII_79.15.ser
HMBCGP CDCl3 {C:\nmrdata\CBDD} Mehman 11
f2 (ppm)
f1 (ppm)

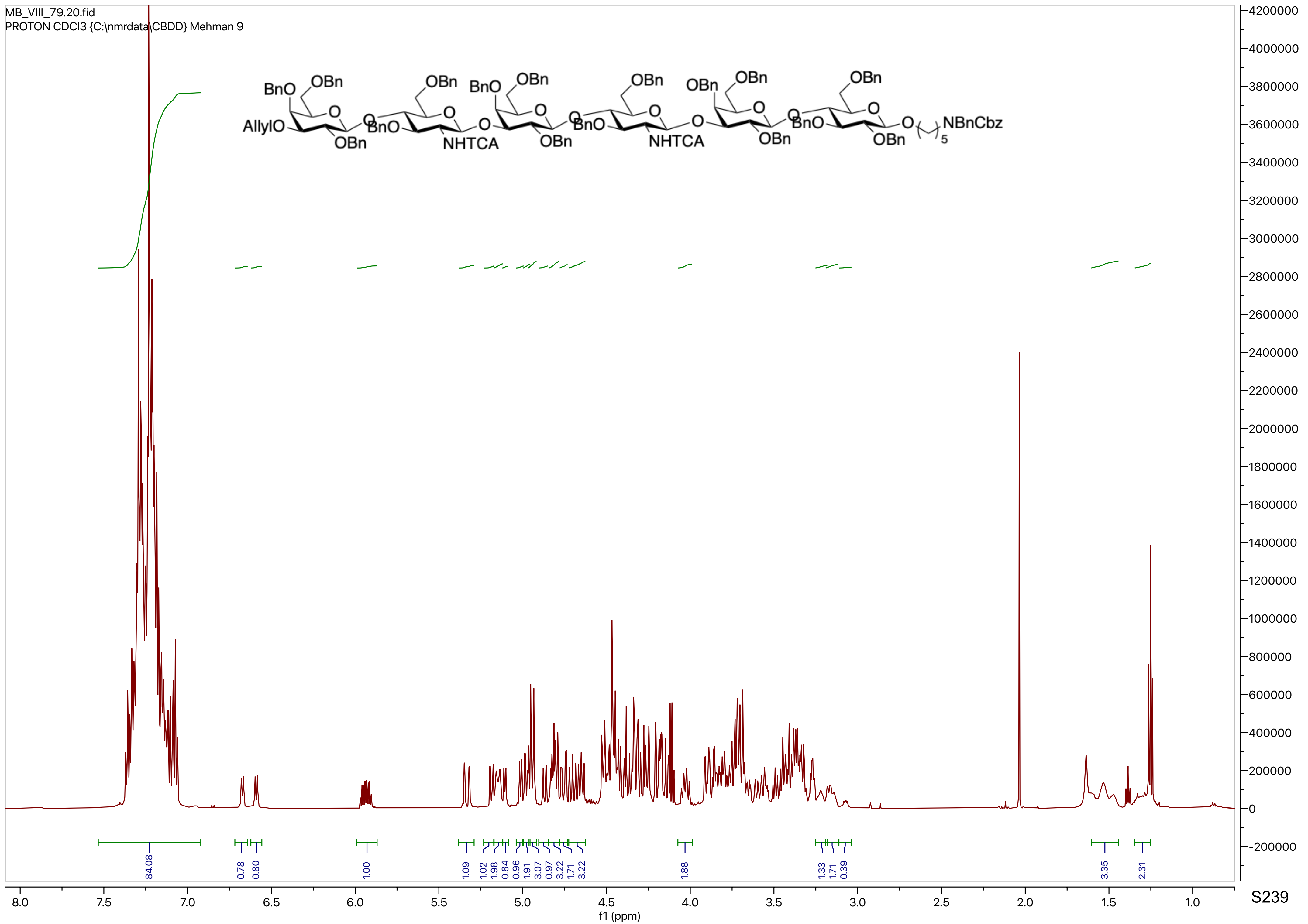

MB_VIII_79.20.fid
PROTON CDCl3 {C:\nmrdata\CBDD} Mehman 9
AllylO
BnO
OBn
NHTCA
NBnCbz
5
84.08
0.78
0.80
1.00
1.09
1.02
1.98
0.84
0.96
1.91
3.07
0.97
3.22
1.71
3.22
1.88
1.33
1.71
0.39
3.35
2.31
f1 (ppm)

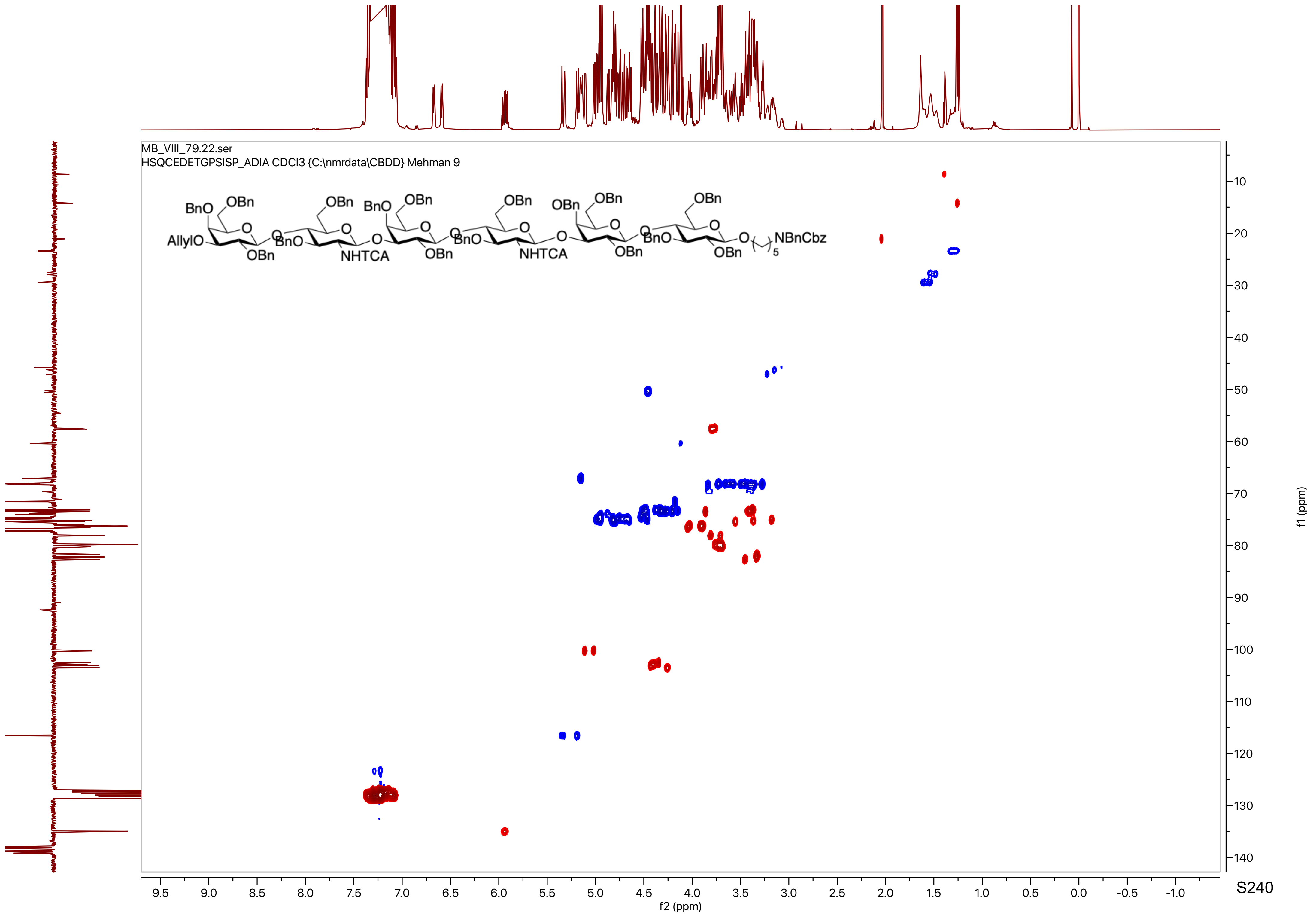
MB_VIII_79.22.ser
HSQCEDETGPSISP_ADIA CDCl3 {C:\nmrdata\CBDD} Mehman 9
AllylO
BnO
OBn
NHTCA
NBnCbz
f2 (ppm)
f1 (ppm)

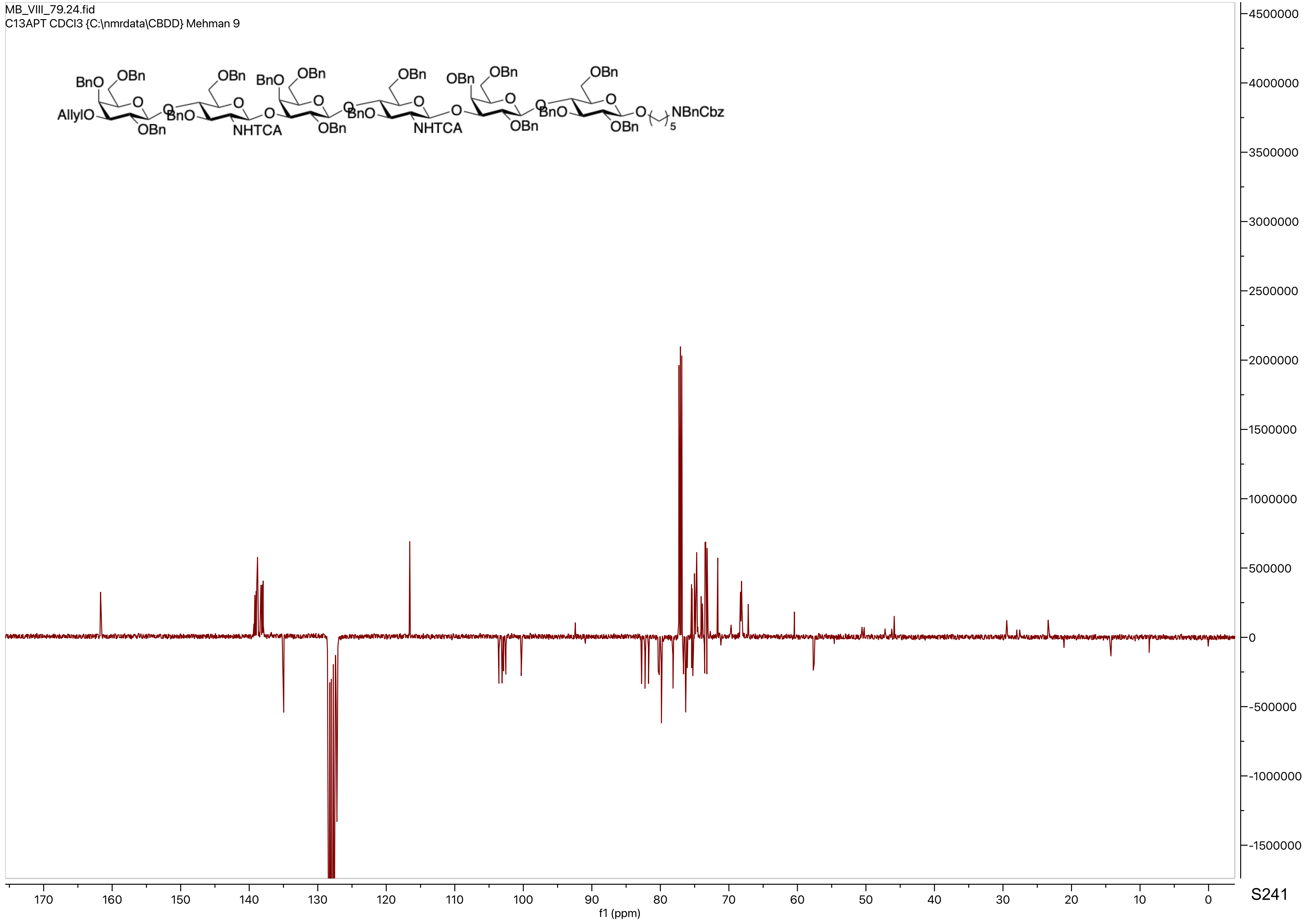
MB_VIII_79.24.fid
C13APT CDCl3 {C:\nmrdata\CBDD} Mehman 9
BnO
OBn
AllylO
OBn
NHTCA
OBn
NHTCA
OBn
OBn
NBnCbz
f1 (ppm)

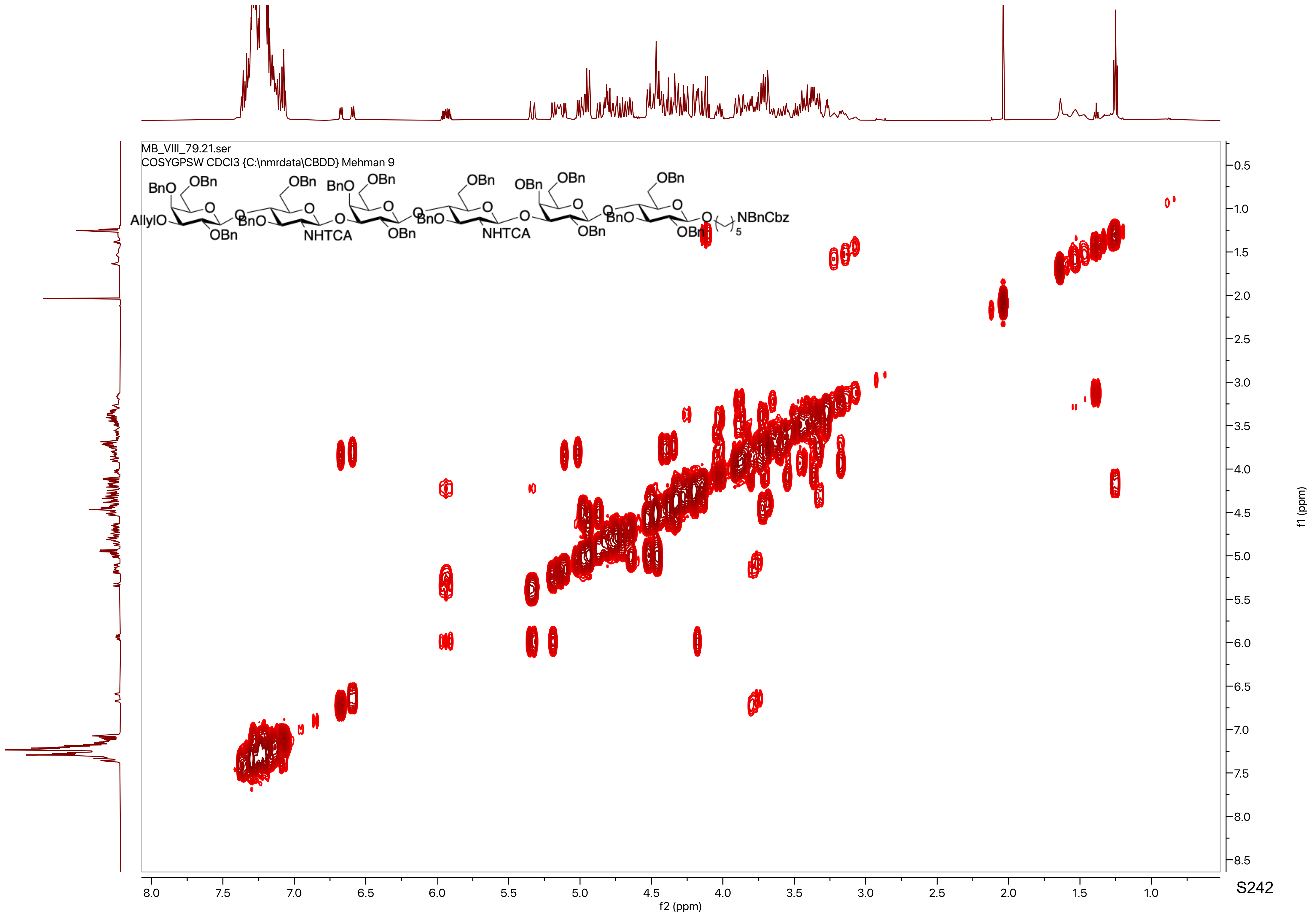
MB_VIII_79.21.ser
COSYGPSW CDCl3 {C:\nmrdata\CBDD} Mehman 9
BnO
OBn
AllylO
O
OBn
BnO
NHTCA
NBnCbz
5
f2 (ppm)
f1 (ppm)

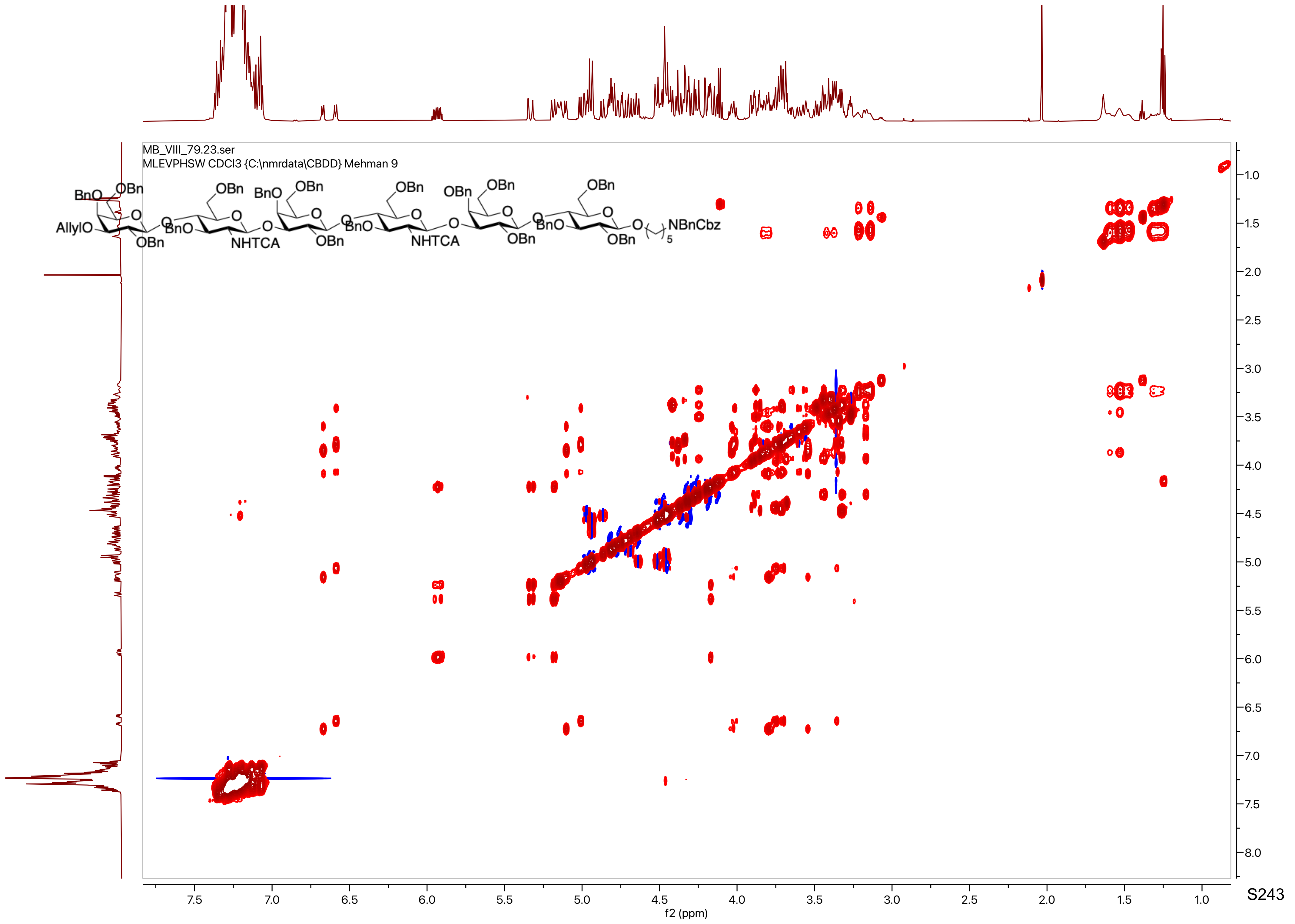
MB_VIII_79.23.ser
MLEVPHSW CDCl3 {C:\nmrdata\CBDD} Mehman 9
BnO
OBn
AllylO
NHTCA
NBnCbz
5
f2 (ppm)
f1 (ppm)

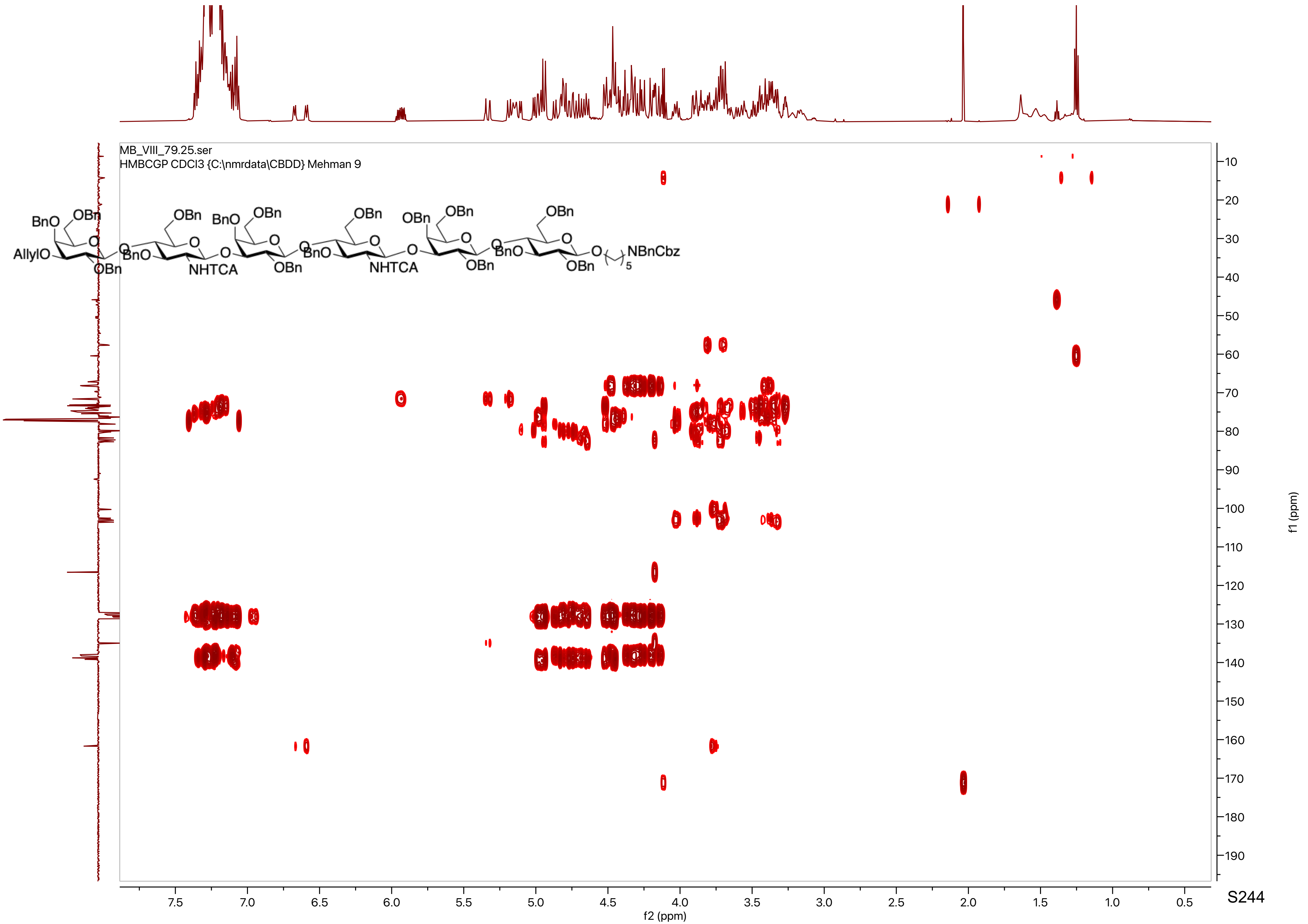
MB_VIII_79.25.ser
HMBCGP CDCl3 {C:\nmrdata\CBDD} Mehman 9
f2 (ppm)
f1 (ppm)

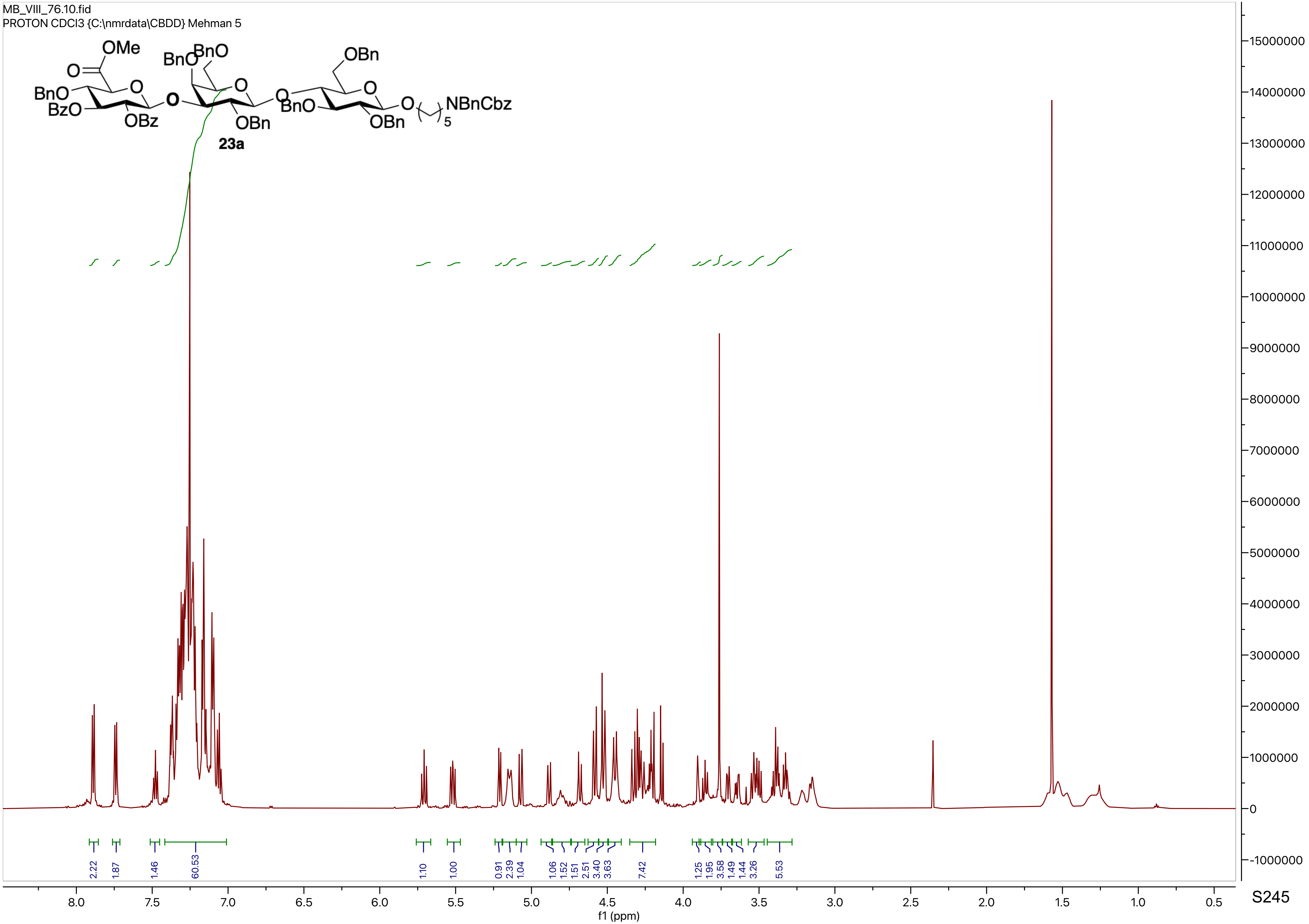

MB_VIII_76.10.fid
PROTON CDCl3 {C:\nmrdata\CBDD} Mehman 5
OMe
BnO
BzO
OBz
BnO
BnO
OBn
OBn
BnO
OBn
NBnCbz
23a
2.22
1.87
1.46
60.53
1.10
1.00
0.91
2.39
1.04
1.06
1.52
1.51
2.51
3.40
3.63
7.42
1.25
1.95
3.58
1.49
1.44
3.26
5.53
f1 (ppm)

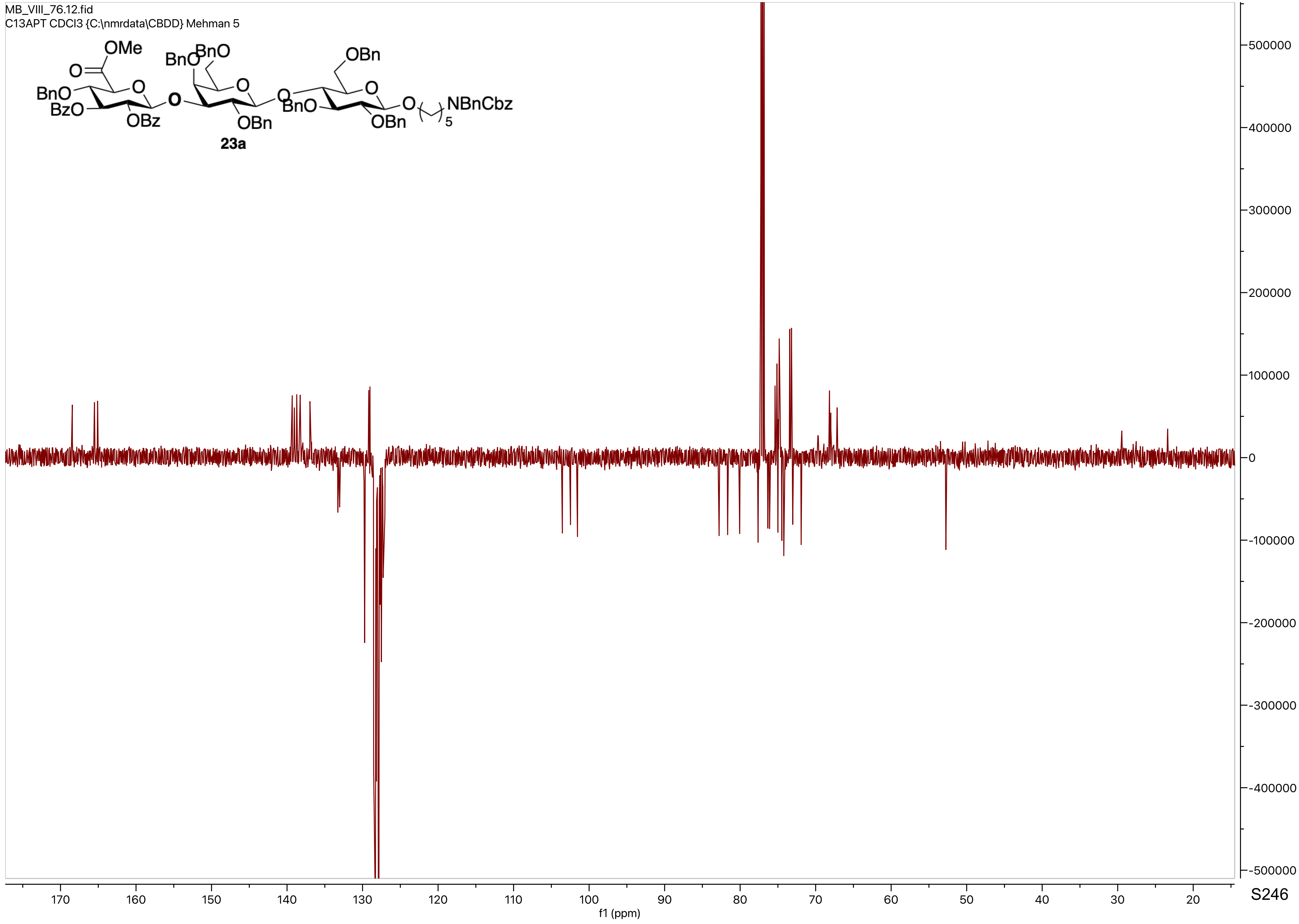
MB_VIII_76.12.fid
C13APT CDCl3 {C:\nmrdata\CBDD} Mehman 5
OMe
O
BnO
BzO
OBz
BnO
BnO
O
O
OBn
OBn
O
BnO
O
OBn
O
5
NBnCbz
23a
500000
400000
300000
200000
100000
0
-100000
-200000
-300000
-400000
-500000
170
160
150
140
130
120
110
100
90
80
70
60
50
40
30
20
f1 (ppm)

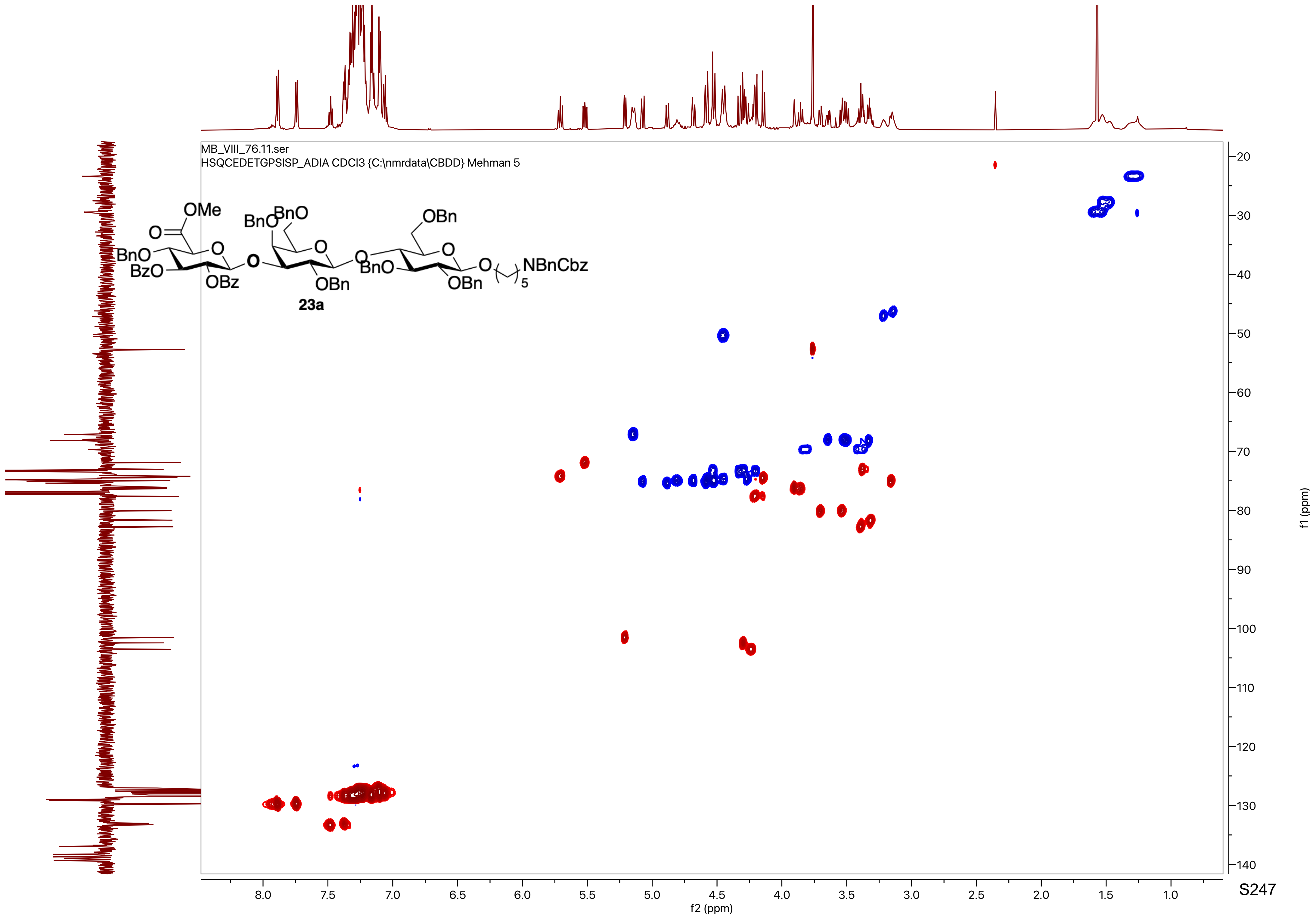
MB_VIII_76.11.ser
HSQCEDETGPSISP_ADIA CDCl3 {C:\nmrdata\CBDD} Mehman 5
OMe
BnO
BzO
OBz
BnO
BnO
OBn
OBn
BnO
OBn
NBnCbz
5
23a
f2 (ppm)
f1 (ppm)

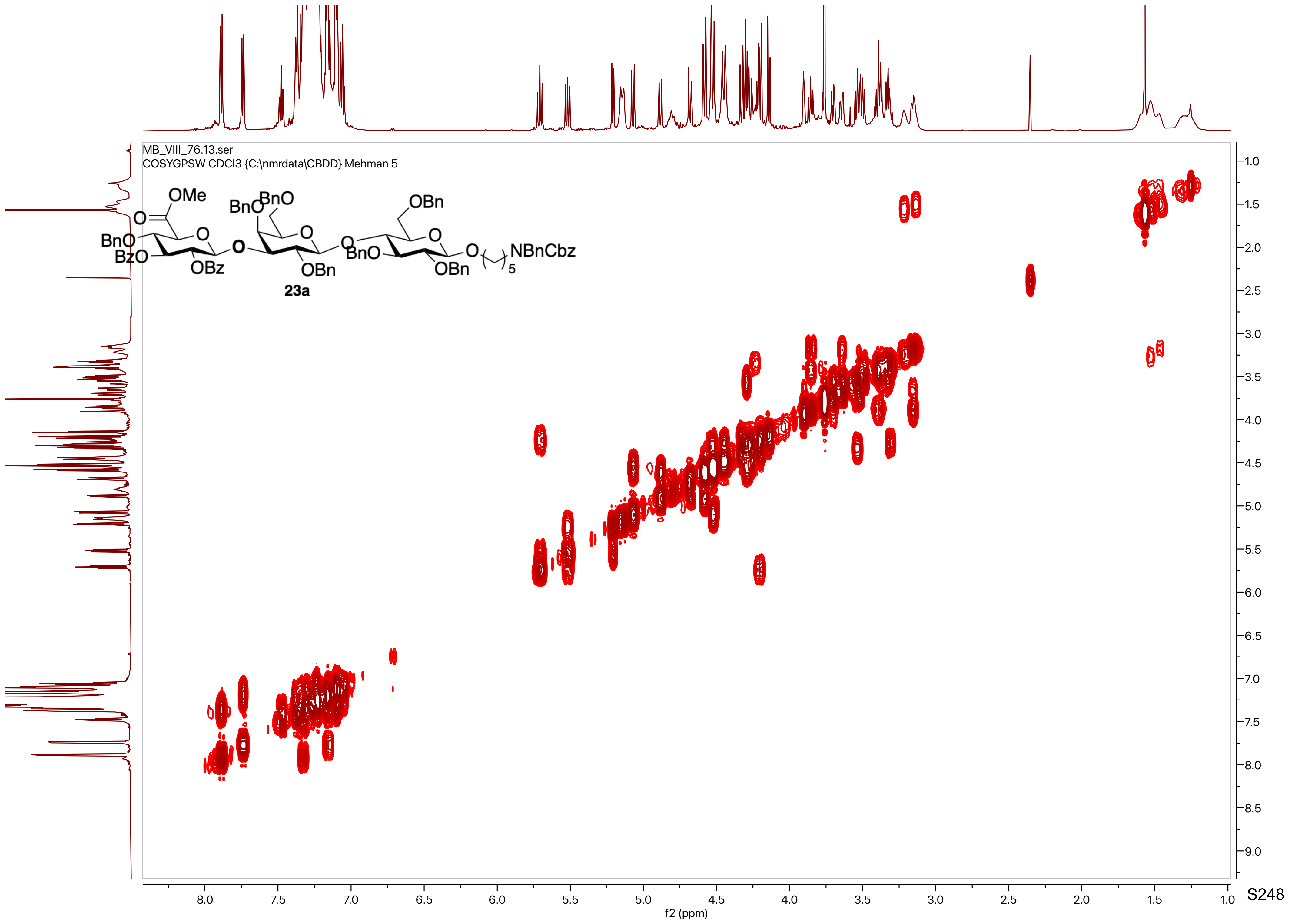
MB_VIII_76.13.ser
COSYGPSW CDCl3 {C:\nmrdata\CBDD} Mehman 5
OMe
BnO
BnO
OBn
BnO
BzO
OBz
OBn
BnO
OBn
NBnCbz
5
23a
f2 (ppm)
f1 (ppm)

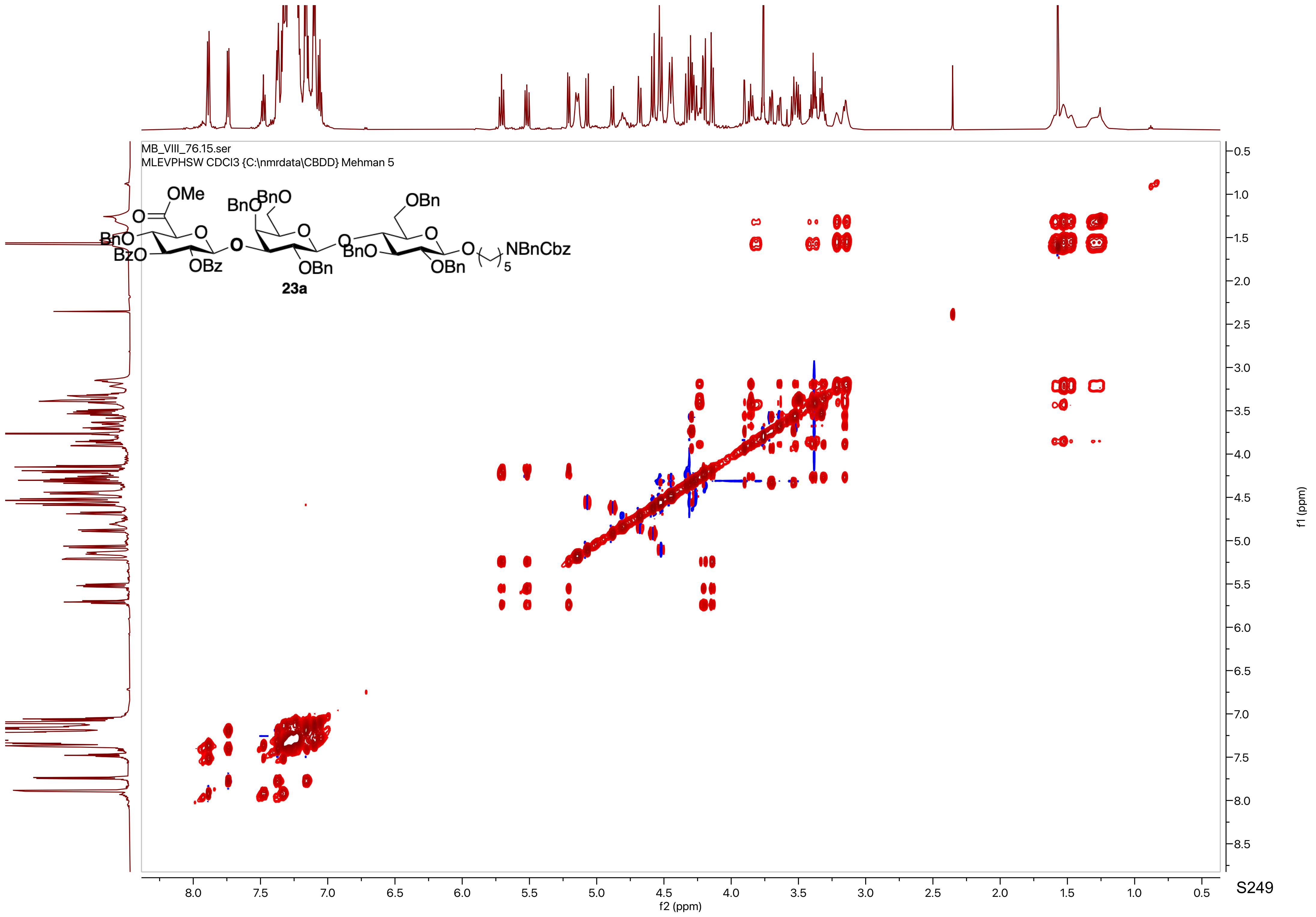
MB_VIII_76.15.ser
MLEVPHSW CDCl3 {C:\nmrdata\CBDD} Mehman 5
OMe
BnO
BnO
OBn
O
BnO
BzO
OBz
O
OBn
BnO
O
OBn
NBnCbz
5
23a
f2 (ppm)
f1 (ppm)

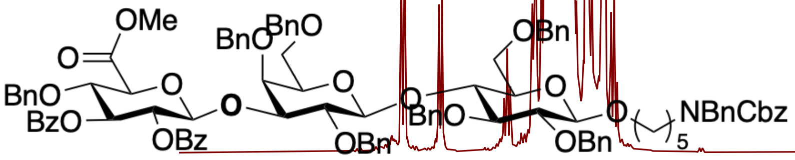
OMe
O
BnO
BzO
O
OBz
O
BnO
BnO
O
OBn
O
BnO
OBn
O
OBn
O
NBnCbz
5

23a

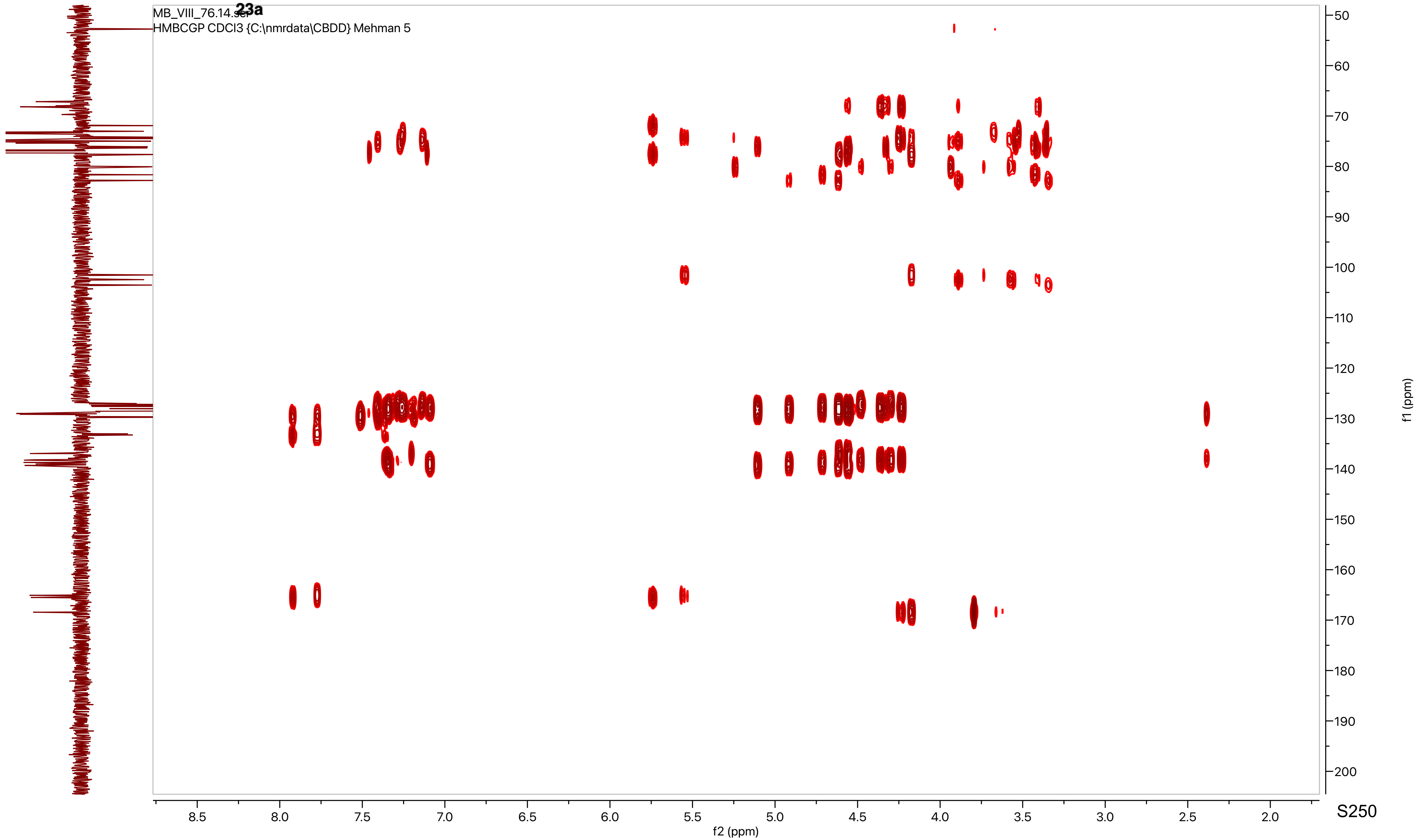
MB_VIII_76.14.ser
HMBCGP CDCl3 {C:\nmrdata\CBDD} Mehman 5
f2 (ppm)
f1 (ppm)

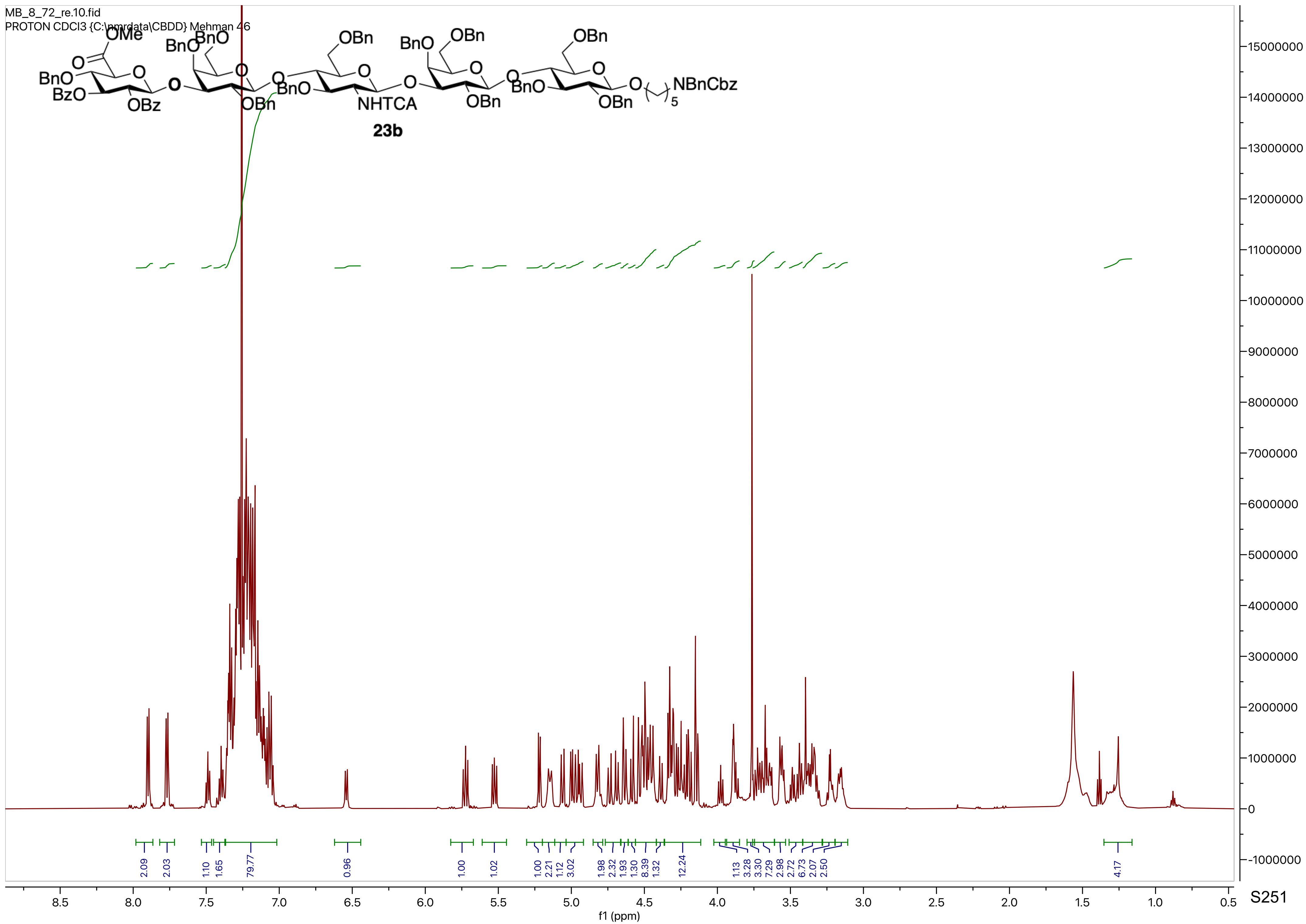

MB_8_72_re.10.fid
PROTON CDCl3 {C:\nmrdata\CBDD} Mehman 46
OMe
BnO
BzO
OBz
BnO
BnO
OBn
OBn
BnO
NHTCA
BnO
OBn
OBn
BnO
OBn
OBn
NBnCbz
5
23b
2.09
2.03
1.10
1.65
79.77
0.96
1.00
1.02
1.00
2.21
1.12
3.02
1.98
2.32
1.93
1.30
8.39
1.32
12.24
1.13
3.28
3.30
7.29
2.98
2.72
6.73
2.07
2.50
4.17
f1 (ppm)

MB_VIII_72.12.fid
C13APT CDCl3 {C:\nmrdata\CBDD} Mehman 28

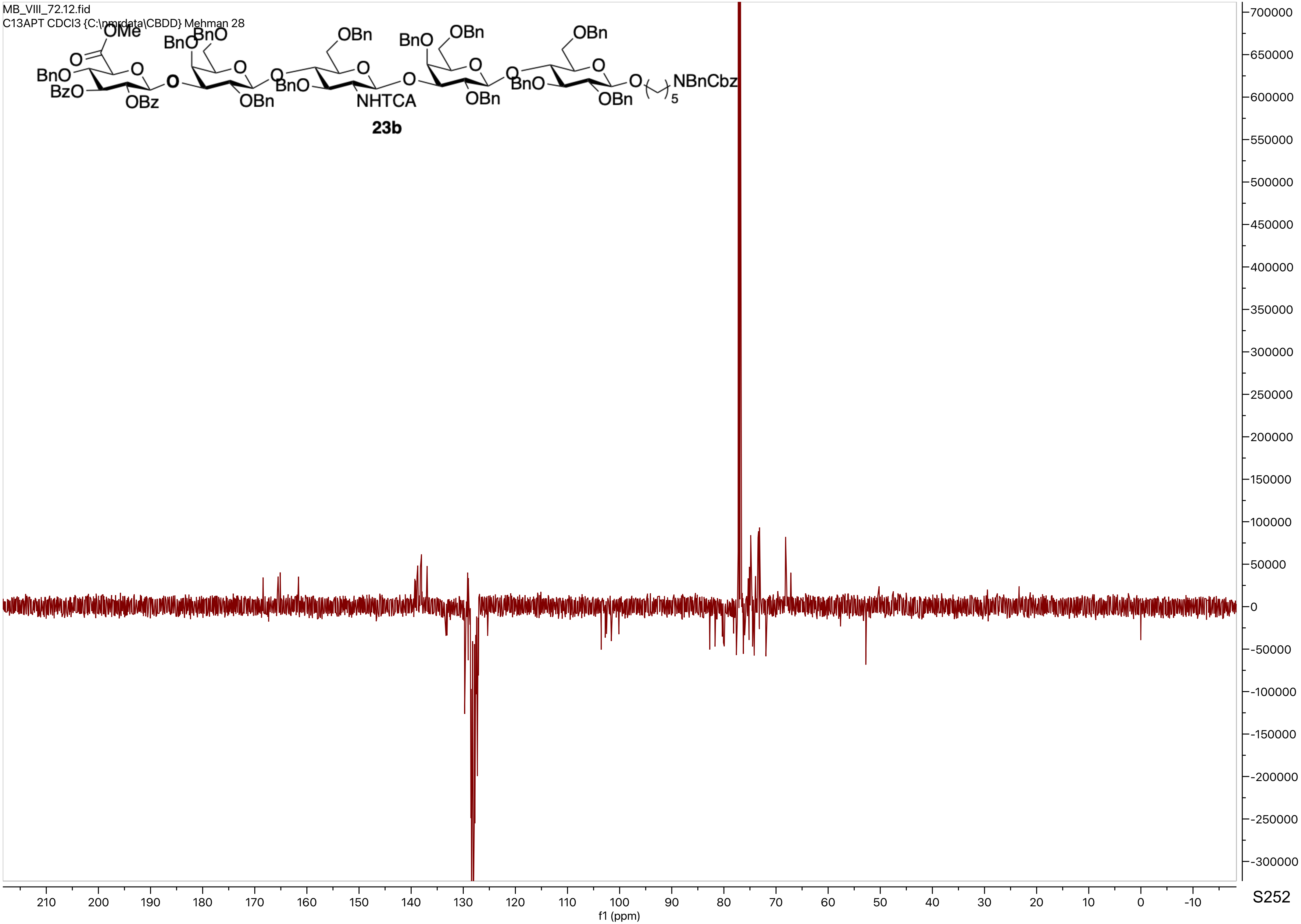


S252

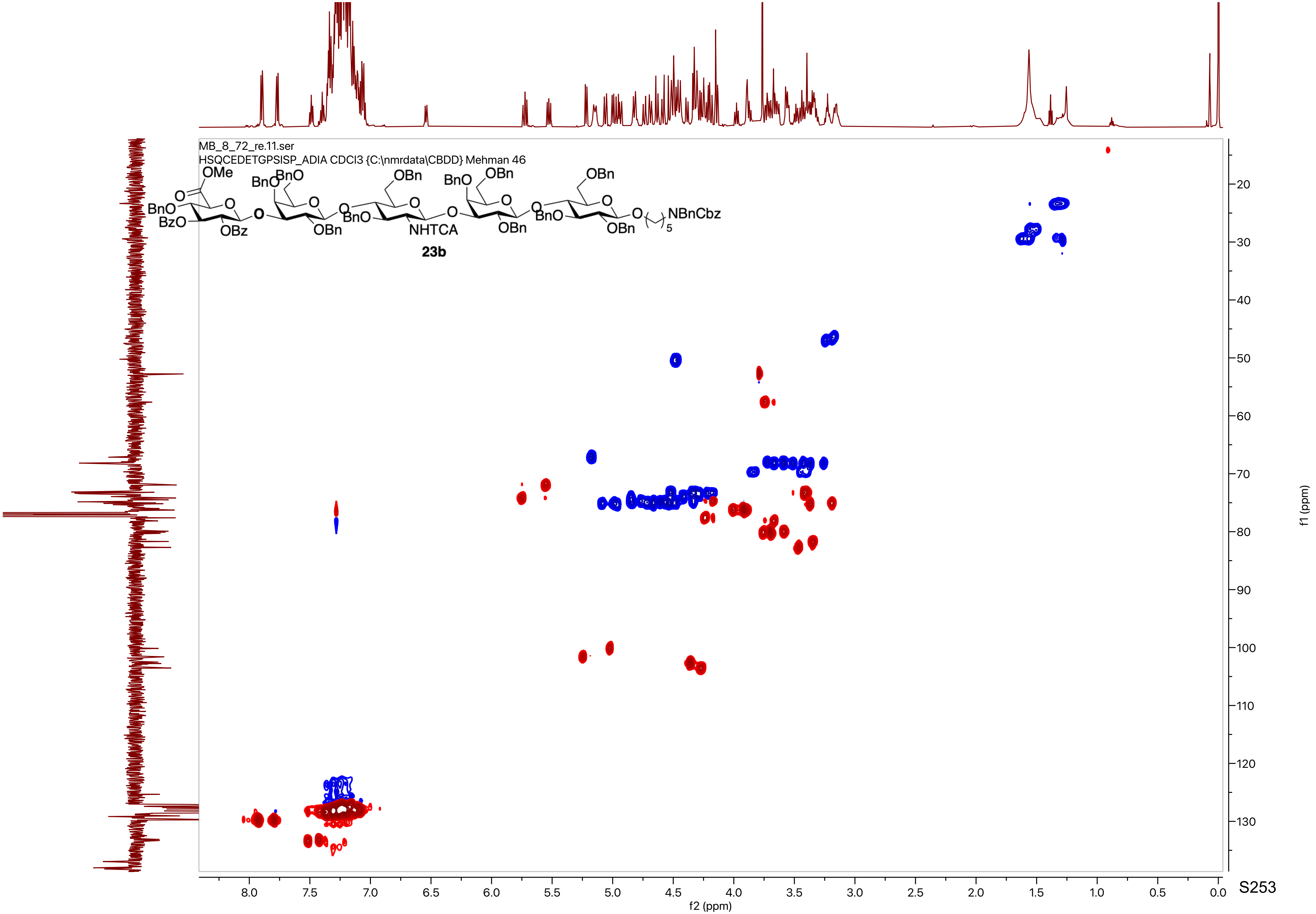

MB_8_72_re.11.ser
HSQCEDETGPSISP_ADIA CDCl3 {C:\nmrdata\CBDD} Mehman 46
OMe
BnO
BzO
OBz
BnO
BnO
OBn
OBn
NHTCA
OBn
BnO
OBn
OBn
NBnCbz
5
23b
f2 (ppm)
f1 (ppm)

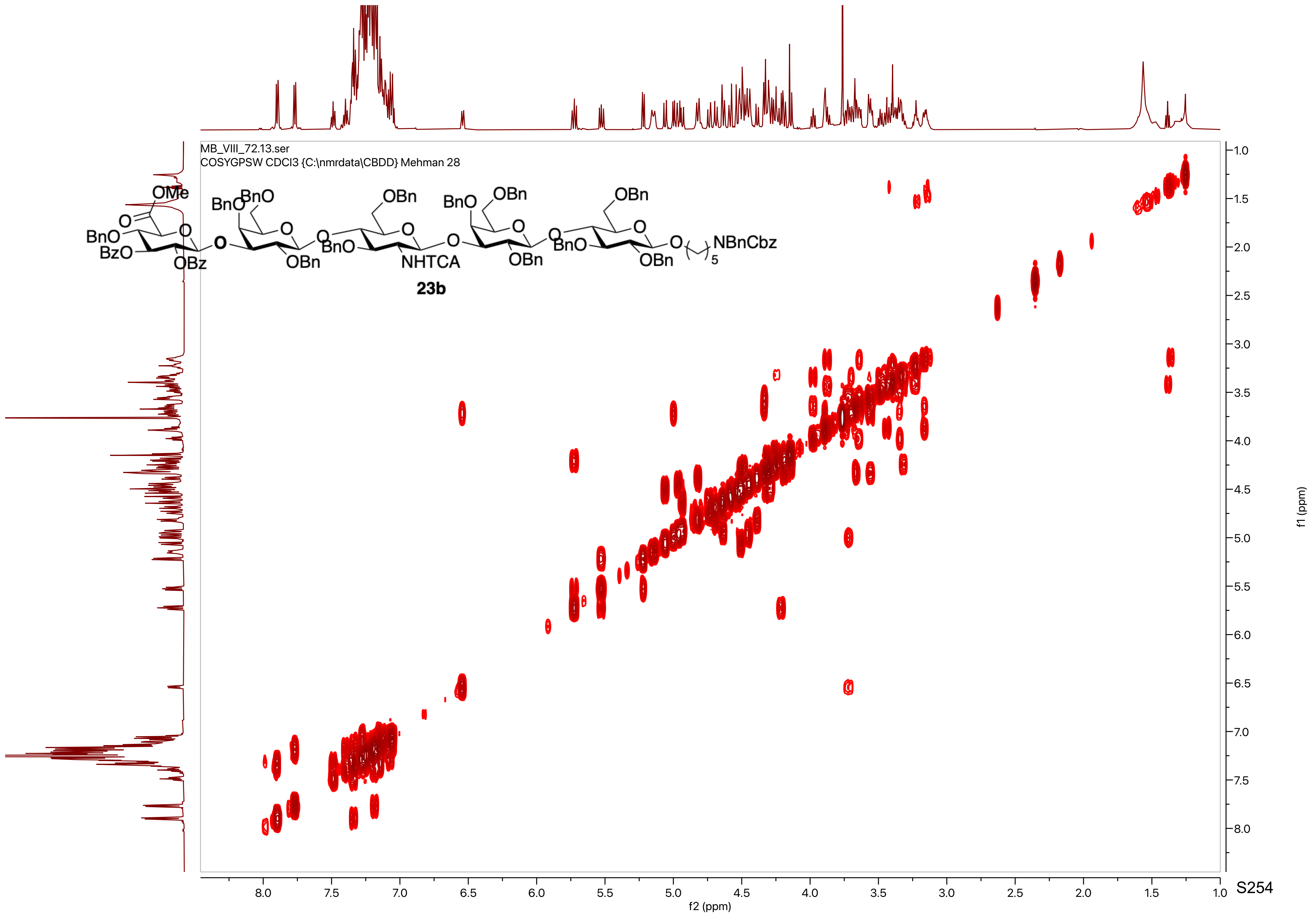
MB_VIII_72.13.ser
COSYGPSW CDCl3 {C:\nmrdata\CBDD} Mehman 28
OMe
BnO
BnO
BzO
OBz
OBn
OBn
NHTCA
NBnCbz
23b
f2 (ppm)
f1 (ppm)

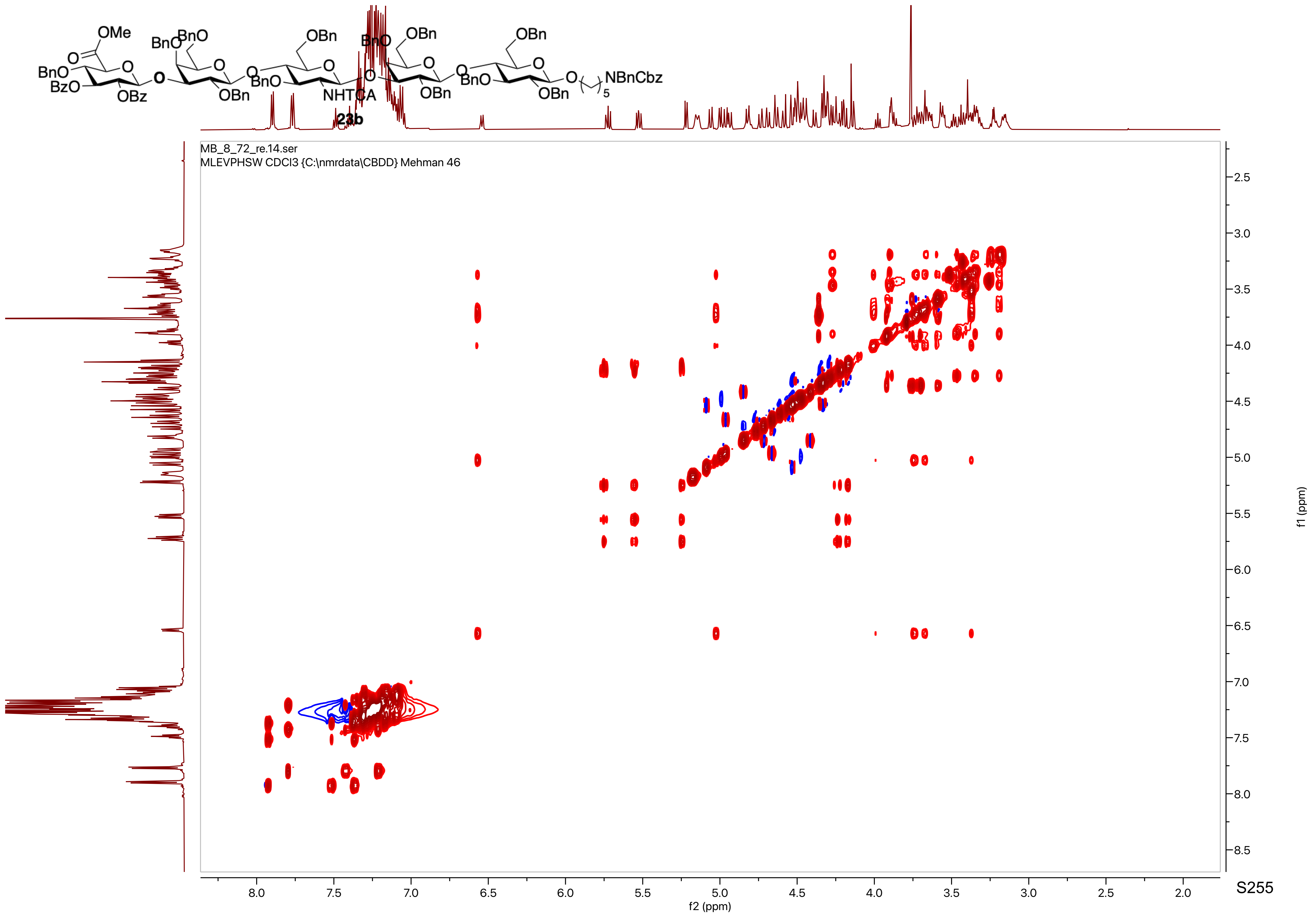

OMe
BnO
BzO
OBz
BnO
BnO
OBn
OBn
BnO
NHTCA
OBn
BnO
OBn
OBn
BnO
OBn
NBnCbz
5
23b
MB_8_72_re.14.ser
MLEVPHSW CDCl3 {C:\nmrdata\CBDD} Mehman 46
f2 (ppm)
f1 (ppm)

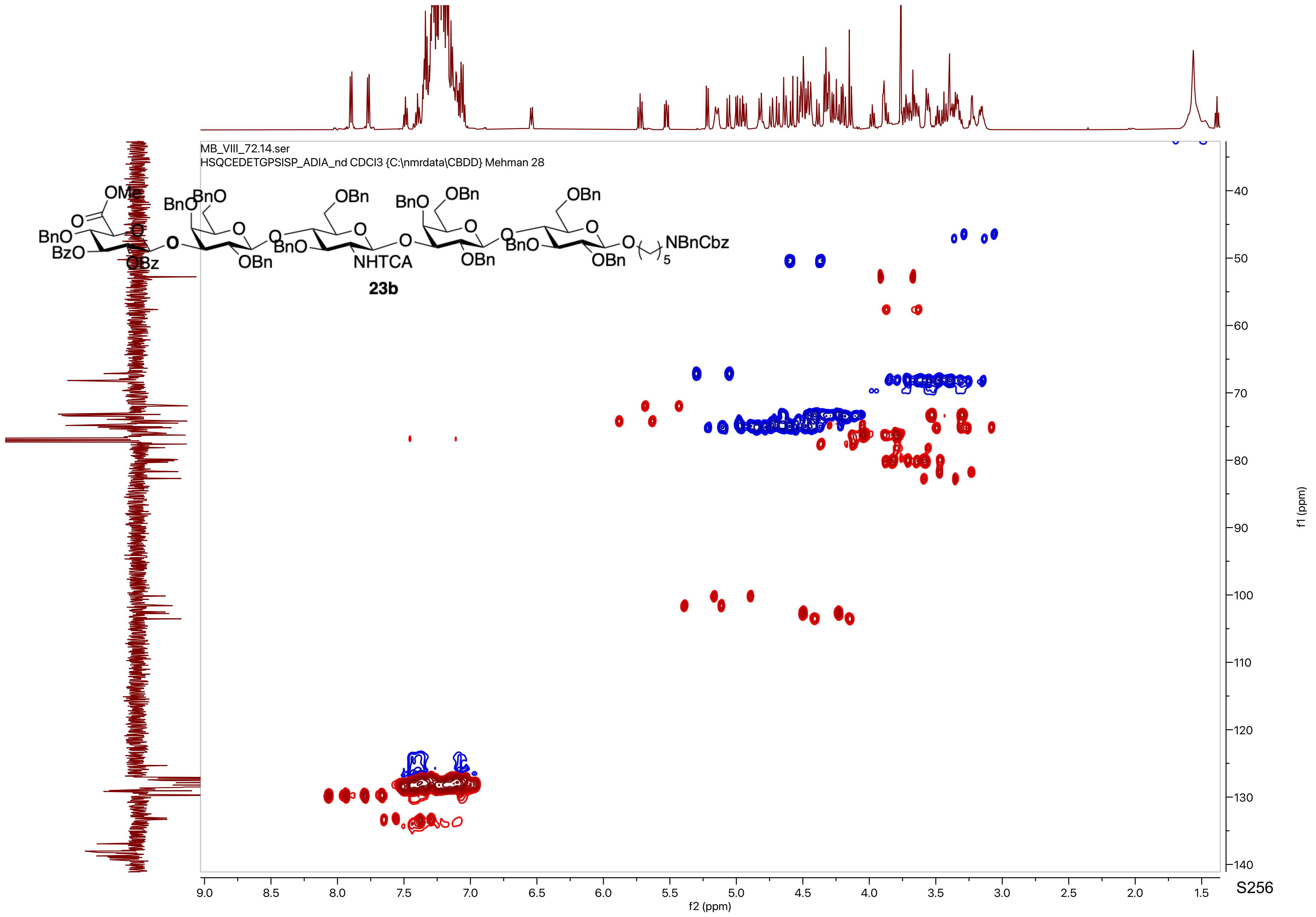
MB_VIII_72.14.ser
HSQCEDETGPSISP_ADIA_nd CDCl3 {C:\nmrdata\CBDD} Mehman 28
23b
f2 (ppm)
f1 (ppm)

MB_VIII_73_GLPG_hepta.10.fid
PROTON CDCl3 {C:\nmrdata\CBDD} Mehman 29

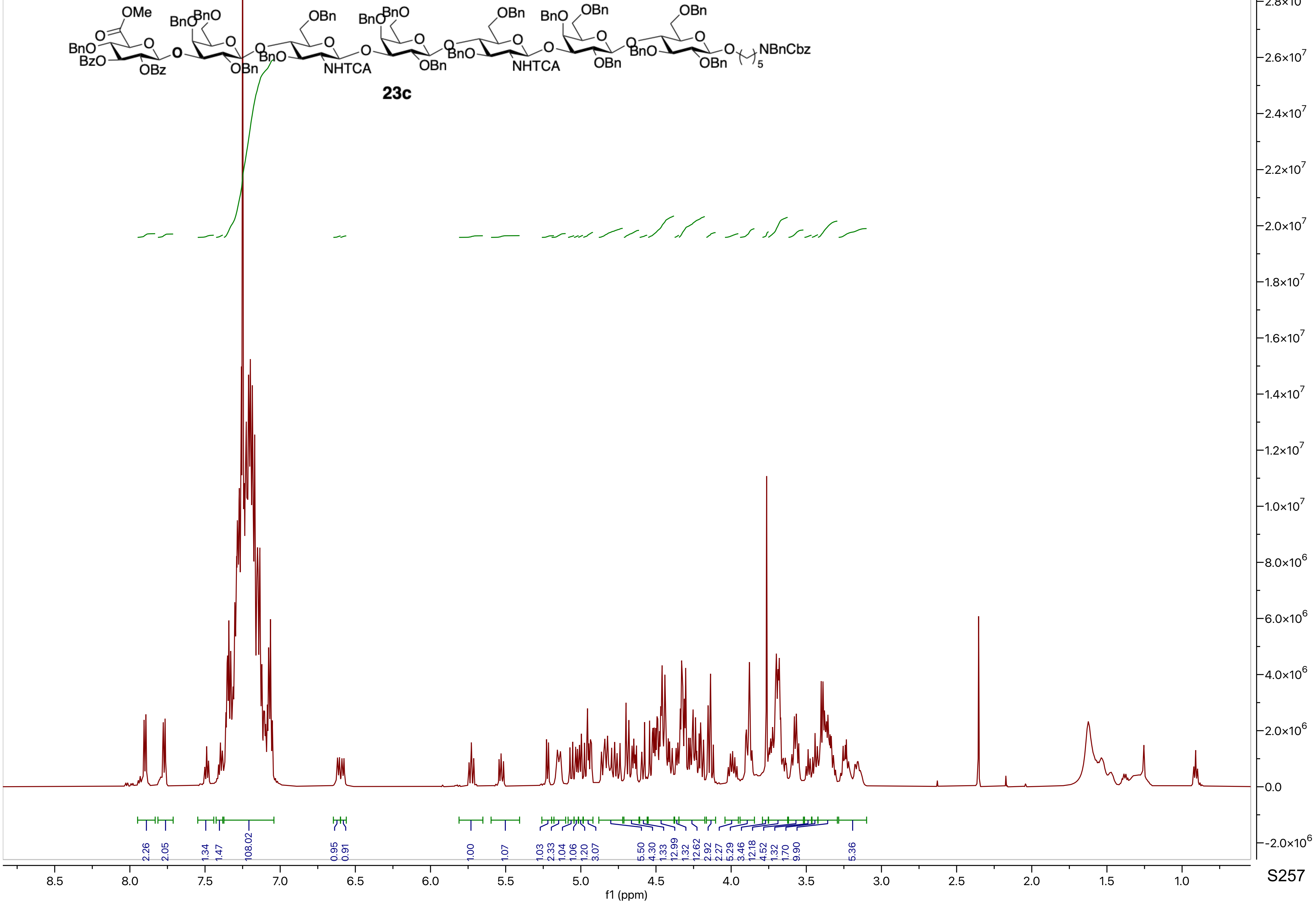

MB_VIII_73_GLGP_hepta.32.fid
C13APT CDCl3 {C:\nmrdata\CBDD} Mehman 24

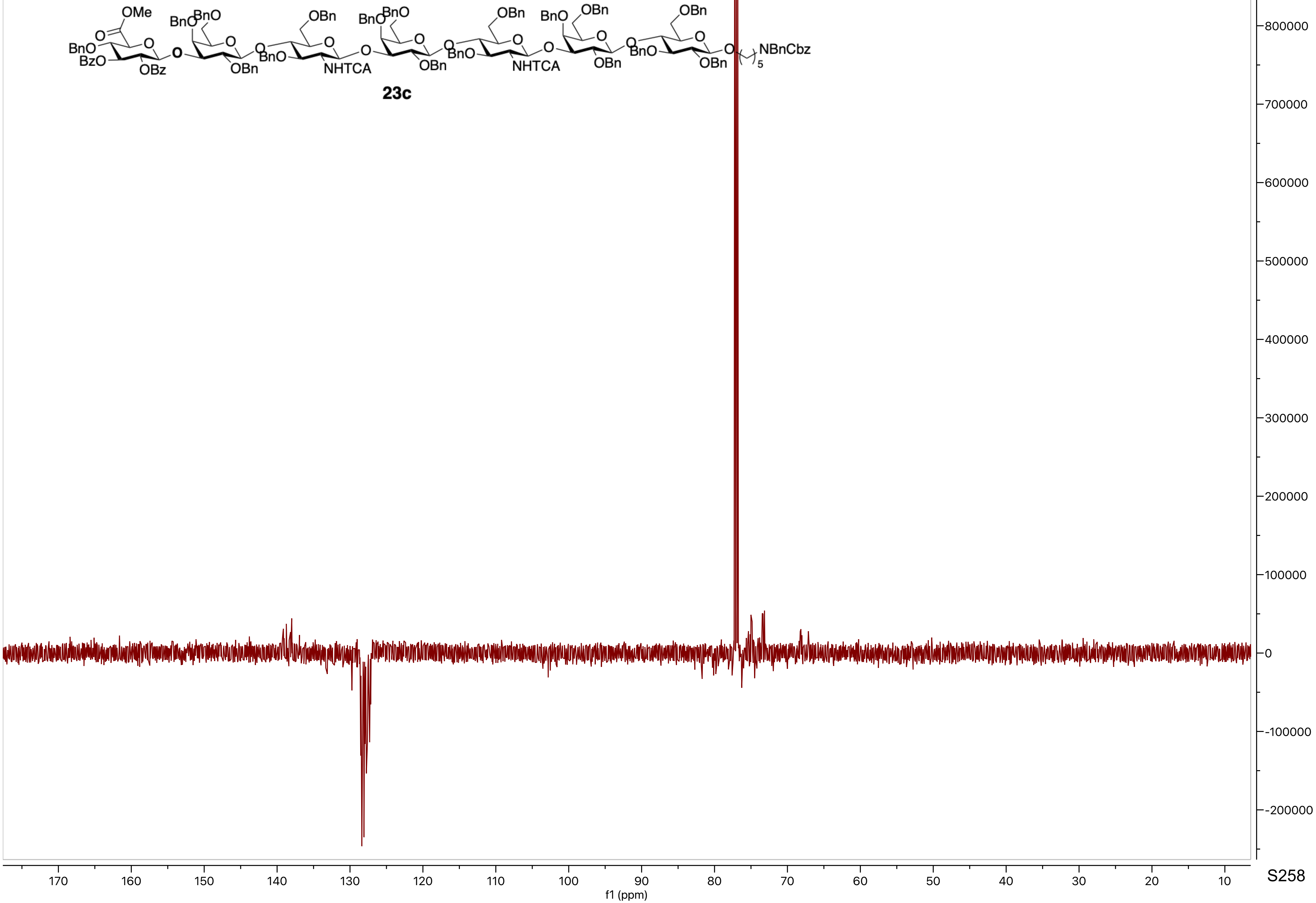

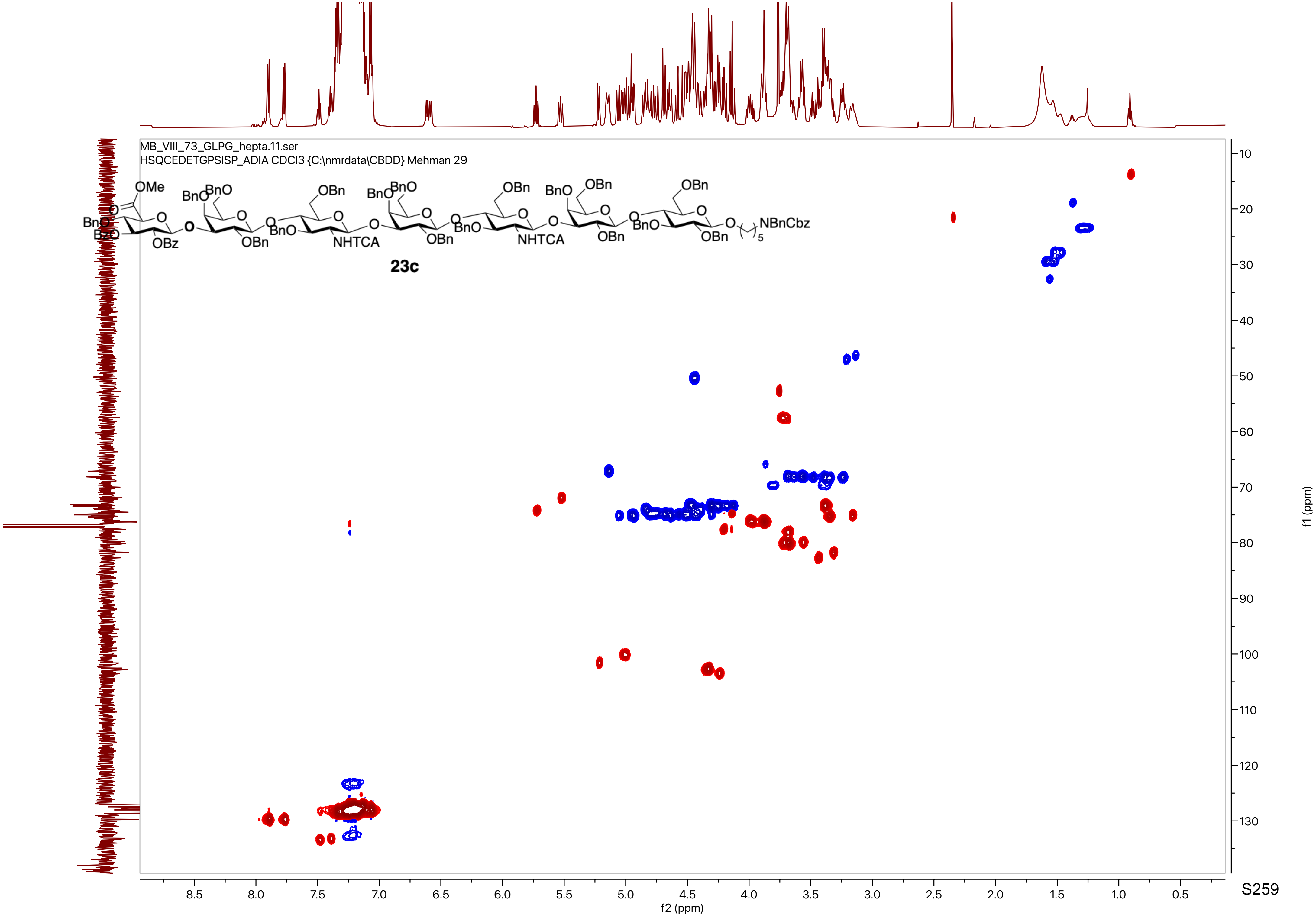
MB_VIII_73_GLPG_hepta.11.ser
HSQCEDETGPSISP_ADIA CDCl3 {C:\nmrdata\CBDD} Mehman 29
23c
f1 (ppm)
f2 (ppm)

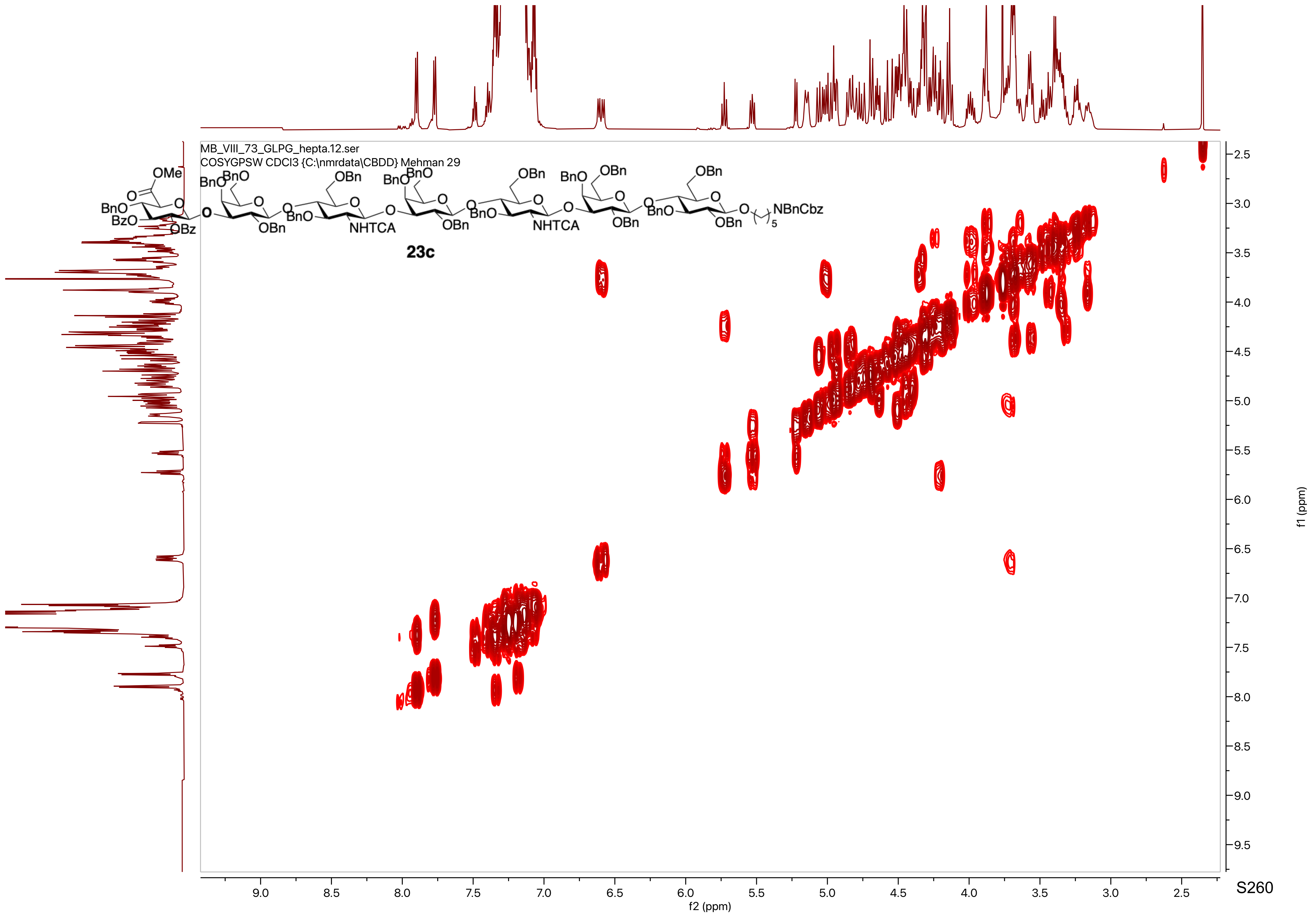
MB_VIII_73_GLPG_hepta.12.ser
COSYGPSW CDCl3 {C:\nmrdata\CBDD} Mehman 29
23c
f2 (ppm)
f1 (ppm)

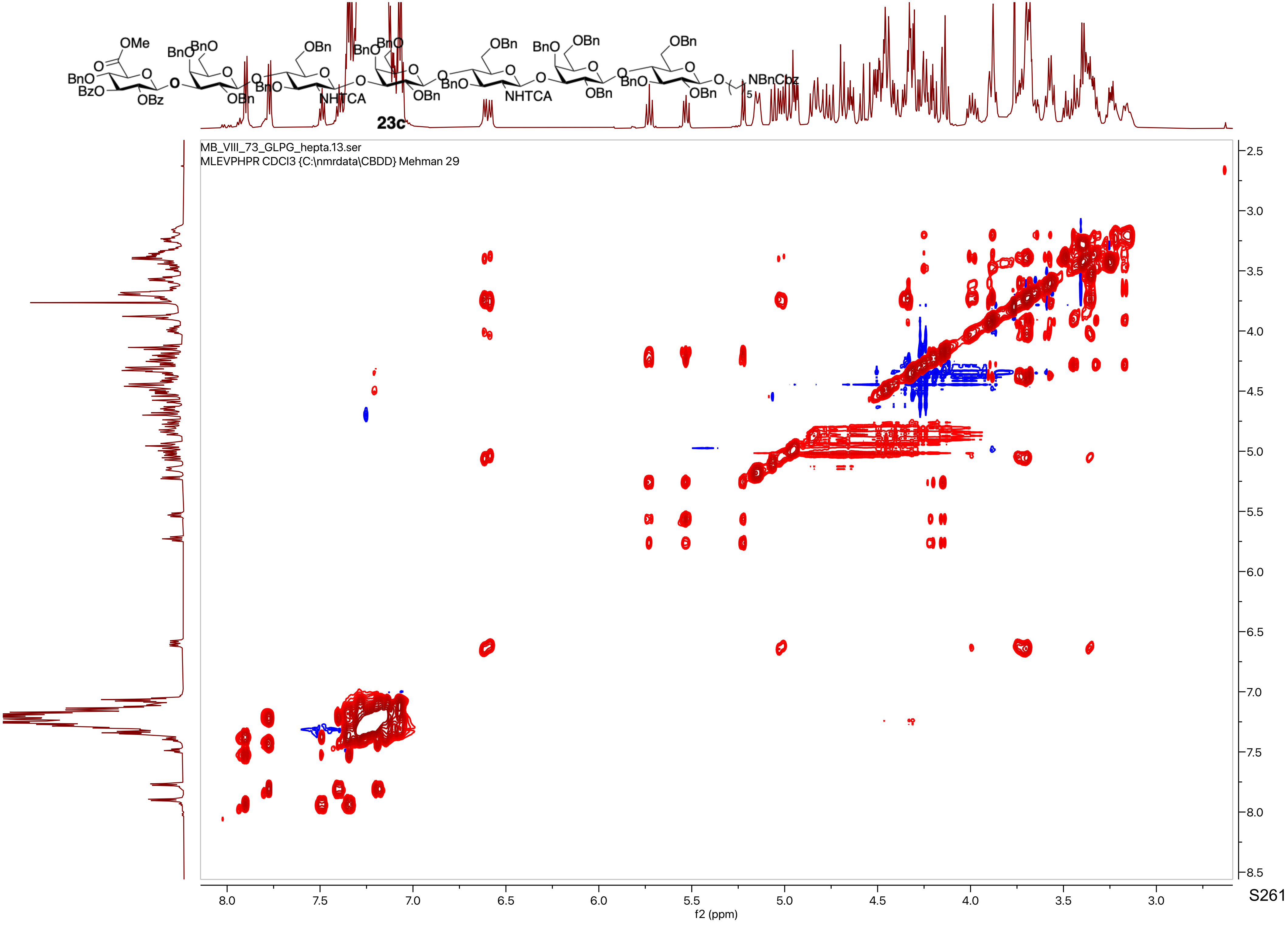

23c
MB_VIII_73_GLPG_hepta.13.ser
MLEVPHPR CDCl3 {C:\nmrdata\CBDD} Mehman 29
f2 (ppm)
f1 (ppm)

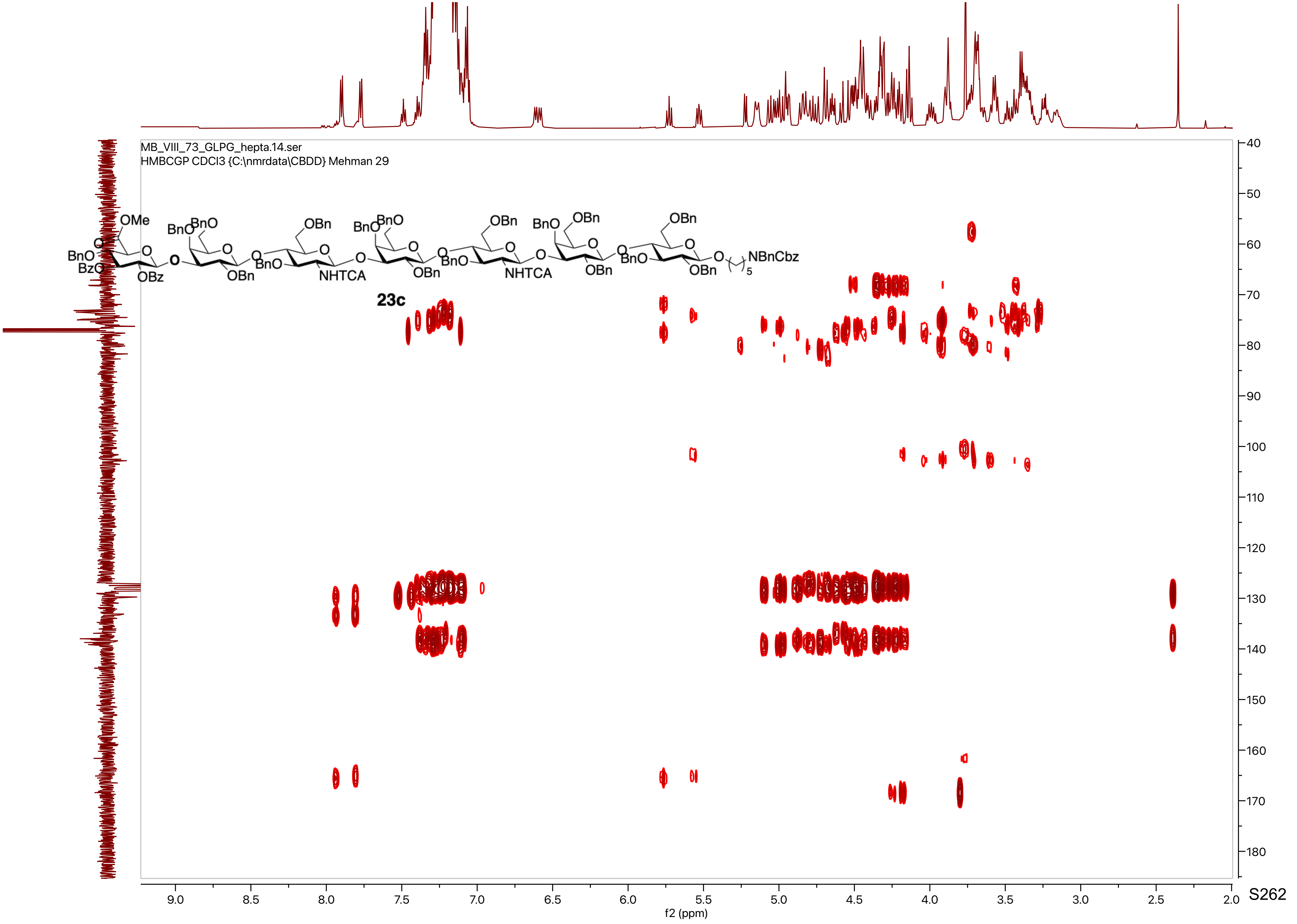
MB_VIII_73_GLPG_hepta.14.ser
HMBCGP CDCl3 {C:\nmrdata\CBDD} Mehman 29
23c
f2 (ppm)
f1 (ppm)

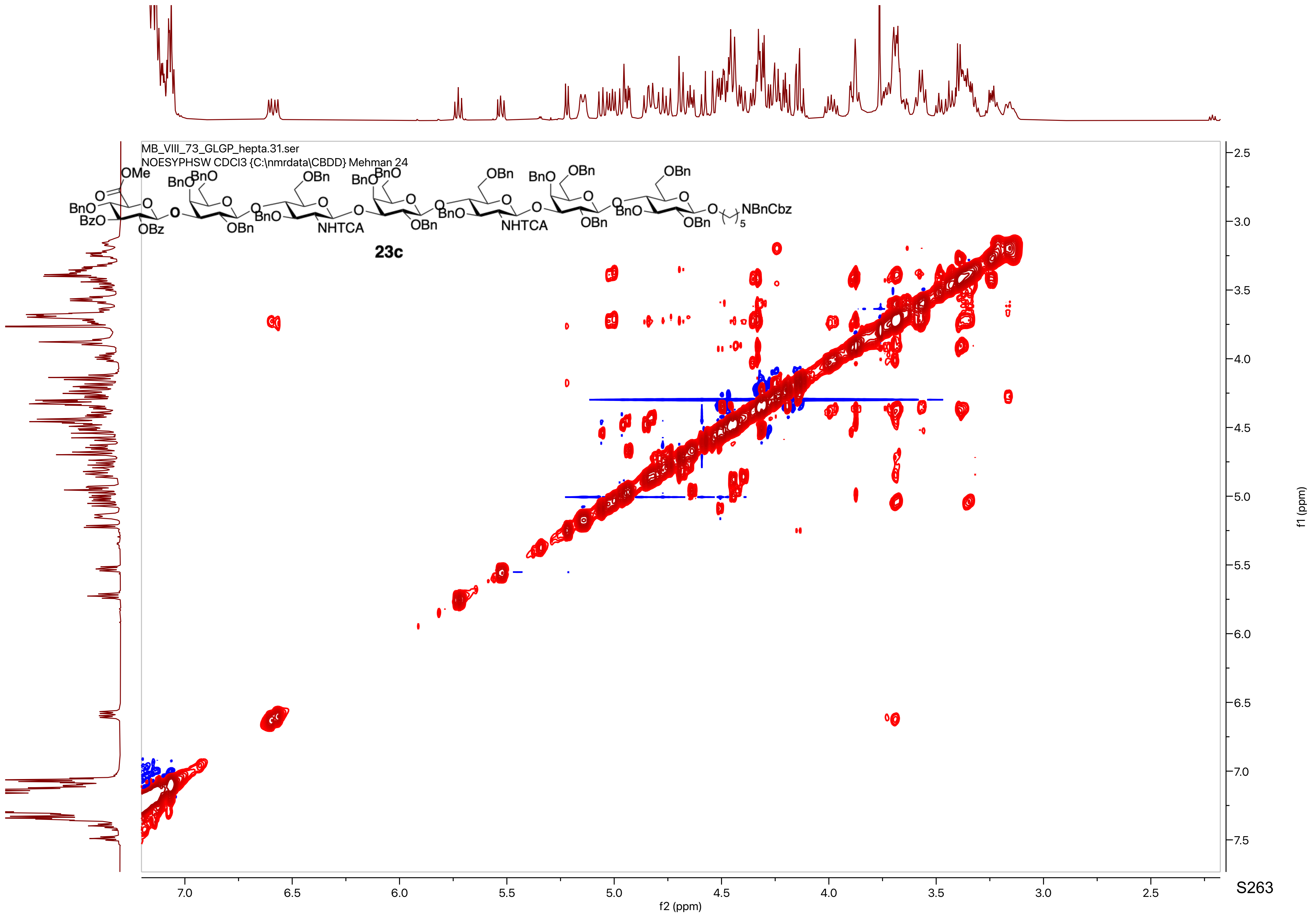
MB_VIII_73_GLGP_hepta.31.ser
NOESYPHSW CDCl3 {C:\nmrdata\CBDD} Mehman 24
23c
f2 (ppm)
f1 (ppm)

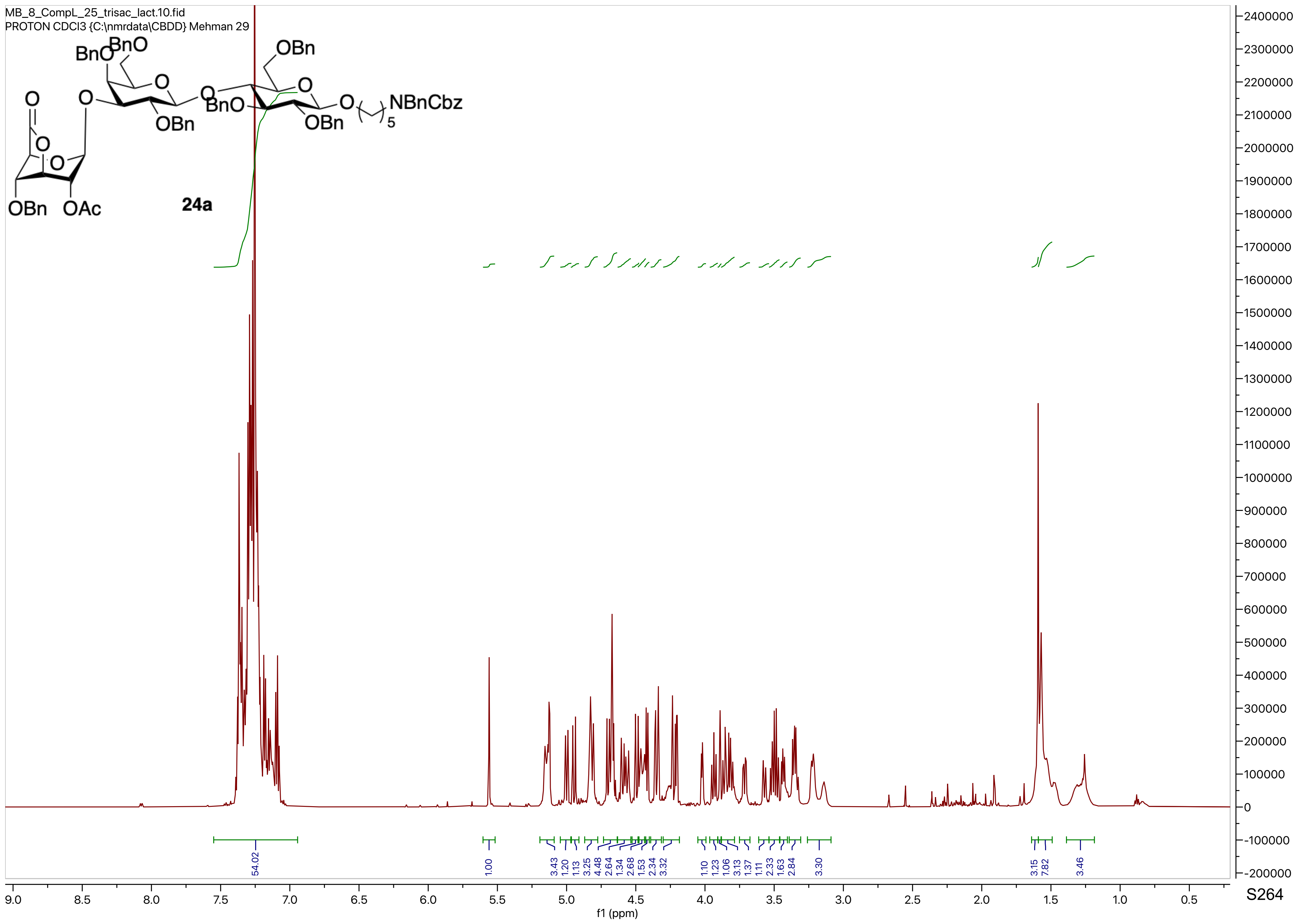
MB_8_CompL_25_trisac_lact.10.fid
PROTON CDCl3 {C:\nmrdata\CBDD} Mehman 29
BnO
BnO
OBn
O
O
O
BnO
O
NBnCbz
5
OBn
OBn
O
O
O
OBn
OAc
24a
54.02
1.00
3.43
1.20
1.13
3.25
4.48
2.64
1.34
2.68
1.53
2.34
3.32
1.10
1.23
1.06
3.13
1.37
1.11
2.33
1.63
2.84
3.30
3.15
7.82
3.46
f1 (ppm)

MB_8_CompL_25_trisac_lact.13.fid
C13APT CDCl3 {C:\nmrdata\CBDD} Mehman 29

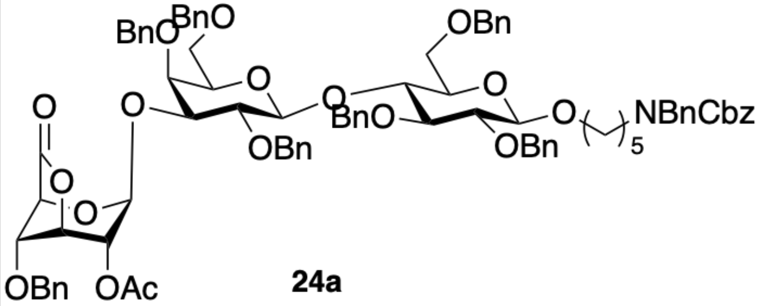


**24a**

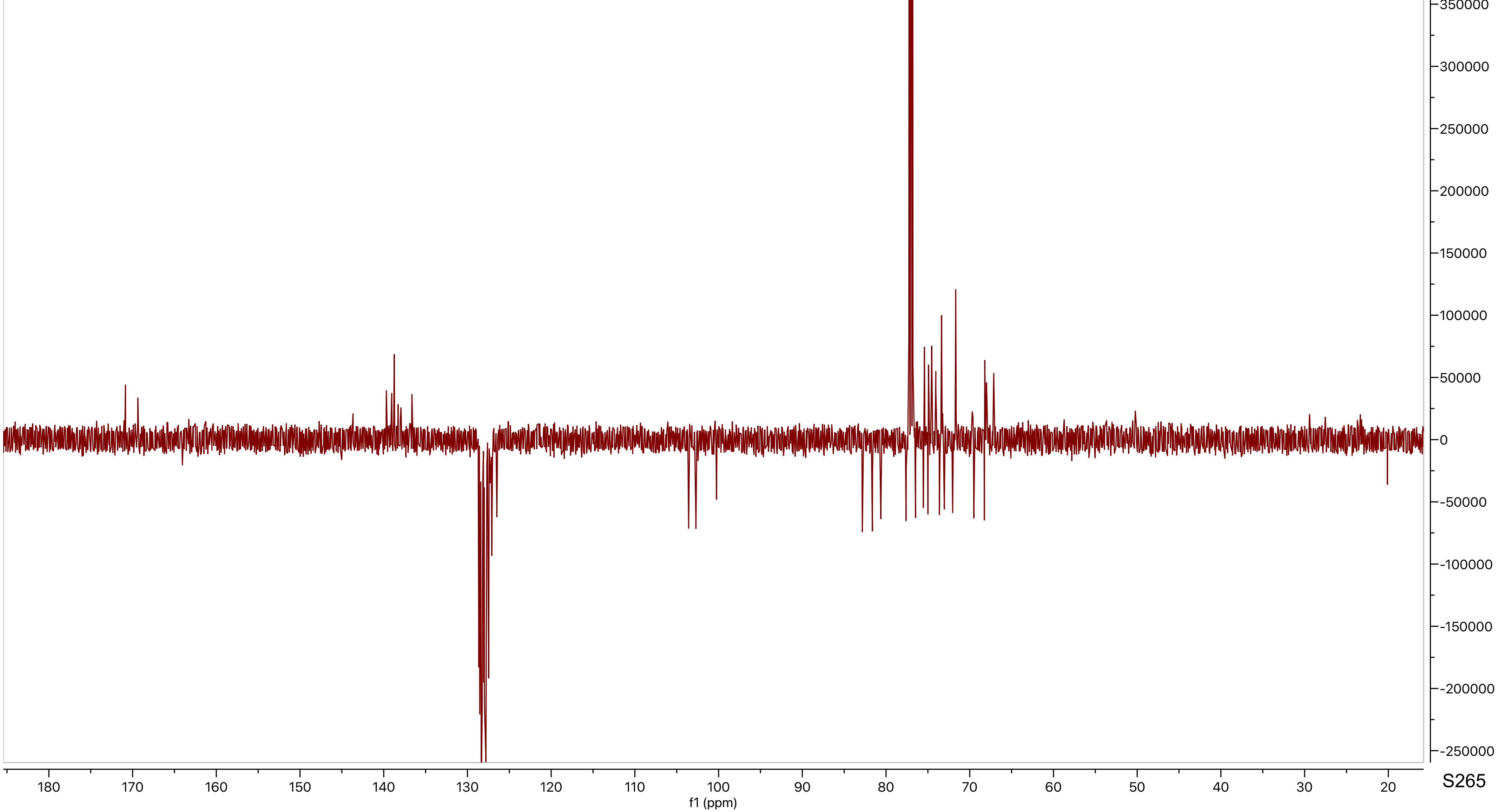

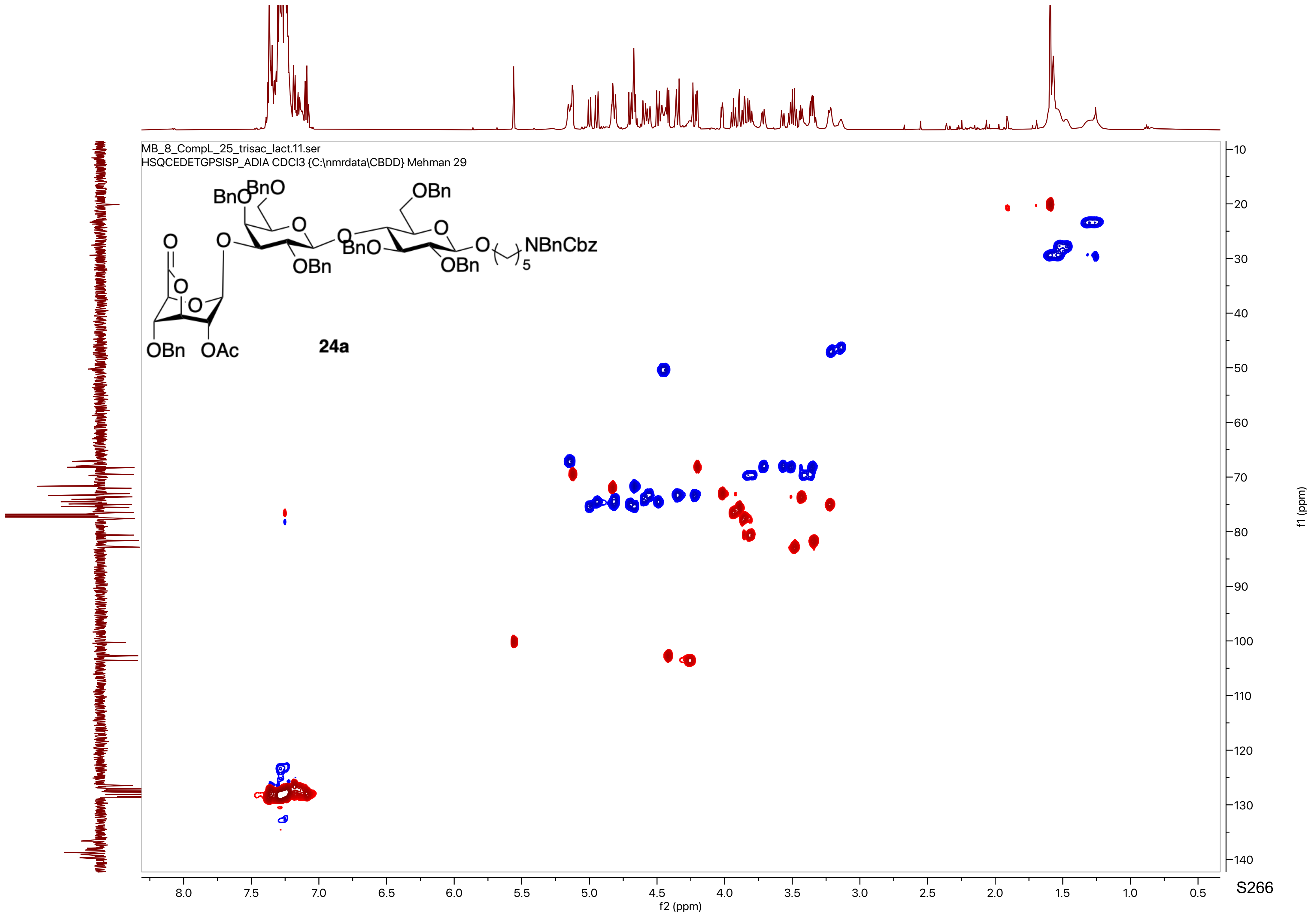

MB_8_CompL_25_trisac_lact.11.ser
HSQCEDETGPSISP_ADIA CDCl3 {C:\nmrdata\CBDD} Mehman 29
BnO
BnO
OBn
O
O
O
O
O
BnO
O
OBn
OBn
O
NBnCbz
5
O
O
OBn
OAc
24a
f2 (ppm)
f1 (ppm)

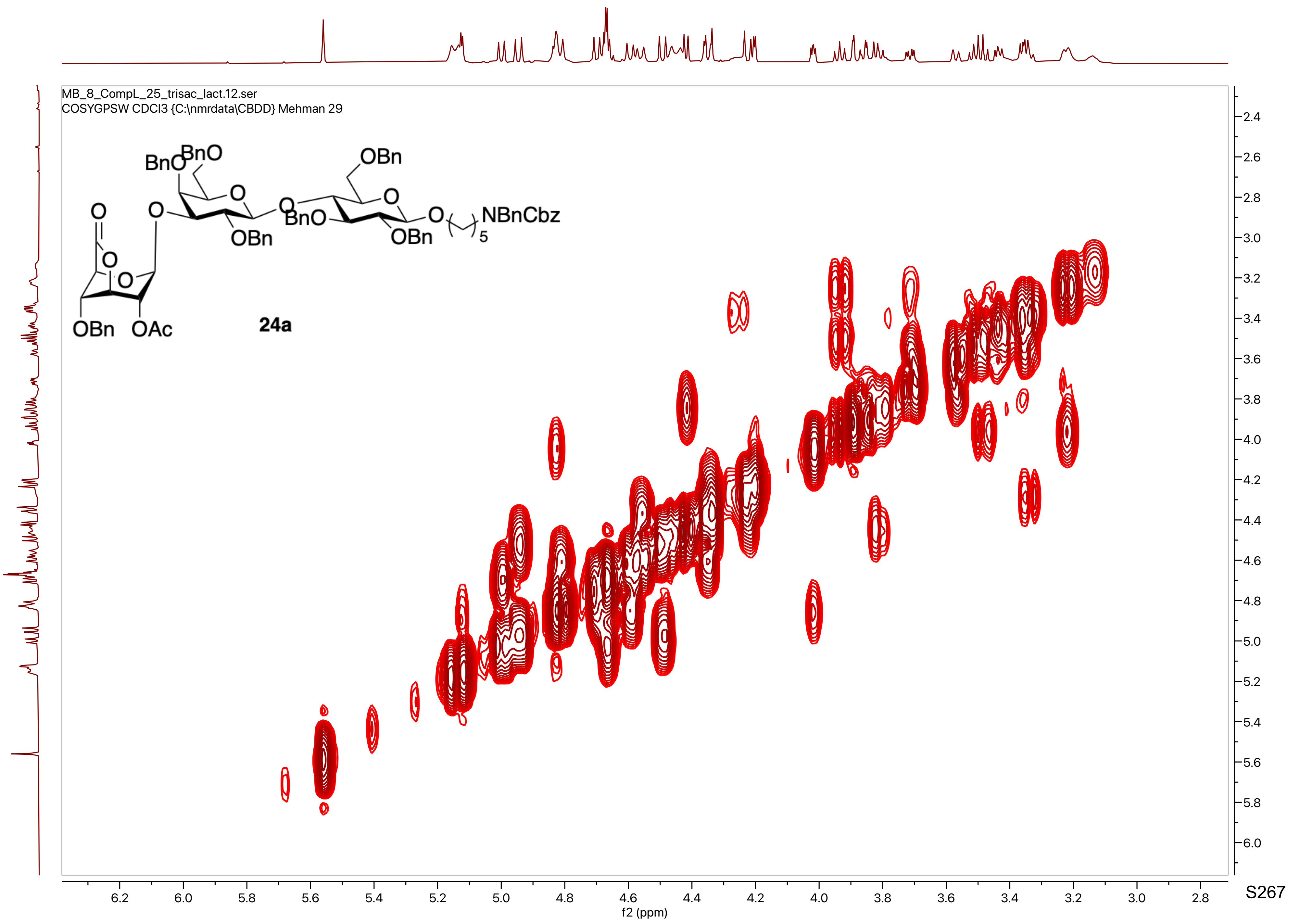
MB_8_CompL_25_trisac_lact.12.ser
COSYGPSW CDCl3 {C:\nmrdata\CBDD} Mehman 29
24a
f2 (ppm)
f1 (ppm)

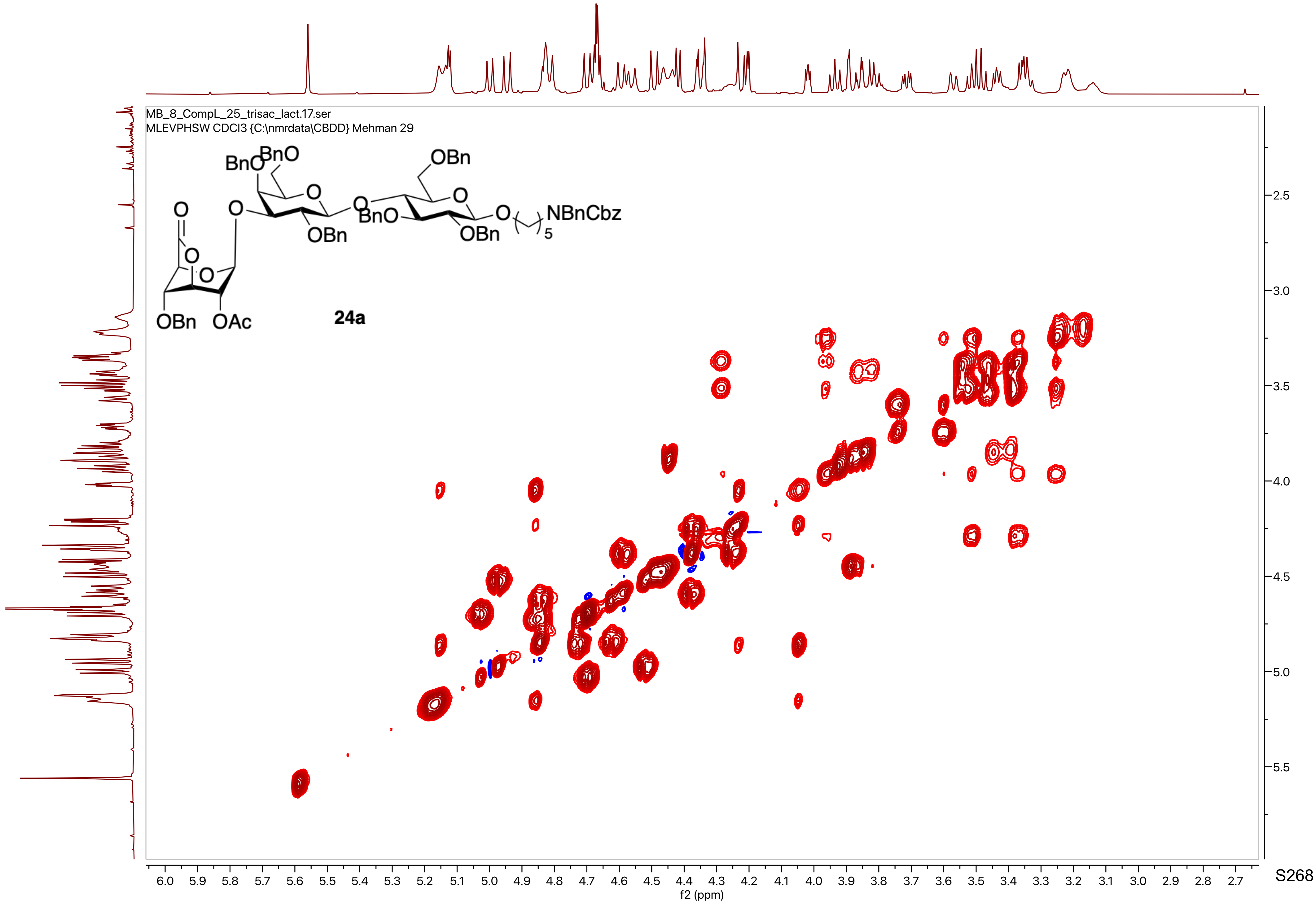

MB_8_CompL_25_trisac_lact.17.ser
MLEVPHSW CDCl3 {C:\nmrdata\CBDD} Mehman 29
BnO
BnO
OBn
O
O
O
O
O
BnO
OBn
OBn
O
NBnCbz
5
O
O
OBn
OAc
24a
f2 (ppm)
f1 (ppm)

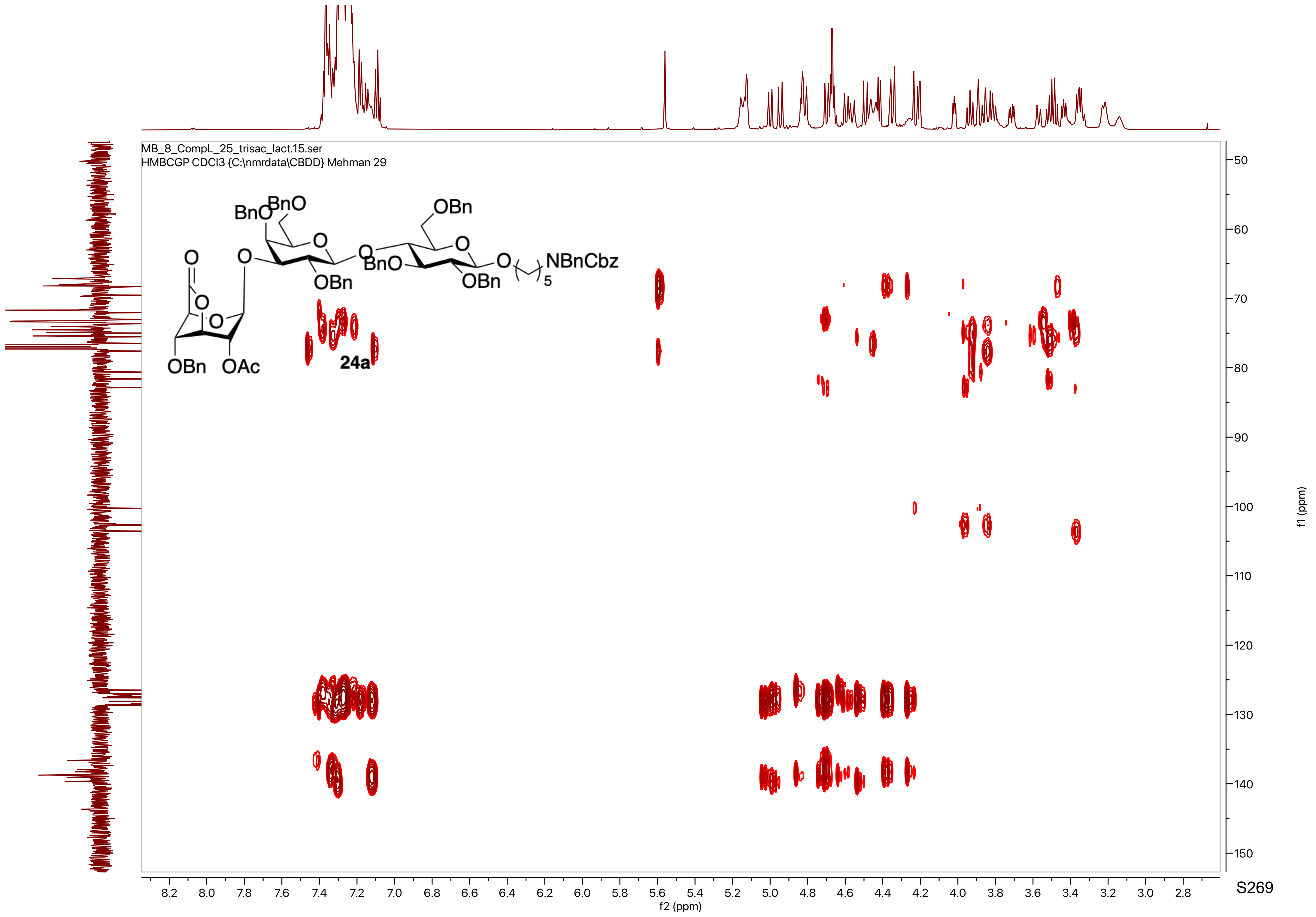
MB_8_CompL_25_trisac_lact.15.ser
HMBCGP CDCl3 {C:\nmrdata\CBDD} Mehman 29
24a
f2 (ppm)
f1 (ppm)

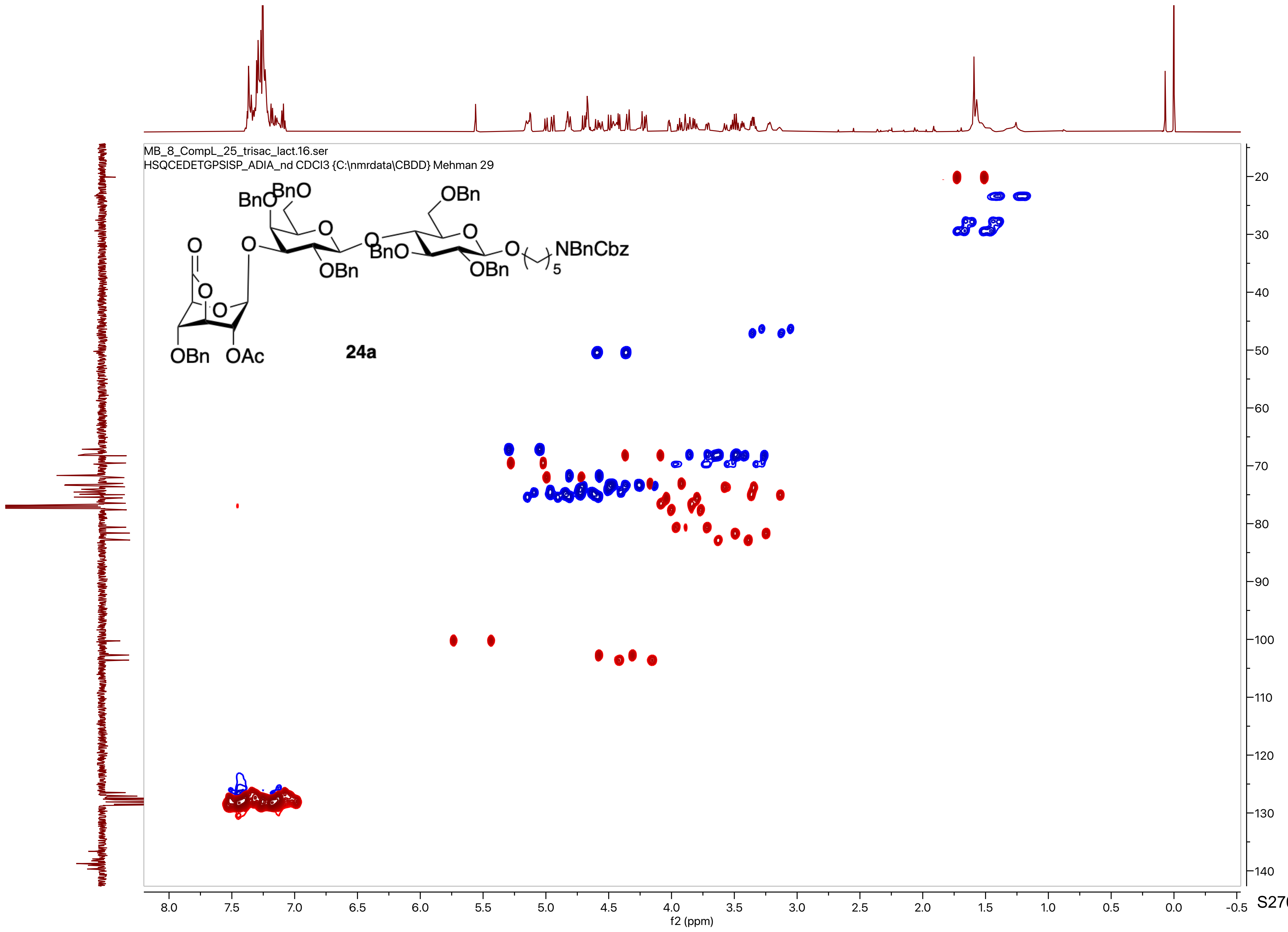
MB_8_CompL_25_trisac_lact.16.ser
HSQCEDETGPSISP_ADIA_nd CDCl3 {C:\nmrdata\CBDD} Mehman 29
BnO
BnO
OBn
O
O
O
O
BnO
OBn
OBn
O
NBnCbz
5
OBn
OAc
24a
f2 (ppm)
f1 (ppm)

MB_8_comp_L_26_penta_lactone.10.fid
PROTON CDCl3 {C:\nmrdata\CBDD} Mehman 46

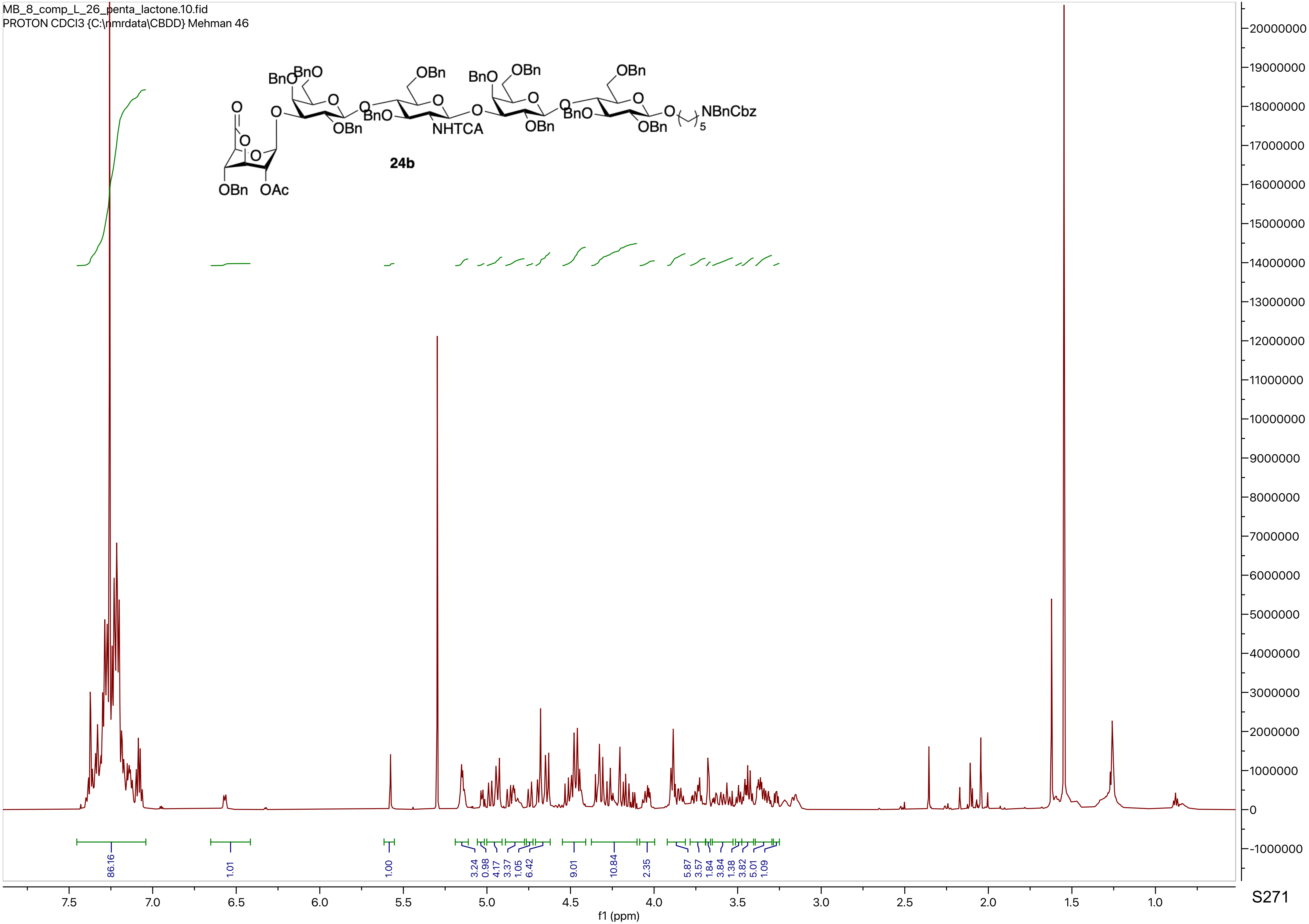

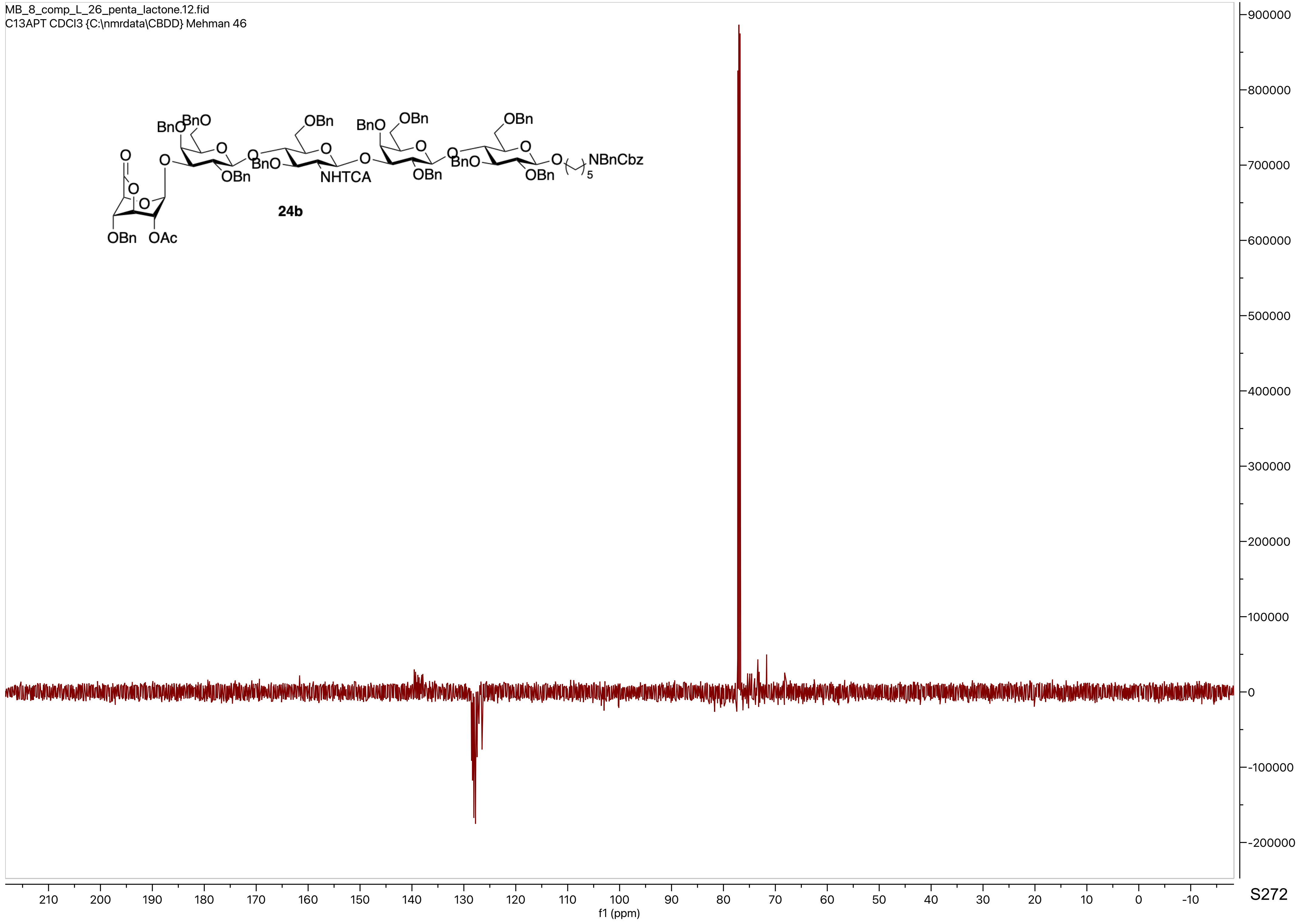
MB_8_comp_L_26_penta_lactone.12.fid
C13APT CDCl3 {C:\nmrdata\CBDD} Mehman 46
BnO BnO O OBn BnO OBn NHTCA BnO OBn OBn BnO OBn OBn NBnCbz 5
O OBn OAc
24b
900000
800000
700000
600000
500000
400000
300000
200000
100000
0
-100000
-200000
210 200 190 180 170 160 150 140 130 120 110 100 90 80 70 60 50 40 30 20 10 0 -10
f1 (ppm)

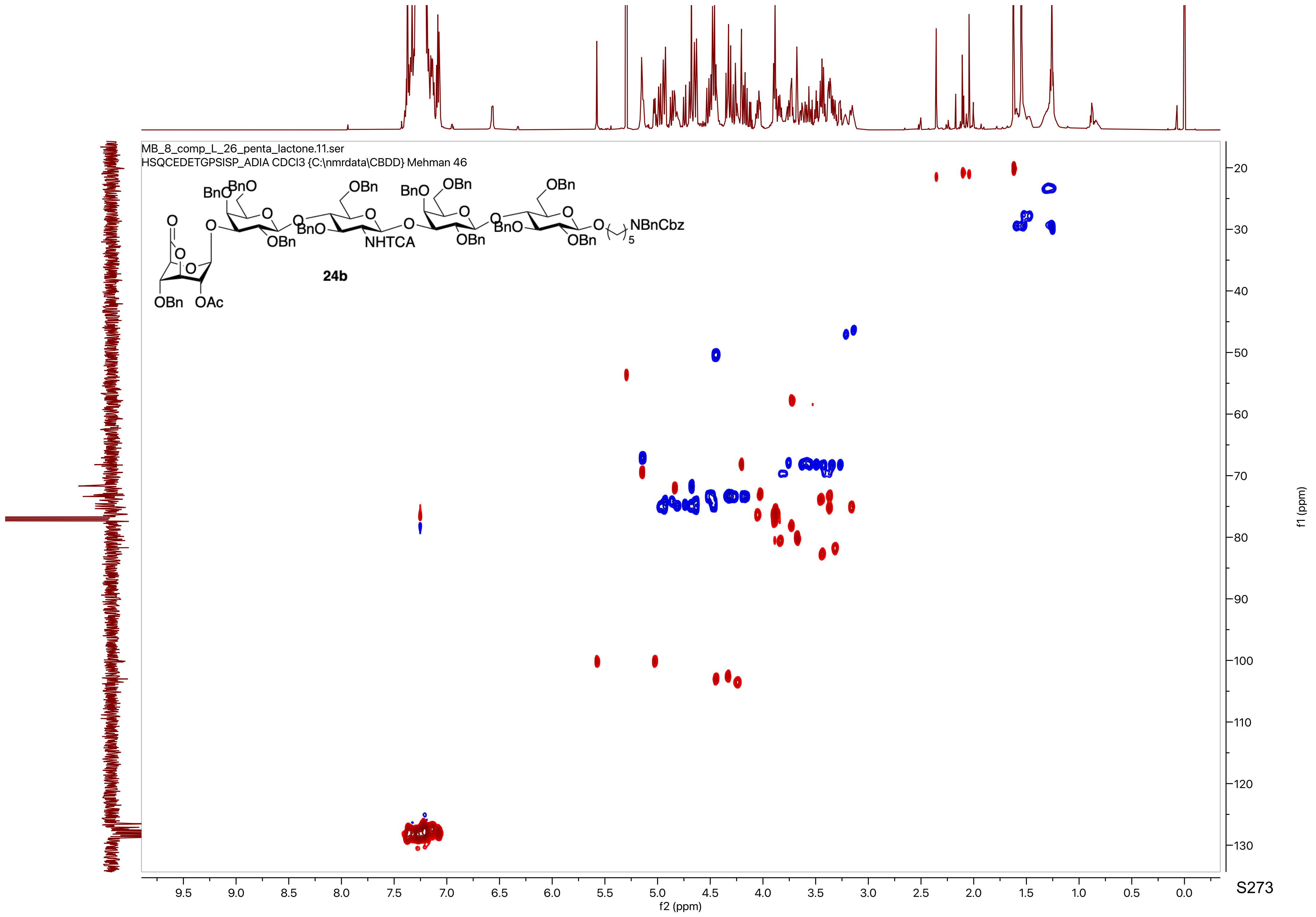

MB_8_comp_L_26_penta_lactone.11.ser
HSQCEDETGPSISP_ADIA CDCl3 {C:\nmrdata\CBDD} Mehman 46
24b
f2 (ppm)
f1 (ppm)

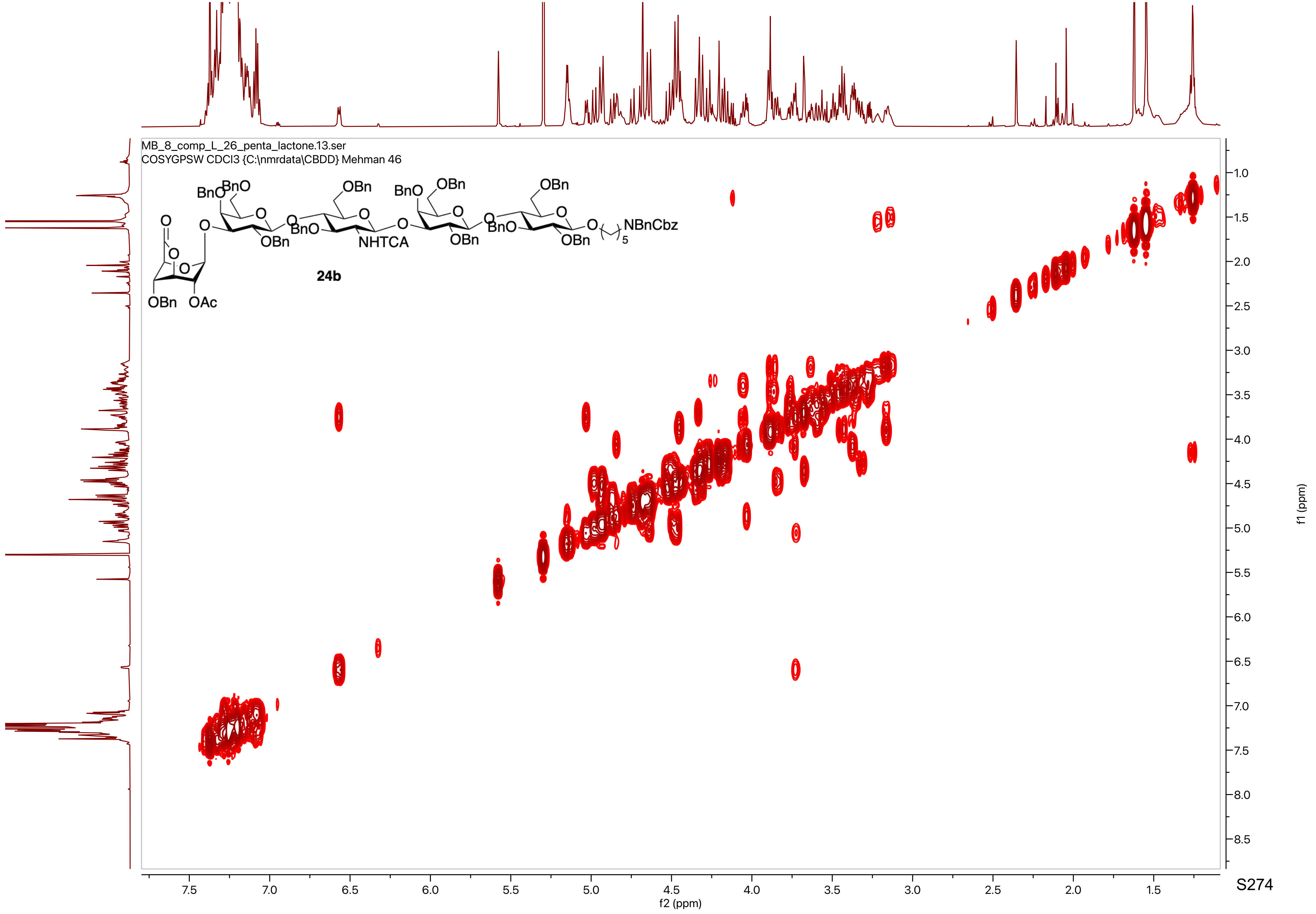

MB_8_comp_L_26_penta_lactone.13.ser
COSYGPSW CDCl3 {C:\nmrdata\CBDD} Mehman 46
24b
f2 (ppm)
f1 (ppm)

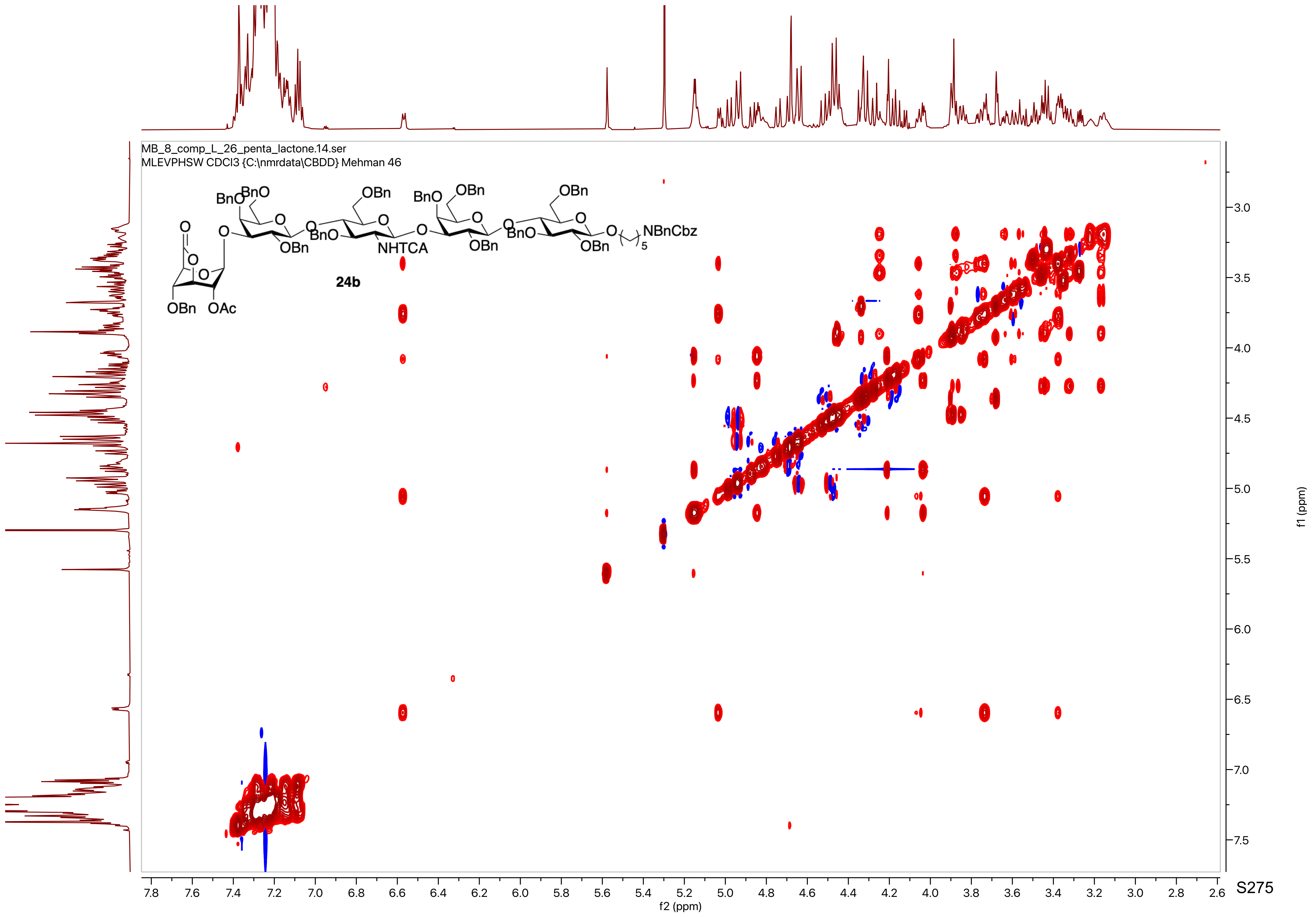
MB_8_comp_L_26_penta_lactone.14.ser
MLEVPHSW CDCl3 {C:\nmrdata\CBDD} Mehman 46
BnO
BnO
OBn
OBn
BnO
OBn
BnO
OBn
BnO
OBn
NHTCA
OBn
NBnCbz
5
OBn
OAc
24b
f2 (ppm)
f1 (ppm)

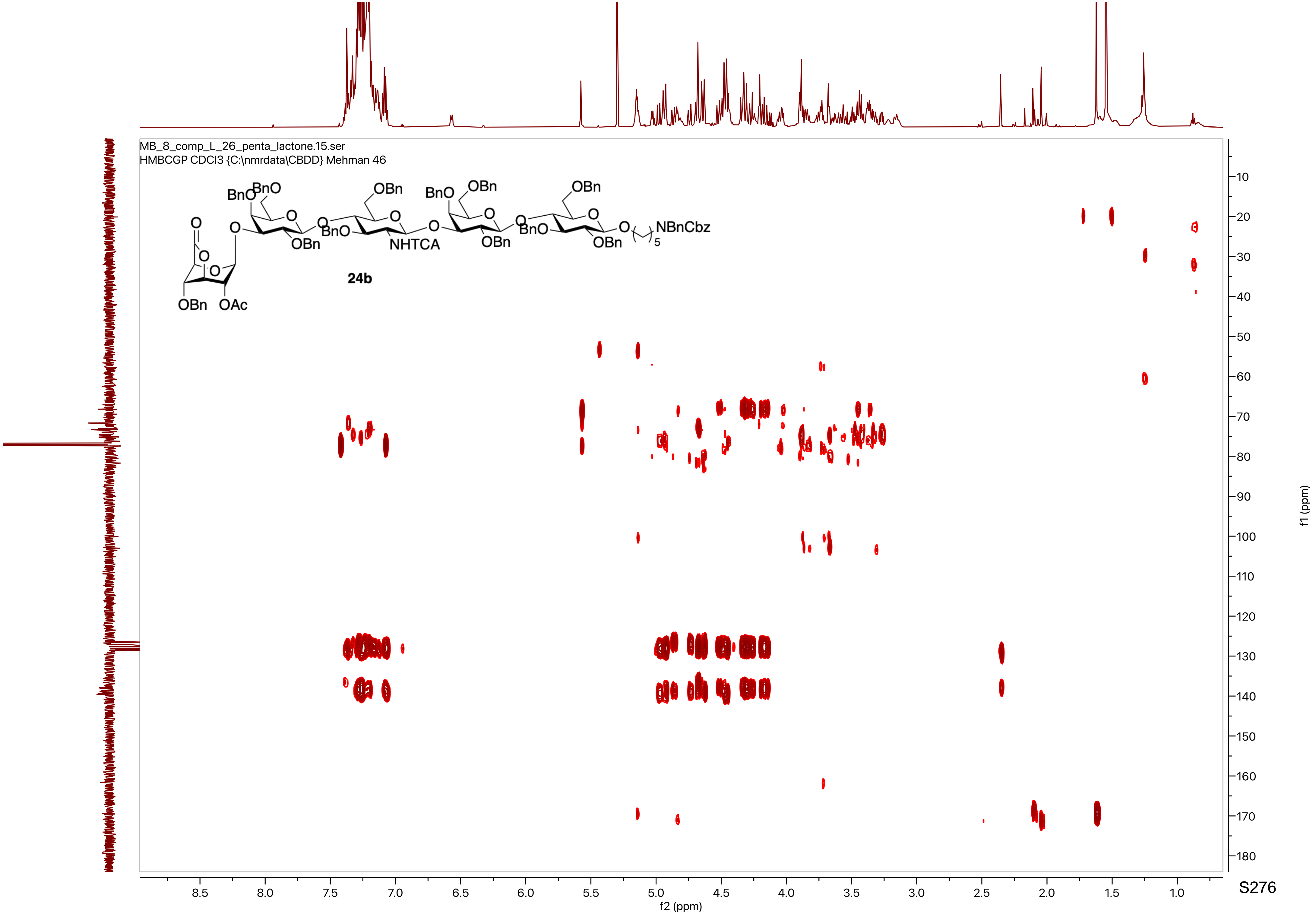
MB_8_comp_L_26_penta_lactone.15.ser
HMBCGP CDCl3 {C:\nmrdata\CBDD} Mehman 46
24b
f2 (ppm)
f1 (ppm)

MB_8_CompL_27.10.fid
PROTON CDCl3 {C:\nmrdata\CBDD} Mehman 46

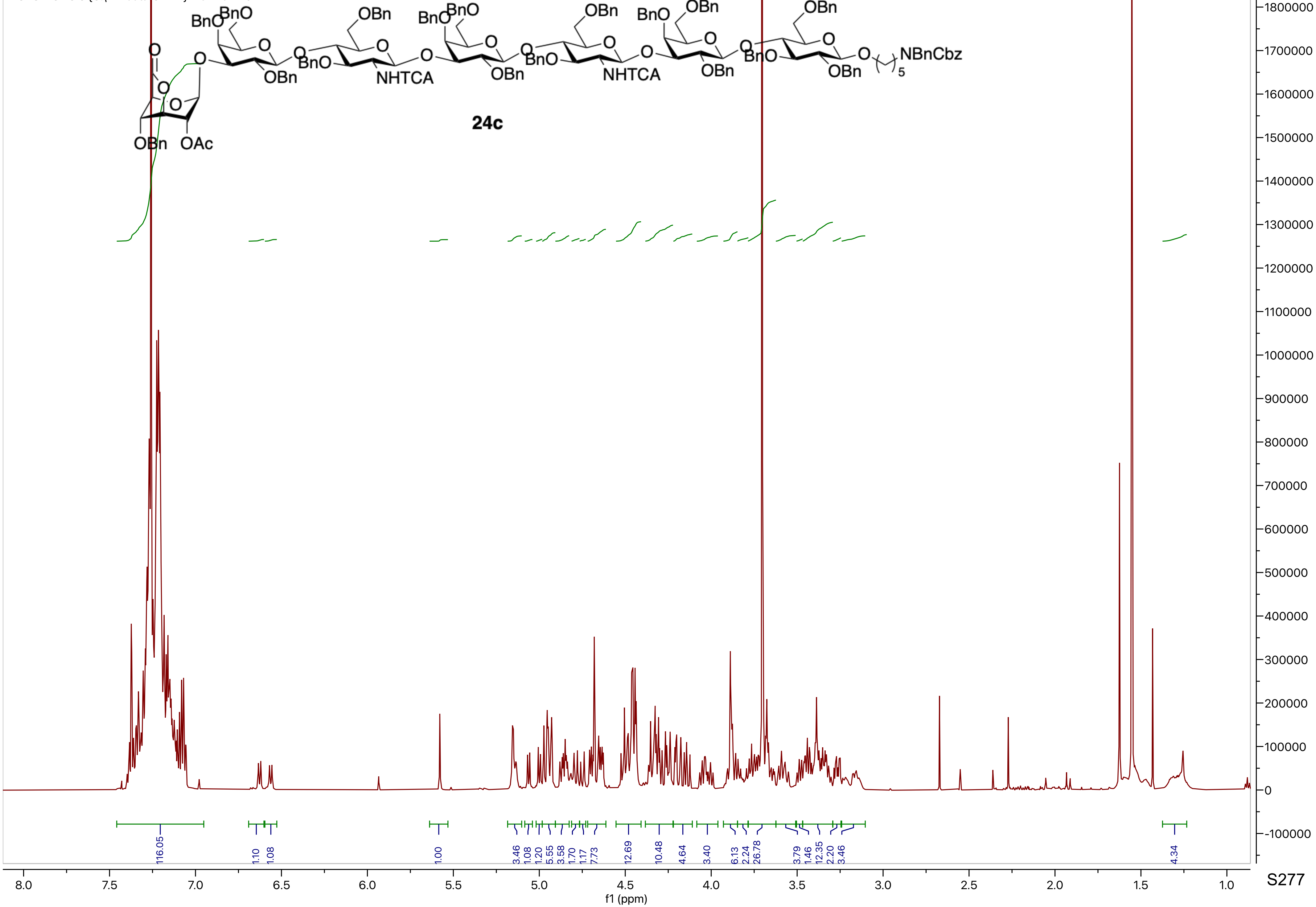

MB_8_CompL_27.12.fid
C13APT CDCl3 {C:\nmrdata\CBDD} Mehman 46

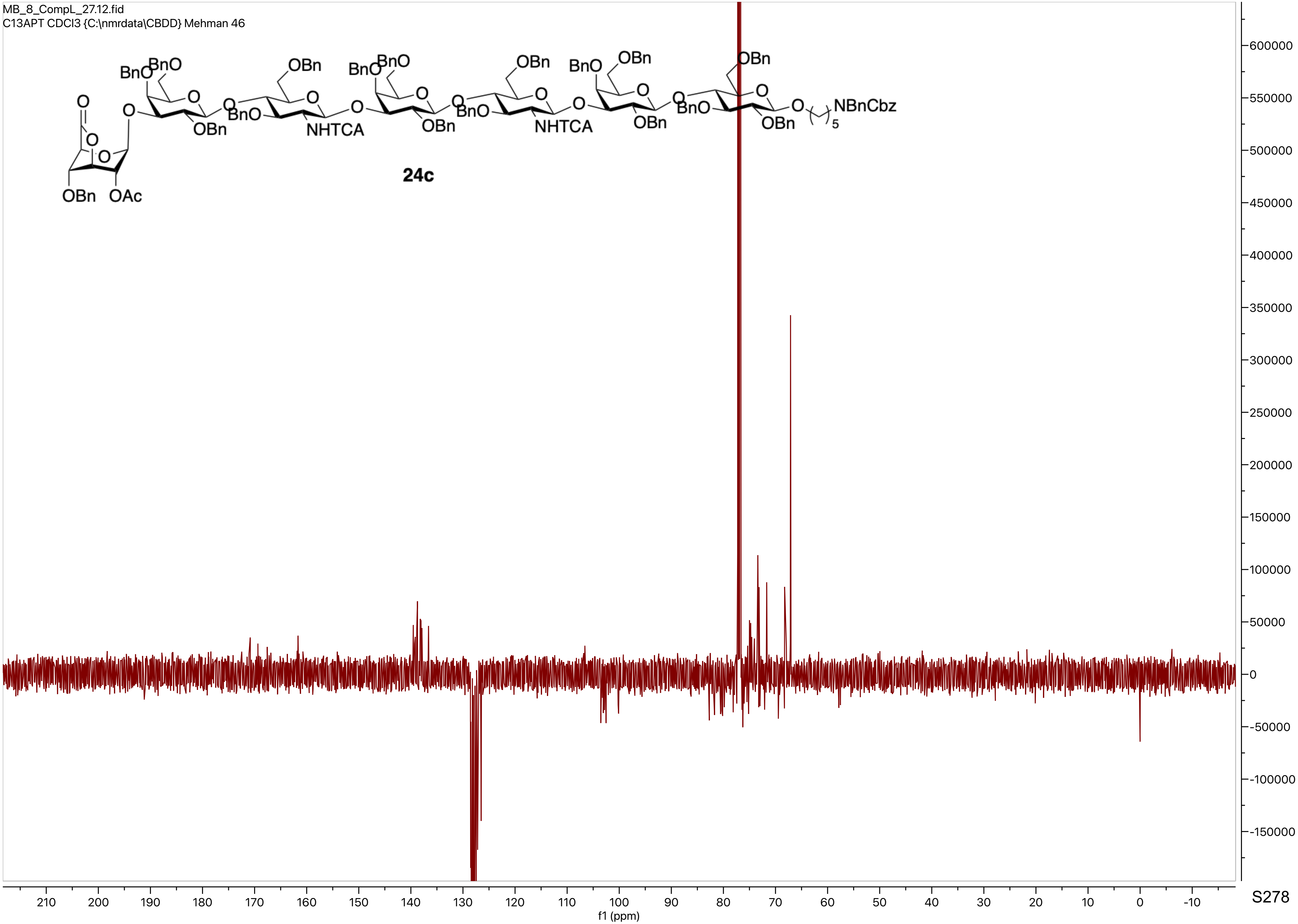

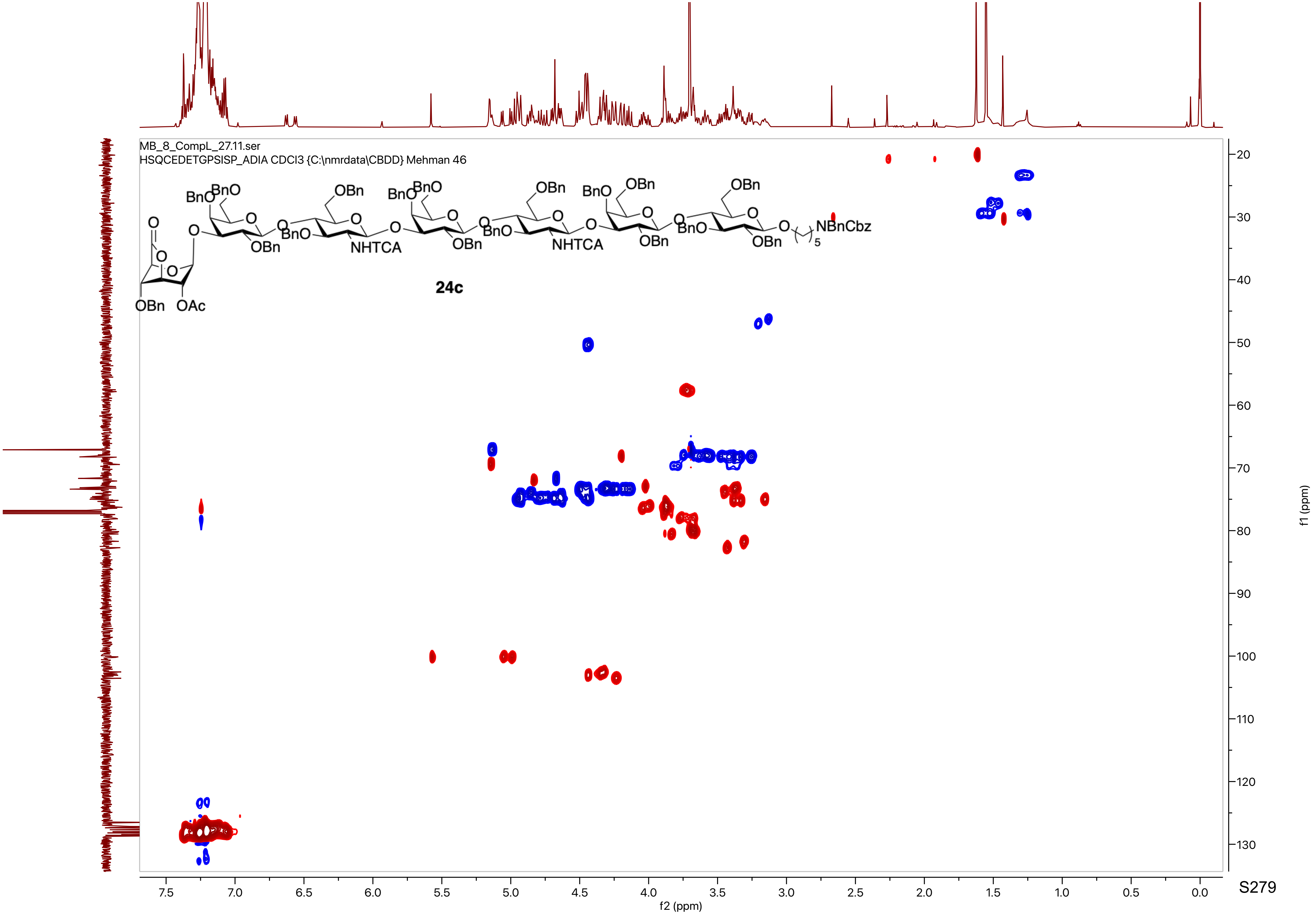
MB_8_CompL_27.11.ser
HSQCEDETGPSISP_ADIA CDCl3 {C:\nmrdata\CBDD} Mehman 46
24c
f2 (ppm)
f1 (ppm)

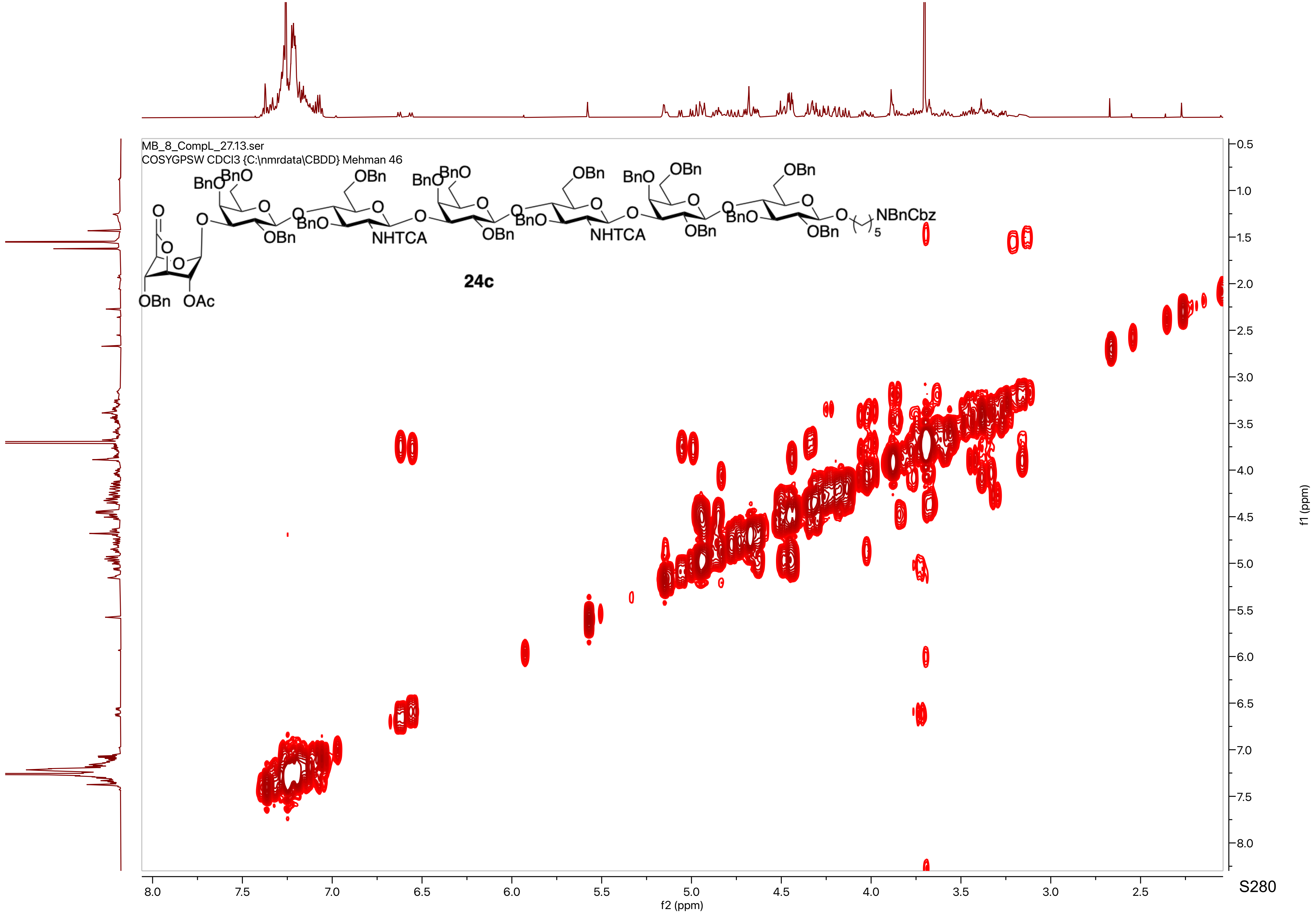
MB_8_CompL_27.13.ser
COSYGPSW CDCl3 {C:\nmrdata\CBDD} Mehman 46
BnO
OBn
O
OAc
NHTCA
NBnCbz
5
24c
f2 (ppm)
f1 (ppm)

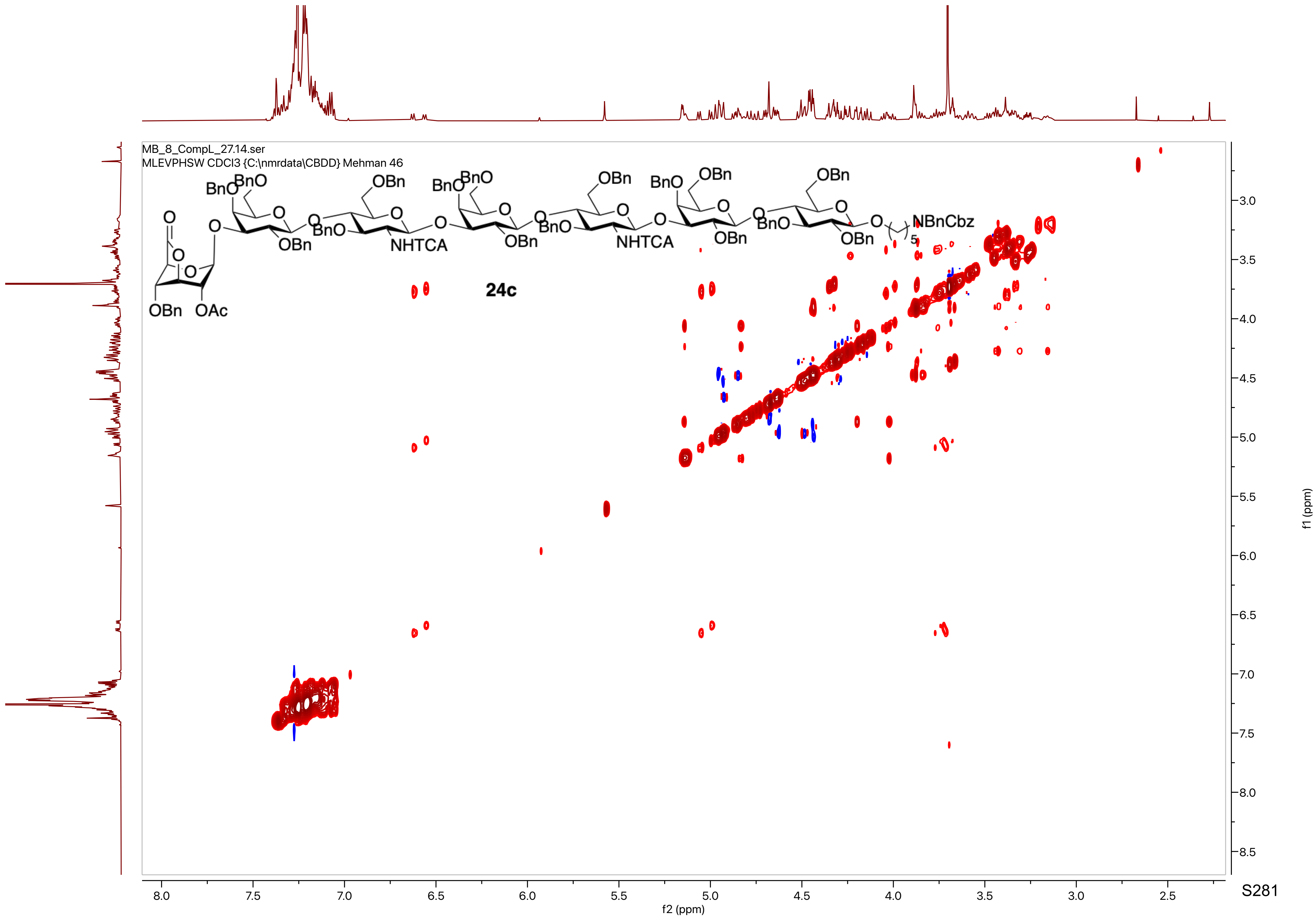

MB_8_CompL_27.14.ser
MLEVPHSW CDCl3 {C:\nmrdata\CBDD} Mehman 46
24c
f2 (ppm)
f1 (ppm)

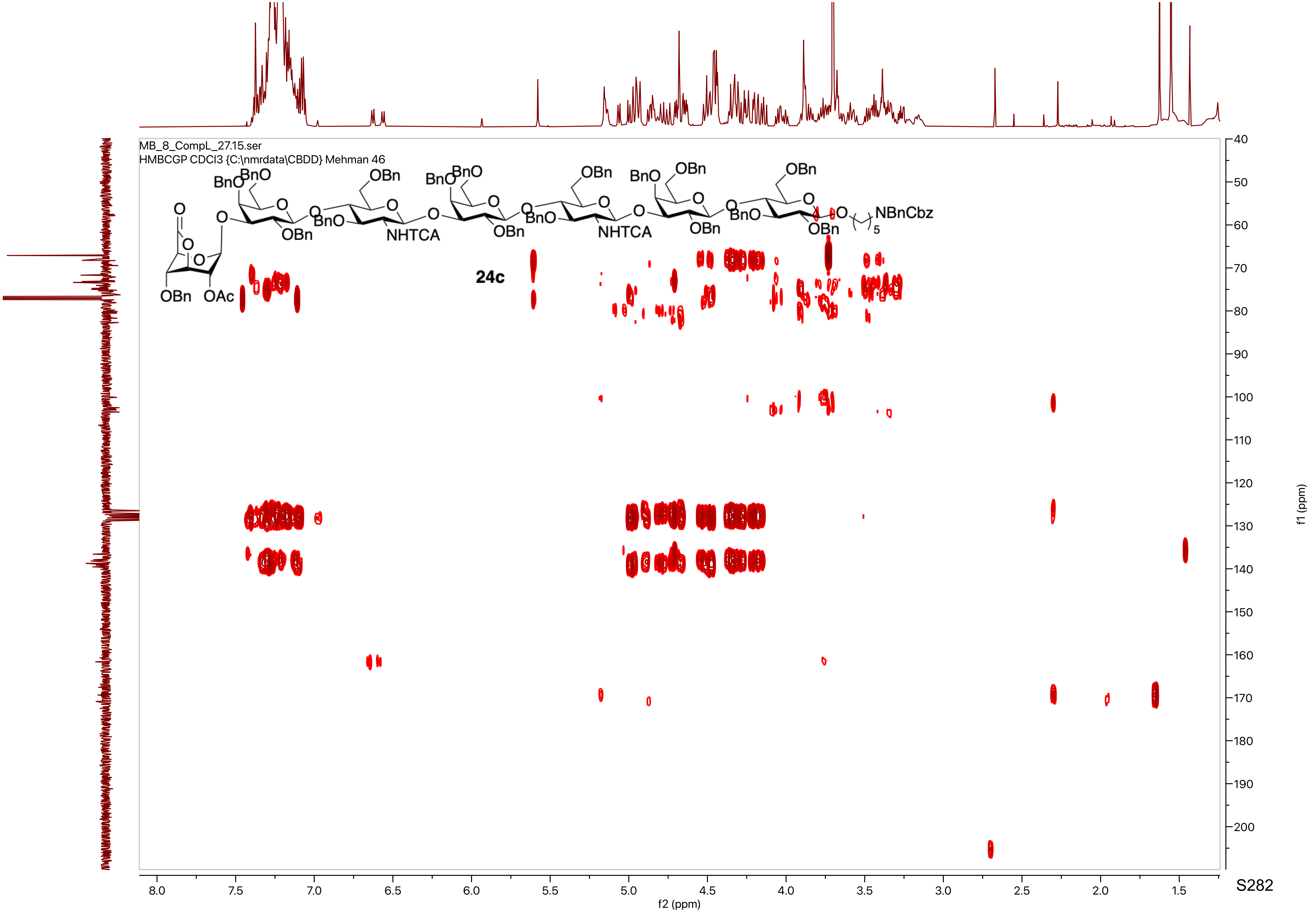
MB_8_CompL_27.15.ser
HMBCGP CDCl3 {C:\nmrdata\CBDD} Mehman 46
24c
f2 (ppm)
f1 (ppm)

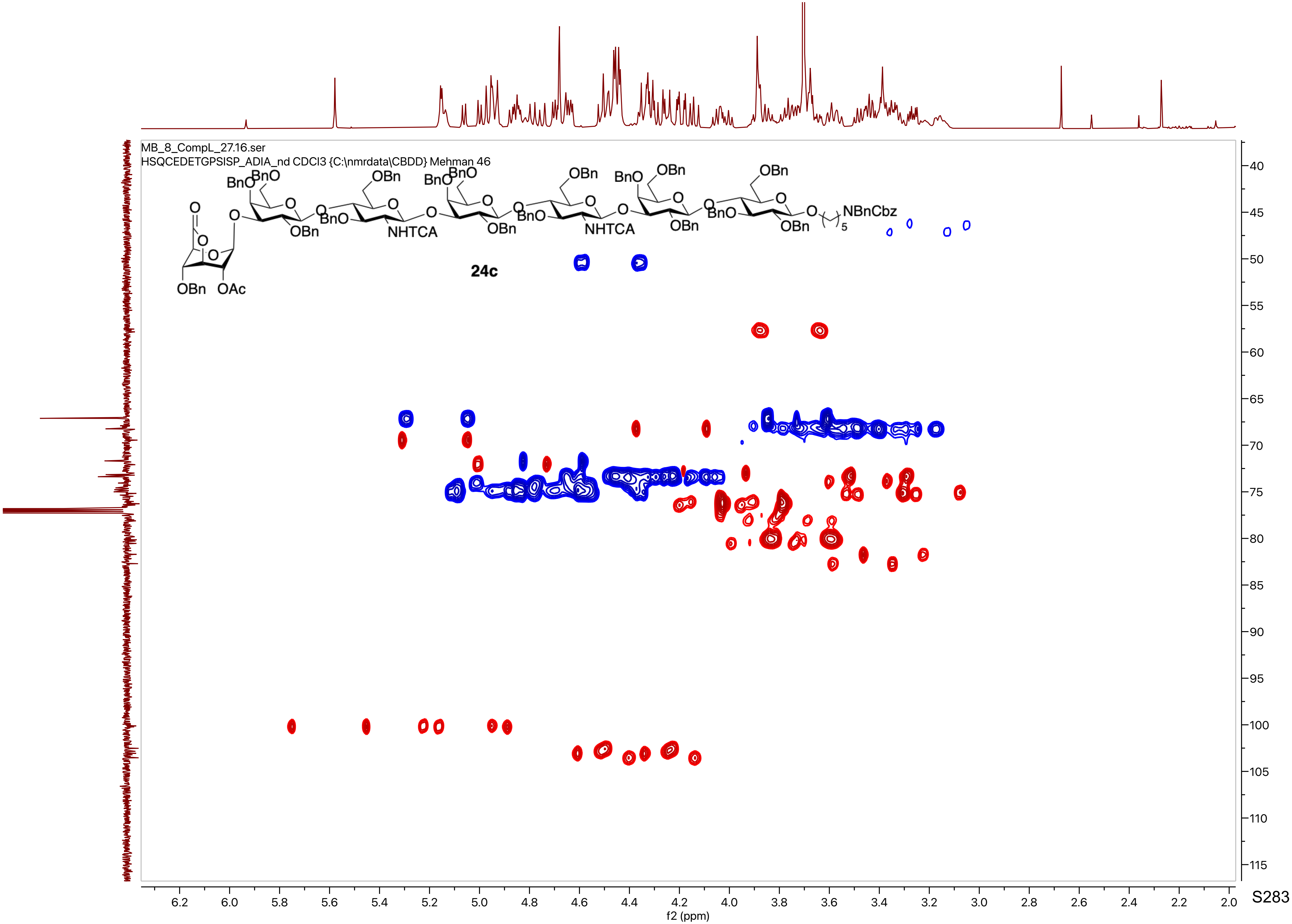
MB_8_CompL_27.16.ser
HSQCEDETGPSISP_ADIA_nd CDCl3 {C:\nmrdata\CBDD} Mehman 46
24c
BnO
OBn
NHTCA
OAc
NBnCbz
f2 (ppm)
f1 (ppm)

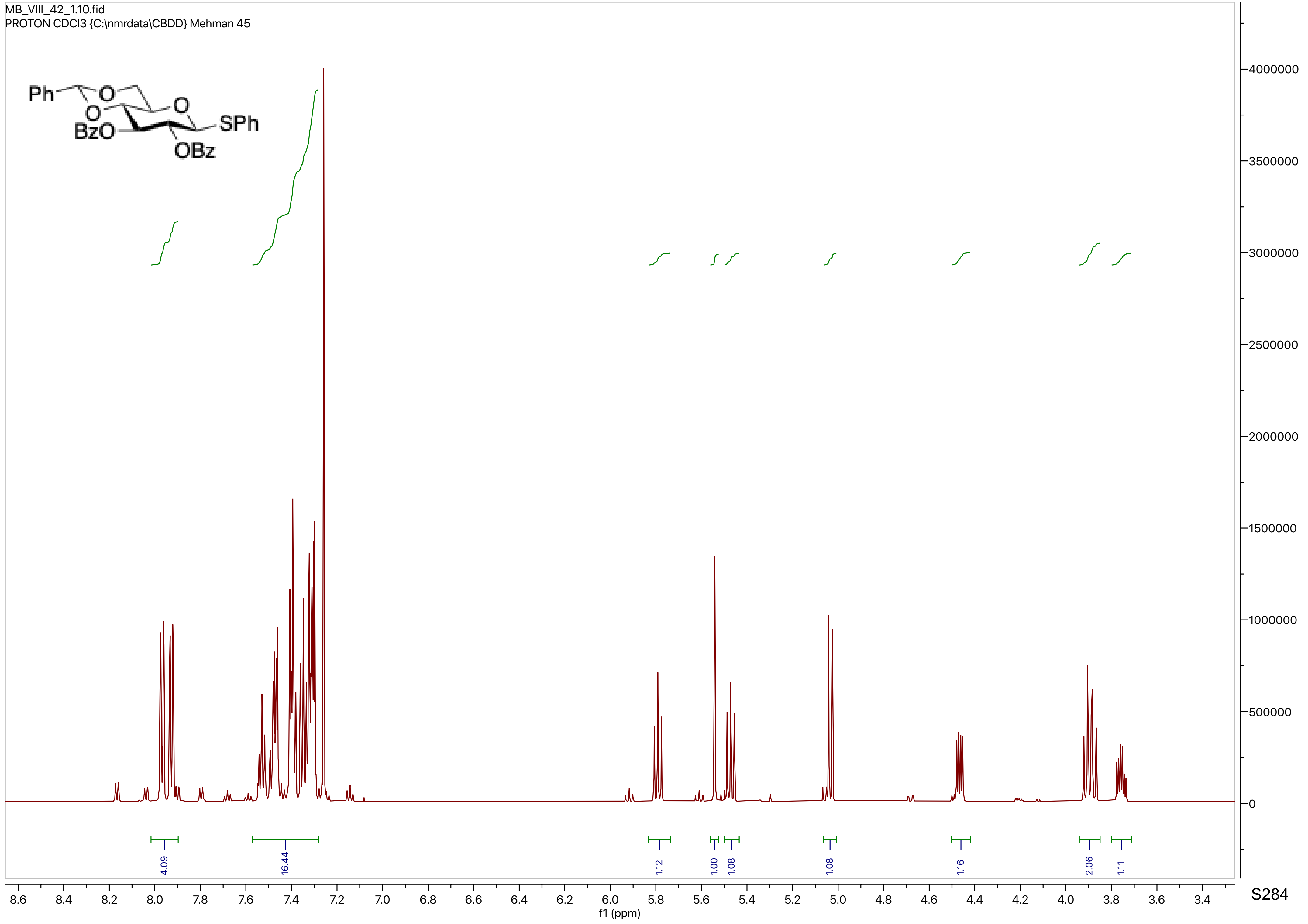

MB_VIII_42_1.10.fid
PROTON CDCl3 {C:\nmrdata\CBDD} Mehman 45
Ph
O
O
O
BzO
SPh
OBz
4000000
3500000
3000000
2500000
2000000
1500000
1000000
500000
0
4.09
16.44
1.12
1.00
1.08
1.08
1.16
2.06
1.11
8.6
8.4
8.2
8.0
7.8
7.6
7.4
7.2
7.0
6.8
6.6
6.4
6.2
6.0
5.8
5.6
5.4
5.2
5.0
4.8
4.6
4.4
4.2
4.0
3.8
3.6
3.4
f1 (ppm)

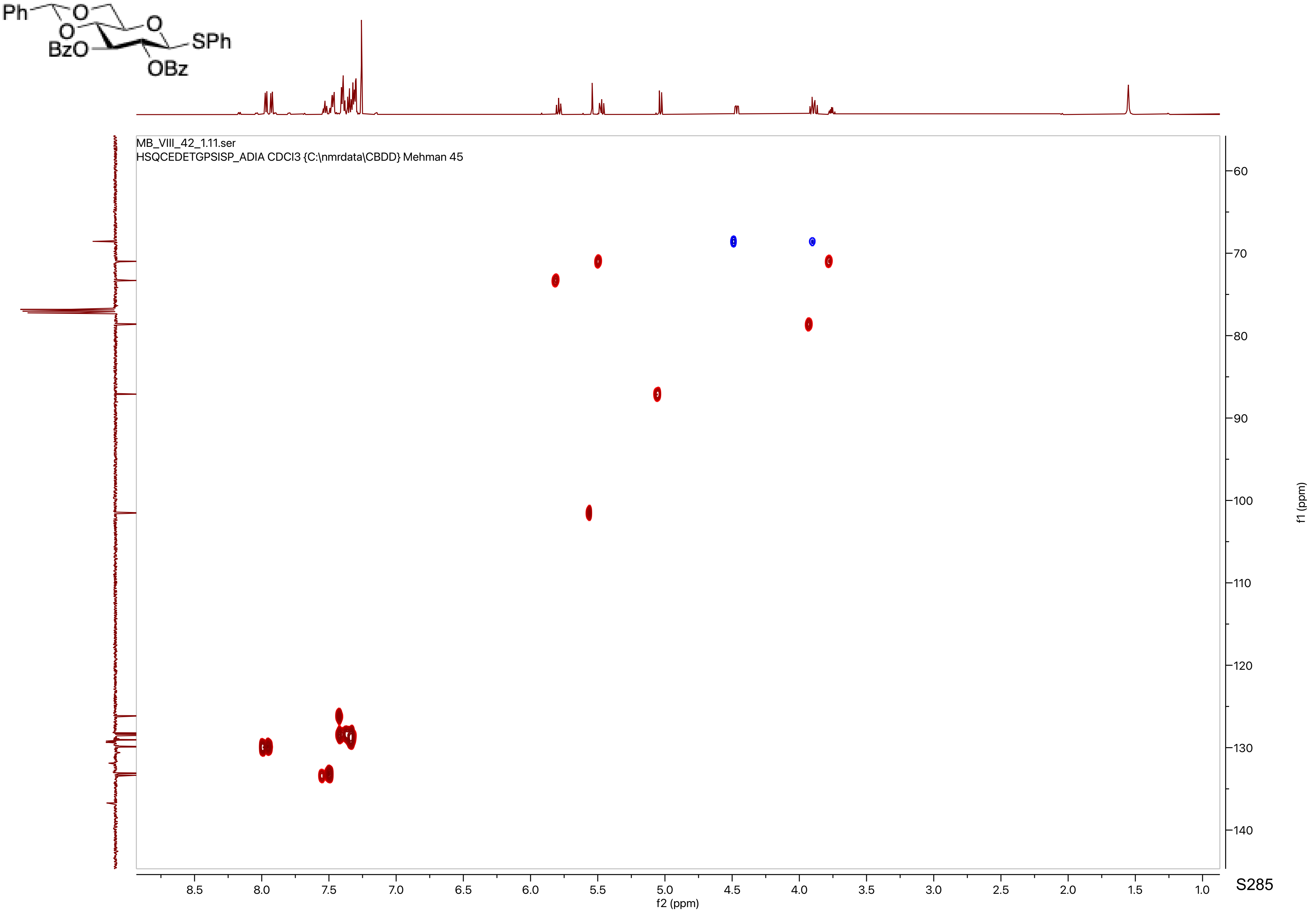
Ph
O
O
O
BzO
SPh
OBz
MB_VIII_42_1.11.ser
HSQCEDETGPSISP_ADIA CDCl3 {C:\nmrdata\CBDD} Mehman 45
f1 (ppm)
f2 (ppm)
8.5
8.0
7.5
7.0
6.5
6.0
5.5
5.0
4.5
4.0
3.5
3.0
2.5
2.0
1.5
1.0
60
70
80
90
100
110
120
130
140

MB_VIII_42_1.13.fid
C13APT CDCl3 {C:\nmrdata\CBDD} Mehman 45

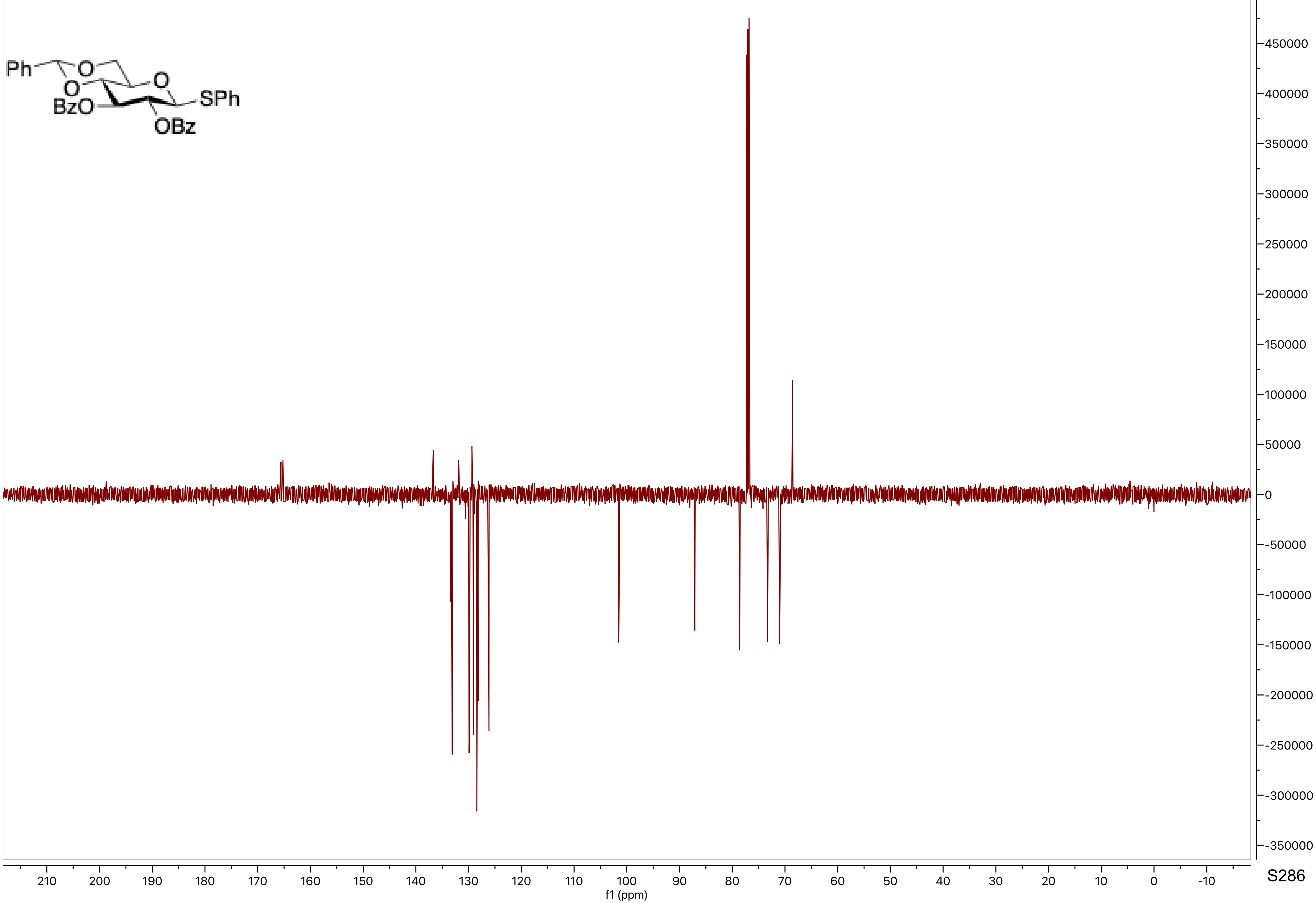

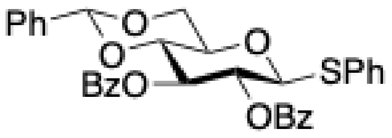
Ph
O
O
O
BzO
SPh
OBz

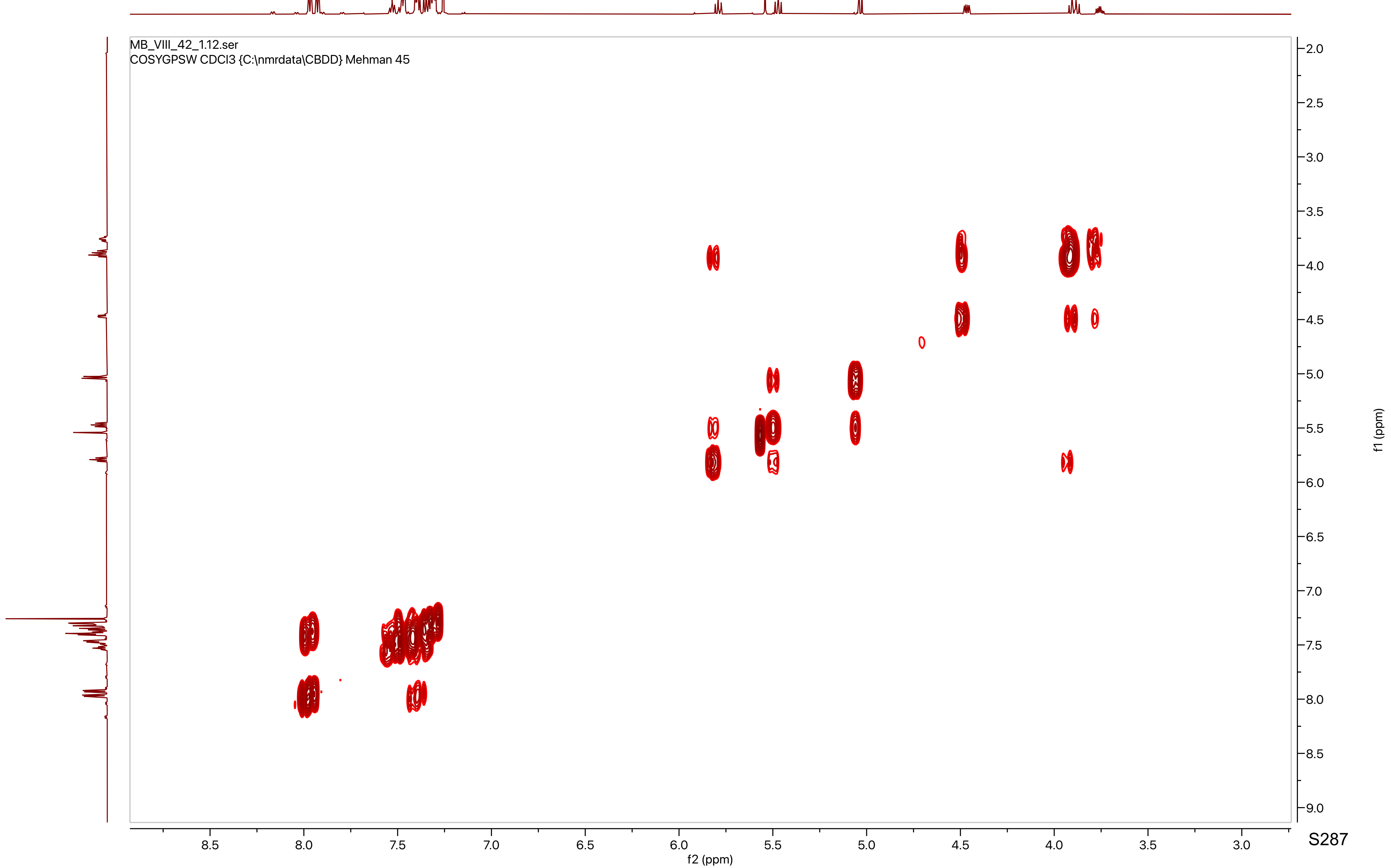
MB_VIII_42_1.12.ser
COSYGPSW CDCl3 {C:\nmrdata\CBDD} Mehman 45
8.5
8.0
7.5
7.0
6.5
6.0
5.5
5.0
4.5
4.0
3.5
3.0
f2 (ppm)
2.0
2.5
3.0
3.5
4.0
4.5
5.0
5.5
6.0
6.5
7.0
7.5
8.0
8.5
9.0
f1 (ppm)

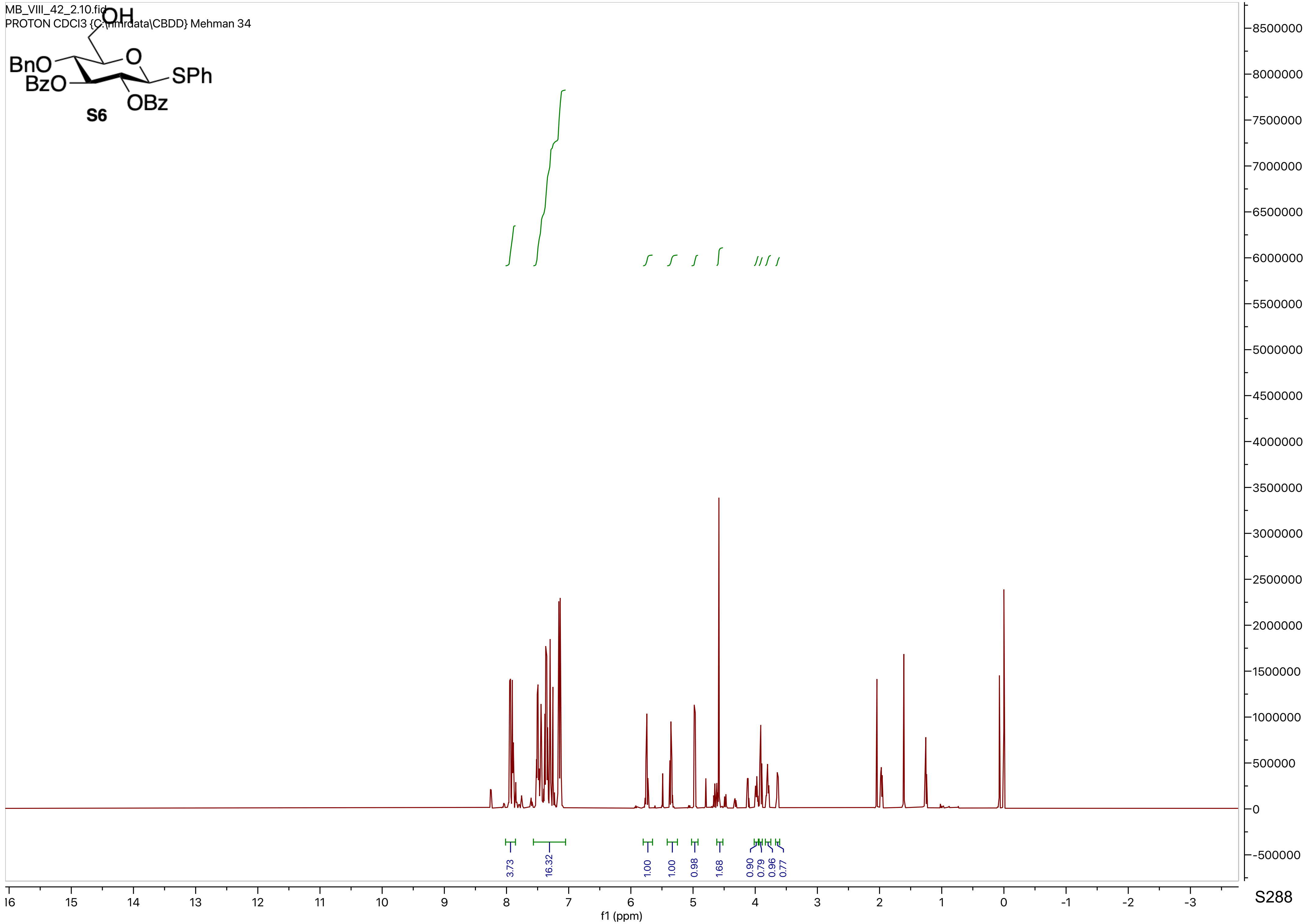
MB_VIII_42_2.10.fid
PROTON CDCl3 {C:\nmrdata\CBDD} Mehman 34
OH
BnO
BzO
O
SPh
OBz
S6
3.73
16.32
1.00
1.00
0.98
1.68
0.90
0.79
0.96
0.77
f1 (ppm)

MB_VIII_42_2.13.fid
C13APT CDCl3 {C:\nmrdata\CBDD} Mehman 34

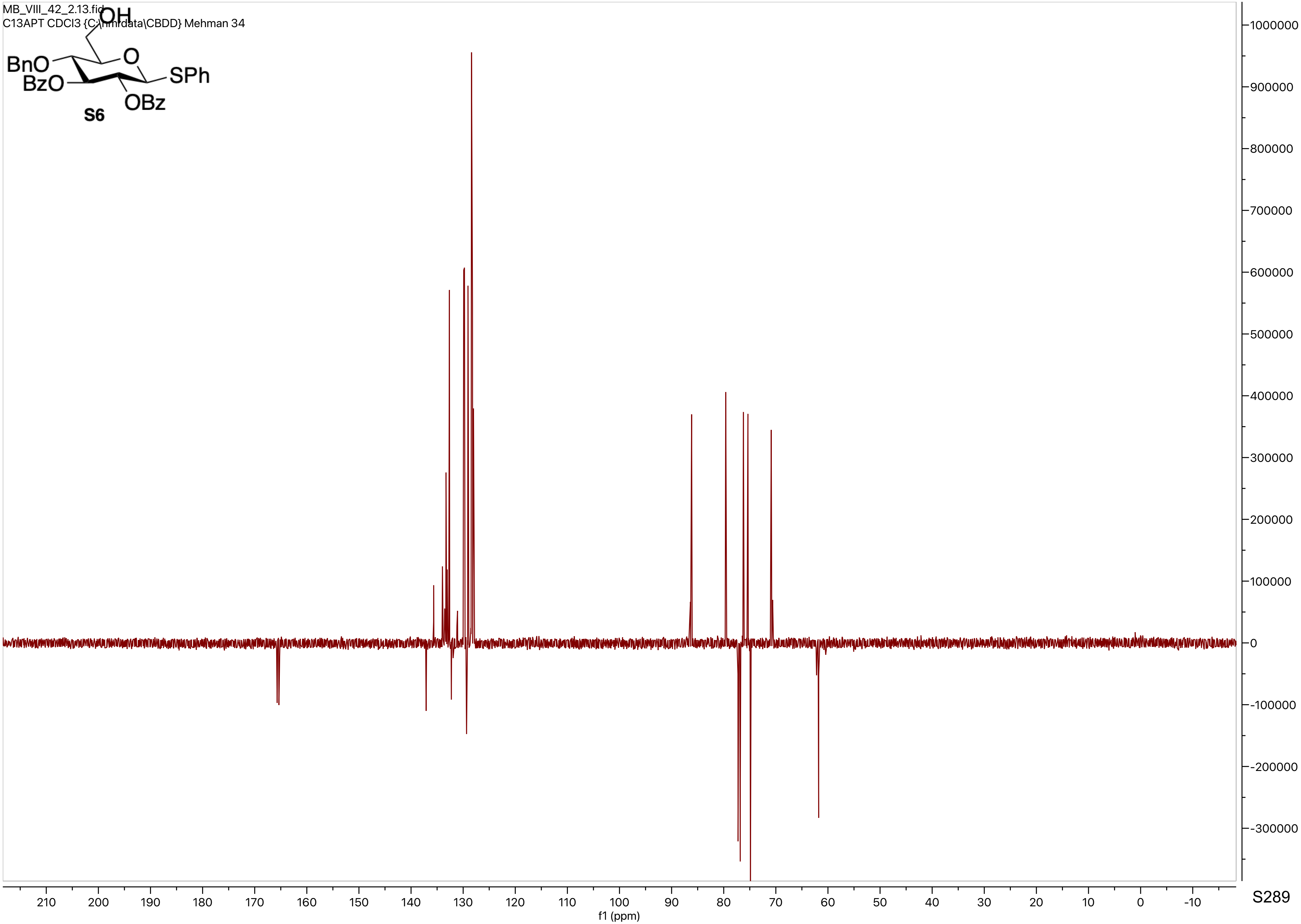

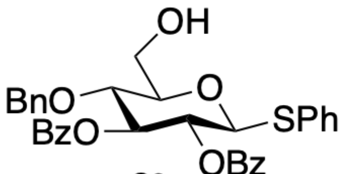
BnO
BzO
OH
O
SPh
OBz

**S6**

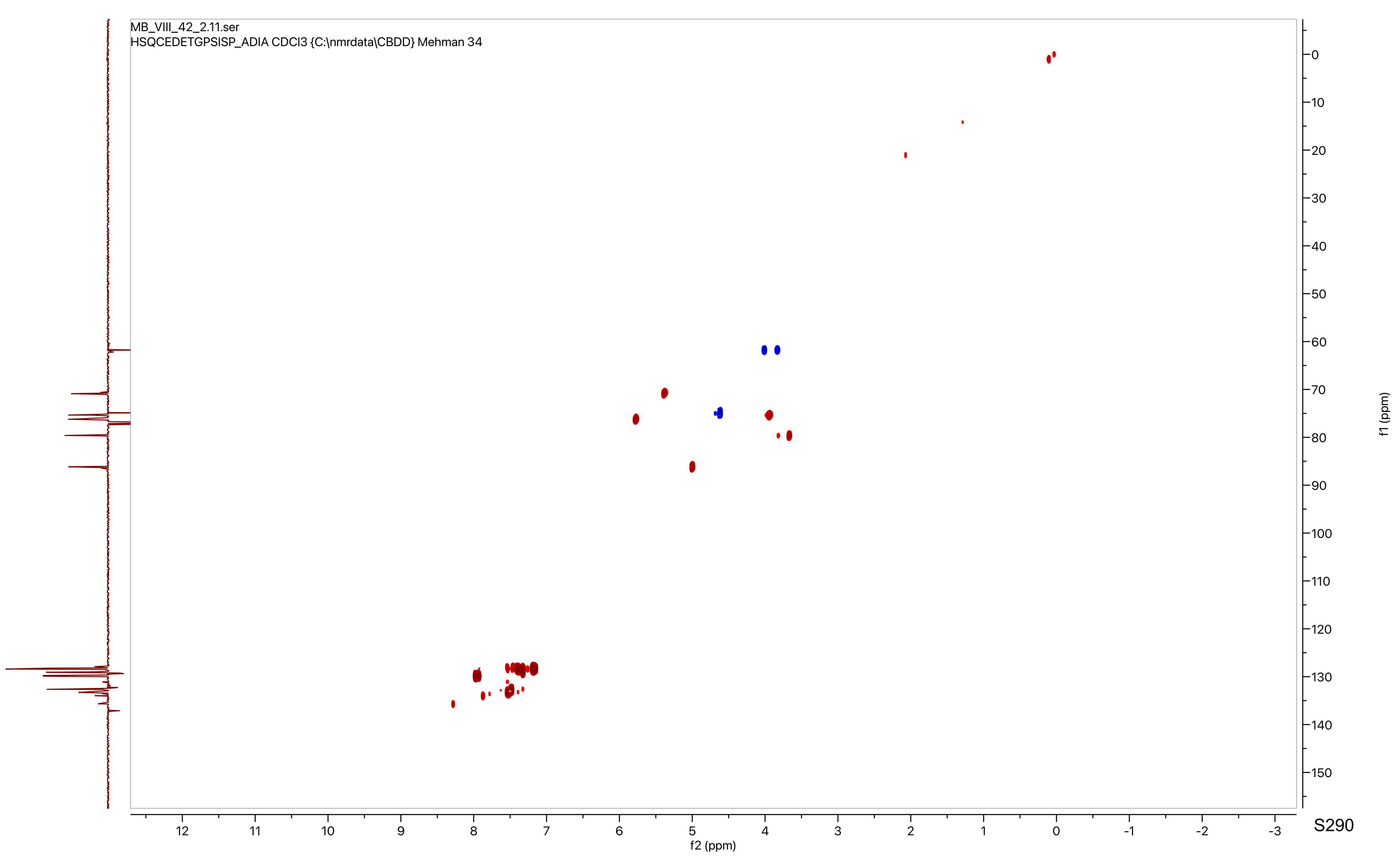
MB_VIII_42_2.11.ser
HSQCEDETGPSISP_ADIA CDCl3 {C:\nmrdata\CBDD} Mehman 34
f2 (ppm)
f1 (ppm)

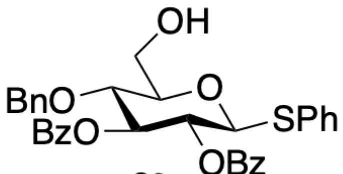

OH
O
BnO
BzO
SPh
OBz


**S6**

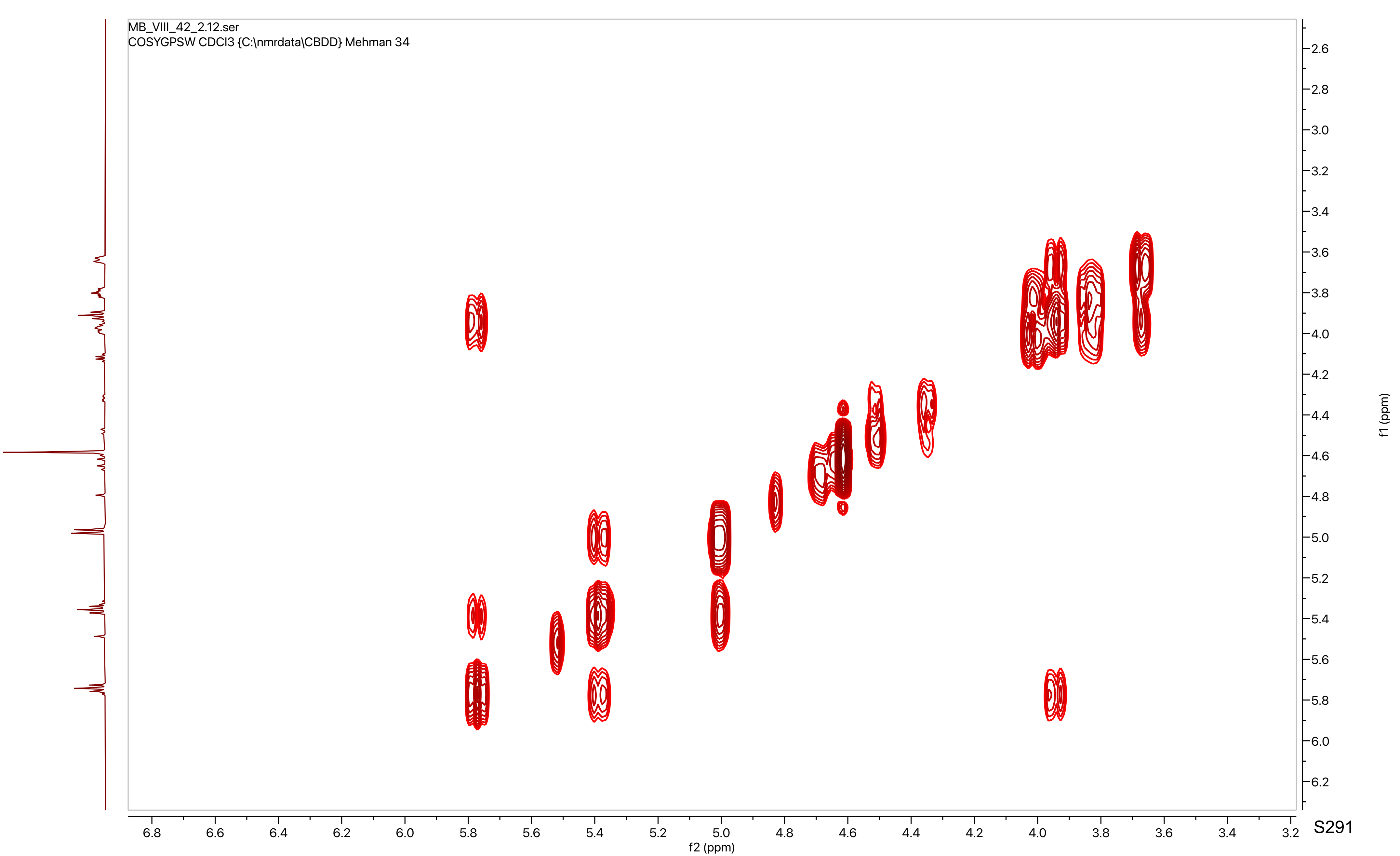

MB_VIII_42_2.12.ser
COSYGPSW CDCl3 {C:\nmrdata\CBDD} Mehman 34
f2 (ppm)
f1 (ppm)
6.8 6.6 6.4 6.2 6.0 5.8 5.6 5.4 5.2 5.0 4.8 4.6 4.4 4.2 4.0 3.8 3.6 3.4 3.2
2.6 2.8 3.0 3.2 3.4 3.6 3.8 4.0 4.2 4.4 4.6 4.8 5.0 5.2 5.4 5.6 5.8 6.0 6.2

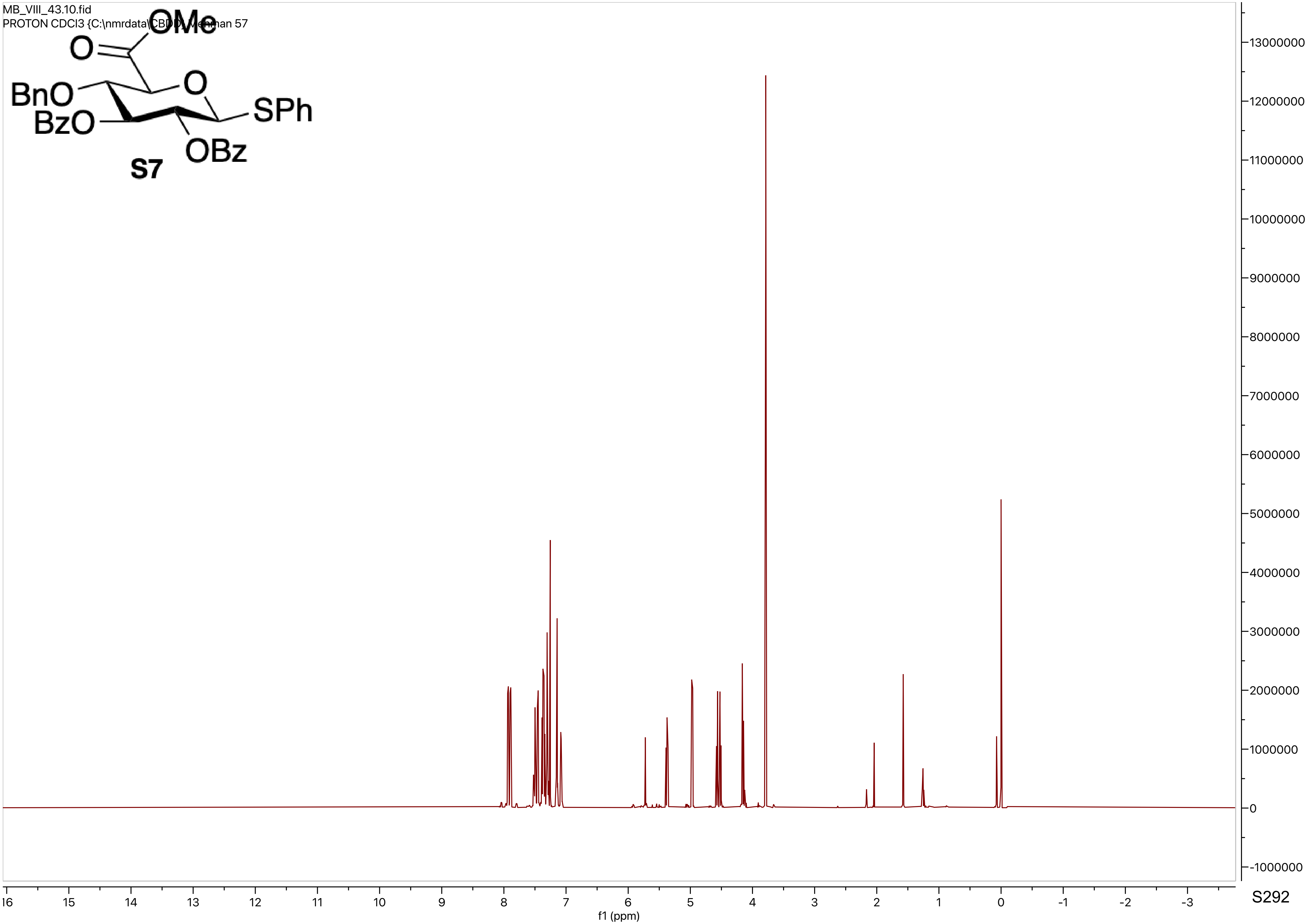
MB_VIII_43.10.fid
PROTON CDCl3 {C:\nmrdata\CBDD} Mehman 57
OMe
O
O
BnO
BzO
SPh
OBz
S7
f1 (ppm)

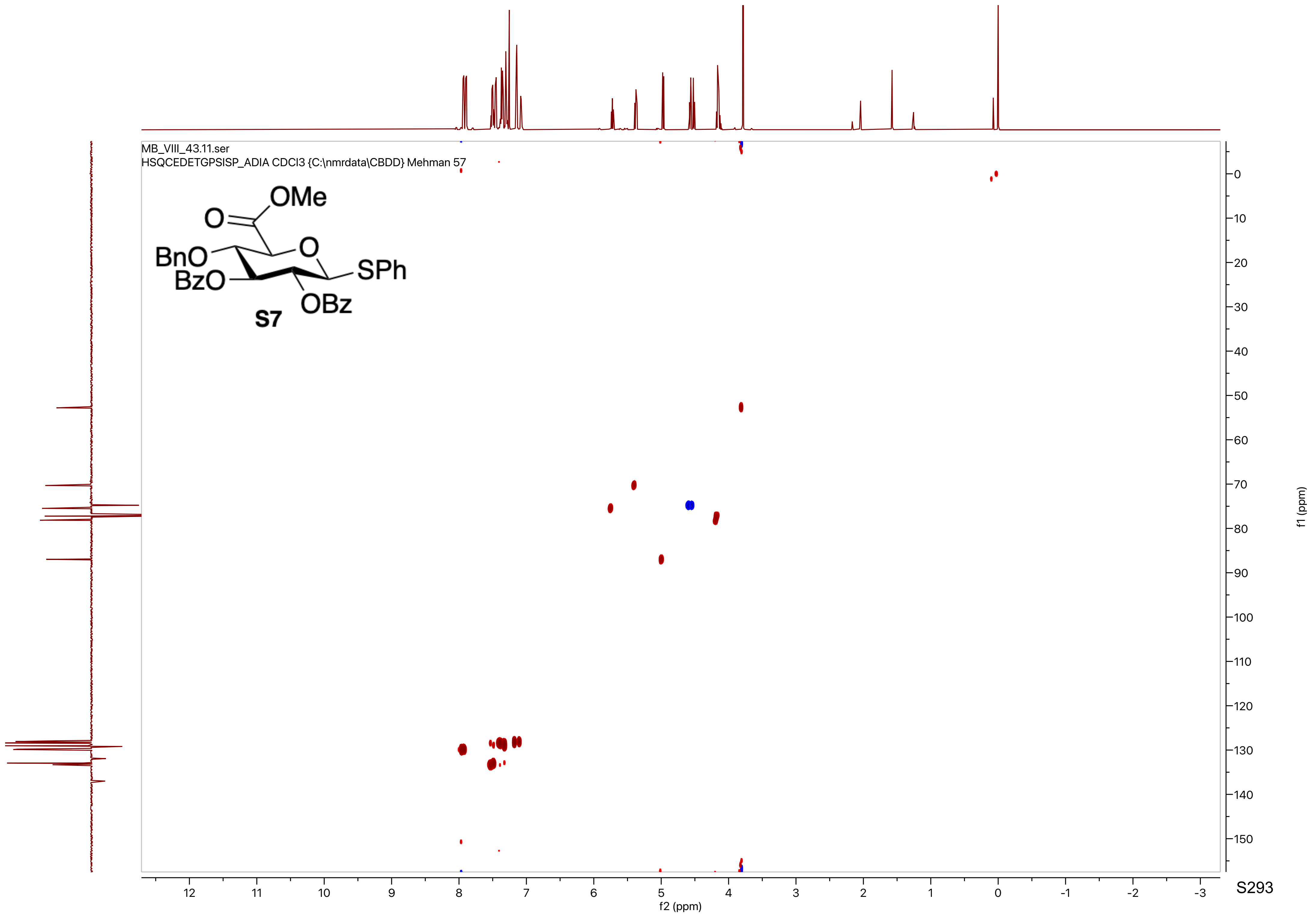
MB_VIII_43.11.ser
HSQCEDETGPSISP_ADIA CDCl3 {C:\nmrdata\CBDD} Mehman 57
OMe
O
O
BnO
BzO
SPh
OBz
S7
f2 (ppm)
f1 (ppm)

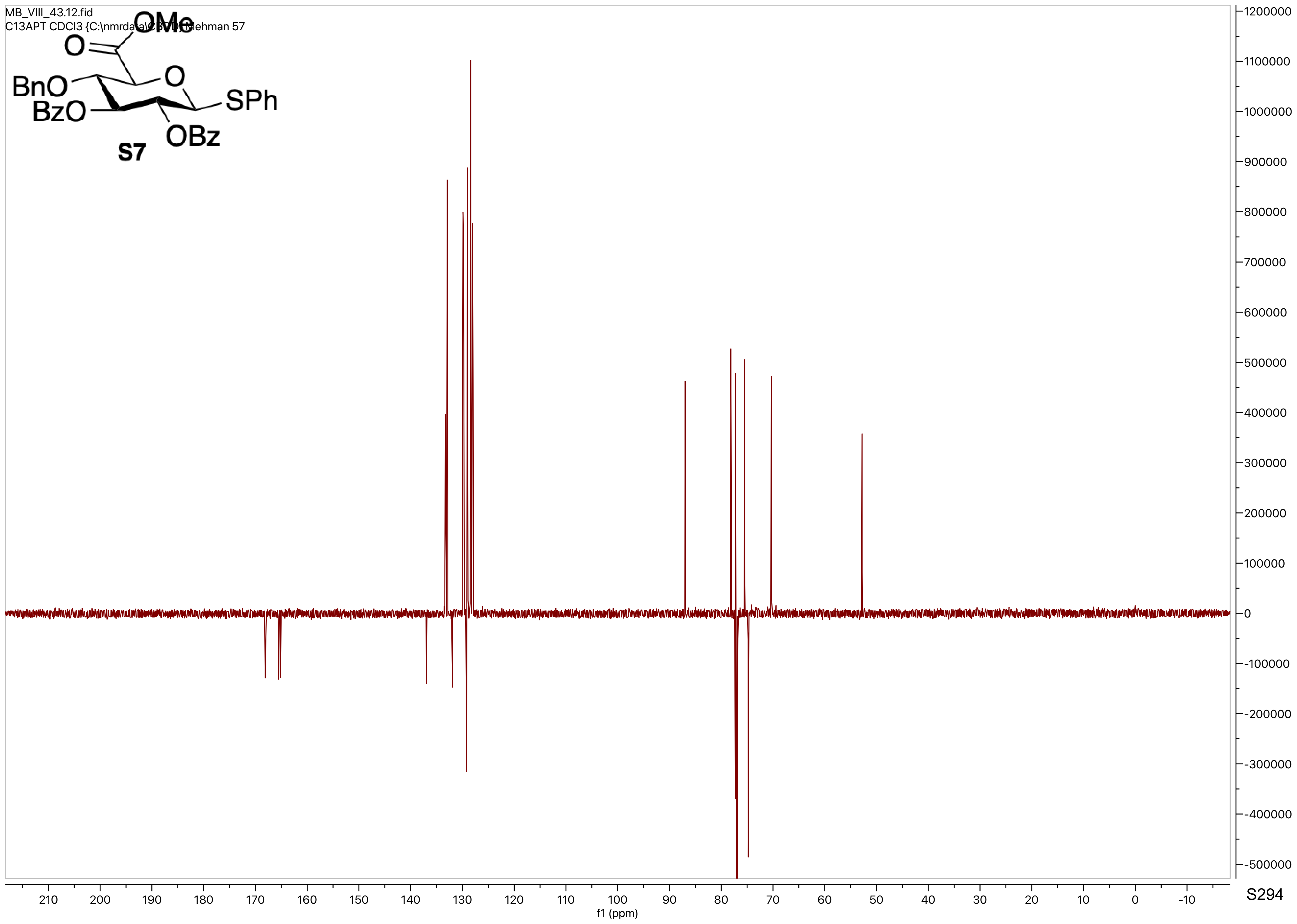

MB_VIII_43.12.fid
C13APT CDCl3 {C:\nmrdata\C3TD} Mehman 57
OMe
O
O
BnO
BzO
SPh
OBz
S7
f1 (ppm)

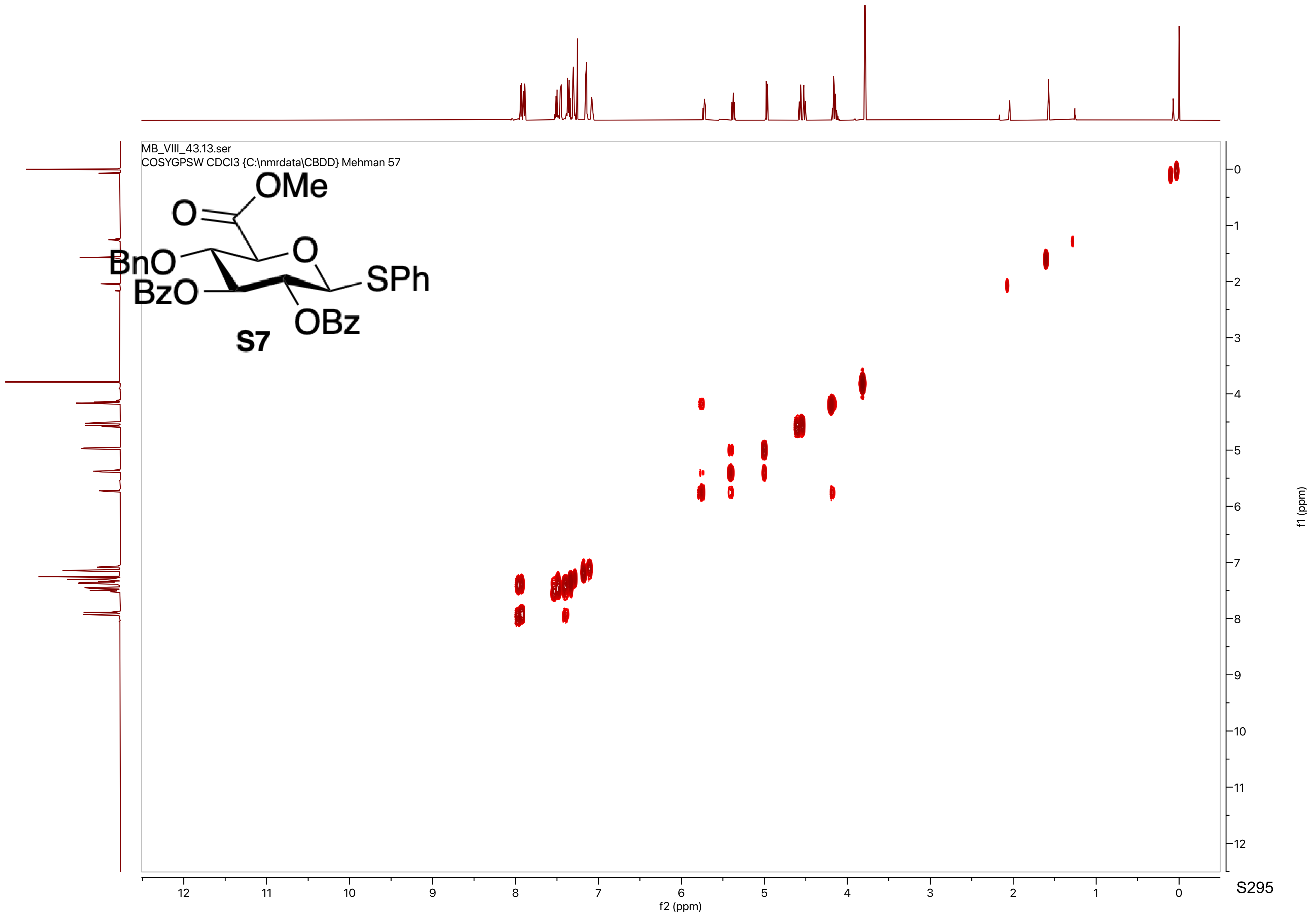
MB_VIII_43.13.ser
COSYGPSW CDCl3 {C:\nmrdata\CBDD} Mehman 57
OMe
O
O
BnO
BzO
SPh
OBz
S7
f1 (ppm)
f2 (ppm)

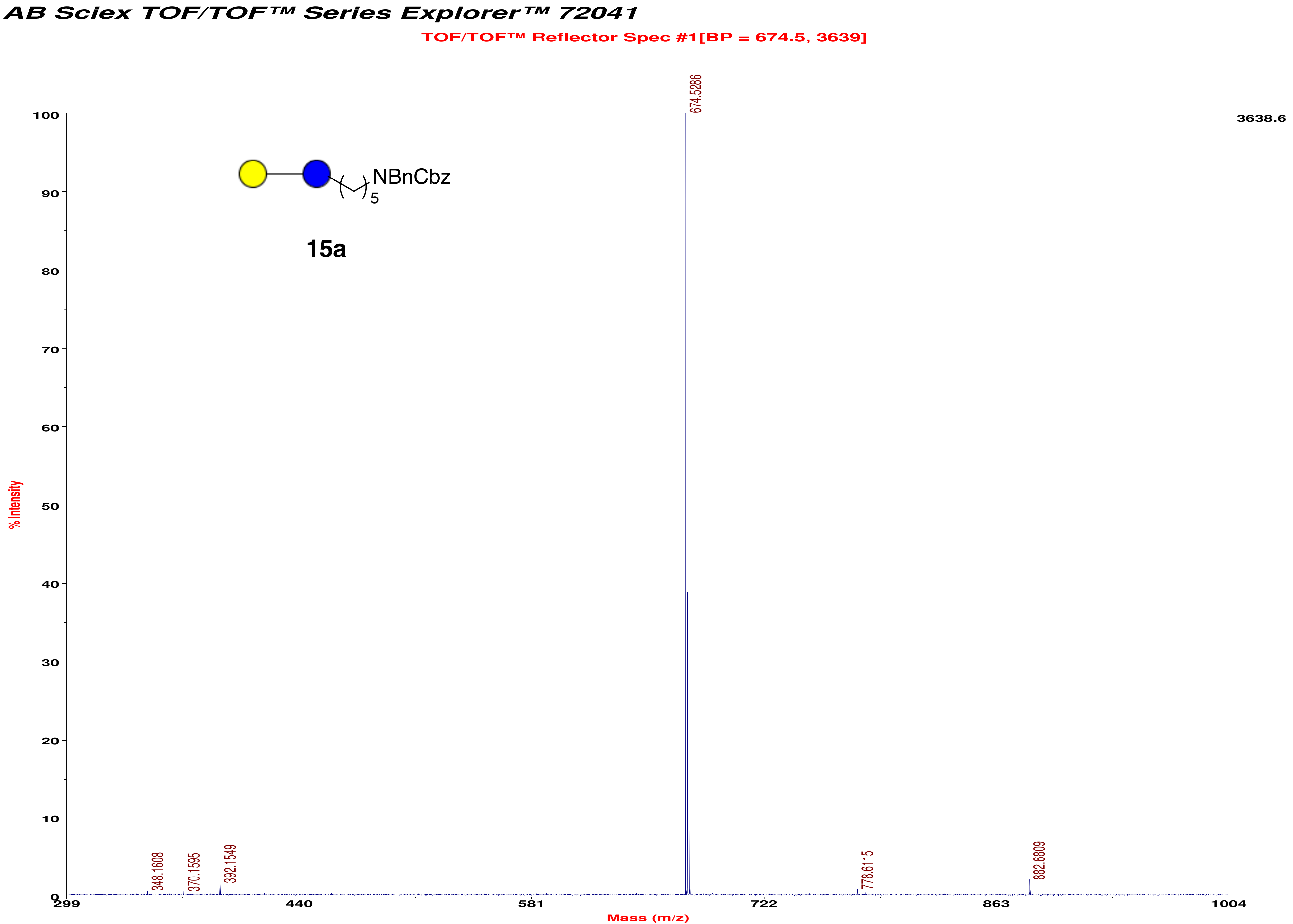
AB Sciex TOF/TOF™ Series Explorer™ 72041
TOF/TOF™ Reflector Spec #1[BP = 674.5, 3639]
NBnCbz
5
15a
674.5286
3638.6
348.1608
370.1595
392.1549
778.6115
882.6809
% Intensity
Mass (m/z)
299
440
581
722
863
1004

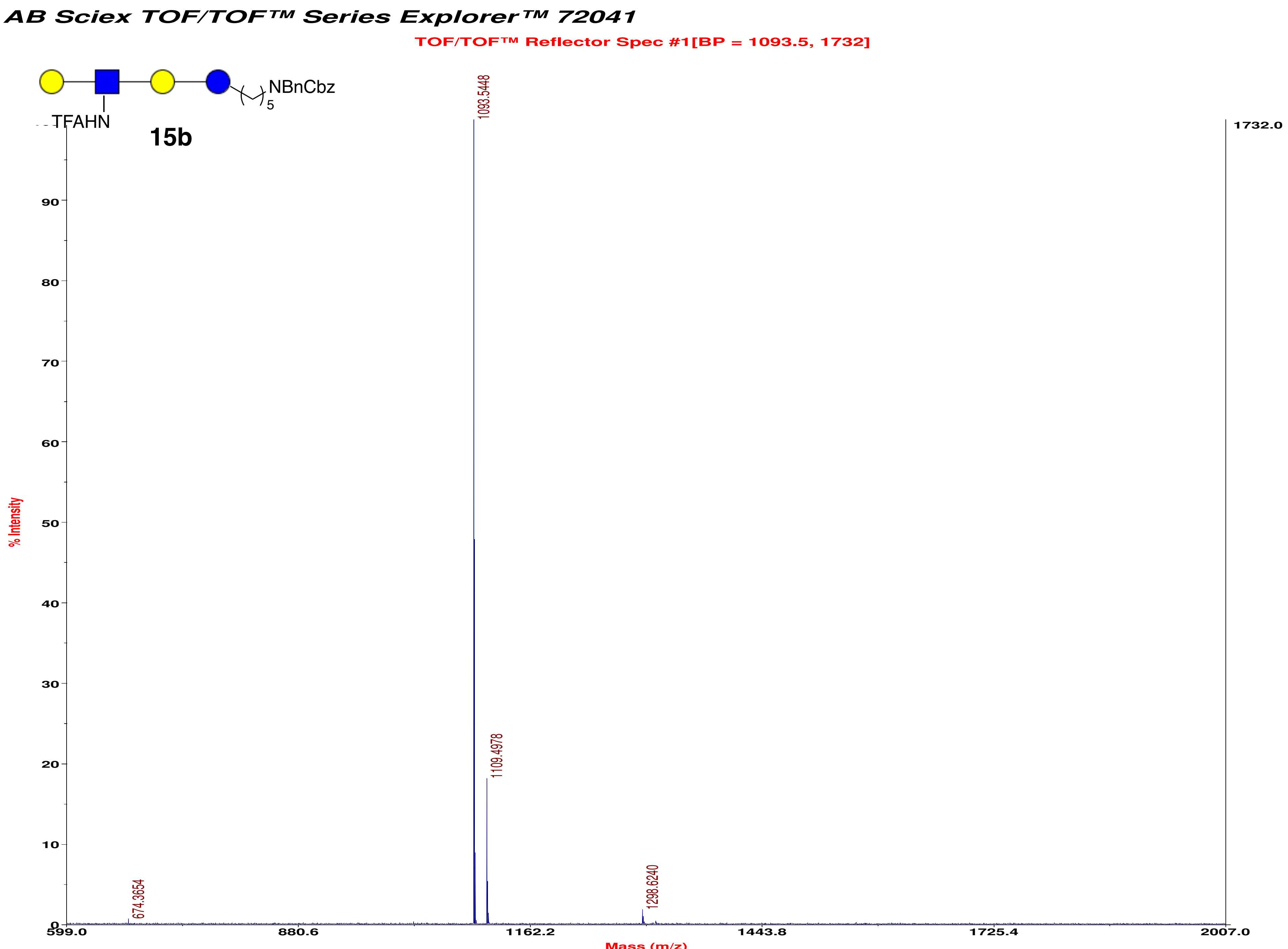
AB Sciex TOF/TOF™ Series Explorer™ 72041
TOF/TOF™ Reflector Spec #1[BP = 1093.5, 1732]
NBnCbz
5
TFAHN
15b
1093.5448
1109.4978
674.3654
1298.6240
1732.0
% Intensity
Mass (m/z)
599.0
880.6
1162.2
1443.8
1725.4
2007.0
0
10
20
30
40
50
60
70
80
90

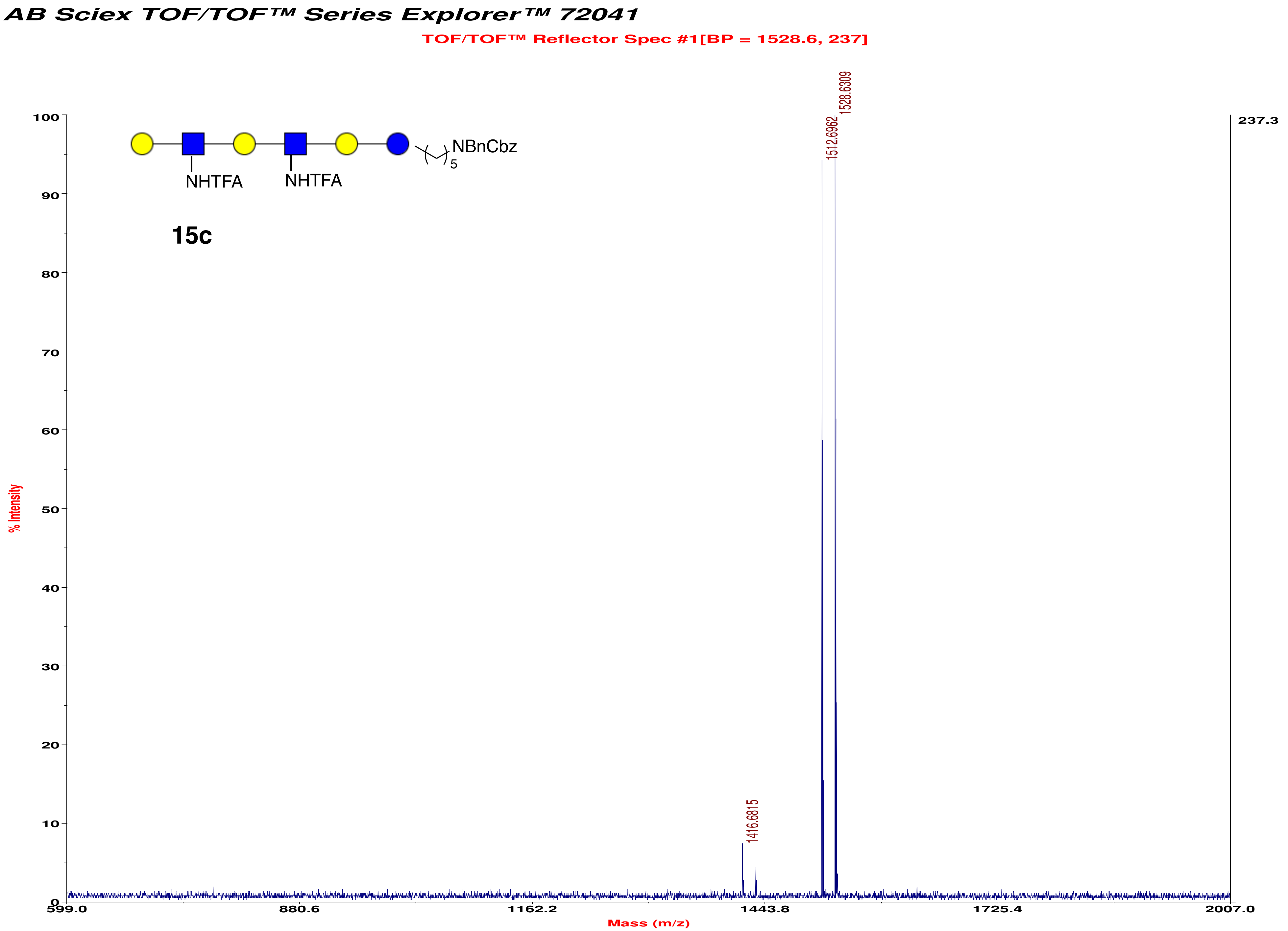
AB Sciex TOF/TOF™ Series Explorer™ 72041
TOF/TOF™ Reflector Spec #1[BP = 1528.6, 237]
NBnCbz
5
NHTFA
NHTFA
15c
1528.6309
1512.6962
1416.6815
237.3
% Intensity
Mass (m/z)
599.0
880.6
1162.2
1443.8
1725.4
2007.0
0
10
20
30
40
50
60
70
80
90
100

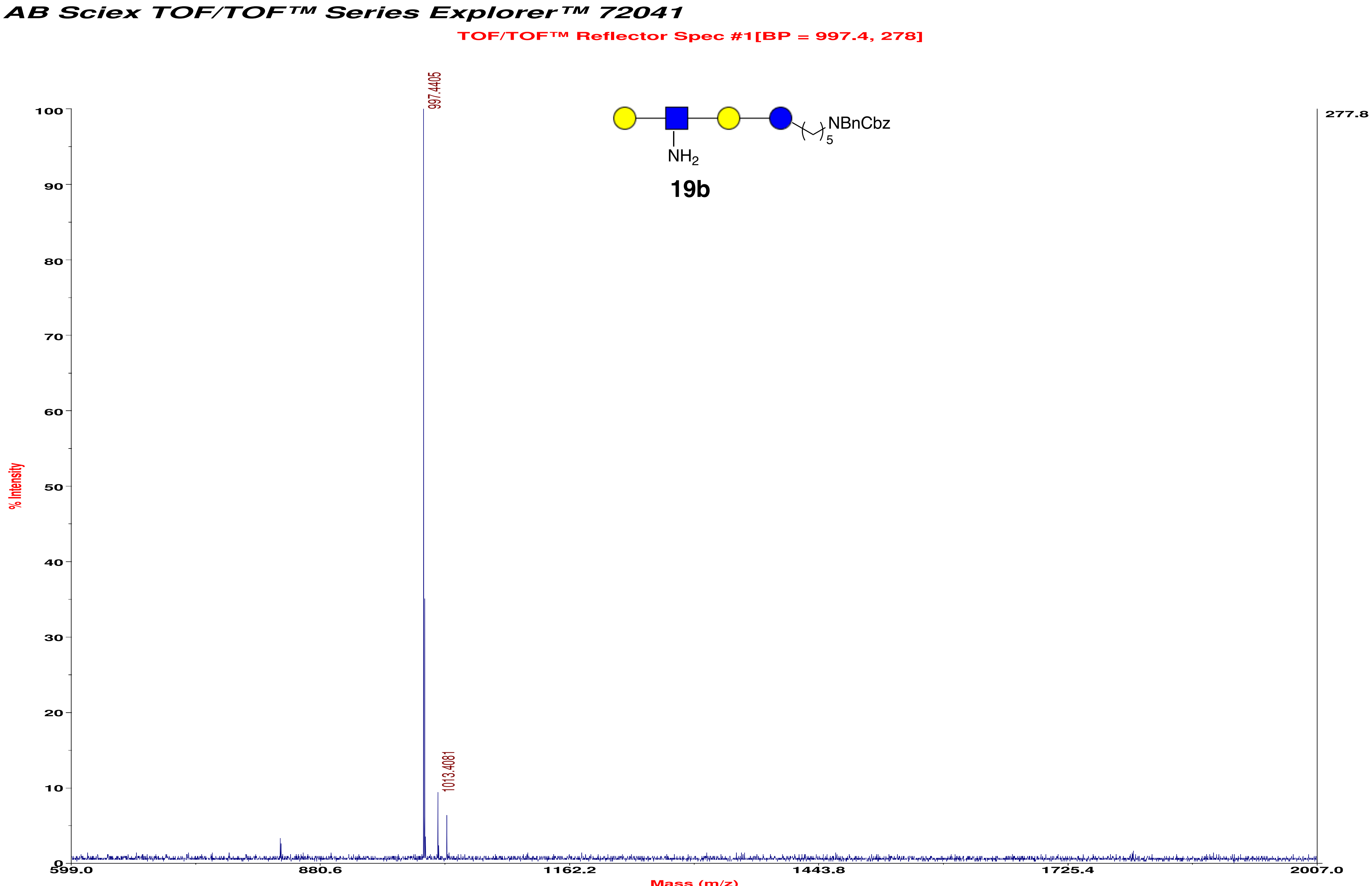

AB Sciex TOF/TOF™ Series Explorer™ 72041
TOF/TOF™ Reflector Spec #1[BP = 997.4, 278]
997.4405
1013.4081
NBnCbz
5
NH2
19b
277.8
% Intensity
100
90
80
70
60
50
40
30
20
10
0
599.0
880.6
1162.2
1443.8
1725.4
2007.0
Mass (m/z)

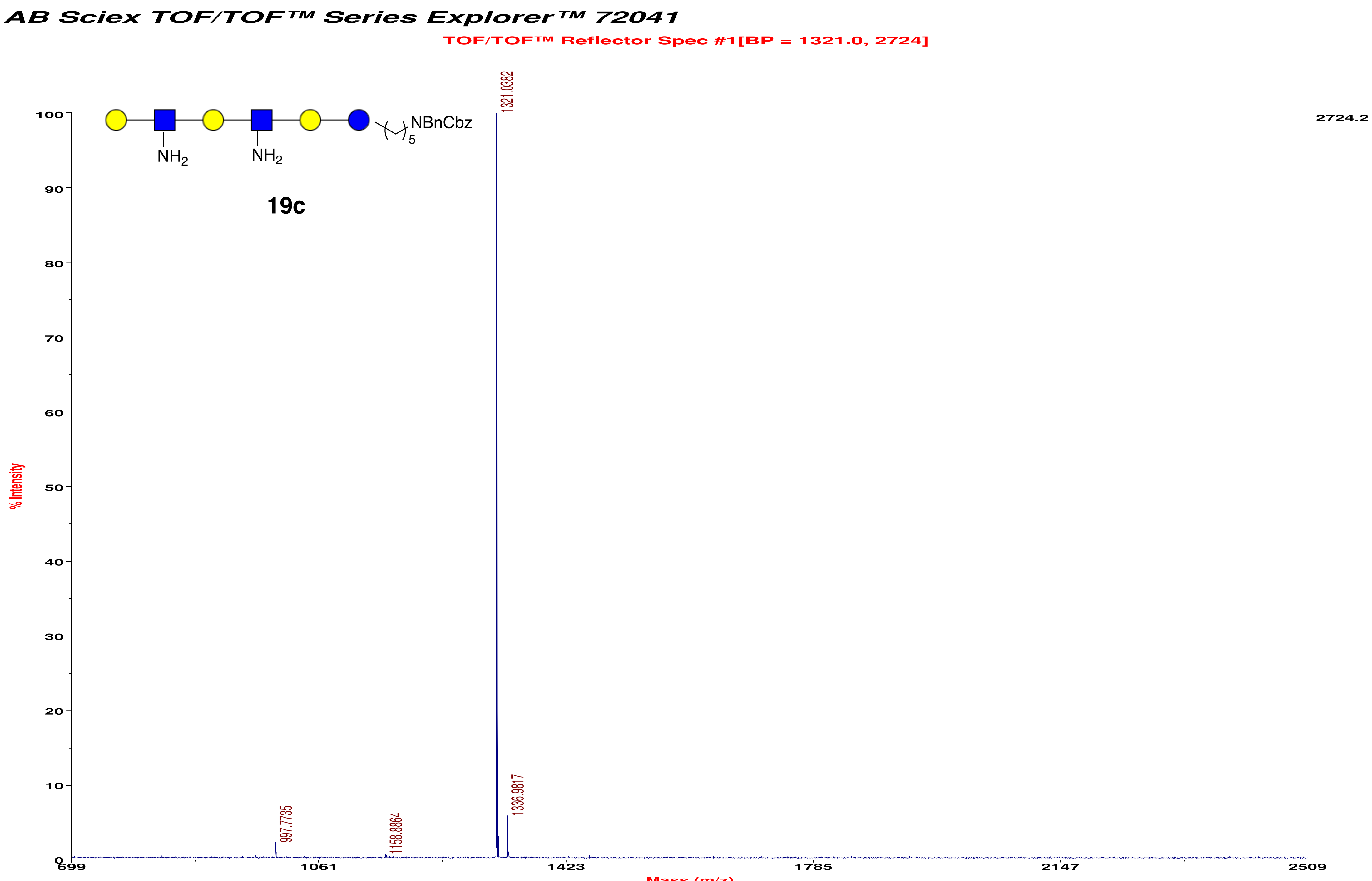

AB Sciex TOF/TOF™ Series Explorer™ 72041
TOF/TOF™ Reflector Spec #1[BP = 1321.0, 2724]
NBnCbz
5
NH2
NH2
19c
1321.0382
1336.9817
997.7735
1158.8864
2724.2
% Intensity
Mass (m/z)

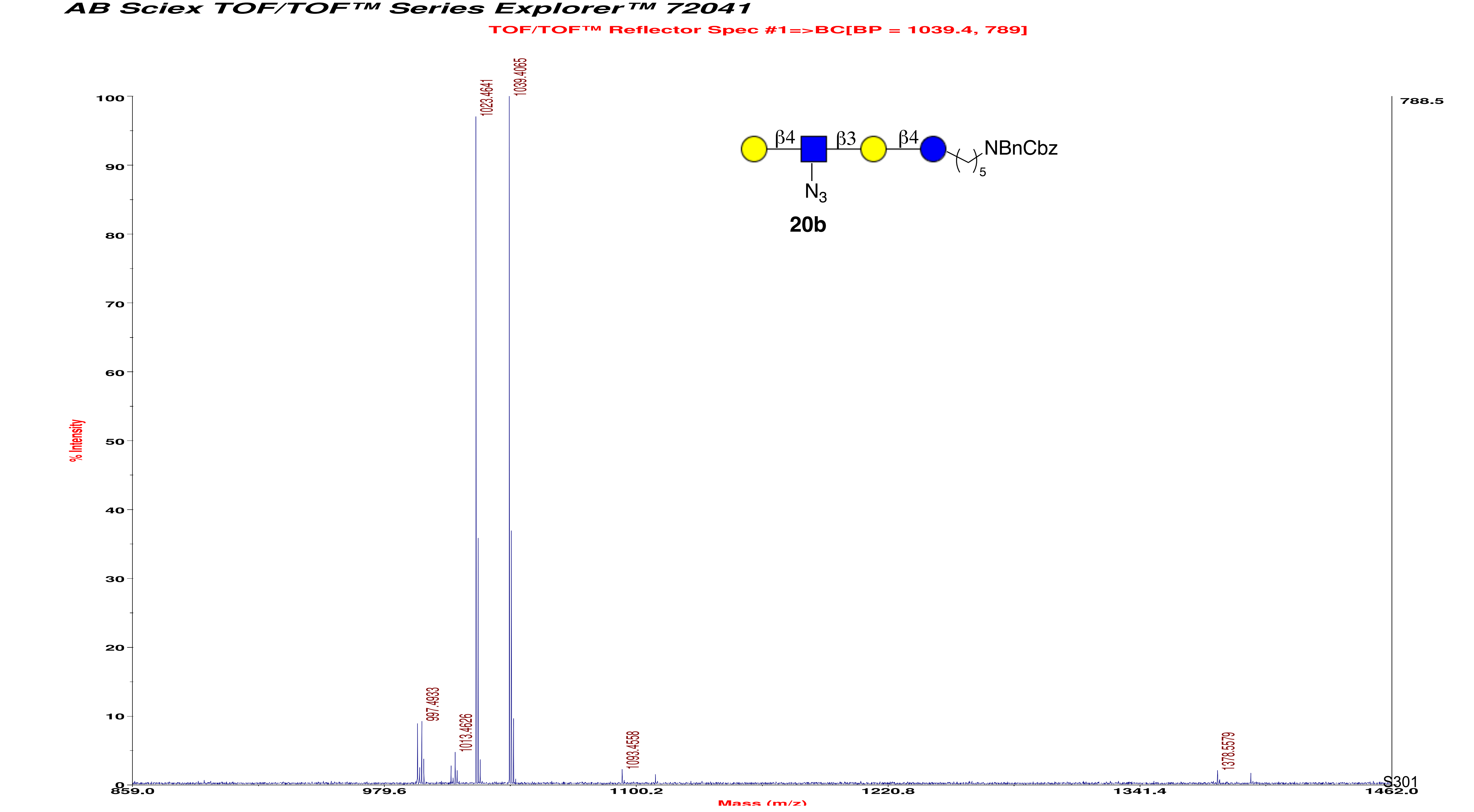
AB Sciex TOF/TOF™ Series Explorer™ 72041
TOF/TOF™ Reflector Spec #1=>BC[BP = 1039.4, 789]
1039.4065
1023.4641
997.4933
1013.4626
1093.4558
1378.5579
788.5
% Intensity
Mass (m/z)
β4
β3
β4
NBnCbz
N3
5
20b
S301

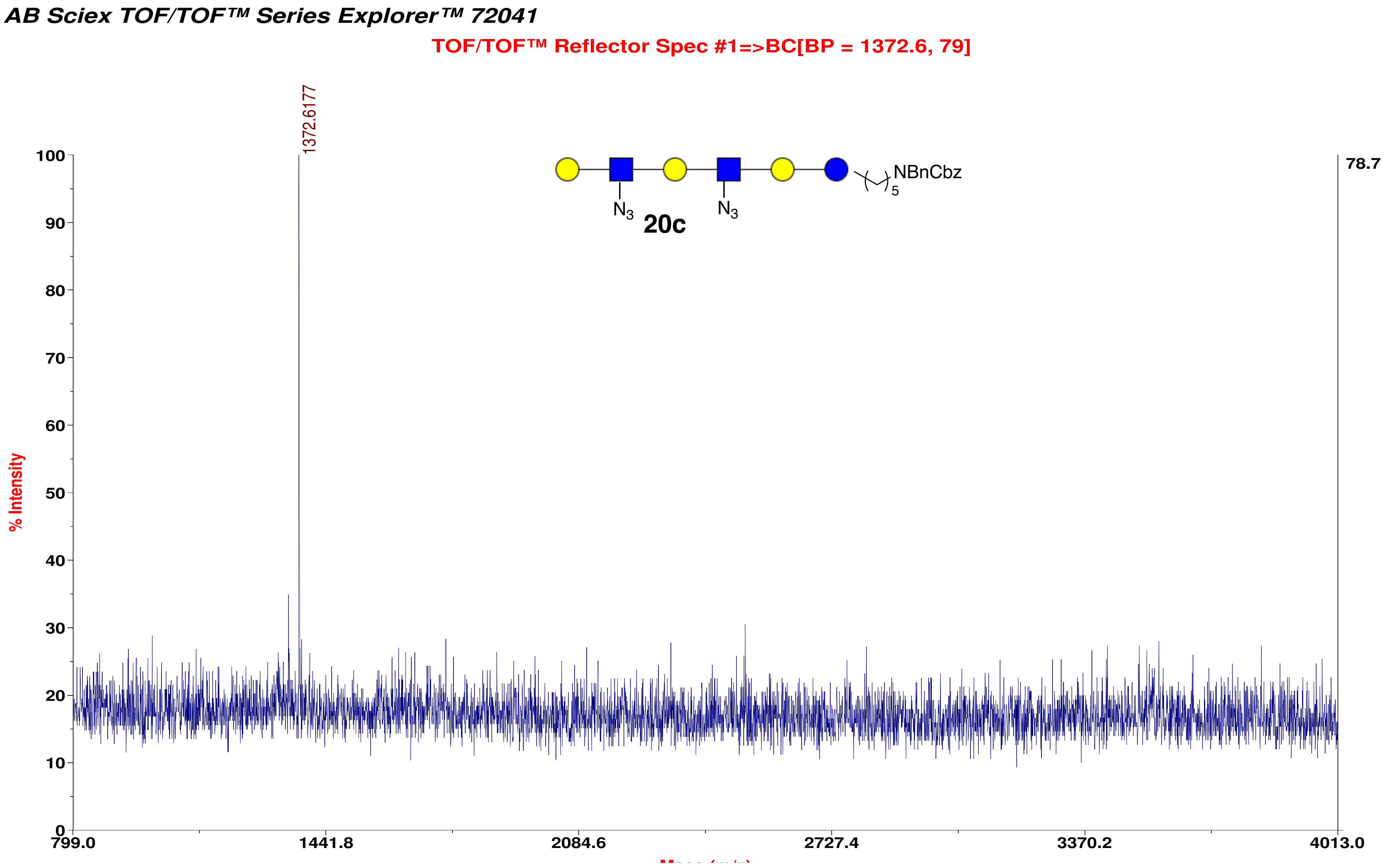

AB Sciex TOF/TOF™ Series Explorer™ 72041
TOF/TOF™ Reflector Spec #1=>BC[BP = 1372.6, 79]
1372.6177
NBnCbz
5
N3
N3
20c
78.7
% Intensity
100
90
80
70
60
50
40
30
20
10
0
799.0
1441.8
2084.6
2727.4
3370.2
4013.0

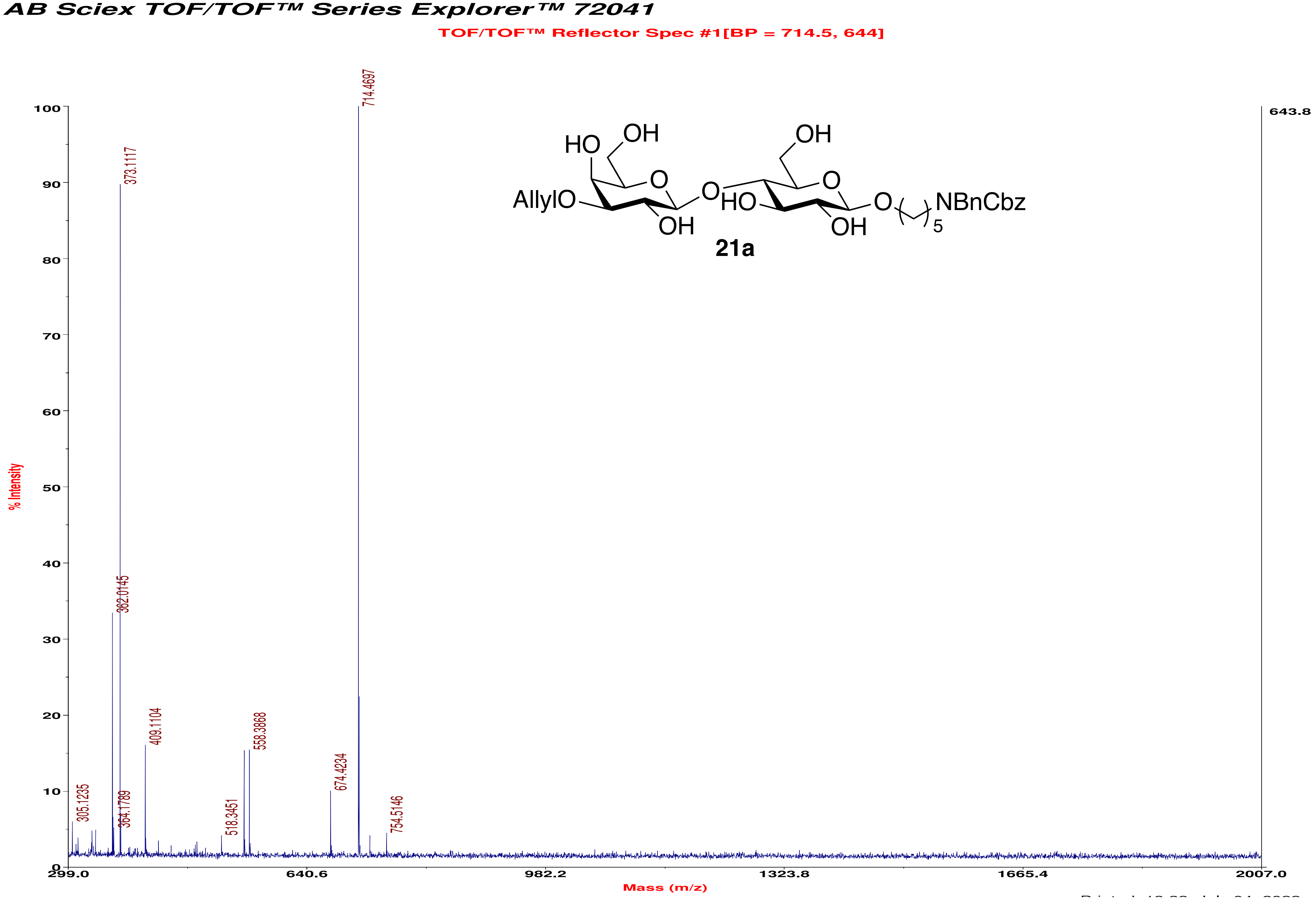
AB Sciex TOF/TOF™ Series Explorer™ 72041
TOF/TOF™ Reflector Spec #1[BP = 714.5, 644]
% Intensity
Mass (m/z)
643.8
714.4697
373.1117
362.0145
409.1104
558.3868
674.4234
305.1235
364.1789
518.3451
754.5146
AllylO
HO
OH
O
HO
OH
O
OH
NBnCbz
5
21a

### AB Sciex TOF/TOF™ Series Explorer™ 72041

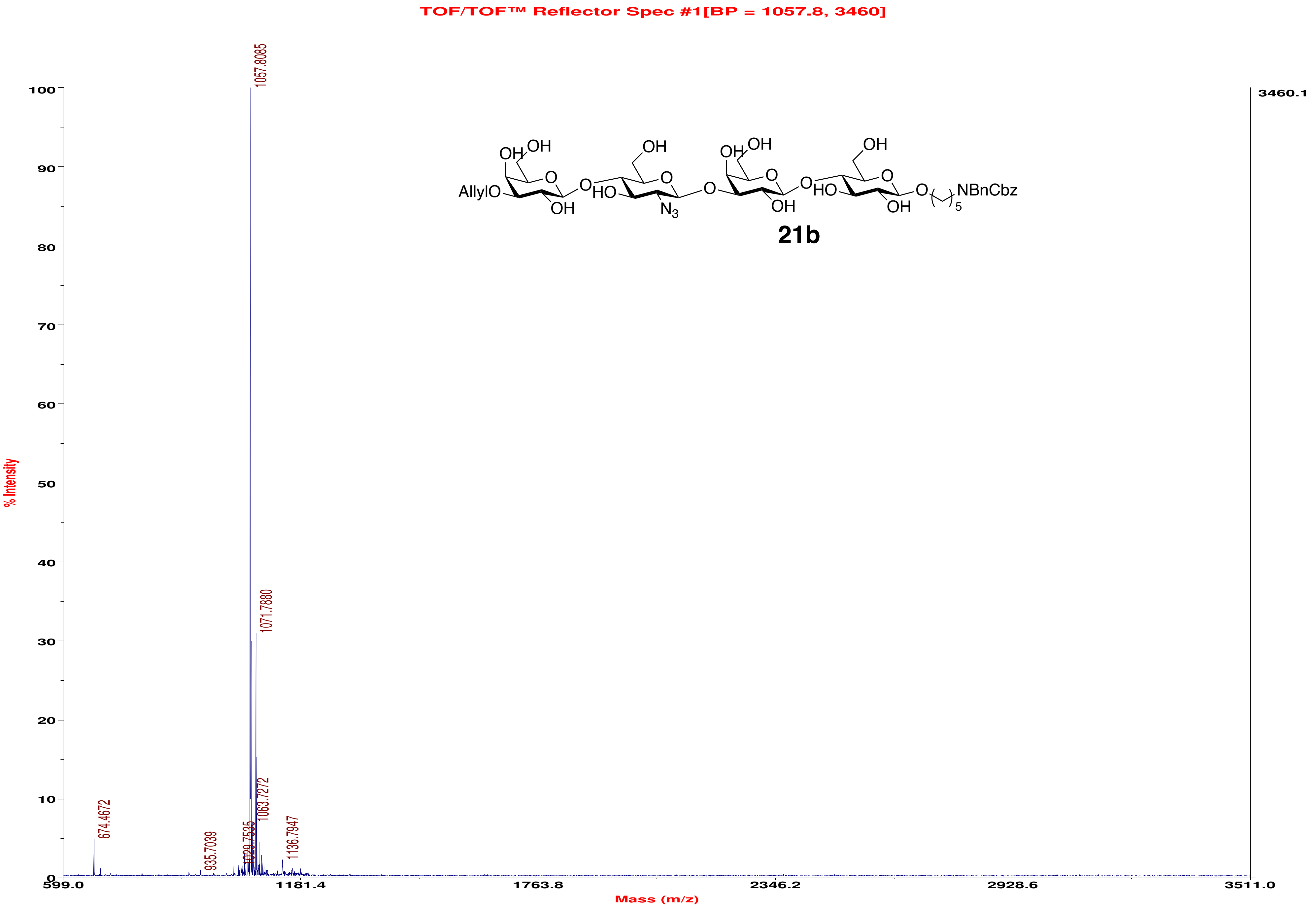

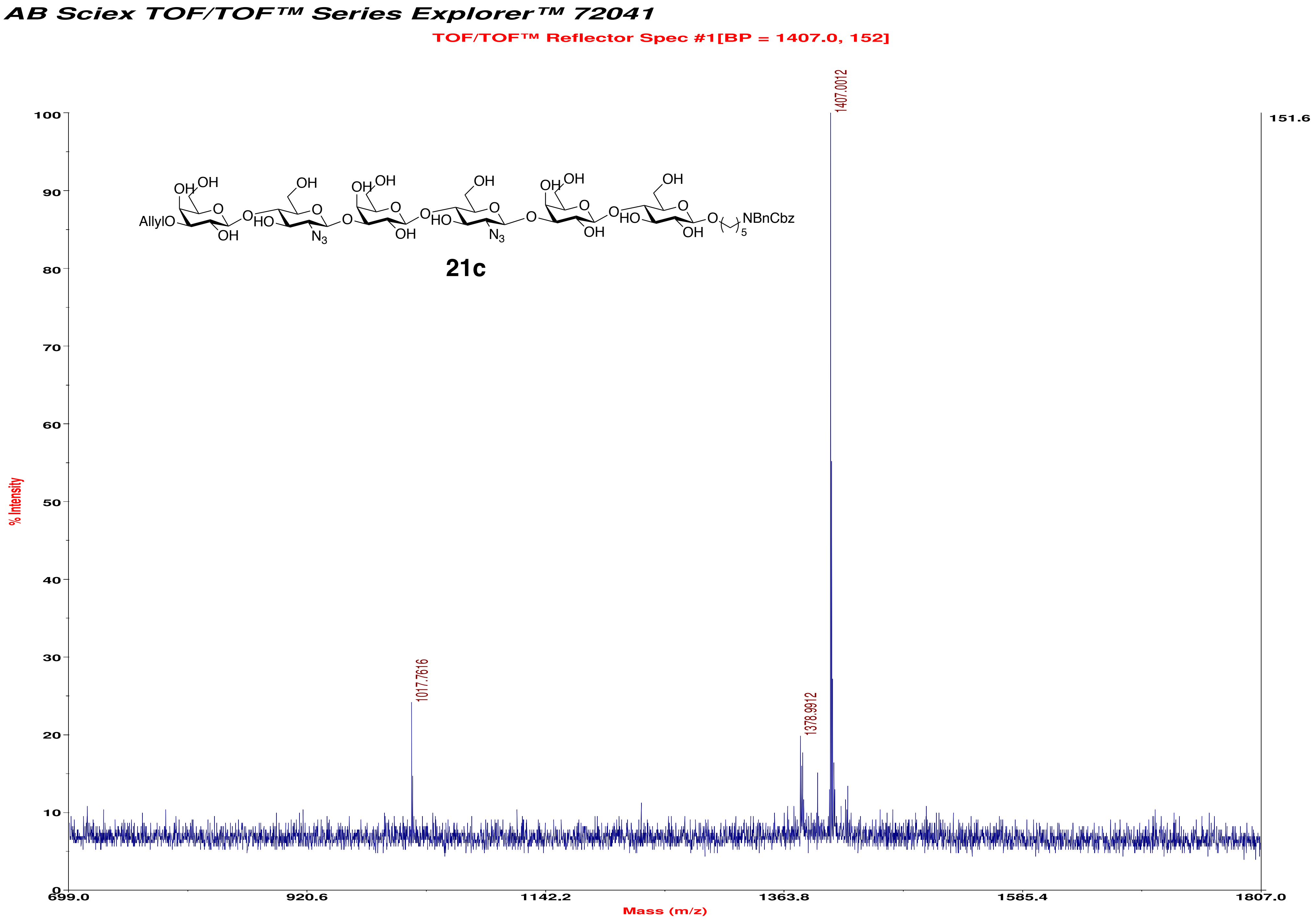
AB Sciex TOF/TOF™ Series Explorer™ 72041
TOF/TOF™ Reflector Spec #1[BP = 1407.0, 152]
% Intensity
Mass (m/z)
1407.0012
1378.9912
1017.7616
151.6
AllylO
NBnCbz
21c

AB Sciex TOF/TOF™ Series Explorer™ 72041

TOF/TOF™ Reflector Spec #1[BP = 1254.8, 1880]

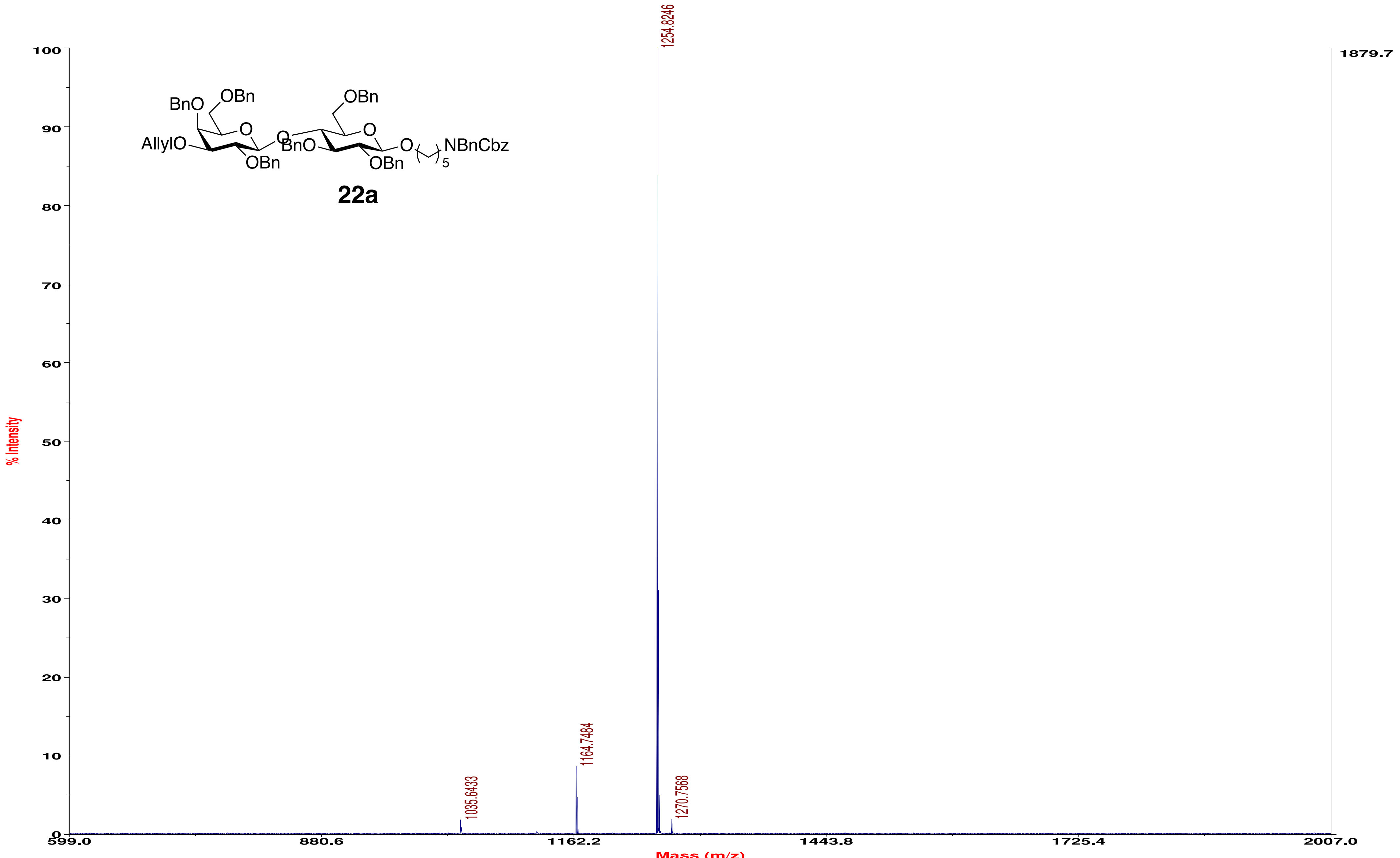

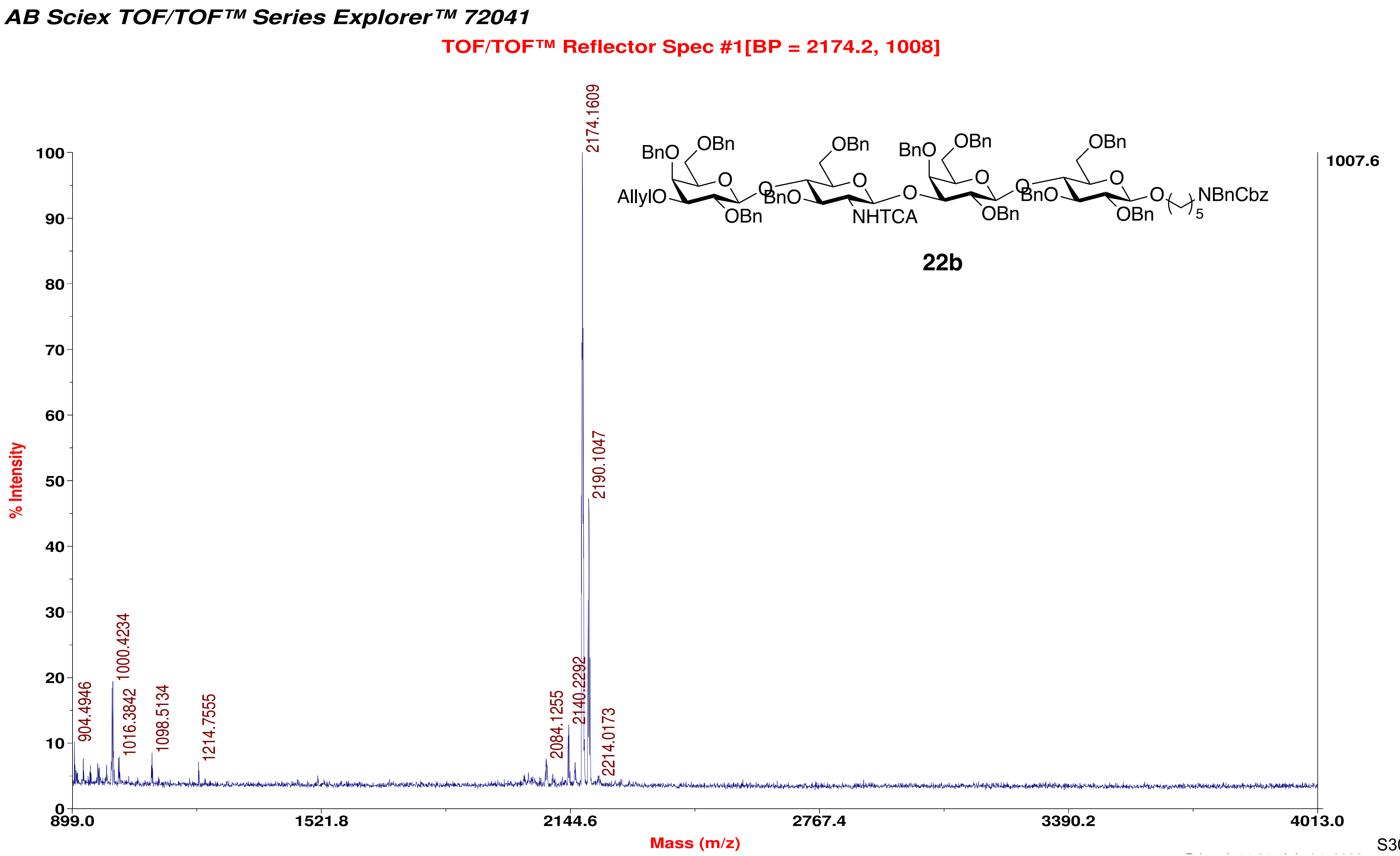
AB Sciex TOF/TOF™ Series Explorer™ 72041
TOF/TOF™ Reflector Spec #1[BP = 2174.2, 1008]
% Intensity
Mass (m/z)
1007.6
2174.1609
2190.1047
1000.4234
904.4946
1016.3842
1098.5134
1214.7555
2084.1255
2140.2292
2214.0173
BnO
OBn
AllylO
O
BnO
NHTCA
NBnCbz
5
22b

*AB Sciex TOF/TOF™ Series Explorer™ 72041*

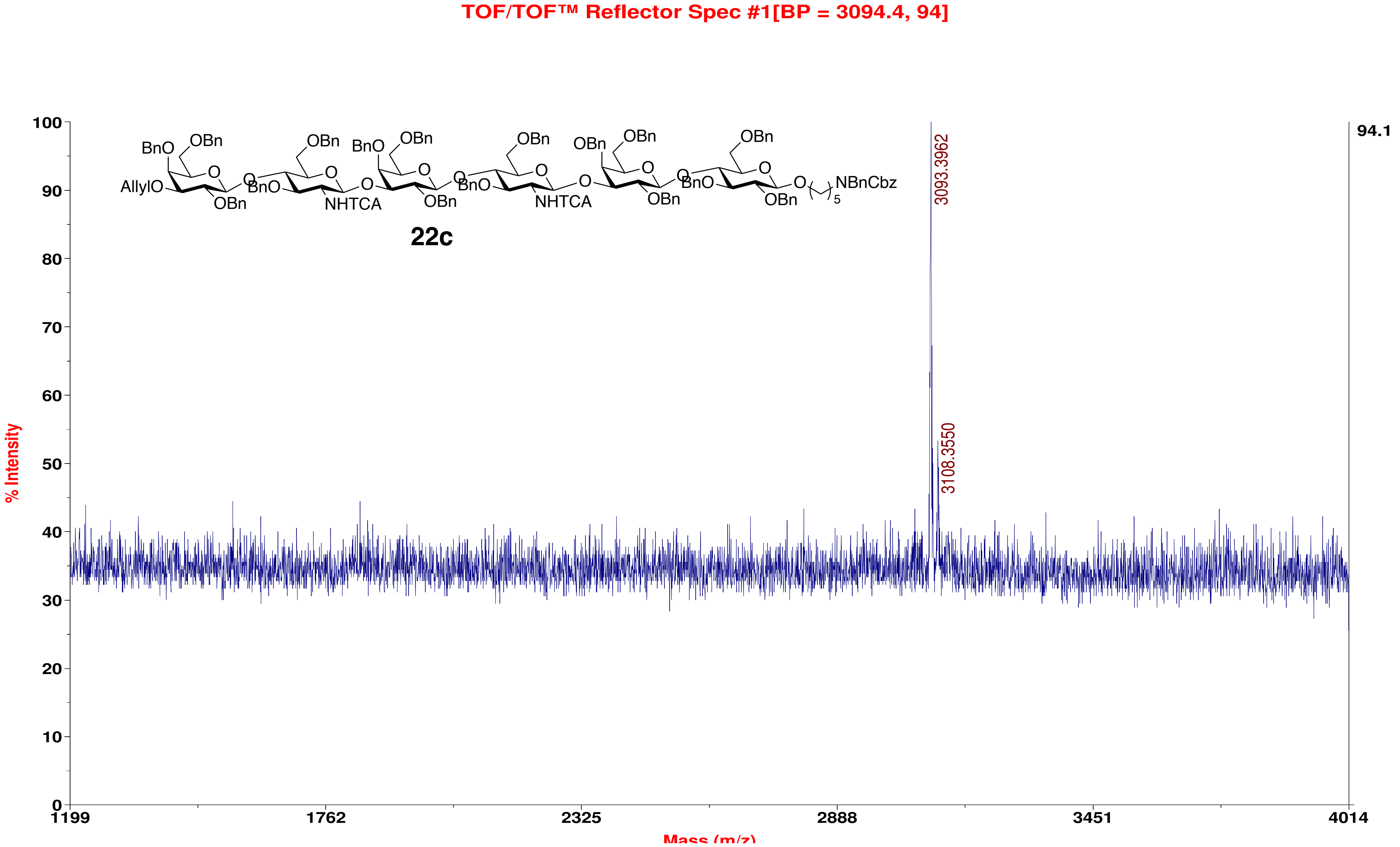

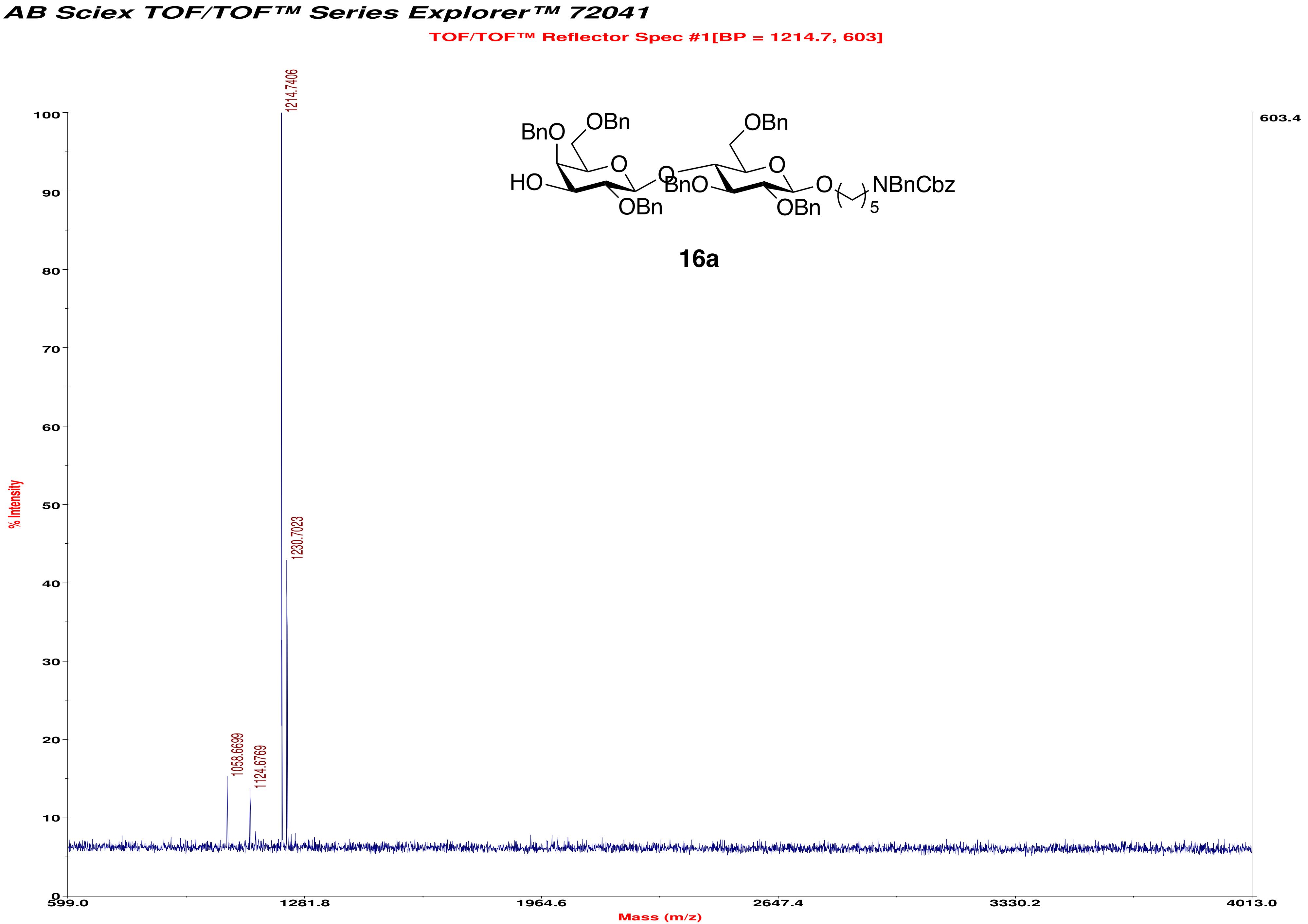
AB Sciex TOF/TOF™ Series Explorer™ 72041
TOF/TOF™ Reflector Spec #1[BP = 1214.7, 603]
1214.7406
1230.7023
1058.6699
1124.6769
603.4
% Intensity
Mass (m/z)
599.0
1281.8
1964.6
2647.4
3330.2
4013.0
BnO
OBn
HO
O
OBn
BnO
O
OBn
NBnCbz
5
16a

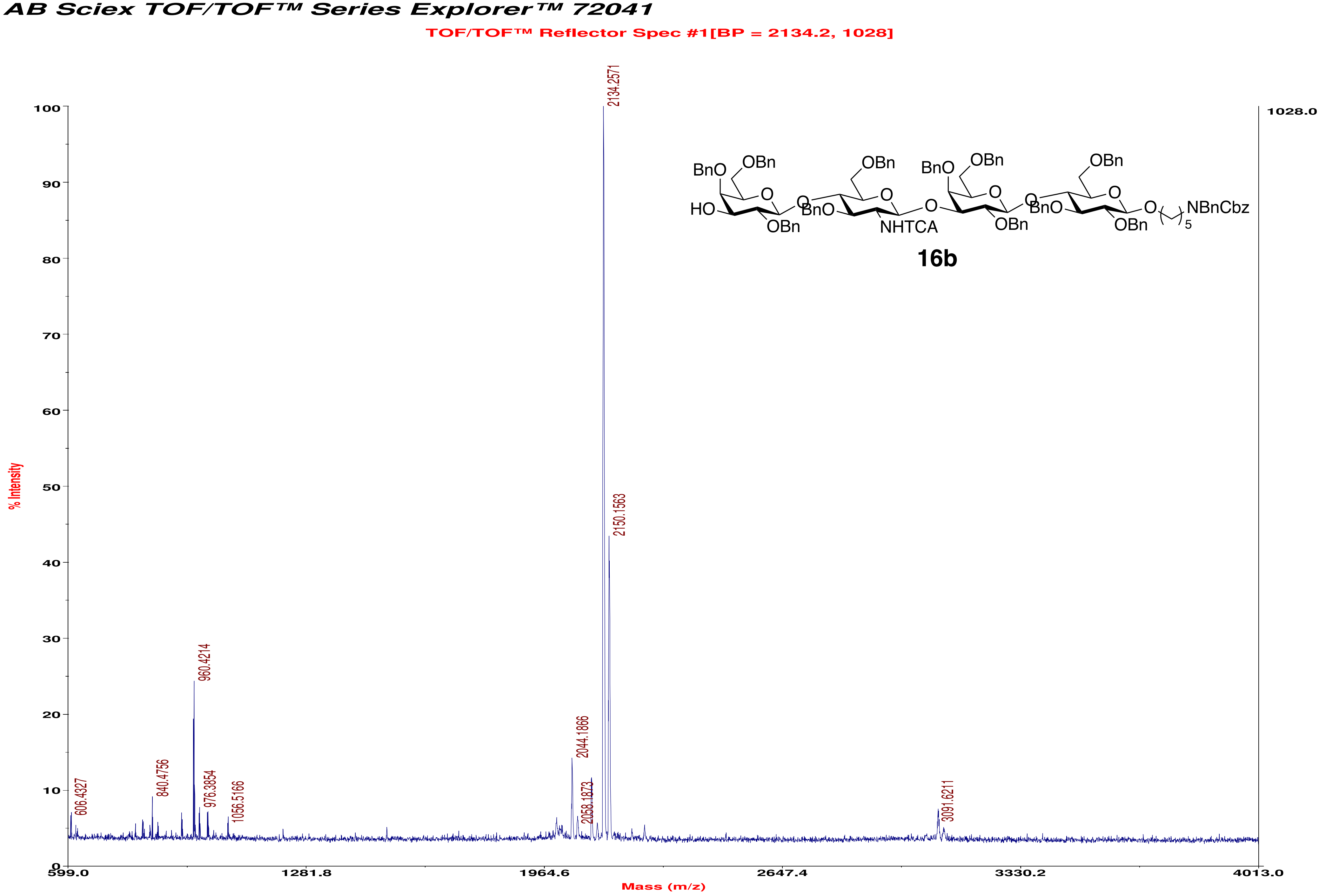
AB Sciex TOF/TOF™ Series Explorer™ 72041
TOF/TOF™ Reflector Spec #1[BP = 2134.2, 1028]
1028.0
% Intensity
Mass (m/z)
0
10
20
30
40
50
60
70
80
90
100
599.0
1281.8
1964.6
2647.4
3330.2
4013.0
606.4327
840.4756
960.4214
976.3854
1056.5166
2044.1866
2058.1873
2134.2571
2150.1563
3091.6211
BnO
OBn
HO
OBn
BnO
NHTCA
BnO
OBn
OBn
BnO
OBn
OBn
NBnCbz
5
16b

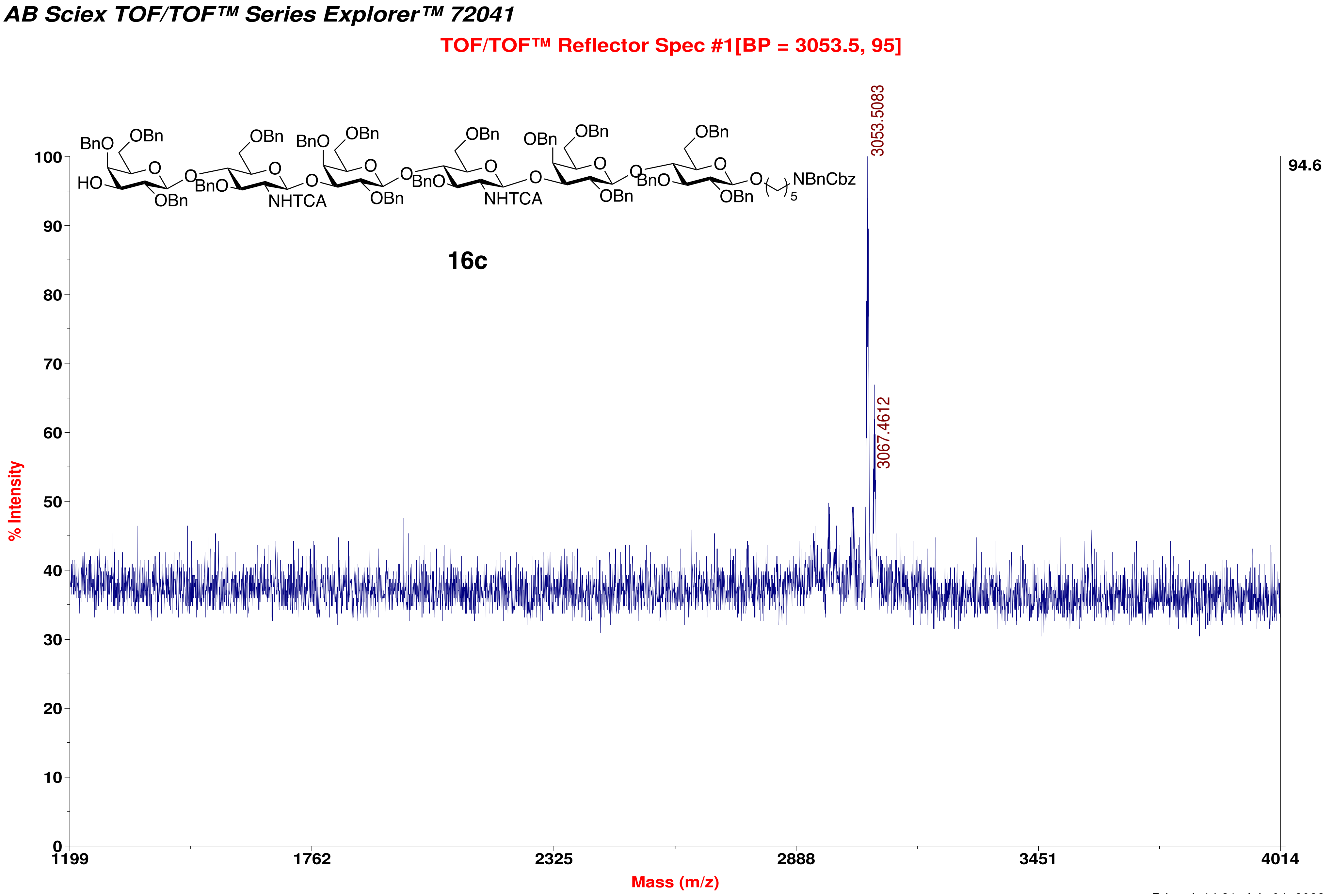
AB Sciex TOF/TOF™ Series Explorer™ 72041
TOF/TOF™ Reflector Spec #1[BP = 3053.5, 95]
16c
3053.5083
3067.4612
94.6
% Intensity
Mass (m/z)

**AB Sciex TOF/TOF™ Series Explorer™ 72041**

TOF/TOF™ Reflector Spec #1[BP = 1704.2, 6892]

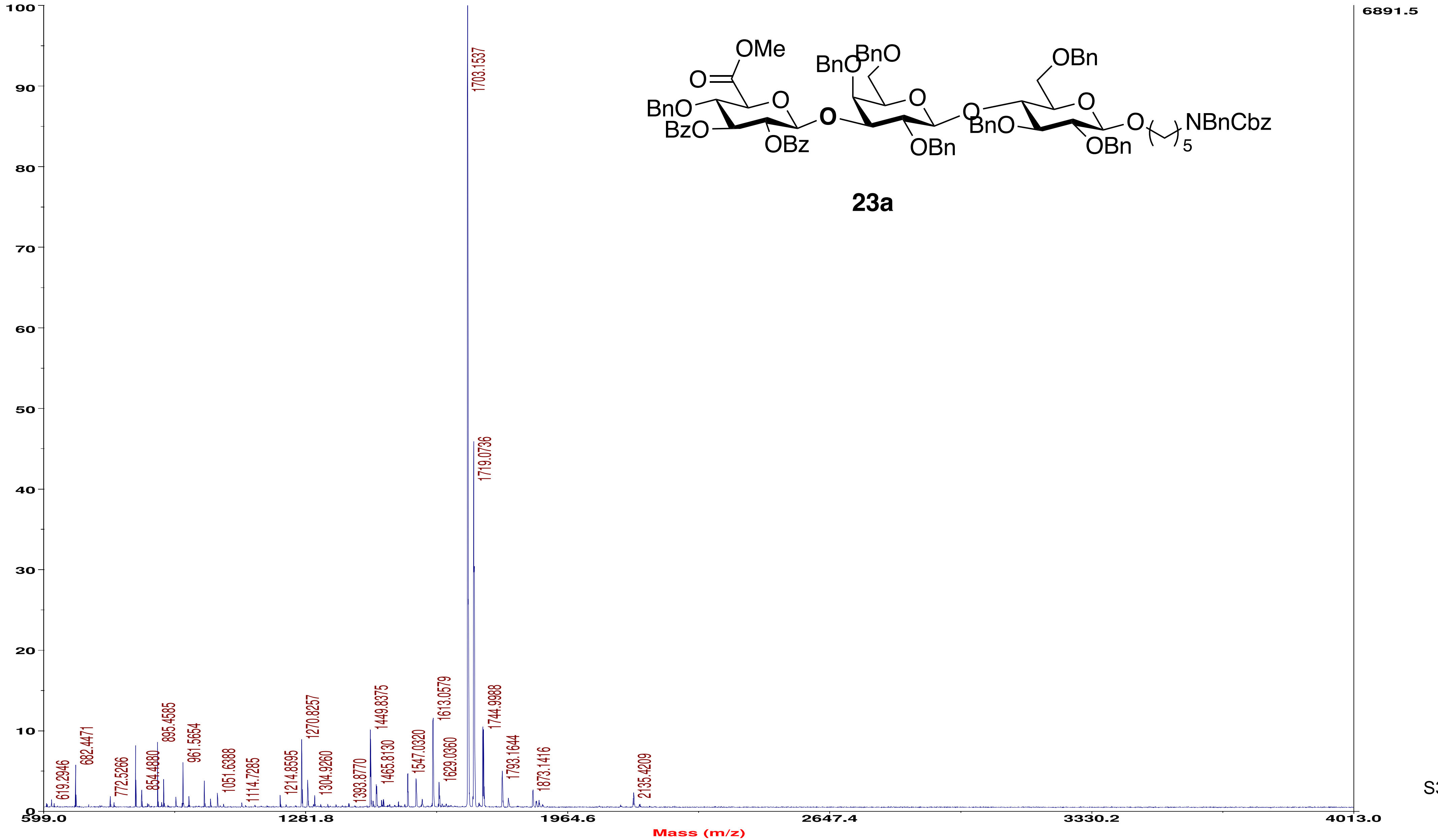


**23a**

AB Sciex TOF/TOF™ Series Explorer™ 72041

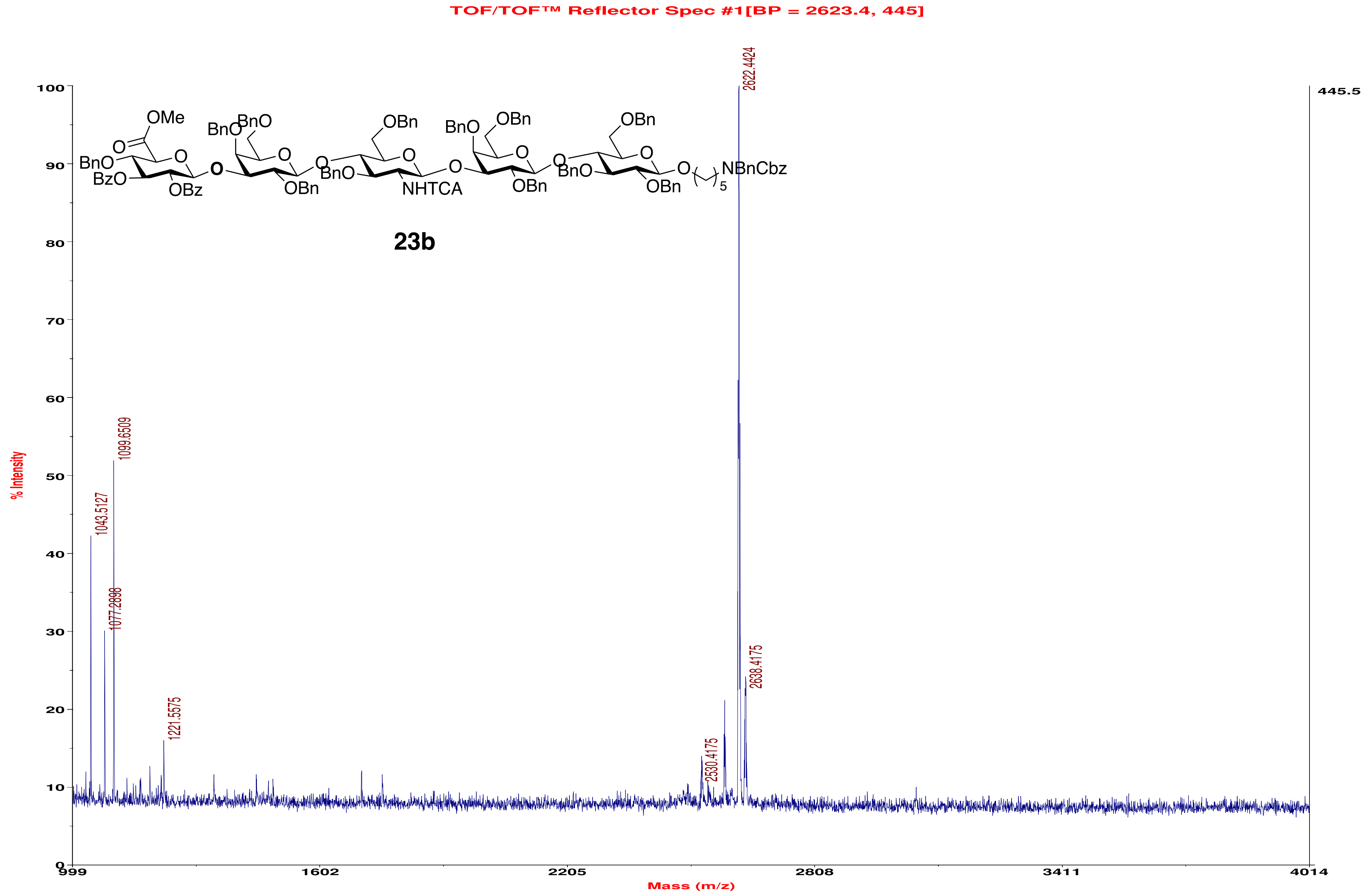

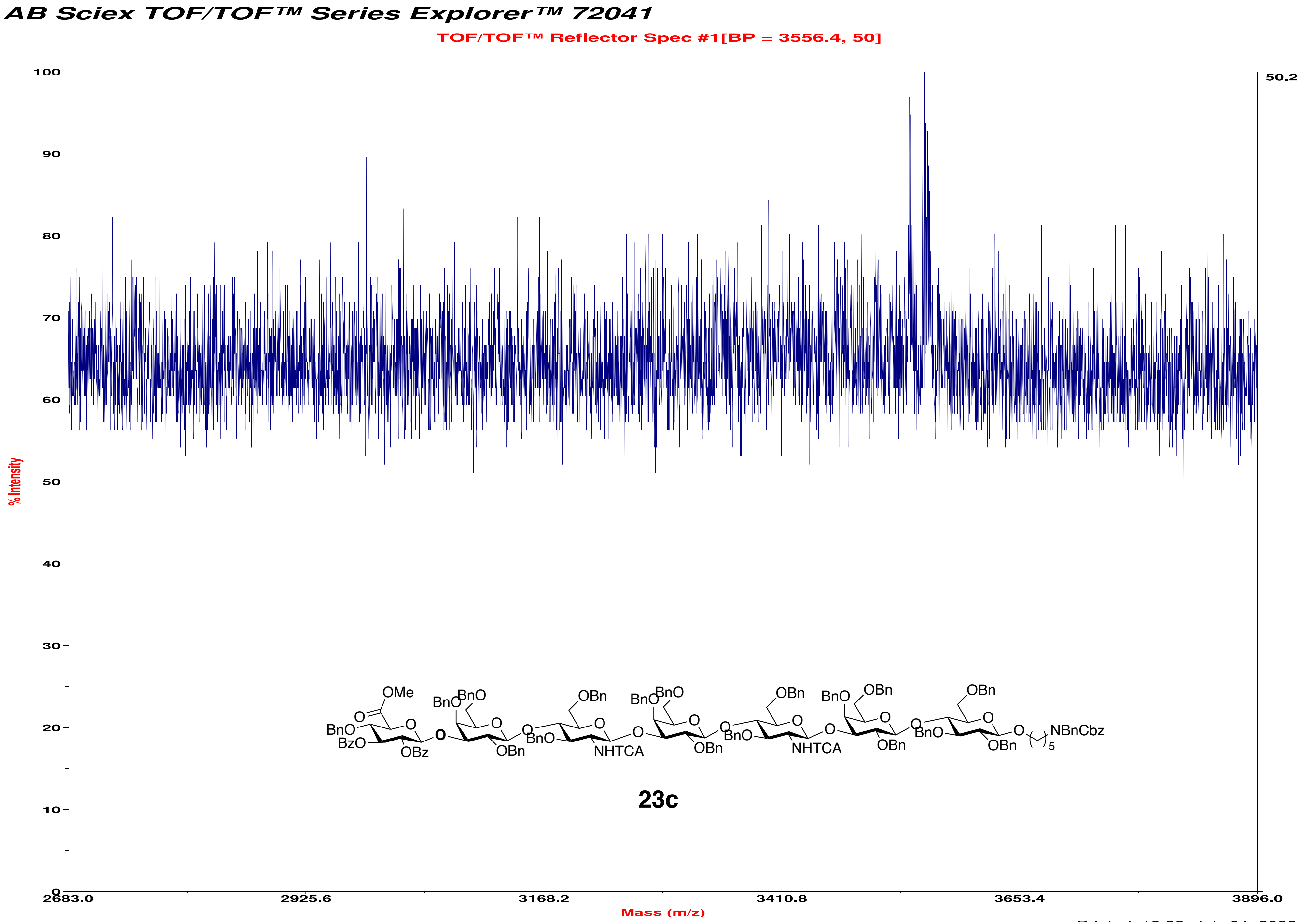
AB Sciex TOF/TOF™ Series Explorer™ 72041
TOF/TOF™ Reflector Spec #1[BP = 3556.4, 50]
% Intensity
Mass (m/z)
50.2
100
90
80
70
60
50
40
30
20
10
0
2683.0
2925.6
3168.2
3410.8
3653.4
3896.0
OMe
BnO
BzO
OBz
BnO
OBn
NHTCA
NBnCbz
5
23c

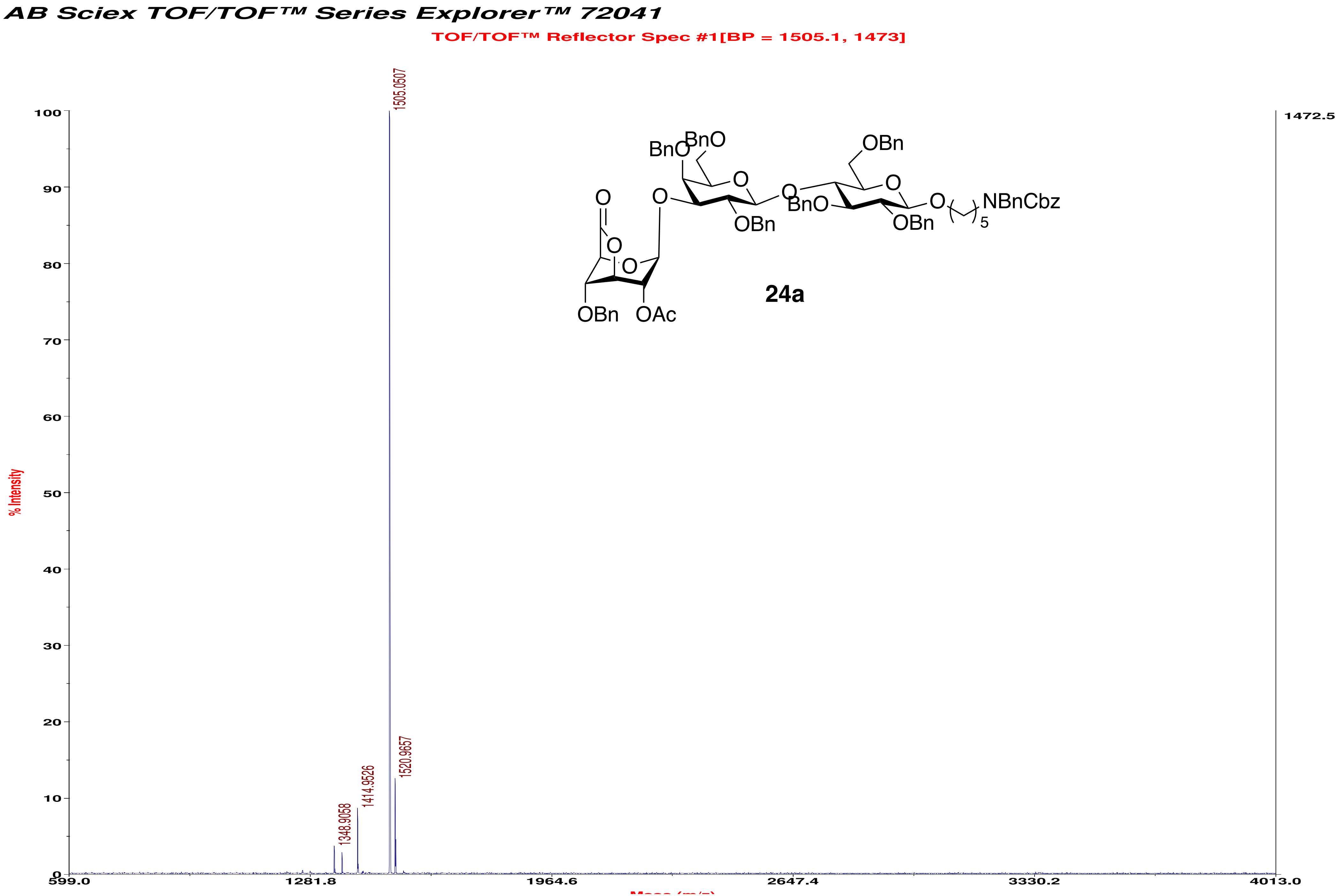
AB Sciex TOF/TOF™ Series Explorer™ 72041
TOF/TOF™ Reflector Spec #1[BP = 1505.1, 1473]
% Intensity
1505.0507
1520.9657
1414.9526
1348.9058
1472.5
599.0
1281.8
1964.6
2647.4
3330.2
4013.0
0
10
20
30
40
50
60
70
80
90
100
BnO
BnO
OBn
O
O
O
O
O
O
O
BnO
OBn
OBn
NBnCbz
5
OBn
OAc
24a

AB Sciex TOF/TOF™ Series Explorer™ 72041

TOF/TOF™ Reflector Spec #1[BP = 2424.2, 302]

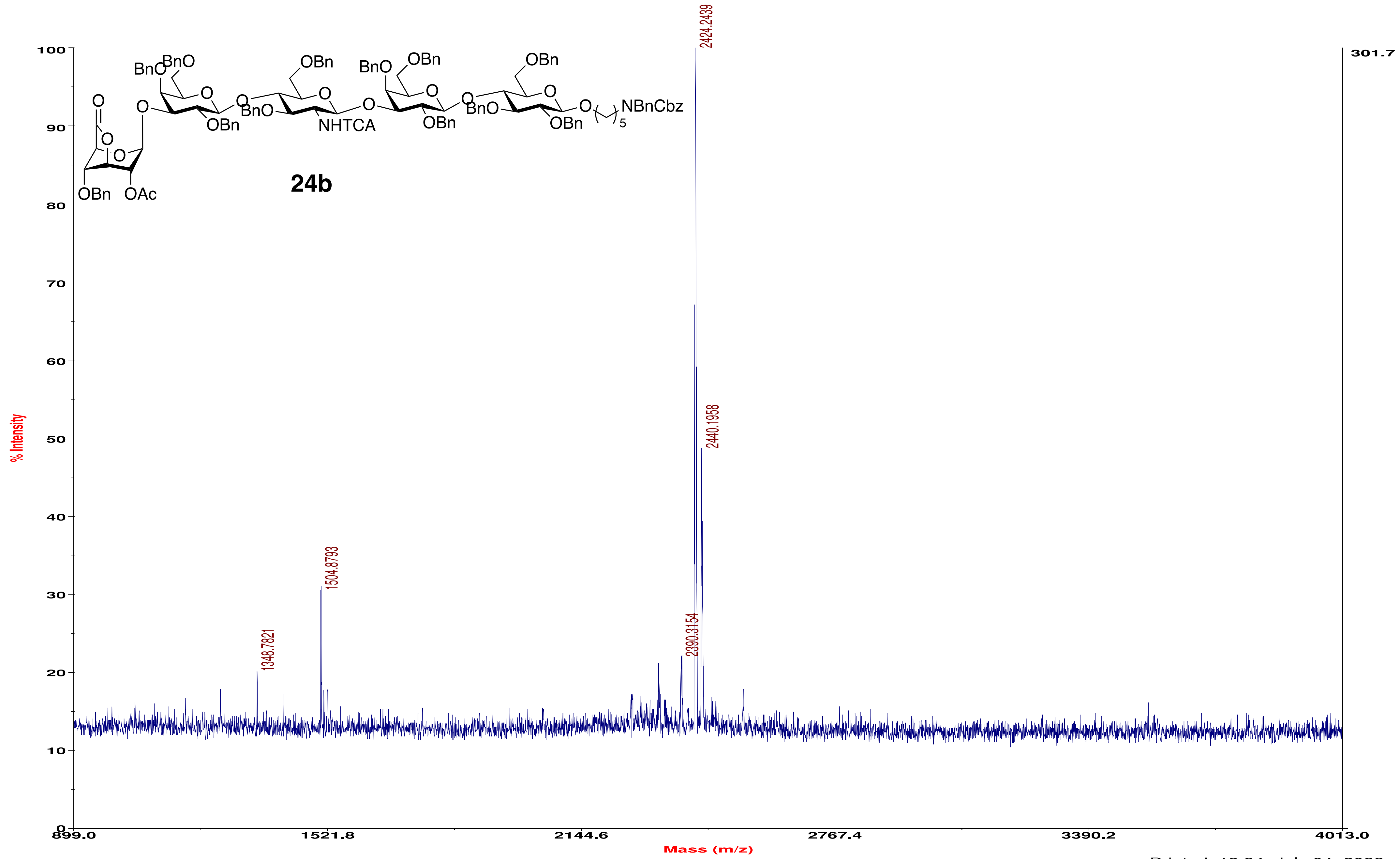

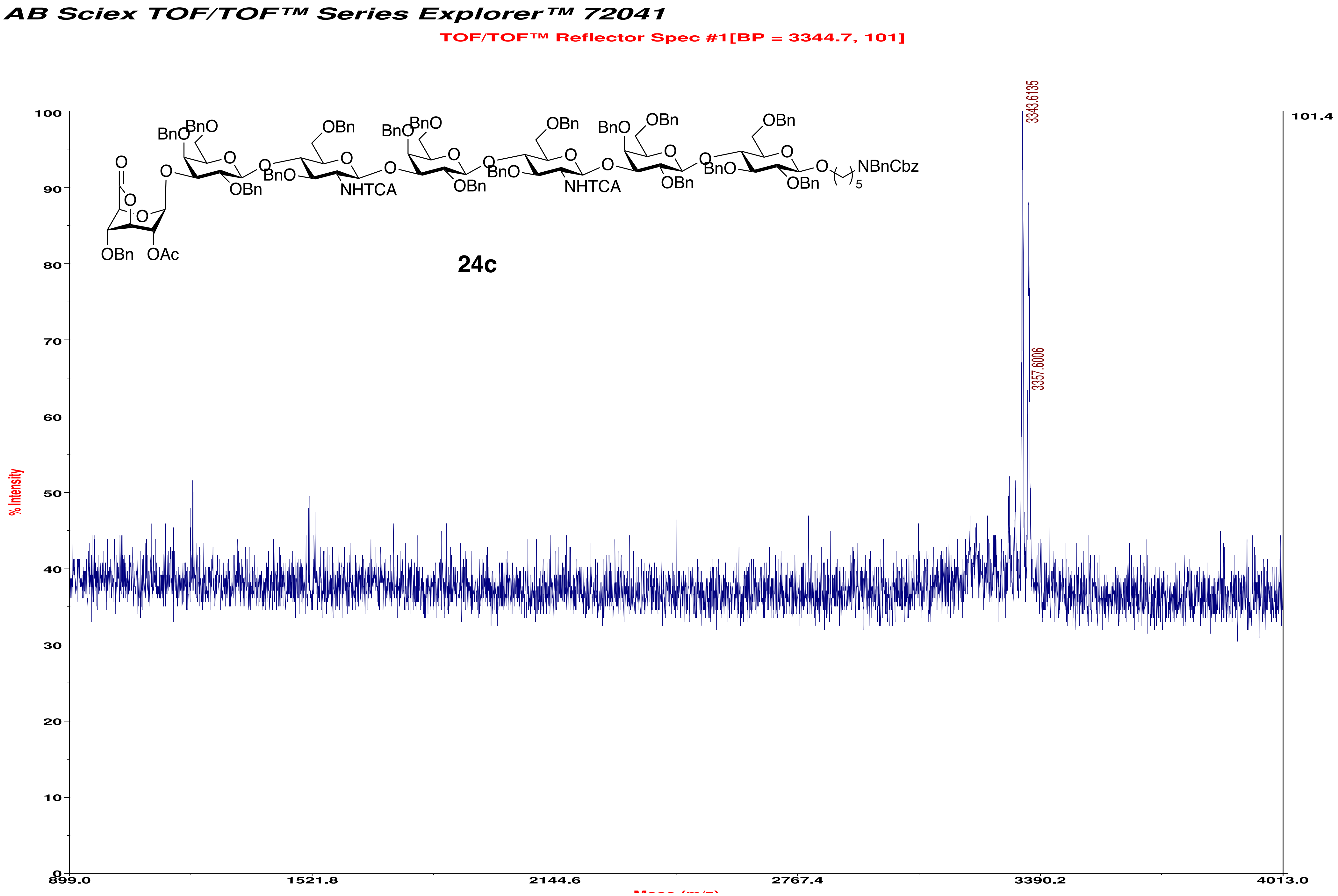
AB Sciex TOF/TOF™ Series Explorer™ 72041
TOF/TOF™ Reflector Spec #1[BP = 3344.7, 101]
3343.6135
3357.6006
101.4
% Intensity
BnO
OBn
O
OAc
BnO
NHTCA
NBnCbz
24c
899.0
1521.8
2144.6
2767.4
3390.2
4013.0

AB Sciex TOF/TOF™ Series Explorer™ 72041

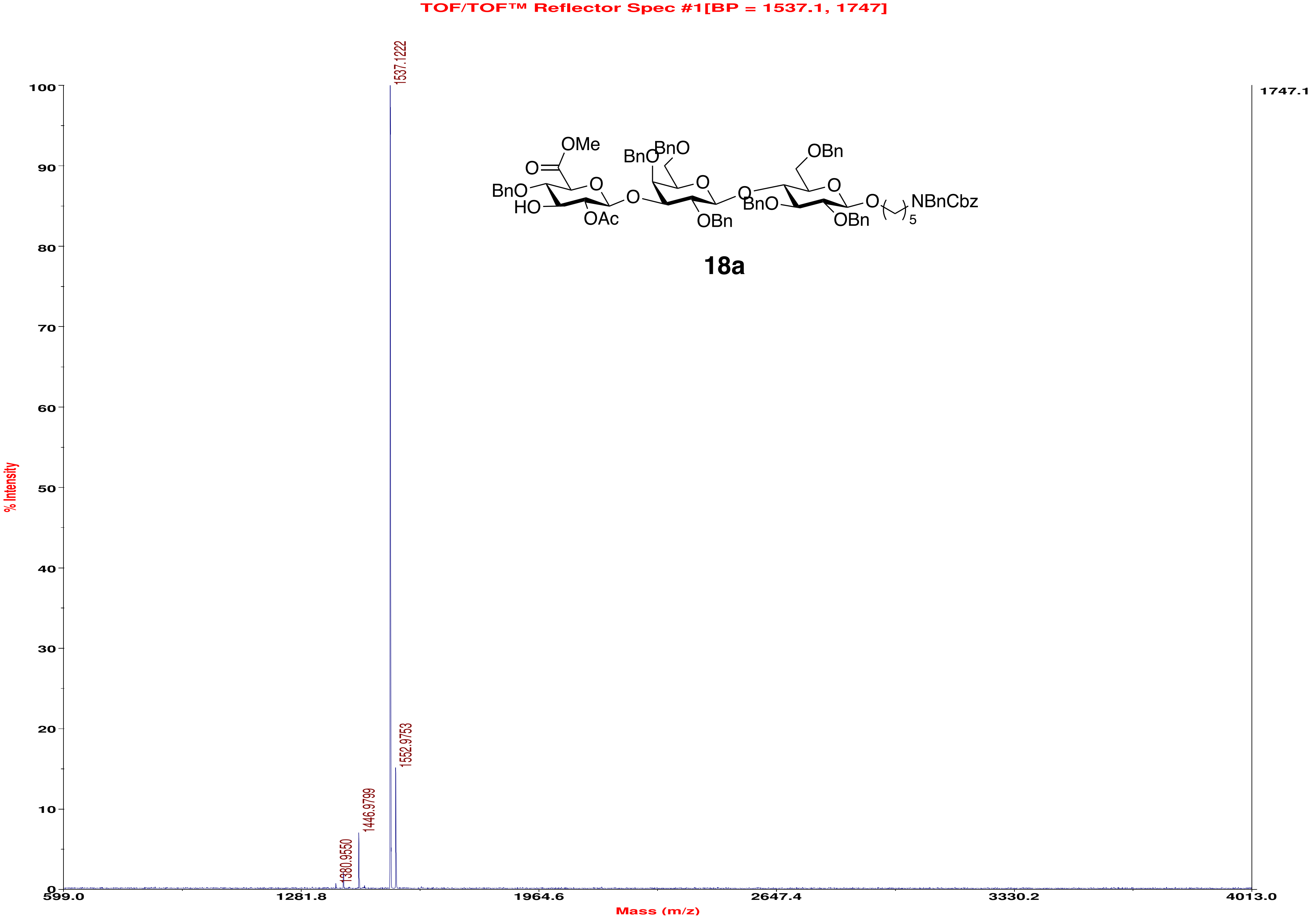

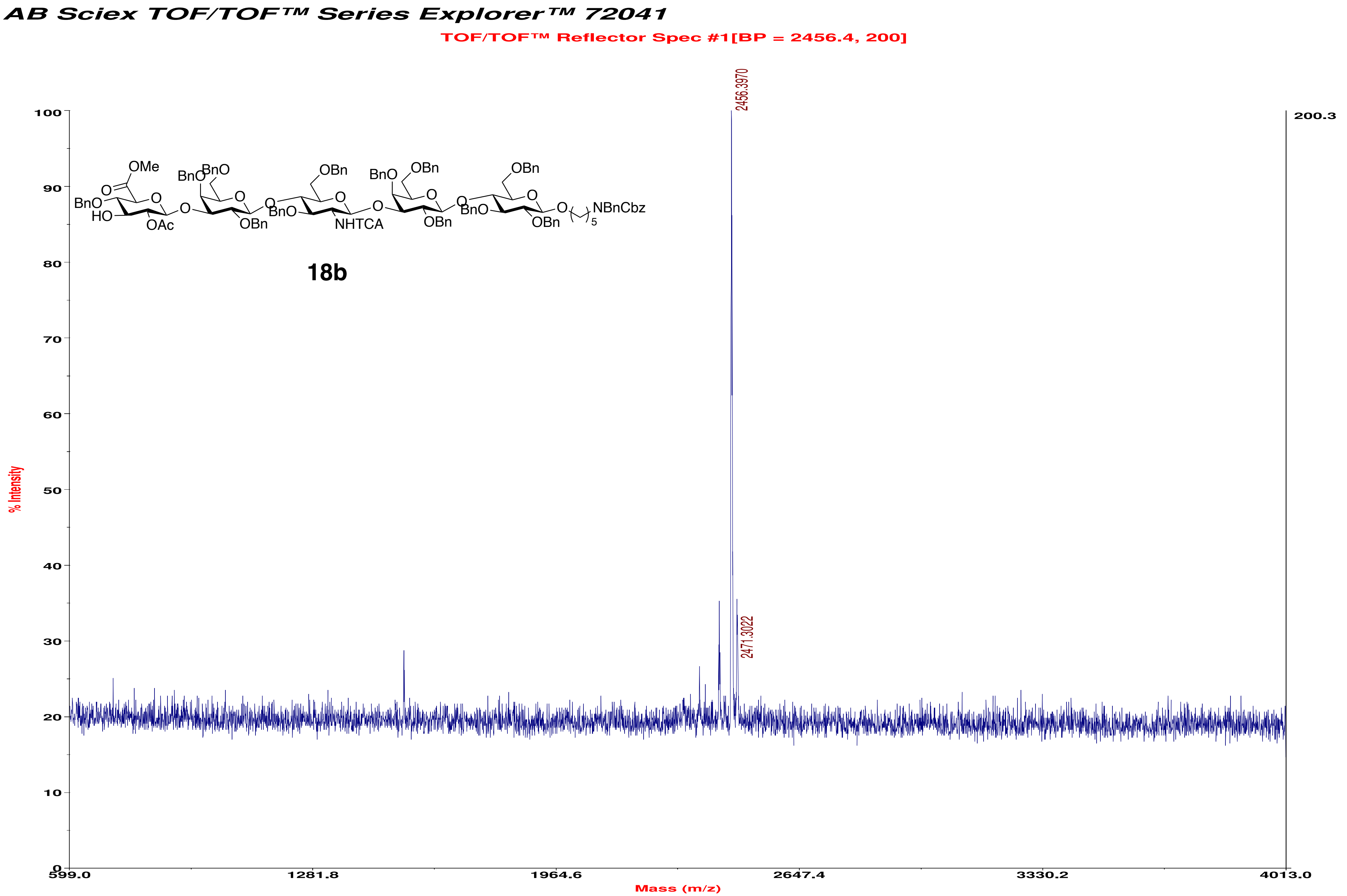
AB Sciex TOF/TOF™ Series Explorer™ 72041
TOF/TOF™ Reflector Spec #1[BP = 2456.4, 200]
2456.3970
2471.3022
200.3
% Intensity
Mass (m/z)
18b
NHTCA
NBnCbz
OMe
OAc

*AB Sciex TOF/TOF™ Series Explorer™ 72041*

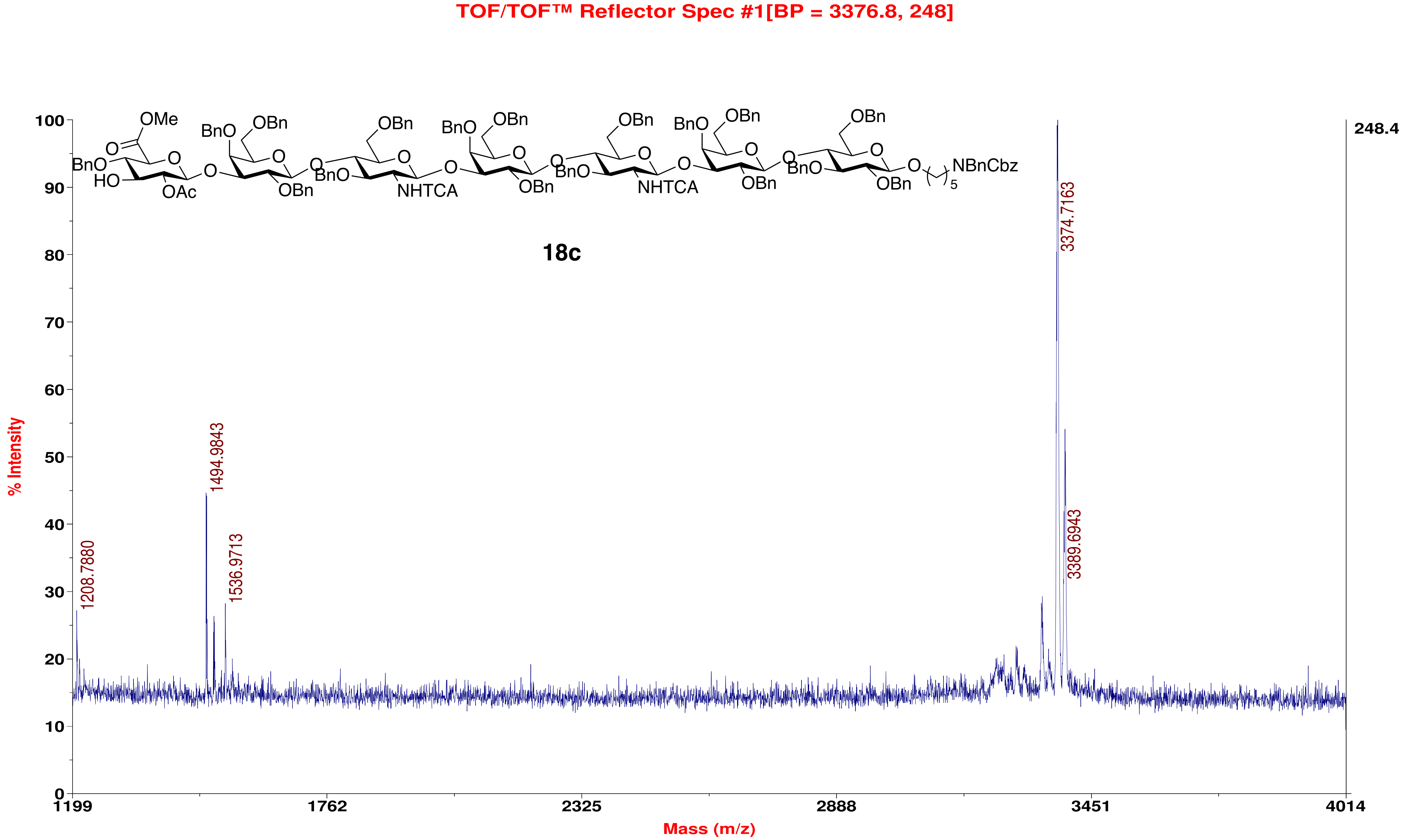

#### AB Sciex TOF/TOF™ Series Explorer™ 72041

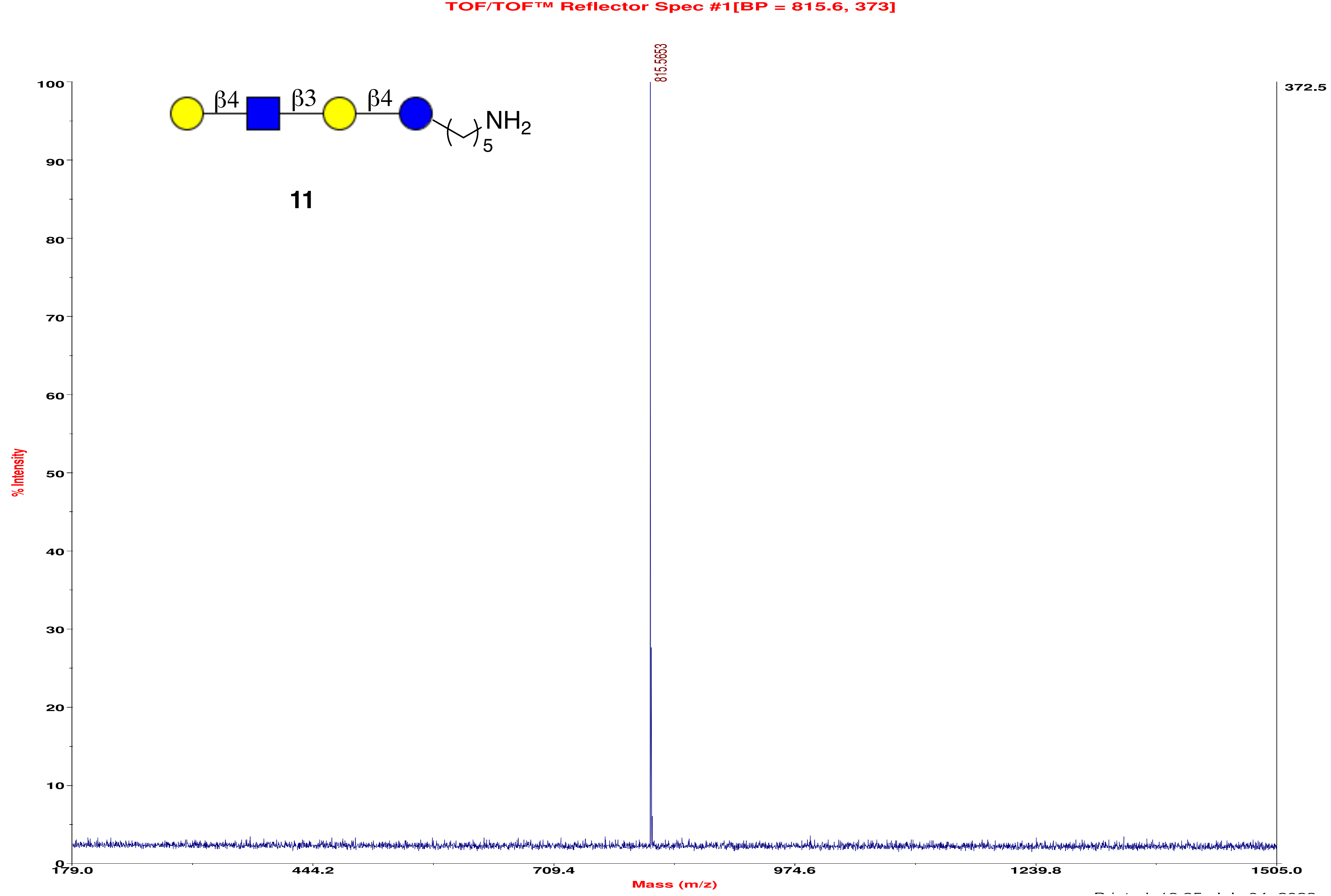

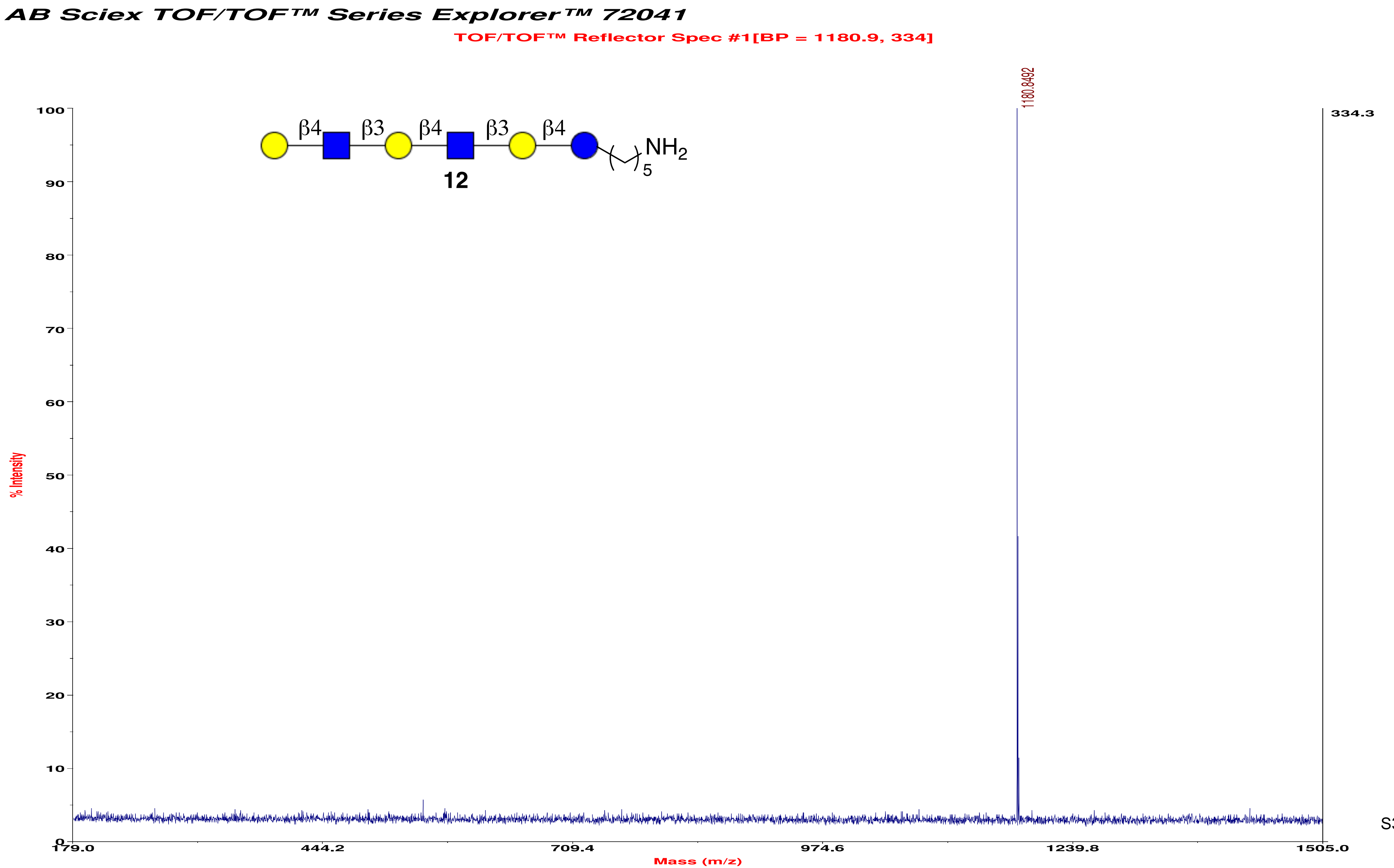
AB Sciex TOF/TOF™ Series Explorer™ 72041
TOF/TOF™ Reflector Spec #1[BP = 1180.9, 334]
1180.8492
334.3
β4
β3
β4
β3
β4
NH2
5
12
% Intensity
Mass (m/z)
179.0
444.2
709.4
974.6
1239.8
1505.0
0
10
20
30
40
50
60
70
80
90
100

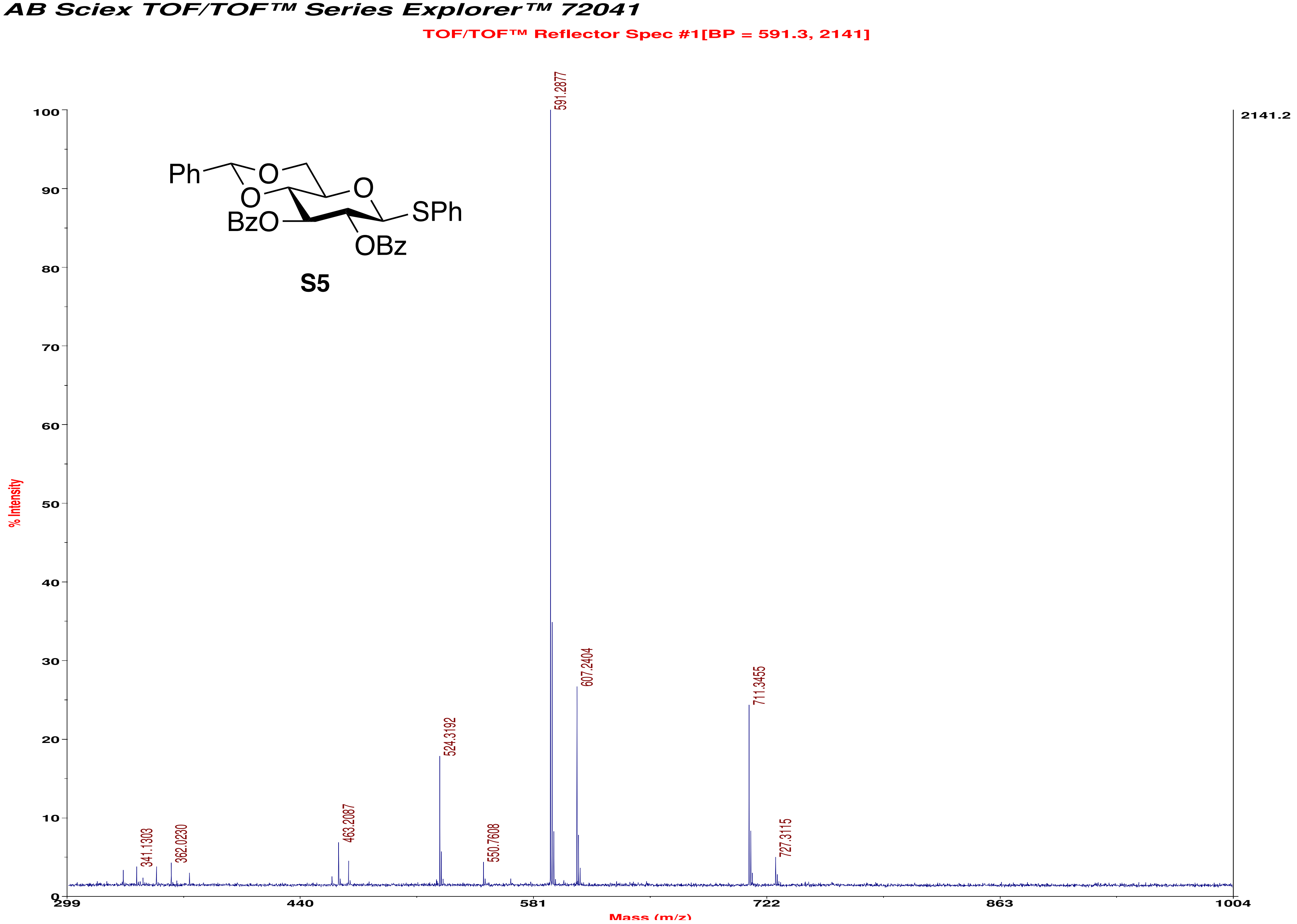
AB Sciex TOF/TOF™ Series Explorer™ 72041
TOF/TOF™ Reflector Spec #1[BP = 591.3, 2141]
591.2877
607.2404
711.3455
524.3192
463.2087
341.1303
362.0230
550.7608
727.3115
2141.2
% Intensity
Mass (m/z)
S5

AB Sciex TOF/TOF™ Series Explorer™ 72041

TOF/TOF™ Reflector Spec #1[BP = 593.3, 16129]

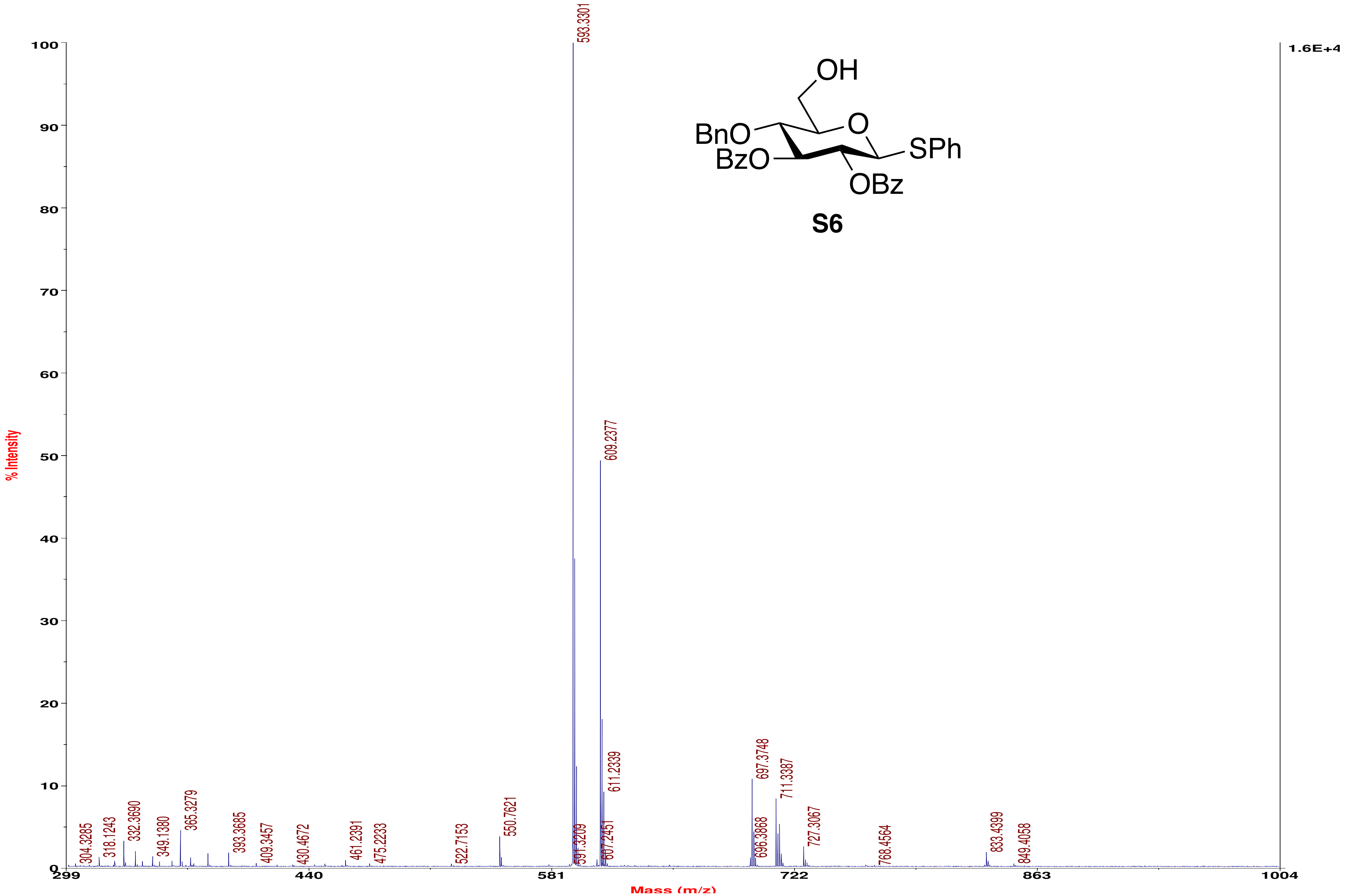

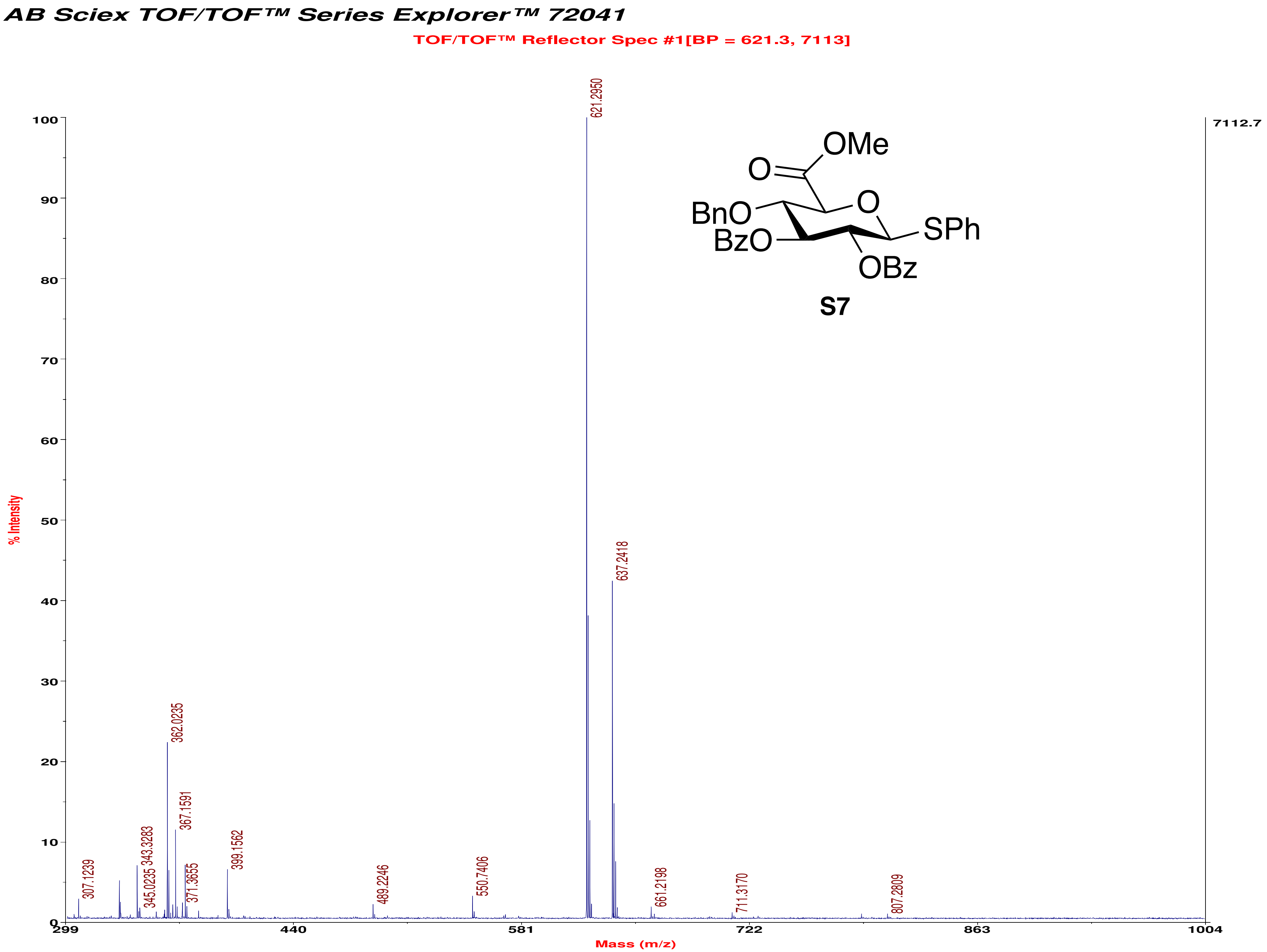
AB Sciex TOF/TOF™ Series Explorer™ 72041
TOF/TOF™ Reflector Spec #1[BP = 621.3, 7113]
% Intensity
Mass (m/z)
7112.7
621.2950
637.2418
362.0235
367.1591
343.3283
345.0235
371.3655
399.1562
307.1239
489.2246
550.7406
661.2198
711.3170
807.2809
OMe
O
O
BnO
BzO
SPh
OBz
S7

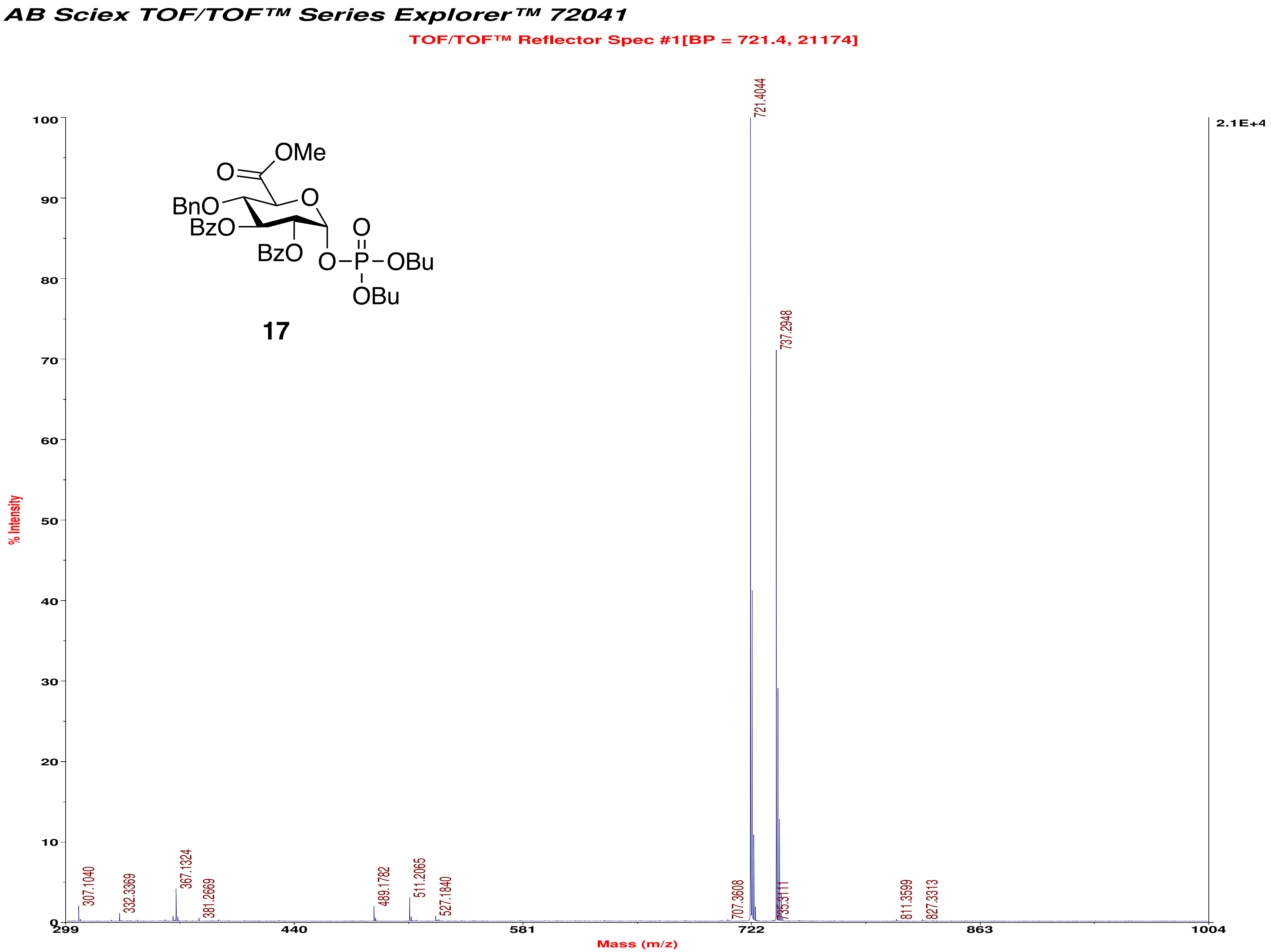
AB Sciex TOF/TOF™ Series Explorer™ 72041
TOF/TOF™ Reflector Spec #1[BP = 721.4, 21174]
OMe
O
BnO
BzO
O
BzO
O
O–P–OBu
OBu
17
% Intensity
2.1E+4
721.4044
737.2948
307.1040
332.3369
367.1324
381.2669
489.1782
511.2065
527.1840
707.3608
811.3599
827.3313
Mass (m/z)

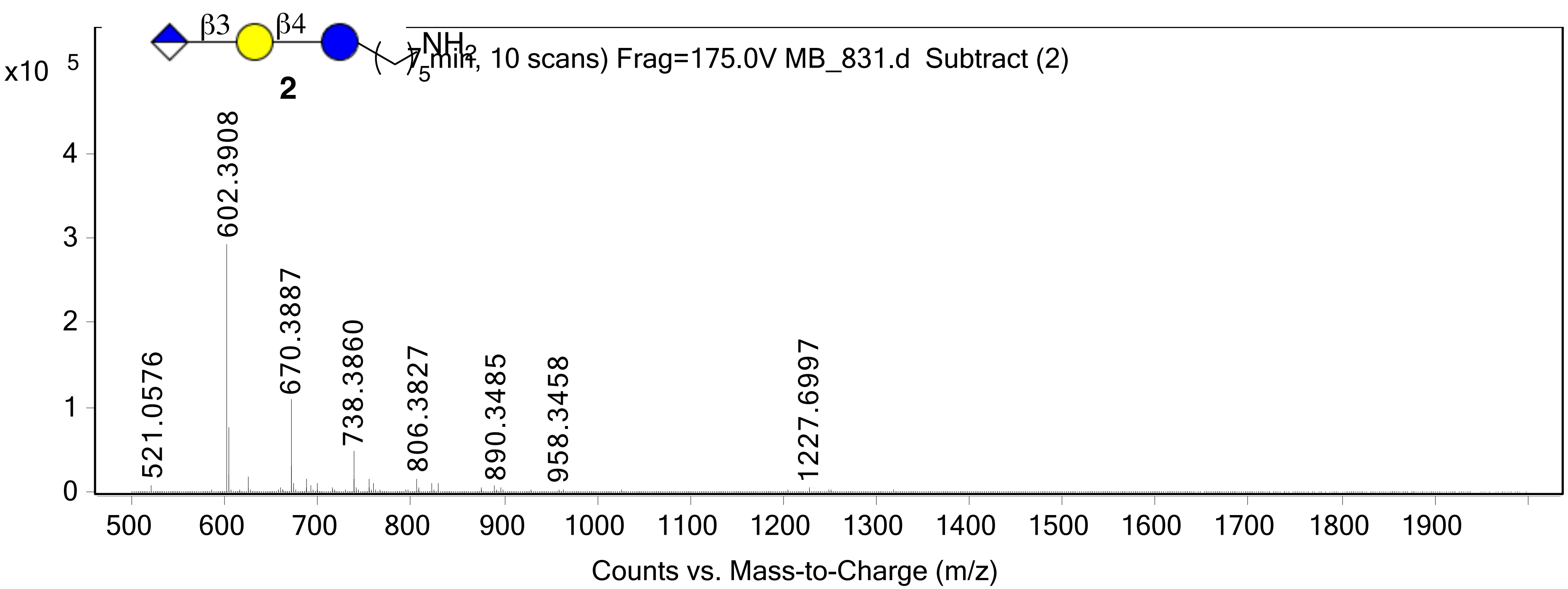

β3
β4
NH2
5
2
x10 5
7 min, 10 scans) Frag=175.0V MB_831.d Subtract (2)
602.3908
670.3887
738.3860
806.3827
890.3485
958.3458
1227.6997
521.0576
0
1
2
3
4
500 600 700 800 900 1000 1100 1200 1300 1400 1500 1600 1700 1800 1900
Counts vs. Mass-to-Charge (m/z)

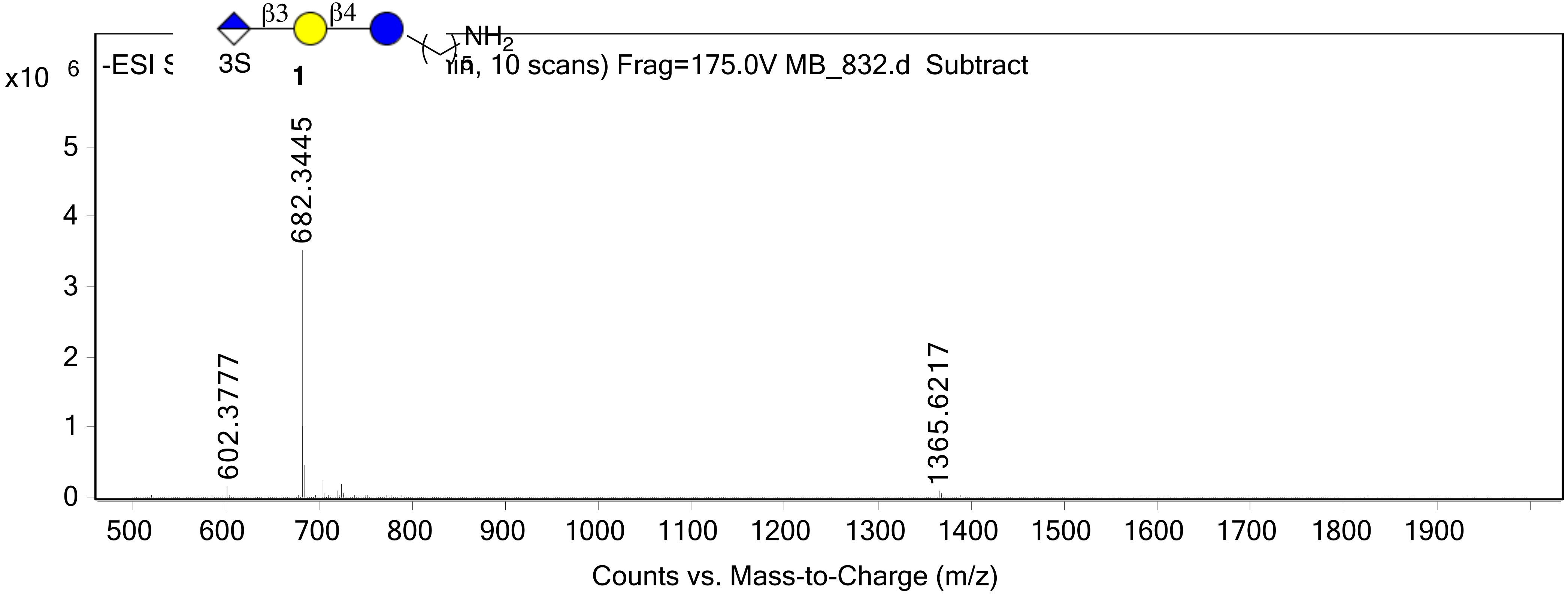
β3
β4
NH2
3S
1
5
x10 6
-ESI S
, 10 scans) Frag=175.0V MB_832.d Subtract
682.3445
602.3777
1365.6217
0
1
2
3
4
5
500
600
700
800
900
1000
1100
1200
1300
1400
1500
1600
1700
1800
1900
Counts vs. Mass-to-Charge (m/z)

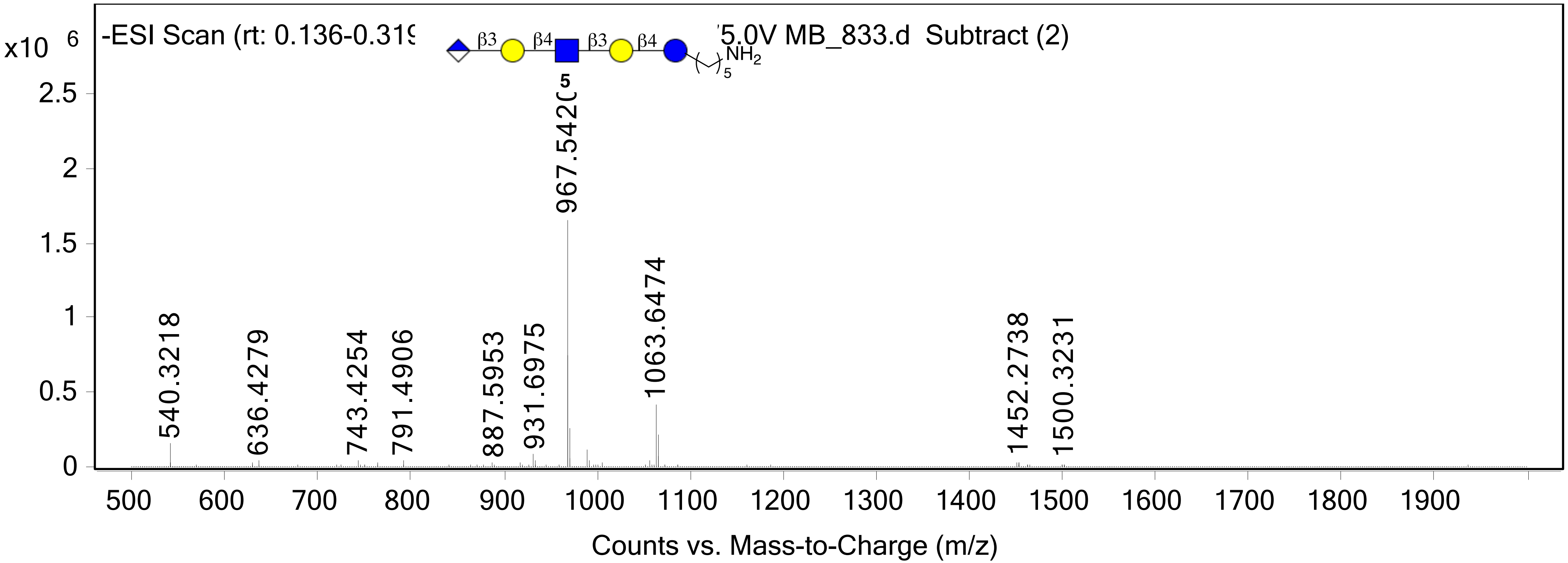

-ESI Scan (rt: 0.136-0.319
5.0V MB_833.d Subtract (2)
x10 6
β3
β4
β3
β4
NH2
5
5
967.5420
1063.6474
540.3218
636.4279
743.4254
791.4906
887.5953
931.6975
1452.2738
1500.3231
2.5
2
1.5
1
0.5
0
500
600
700
800
900
1000
1100
1200
1300
1400
1500
1600
1700
1800
1900
Counts vs. Mass-to-Charge (m/z)

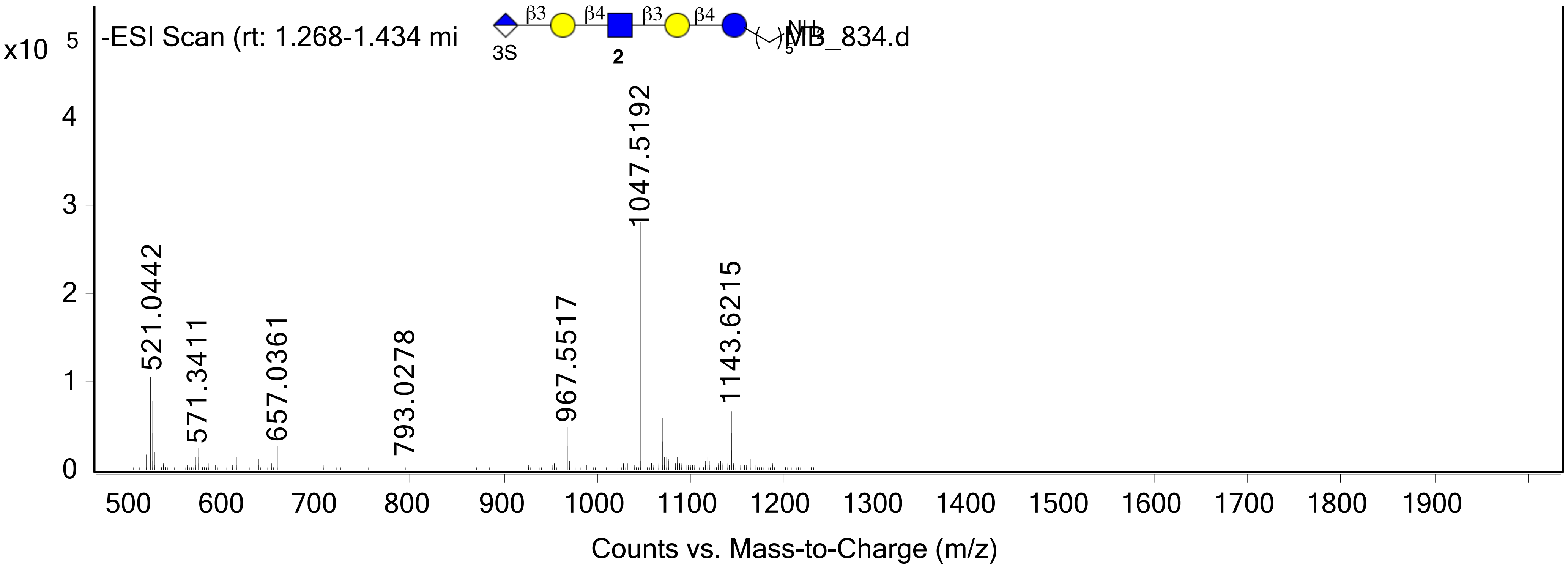

-ESI Scan (rt: 1.268-1.434 mi
MB_834.d
x10 5
β3
β4
β3
β4
3S
2
NH
5
521.0442
571.3411
657.0361
793.0278
967.5517
1047.5192
1143.6215
0
1
2
3
4
500
600
700
800
900
1000
1100
1200
1300
1400
1500
1600
1700
1800
1900
Counts vs. Mass-to-Charge (m/z)

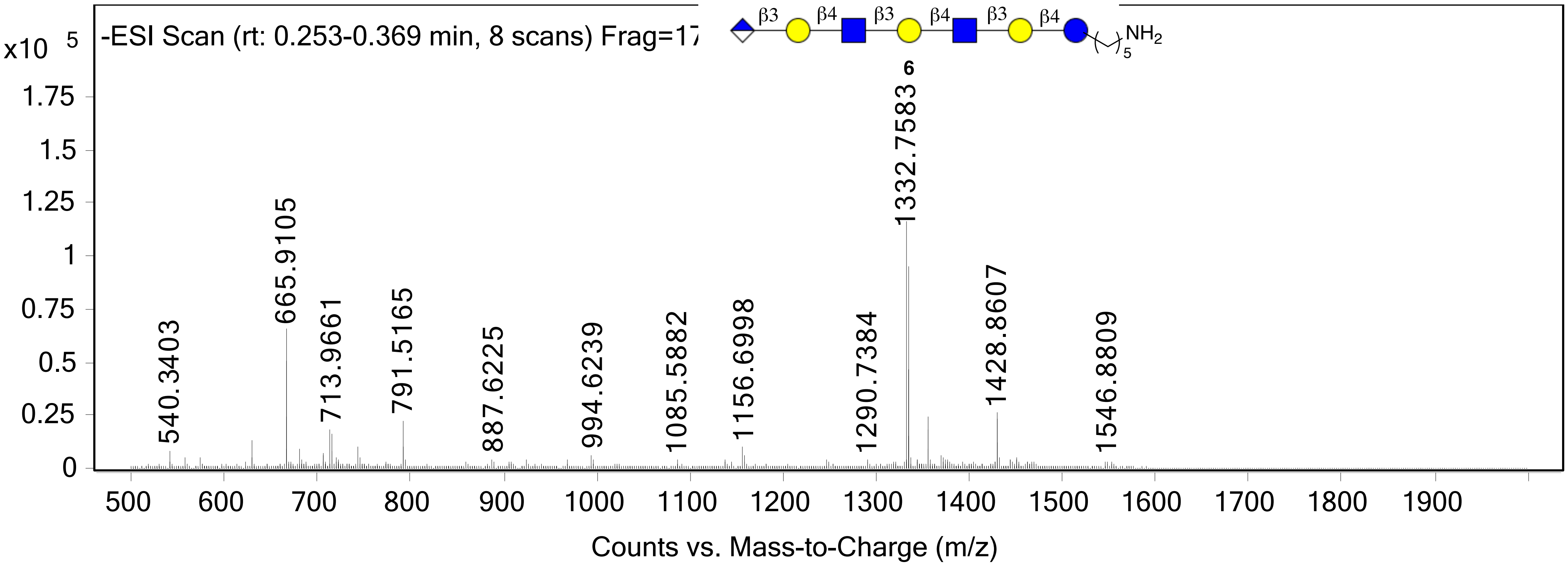
-ESI Scan (rt: 0.253-0.369 min, 8 scans) Frag=17
x10 5
β3
β4
β3
β4
β3
β4
NH2
5
6
1332.7583
665.9105
540.3403
713.9661
791.5165
887.6225
994.6239
1085.5882
1156.6998
1290.7384
1428.8607
1546.8809
Counts vs. Mass-to-Charge (m/z)

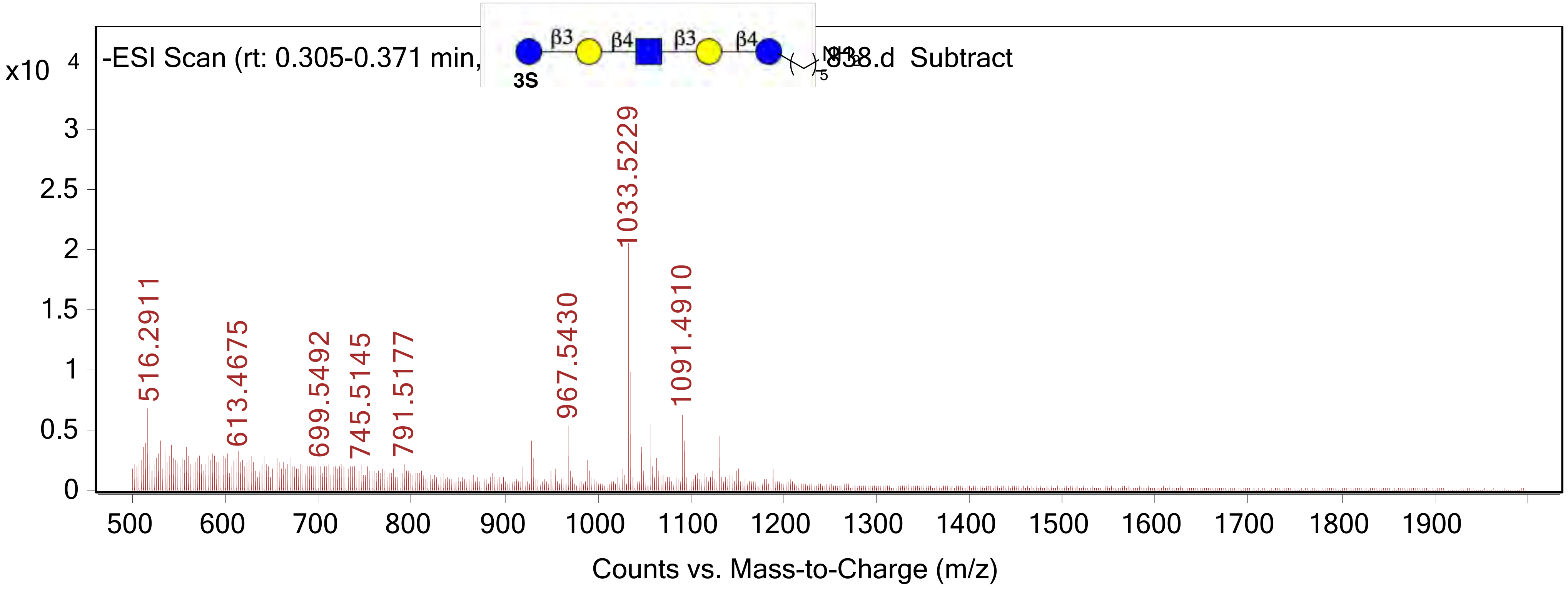

-ESI Scan (rt: 0.305-0.371 min,
838.d Subtract
β3
β4
β3
β4
3S
5
x10 4
3
2.5
2
1.5
1
0.5
0
516.2911
613.4675
699.5492
745.5145
791.5177
967.5430
1033.5229
1091.4910
500
600
700
800
900
1000
1100
1200
1300
1400
1500
1600
1700
1800
1900
Counts vs. Mass-to-Charge (m/z)

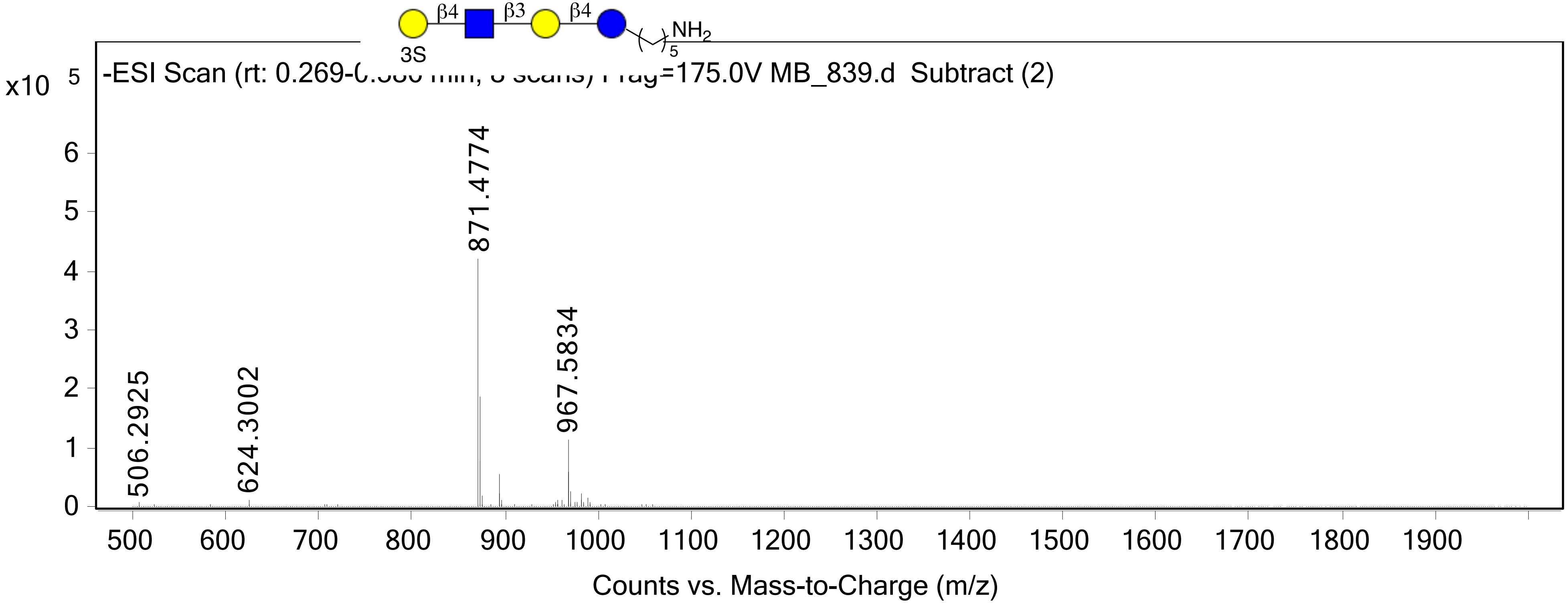

β4
β3
β4
3S
NH2
5
x10 5
-ESI Scan (rt: 0.269-
=175.0V MB_839.d Subtract (2)
871.4774
967.5834
506.2925
624.3002
0
1
2
3
4
5
6
500
600
700
800
900
1000
1100
1200
1300
1400
1500
1600
1700
1800
1900
Counts vs. Mass-to-Charge (m/z)

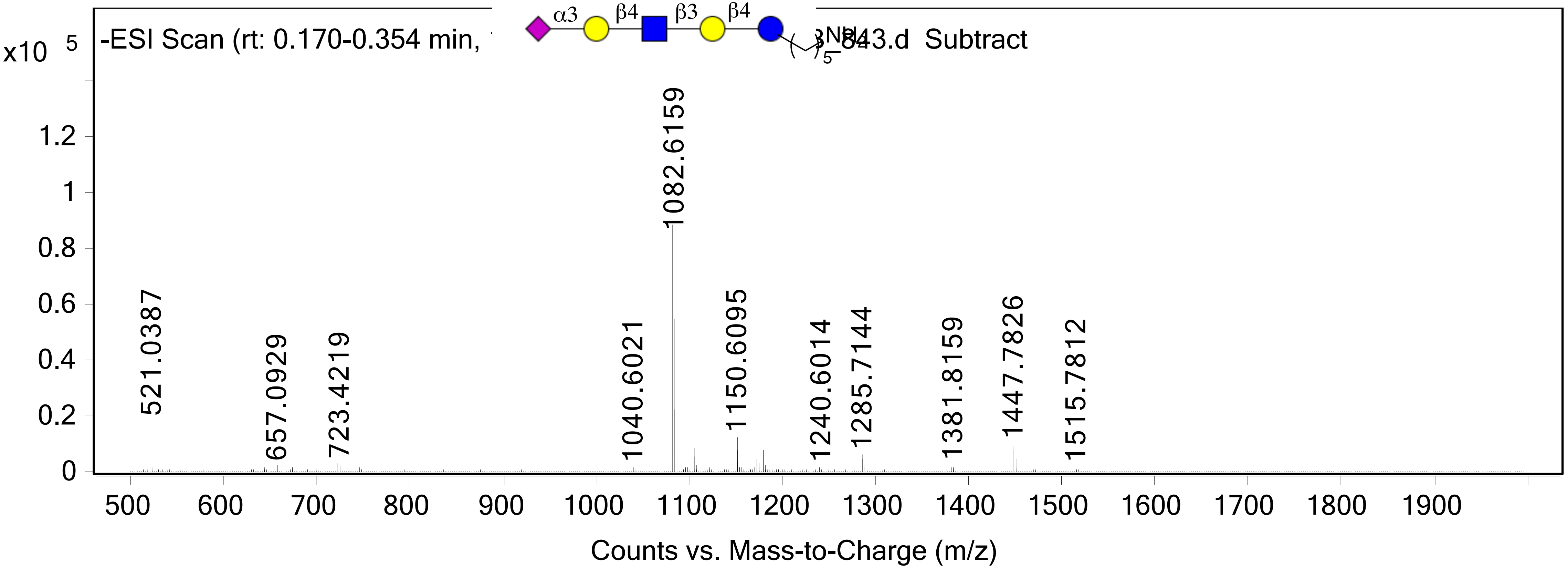

x10 5
-ESI Scan (rt: 0.170-0.354 min,
843.d Subtract
α3
β4
β3
β4
5
1.2
1
0.8
0.6
0.4
0.2
0
521.0387
657.0929
723.4219
1040.6021
1082.6159
1150.6095
1240.6014
1285.7144
1381.8159
1447.7826
1515.7812
500 600 700 800 900 1000 1100 1200 1300 1400 1500 1600 1700 1800 1900
Counts vs. Mass-to-Charge (m/z)

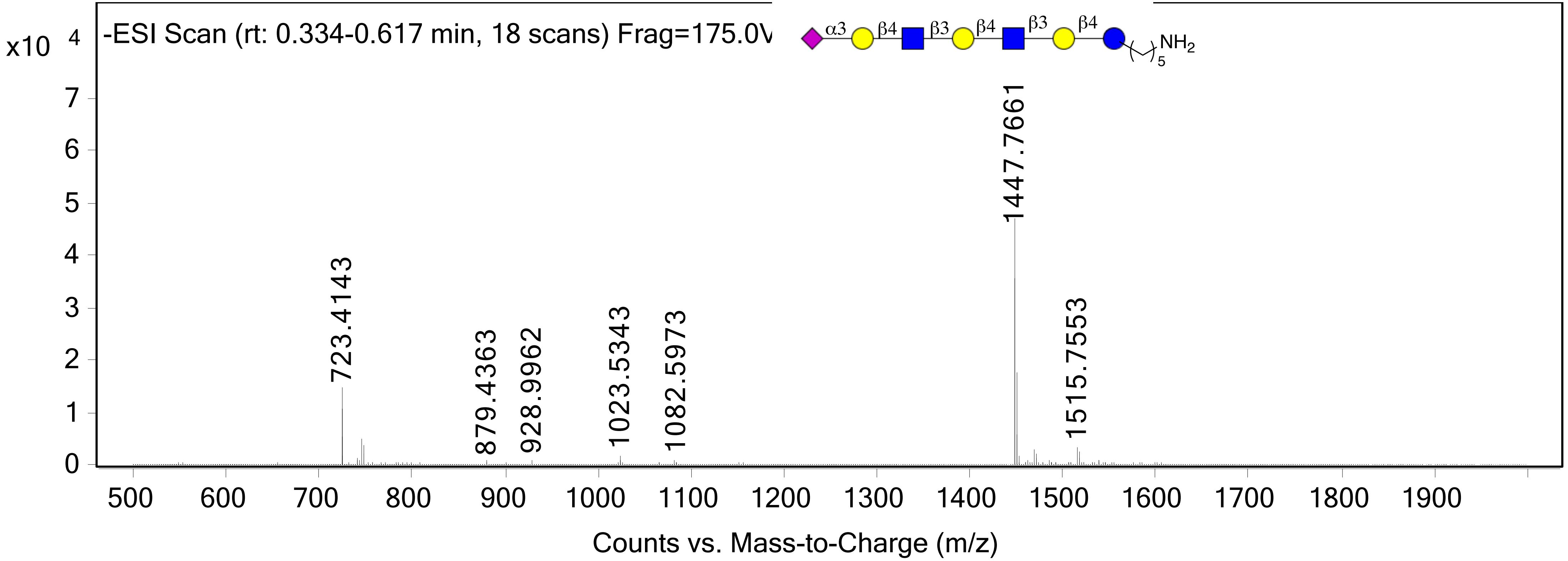

x10 4
-ESI Scan (rt: 0.334-0.617 min, 18 scans) Frag=175.0V
α3
β4
β3
β4
β3
β4
NH2
5
7
6
5
4
3
2
1
0
723.4143
879.4363
928.9962
1023.5343
1082.5973
1447.7661
1515.7553
500
600
700
800
900
1000
1100
1200
1300
1400
1500
1600
1700
1800
1900
Counts vs. Mass-to-Charge (m/z)
